# Are railways detrimental to bird populations? A BDACI study on the construction of the Bothnia Line Railway

**DOI:** 10.1101/691535

**Authors:** Adriaan de Jong

## Abstract

Railways, like roads, are commonly considered bad for birdlife, but supporting data are very scarce. We conducted a 14-year before-during-after control-impact (BDACI) study of birds in agricultural landscapes affected by the construction of the Bothnia Line Railway (BLR) in northern Sweden. The “during” phase was split into two phases, one for the true construction period and one for the years when the railway was ready but not trafficked. Avian biodiversity increased in impact sites (N=13), but not in control sites (N=6). The numbers of breeding territories decreased correspondingly in impact and control sites, but trends differed between species and sites. Developments in the Degernäs site demonstrated that mitigation could be successful. Finally, there was no support for a shying-away effect of the BLR. Territory midpoints moved closer rather than away from the railway, albeit with variable patterns for individual sites and species. Mixed effect models showed no differences in avian biodiversity between the *Construction*, *Ready* and *Traffic* phases compared with *Before*, but relative increases in numbers of territories and decreases in territory midpoint distances to the BLR. The results do not support a general detrimental effect of railway construction on bird populations in terms of biodiversity loss, reduced abundances or shying-away from train traffic. More studies and the development of “Railway Ecology” are badly needed.

## Background

In line with how we foster our children and value our sleeping environment, we generally consider roads and railways to be harmful and disturbing for humans, birds and mammals alike. In a meta-analysis of the impacts of infrastructure on bird populations, Benítez-López *et al.* (2010) estimated a mean species abundance (MSA) of 0.678 (equivalent to 32% less birds) over a maximum effect distance of 2.6 km. Their dataset included 21 studies of the effect of roads, but none of railways. Popp & Boyle (2017) confirm this deficiency of railway studies, and advocate a long overdue development of a “Railway Ecology” field of R&D. They found only three railway-bird studies of which two targeted behaviour (Trembley & St. Clair 2009, Ge *et al.* 2011) and one abundances (Li *et al.* 2010). In this latter study, avian biodiversity and abundances along 150 m line transects were higher between the existing Qinghai railway and parallel highway in Tibet than transects 300-1200 m away. Our general perception of the damage done by railways on avifauna appears to lack supporting data.

During the planning of the Bothnia Line Railway (BLR), universities in northern Sweden were invited to propose follow-up studies. The department of Wildlife, Fish, and Environmental Studies (formerly the department of Forest Animal Ecology) of the Swedish University of Agricultural Sciences (SLU) proposed, and was granted a study of potential effects on birdlife. The author was assigned to design and conduct the breeding bird part of this study. We chose to focus on birds in agricultural landscapes, because agricultural land is a minority habitat in the boreal landscape through which the railway was to be built, and thus, its inhabitants prone to suffer relatively more from habitat loss and fragmentation than birds in the forest matrix. In addition, birds of agricultural landscapes (“farmland birds”) had shown a steeper decline than any other bird index in Sweden and Europe (Green et al. 2016, European Environment Agency 2018), and consequently were of conservation interest. Finally, bird species in this largely man-made habitat contribute significantly to the avian biodiversity of the region (Svensson *et al.* 1999, Olsson & Wiklund 1999, Ottosson *et al.* 2012).

In order to investigate the potential effects of the construction of the BLR on birdlife, we will apply a Before-During-After Control-Impact (BDACI) design, as recommended by Fahrig & Rytwinski (2009) and Roedenbeck *et al.* (2007). Due to the lengthy construction process, we will divide the *During* phase into two sub-phases corresponding to the true construction phase and the period between the construction and the onset of full-scale train traffic. Obviously, the timing and duration of these phases can not be controlled for the purpose this study.

We will address three aspects of the bird fauna: (1) avian biodiversity, (2) abundances of a selection of farmland bird species and (3) the spatial arrangement of their territories. For this, we will document (1) numbers of observed bird species, (2) numbers of mapped breeding territories and (3) nearest distances between territory midpoints to the railway or (for Control sites) an arbitrary baseline.

Because potential effects are expected to be expressed at various spatial scales, we will analyse the data for (a) the whole region along the 190 km railway (“regional scale”), (b) the local landscapes of individual study sites (“landscape scale”) and (c) the locations of breeding bird territories (“territory scale”). We will complement temporal trend analyses and comparisons between Impact and Control sites with mixed effect modelling of the four sequential phases of railway construction. We will test observed changes against the null hypothesis of no change.

## Methods and materials

### Study area and sites

The 190 km new Bothnia Line Railway (BLR) was built between Nyland (63.0°N, 17.8°E) and Umeå (63.8°N, 20.3°E) through the boreal forest landscapes along the Gulf of Bothnia in northern Sweden (Fig. 1). In the boreal landscape matrix of this region, patches of farmland were found on sediment soils in valleys and deltas. Within 10 km from the Bothnia Line Railway, farmed land covered 7% of area and forests 67%, while the remaining 26% were wetlands, lakes, rivers, infrastructure and settlements. Dairy production dominated the farming industry, and ley (sown perennial grasses) and spring-sown barley *Hordeum vulgare* were the main fodder crops.

**Figure 1.**
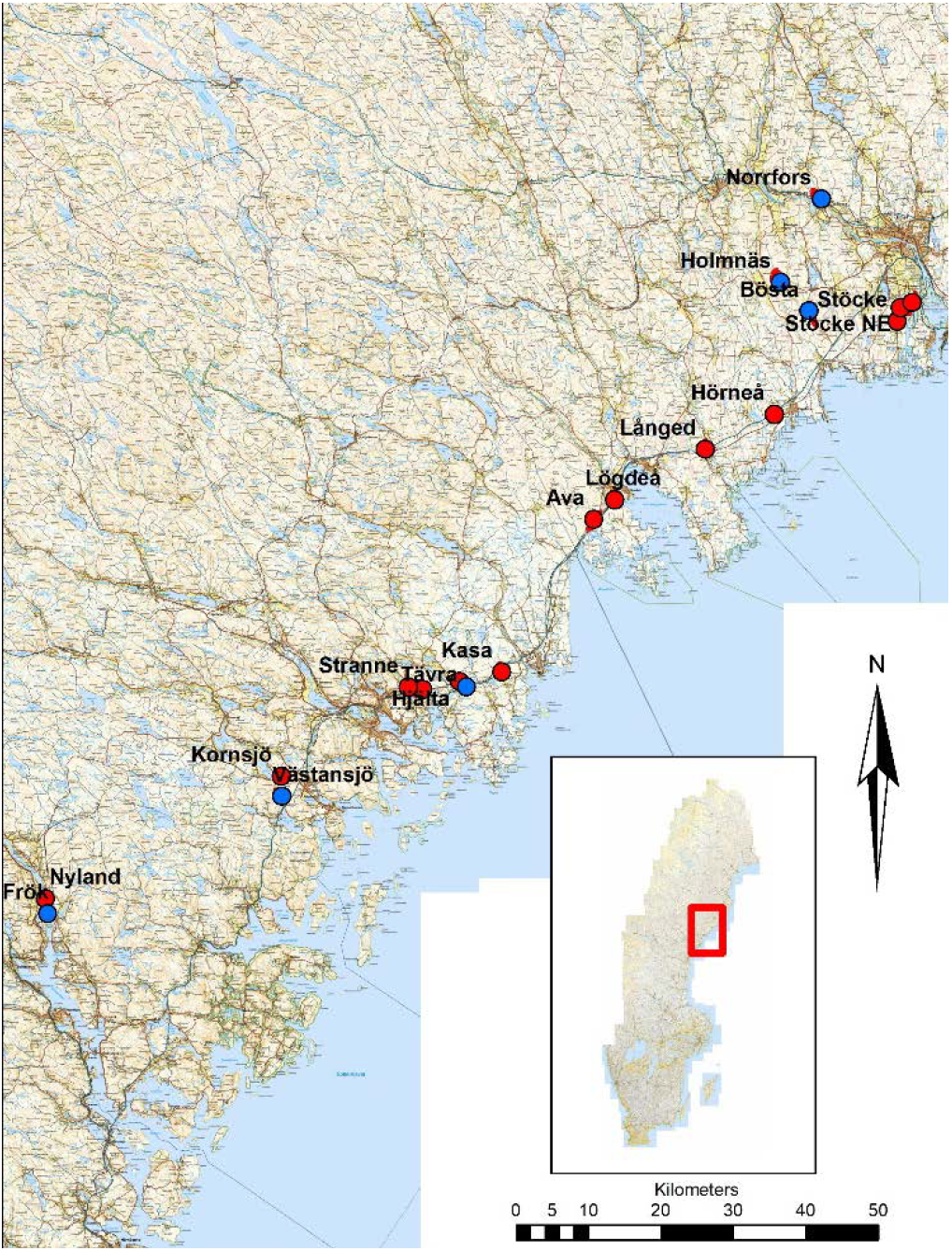
Location of study sites in the region along the Gulf of Bothnia in northern Sweden. Impact sites in red and Control sites in blue. The label of the Degernäs sites is blocked by the Stöcke label.

We included as Impact sites all 13 patches of agricultural land over which the BLR was built (Table 1). A few small (< 10 ha) patches were not included because (a) the precise trajectory of the BLR was initially unknown and (b) these small patches were expected to host low numbers of focal species. Preferably, we included the entire cohesive patch of agricultural land surrounded by other habitat, but we had to truncate large patches to fit the budget of the study (Table 1). More than 90% of the area of the Impact sites was within 800 m from the BLR, but distances up to 1725 m occurred (Fig. S1). The overall length of BLR transects crossing over agricultural land in Impact sites was 9360 m. For comparisons, we chose six resembling Control sites in nearby landscapes (Fig. 1). The total acreage was 983 ha for Impact sites and 731 ha for Control sites. All sites were in rural environments where various human activities took place alongside farming (Suppl. Mat. Shape-files for details).

**Table 1.** Location, delimitation, area, transect and nearest distance to the Bothnia Line Railway of Impact (I) and Control (C) sites. The sites Holmnäs and Norrfors were surveyed by Marianne de Boom, all others by the author.

| Site name |  | °N | °E | Delimitation<sup>1)</sup> | Area (ha) | Transect (m) | Distance BLR (km) |
| --- | --- | --- | --- | --- | --- | --- | --- |
| Nyland | I | 63.07 | 17.78 | P | 33.6 | 390 | 0 |
| Frök | C | 63.06 | 17.78 | P | 51.5 | - | 0.6 |
| Västansjö | C | 63.19 | 18.43 | P | 66.9 | - | 1.2 |
| Kornsjö | I | 63.22 | 18.44 | E | 82.8 | 400 | 0 |
| Stranne | I | 63.31 | 18.80 | E | 37.0 | 250 | 0 |
| Strandnyland | I | 63.31 | 18.84 | E | 69.2 | 890<sup>2)</sup> | 0 |
| Hjälta | I | 63.32 | 18.93 | P | 91.1 | 850 | 0 |
| Tävra | C | 63.31 | 18.96 | P | 57.1 | - | 0.1 |
| Kasa | I | 63.33 | 19.06 | E | 98.3 | 1200 | 0 |
| Ava | I | 63.51 | 19.34 | E | 109.5 | 1460 | 0 |
| Lögdeå | I | 63.54 | 19.39 | P | 67.4 | 380 | 0 |
| Långed | I | 63.59 | 19.66 | E | 34.2 | 200 | 0 |
| Hörneå | I | 63.62 | 19.87 | E | 48.9 | 600 | 0 |
| Stöcke | I | 63.73 | 20.23 | E | 208.4 | 1190 | 0 |
| Stöcke NE | I | 63.74 | 20.26 | E | 33.9 | 525 | 0 |
| Degernäs | I | 63.75 | 20.27 | P | 68.8 | 1025 | 0 |
| Bösta | C | 63.74 | 19.98 | P | 121.7 | - | 5.0 |
| Holmnäs | C | 63.79 | 19.90 | P | 248.7 | - | 12.6 |
| Norrfors | C | 63.89 | 20.03 | P | 185.0 | - | 18.8 |

1) E = the entire patch of farmland was included, P = the farmland patch was only partially included.
2) of which 410 m under ground

## Railway construction

The BLR was built in sections between 1999 and 2010 in a stepwise process with distinct subsequent phases: *Before*, *Construction*, *Ready* and *Traffic* (Fig. 2). The *Before* phase was the initial state of a study site before the construction of the railway started. The *Construction* phase encompassed the period during which the major ground- and construction works took place. These works included extensive digging, blasting and transportation, with large numbers of heavy machinery and people involved. Support roads, storage areas and temporary housing facilities were also used during this phase. The *Ready* phase was the period between the end of the *Construction* phase and the start of scheduled train traffic in the autumn of 2010. Low intensity works occurred during short periods of this *Ready* phase, e.g. the laying of the rails and the installation of wires and signal systems. During the summer of 2010, test-driving was performed. Although the BLR was officially inaugurated on the 28^th^ of August 2010, the survey of Impact sites was paused as a result of the significantly reduced timetable for 2011 and 2012. The survey was reactivated in 2013 for a three-year *Traffic* phase when train traffic had reached intended levels. This whole process created a complex temporal pattern with phases of different lengths and timing along the length of the BLR (Fig. 2).

**Figure 2.**
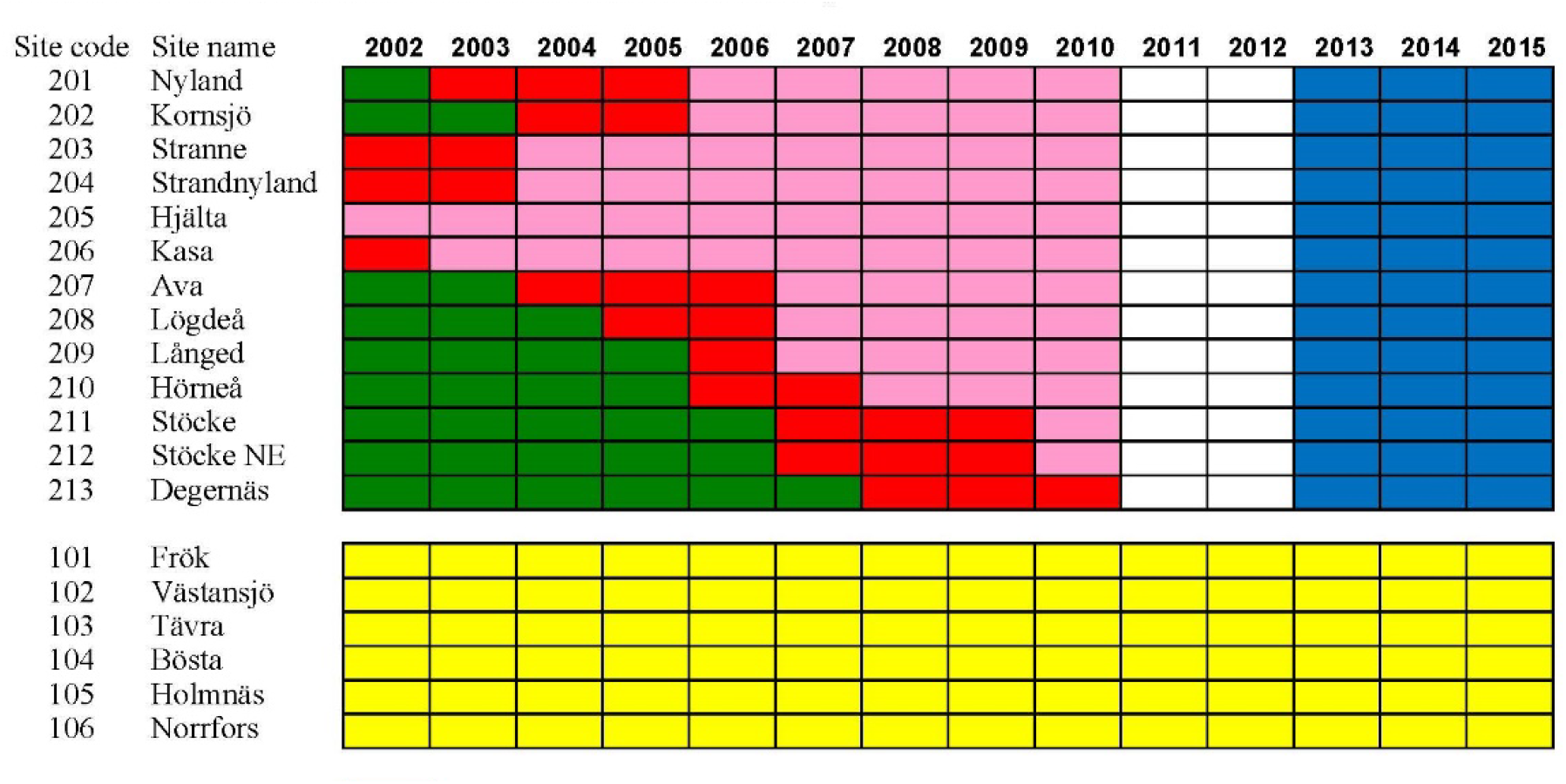
Construction schedule of the Bothnia Line Railway with *Before* (green), *Construction* (red), *Ready* (pink) and *Traffic* (blue) phases during the breeding season presented per site. Control sites marked yellow.

## Focal species

The study targeted avian biodiversity and a selection of focal species during the main breeding period May 1^st^ – July 15^th^. The focal species represent an array of life history traits (Table 2), and make up the majority of the farmland bird species in the region. Eurasian Curlew *Numenius arquata*, Eurasian Golden Plover *Pluvialis apricaria*, Eurasian Skylark *Alauda arvensis*, Meadow Pipit *Anthus pratensis* and Northern Lapwing *Vanellus vanellus* are ground-nesting birds, within the study sites confined to agricultural land. Common Rosefinch *Carpodacus erythrinus*, Red-backed Shrike *Lanius collurio*, Western Yellow Wagtail *Motacilla flava* and Whinchat *Saxicola rubetra* inhabit open places with shrubs and trees, including edges around the farmed fields. Barn Swallow *Hirundo rustica* and Common Starling *Sturnus vulgaris* are strongly associated with livestock husbandry and farmyards, while Common Snipe *Gallinago gallinago* and Green Sandpiper *Tringa ochropus* are found in wetland habitats in the agricultural landscape. Finally, Little Ringed Plovers *Charadrius dubius* breed and forage on patches of barren soil: tilled fields, gravel pits and construction sites. Most species feed on invertebrates, and so do nestlings and juveniles of herbivores. Due to the latitude of the study area, all the study species are strictly migratory, but their winter quarters range from NW Europe (e.g. Northern Lapwing) to southern tropical Africa (e.g. Barn Swallow). This implies that individual birds decide where to settle when they arrive in the breeding area in spring.

**Table 2.** Total number of territories in this study (N) and a schematic overview over some ecological features of the focal species (Cramp 1983, 1988 and 1993, Cramp & Perrins 1994a and 1994b, Svensson *et al.* 1999). Abbr. column presents the acronym for scientific names used in this text.

| Species | Abbr. | N | Habitat* | Nest site† | Food‡ | Migration¶ |
| --- | --- | --- | --- | --- | --- | --- |
| Barn Swallow | Hr | 1412 | P | B | I | T |
| Common Rosefinch | Ce | 66 | S | T | P | T |
| Common Snipe | Gg | 95 | W | G | S | E |
| Common Starling | Sv | 220 | P | T | I | E |
| Eurasian Curlew | Na | 1001 | F | F | S | E |
| Eurasian Golden Plover | Pa | 9 | F | F | I | E |
| Eurasian Skylark | Aa | 807 | F | F | I/P | E |
| Green Sandpiper | To | 46 | W | T | S | T |
| Little Ringed Plover | Cd | 21 | B | S | I | T |
| Meadow Pipit | Ap | 163 | F | G | I | E |
| Northern Lapwing | Vv | 997 | F | F | I | E |
| Ortolan Bunting | Eh | 14 | S | G | P | T |
| Red-backed Shrike | Lc | 17 | S | T | V | T |
| Western Yellow Wagtail | Mf | 342 | S | G | I | T |
| Whinchat | Sr | 548 | S | T | I | T |

\* B = barren terrain, F = farmed fields, P = pasture, S = shrubs and trees, W = wetland
† B = buildings, F = farmed fields, G = ground other than farmed fields, S = barren soil, T = trees and bushes
‡ I = invertebrates, P = plants and seeds (adults), S = soil living invertebrates, V = small vertebrates and insects
¶ E = short distance migrant (winters mainly in Europe), T = tropical migrant

## Data collection

The author and his wife Marianne de Boom did all the fieldwork. Both were experienced ornithologists and well familiar with the habitat, the species and the survey methods. The full-scale study started in 2002, after a pilot study in 2000 and 2001 which included 53% and 84% of the final number of sites, respectively. We surveyed sites yearly in 2002-2015, but Impact sites were not surveyed in 2011 and 2012. Mean effort across all sites was 189 h/year (sd. 8.8 h) (Fig. S2).

### Territory mapping

We surveyed the abundancies of the 15 focal species (Table 2) by standardized territory mapping based on four visits (Bibby *et al.* 2000, Svensson 2001, Sutherland 2006). By focusing on a limited number of focal species, we could allocate more attention to these species during fieldwork, and thus, improve the quality of the data. The author did all the interpretations from observations to territories. This process was initially performed after each breeding season, but the final interpretations were made after the 2015 season. This way, we avoided variation and drift in the interpretations.

### Avian biodiversity

During territory mapping, we also documented all the observed species for each visit and site. Observations were marked on a pre-print protocol and unusual species added manually. All species were included regardless of their position and activity in the landscape, i.e. a species flying over on migration was included as well as a local breeder. Before the analyses, we pooled the observations from the four yearly visits per site. In order to minimize potential observer bias, we only used observations made by the author; these data originated from all Impact sites and four out of the six Control sites.

### Territory midpoint distances

From the territory maps, we visually estimated the midpoint (point of gravity) of each territory. For known nest sites, we used the position of the nest as the territory midpoint. For Impact sites, we calculated the nearest distances between the territory midpoints and the Bothnia Line Railway (TMtR distances) in ArcGIS (ESRI, Redlands, Ca). Please note that this was done *a posteriori*, when the location of the railway track was known. For Control sites, we created a linear baseline just outside each study site and calculated the nearest distances between the territory midpoints and this baseline (TMtB distances).

## Statistical analyses

For clarity, we used capitalized variable and category names throughout the text (e.g. Year and Impact site), and described derived and composite variables in the context of their appearance. We marked the phases of the construction process in the Status variable with the level names in italic.

Because the number of data points was limited (N = 12 or N = 14), we used non-parametric Kendall’s rank correlation between Year and the relevant measurement in all temporal trend analyses. For comparisons between Impact and Control sites, we used Kendall’s rank correlation between Year and the numerical difference between yearly values of the measurements for these two types of sites. For transparency, we added plots for all trend analyses.

We used mixed effects modelling to unravel potential differences between the phases of railway construction under repeated measurements. For the analyses of TMtR distances, we used function lme in package nlme with Site as random factor. We compared models with Status (levels *Before*, *Construction*, *Ready* and *Traffic*) as fixed effect with models without a fixed effect. We then evaluated estimates of Status levels in relation to the intercept (usually Status *Before*).

Functions in packages nlme, lme4, GLMTMB and GLMM proved unable to handle the complex structure of the avian biodiversity and abundance data sets from an unbalanced design in combination with cases of empty treatment levels and low sample size. After extensive testing we chose to use the GLMMadaptive package (Rizopoulos 2019). For the analyses of avian biodiversity, we used function mixed_model (family = poisson) with Site as random factor. In addition to Status, we used fixed effects YearS (= calendar year – 2001) for an overall (linear) temporal trend and log(Effort/ mean Effort) for the effect of sampling effort. For the analyses of abundances, we used a very similar approach, but with Area instead of Effort. We chose these sets of variables after having compared a large number of ecologically/methodologically relevant variable combinations (including composite variables and interactions).

For statistical analyses and plotting, we used R version 3.5.3 (64 bit) (R Development Core Team 2018) with packages nlme v 3.1-137 (Pinheiro *et al.* 2018), AICcmodavg v. 2.2-2 (Mazerolle 2019), GLMMadaptive v. 0.5-1 (Rizopoulos 2019) and dependencies. We used Microsoft Office Excel 2013 for initial data handling and complementary plotting. Finally, we used ArcGIS 10.6 for (1) mapping study sites and railway & baseline trajectories, (2) generating random points and (3) measuring nearest distances.

## Results

### Avian Biodiversity

In total 169 species were observed during the study, and the yearly numbers of observed species increased over the course of the study period (Kendall rank correlation tau = 0.45, z_11_=2.00, *P*<0.05, Fig. S3). The numbers decreased for the four Control sites (tau = −0.25, z_13_=-1.22, *P*>0.05) and increased for the 13 Impact sites (tau = 0.56, z_11_=2.45, *P*<0.05) (Fig. 3). Differences in average numbers between Impact and Control sites also increased (tau = 0.70, T_11_=56, *P*<0.001), but there was no significant correlation between yearly numbers in Impact and Control sites (tau = 0.14, z_11_ = 0.57, *P* >0.05). For the Impact sites combined, the increase in avian biodiversity between 2002 and 2015 was c. 10 species.

**Figure 3.**
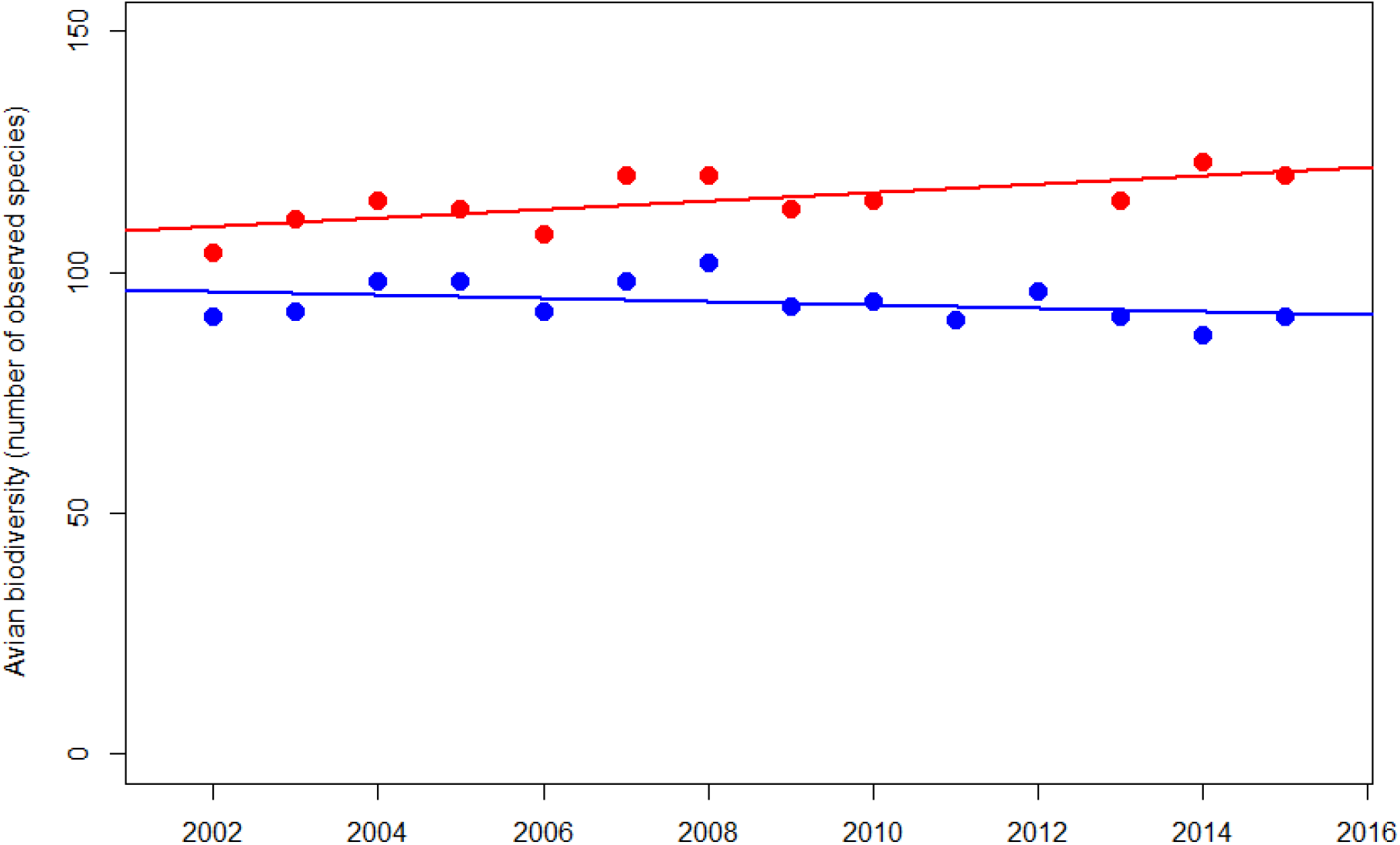
Yearly numbers of bird species observed in 13 Impact sites (red) and four Control sites (blue). Linear regression lines added for visualization.

Sites differed in total number of species (ANOVA F_1_=9.2, Pr(>F) = 0.003, Fig. S4), but none of them showed a significant temporal trend in numbers of observed species (Table 3, Suppl. Mat. AB1).

**Table 3.**
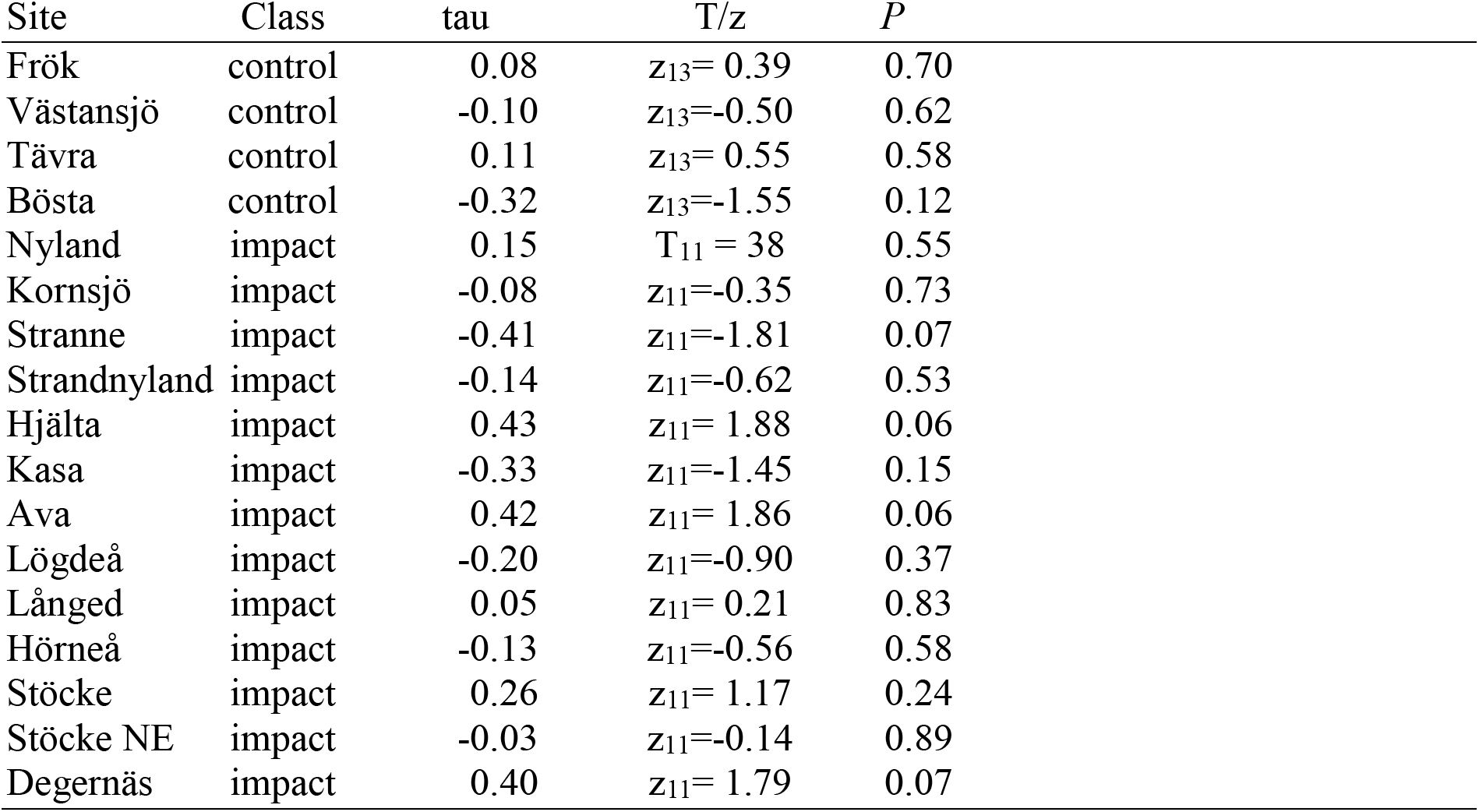
Kendall rank-order correlation test statistics for analyses of yearly numbers of observed bird species per site over the study period 2002-2015. Impact sites were not surveyed in 2011 and 2012.

The Status variable improved the GLMMadaptive generalized mixed effects model (ΔAIC = −12, ANOVA LR_5_ = 21.8, *P* < 0.001), but none of the subsequent phases were significantly different from Status = *Before* (Table 4). The lack of consistent differences between the phases of the railway construction process was also expressed in Table S1 and visualized in Fig. 4.

**Table 4.** Model estimates output for the Poisson mixed effects model (function glmer) including Status, YearS^1)^ and log(NEffort)^2)^ as fixed effects and Site as random effect.

| Fixed effect variables and levels | Estimate | Std.Err | z-value | p-value |
| --- | --- | --- | --- | --- |
| Status Before (Intercept) | 4.05 | 0.040 | 101.40 | < 0.001 |
| Status Construction | 0.02 | 0.040 | 0.42 | 0.674 |
| Status Ready | 0.03 | 0.048 | 0.61 | 0.542 |
| Status Traffic | -0.01 | 0.082 | -0.07 | 0.945 |
| YearS | -0.00 | 0.007 | -0.05 | 0.961 |
| log(NEffort) | 0.33 | 0.054 | 6.07 | < 0.001 |

<sup>1)</sup> YearS = Year - 2001
<sup>2)</sup> NEffort = Effort / mean(Effort)

**Figure 4.**
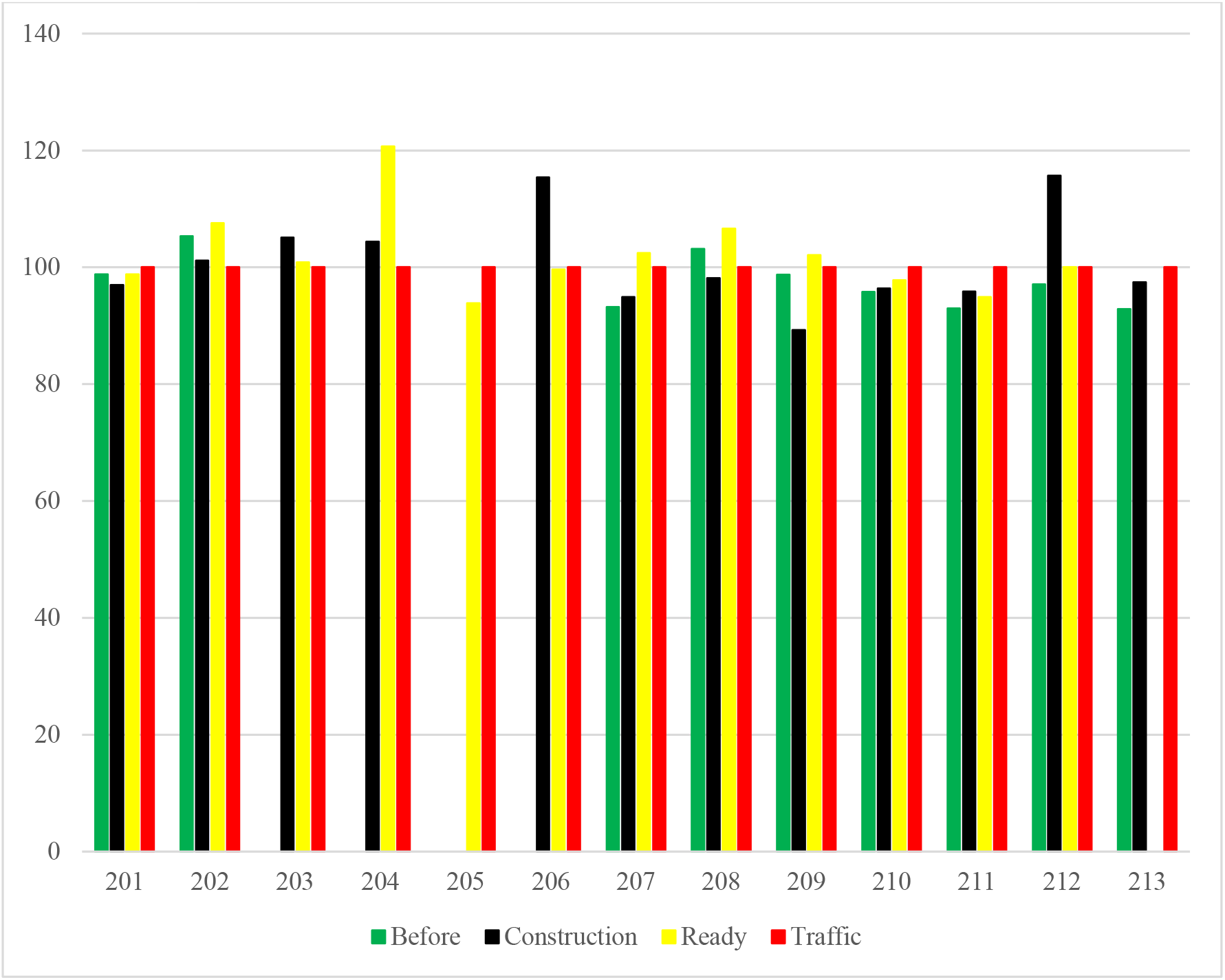
Avian biodiversity index (*Traffic* = 100) for all 13 Impact sites (#201-213) during the various phases of the railway construction process.

## Abundances

The numbers of territories in all 19 sites (N = 5758) decreased over the course of the study period (tau = −0.55, T_11_ = 15, *P*<0.05), and this negative trend was expressed in both Control sites (tau = −0.55, T_11_ = 15, *P*<0.05) and Impact sites (tau = −0.50, z_11_ = −2.27, *P*<0.05) (Fig. 5). The similarity between the trends for Control and Impact sites is shown by the absence of a temporal trend in the numeric differences between these types of sites (tau = −0.14, z_11_ = −0.62, *P*>0.05).

**Figure 5.**
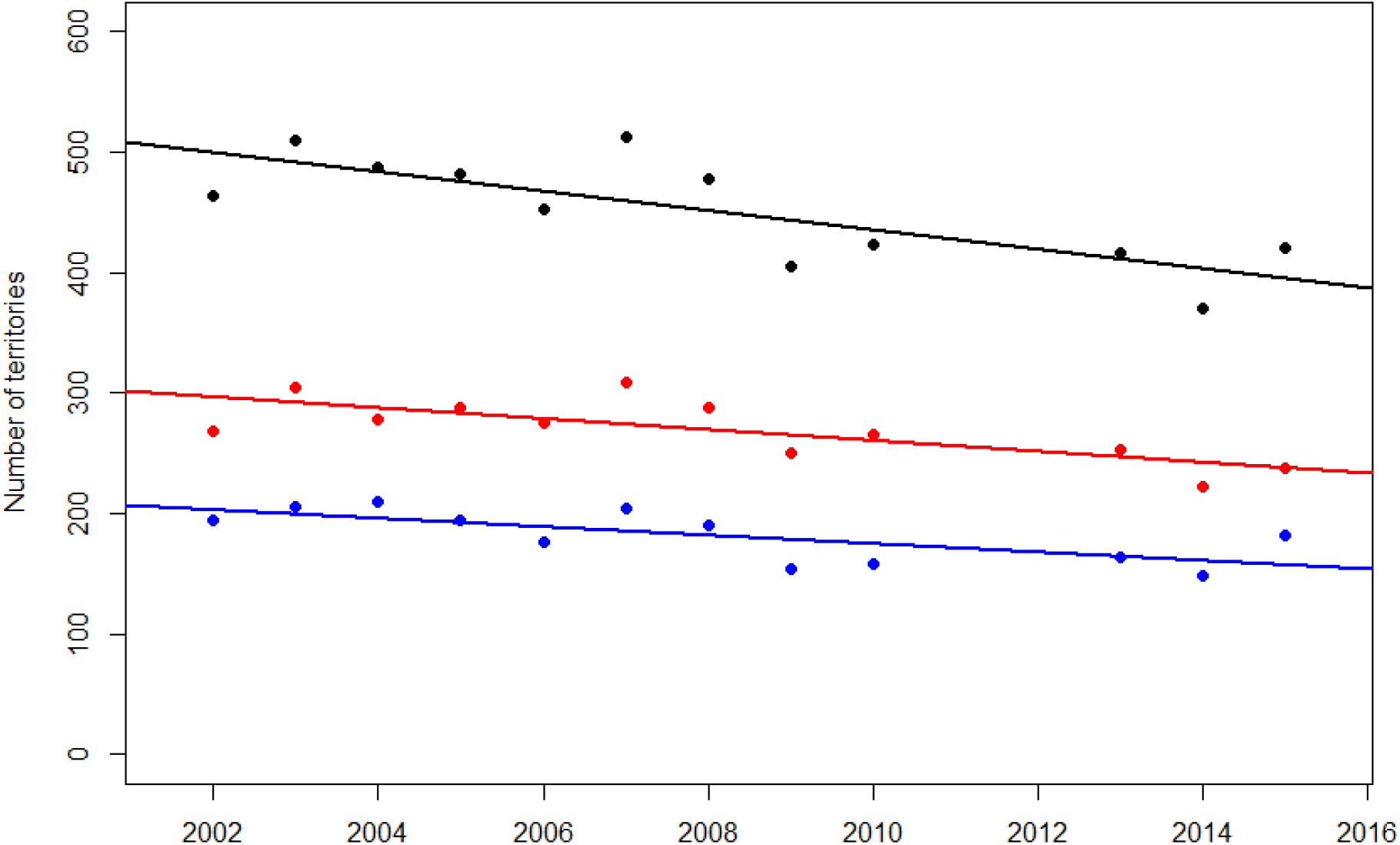
Numbers of territories of all study species in Impact sites (red), Control sites (blue) and all sites combined (black). Simple linear regressions lines in matching colours added for visualization.

Trends varied among species (Table 5, Suppl. Mat. Terr1), but negative trends prevailed. Eurasian Curlew, Eurasian Skylark and Western Yellow Wagtail showed significant negative trends throughout, in harmony with their national trends (Green et al. 2016). No species showed consistently positive trends, but Common Snipe and Whinchat came close (Table 5). In Meadow Pipit, the negative trend in Control sites (in line with the national trend) was absent in Impact sites. Territory numbers of Little Ringed Plover and Common Rosefinch decreased in Impact sites but not in Control sites. For Common Rosefinch, this trend reflects the national trend (Green *et al.* 2016). For the Little Ringed Plover, the linear trend does not account for the complete absence of this species prior to the construction of the railway (Status = *Before*) and the trend merely expresses the local extinction after the boost in numbers as a result of construction works (Suppl. Mat. Terr2 for site-level details).

**Table 5.** Temporal trends for numbers of territories of 14 study species in All, Control and Impact sites, expressed as Kendall rank correlation tau coefficients. Significance levels: ^NS^ = P>0.05, * = P < 0.05, ** = P < 0.01 and *** = P < 0.001. (Suppl. Mat. Terr1 for details and plots). National trends for 1998-2015 from Green *et al.* (2016).

| Species | All | Control | Impact | National trend (%/year) |
| --- | --- | --- | --- | --- |
| Barn Swallow | 0.20<sup>NS</sup> | 0.28<sup>NS</sup> | -0.06<sup>NS</sup> | 1.5*** |
| Common Rosefinch | -0.58* | 0.20<sup>NS</sup> | -0.58* | -4.3*** |
| Common Snipe | 0.45<sup>NS</sup> | 0.51* | 0.35<sup>NS</sup> | 1.1** |
| Common Starling | -0.36<sup>NS</sup> | -0.38<sup>NS</sup> | -0.10<sup>NS</sup> | -3.3*** |
| Eurasian Curlew | -0.71*** | -0.76*** | -0.63*** | -2.0*** |
| Eurasian Golden Plover<sup>1)</sup> | NA | NA | NA | 1.1* |
| Eurasian Skylark | -0.74*** | -0.74** | -0.79*** | -1.7*** |
| Green Sandpiper | -0.61** | -0.31<sup>NS</sup> | -0.32<sup>NS</sup> | 3.8*** |
| Little Ringed Plover | -0.48* | -0.36<sup>NS</sup> | -0.48* | 1.6<sup>NS</sup> |
| Meadow Pipit | -0.22<sup>NS</sup> | -0.46* | 0.38<sup>NS</sup> | -1.2*** |
| Northern Lapwing | -0.02<sup>NS</sup> | 0.09<sup>NS</sup> | -0.09<sup>NS</sup> | -0.7<sup>NS</sup> |
| Ortolan Bunting | -0.17<sup>NS</sup> | 0.08<sup>NS</sup> | -0.33<sup>NS</sup> | -7.1*** |
| Red-backed Shrike | 0.29<sup>NS</sup> | 0.17<sup>NS</sup> | 0.19<sup>NS</sup> | -0.2<sup>NS</sup> |
| Western Yellow Wagtail<sup>2)</sup> | -0.60** | -0.56* | -0.57* | -0.5<sup>NS</sup> |
| Whinchat | 0.35<sup>NS</sup> | 0.49* | 0.25<sup>NS</sup> | -1.5*** |

<sup>1)</sup> Eurasian Golden Plover only occurred in a single Control site.
<sup>2)</sup> National trend given for Western Yellow Wagtail subspecies *M. f. thunbergi* (the dominant subspecies in northern Sweden), but currently, the study area is in an intermediate zone where *thunbergi*, *flava* and hybrids coexist. The national trend for *M. f. flava* was 5.0\*\*\*.

Analyses of numbers of territories per site (landscape level) resulted in negative trends in a majority of cases with the trend for Degernäs as a striking exception to the rule (Table 6, Suppl. Mat. Terr2).

**Table 6.**
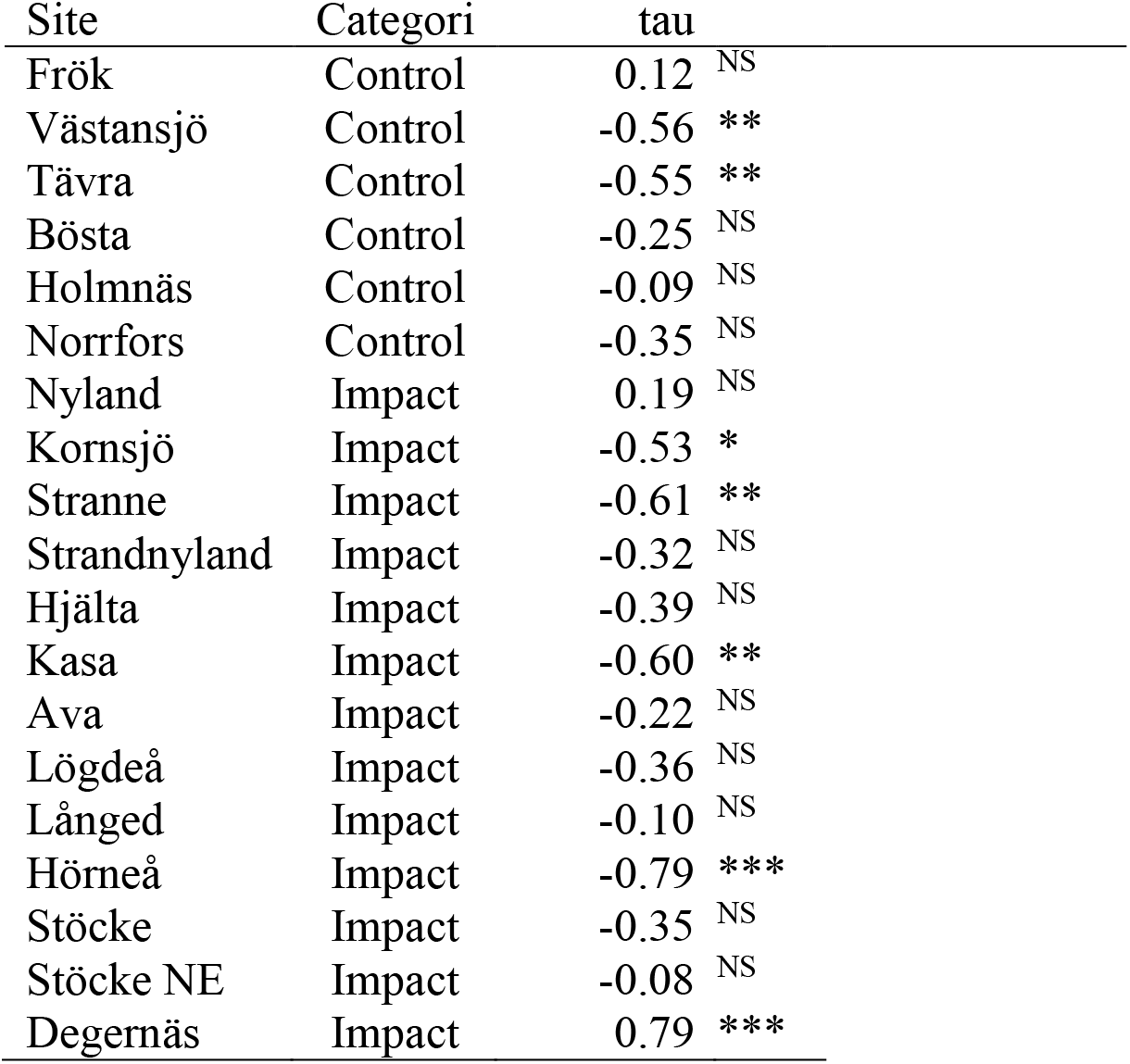
Kendall’s rank correlation tau for numbers of territories per study site. Significance levels: ^NS^ = P> 0.05, * = P < 0.05, ** = P < 0.01 and *** = P < 0.001. (Suppl. Mat. Terr2 for details and plots).

Split up to species level, low numbers of territories per site prevented relevant trend analyses in many cases (Table 7). Negative correlations were expressed in 67% of the tests with *P*<0.05 tau estimates; in Control sites 79% and in Impact sites 63%. These numbers indicate that, overall, the species trends were negative, but not more severely so in Impact sites than in Control sites.

**Table 7.**
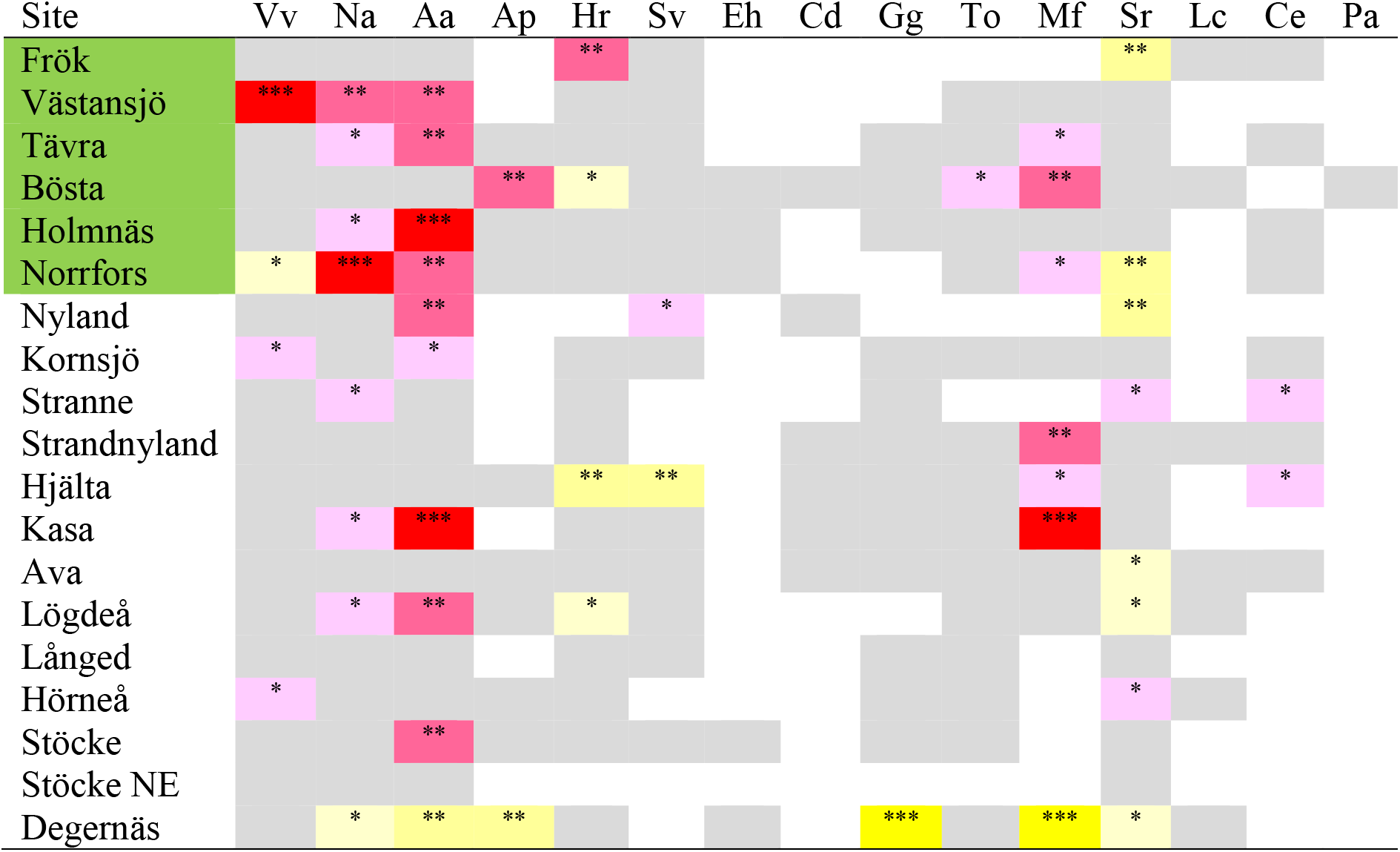
Summary of temporal trends per site and per species expressed as P-value classes of Kendall rank correlation tests between number of territories and year. Positive Kendall’s rank number correlations in yellow and negative ones in red. Grey background = no significant correlation. * = P<0.05, ** = P < 0.01 and *** = P < 0.001. Empty cells for cases where low numbers of territories did not allow for correlation tests. Table 2 for species name abbreviations.

The trends for Eurasian Curlew (Na), Eurasian Skylark (Aa) and Western Yellow Wagtail (Mf) were negative in many sites, 37%, 47% and 50% of sites with valid correlation models, respectively. A high proportion (32%) of positive models was found in Whinchat (Sr) (Table R6). Table 7 also contains trend evaluation of the Eurasian Golden Plover (Pa), a species that only occurred in a single Control site (Suppl. Mat. Terr2).

The generalized mixed effects model results (Table 8) showed that, under the conditional mean, numbers of territories of all species combined were higher during the *Construction*, *Ready* and *Traffic* phases than before the onset of railway construction (Status *Before*). Effects of survey year (calendar year – 2001) and log(normalized site area) were also significant (negative and positive, respectively) and thus, were important for model improvement.

**Table 8.** Model estimates output for the Poisson generalized mixed effects model (function GLMMadaptive) including Status, YearS^1)^ and log(NArea)^2)^ as fixed effects and Site as random effect for numbers of territories for all focal species in Impact sites.

|  | Estimate | Std.Err | z-value | p-value |
| --- | --- | --- | --- | --- |
| Status *Before* (Intercept) | 2.96 | 0.135 | 22.01 | < 0.001 |
| Status *Construction* | 0.13 | 0.065 | 2.07 | < 0.05 |
| Status *Ready* | 0.28 | 0.083 | 3.33 | < 0.001 |
| Status *Traffic* | 0.28 | 0.137 | 2.04 | < 0.05 |
| YearS | -0.04 | 0.011 | -3.21 | < 0.01 |
| log(NArea) | 1.59 | 0.236 | 6.77 | < 0.001 |

<sup>1)</sup> YearS = Year - 2001
<sup>2)</sup> NArea = Area / mean(Area)

For most species, equivalent GLMMadaptive models did not show significant (*P* < 0.05) differences between Status classes, but for Eurasian Skylark numbers were higher during *Construction* and for Western Yellow Wagtail numbers were higher during all subsequent phases compared with *Before* (Suppl. Mat. Terr3).

## Territory positions

For the 13 Impact sites, the overall median TMtR distance was 364 m (N = 3245), and over the study period, the yearly median TMtR distance decreased (tau = −0.48, T_11_ = 17, *P*<0.05) (Fig. 6). For the six Control sites, the overall median TMtB distance was 806 m (N = 2520) and did not change significantly over time (tau = 0.20, z_13_ = 0.97, P>0.05) (Fig. 6).

**Figure 6.**
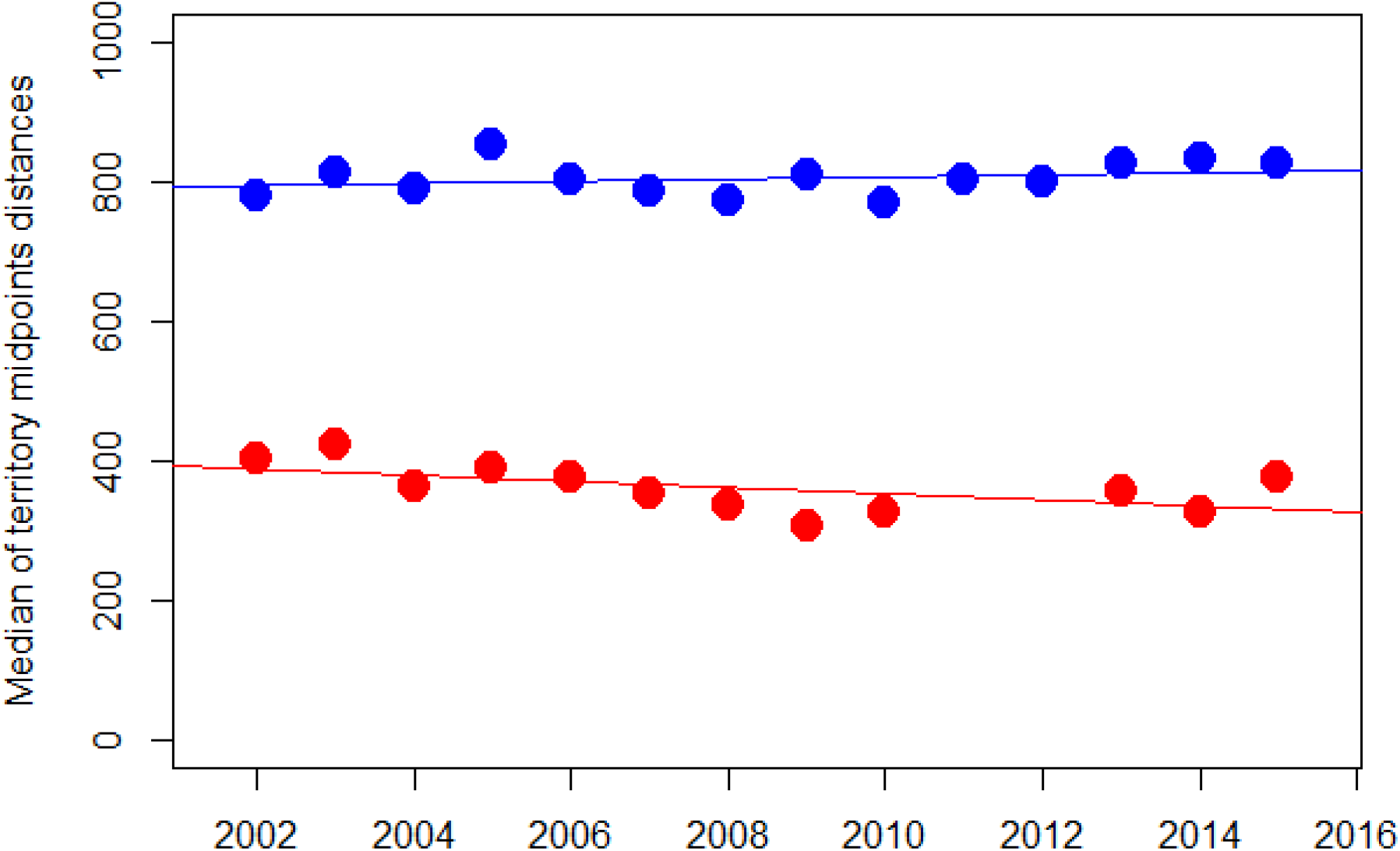
Yearly median distances from territory midpoints to the railway for Impact sites (red) and to an arbitrary baseline for Control sites (blue). Linear regression lines added for visualization.

At species level, Western Yellow Wagtail and Whinchat showed decreasing TMtR distances, indicative of an attraction towards habitats near the railway. In all other species, no significant temporal change could be detected (Suppl. Mat. Dist1). In Control sites, the only significant change in TMtB distances occurred in Eurasian Skylark (increase) (Suppl. Mat. Dist1).

Kendall correlation tests for individual species and Impact sites revealed four (5 %) valid tests where yearly values increased over time and equally many (5 %) where these decreased (Table S2). The vast majority (90 %) of 83 cases for which correlations coefficients could be calculated revealed no significant change over the study period.

The distribution of territory midpoint positions in relation to the BLR was similar to the distribution of random points (Fig. 7). More than 90 % of the territory midpoints and the random points were located within 800 m from the railway (Fig. 8, Suppl. Mat. Dist2 for separate study species). Positions of territory midpoints in the 0 - 200 m distance range and beyond 800 m appeared to be underrepresented, and distances between 400 and 750 m overrepresented (Fig. 7). Complete depletion of territories within 100 m from the railway did not occur, except for Little Ringed Plover, Ortolan Bunting and Common Rosefinch, but these species completely disappeared from the entire study area. The Meadow Pipit showed a marked increase in numbers of territories within 100 m from the railway once train traffic had started (Suppl. Mat. Dist2).

**Figure 7.**
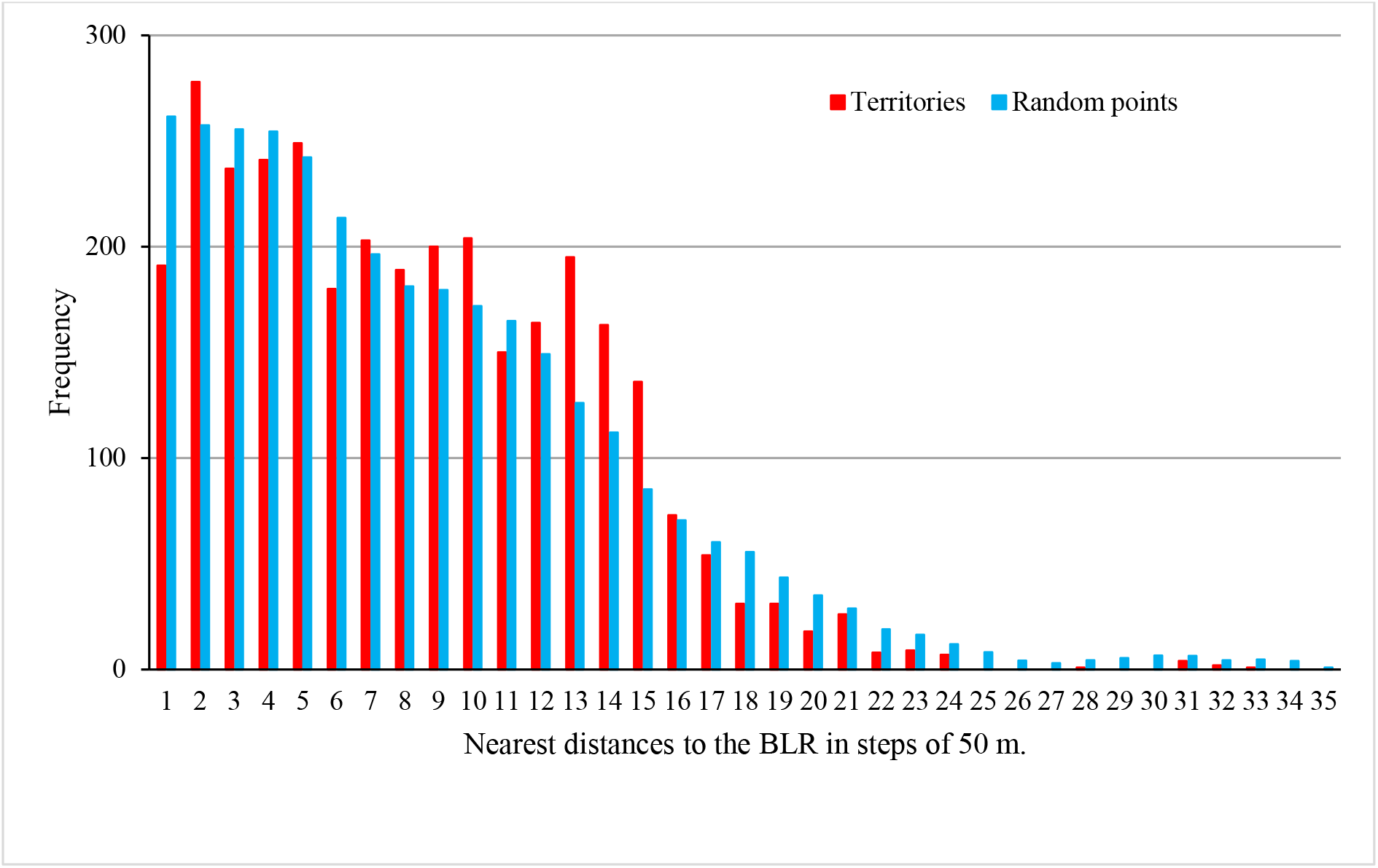
Frequency distribution of nearest distances from territory midpoints (all species combined) and random points to the Bothnia Line Railway. Total frequencies are harmonized.

**Figure 8.**
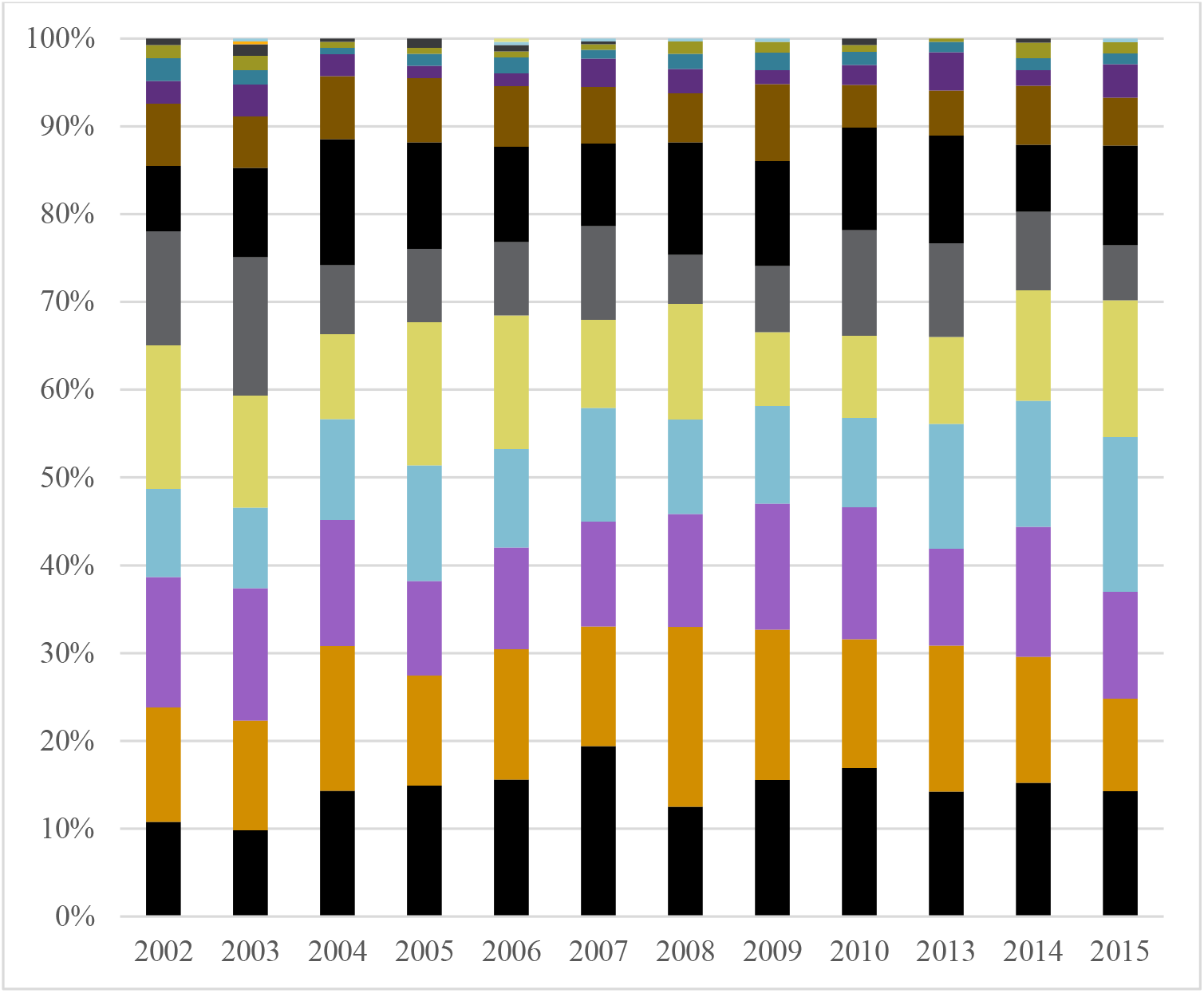
Yearly percentage of territory midpoints for all focal species within 100 m zones from the BLR; from shortest (0 – 100 m) distances at the base of the columns to zone 17 (1600 – 1700 m) on top.

Focusing on the temporal pattern of the proportion of territory midpoints **near** the railway (within 300 and 500 m, respectively) for individual species, a decreasing proportion was only found in Eurasian Curlew (500 m), indicative of shying away from the railway. For Meadow Pipit (300 m) and Western Yellow Wagtail (300 and 500 m) the proportion of territory midpoints near the railway increased over the study period, indicating a preference for habitats close to the railway. For all other species, no significant changes could be detected (Suppl. Mat. Dist3 for details).

Visual inspection of per site distances of individual species revealed a complex pattern of responses to the *Construction*, *Ready* and *Traffic* phases (Fig 9, Suppl. Mat. Dist4). For many species, numbers per site were too small for trend analysis; especially for Ortolan Bunting and Common Rosefinch, which were in widespread strong decline during the study period (Table 5, Green et al. 2016). Occurrences of Green sandpiper, Little Ringed Plover and Red-backed Shrike were almost entirely linked to the Construction and Ready phases of the railway construction process, which is in line with the ecology of these species.

**Figure 9.**
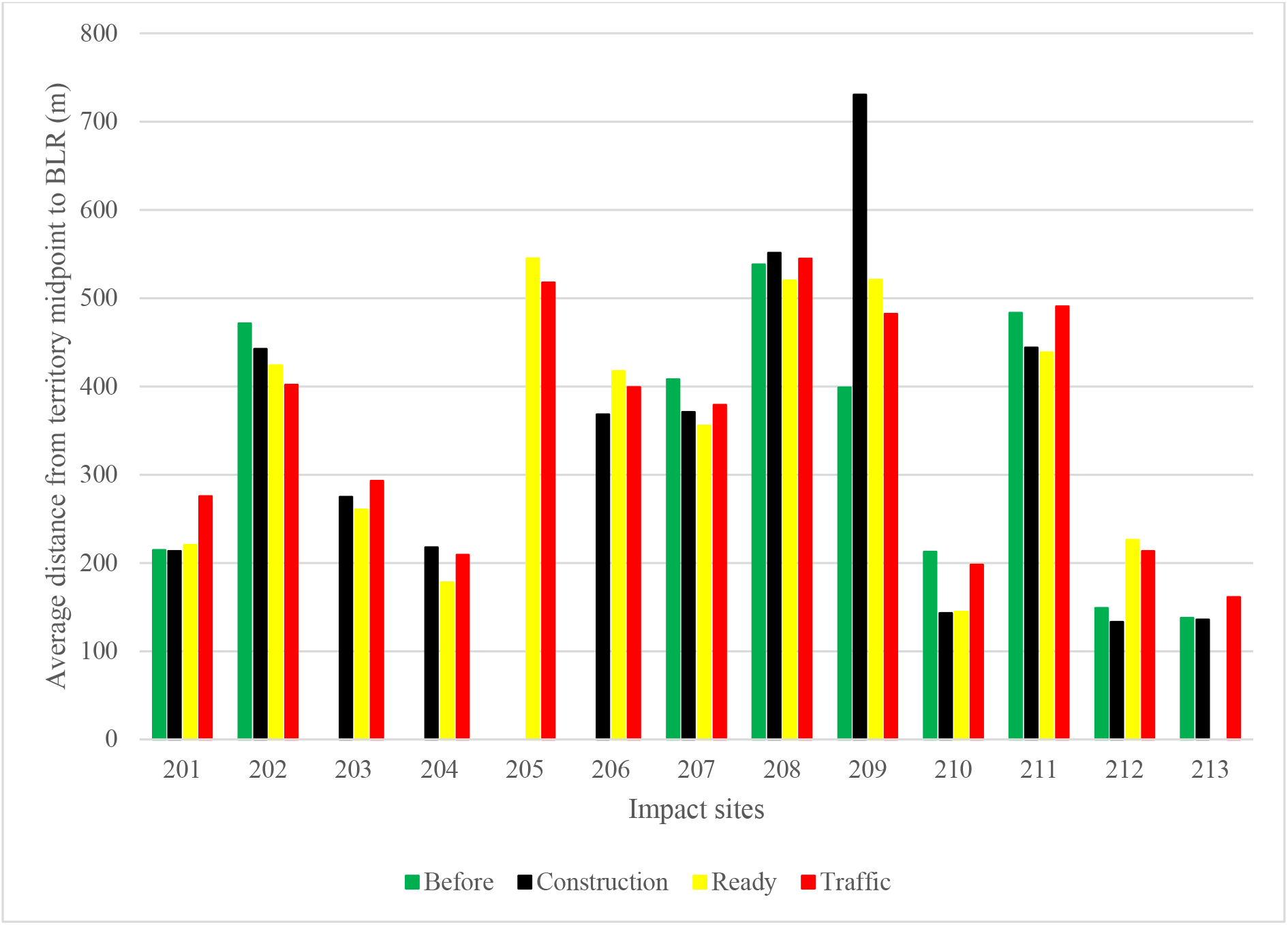
Average TMtR distances under subsequent phases of the railway construction process for all species in Impact sites 201-213. Fig. 2 for site codes.

The empty mixed effect model for TMtR distances of all species combined (lme formula: fixed = response ~ 1, random = ~ 1|Site) had AICc = 44425. Adding the Status variable as fixed effect improved the model (ΔAICc=-18), but none of the phases did differ significantly from the conditioned mean of distances on Status *Before* (the intercept) (Table 9). At species level, significant (*P* < 0.05) effects of *Construction*, *Ready* and/or *Traffic* phases of the BLR construction process were only expressed in mixed effects models for Common Starling, Western Yellow Wagtail and Whinchat (Suppl. Mat. Dist5). In all cases, this effect showed a decrease in TMtR distances relative distances observed before the construction process started; the territories moved closer to the railway trajectory.

**Table 9.**
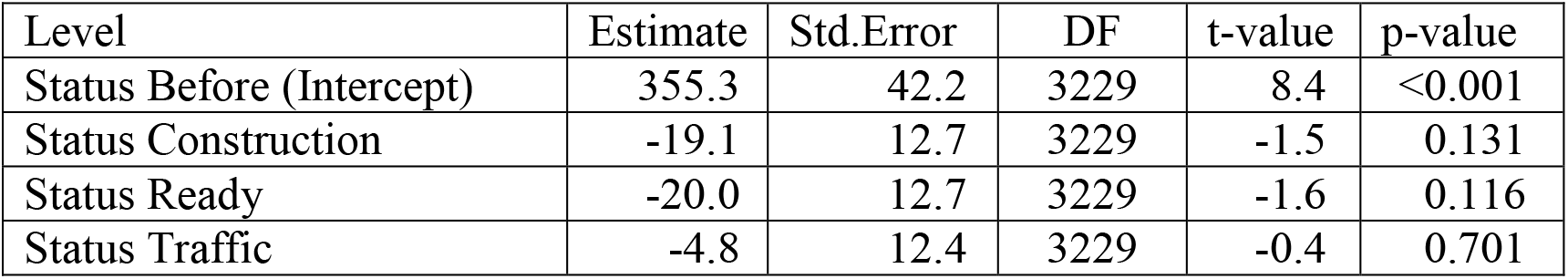
Model estimates output for the Poisson mixed effects model (function lme) including Status as fixed effect and Site as random effect for all TMtR distances in Impact sites.

## Conclusion

The results of this study do not support the perception of a widespread detrimental effect of railways *sensu* Benítez-López et al. (2010).

Avian biodiversity did not decrease, neither at regional nor at landscape level. Compared with Control sites, the numbers of observed species in Impact sites increased over the study period, but this positive trend was not expressed in the generalized mixed effects analyses of the railway construction process.

Numbers of territories did decrease, but similarly in Impact and Control sites at regional and landscape scale. This decrease was largely driven by negative trends in Eurasian Curlew, Eurasian Skylark and Western Yellow Wagtail. Trends for individual species in individual sites showed there were winners and loosers, but mainly neutrals. This variable pattern needs to be interpreted in the light of large-scale (national) trends, ecological traits and site conditions. The development of abundances in the Degernäs site demonstrates that successful mitigation is possible even for a category of birds in widespread decline. A major compensation programme in favour of birds in open landscapes was implemented in and around this site.

Focal species in Impact sites moved their territories closer to the BLR, but the inhabitants of Control sites did not change the position of their territories in relation to an arbitrary baseline. This tendency of decreasing distances to the BLR was also expressed in the mixed effects analyses; territories were significant closer to the railway during *Construction*, *Ready* and *Traffic* phases than *Before*. Avoidance of areas near the BLR (< 500 m) was detected only in a few species and specific sites.

In a single-site Before-After study of the construction of a combined motorway and railway outside Helsinki, Finland, Yrjölä & Santahurja (2015) found increasing numbers of territories of birds, and a tendency to cluster near the new motor/railway. From this, they concluded that the construction project had had a positive effect on breeding birds. The study of Li et al. (2010) also indicates a positive effect of a combined motor/railway on the ground-dwelling birds in their study. I am reluctant to proclaim that the building of the Bothnia Line Railway was positive for breeding birds in agricultural landscapes, but it is safe to say that our results do not support the hypothesis of negative effects. I strongly suggest further studies of the effects of railways on birdlife, BDACI studies in particular (c.f. Smith 2002). I also support Popp & Boyle’s appeal for the rise of “Railway Ecology”.

A manuscript with full background and discussion sections is under preparation and will be submitted for scientific publication asap.

## Supporting information

Avian biodiversity data

Territory midpoint to baseline distances

Territory midpoint to BLR distances

Random point distances Control sites

Rando point distances Impact sites

Territorty numbers data

Trends in avian biodiversity

Trends in TM distances Impact vs Control

Trends in proportions of distances to BLR

Trends in distances in 300 and 500 m zones to BLR

Trends of distances to BLR per site and species

Mixed effects model output for TM to BLR distances

Trends of numbers of territories for Control, Impact and all sites

Trends of territory numbers per site och species

Mixed effects model output for territory number data

All shape files

## Acknowledgements

I want to thank Marianne de Boom for taking part in the fieldwork and Professor emeritus Kjell Sjöberg for his role in the initiation of this project. Thanks also to all the farmers, inhabitants and railway workers who tolerated or even supported us while working in their environment. Last but not least, I want to thank Jan Olof Helldin, Swedish Biodiversity Centre CBM/SLU, for inviting me to TRIEKOL, and for his support and interest.

This study received financial support from the Swedish Transport Administration. This support was unconditioned in terms of study design and methodology. The final analyses and writing-up process was financed by TRIEKOL III.

## Supplementary material

**Supplementary figures and tables**

Supplementary figures and tables referred to in the main text.

**Supplementary material Data**

Data files containing the original data on avian biodiversity, numbers of territories of focal species and territory midpoint distances (including random point distances).

**Supplementary material Shape-files**

Shape files for study sites, arbitrary baselines and relevant parts of the Bothnia Line Railway.

**Supplementary material AB1**

Yearly numbers of observed species per site, colour coded for Status classes *Before*, *Construction*, *Ready* and *Traffic*.

**Supplementary material Terr1**

Temporal trends for numbers of territories of individual study species in Impact sites, Control sites and all sites combined.

**Supplementary material Terr2**

Temporal trends per study site for numbers of territories of individual species and all species combined.

**Supplementary material Terr3**

Mixed effects model output for numbers of territories of individual species across the phases of the railway construction process.

**Supplementary material Dist1**

Temporal trends in annual mean distances between territory midpoints and an arbitrary baseline (Control sites) or the BLR (Impact sites).

**Supplementary material Dist2**

Temporal patterns of numbers and proportions (%) of territory midpoints within 100 m zones from the Bothnia Line Railway. All study species combined and individual species.

**Supplementary material Dist3**

Temporal trends and Kendall’s rank correlation tests for the proportions of territories of individual study species within 300 and 500 m from the Bothnia Line Railway.

**Supplementary material Dist4**

Temporal trends for distances between territory midpoints and the Bothnia Line Railway per site and per species.

**Supplementary material Dist5**

Mixed effect modelling for distances between territory midpoints and the Bothnia Line Railway across phases of railway construction for all species combined and for individual species.

