## Supplementary material for "Are railways detrimental to bird populations? A BDACI study on the construction of the Bothnia Line Railway": Trends in avian biodiversity

Supplementary material **AB1**

### Yearly numbers of observed species per site, colour coded for Status classes *Before* (green), *Construction* (blue), *Ready* (purple) and *Traffic* (red)

Most correlation tests generated warning messages due to ties (data points with the same Y-value). The warning texts were omitted from the output text for readability, but the tau value estimates marked with a * character. For models with ties, exact P-values could not be calculated and the estimated P-values should be treated with caution.

Contents


| Site | page |
| --- | ---: |
| Nyland | 2 |
| Frök | 3 |
| Västansjö | 4 |
| Kornsjö | 5 |
| Stranne | 6 |
| Strandnyland | 7 |
| Hjälta | 8 |
| Tävra | 9 |
| Kasa | 10 |
| Ava | 11 |
| Lögdeå | 12 |
| Långed | 13 |
| Hörneå | 14 |
| Stöcke | 15 |
| Stöcke NE | 16 |
| Degernäs | 17 |
| Bösta | 18 |

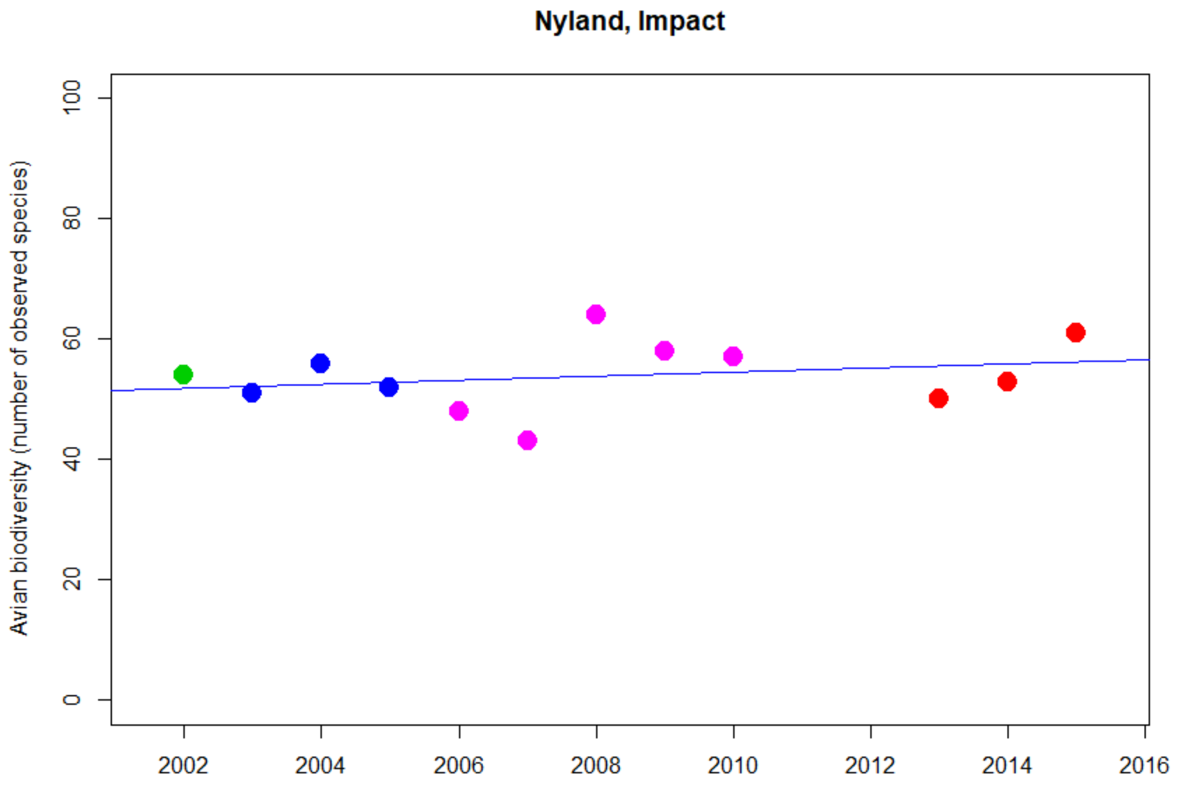


```
Kendall's rank correlation tau = 0.152
T = 38, p-value = 0.545
```

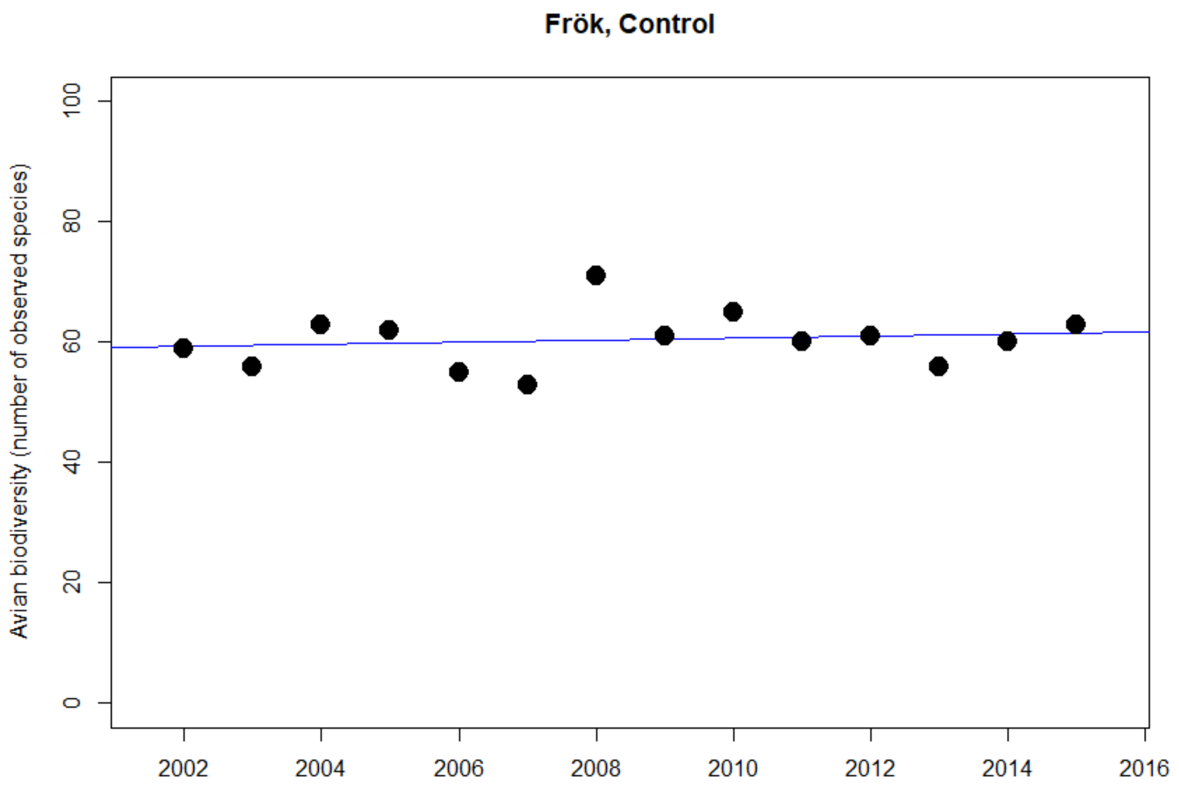


```
> cor.test(SFrok$Year,SFrok$Arter, method="kendall")

Kendall's rank correlation tau = 0.079
z = 0.38553, p-value = 0.700 (estimated due to ties)
```

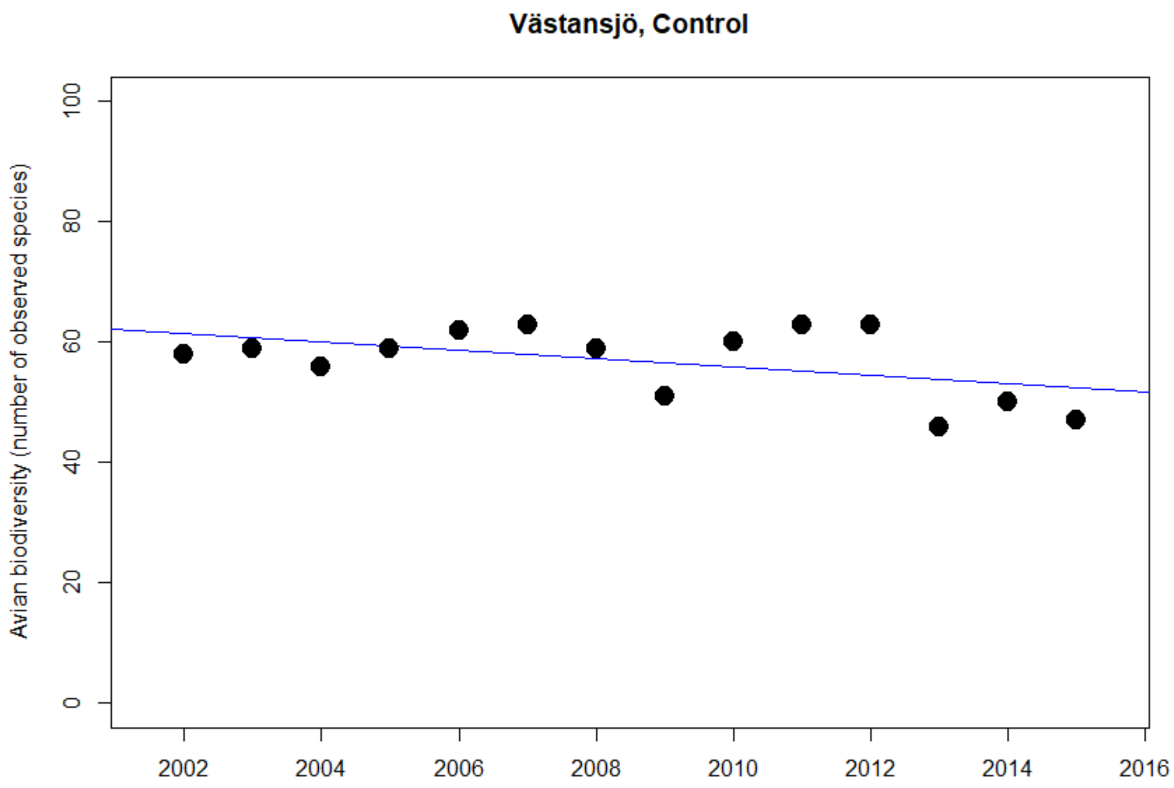


```
Kendall's rank correlation tau = -0.102
z = -0.49821, p-value = 0.618 (estimated due to ties)
```

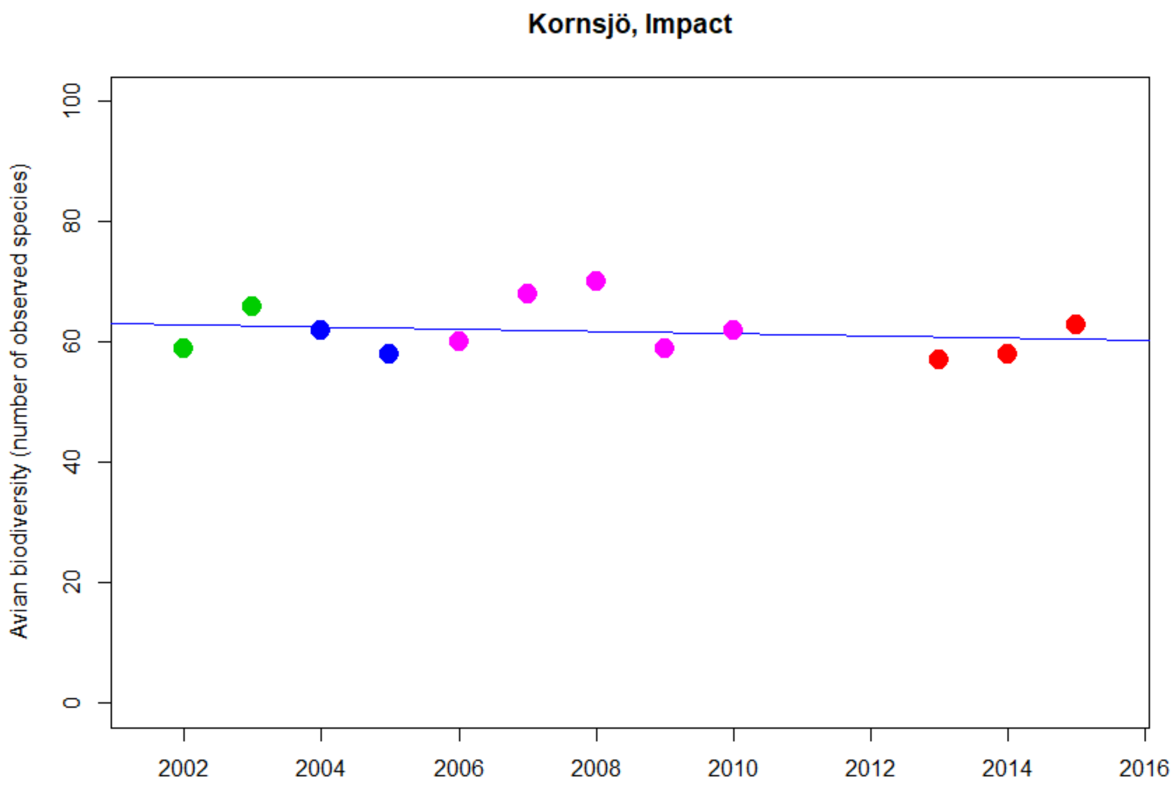


```
Kendall's rank correlation tau = -0.078
z = -0.34531, p-value = 0.730 (estimated due to ties)
```

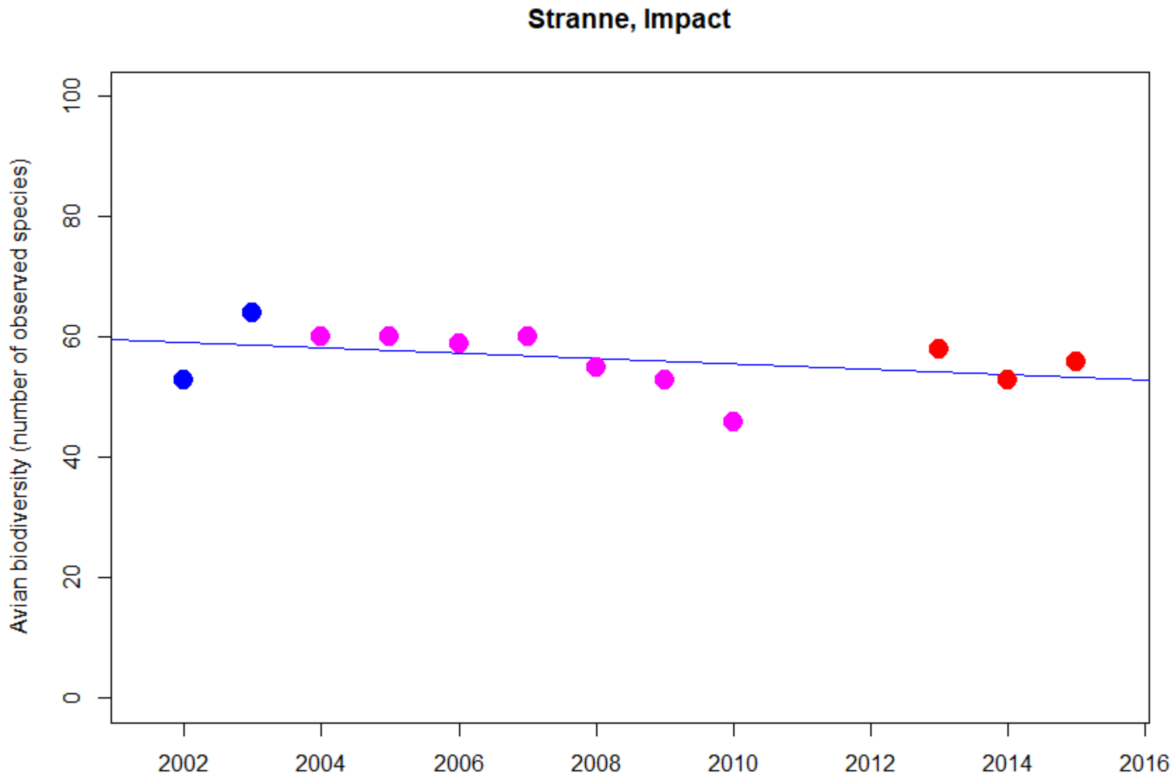


```
Kendall's rank correlation tau = -0.413
z = -1.8144, p-value = 0.070 (estimated due to ties)
```

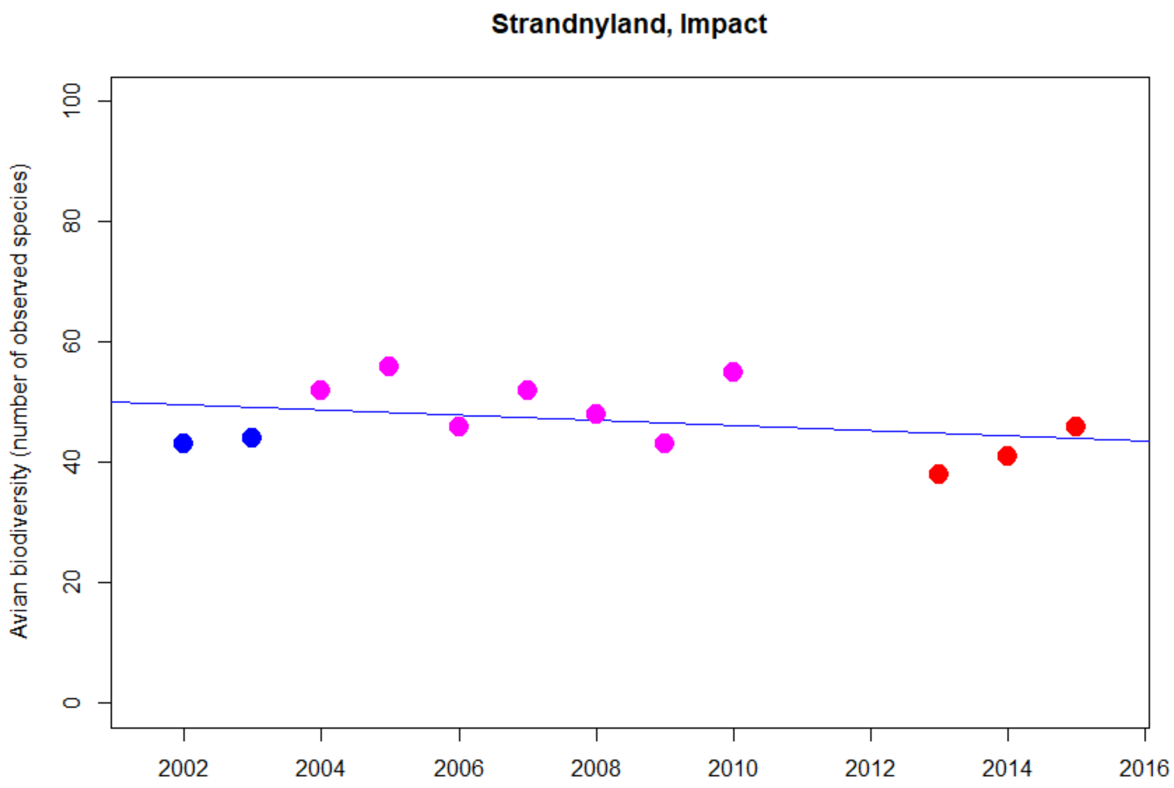


```
Kendall's rank correlation tau = -0.140
z = -0.62155, p-value = 0.534 (estimated due to ties)
```

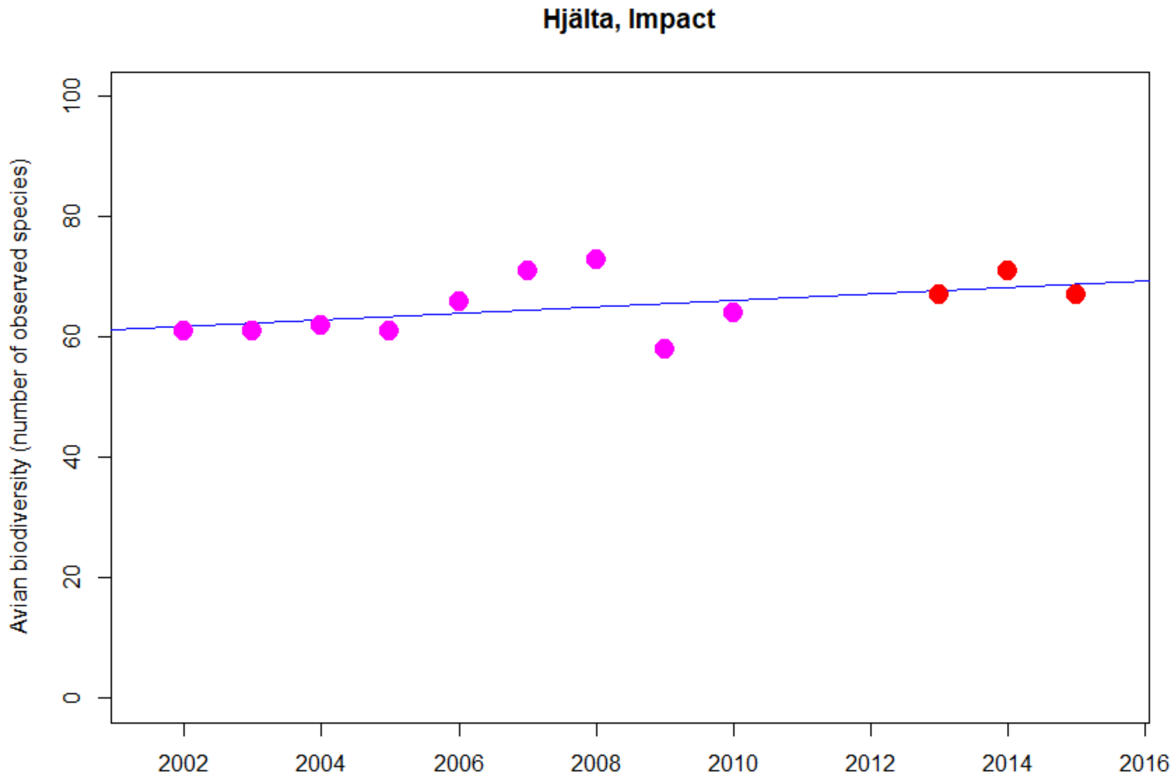


Kendall's rank correlation tau = 0.426
z = 1.8766, p-value = 0.061 (estimated due to ties)

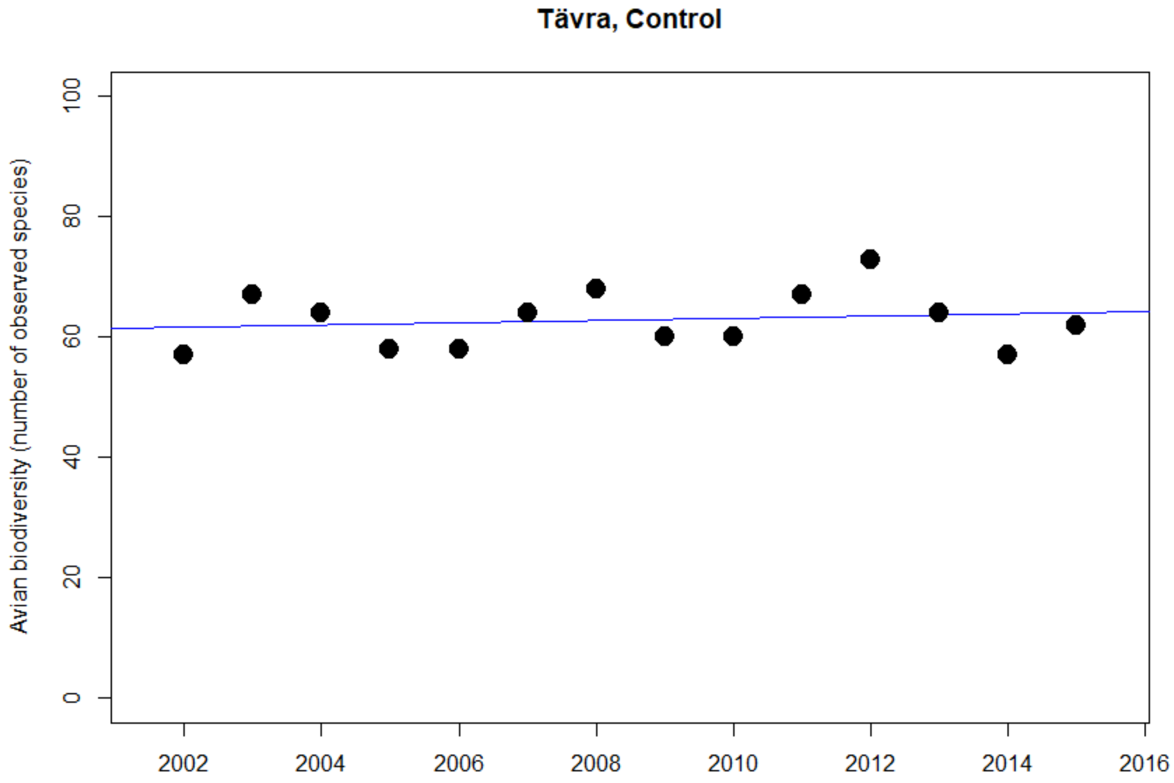


```
Kendall's rank correlation tau = 0.114
z = 0.55385, p-value = 0.580 (estimated due to ties)
```

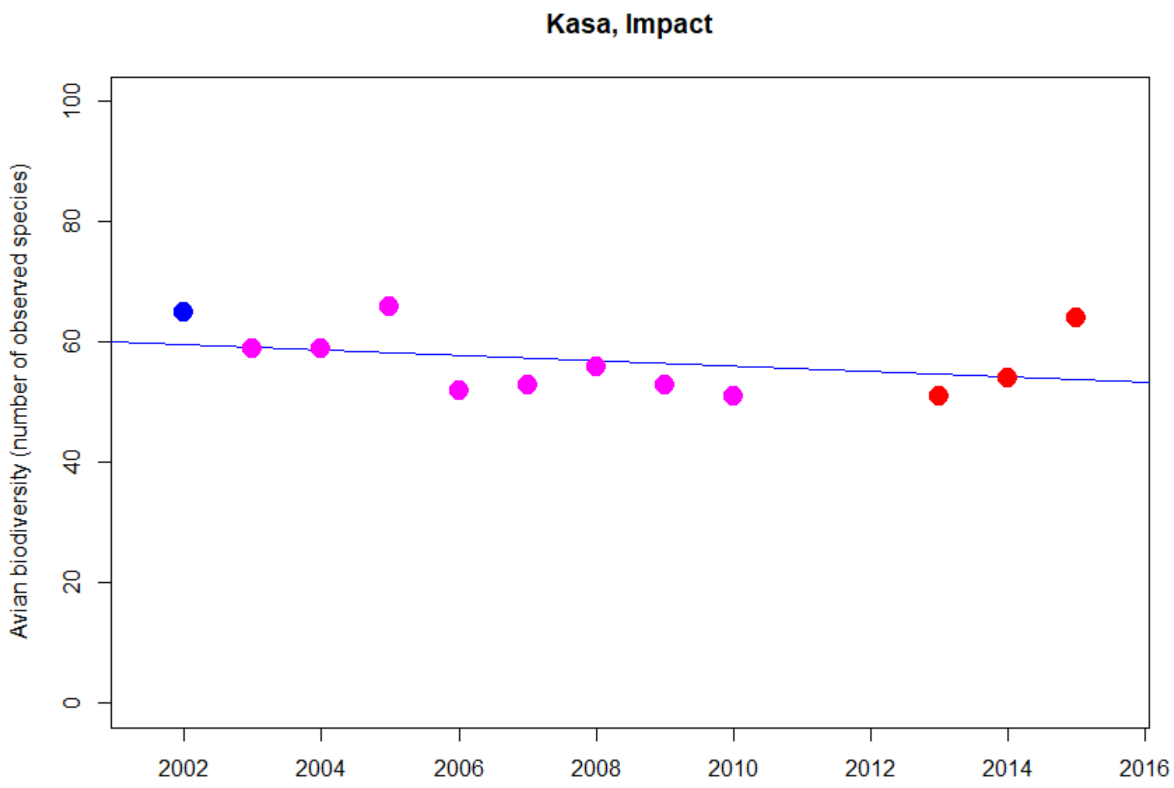


```
Kendall's rank correlation tau = -0.326
z = -1.4503, p-value = 0.147 (estimated due to ties)
```

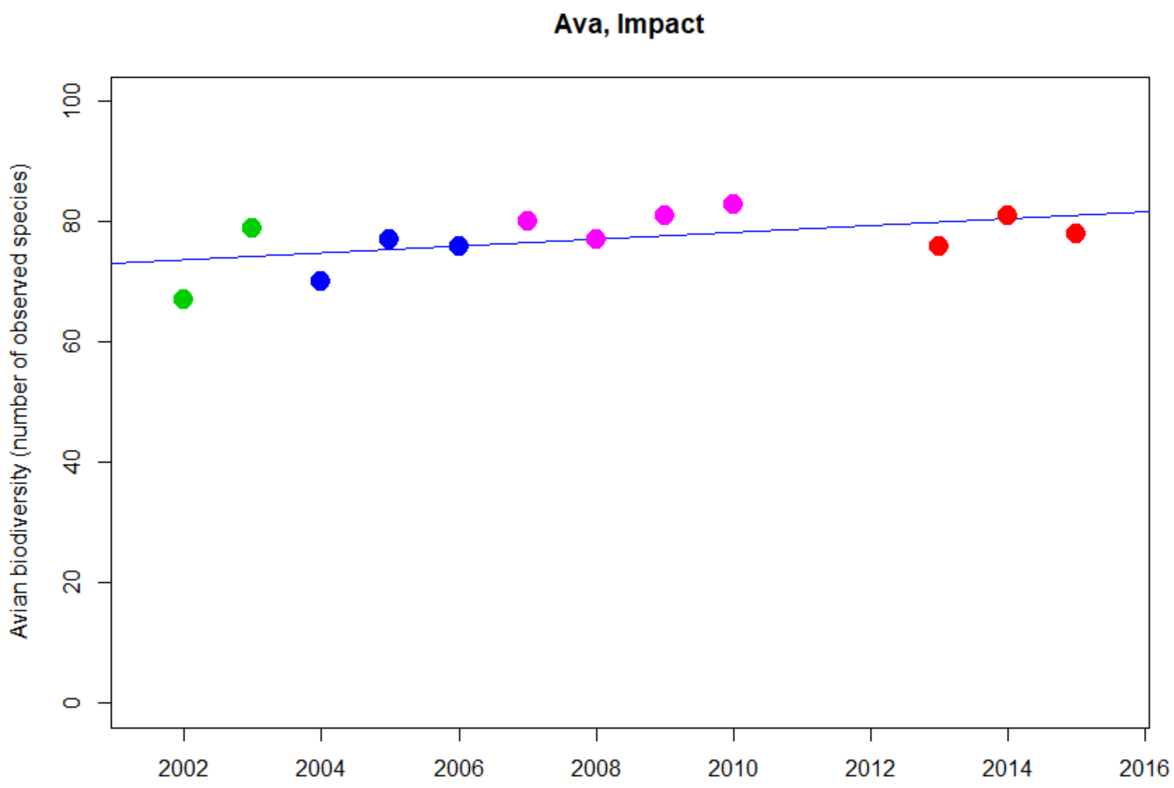


```
Kendall's rank correlation tau = 0.419
z = 1.8647, p-value = 0.062 (estimated due to ties)
```

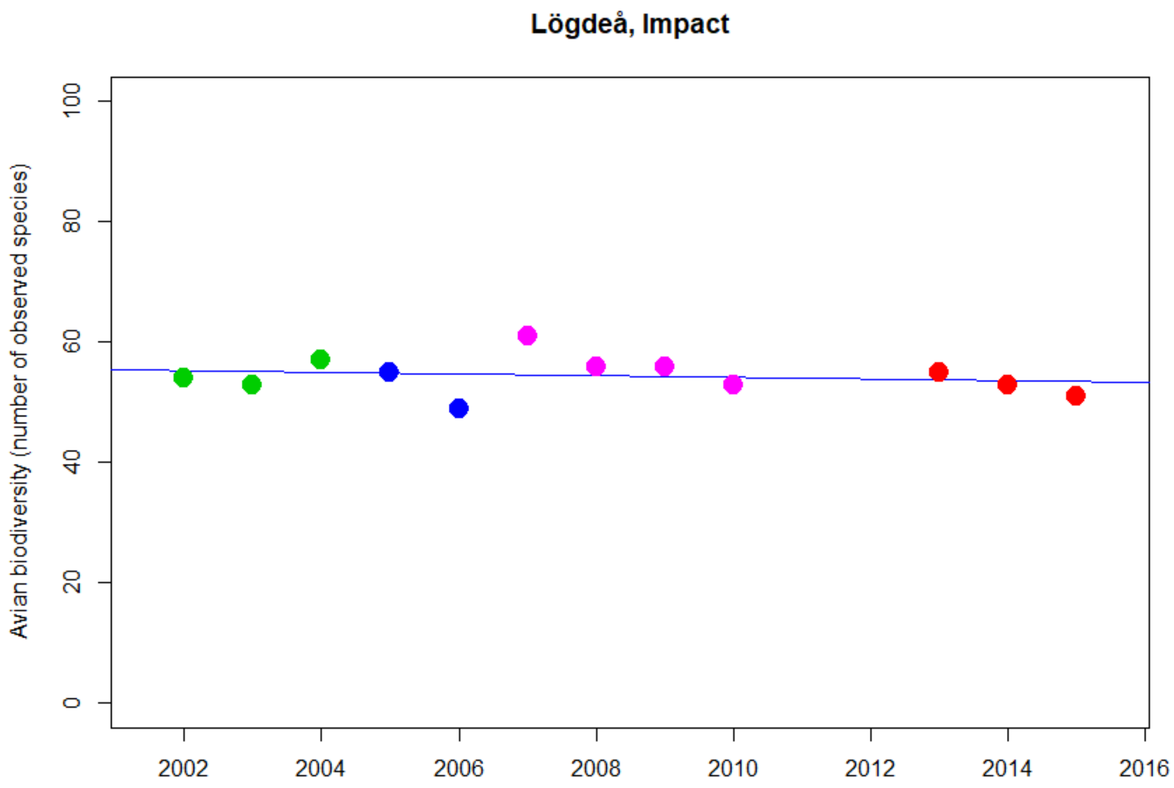


```
Kendall's rank correlation tau = -0.205
z = -0.90356, p-value = 0.366 (estimated due to ties)
```

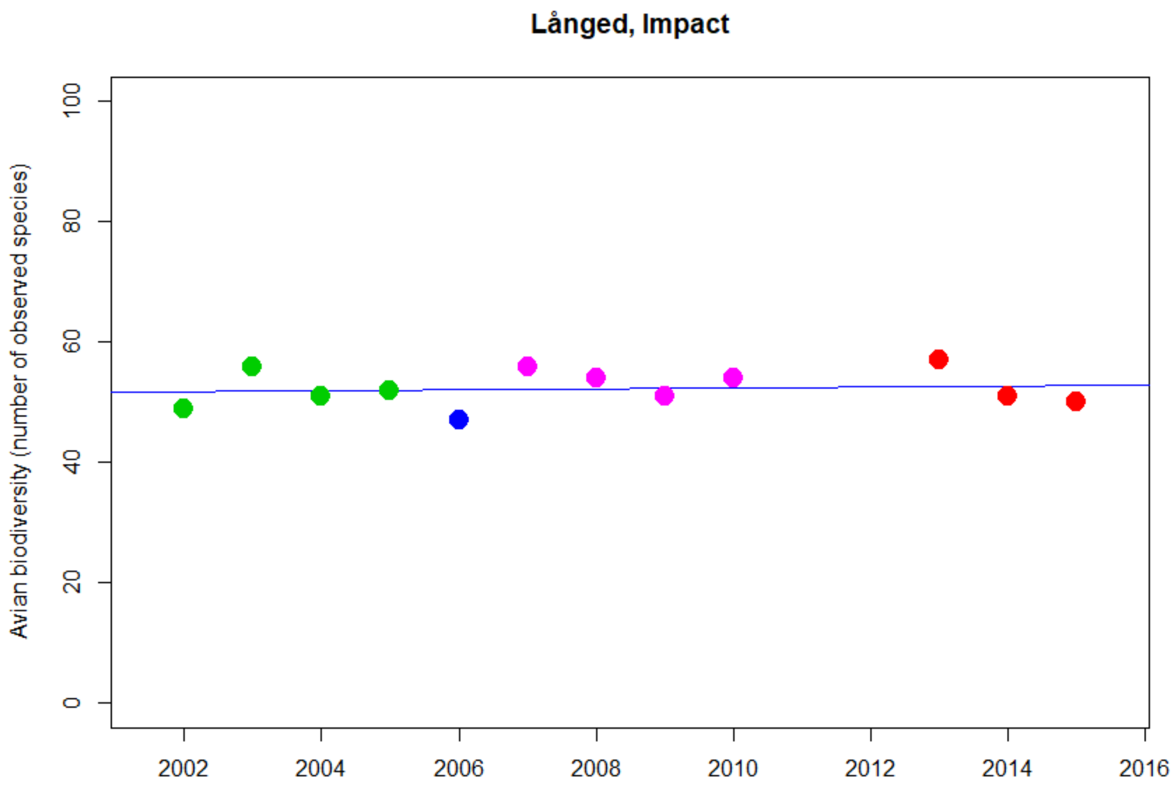


```
Kendall's rank correlation tau = 0.047
z = 0.20851, p-value = 0.835 (estimated due to ties)
```

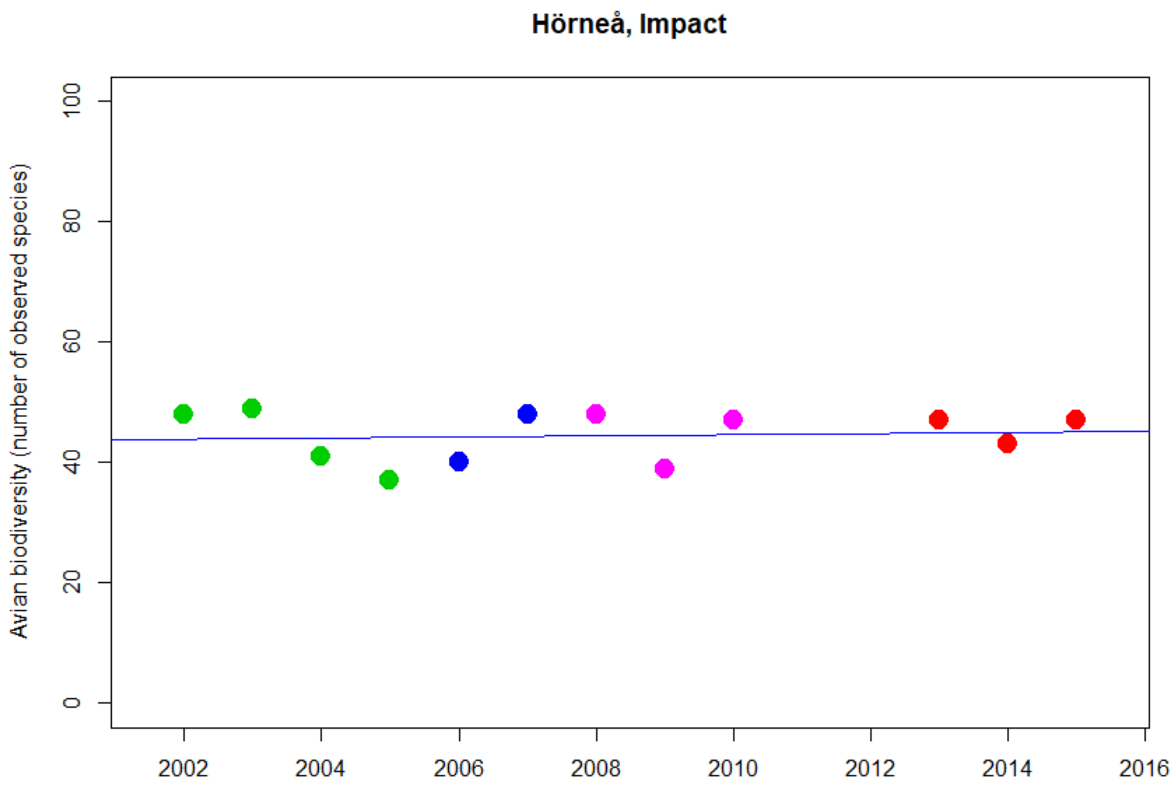


```
Kendall's rank correlation tau = -0.127
z = -0.55829, p-value = 0.577 (estimated due to ties)
```

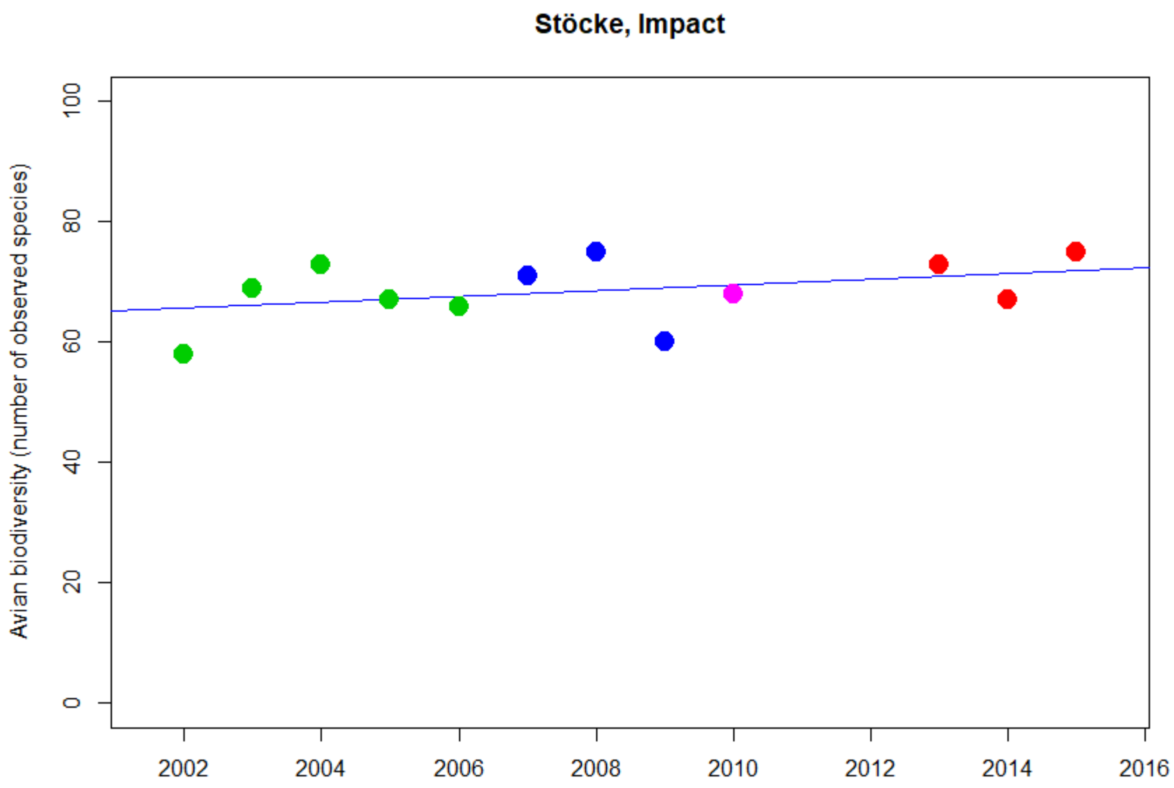


```
Kendall's rank correlation tau = 0.264
z = 1.174, p-value = 0.240 (estimated due to ties)
```

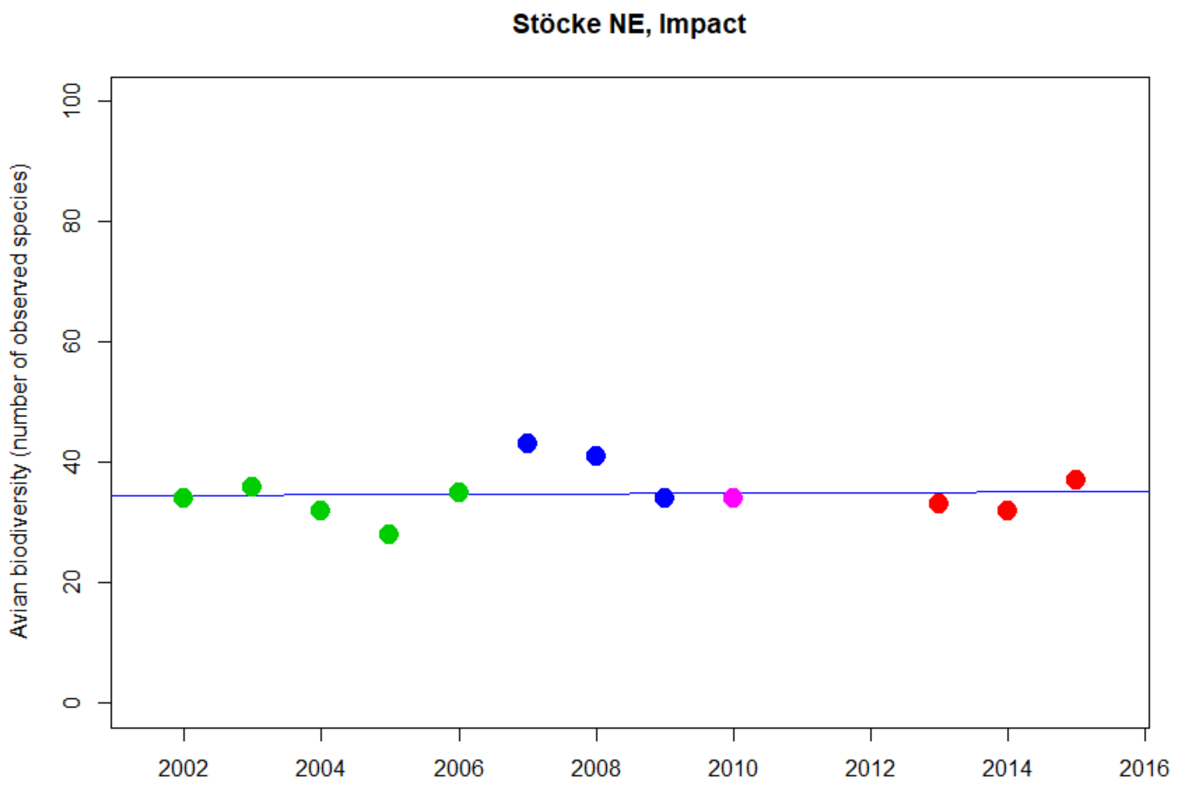


```
Kendall's rank correlation tau = -0.031
z = -0.13868, p-value = 0.890 (estimated due to ties)
```

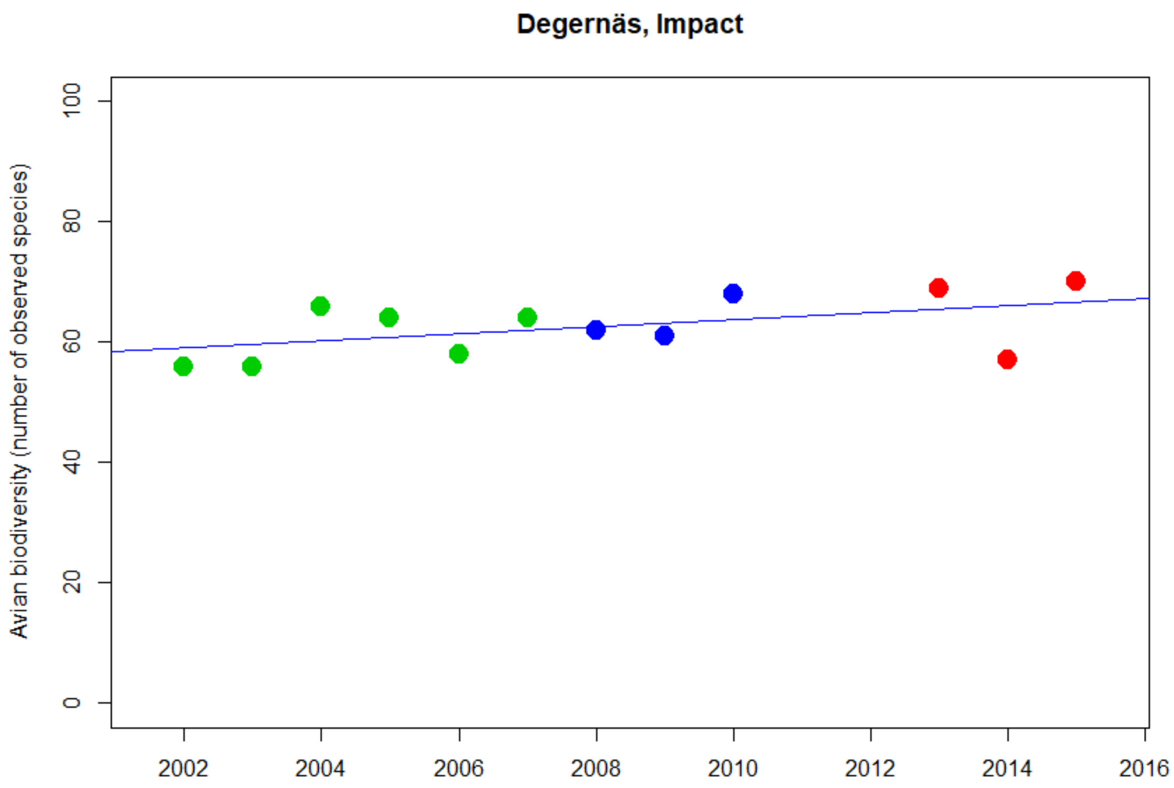


```
Kendall's rank correlation tau = 0.400
z = 1.7913, p-value = 0.073 (estimated due to ties)
```

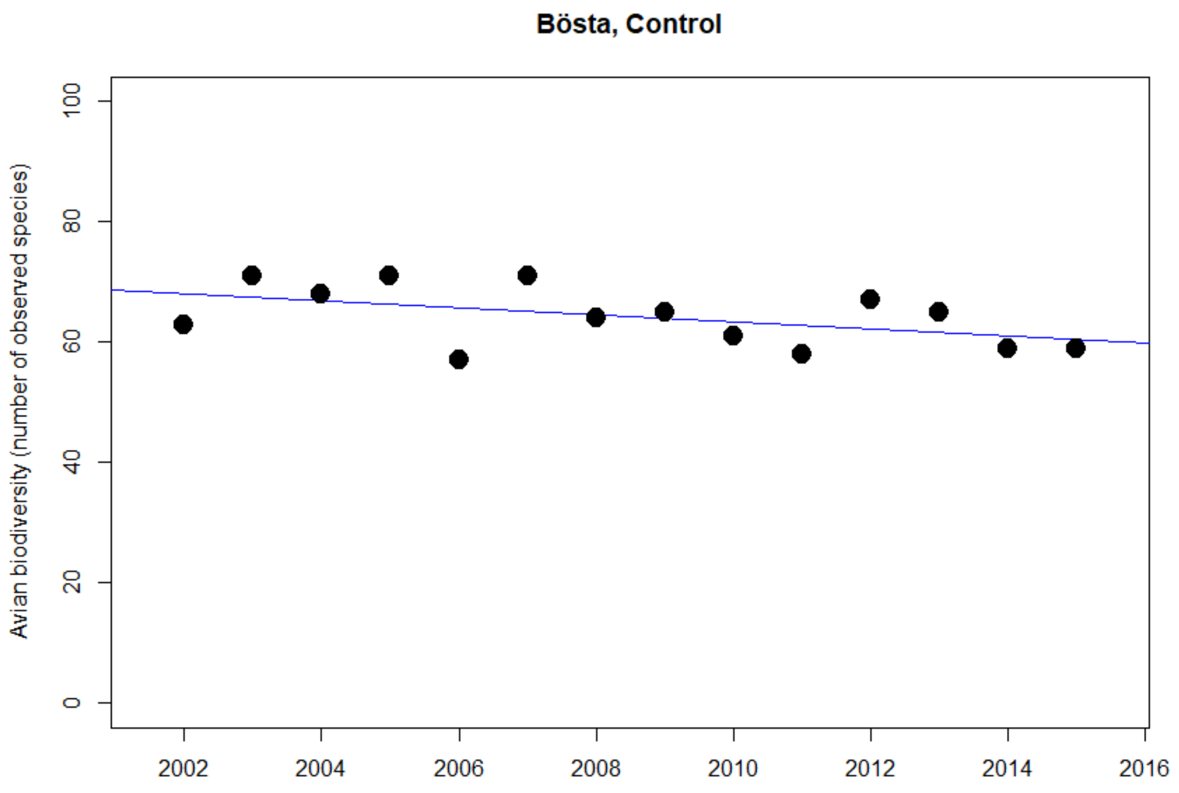


```
Kendall's rank correlation tau = -0.317
z = -1.546, p-value = 0.122 (estimated due to ties)
```
