## Supplementary material for "Are railways detrimental to bird populations? A BDACI study on the construction of the Bothnia Line Railway": Trends in TM distances Impact vs Control

Supplementary material **Dist1**

### Temporal trends in annual mean distances between territory midpoints and an arbitrary baseline (Control sites, blue) or the BLR (Impact sites, red).

Plots and Kendall’s rank correlation tests.

Some correlation tests generated warning messages due to ties (data points with the same Y-value). These warning texts were omitted for improved readability.

*“NA” marks cases where the number of data points was too small for meaningful statistical analyses.*

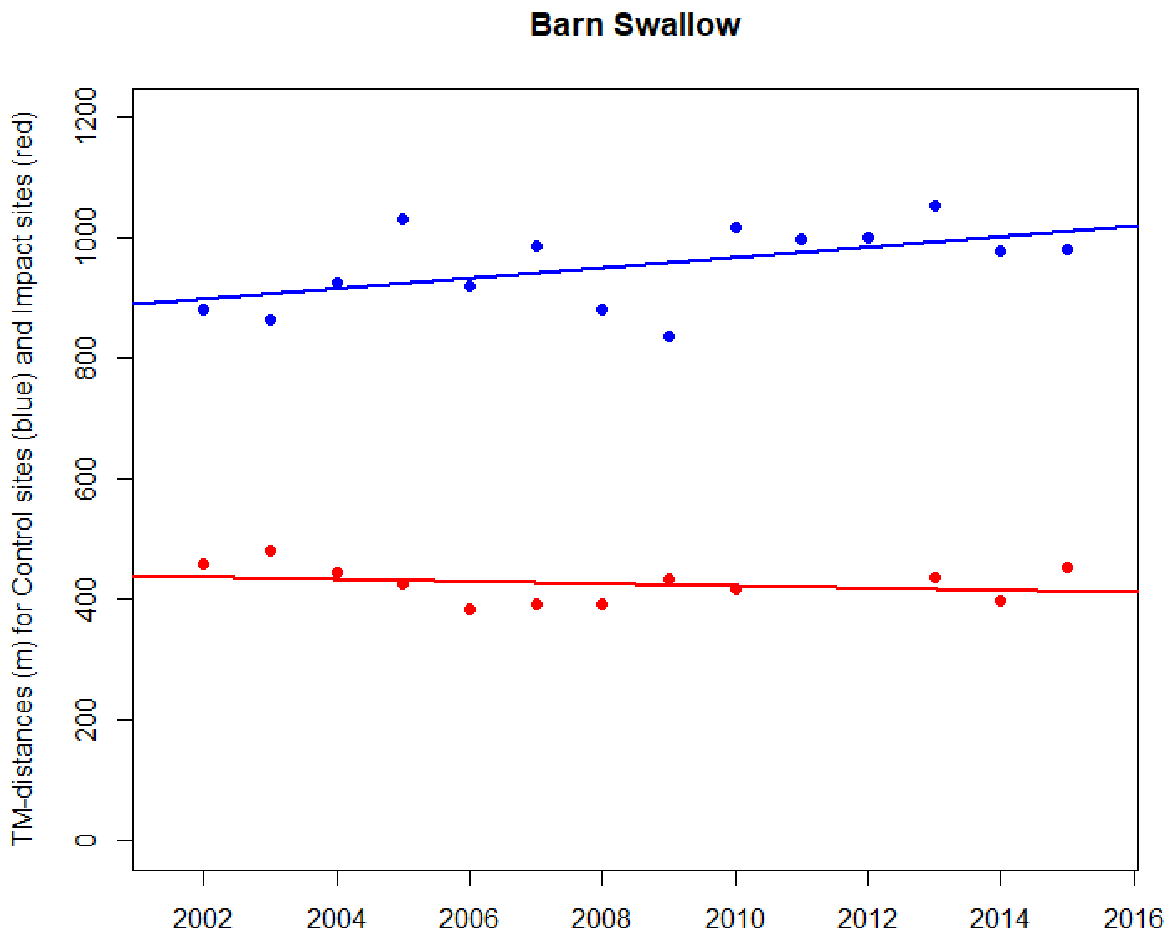


**Impact sites, TMtR distances**

Kendall's rank correlation tau = -0.121

T = 29, p-value = 0.638

**Control sites, TMtB distances**

Kendall's rank correlation tau = 0.275

T = 58, p-value = 0.193

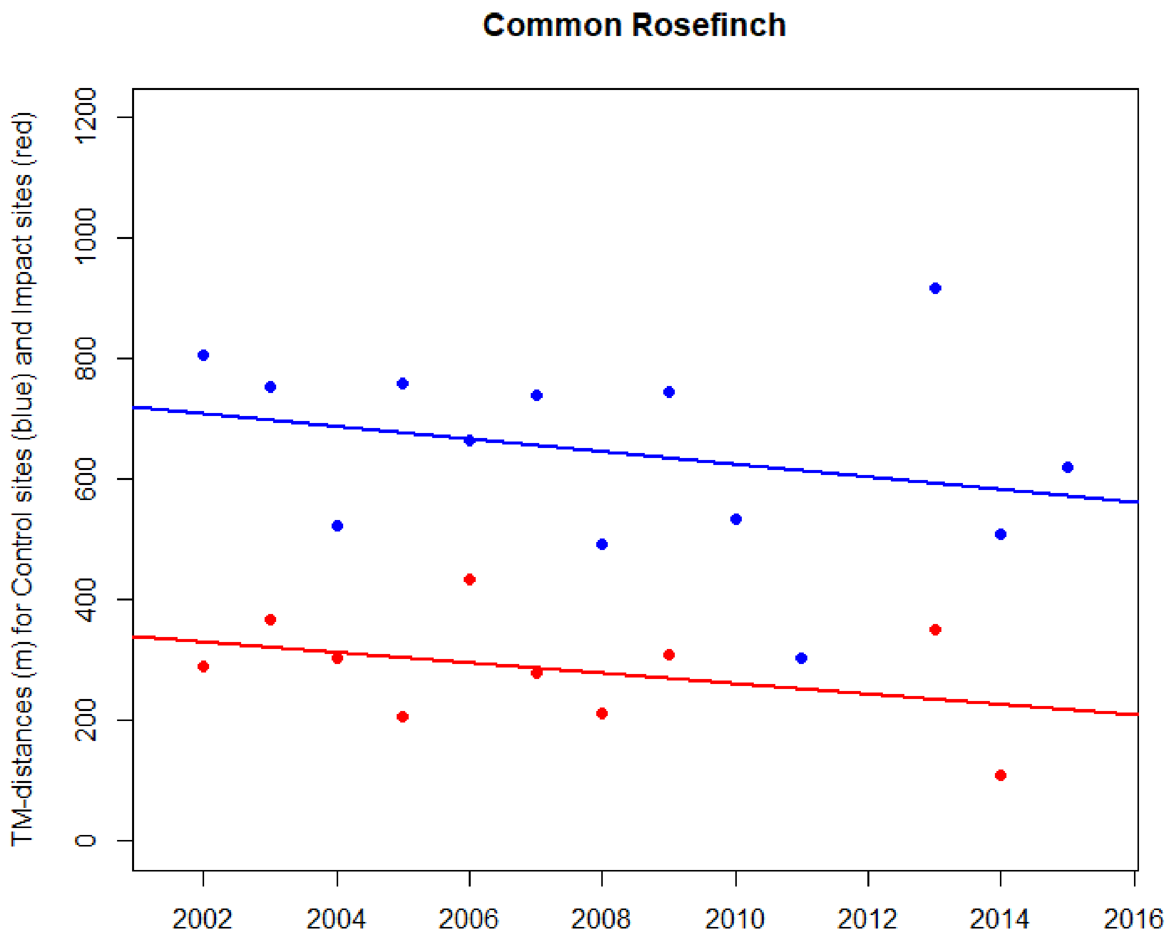


**Impact sites, TMtR distances**

Kendall's rank correlation tau = -0.156

T = 19, p-value = 0.601

**Control sites, TMtB distances**

Kendall's rank correlation tau = -0.282

T = 28, p-value = 0.204

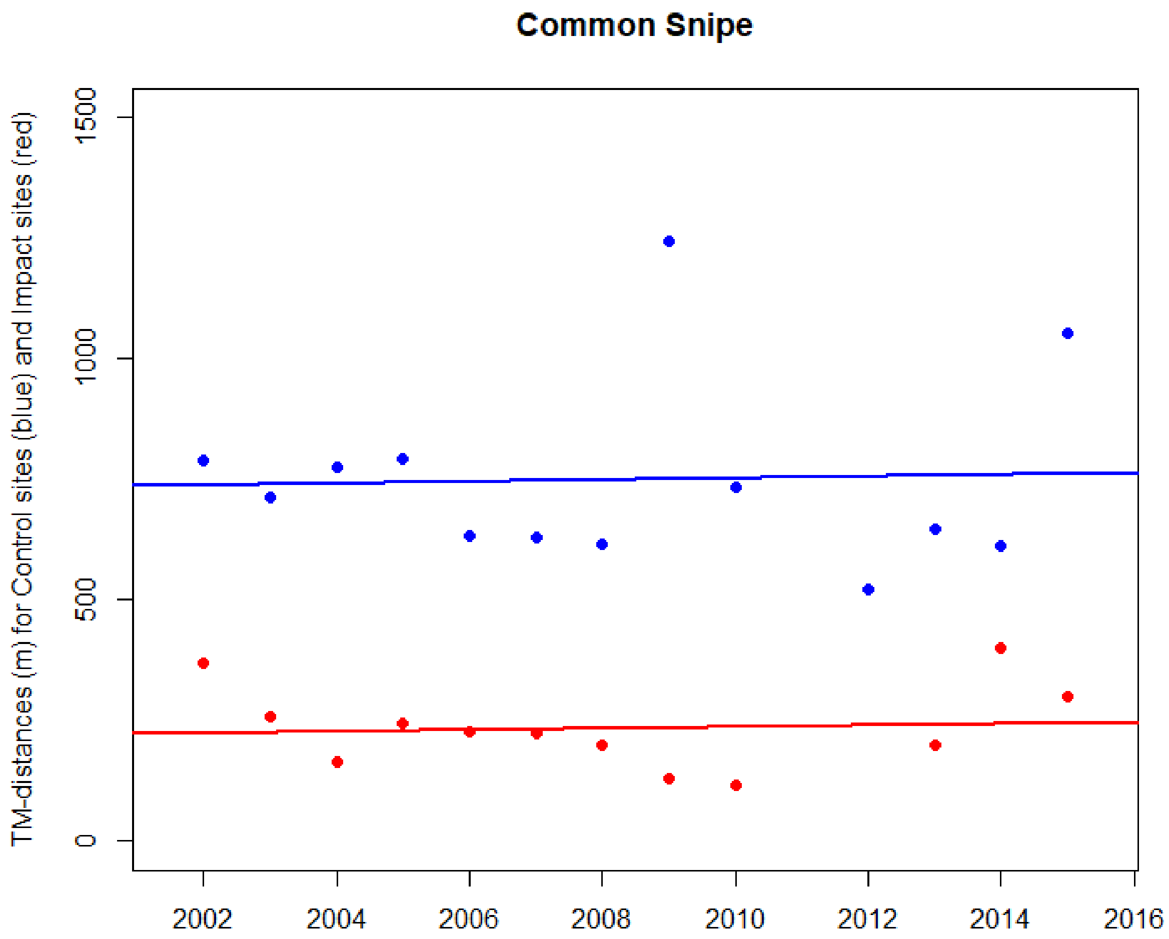


**Impact sites, TMtR distances**

Kendall's rank correlation tau = -0.182

T = 27, p-value = 0.459

**Control sites, TMtB distances**

Kendall's rank correlation tau = -0.205

T = 31, p-value = 0.367

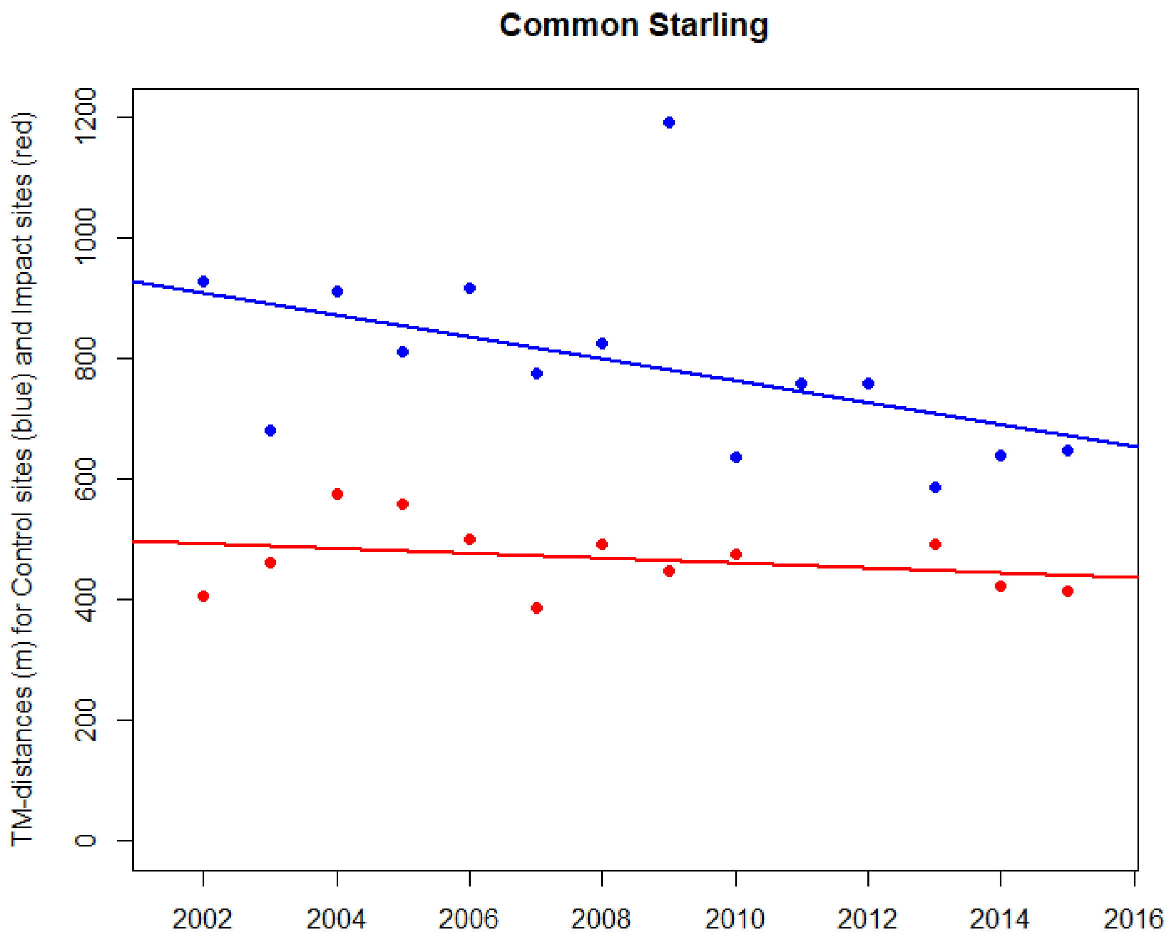


**Impact sites, TMtR distances**

Kendall's rank correlation tau = -0.242

T = 25, p-value = 0.311

**Control sites, TMtB distances**

Kendall's rank correlation tau = -0.429

T = 26, p-value = 0.036*

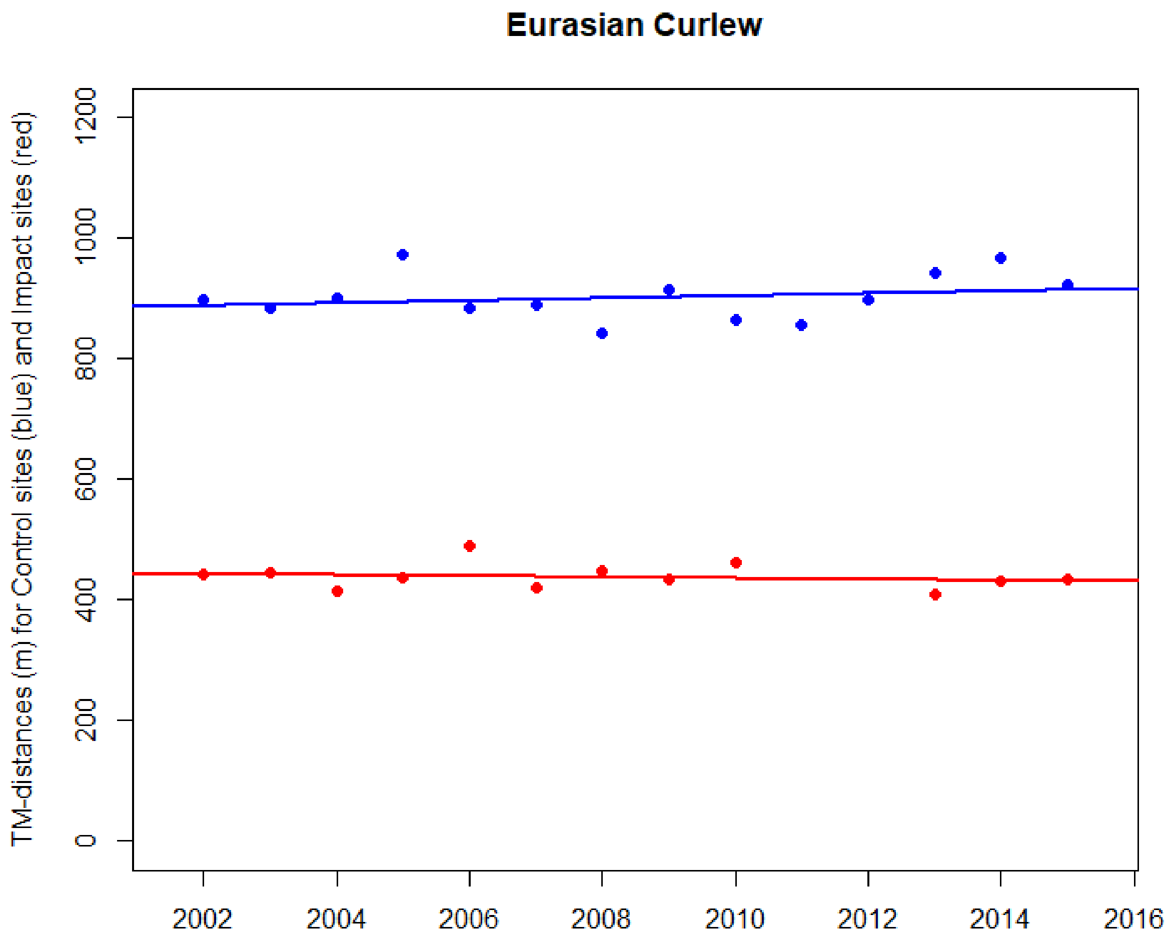


**Impact sites TMtR distances**

Kendall's rank correlation tau = -0.121

T = 29, p-value = 0.6384

**Control sites TMtB distances**

Kendall's rank correlation tau = 0.187

T = 54, p-value = 0.388

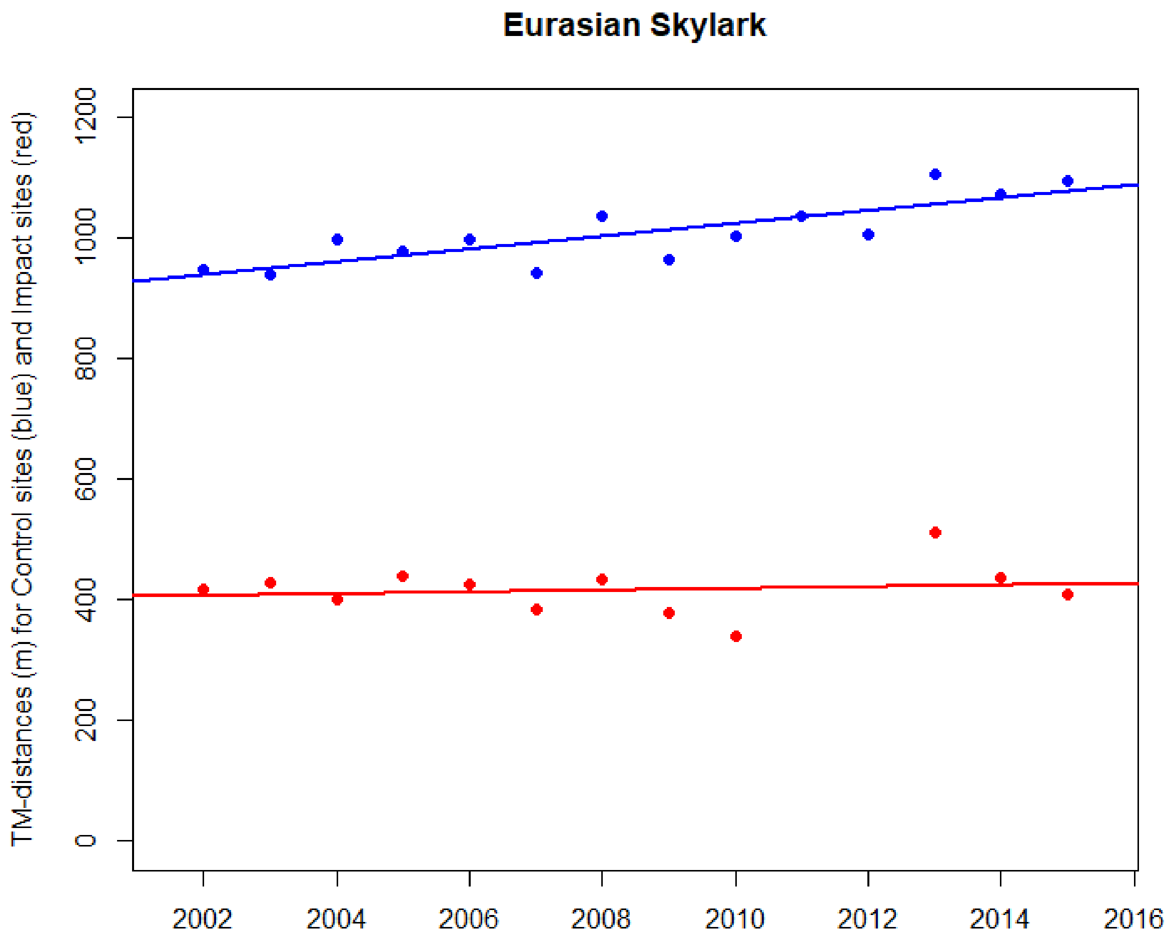


**Impact sites, TMtR distances**

Kendall's rank correlation tau = -0.030

T = 32, p-value = 0.947

**Control sites, TMtB distances**

Kendall's rank correlation tau = 0.670

T = 76, p-value < 0.001***

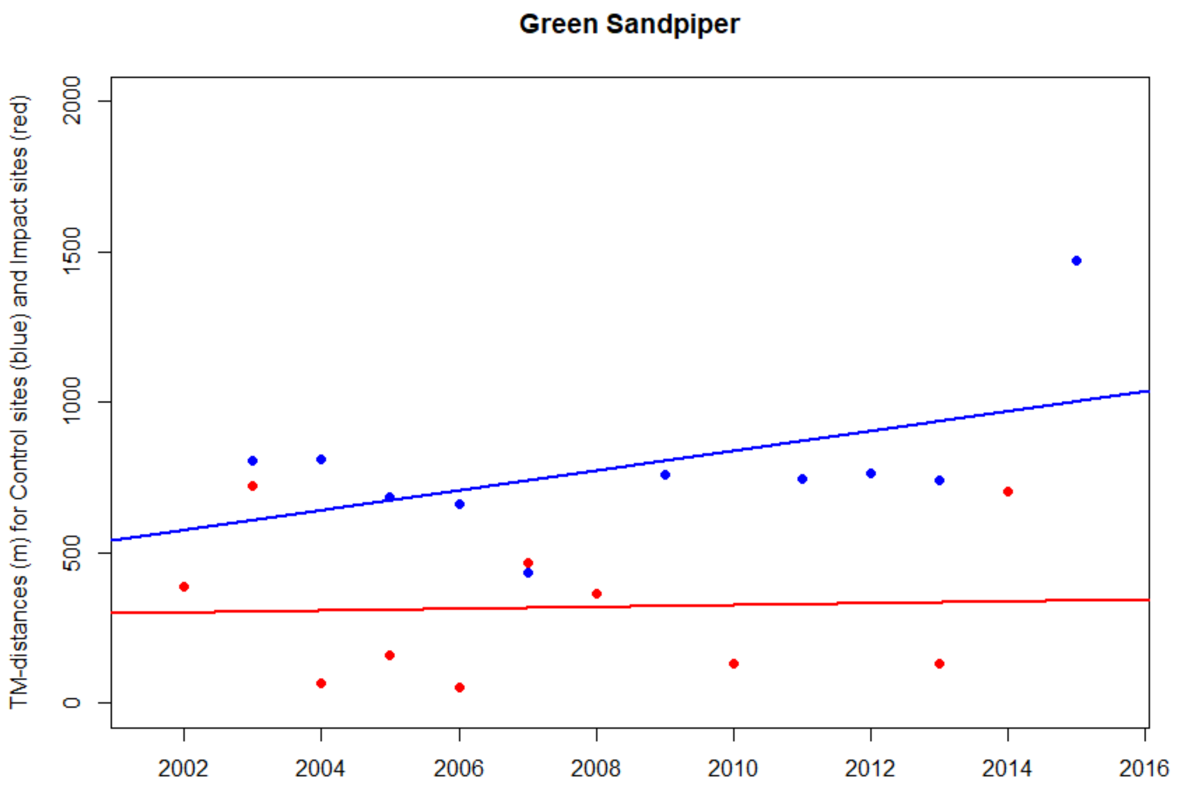


**Impact sites, TMtR distances**

Kendall's rank correlation tau = -0.022

T = 22, p-value = 1

**Control sites, TMtB distances**

Kendall's rank correlation tau = 0.067

T = 24, p-value = 0.862

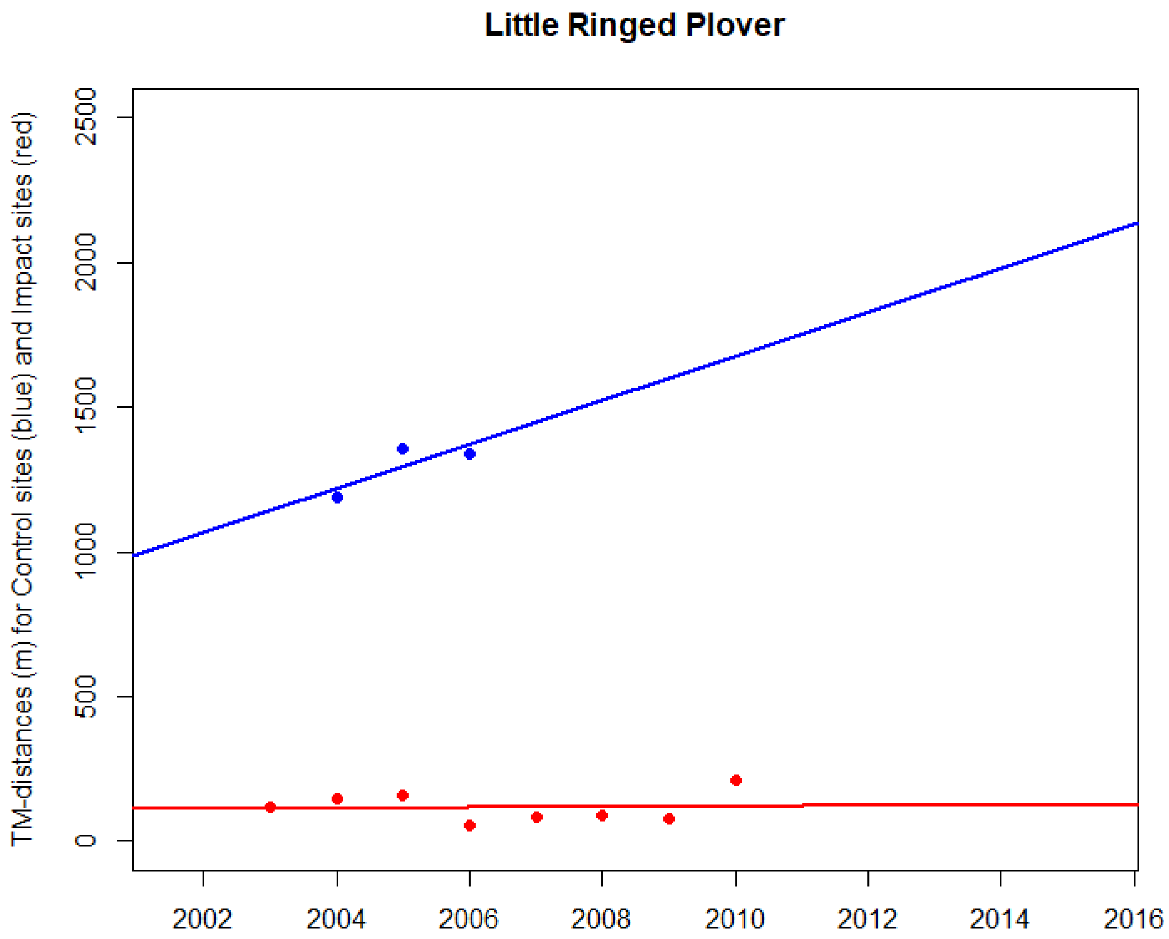


**Impact sites, TMtR distances**

Kendall's rank correlation tau = 0.0

T = 14, p-value = 1

**Control sites, TMtB distances**

Kendall's rank correlation tau = NA

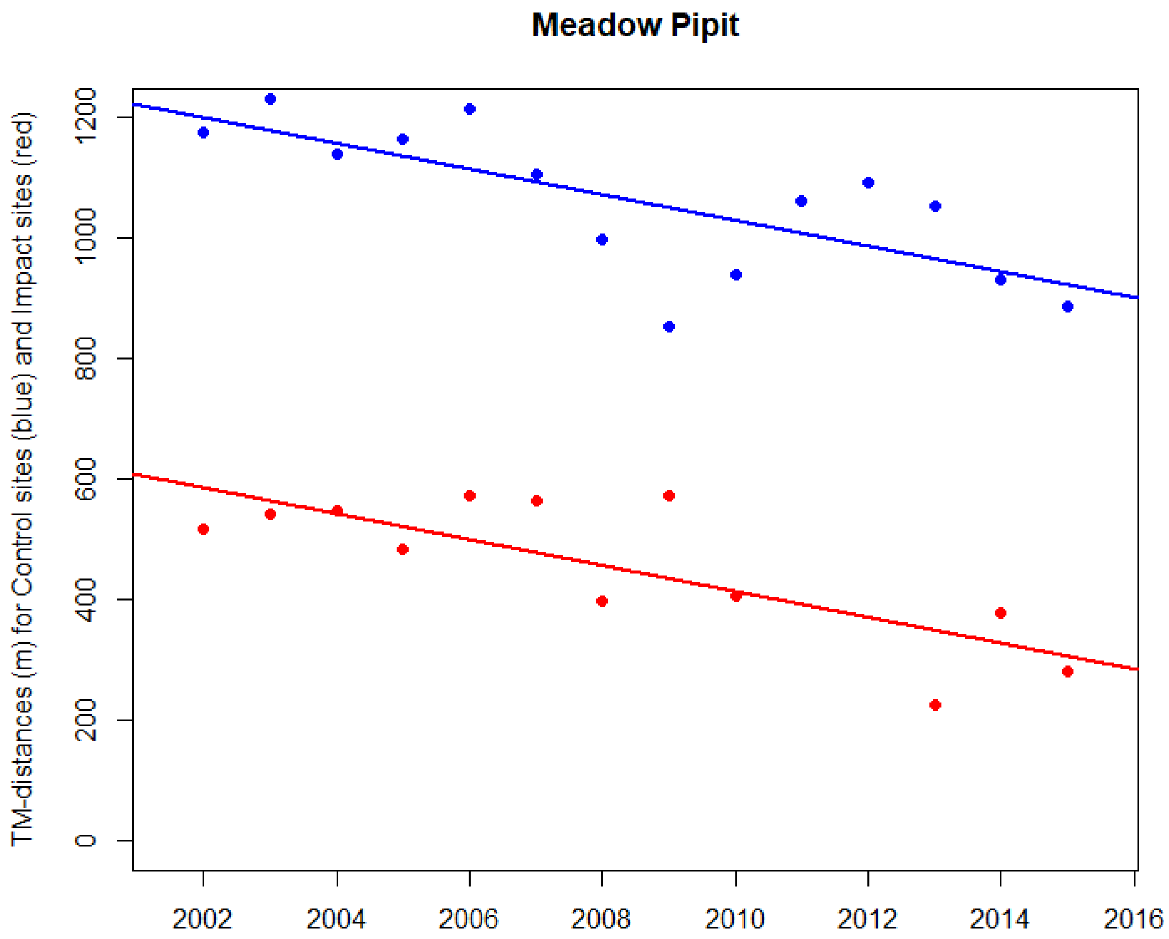


**Impact sites, TMtR distances**

Kendall's rank correlation tau = -0.364

T = 21, p-value = 0.116

**Control sites, TMtB distances**

Kendall's rank correlation tau = -0.604

T = 18, p-value = 0.002**

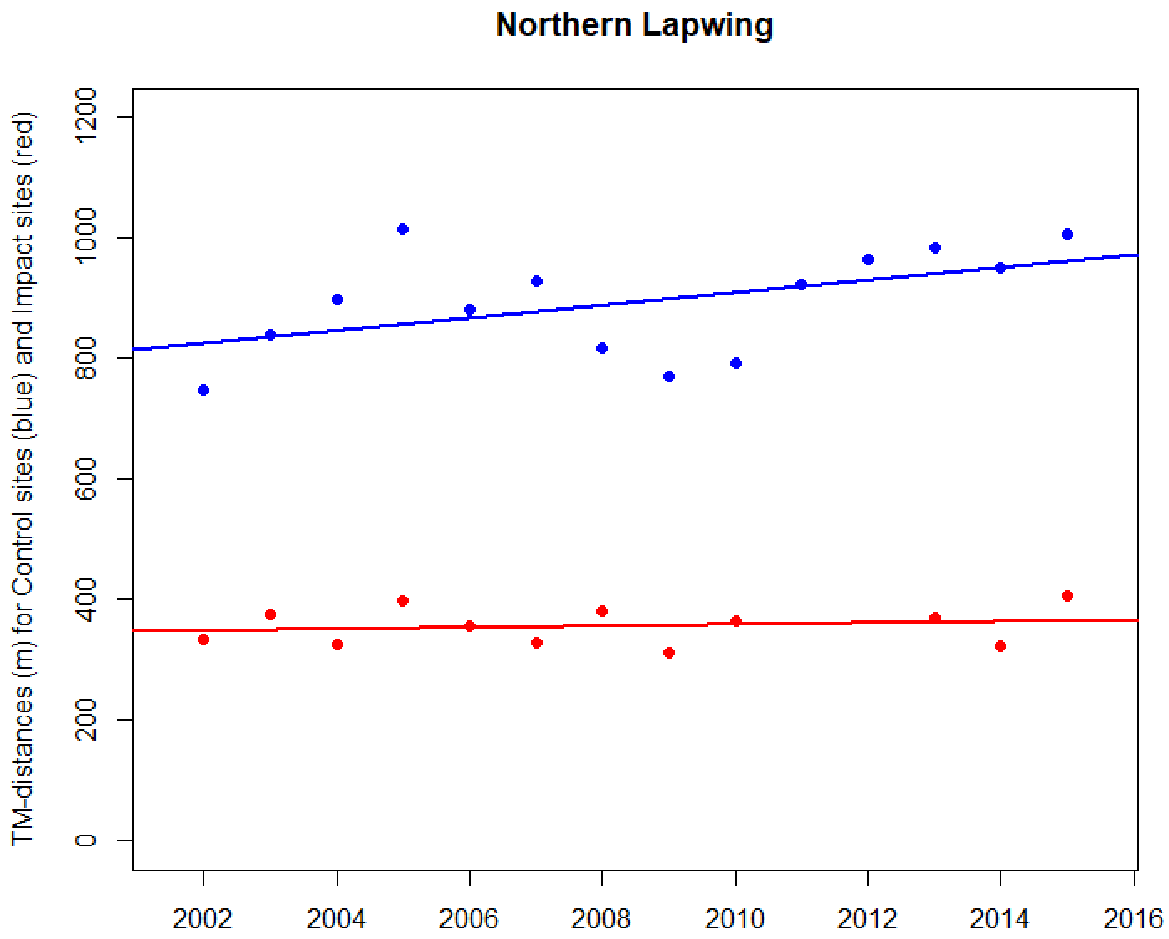


**Impact sites, TMtR distances**

Kendall's rank correlation tau = 0.061

T = 35, p-value = 0.841

**Control sites, TMtB distances**

Kendall's rank correlation tau = 0.385

T = 63, p-value = 0.062

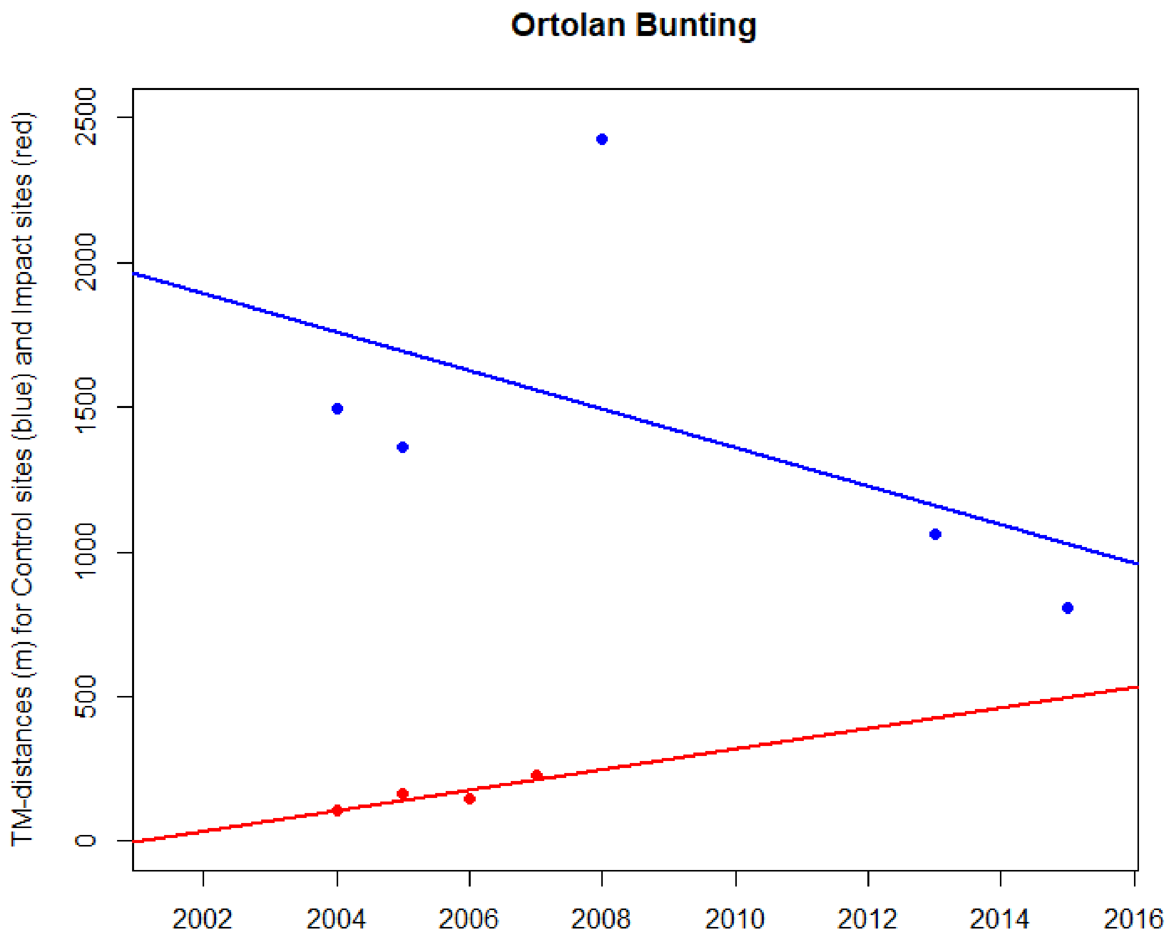


**Impact sites, TMtR distances**

Kendall's rank correlation tau = 0.667

T = 5, p-value = 0.333

**Control sites, TMtB distances**

Kendall's rank correlation tau = -0.6

T = 2, p-value = 0.233

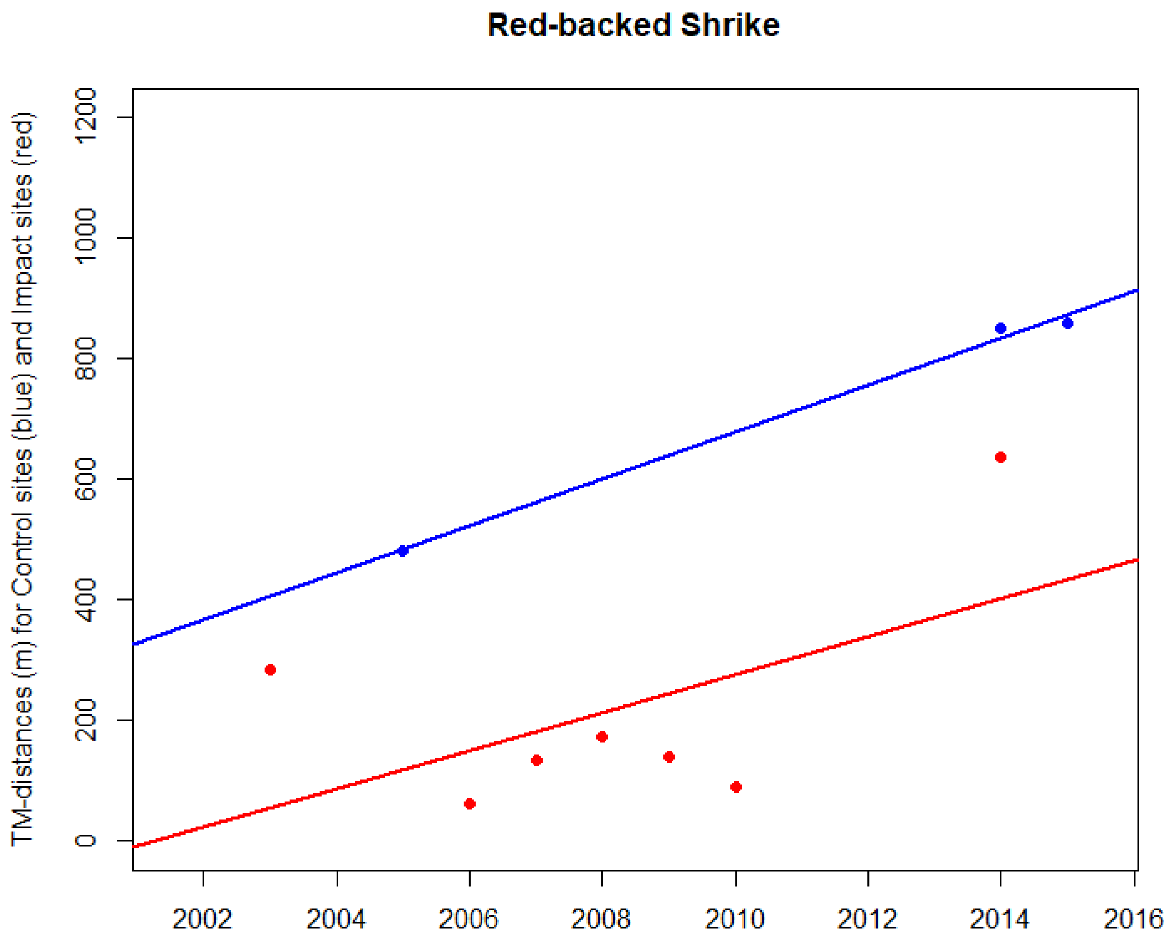


**Impact sites, TMtR distances**

Kendall's rank correlation tau = 0.143

T = 12, p-value = 0.773

**Control sites, TMtB distances**

Kendall's rank correlation tau = NA

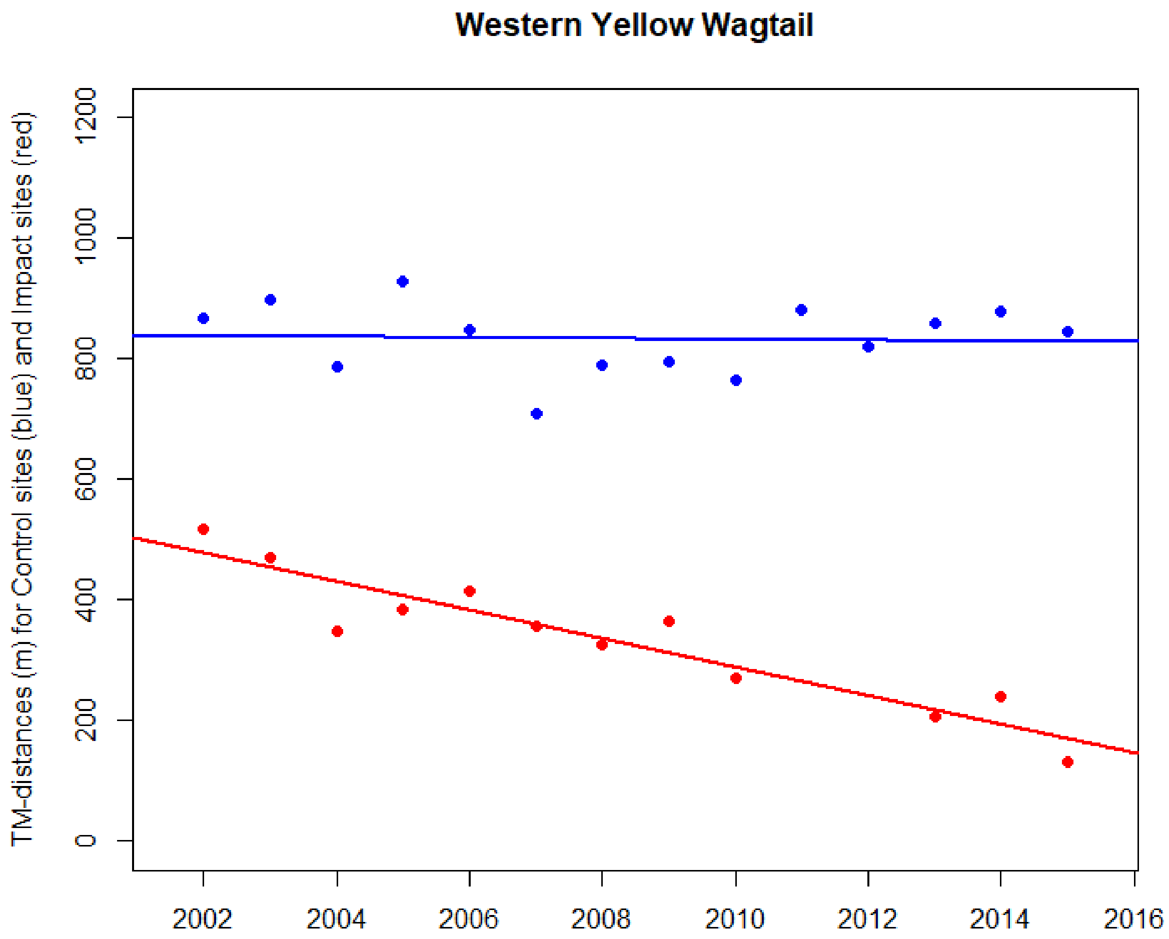


**Impact sites, TMtR distances**

Kendall's rank correlation tau = -0.758

T = 8, p-value < 0.001***

**Control sites, TMtB distances**

Kendall's rank correlation tau = -0.011

T = 45, p-value = 1

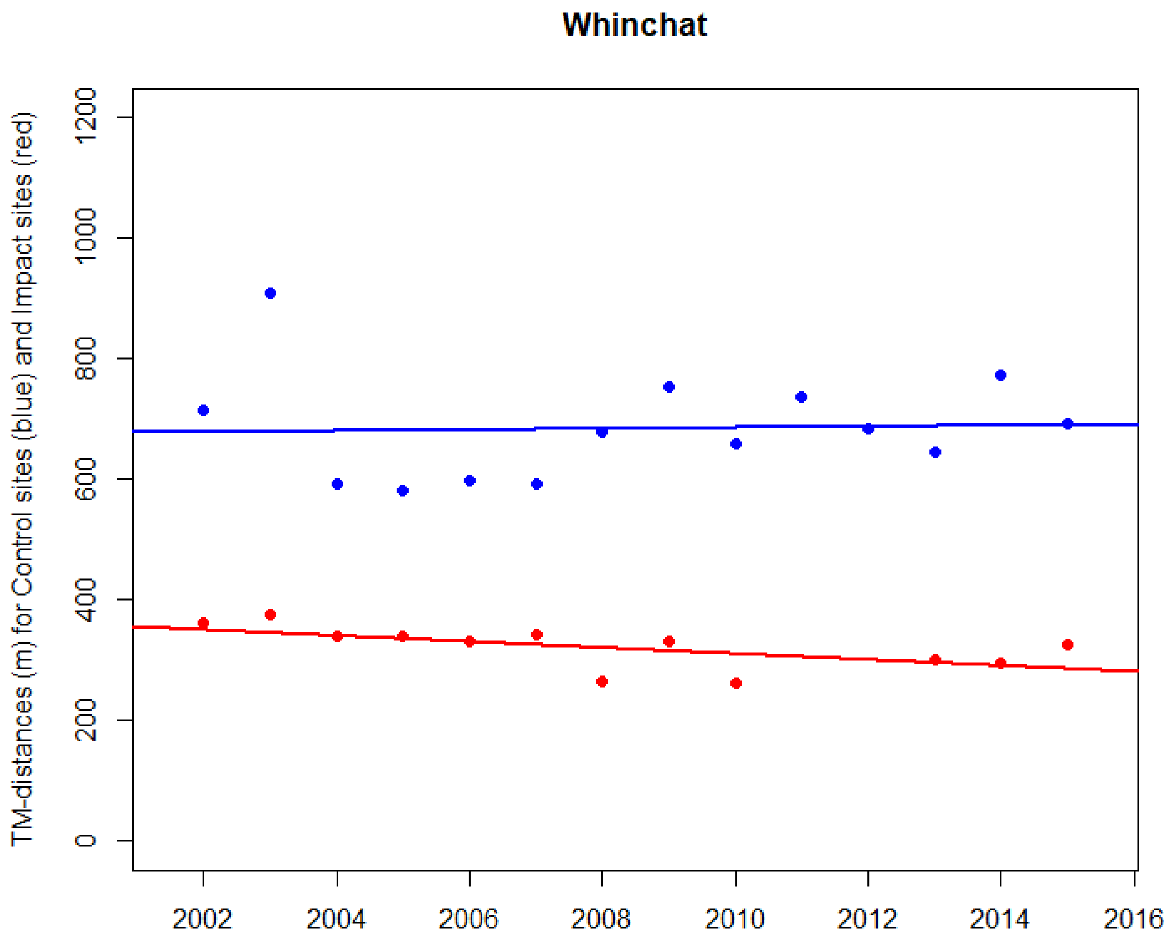


**Impact sites, TMtR distances**

Kendall's rank correlation tau = -0.545

T = 15, p-value = 0.014*

**Control sites, TMtB distances**

Kendall's rank correlation tau = 0.209

T = 55, p-value = 0.331
