## Supplementary material for "Are railways detrimental to bird populations? A BDACI study on the construction of the Bothnia Line Railway": Trends in proportions of distances to BLR

Supplementary material **Dist2**

### Temporal patterns of numbers and proportions (%) of territory midpoints within 100 m zones from the Bothnia Line Railway. All study species combined and individual species.

The 100 m zones (N = max. 17) are depicted from bottom up.

Plots produced in Microsoft Excel.

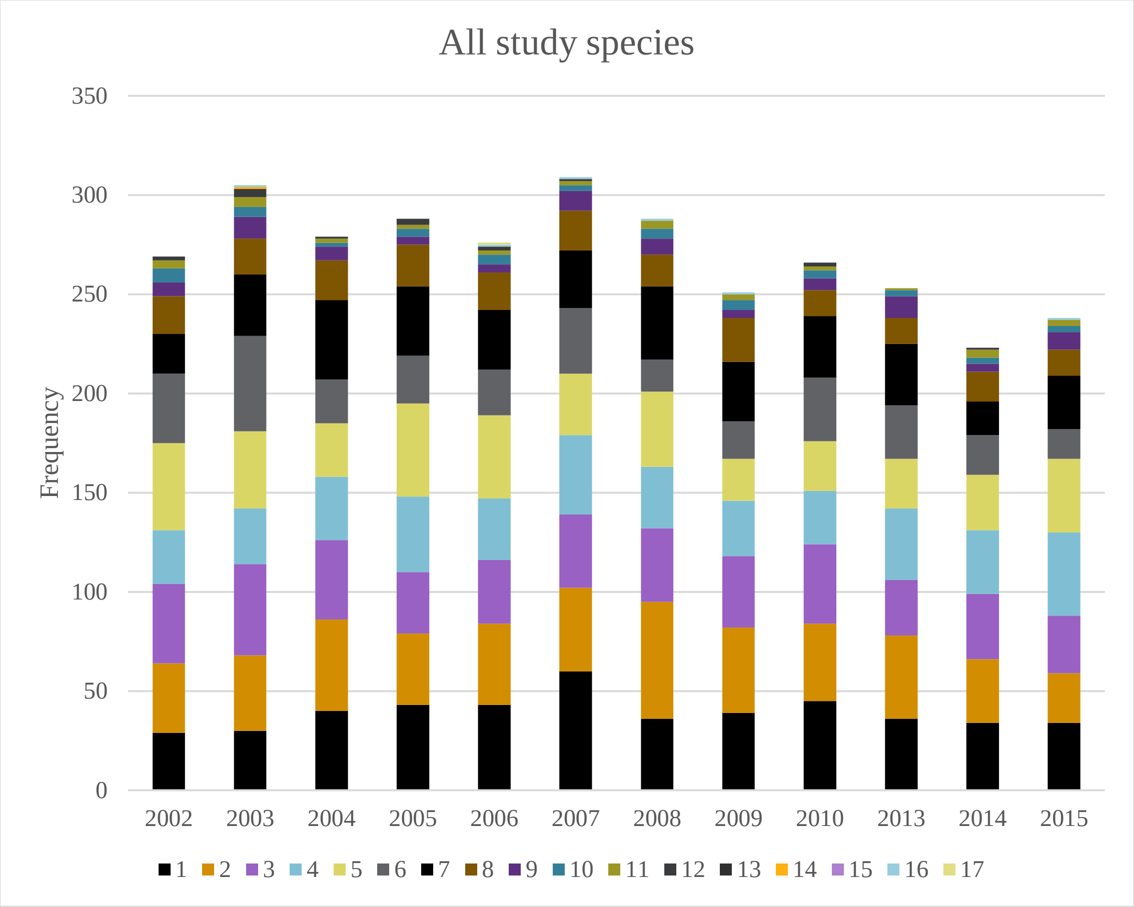
All study species
Frequency
350
300
250
200
150
100
50
0
2002
2003
2004
2005
2006
2007
2008
2009
2010
2013
2014
2015
1 2 3 4 5 6 7 8 9 10 11 12 13 14 15 16 17

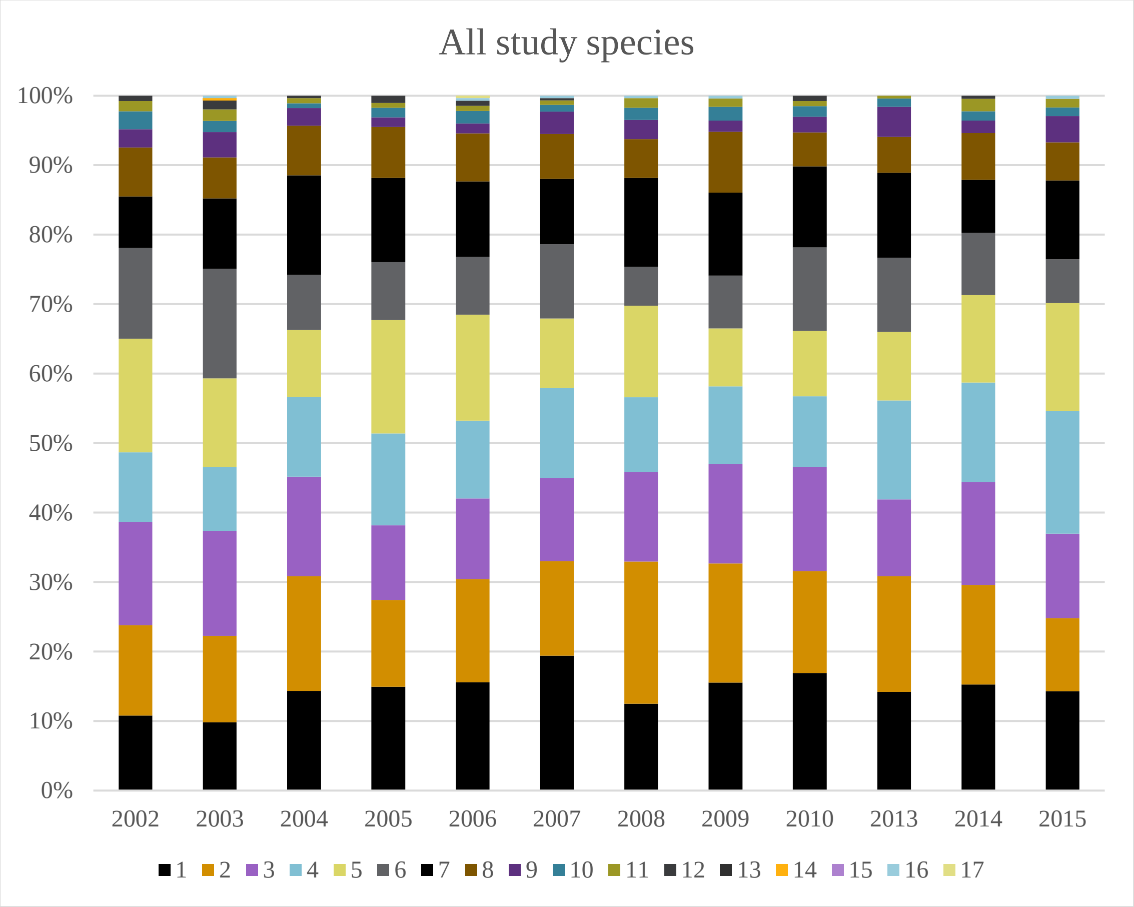
All study species
100%
90%
80%
70%
60%
50%
40%
30%
20%
10%
0%
2002
2003
2004
2005
2006
2007
2008
2009
2010
2013
2014
2015
1 2 3 4 5 6 7 8 9 10 11 12 13 14 15 16 17

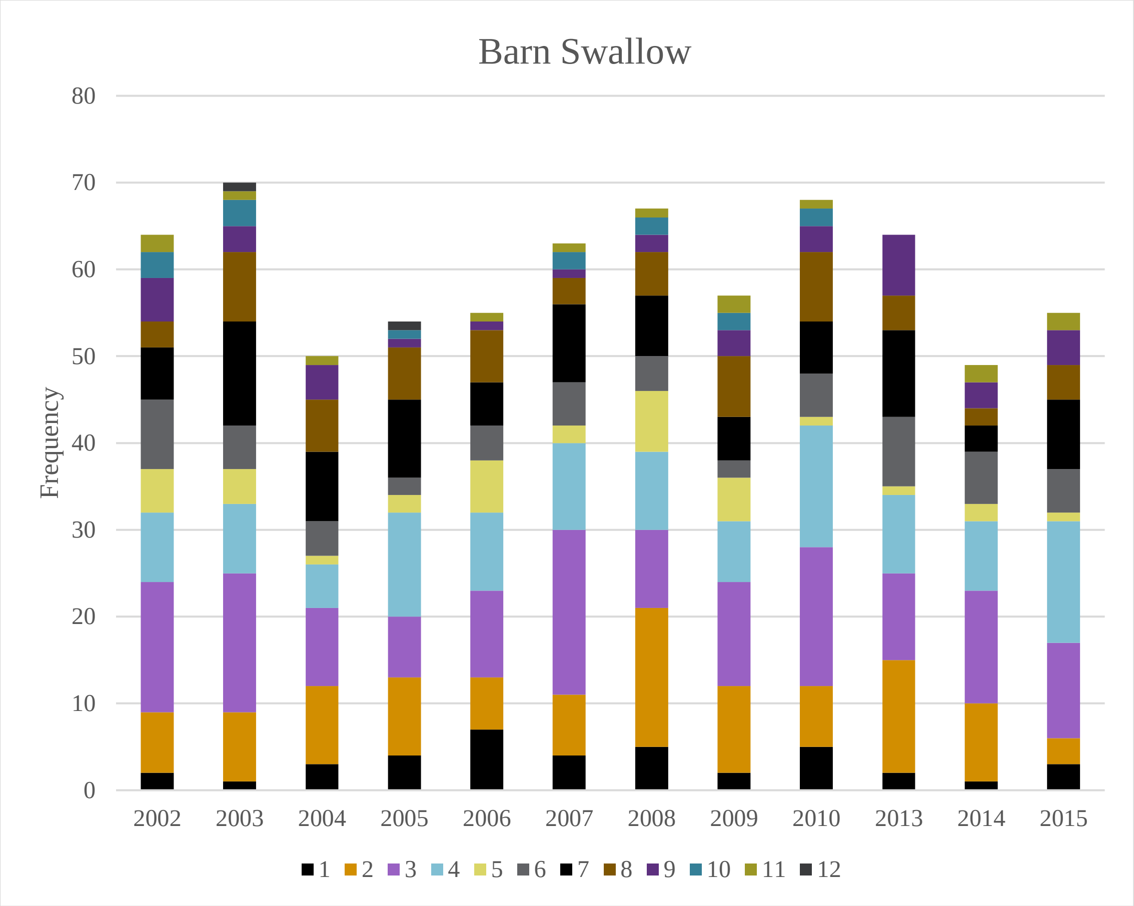
Barn Swallow
Frequency
0
10
20
30
40
50
60
70
80
2002
2003
2004
2005
2006
2007
2008
2009
2010
2013
2014
2015
1 2 3 4 5 6 7 8 9 10 11 12

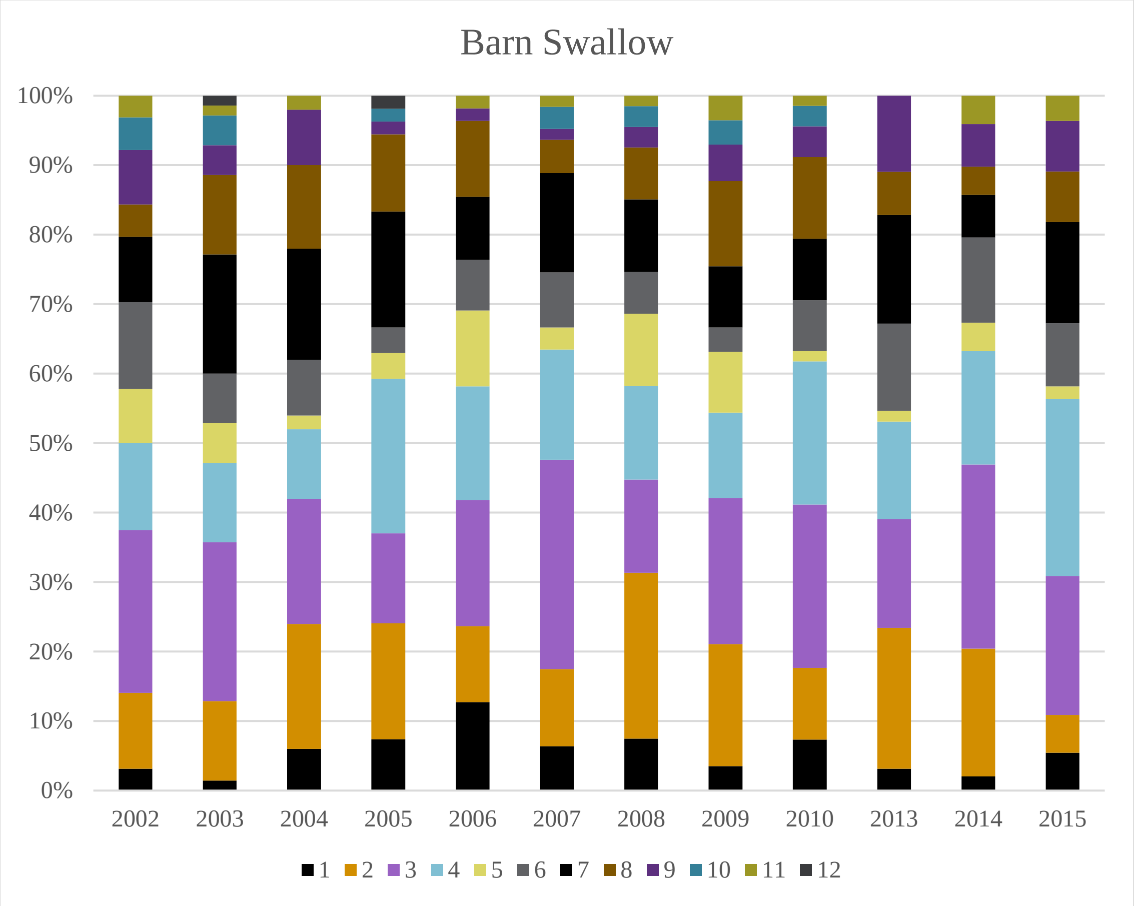
Barn Swallow
0%
10%
20%
30%
40%
50%
60%
70%
80%
90%
100%
2002
2003
2004
2005
2006
2007
2008
2009
2010
2013
2014
2015
1 2 3 4 5 6 7 8 9 10 11 12

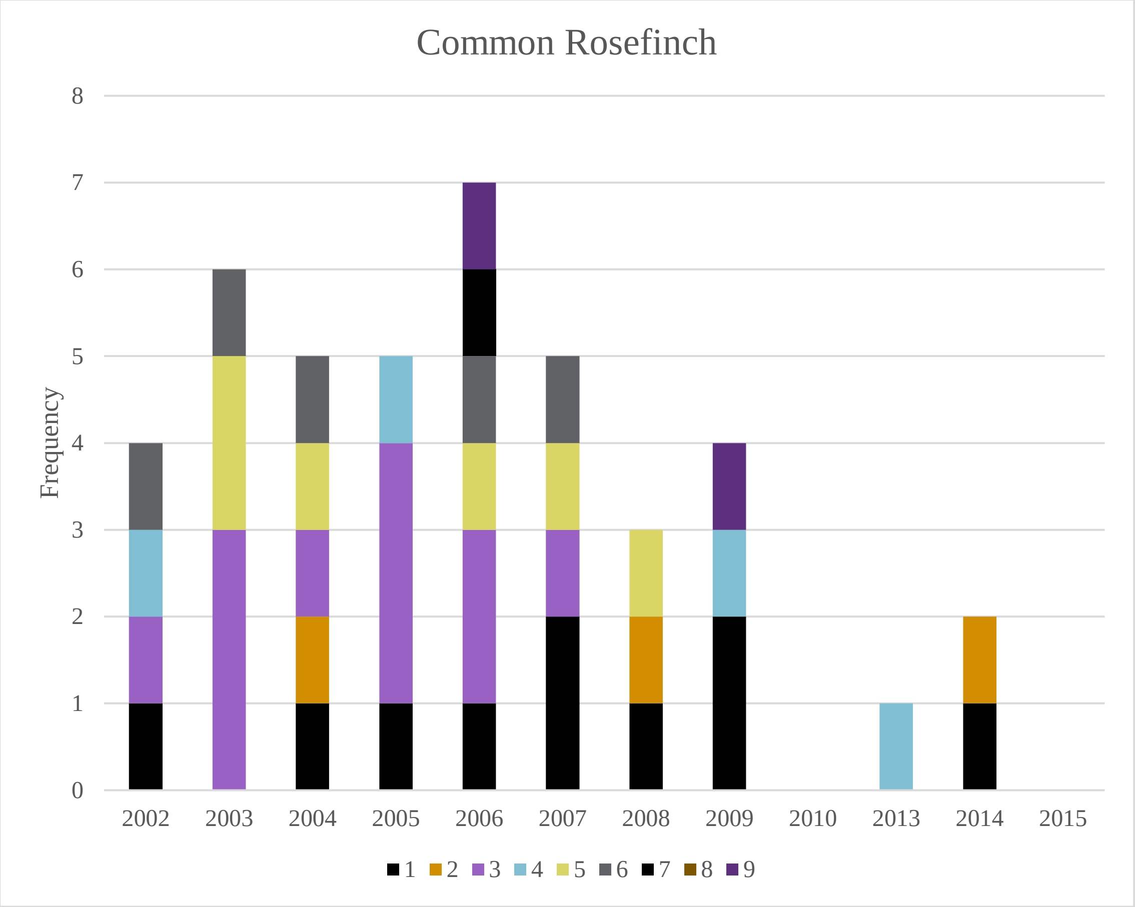
Common Rosefinch
Frequency
0
1
2
3
4
5
6
7
8
2002
2003
2004
2005
2006
2007
2008
2009
2010
2013
2014
2015
1 2 3 4 5 6 7 8 9

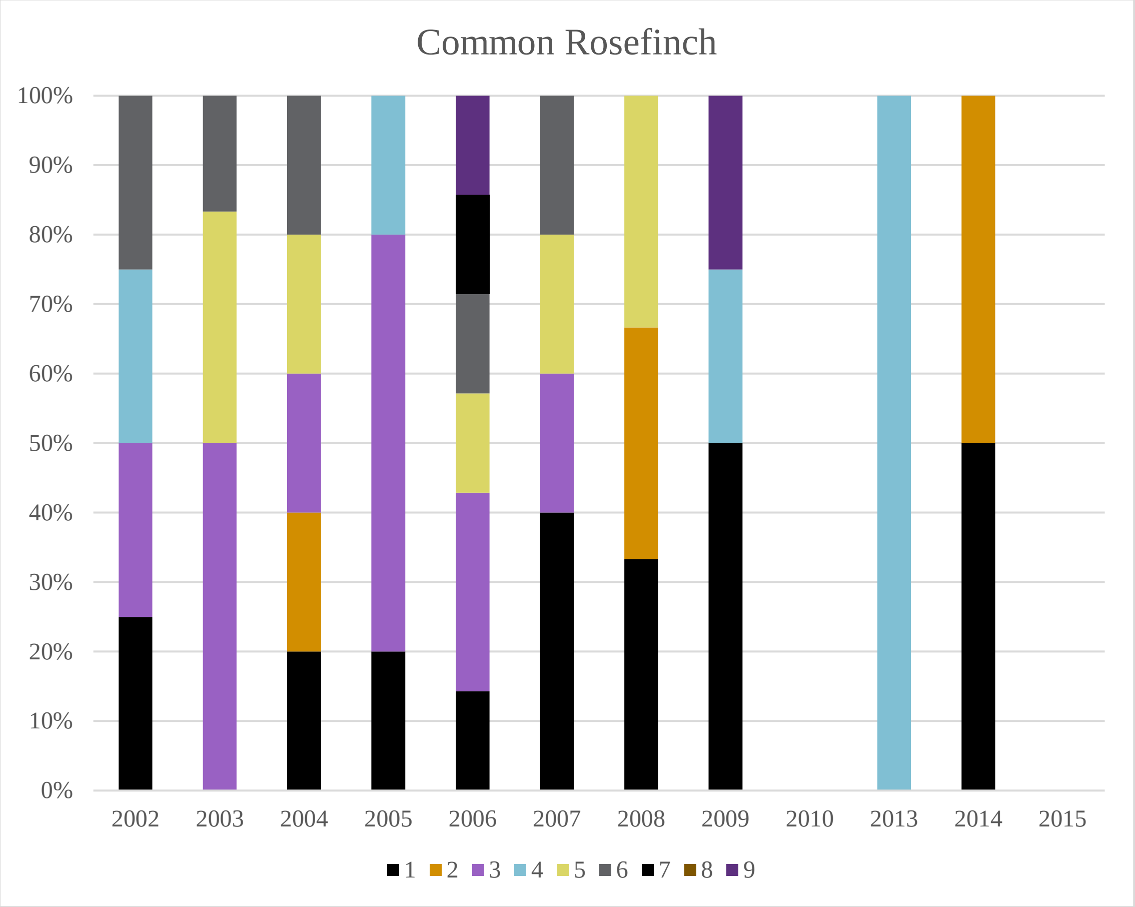
Common Rosefinch
0%
10%
20%
30%
40%
50%
60%
70%
80%
90%
100%
2002
2003
2004
2005
2006
2007
2008
2009
2010
2013
2014
2015
1 2 3 4 5 6 7 8 9

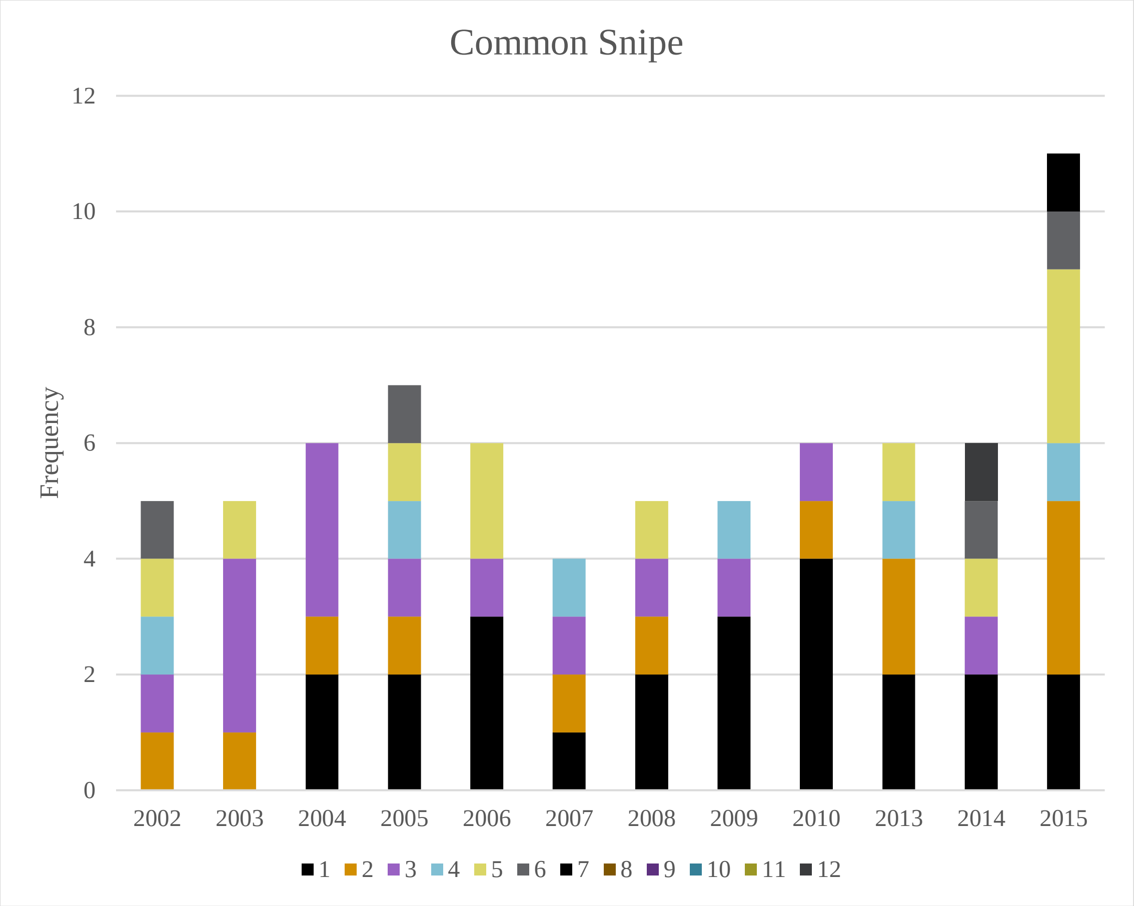
Common Snipe
Frequency
12
10
8
6
4
2
0
2002
2003
2004
2005
2006
2007
2008
2009
2010
2013
2014
2015
1 2 3 4 5 6 7 8 9 10 11 12

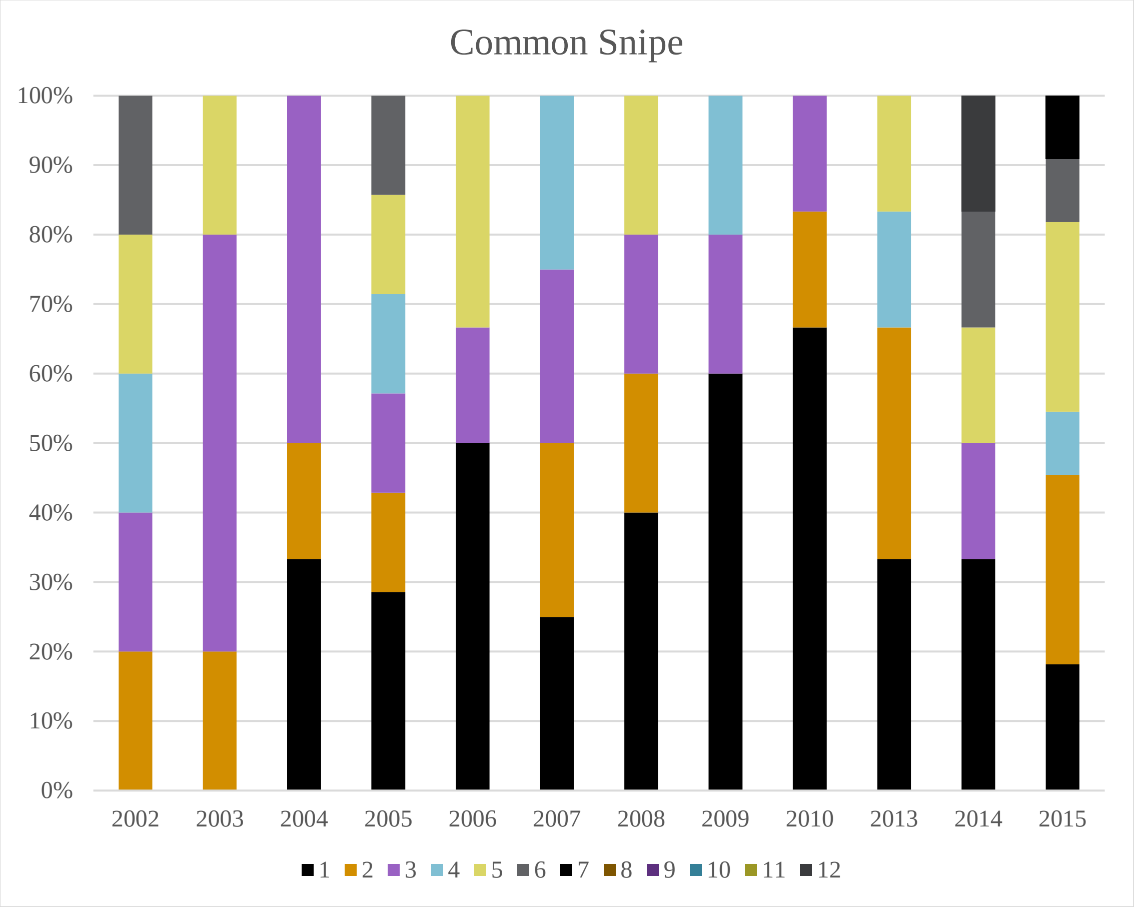
Common Snipe
100%
90%
80%
70%
60%
50%
40%
30%
20%
10%
0%
2002
2003
2004
2005
2006
2007
2008
2009
2010
2013
2014
2015
1 2 3 4 5 6 7 8 9 10 11 12

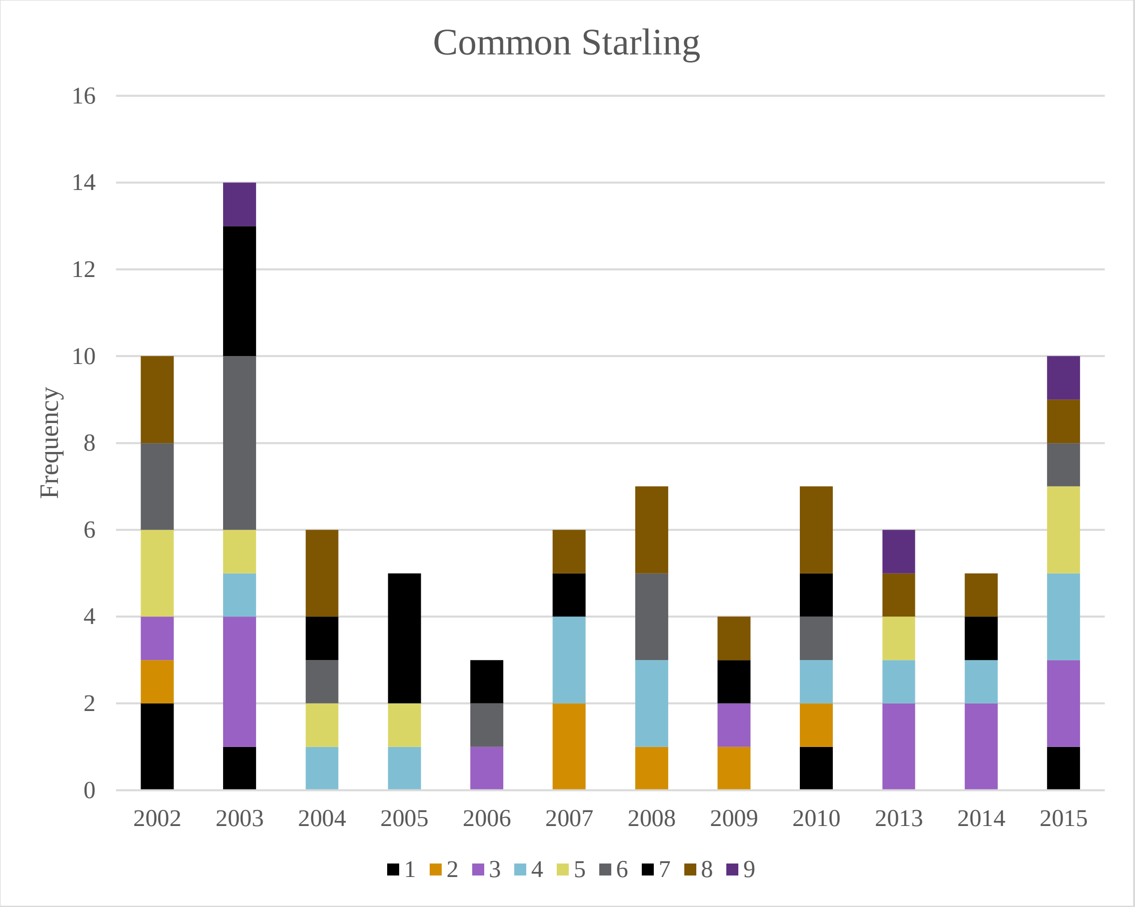
Common Starling
Frequency
16
14
12
10
8
6
4
2
0
2002
2003
2004
2005
2006
2007
2008
2009
2010
2013
2014
2015
1 2 3 4 5 6 7 8 9

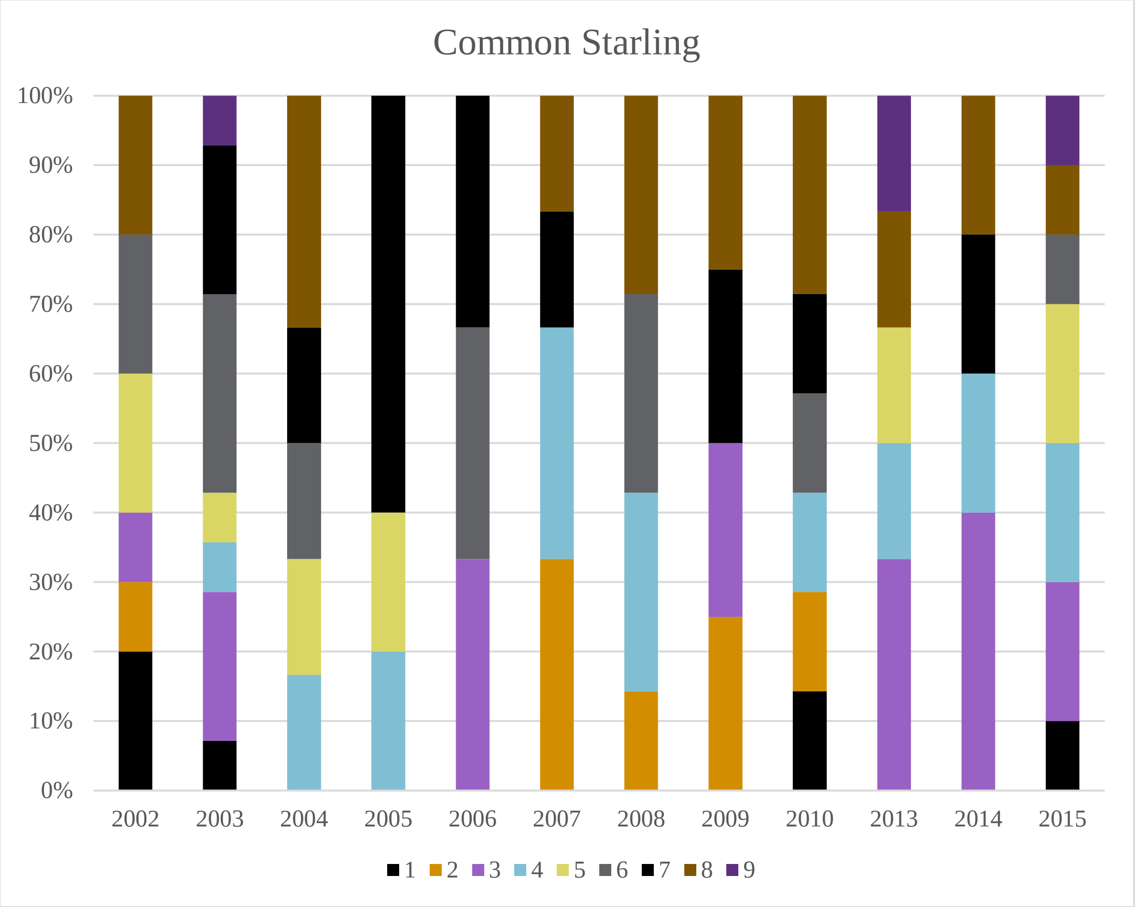
Common Starling
100%
90%
80%
70%
60%
50%
40%
30%
20%
10%
0%
2002
2003
2004
2005
2006
2007
2008
2009
2010
2013
2014
2015
1 2 3 4 5 6 7 8 9

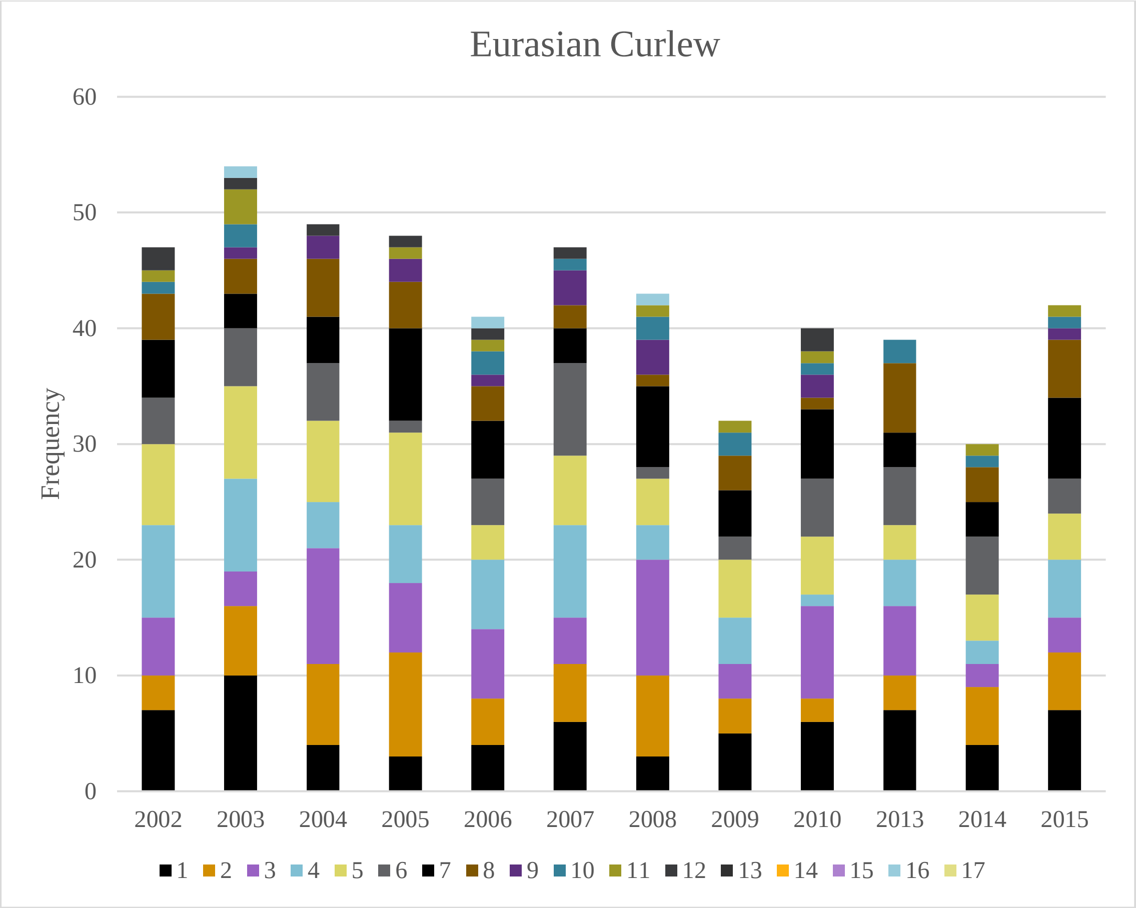

Eurasian Curlew
Frequency
0
10
20
30
40
50
60
2002
2003
2004
2005
2006
2007
2008
2009
2010
2013
2014
2015
1 2 3 4 5 6 7 8 9 10 11 12 13 14 15 16 17


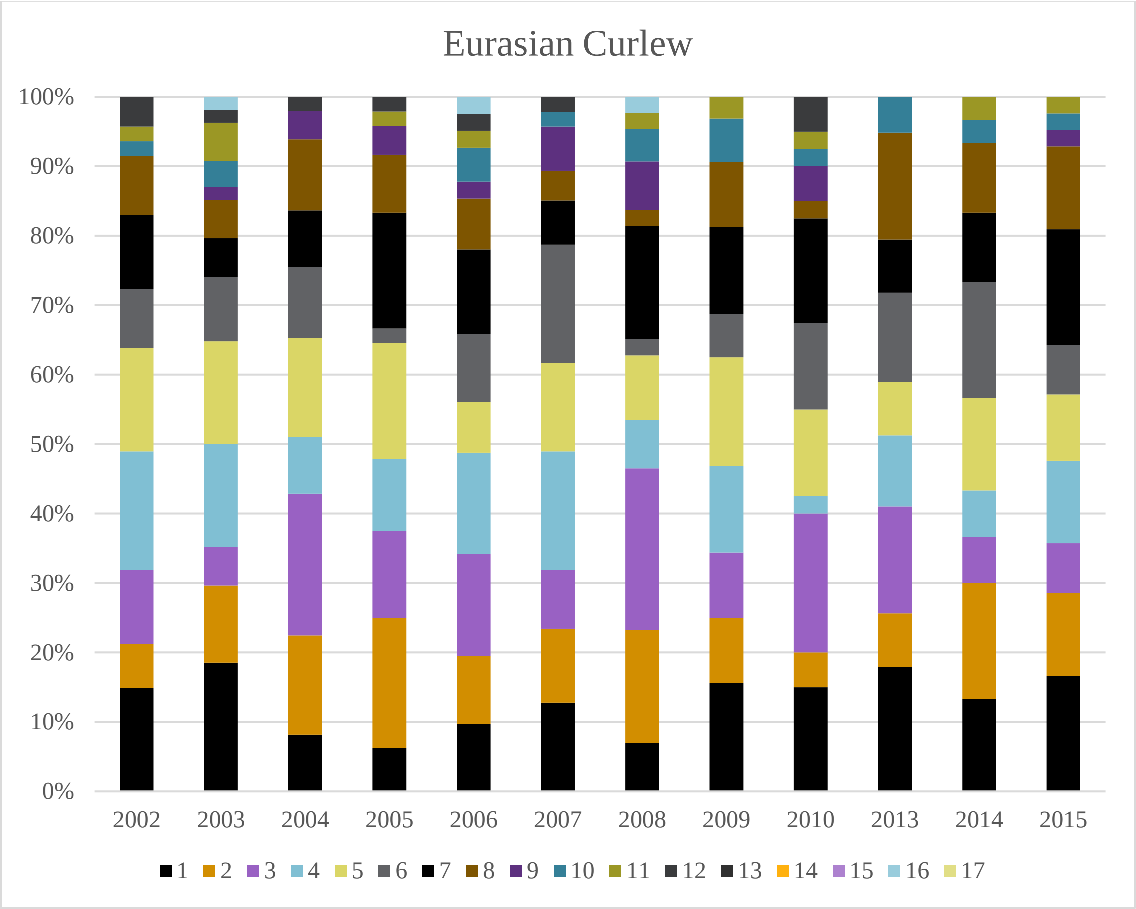

Eurasian Curlew
0%
10%
20%
30%
40%
50%
60%
70%
80%
90%
100%
2002
2003
2004
2005
2006
2007
2008
2009
2010
2013
2014
2015
1 2 3 4 5 6 7 8 9 10 11 12 13 14 15 16 17

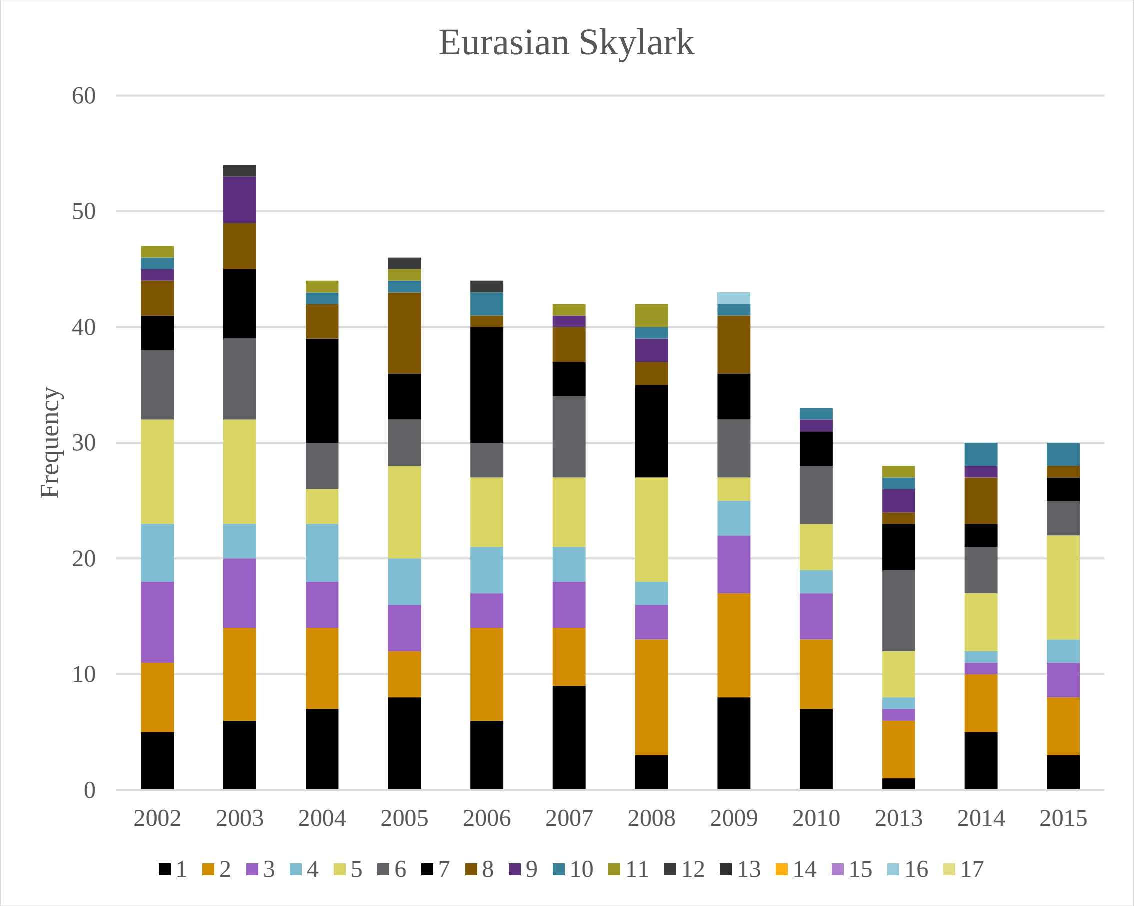
Eurasian Skylark
Frequency
60
50
40
30
20
10
0
2002
2003
2004
2005
2006
2007
2008
2009
2010
2013
2014
2015
1 2 3 4 5 6 7 8 9 10 11 12 13 14 15 16 17

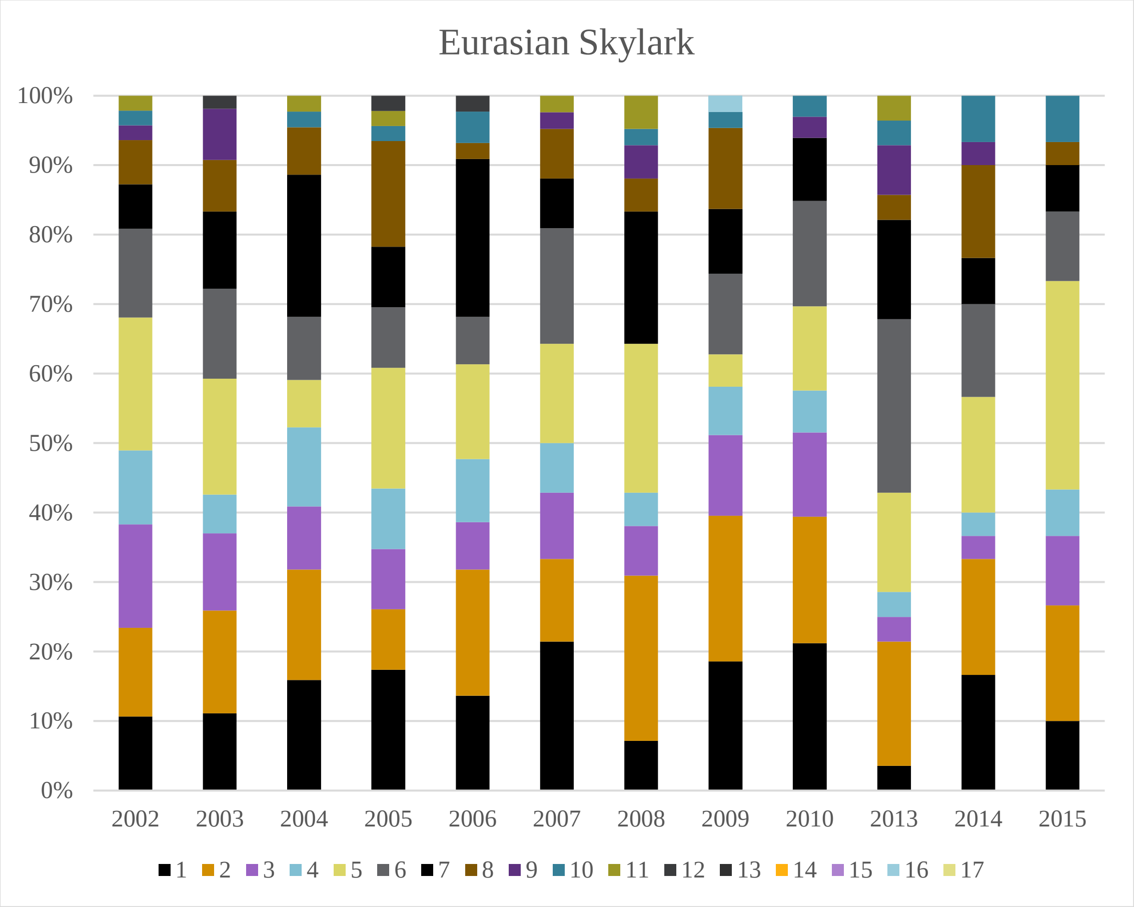
Eurasian Skylark
100%
90%
80%
70%
60%
50%
40%
30%
20%
10%
0%
2002
2003
2004
2005
2006
2007
2008
2009
2010
2013
2014
2015
1 2 3 4 5 6 7 8 9 10 11 12 13 14 15 16 17

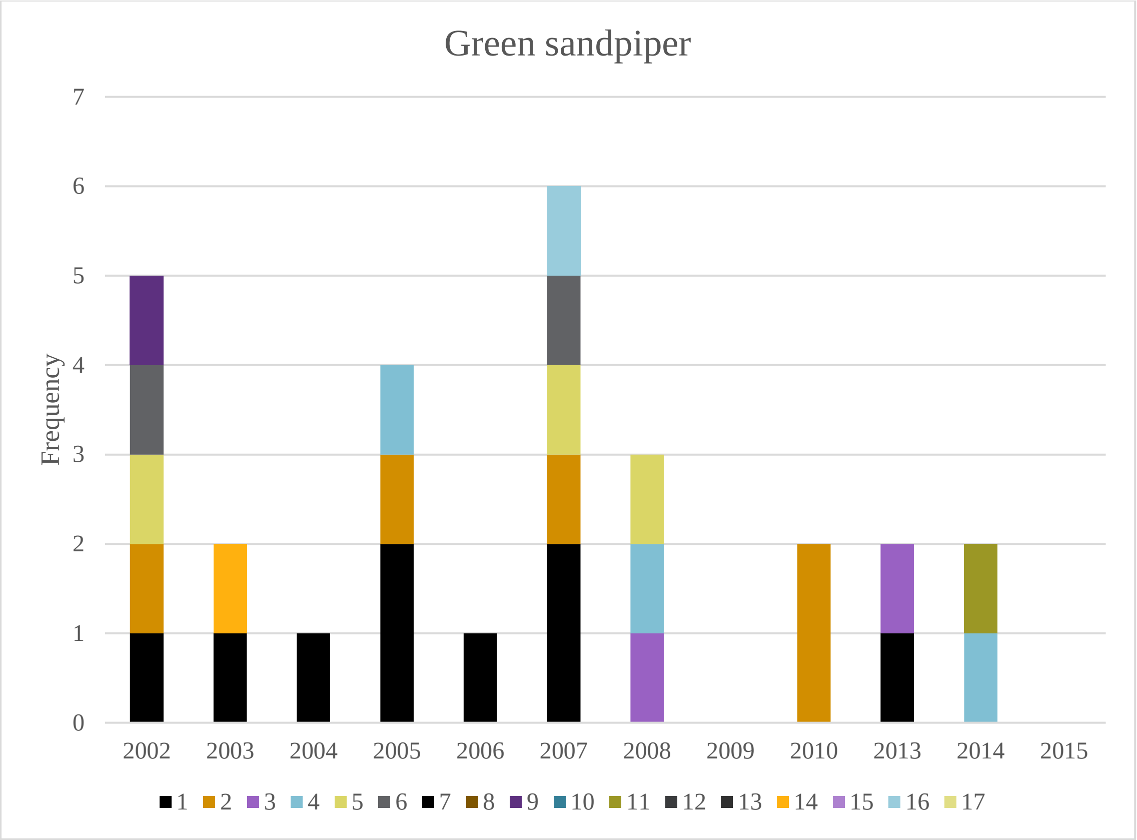
Green sandpiper
Frequency
0
1
2
3
4
5
6
7
2002
2003
2004
2005
2006
2007
2008
2009
2010
2013
2014
2015
1 2 3 4 5 6 7 8 9 10 11 12 13 14 15 16 17

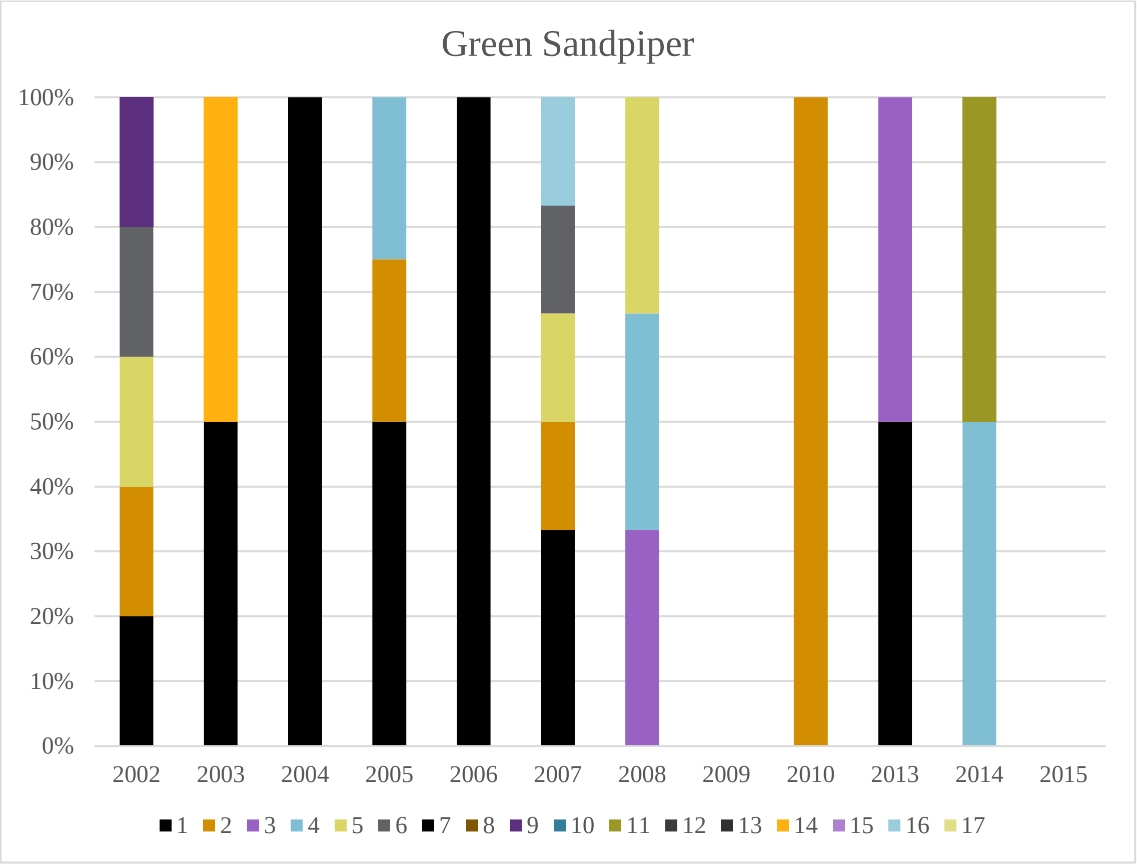
Green Sandpiper
0%
10%
20%
30%
40%
50%
60%
70%
80%
90%
100%
2002
2003
2004
2005
2006
2007
2008
2009
2010
2013
2014
2015
1 2 3 4 5 6 7 8 9 10 11 12 13 14 15 16 17

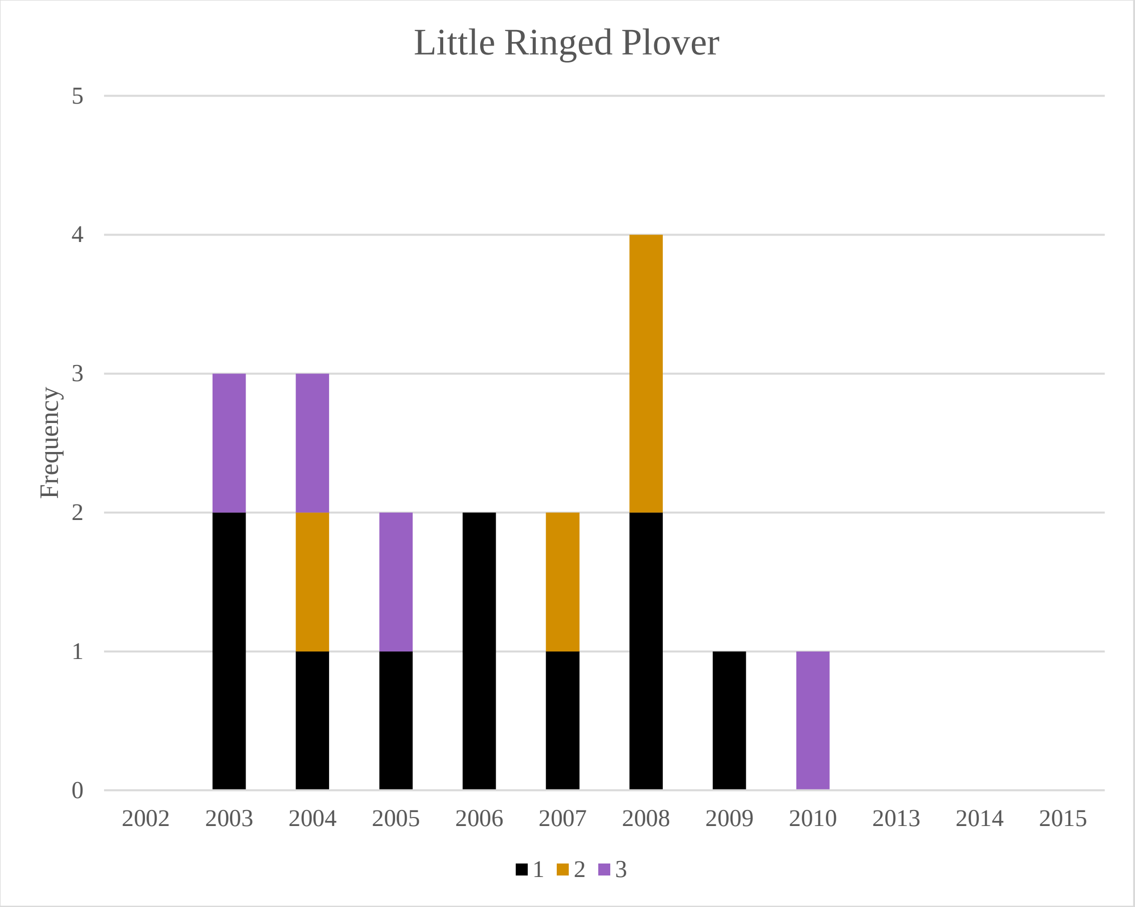
Little Ringed Plover
Frequency
5
4
3
2
1
0
2002
2003
2004
2005
2006
2007
2008
2009
2010
2013
2014
2015
1 2 3

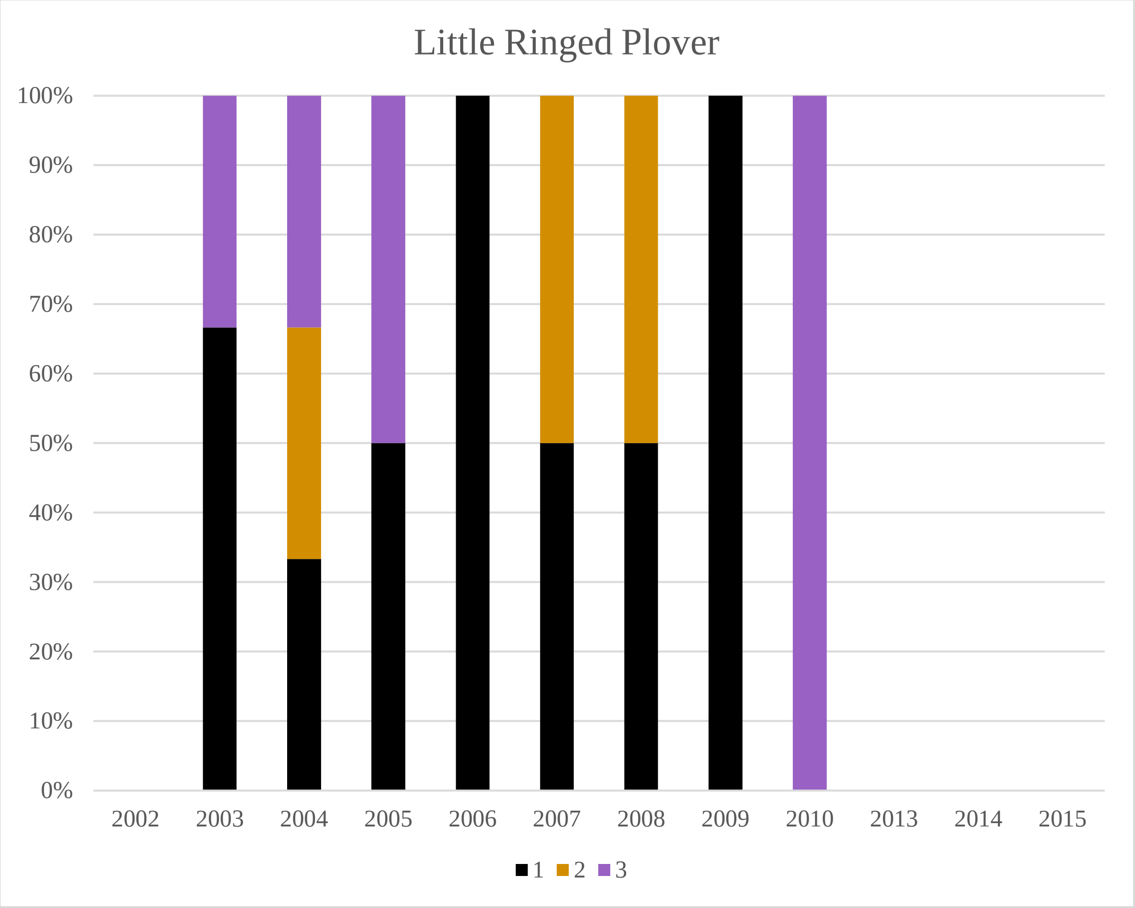
Little Ringed Plover
100%
90%
80%
70%
60%
50%
40%
30%
20%
10%
0%
2002
2003
2004
2005
2006
2007
2008
2009
2010
2013
2014
2015
1 2 3

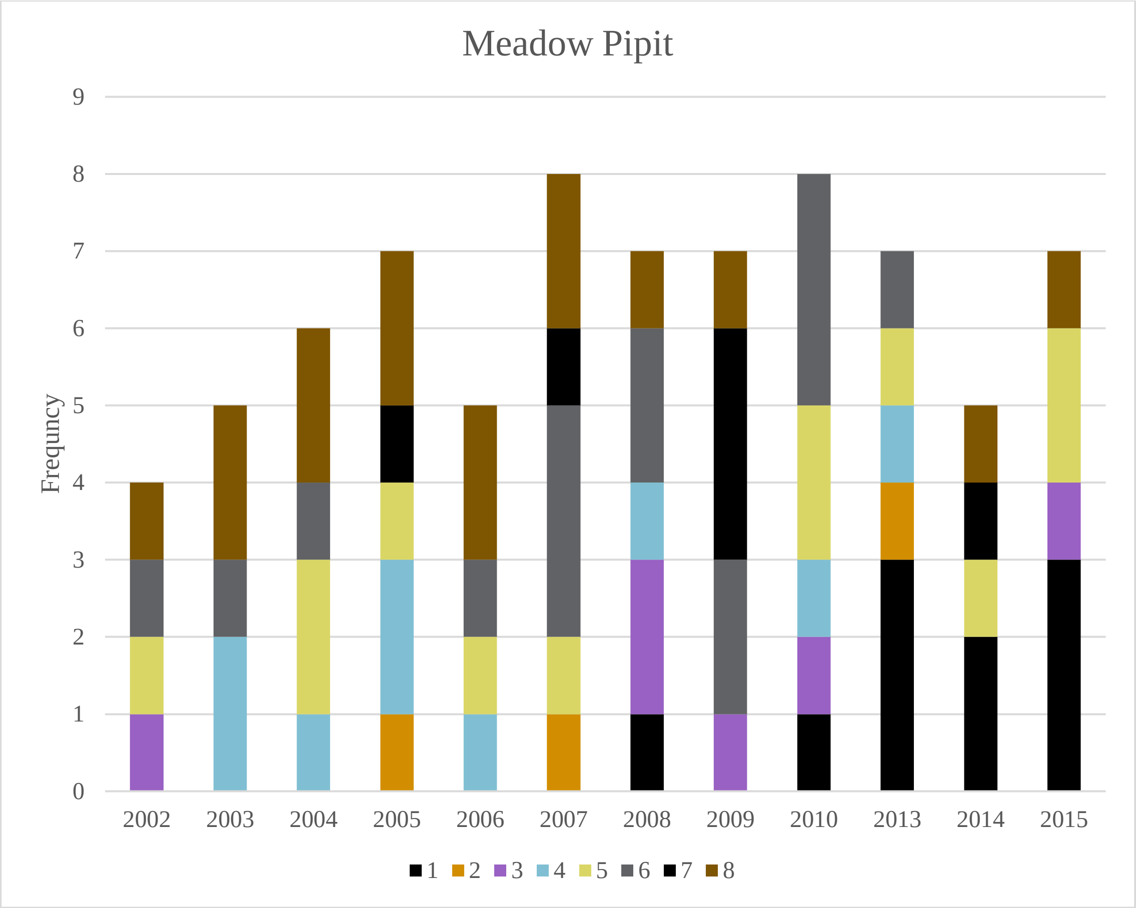
Meadow Pipit
Frequncy
9
8
7
6
5
4
3
2
1
0
2002
2003
2004
2005
2006
2007
2008
2009
2010
2013
2014
2015
1 2 3 4 5 6 7 8

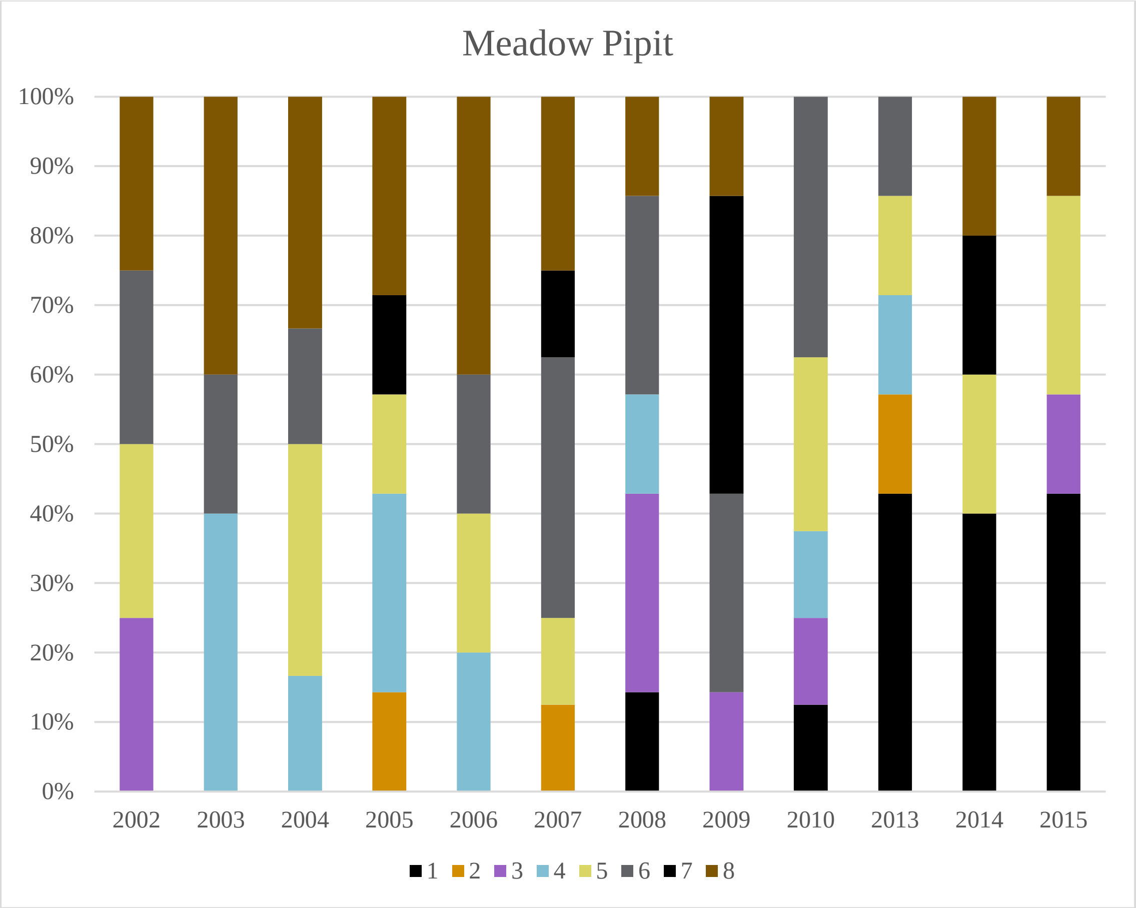
Meadow Pipit
100%
90%
80%
70%
60%
50%
40%
30%
20%
10%
0%
2002
2003
2004
2005
2006
2007
2008
2009
2010
2013
2014
2015
1 2 3 4 5 6 7 8

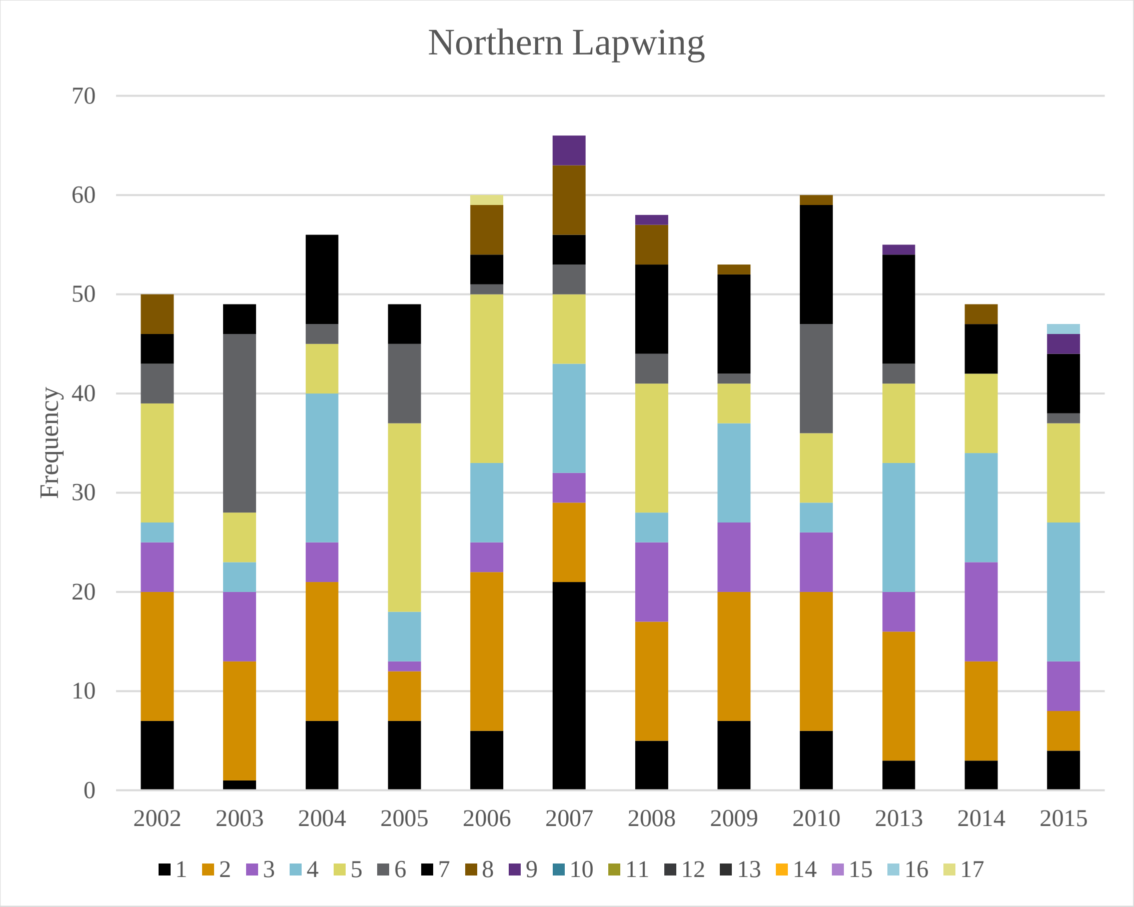
Northern Lapwing
Frequency
70
60
50
40
30
20
10
0
2002
2003
2004
2005
2006
2007
2008
2009
2010
2013
2014
2015
1 2 3 4 5 6 7 8 9 10 11 12 13 14 15 16 17

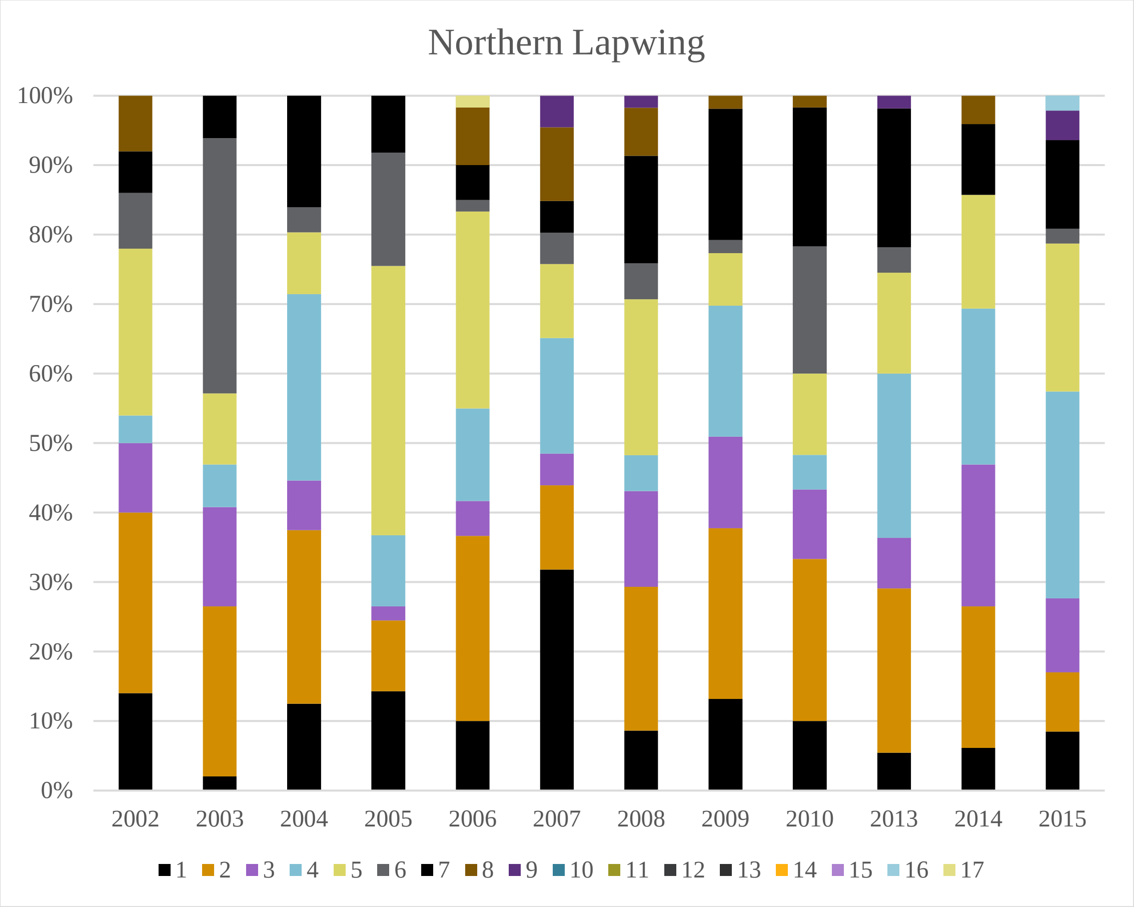
Northern Lapwing
100%
90%
80%
70%
60%
50%
40%
30%
20%
10%
0%
2002
2003
2004
2005
2006
2007
2008
2009
2010
2013
2014
2015
1 2 3 4 5 6 7 8 9 10 11 12 13 14 15 16 17

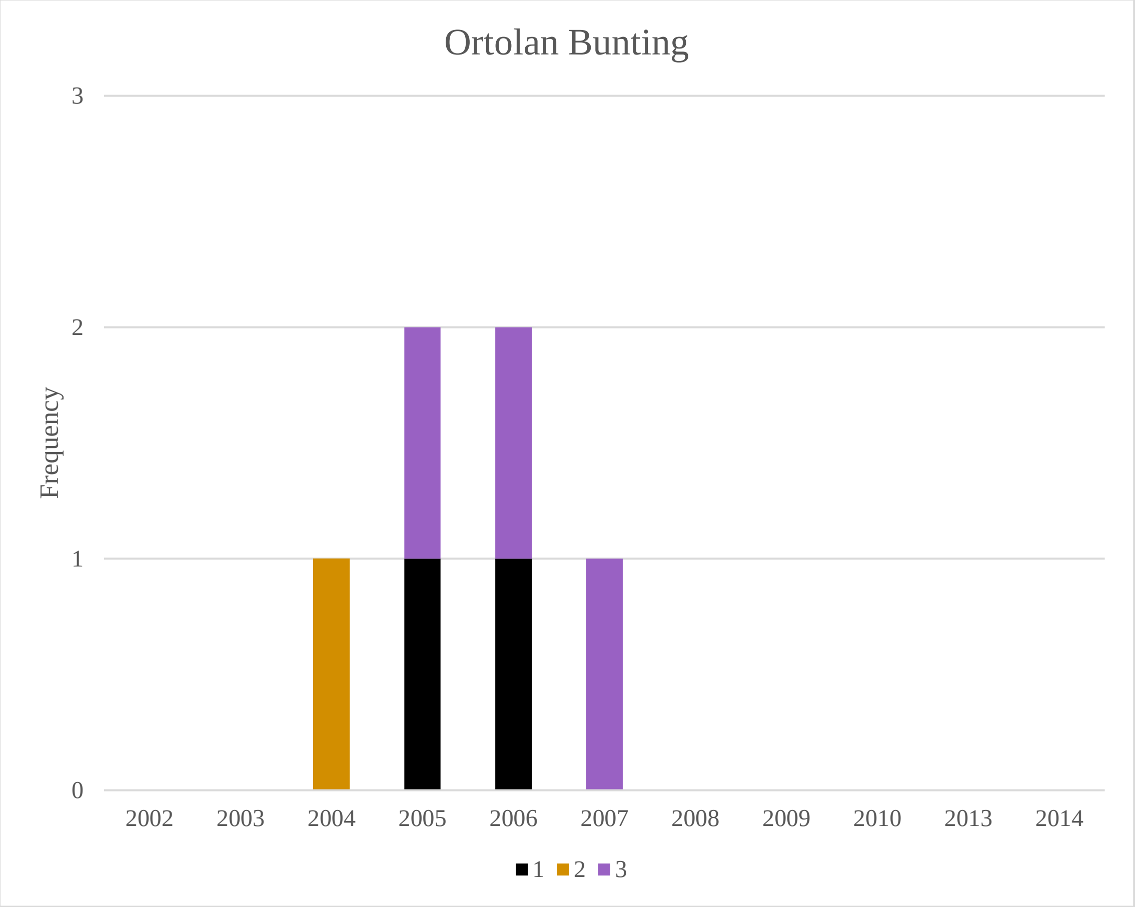
Ortolan Bunting
Frequency
3
2
1
0
2002
2003
2004
2005
2006
2007
2008
2009
2010
2013
2014
1
2
3

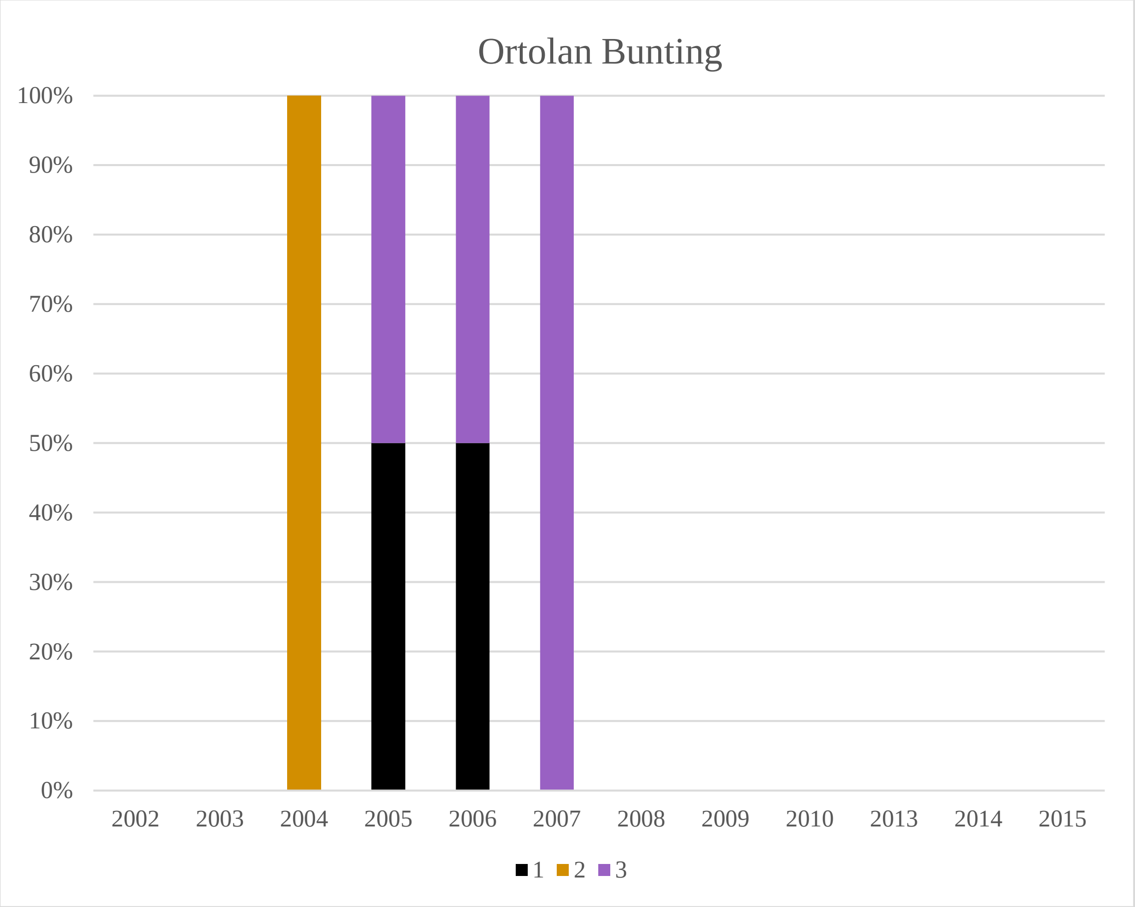
Ortolan Bunting
100%
90%
80%
70%
60%
50%
40%
30%
20%
10%
0%
2002
2003
2004
2005
2006
2007
2008
2009
2010
2013
2014
2015
1
2
3

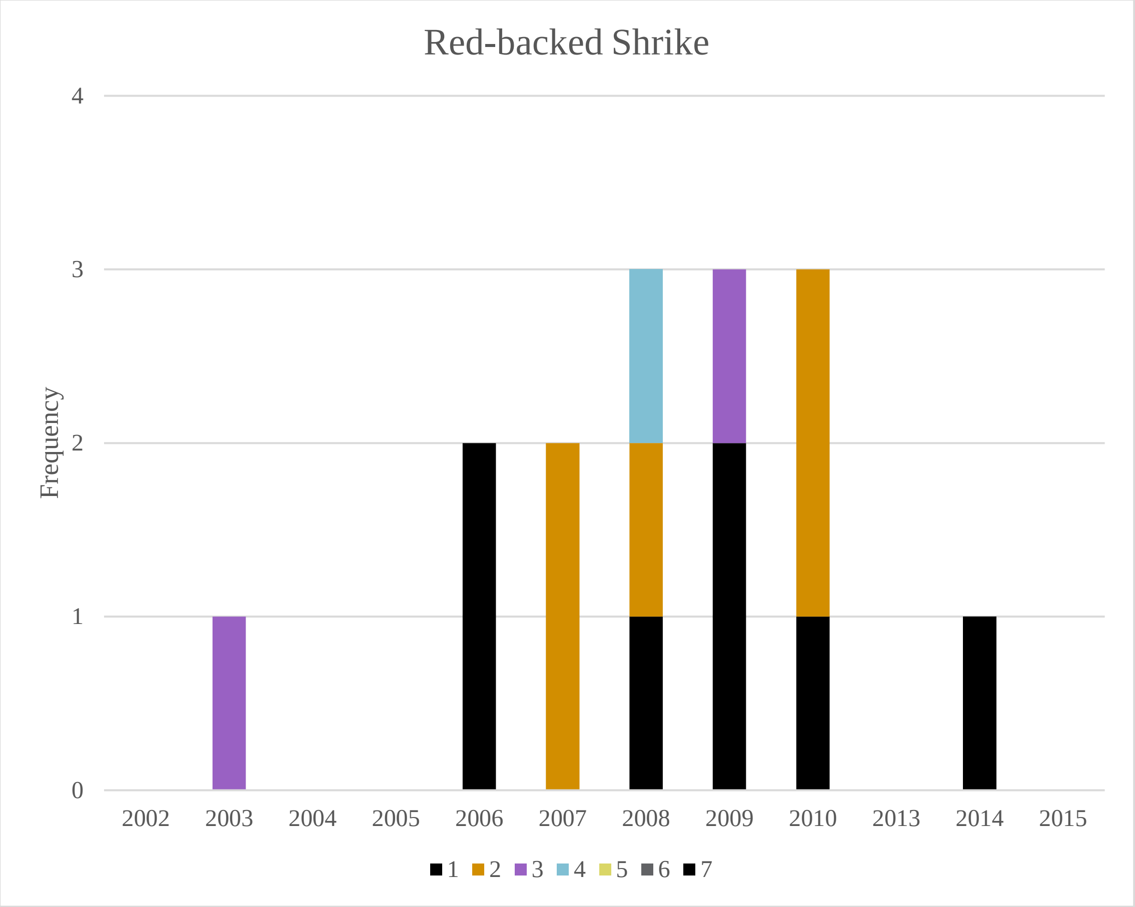
Red-backed Shrike
Frequency
0
1
2
3
4
2002
2003
2004
2005
2006
2007
2008
2009
2010
2013
2014
2015
1
2
3
4
5
6
7

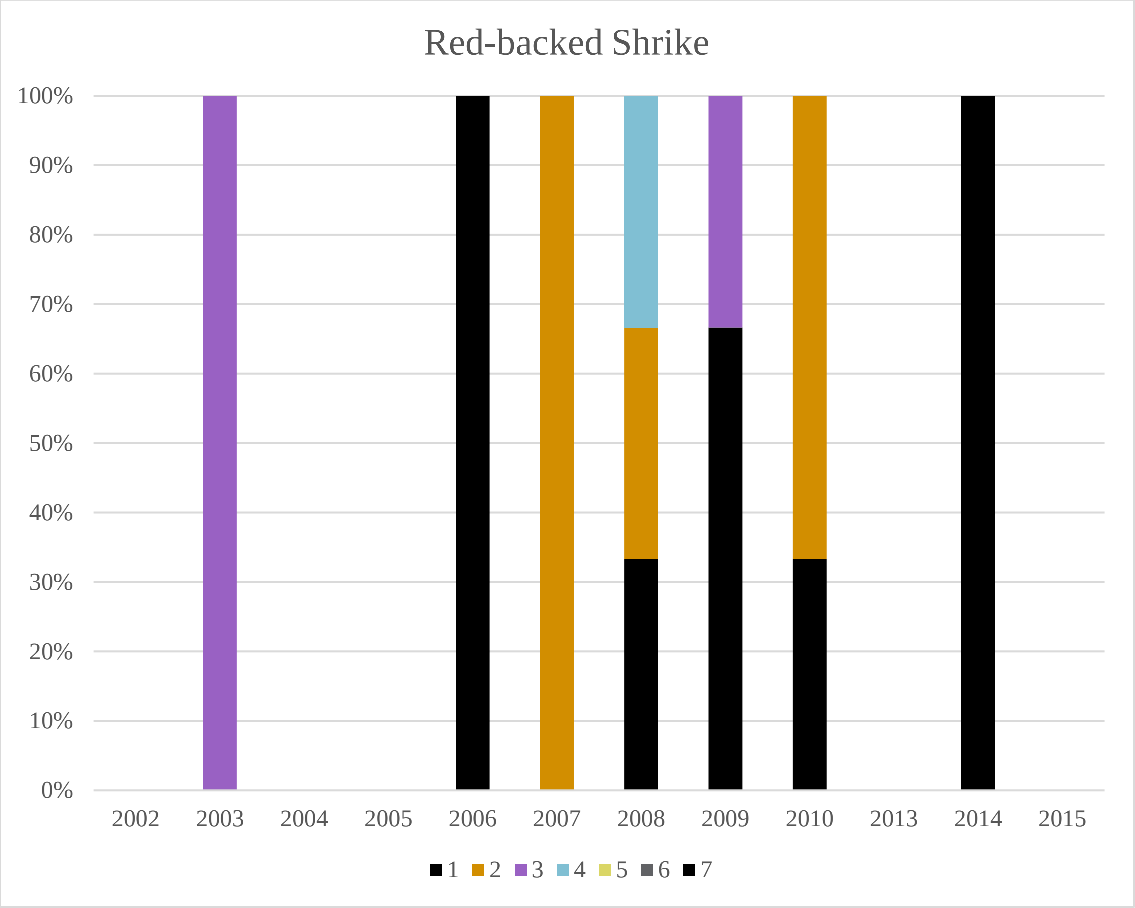
Red-backed Shrike
0%
10%
20%
30%
40%
50%
60%
70%
80%
90%
100%
2002
2003
2004
2005
2006
2007
2008
2009
2010
2013
2014
2015
1
2
3
4
5
6
7

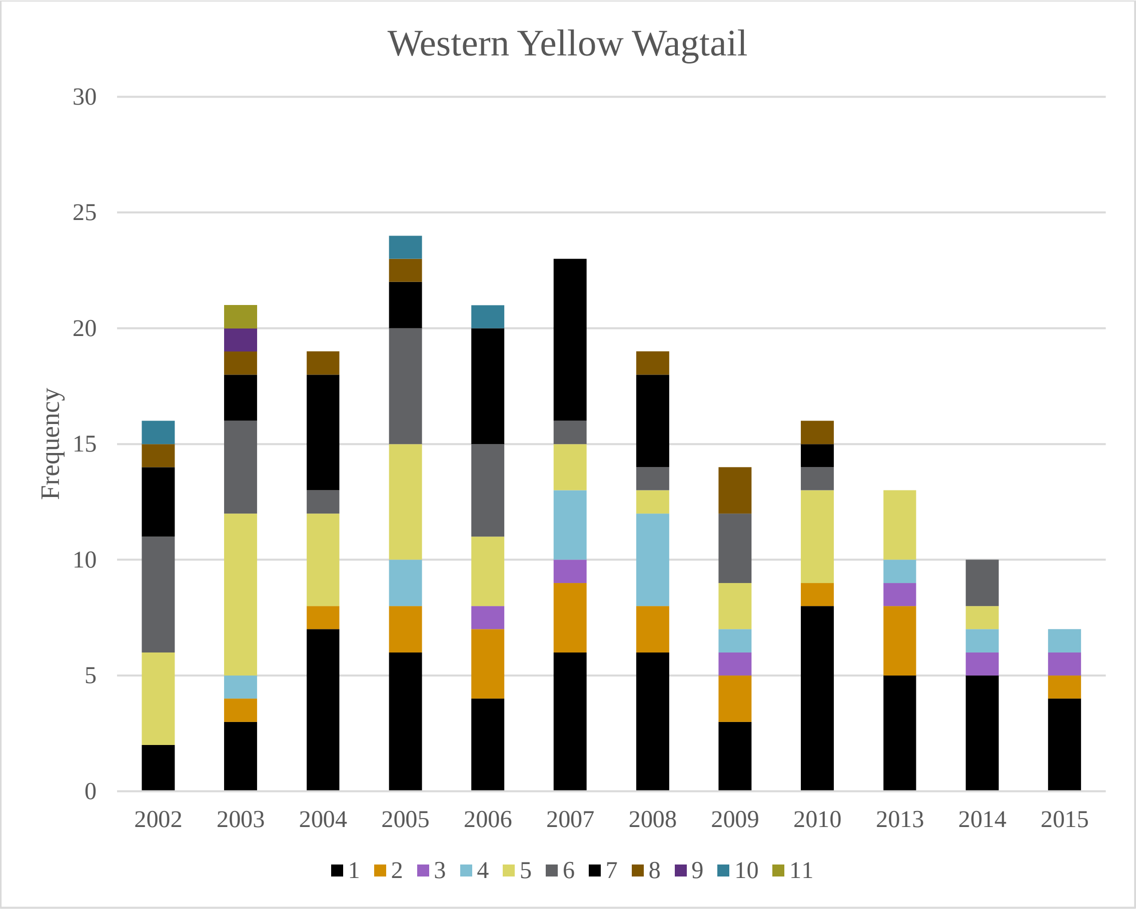
Western Yellow Wagtail
Frequency
30
25
20
15
10
5
0
2002
2003
2004
2005
2006
2007
2008
2009
2010
2013
2014
2015
1 2 3 4 5 6 7 8 9 10 11

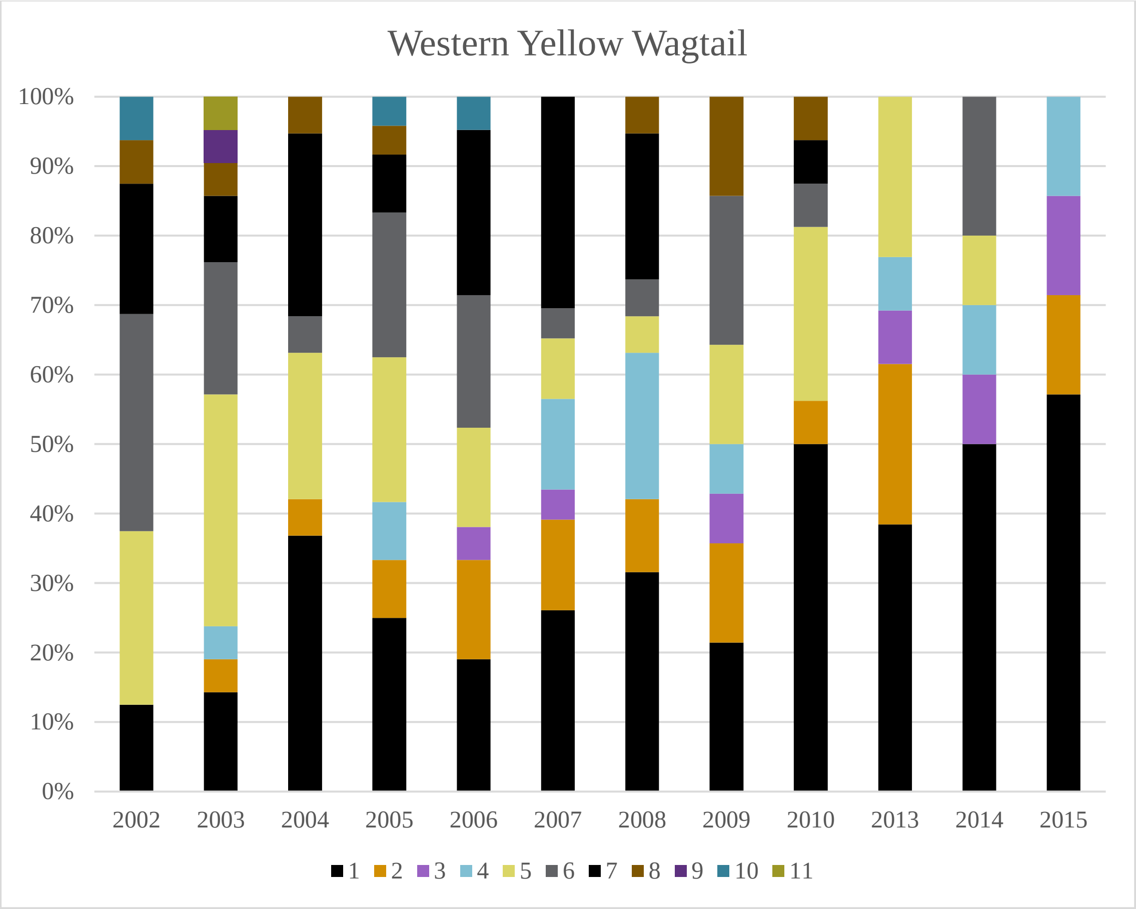
Western Yellow Wagtail
100%
90%
80%
70%
60%
50%
40%
30%
20%
10%
0%
2002
2003
2004
2005
2006
2007
2008
2009
2010
2013
2014
2015
1 2 3 4 5 6 7 8 9 10 11

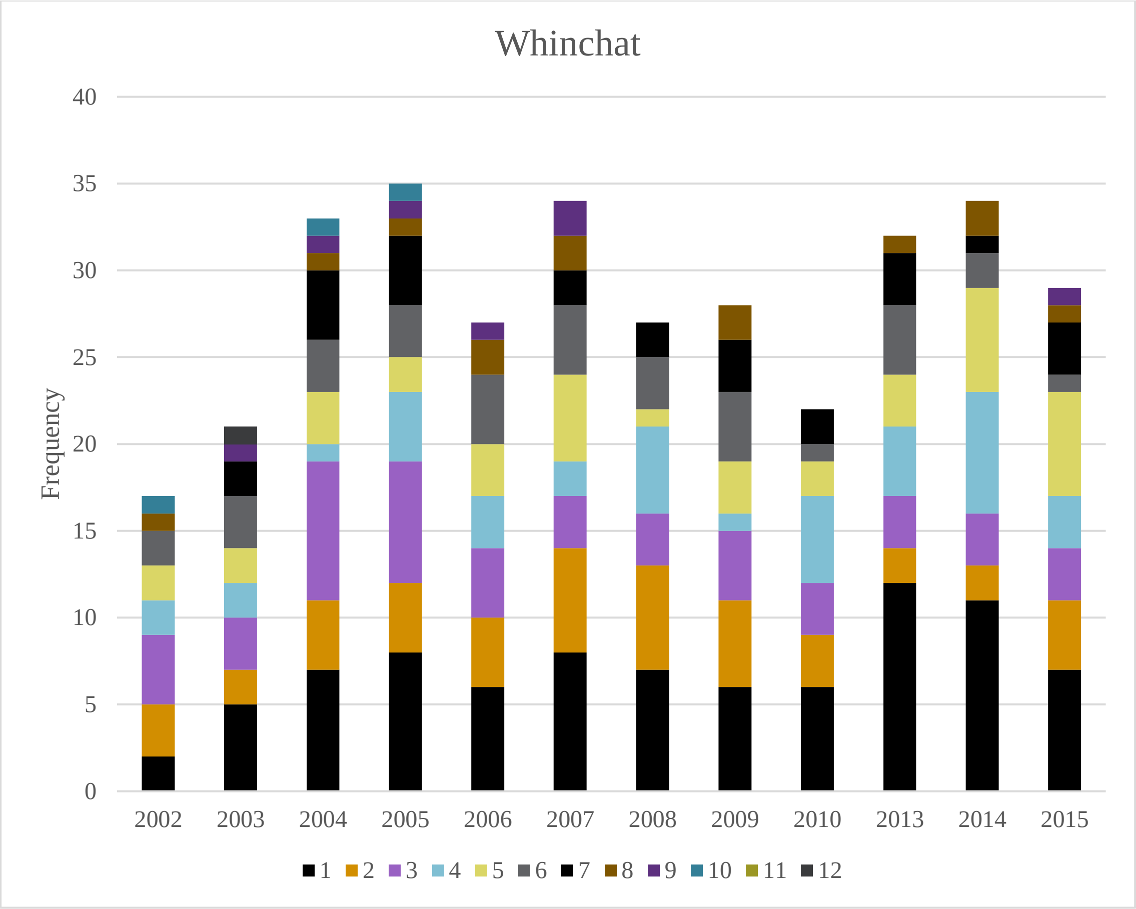
Whinchat
Frequency
40
35
30
25
20
15
10
5
0
2002 2003 2004 2005 2006 2007 2008 2009 2010 2013 2014 2015
1 2 3 4 5 6 7 8 9 10 11 12

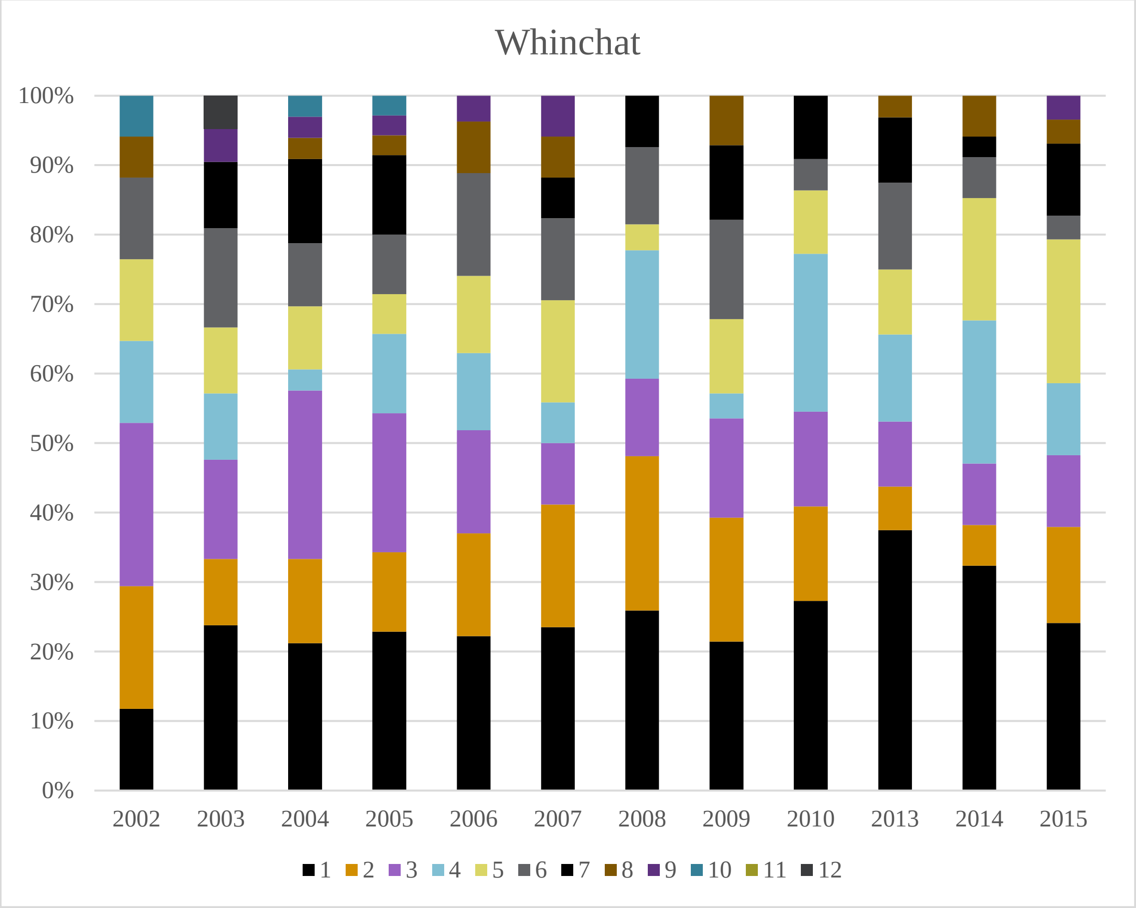
Whinchat
100%
90%
80%
70%
60%
50%
40%
30%
20%
10%
0%
2002 2003 2004 2005 2006 2007 2008 2009 2010 2013 2014 2015
1 2 3 4 5 6 7 8 9 10 11 12

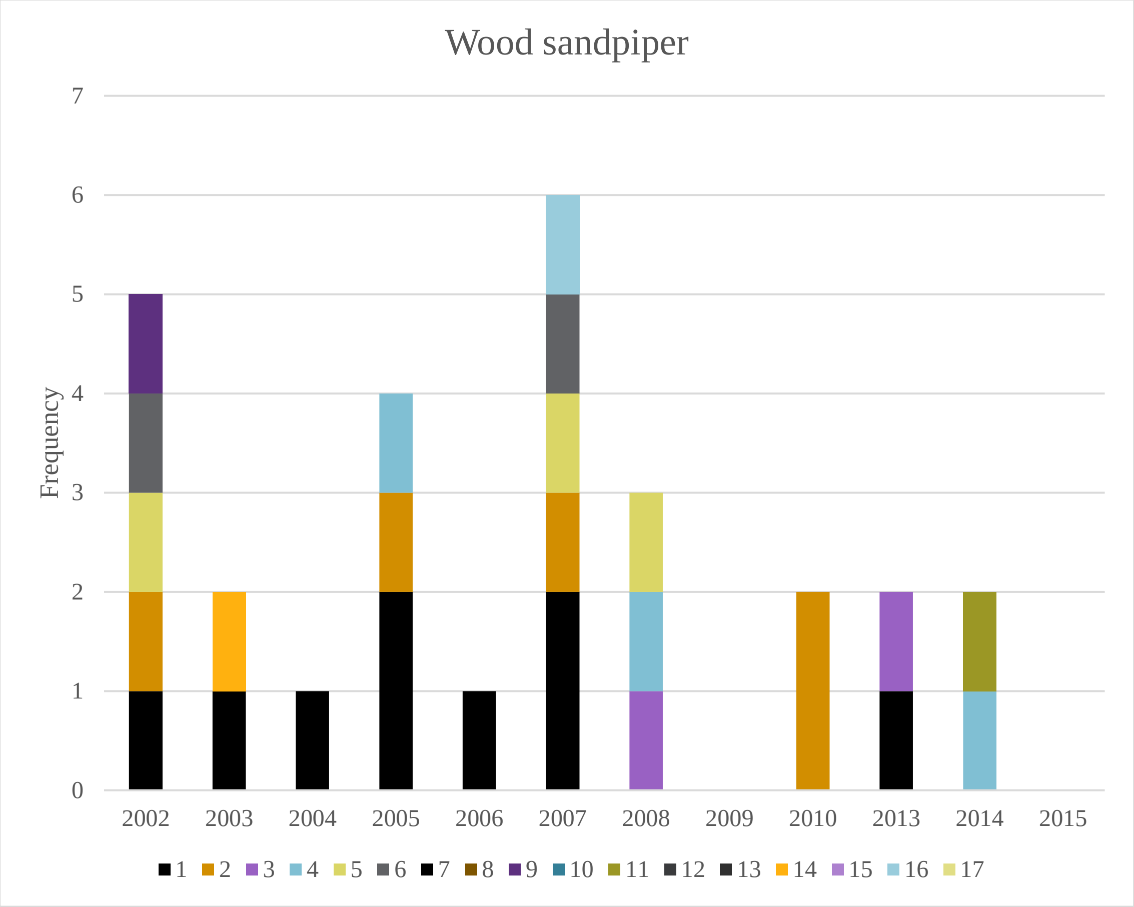
Wood sandpiper
Frequency
0
1
2
3
4
5
6
7
2002
2003
2004
2005
2006
2007
2008
2009
2010
2013
2014
2015
1 2 3 4 5 6 7 8 9 10 11 12 13 14 15 16 17

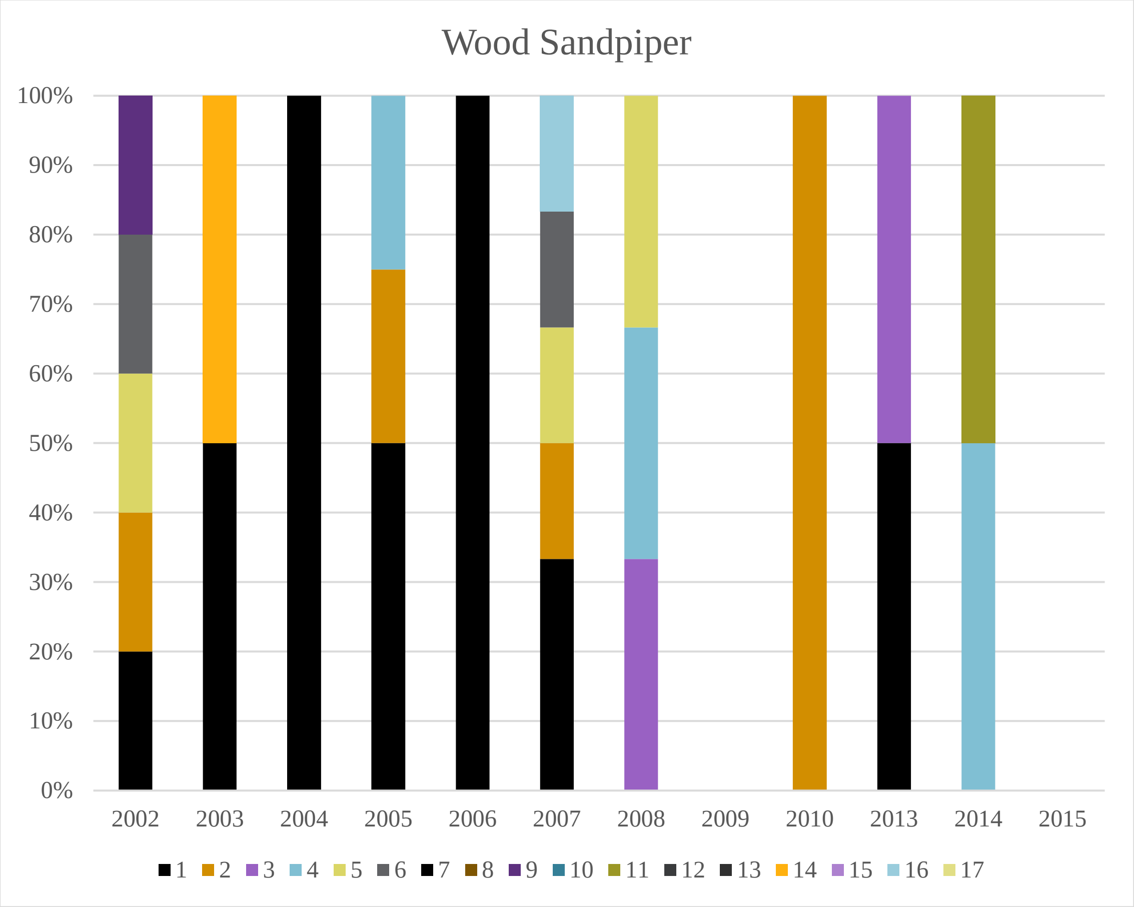
Wood Sandpiper
0%
10%
20%
30%
40%
50%
60%
70%
80%
90%
100%
2002
2003
2004
2005
2006
2007
2008
2009
2010
2013
2014
2015
1 2 3 4 5 6 7 8 9 10 11 12 13 14 15 16 17
