## Supplementary material for "Are railways detrimental to bird populations? A BDACI study on the construction of the Bothnia Line Railway": Trends in distances in 300 and 500 m zones to BLR

Supplementary material **Dist3**

### Temporal trends and Kendall's rank correlation tests for the proportions of territories of individual study species within 300 and 500 m from the Bothnia Line Railway.

Most correlation tests generated warning messages due to ties (data points with the same Y-value). The warning texts were omitted from the output text for readability.

Overlapping data points were jittered along the x-axis.

Significant (*P*<0.05) correlations are marked with yellow background.

Species with insufficient numbers of data points were omitted.

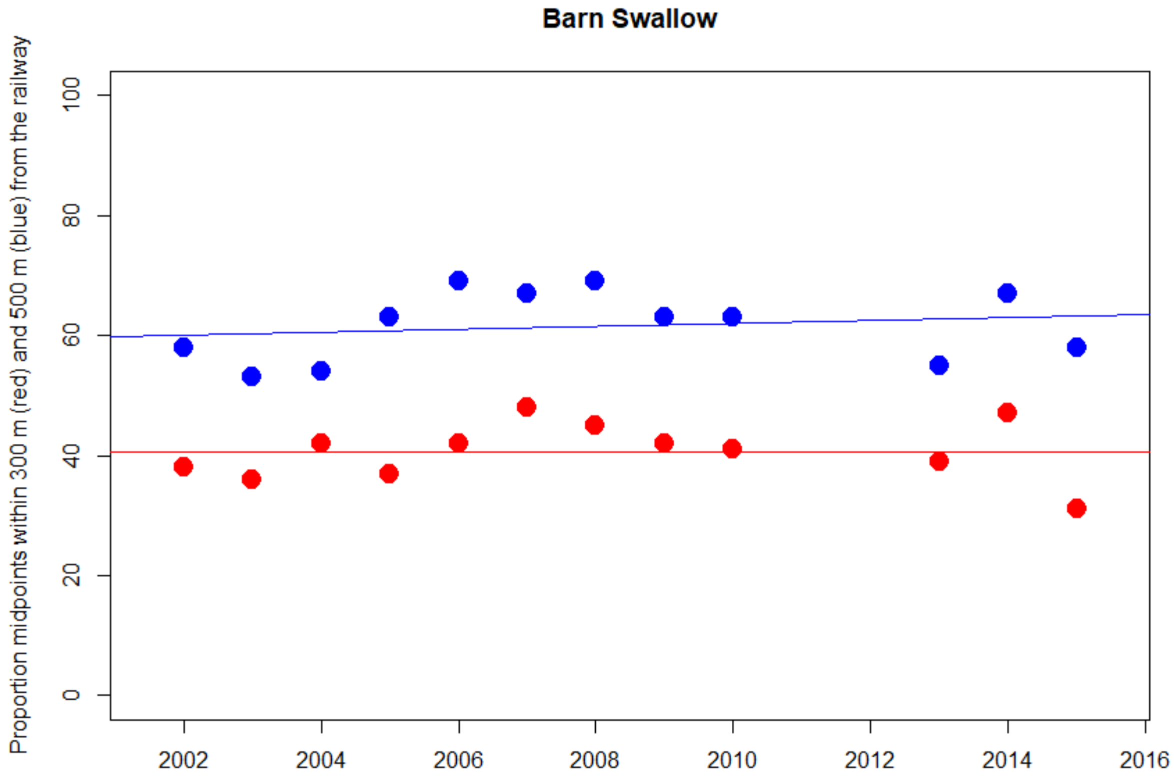


```
300 m. red
Kendall's rank correlation tau = 0.078
z = 0.346, p-value = 0.730

500 m. blue
Kendall's rank correlation tau = 0.159
z = 0.697, p-value = 0.486
```

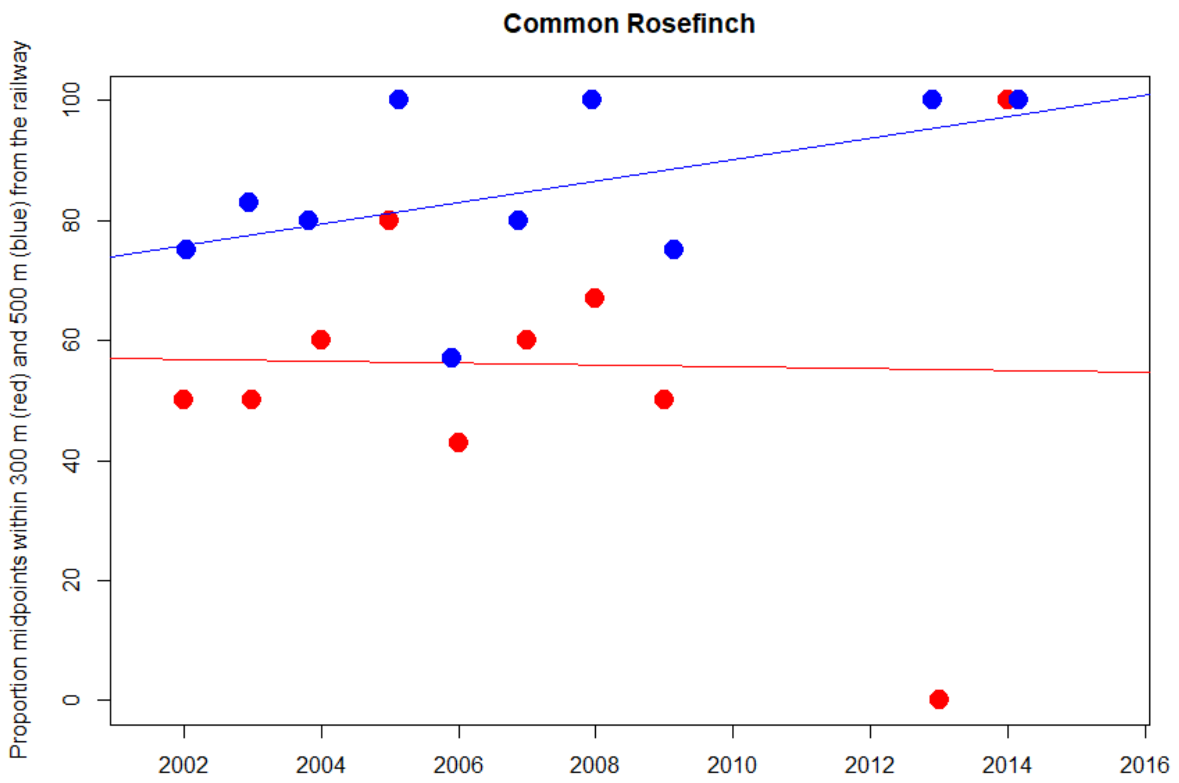


```
300 m. red
Kendall's rank correlation tau = 0.116
z = 0.456, p-value = 0.649

500 m. blue
Kendall's rank correlation tau = 0.319
z = 1.216, p-value = 0.224
```

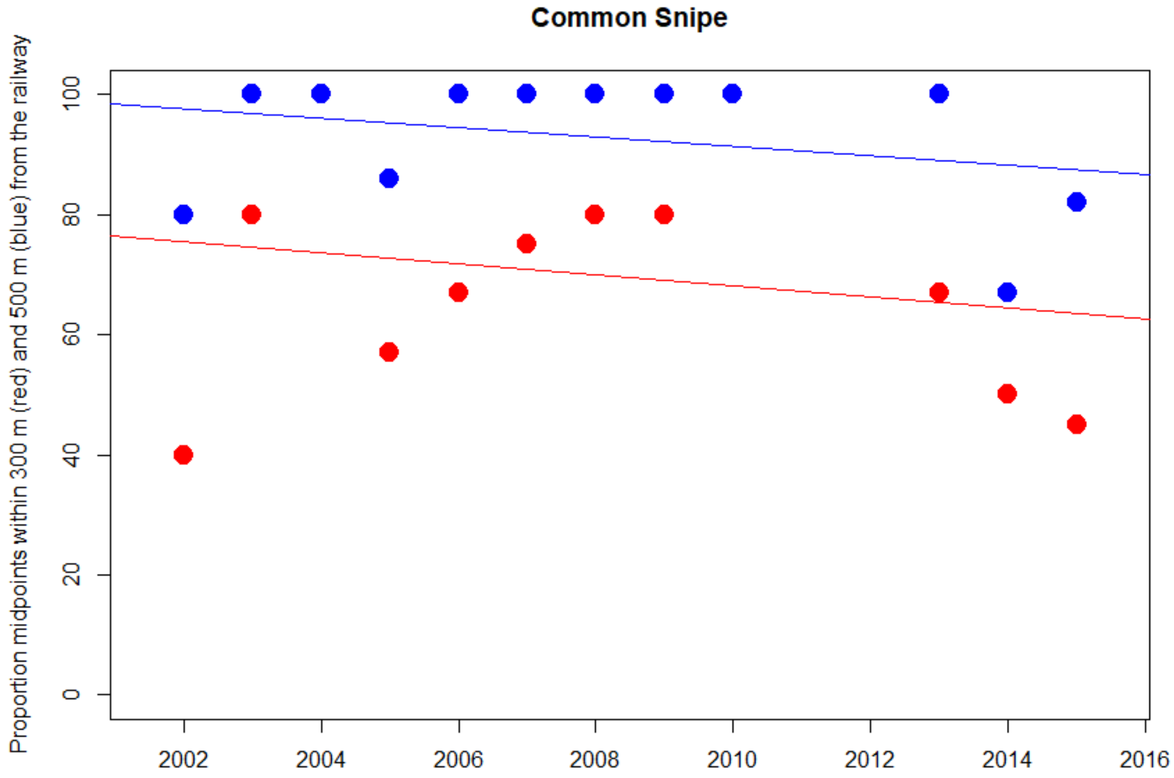


```
300 m. red
Kendall's rank correlation tau = -0.079
z = -0.348, p-value = 0.728

500 m. blue
Kendall's rank correlation tau = -0.080
z = -0.330, p-value = 0.742
```

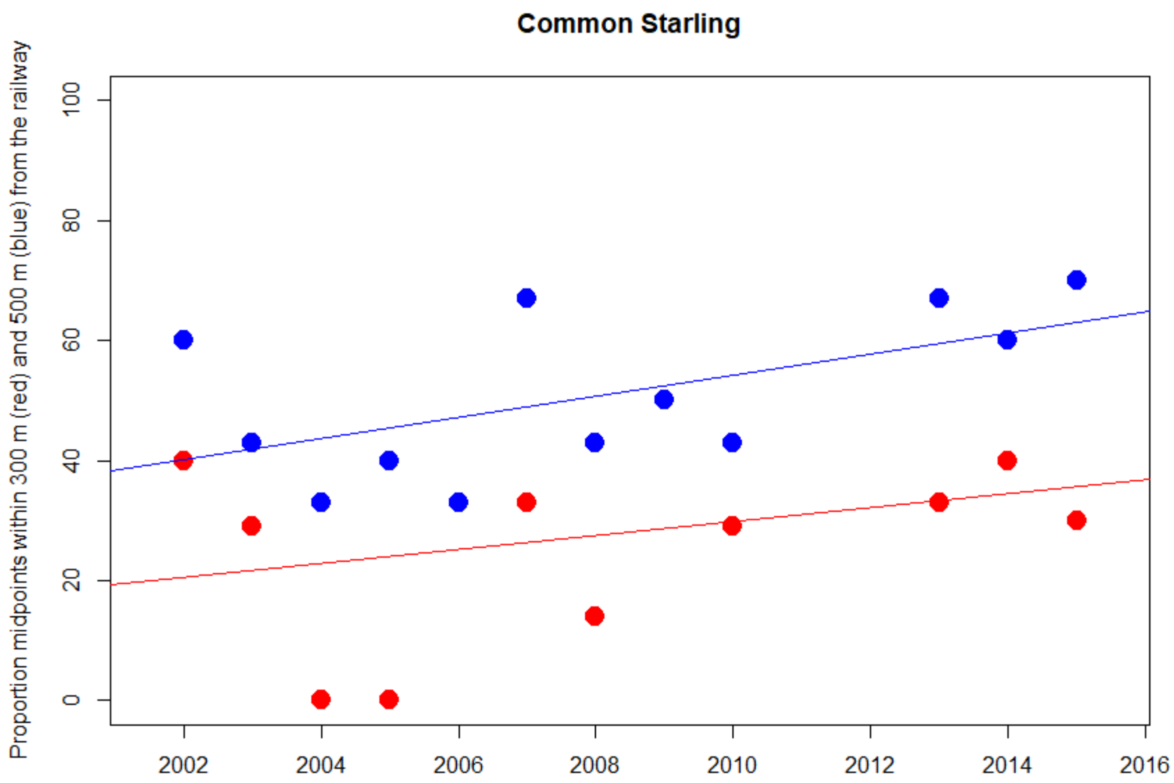


```
300 m. red
Kendall's rank correlation tau = 0.191
z = 0.836, p-value = 0.403

500 m. blue
Kendall's rank correlation tau = 0.413
z = 1.812, p-value = 0.070
```

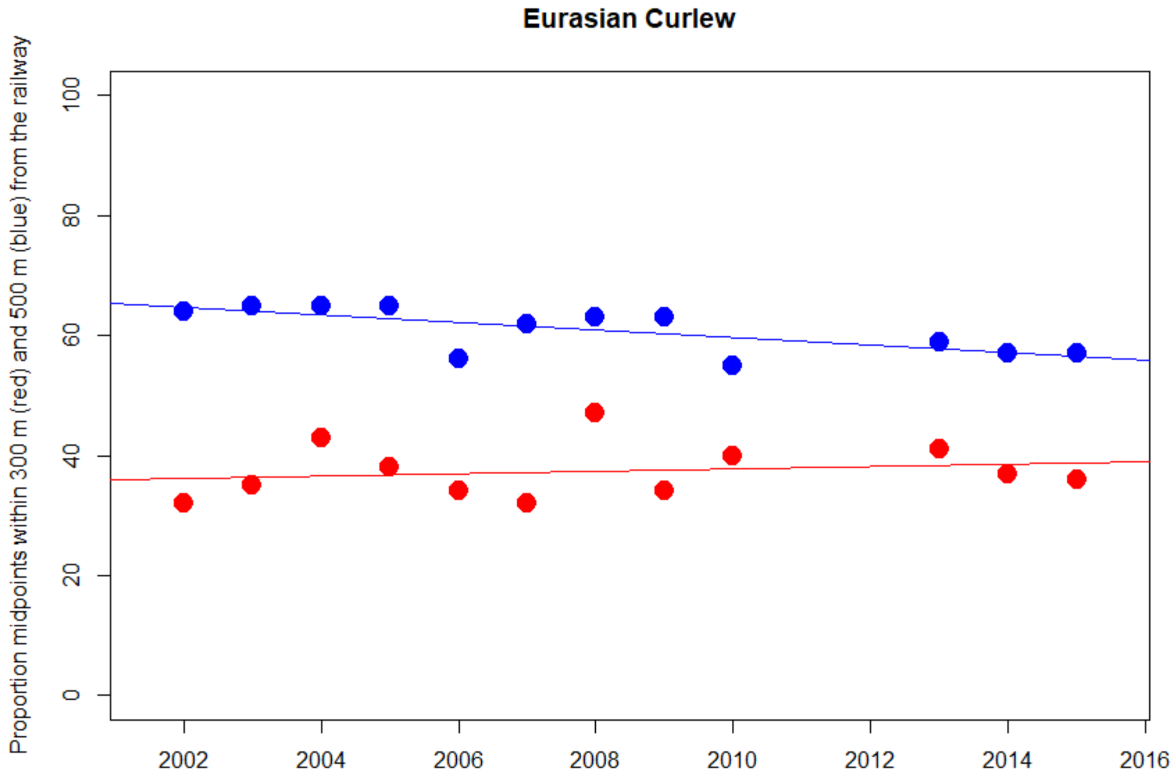


300 m. red
Kendall's rank correlation tau = 0.154
z = 0.689, p-value = 0.491

500 m. blue
Kendall's rank correlation tau = -0.520
z = -2.294, p-value = 0.022

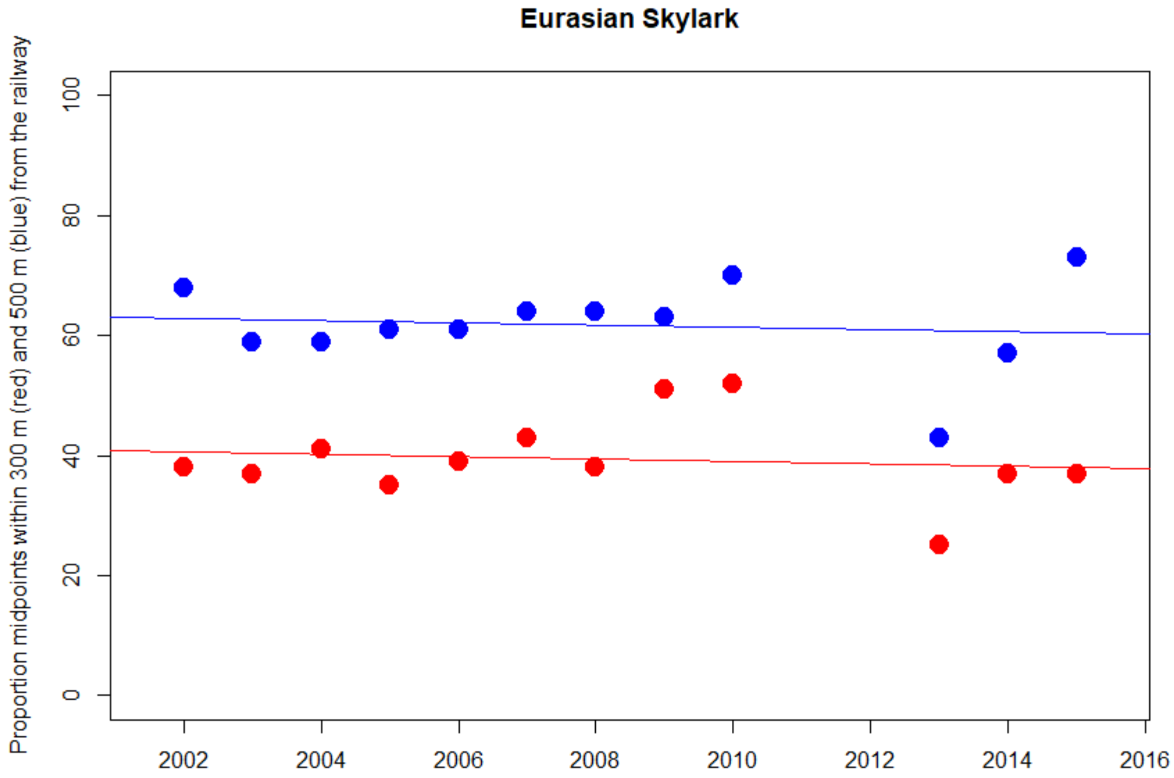


```
300 m. red
Kendall's rank correlation tau =   0.000
z = 0, p-value = 1

500 m. blue
Kendall's rank correlation tau = 0.140
z = 0.622, p-value = 0.534
```

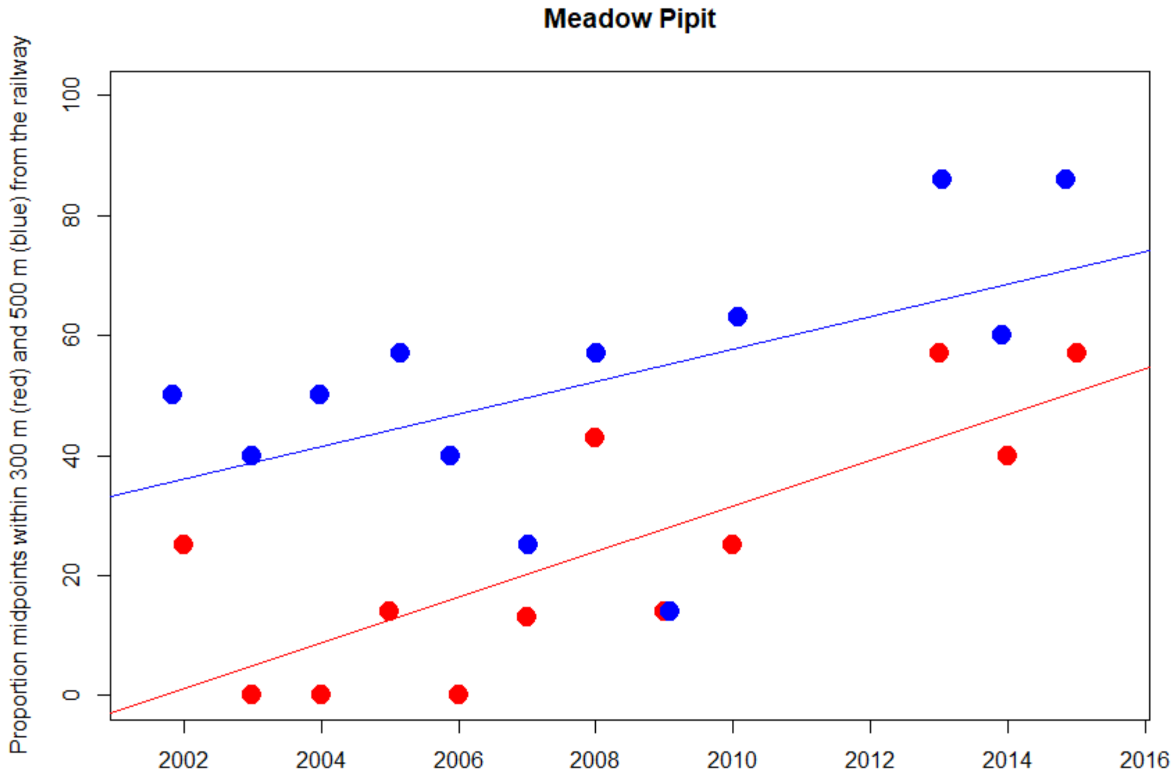


300 m. red
Kendall's rank correlation tau = 0.572
z = 2.508, p-value = 0.012

500 m. blue
Kendall's rank correlation tau = 0.4064485
z = 1.7999, p-value = 0.07188

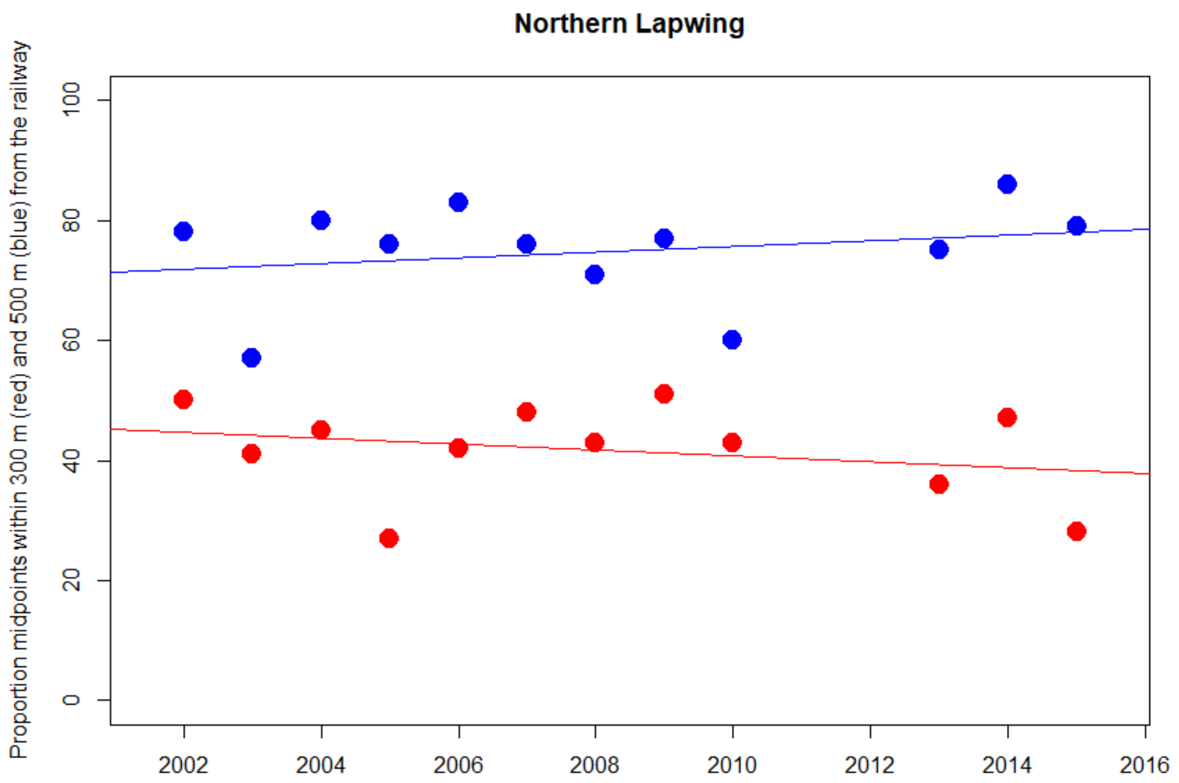


300 m. red
Kendall's rank correlation tau = -0.107
z = -0.481, p-value = 0.630

500 m. blue
Kendall's rank correlation tau = 0.076
z = 0.344, p-value = 0.731

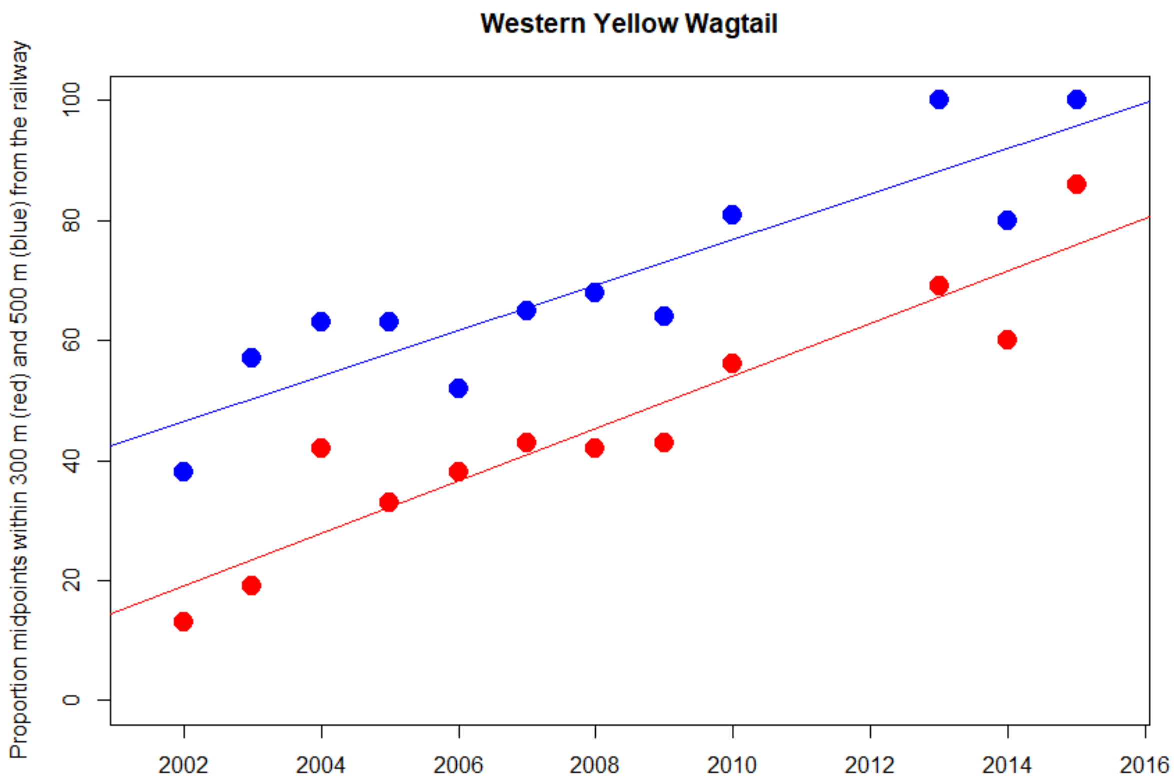


```
300 m. red
Kendall's rank correlation tau = 0.862
z = 3.858, p-value < 0.001

500 m. blue
Kendall's rank correlation tau = 0.769
z = 3.445, p-value < 0.001
```

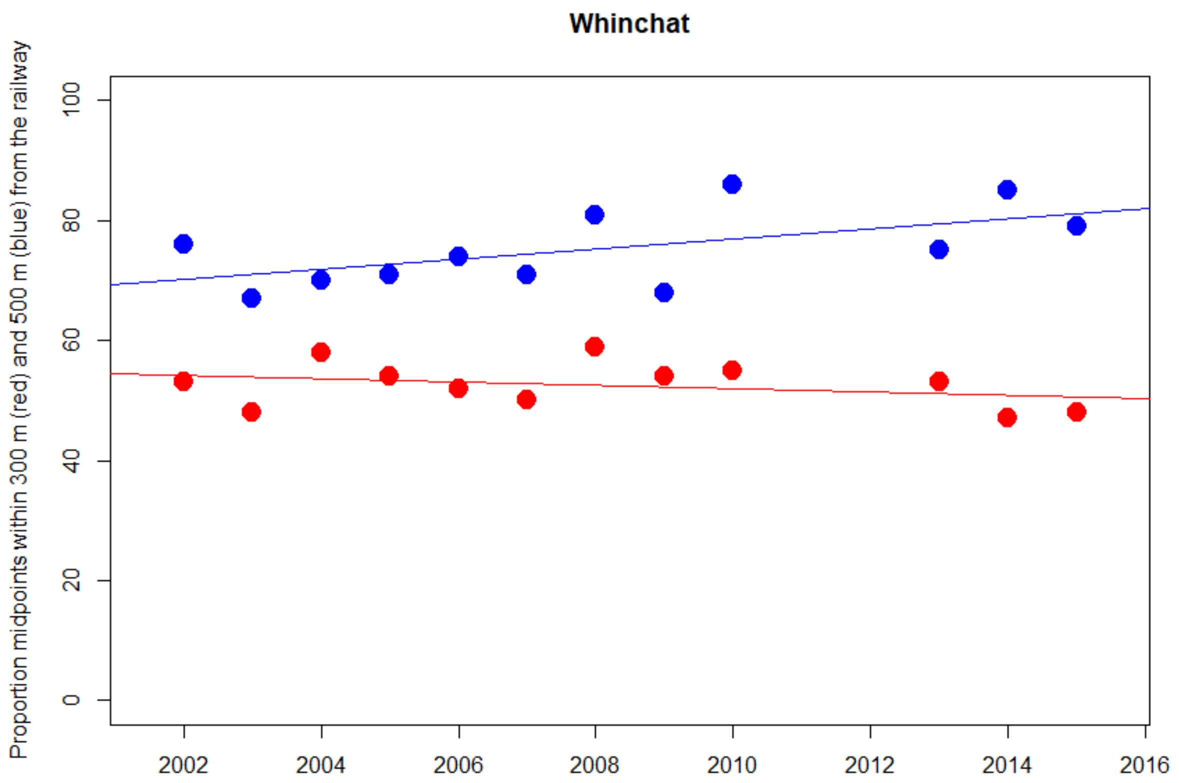


300 m. red
Kendall's rank correlation tau = -0.171
z = -0.760, p-value = 0.447

500 m. blue
Kendall's rank correlation tau = 0.412
z = 1.856, p-value = 0.063
