## Supplementary material for "Are railways detrimental to bird populations? A BDACI study on the construction of the Bothnia Line Railway": Trends of distances to BLR per site and species

Supplementary material **Dist4**

### Temporal trends for distances between territory midpoints and the Bothnia Line Railway per site and per species.

Plots and Kendall's rank correlation tests. Output from R-script presented *as is*.

Colour coding: *Before* = green, *Construction* = blue, *Ready* = purple and *Traffic* = red.

Many correlation tests generated warning messages due to ties (data points with the same Y-value). The warning texts were omitted from the output text for readability.

Significant (*P*<0.05) correlations are marked with yellow background.

NA denotes cases with less than four data points and thus, irrelevant correlation test.

Content:


| Site | Pages |
|---|---:|
| Nyland | 2 - 8 |
| Kornsjö | 9 - 17 |
| Stranne | 18 - 22 |
| Strandnyland | 23 - 33 |
| Hjälta | 34 - 45 |
| Kasa | 46 - 55 |
| Ava | 56 - 67 |
| Lögdeå | 68 - 74 |
| Långed | 75 - 81 |
| Hörneå | 82 - 88 |
| Stöcke | 89 - 98 |
| Stöcke NE | 99 - 100 |
| Degernäs | 101 - 110 |

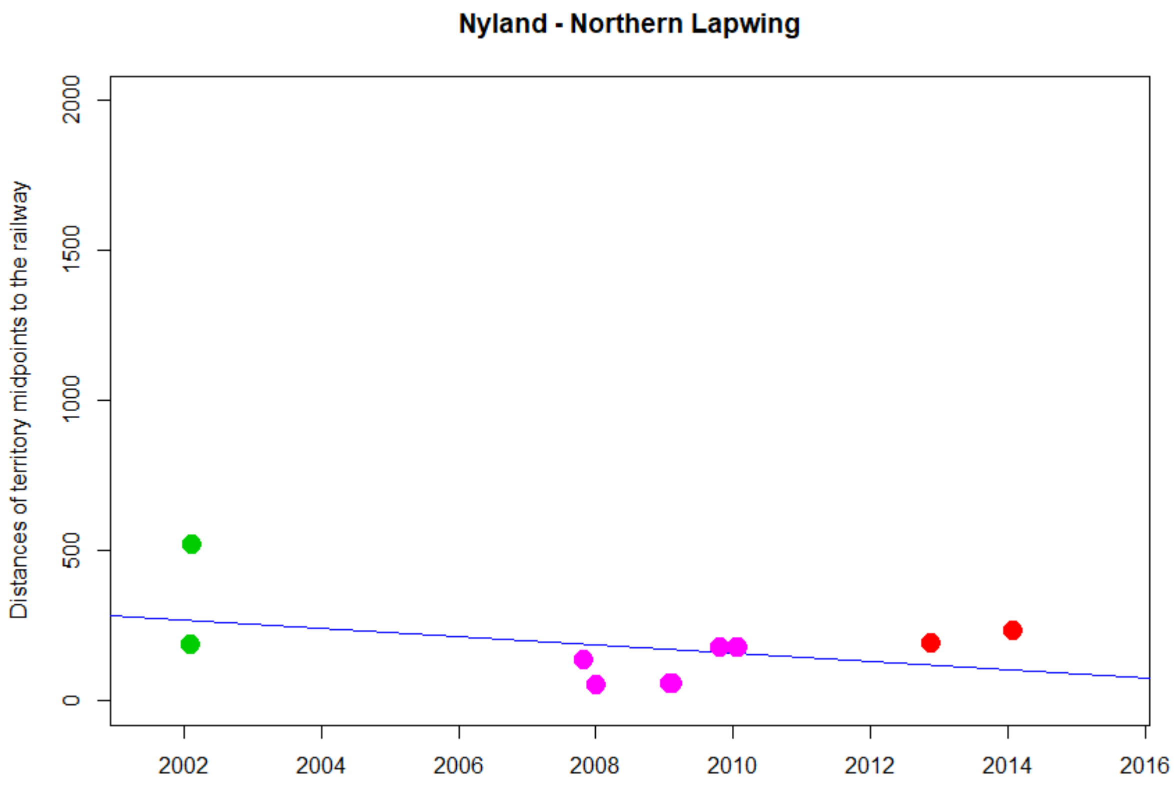


```
> cor.test(DNyland$Year,DNyland$NEAR_DIST_Botnia, method="kendall")

        Kendall's rank correlation tau

data:  DNyland$Year and DNyland$NEAR_DIST_Botnia
z = 1.0472, p-value = 0.295
alternative hypothesis: true tau is not equal to 0
sample estimates:
     tau
0.260052
```

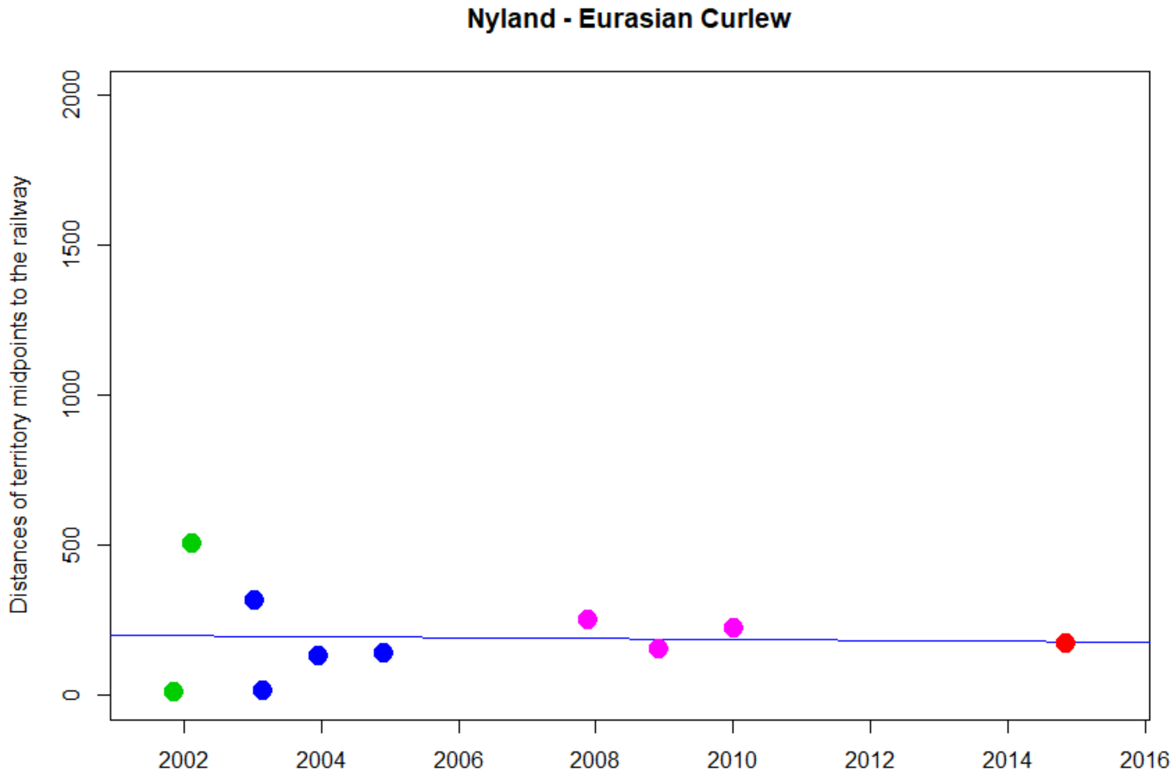


```
> cor.test(DNyland$Year,DNyland$NEAR_DIST_Botnia, method="kendall")

        Kendall's rank correlation tau

data:  DNyland$Year and DNyland$NEAR_DIST_Botnia
z = 0.63117, p-value = 0.5279
alternative hypothesis: true tau is not equal to 0
sample estimates:
     tau 
0.159132 
```

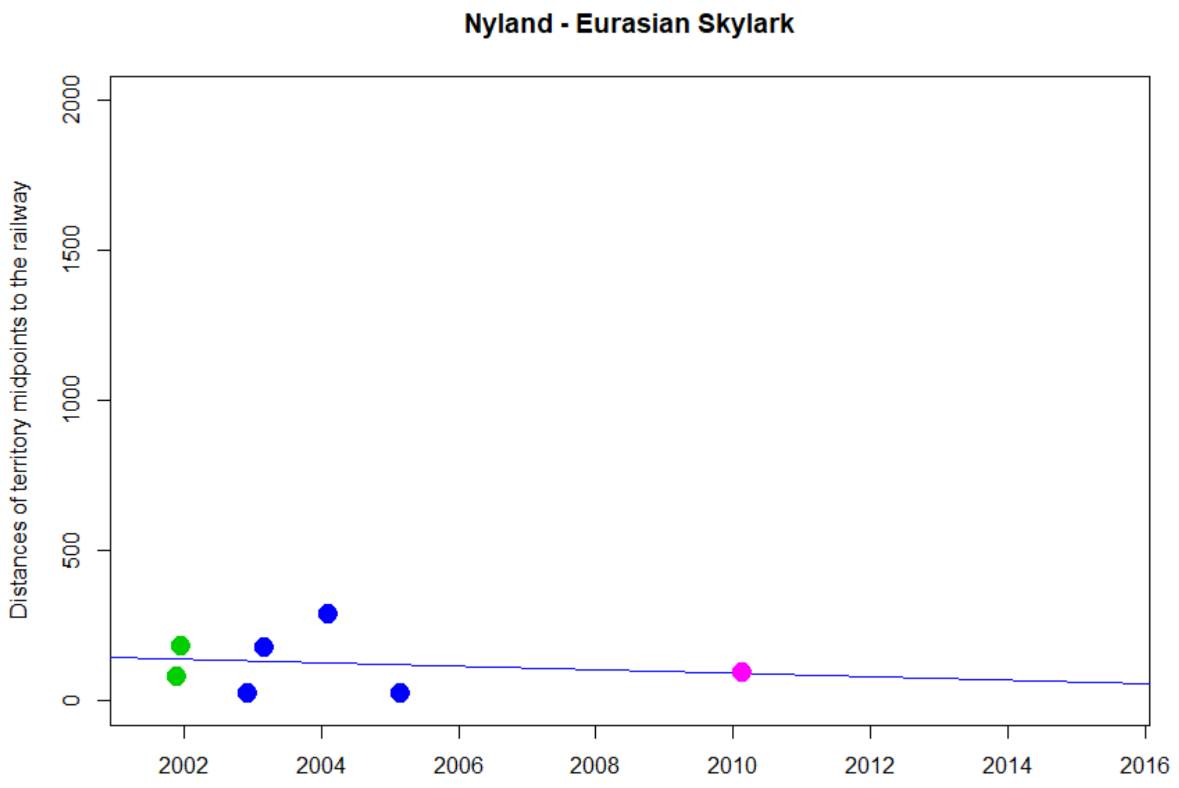


```
> cor.test(DNyland$Year,DNyland$NEAR_DIST_Botnia, method="kendall")

        Kendall's rank correlation tau

data:  DNyland$Year and DNyland$NEAR_DIST_Botnia
z = -0.15369, p-value = 0.8779
alternative hypothesis: true tau is not equal to 0
sample estimates:
        tau
-0.05006262
```

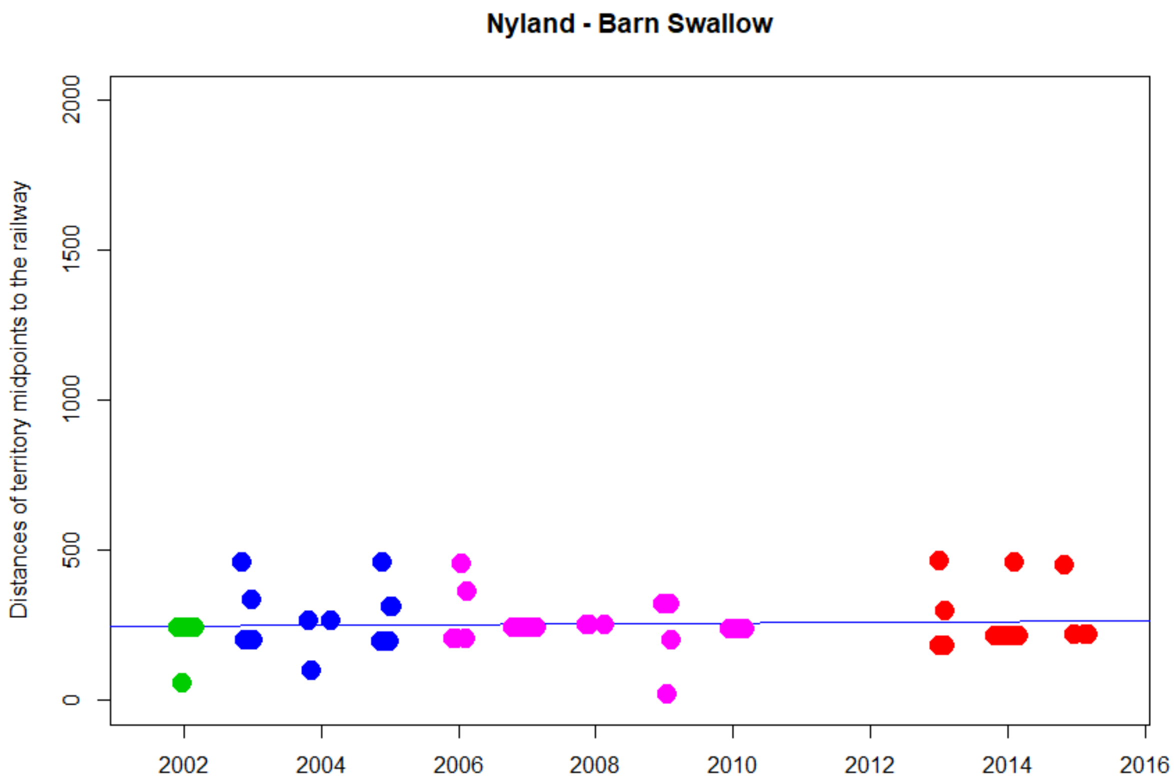


```
> cor.test(DNyland$Year,DNyland$NEAR_DIST_Botnia, method="kendall")

        Kendall's rank correlation tau

data:  DNyland$Year and DNyland$NEAR_DIST_Botnia
z = -0.49427, p-value = 0.6211
alternative hypothesis: true tau is not equal to 0
sample estimates:
        tau
-0.04906824
```

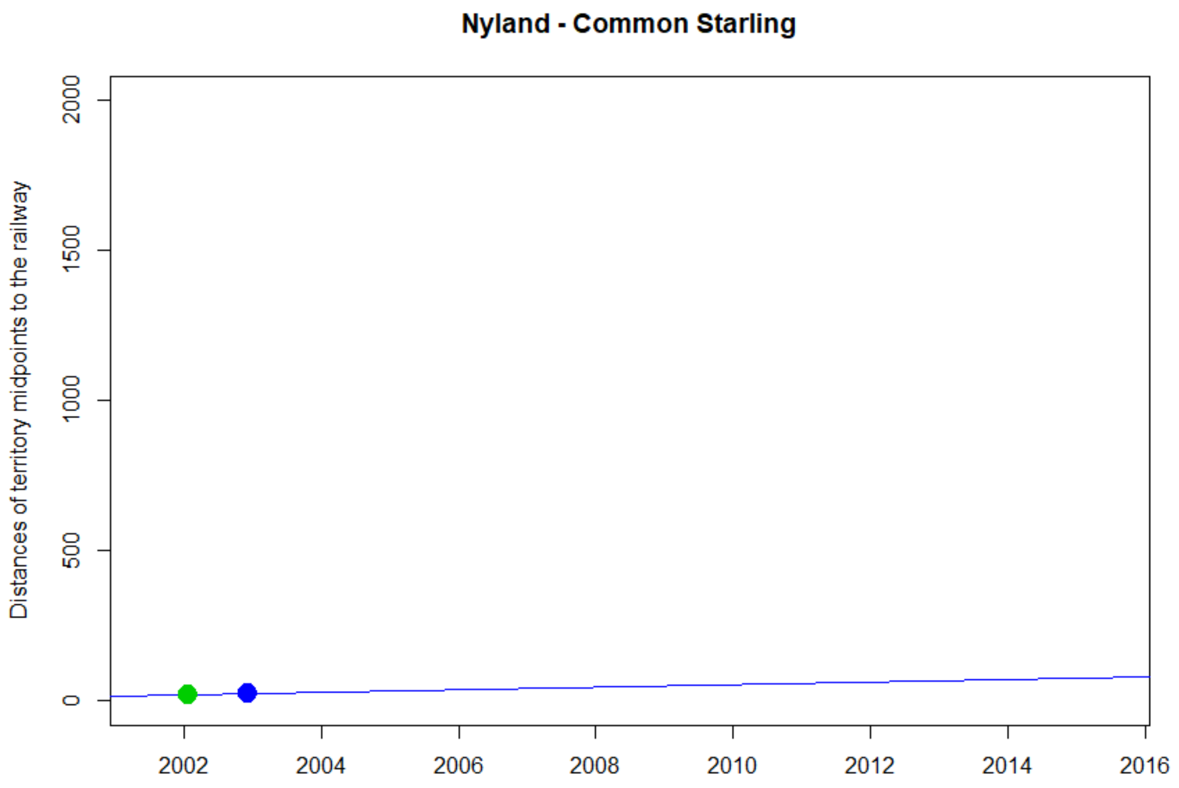

Nyland - Common Starling
Distances of territory midpoints to the railway
0
500
1000
1500
2000
2002
2004
2006
2008
2010
2012
2014
2016


NA

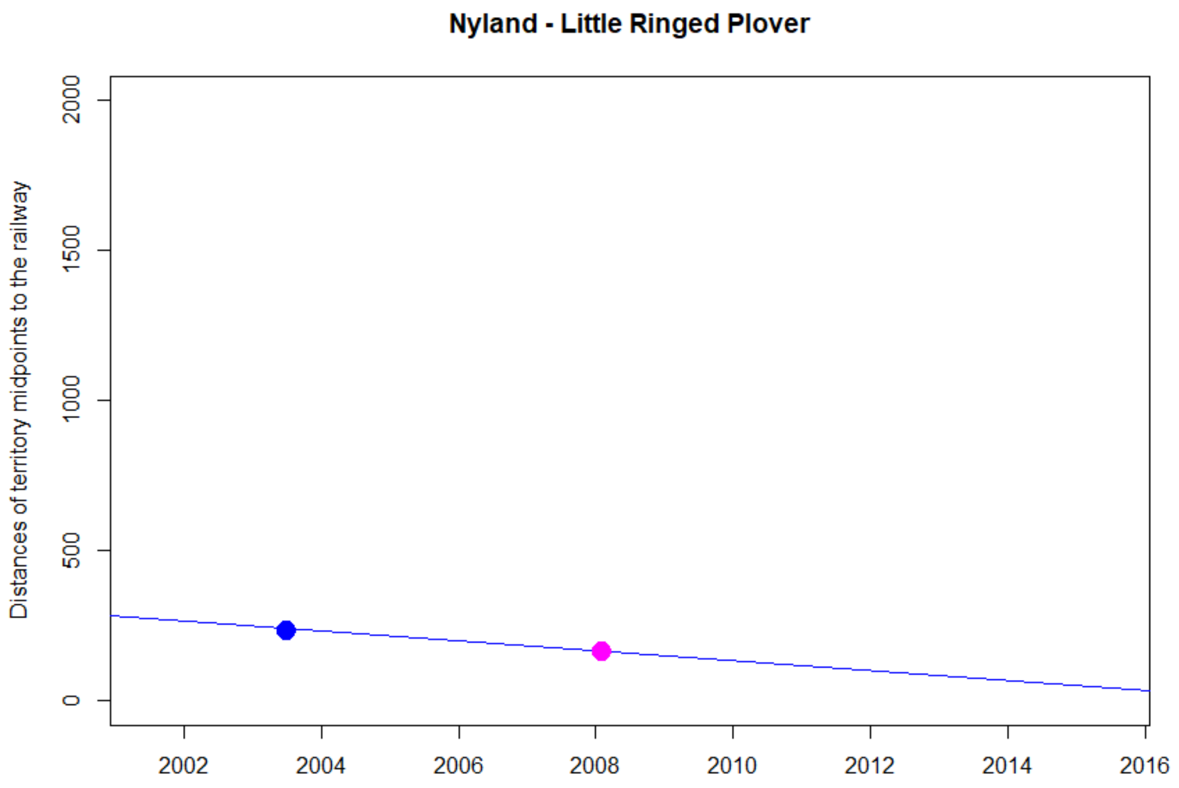
Nyland - Little Ringed Plover
Distances of territory midpoints to the railway
0
500
1000
1500
2000
2002
2004
2006
2008
2010
2012
2014
2016

NA

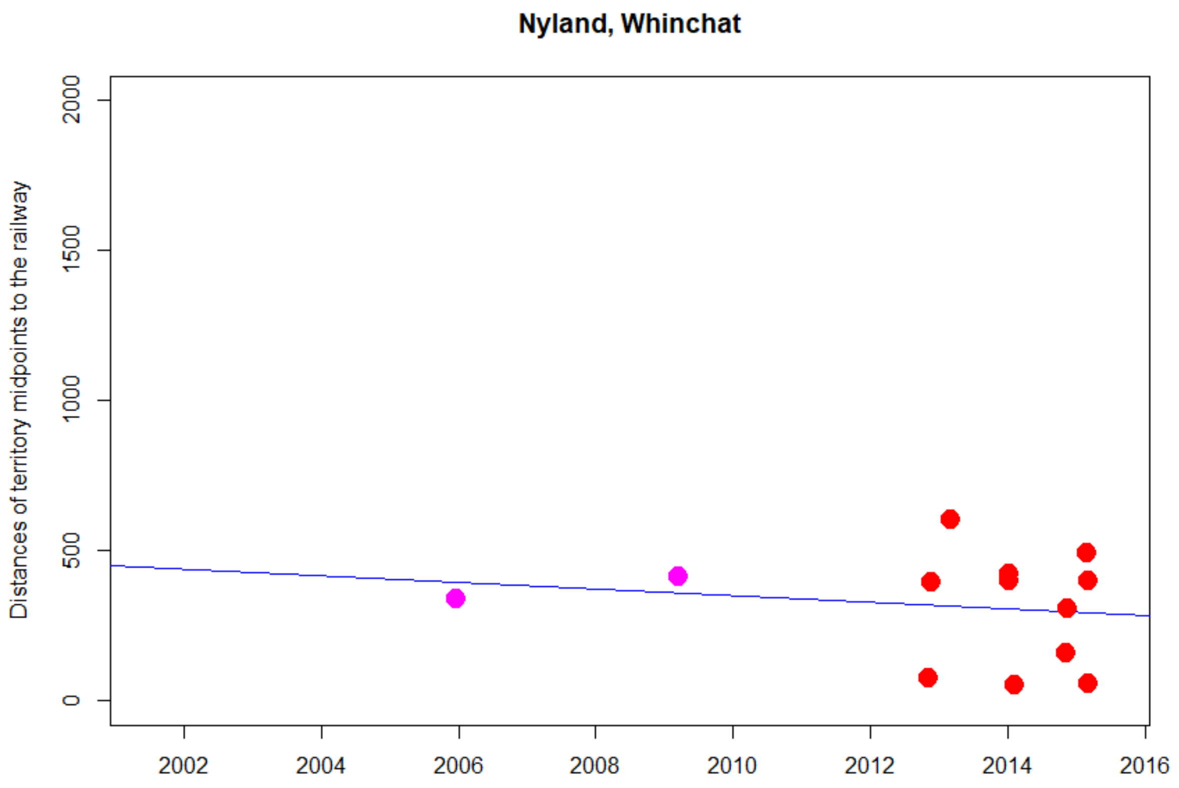


```
> cor.test(DNyland$Year,DNyland$NEAR_DIST_Botnia, method="kendall")

        Kendall's rank correlation tau

data:  DNyland$Year and DNyland$NEAR_DIST_Botnia
z = -0.51145, p-value = 0.609
alternative hypothesis: true tau is not equal to 0
sample estimates:
       tau 
-0.1150395 
```

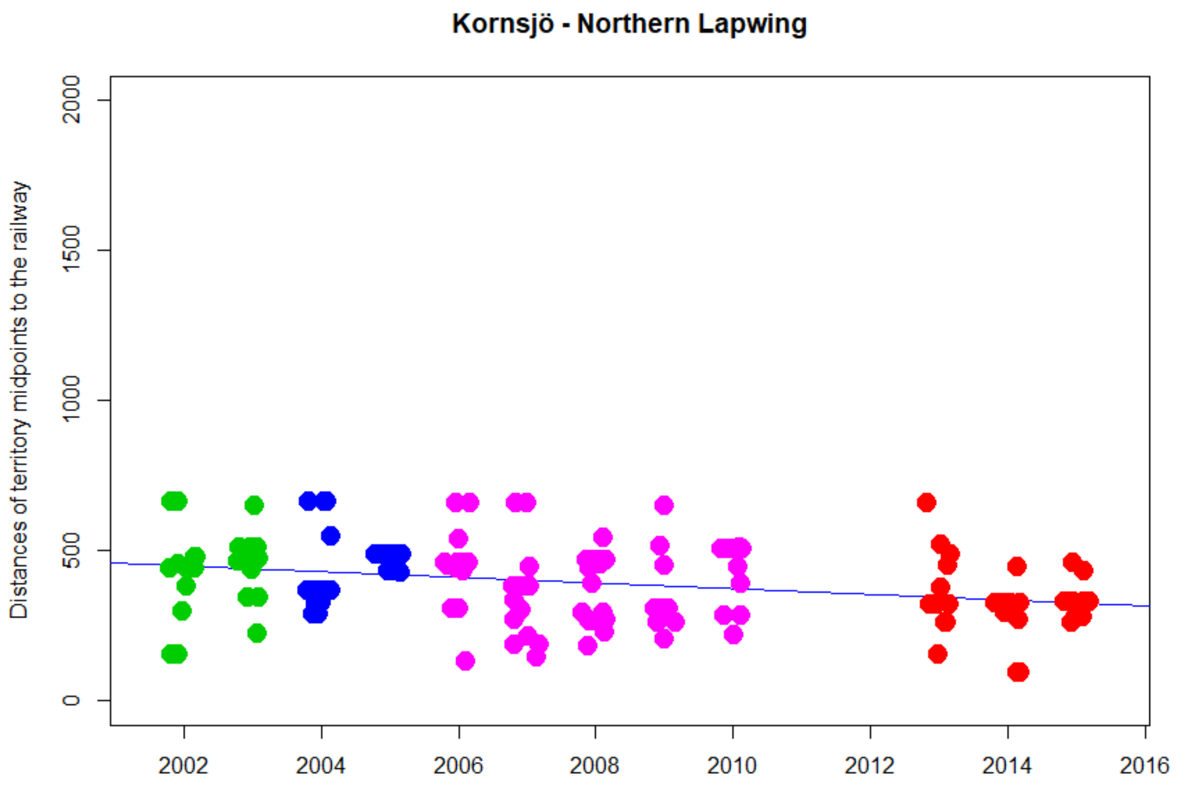


```
> cor.test(DNyland$Year,DNyland$NEAR_DIST_Botnia, method="kendall")

        Kendall's rank correlation tau

data:  DNyland$Year and DNyland$NEAR_DIST_Botnia
z = -3.443, p-value = 0.0005754
alternative hypothesis: true tau is not equal to 0
sample estimates:
       tau
-0.1890802
```

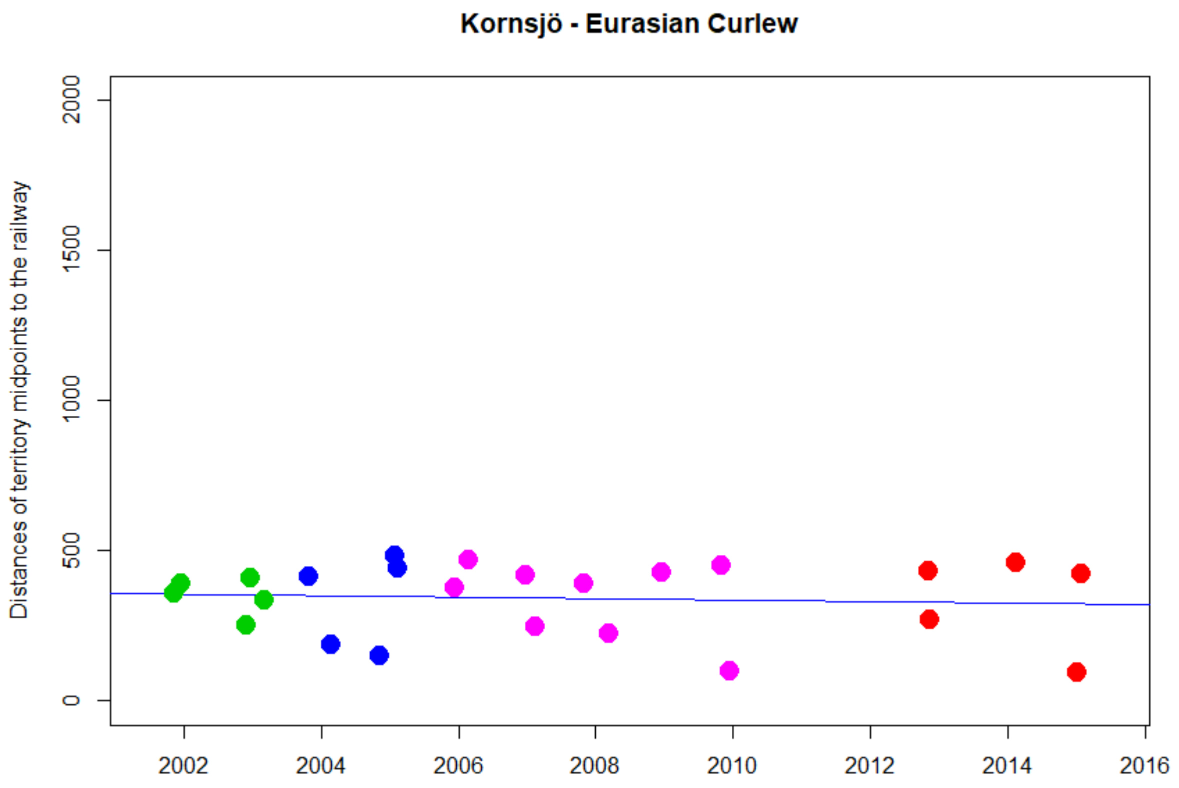


```
> cor.test(DNyland$Year,DNyland$NEAR_DIST_Botnia, method="kendall")

        Kendall's rank correlation tau

data:  DNyland$Year and DNyland$NEAR_DIST_Botnia
z = 0.34891, p-value = 0.7272
alternative hypothesis: true tau is not equal to 0
sample estimates:
       tau
0.05206224
```

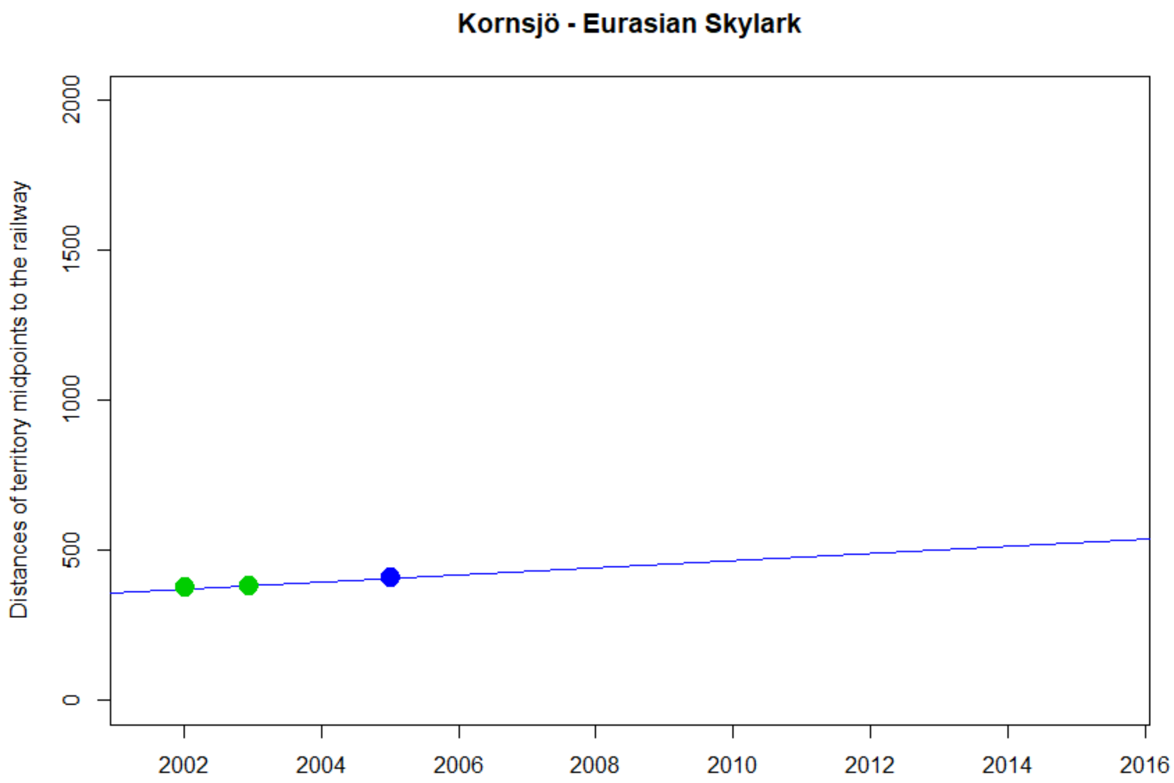

Kornsjö - Eurasian Skylark
Distances of territory midpoints to the railway
0
500
1000
1500
2000
2002
2004
2006
2008
2010
2012
2014
2016


NA

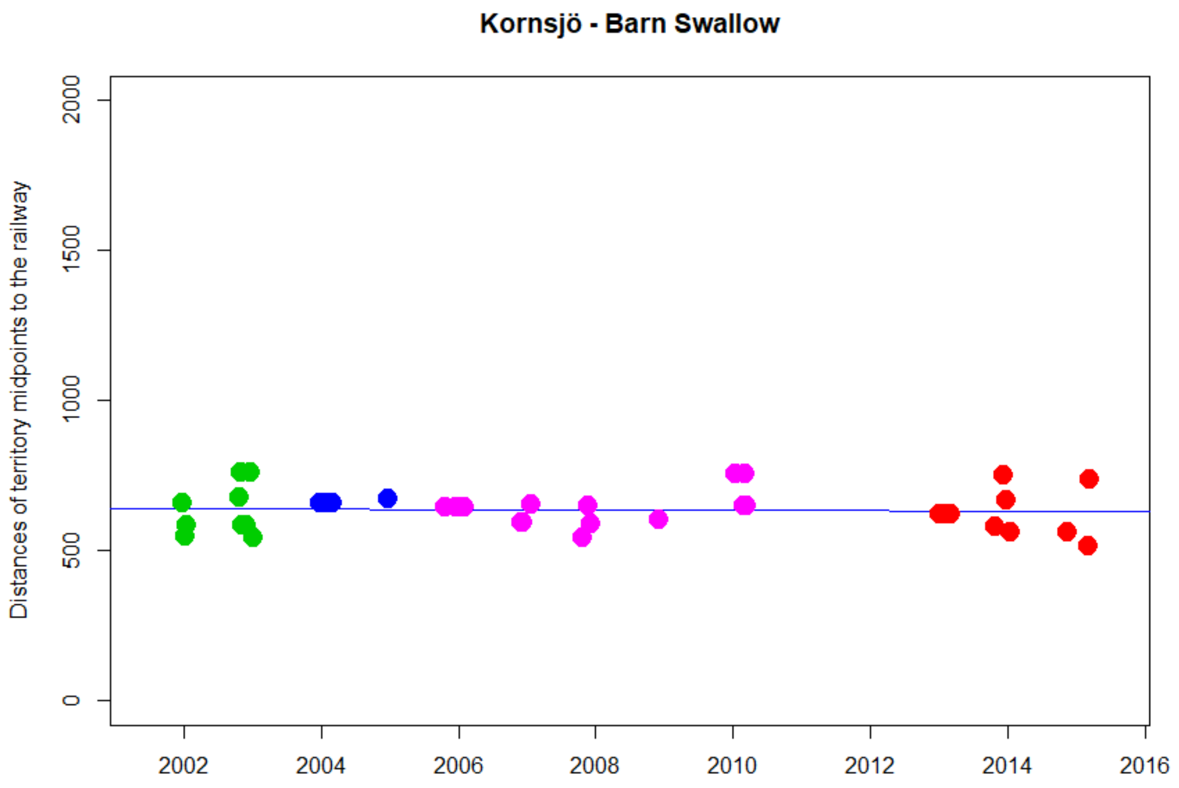


```
> cor.test(DNyland$Year,DNyland$NEAR_DIST_Botnia, method="kendall")

        Kendall's rank correlation tau

data:  DNyland$Year and DNyland$NEAR_DIST_Botnia
z = -0.76011, p-value = 0.4472
alternative hypothesis: true tau is not equal to 0
sample estimates:
        tau
-0.08971507
```

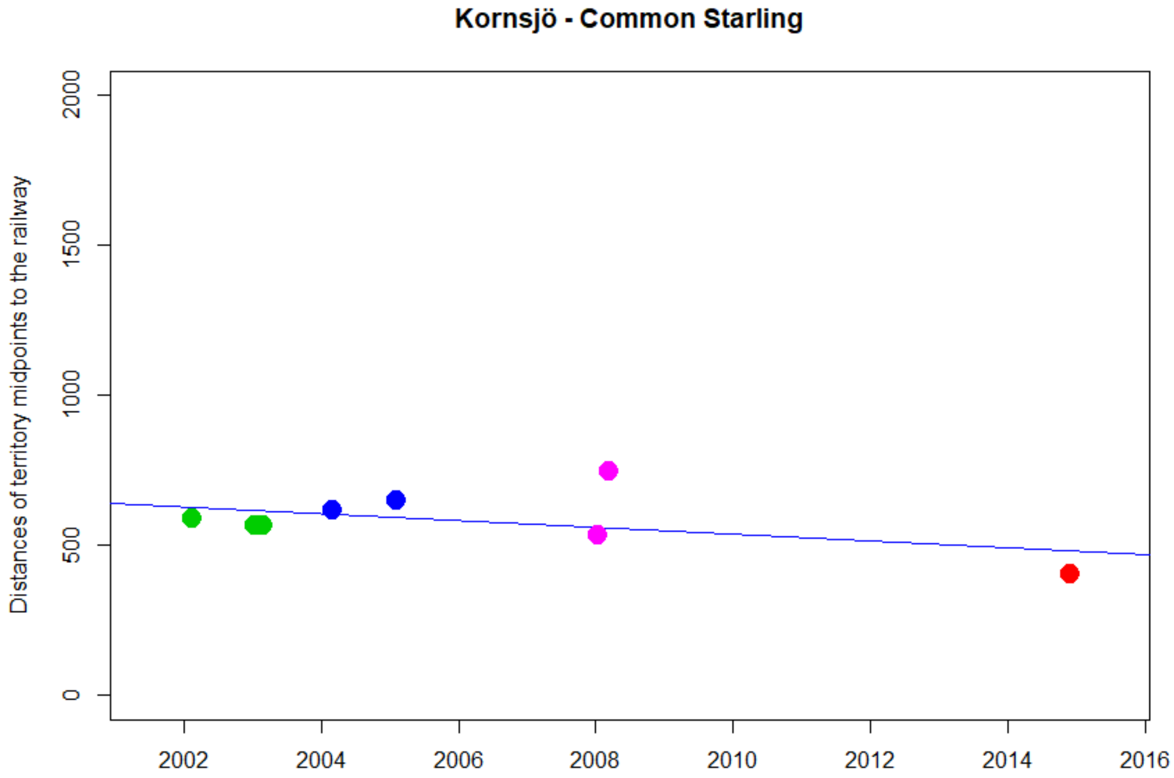


```
> cor.test(DNyland$Year,DNyland$NEAR_DIST_Botnia, method="kendall")

        Kendall's rank correlation tau

data:  DNyland$Year and DNyland$NEAR_DIST_Botnia
z = -0.25318, p-value = 0.8001
alternative hypothesis: true tau is not equal to 0
sample estimates:
        tau
-0.07548514
```

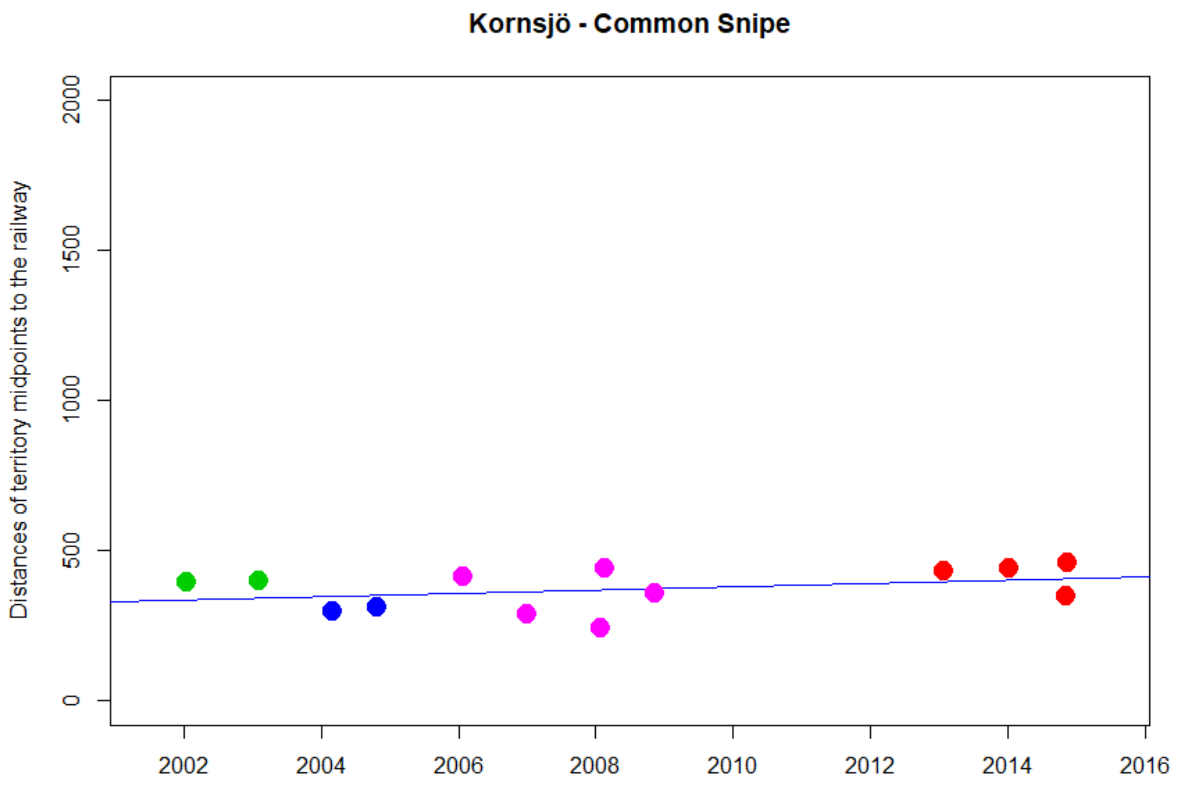


```
> cor.test(DNyland$Year,DNyland$NEAR_DIST_Botnia, method="kendall")

        Kendall's rank correlation tau

data:  DNyland$Year and DNyland$NEAR_DIST_Botnia
z = 1.3472, p-value = 0.1779
alternative hypothesis: true tau is not equal to 0
sample estimates:
      tau
0.2857384
```

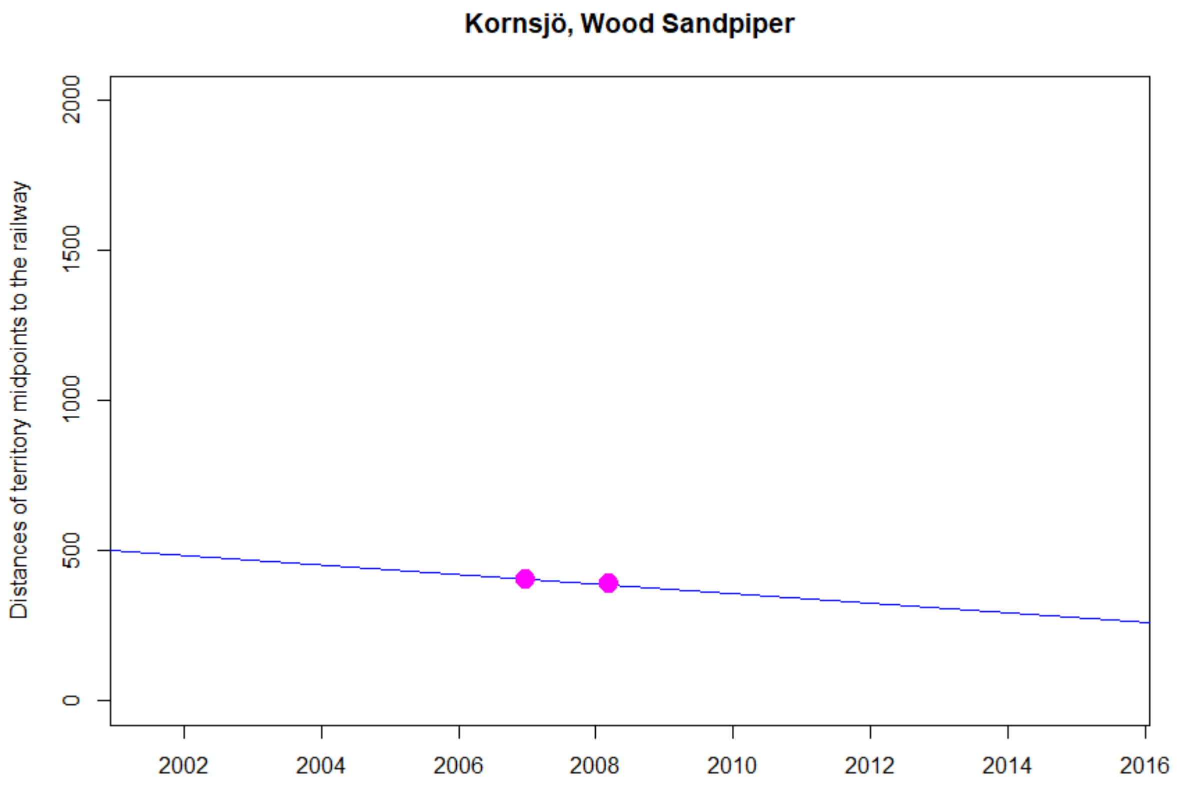
Kornsjö, Wood Sandpiper
Distances of territory midpoints to the railway
0
500
1000
1500
2000
2002
2004
2006
2008
2010
2012
2014
2016

NA

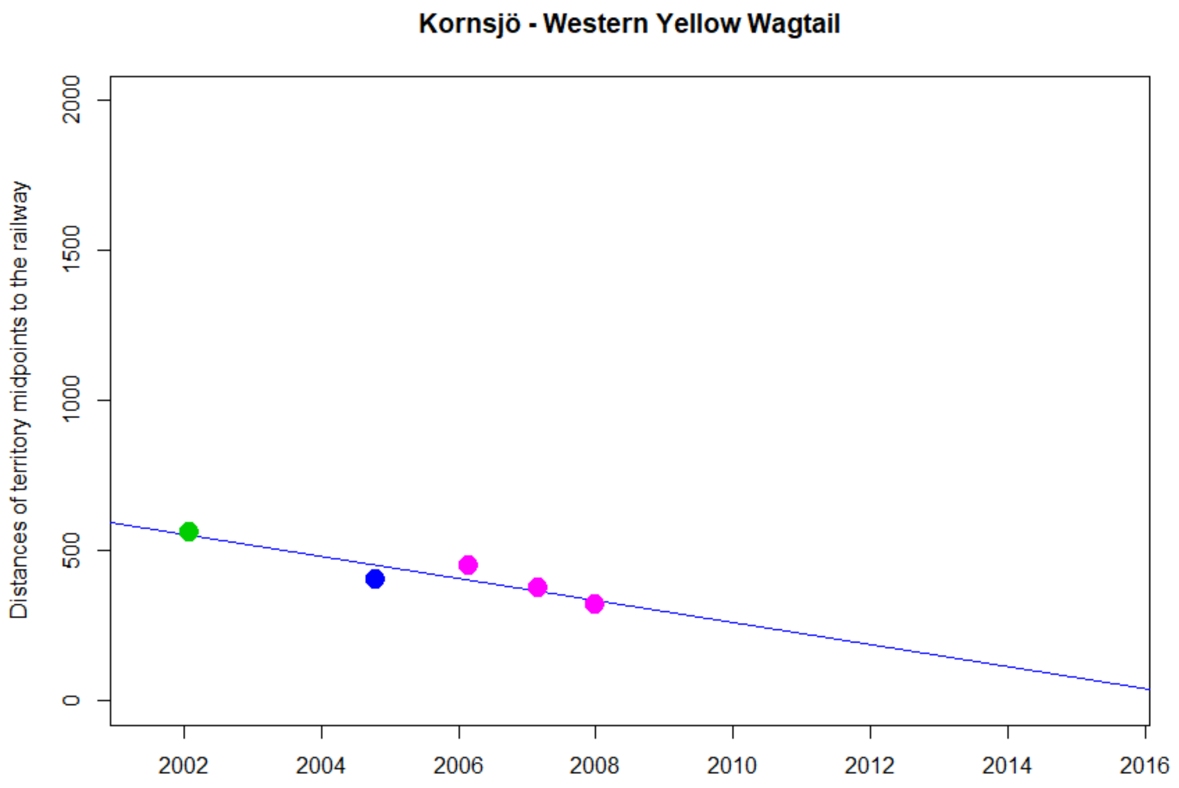


```
> cor.test(DNyland$Year,DNyland$NEAR_DIST_Botnia, method="kendall")

        Kendall's rank correlation tau

data:  DNyland$Year and DNyland$NEAR_DIST_Botnia
T = 1, p-value = 0.08333
alternative hypothesis: true tau is not equal to 0
sample estimates:
 tau 
-0.8 
```

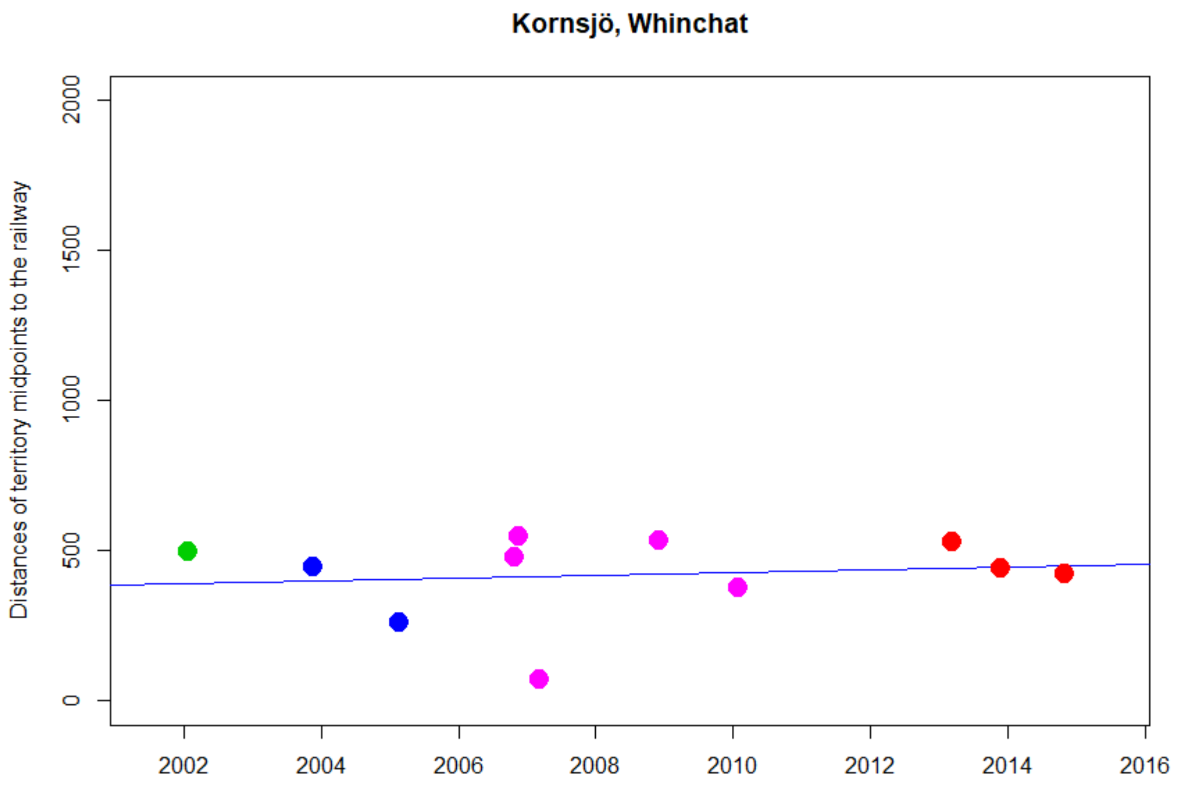


```
> cor.test(DNyland$Year,DNyland$NEAR_DIST_Botnia, method="kendall")

        Kendall's rank correlation tau

data:  DNyland$Year and DNyland$NEAR_DIST_Botnia
z = -0.31492, p-value = 0.7528
alternative hypothesis: true tau is not equal to 0
sample estimates:
        tau
-0.07479576
```

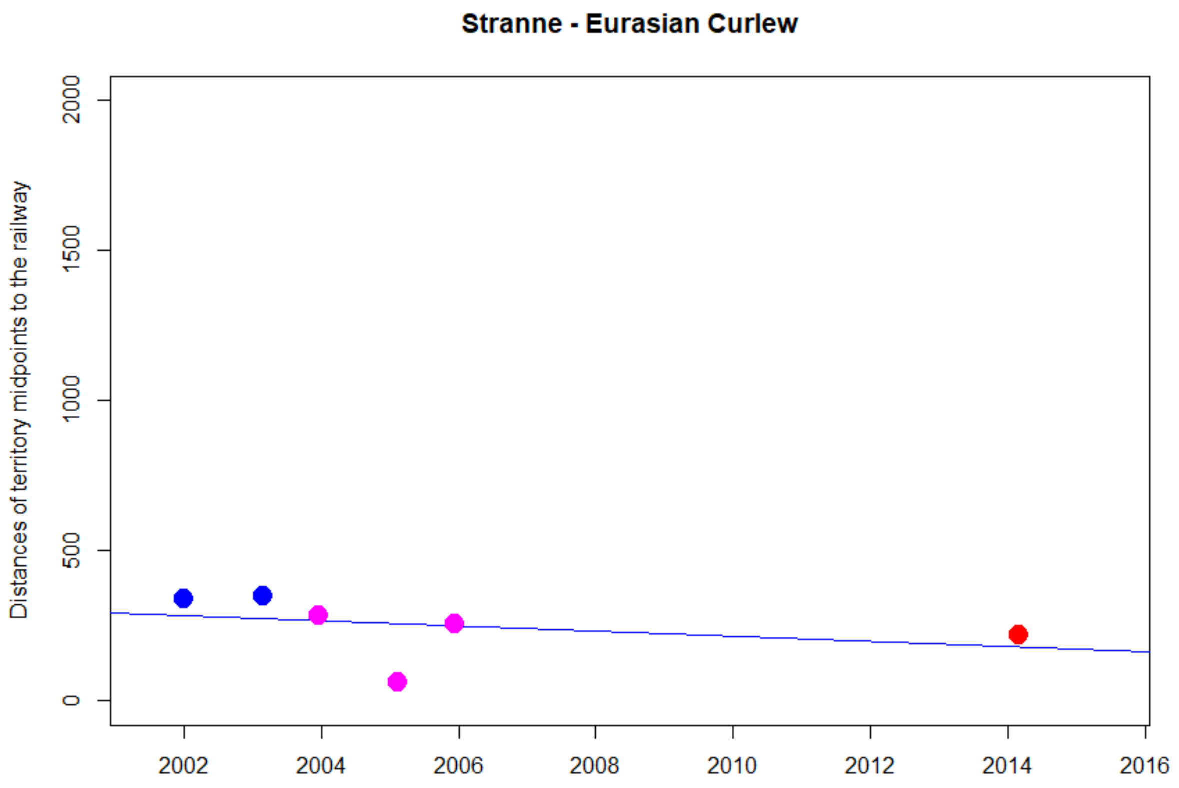


```
> cor.test(DNyland$Year,DNyland$NEAR_DIST_Botnia, method="kendall")

        Kendall's rank correlation tau

data:  DNyland$Year and DNyland$NEAR_DIST_Botnia
T = 3, p-value = 0.1361
alternative hypothesis: true tau is not equal to 0
sample estimates:
 tau
-0.6
```

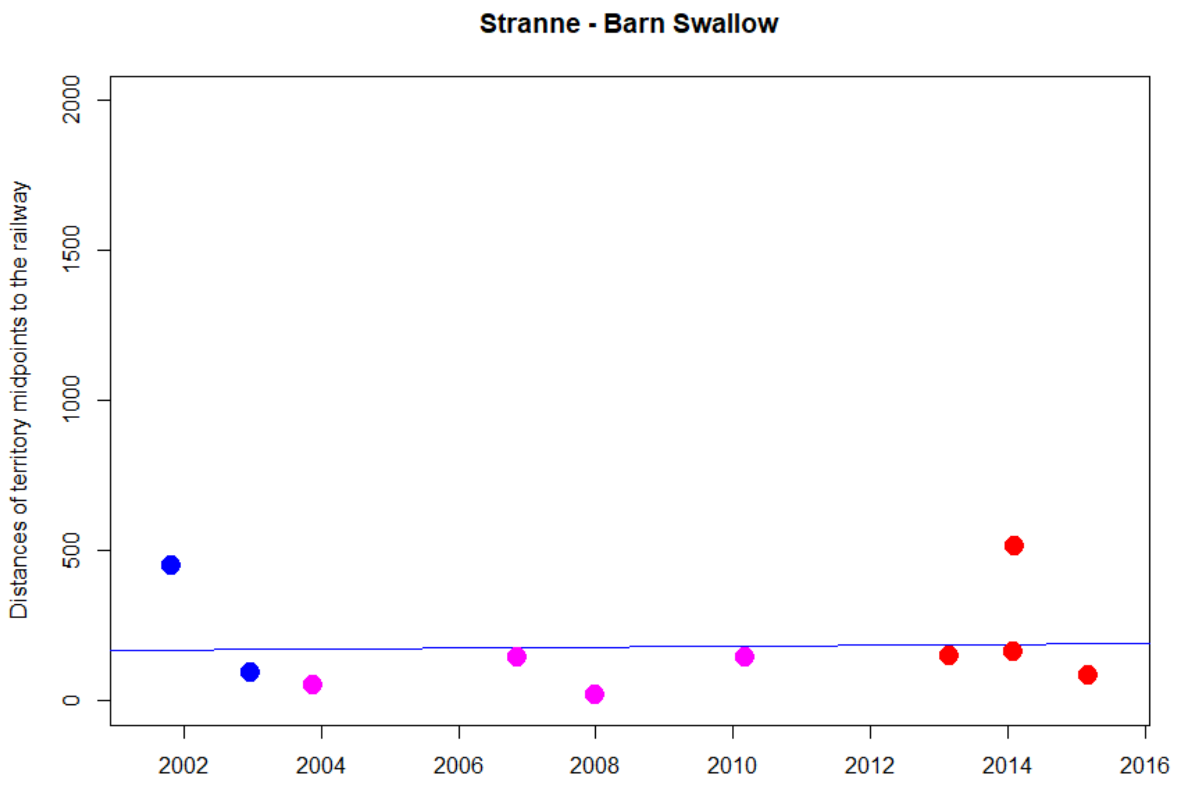


```
> cor.test(DNyland$Year,DNyland$NEAR_DIST_Botnia, method="kendall")

        Kendall's rank correlation tau

data:  DNyland$Year and DNyland$NEAR_DIST_Botnia
z = 0.71842, p-value = 0.4725
alternative hypothesis: true tau is not equal to 0
sample estimates:
      tau
0.1797866
```

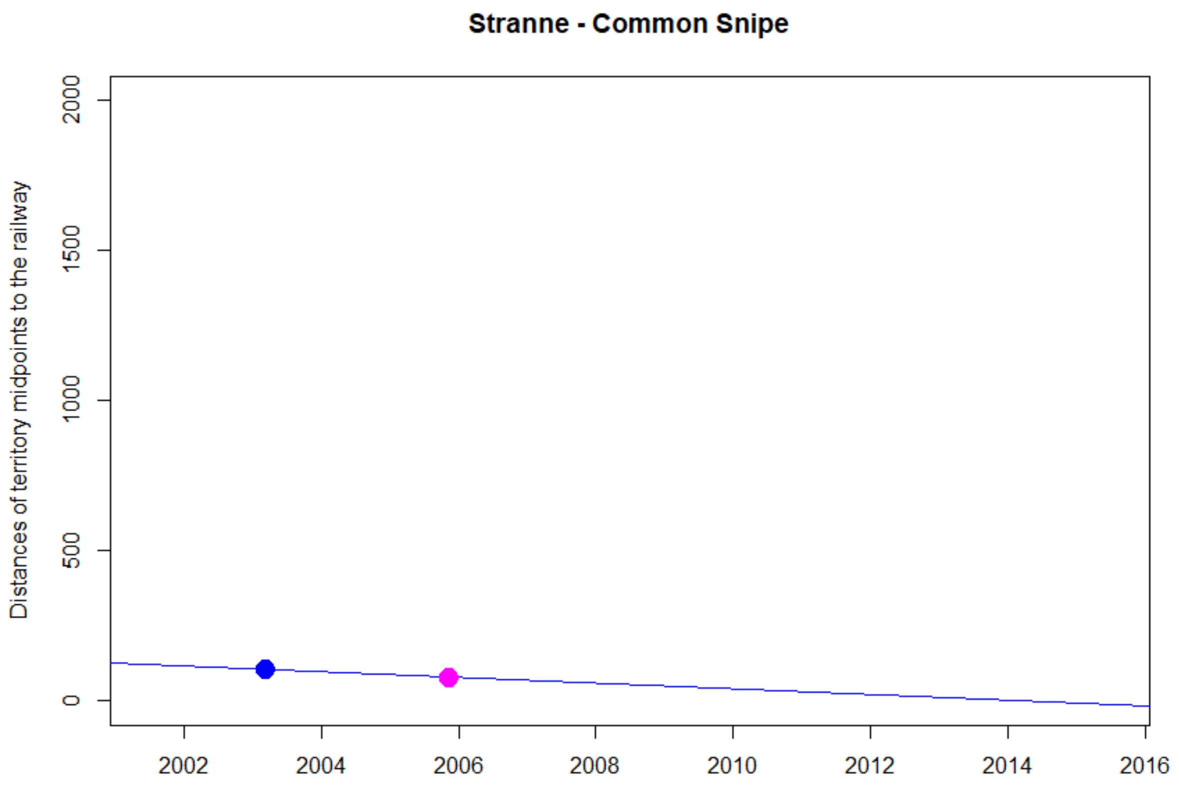
Stranne - Common Snipe
Distances of territory midpoints to the railway
0
500
1000
1500
2000
2002
2004
2006
2008
2010
2012
2014
2016

NA

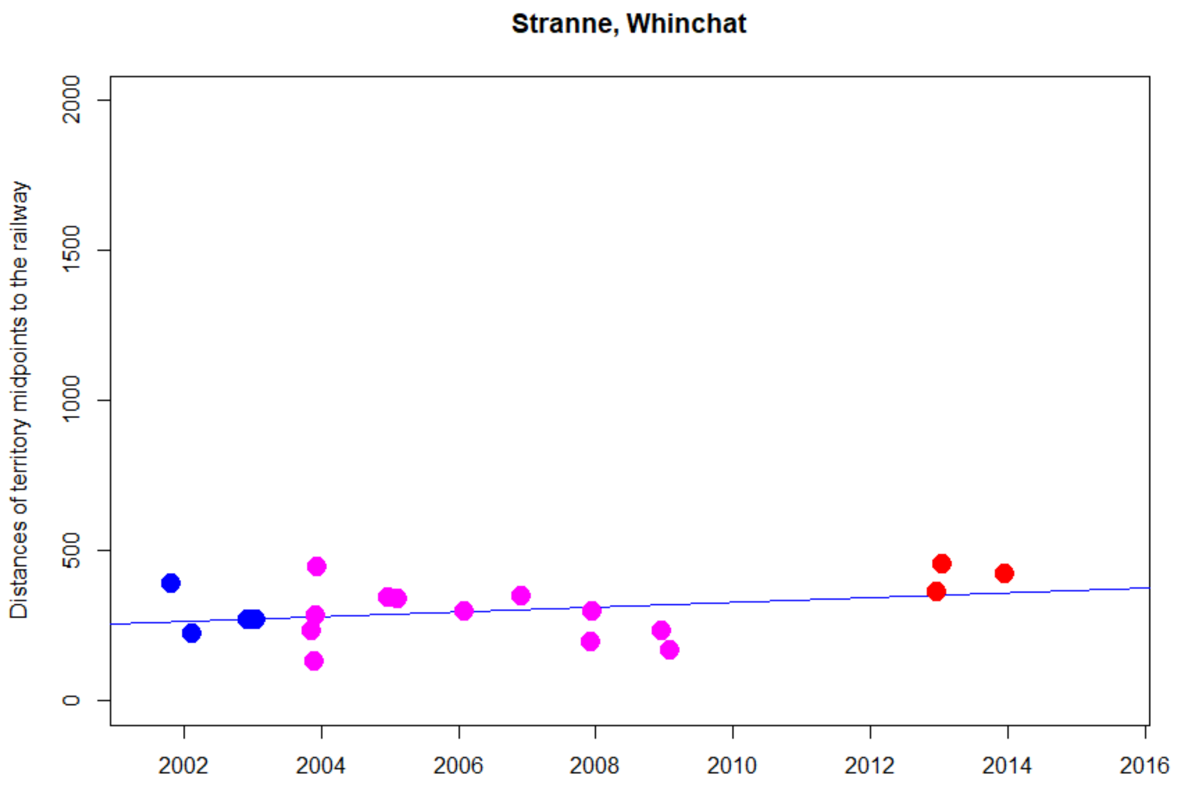


```
> cor.test(DNyland$Year,DNyland$NEAR_DIST_Botnia, method="kendall")

        Kendall's rank correlation tau

data:  DNyland$Year and DNyland$NEAR_DIST_Botnia
z = 1.0944, p-value = 0.2738
alternative hypothesis: true tau is not equal to 0
sample estimates:
      tau
0.1880031
```

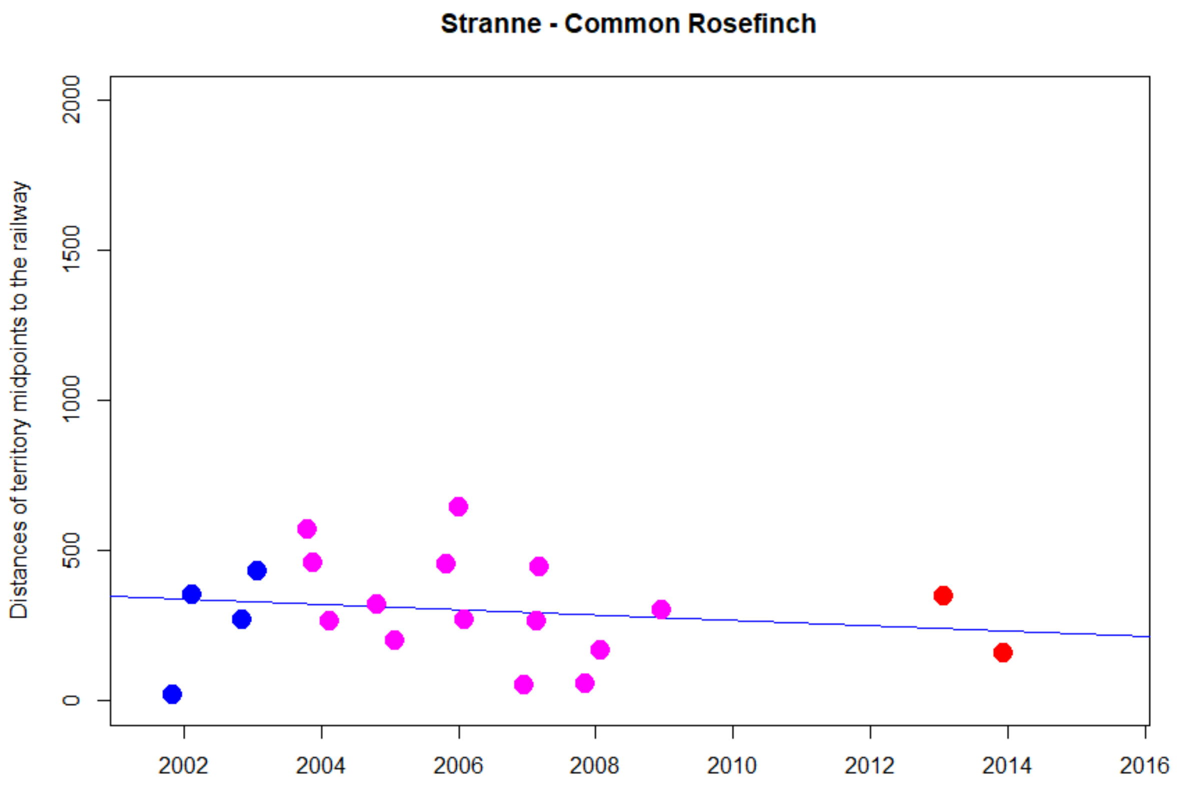


```
> cor.test(DNyland$Year,DNyland$NEAR_DIST_Botnia, method="kendall")

        Kendall's rank correlation tau

data:  DNyland$Year and DNyland$NEAR_DIST_Botnia
z = -0.9484, p-value = 0.3429
alternative hypothesis: true tau is not equal to 0
sample estimates:
       tau
-0.1581374
```

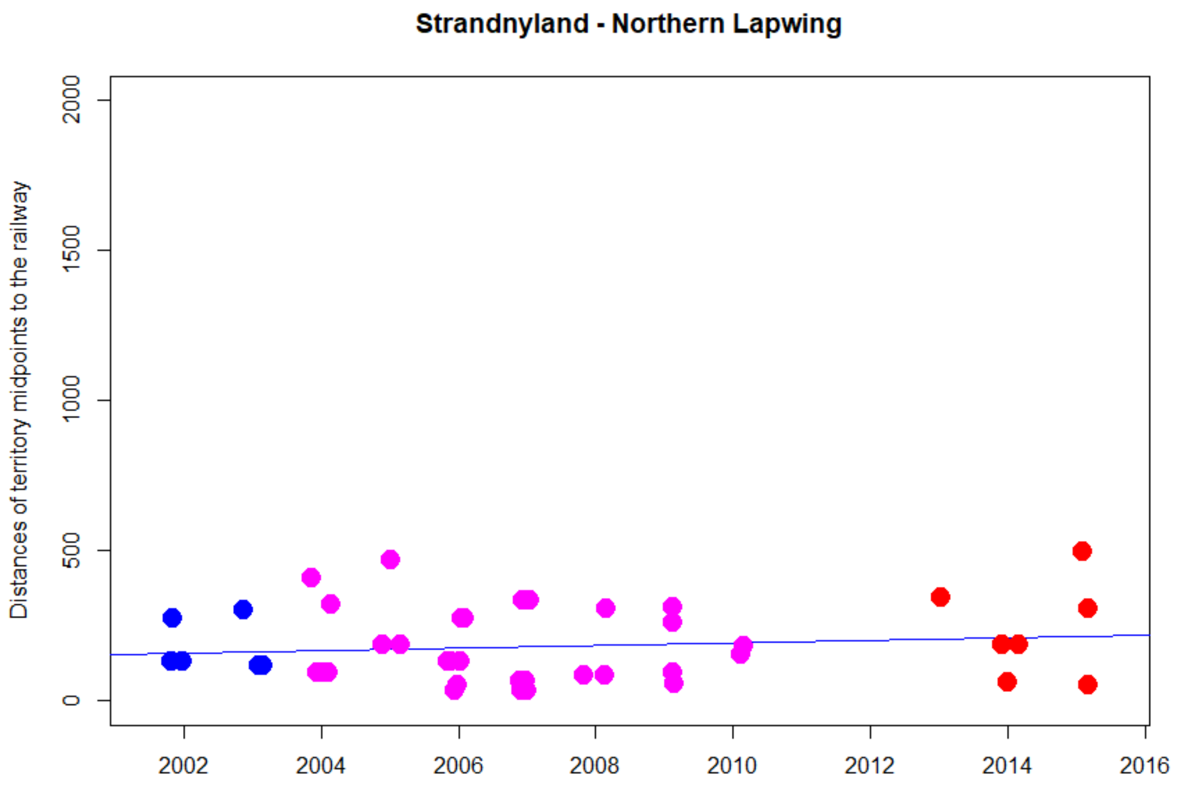


```
> cor.test(DNyland$Year,DNyland$NEAR_DIST_Botnia, method="kendall")

        Kendall's rank correlation tau

data:  DNyland$Year and DNyland$NEAR_DIST_Botnia
z = 0.095471, p-value = 0.9239
alternative hypothesis: true tau is not equal to 0
sample estimates:
       tau
0.01020382
```

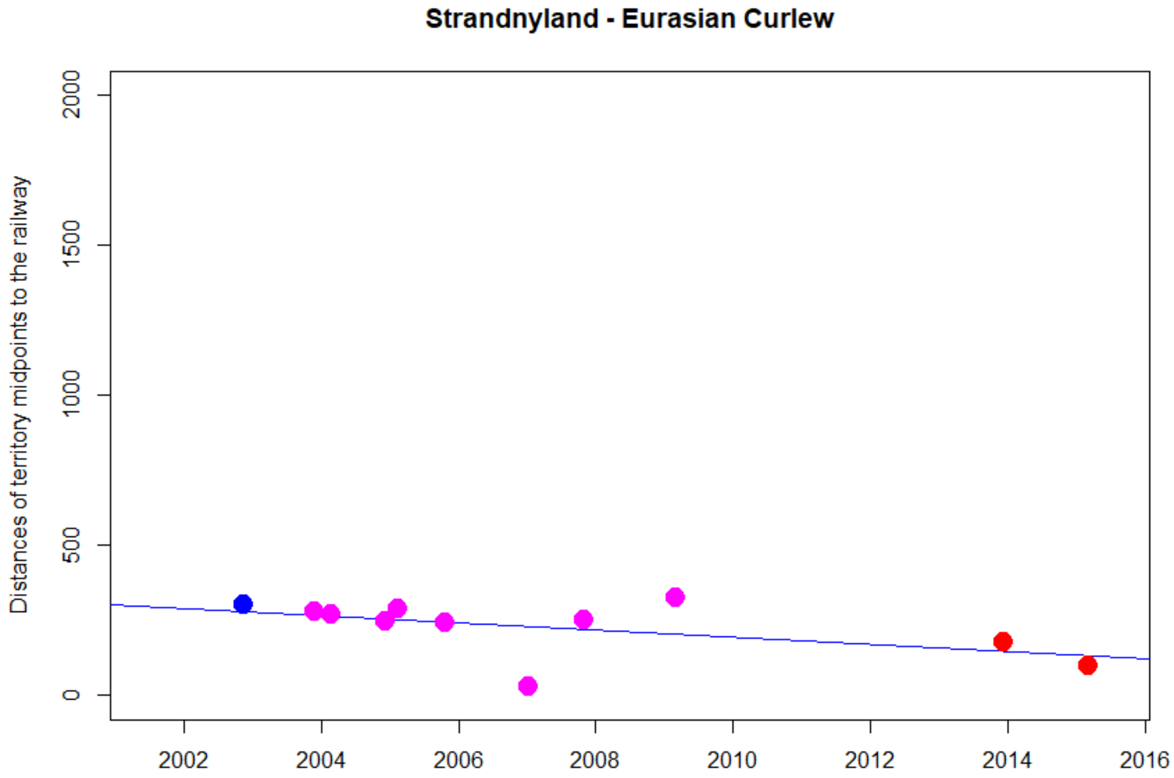


```
> cor.test(DNyland$Year,DNyland$NEAR_DIST_Botnia, method="kendall")

        Kendall's rank correlation tau

data:  DNyland$Year and DNyland$NEAR_DIST_Botnia
z = -1.8015, p-value = 0.07162
alternative hypothesis: true tau is not equal to 0
sample estimates:
      tau
-0.425999
```

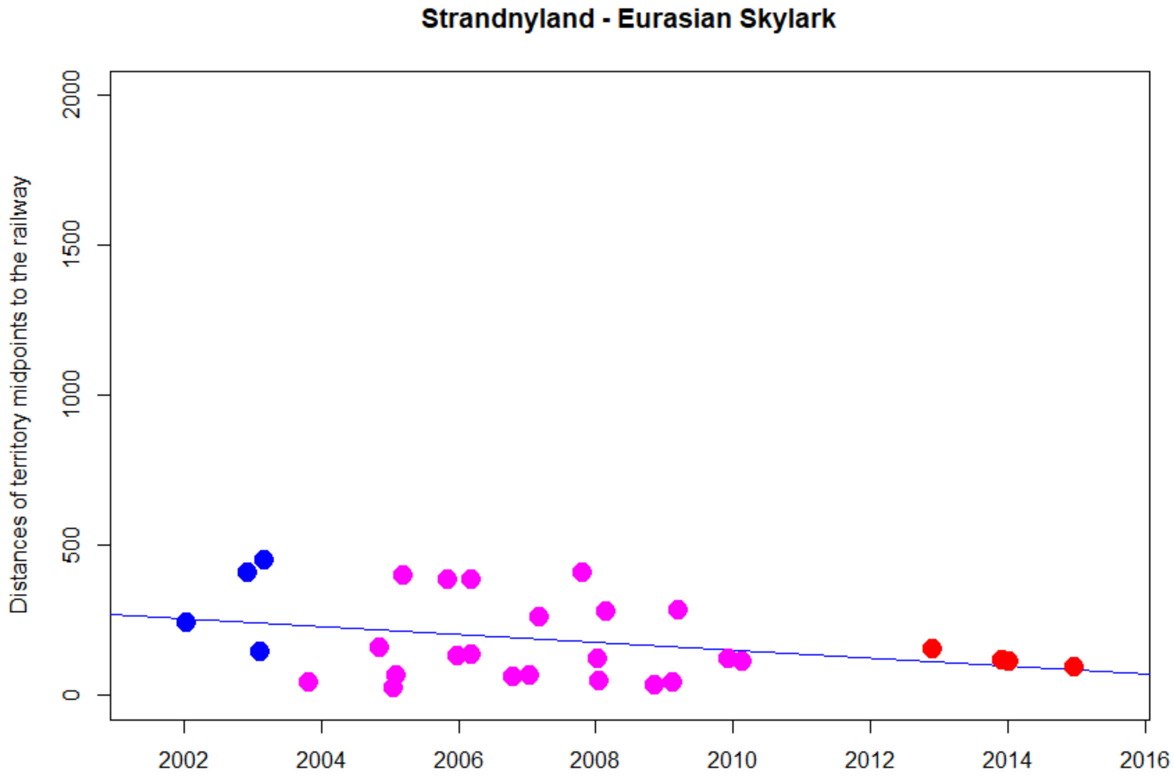


```
> cor.test(DNyland$Year,DNyland$NEAR_DIST_Botnia, method="kendall")

        Kendall's rank correlation tau

data:  DNyland$Year and DNyland$NEAR_DIST_Botnia
z = -1.6813, p-value = 0.0927
alternative hypothesis: true tau is not equal to 0
sample estimates:
       tau
-0.2277675
```

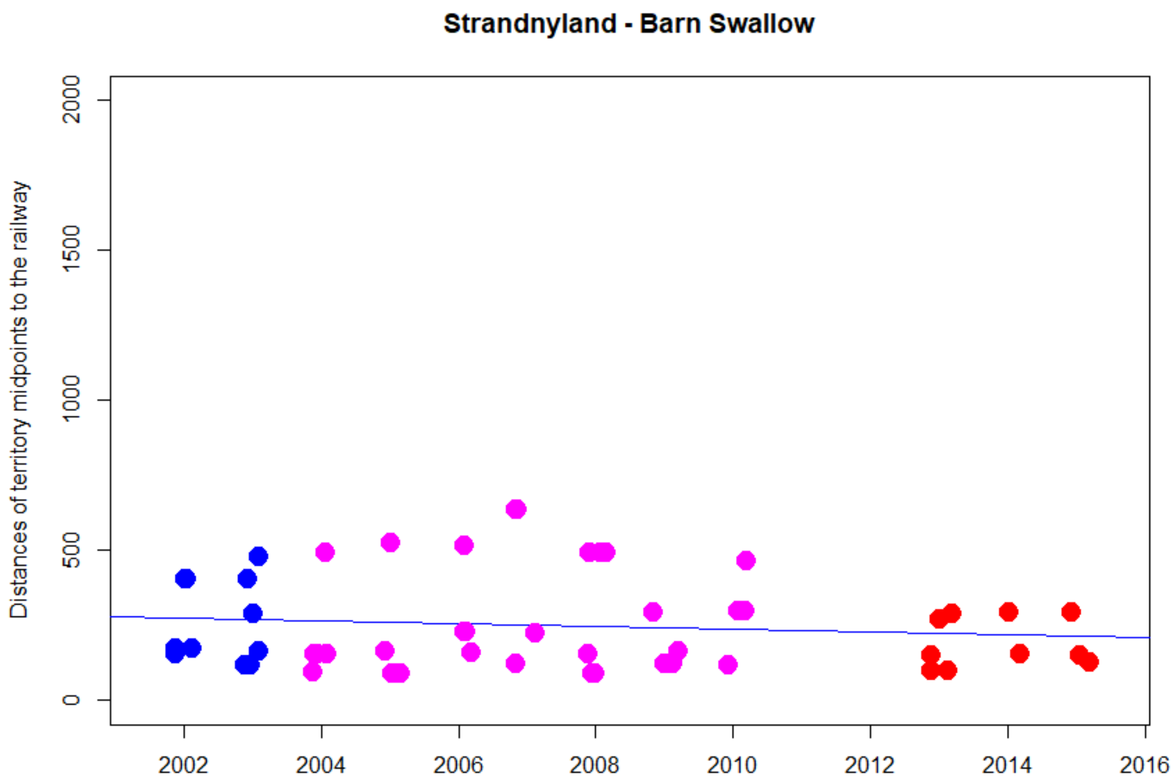


```
> cor.test(DNyland$Year,DNyland$NEAR_DIST_Botnia, method="kendall")

        Kendall's rank correlation tau

data:  DNyland$Year and DNyland$NEAR_DIST_Botnia
z = -0.6171, p-value = 0.5372
alternative hypothesis: true tau is not equal to 0
sample estimates:
        tau
-0.06068133
```

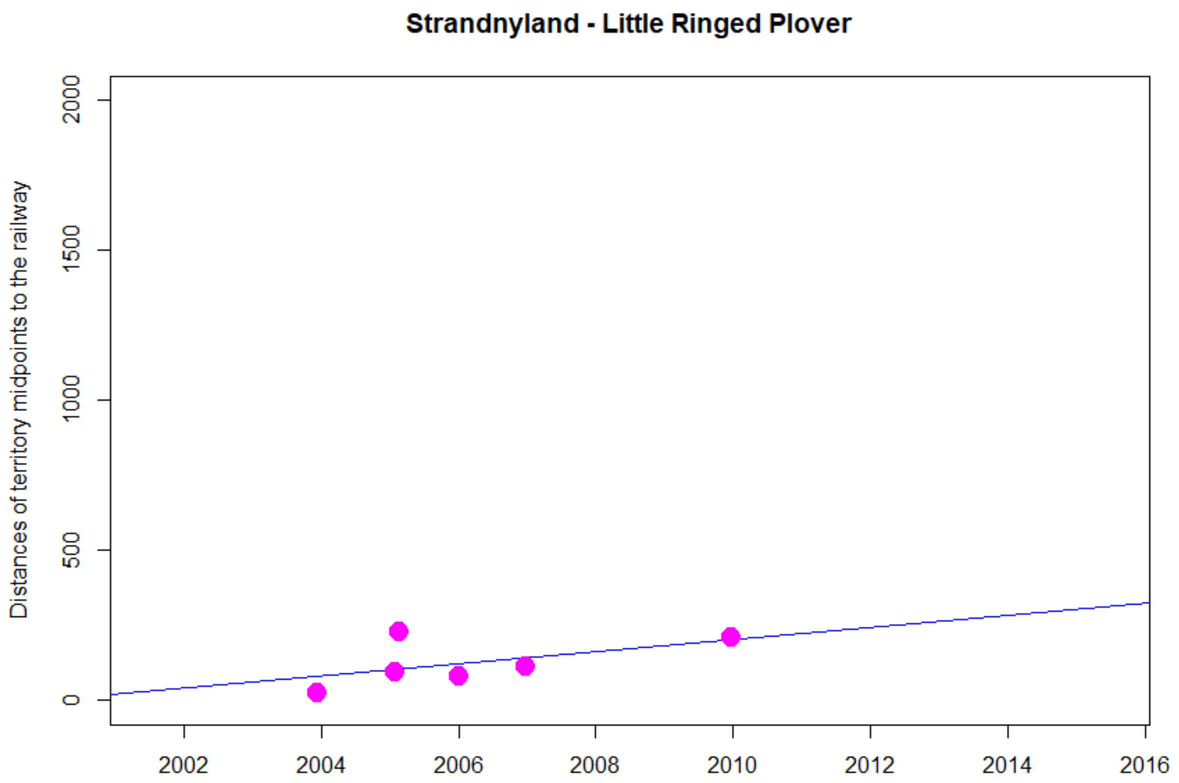


```
> cor.test(DNyland$Year,DNyland$NEAR_DIST_Botnia, method="kendall")

        Kendall's rank correlation tau

data:  DNyland$Year and DNyland$NEAR_DIST_Botnia
z = 1.1476, p-value = 0.2511
alternative hypothesis: true tau is not equal to 0
sample estimates:
      tau
0.4140393
```

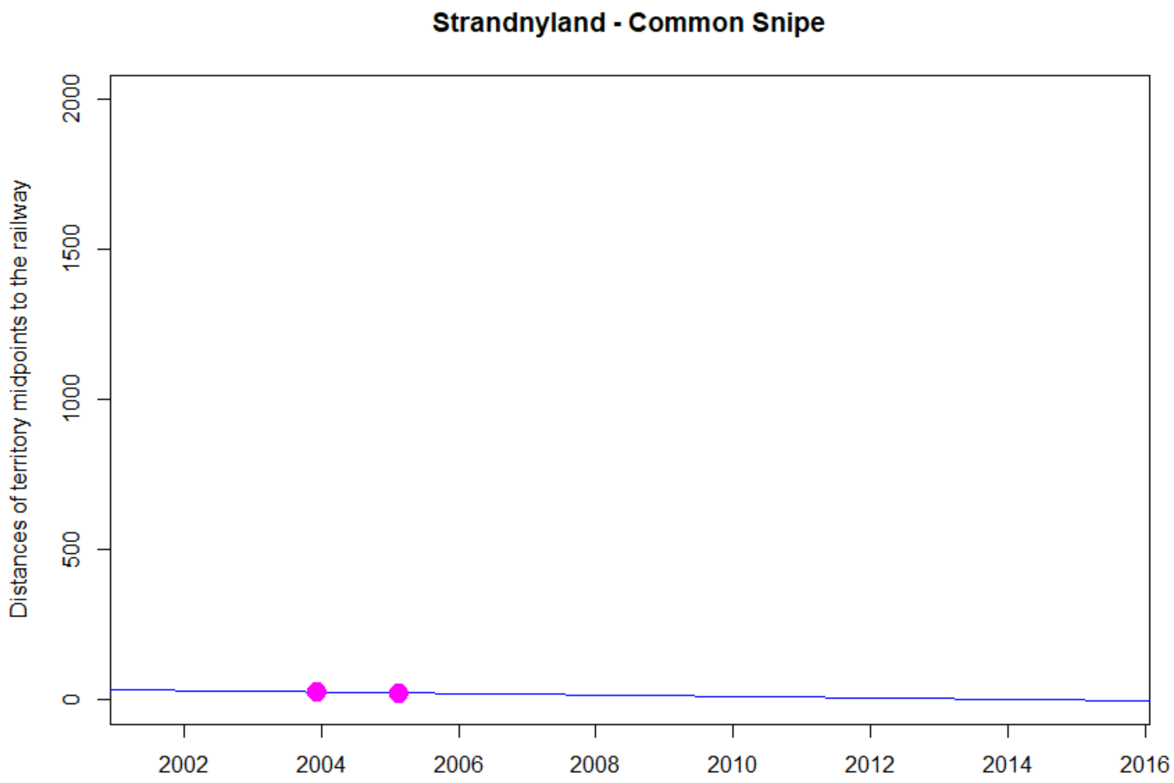
Strandnyland - Common Snipe
Distances of territory midpoints to the railway
0
500
1000
1500
2000
2002
2004
2006
2008
2010
2012
2014
2016

NA

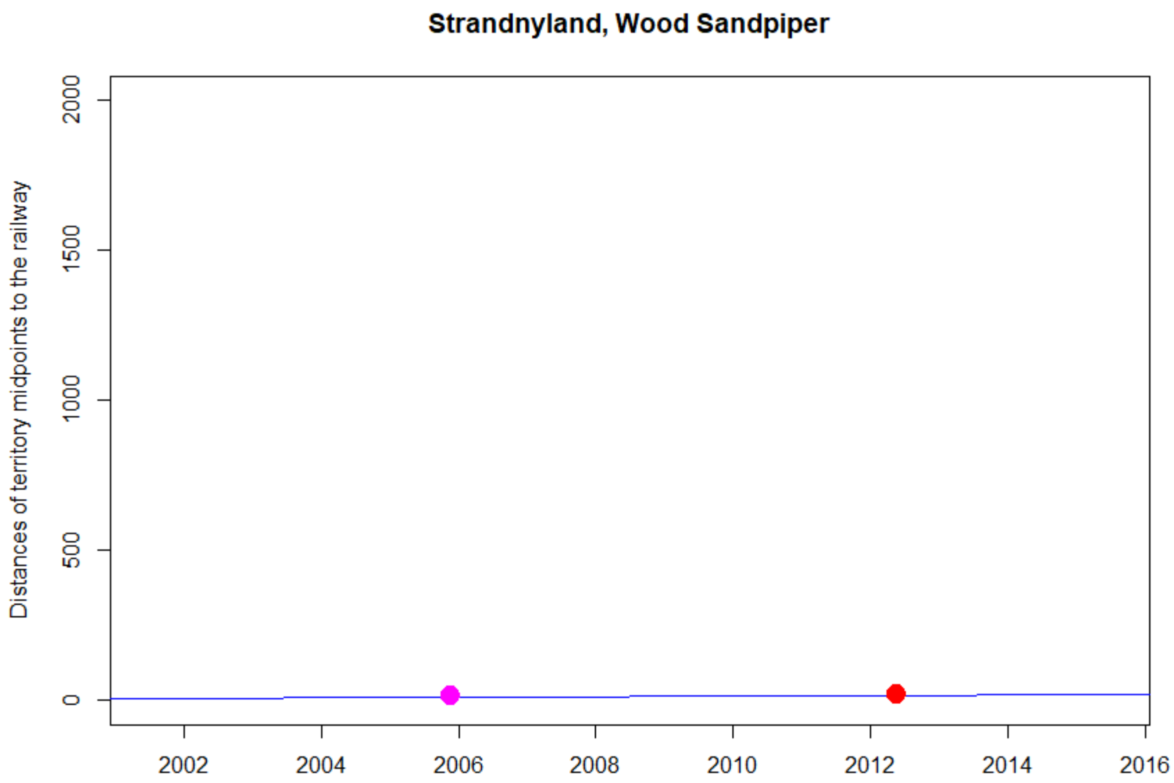

Strandnyland, Wood Sandpiper
Distances of territory midpoints to the railway
0
500
1000
1500
2000
2002
2004
2006
2008
2010
2012
2014
2016


NA

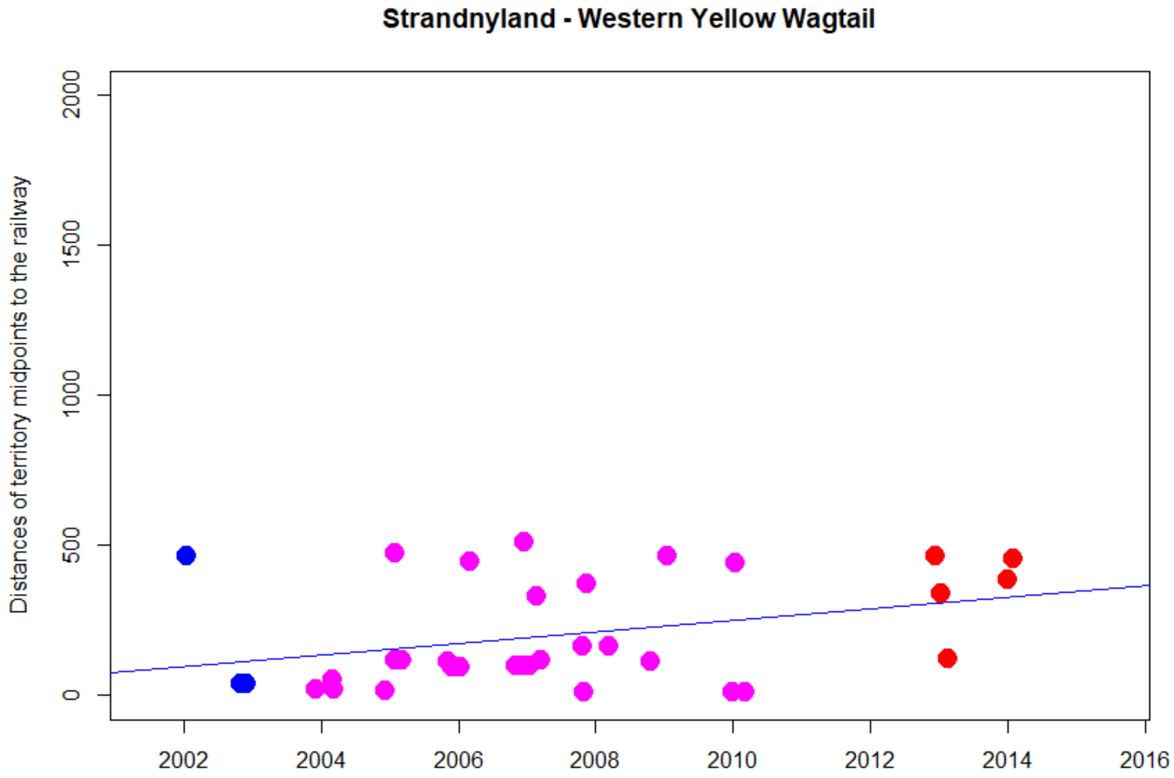


```
> cor.test(DNyland$Year,DNyland$NEAR_DIST_Botnia, method="kendall")

        Kendall's rank correlation tau

data:  DNyland$Year and DNyland$NEAR_DIST_Botnia
z = 2.0335, p-value = 0.04201
alternative hypothesis: true tau is not equal to 0
sample estimates:
      tau
0.2548265
```

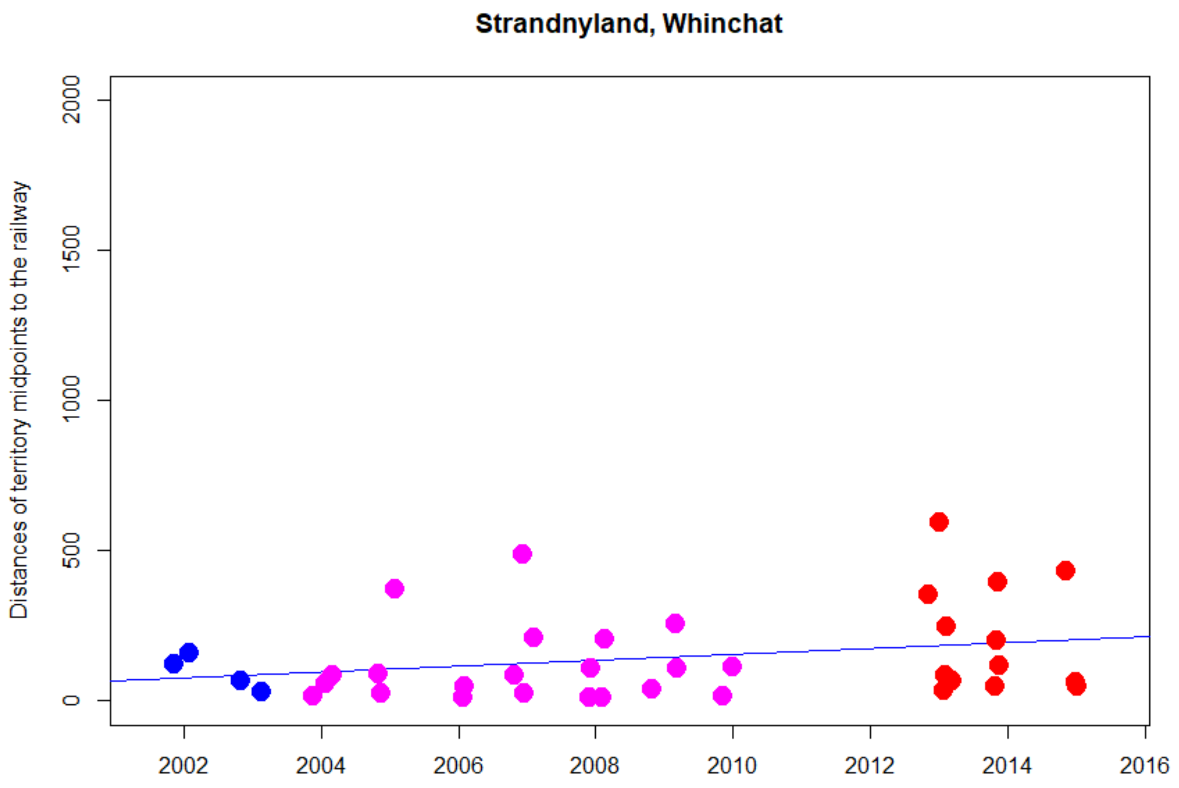


```
> cor.test(DNyland$Year,DNyland$NEAR_DIST_Botnia, method="kendall")

        Kendall's rank correlation tau

data:  DNyland$Year and DNyland$NEAR_DIST_Botnia
z = 1.214, p-value = 0.2247
alternative hypothesis: true tau is not equal to 0
sample estimates:
      tau
0.1416817
```

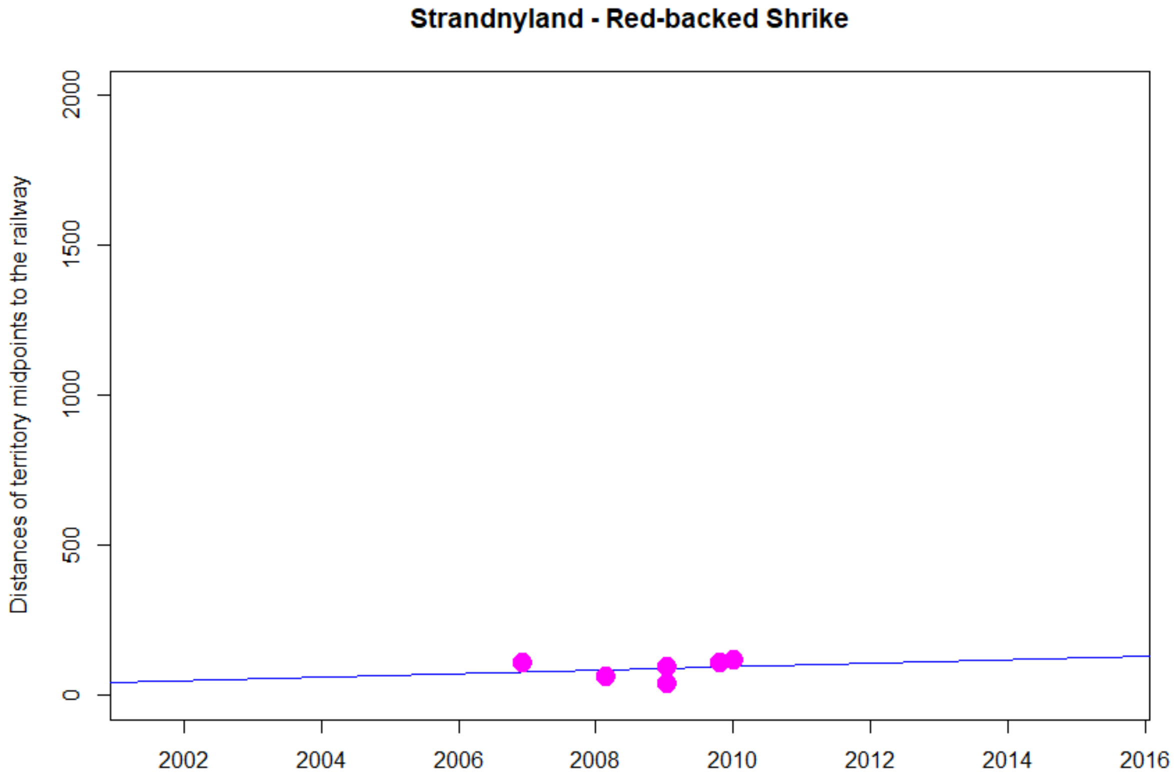


```
> cor.test(DNyland$Year,DNyland$NEAR_DIST_Botnia, method="kendall")

        Kendall's rank correlation tau

data:  DNyland$Year and DNyland$NEAR_DIST_Botnia
z = 0.97435, p-value = 0.3299
alternative hypothesis: true tau is not equal to 0
sample estimates:
      tau
0.3580574
```

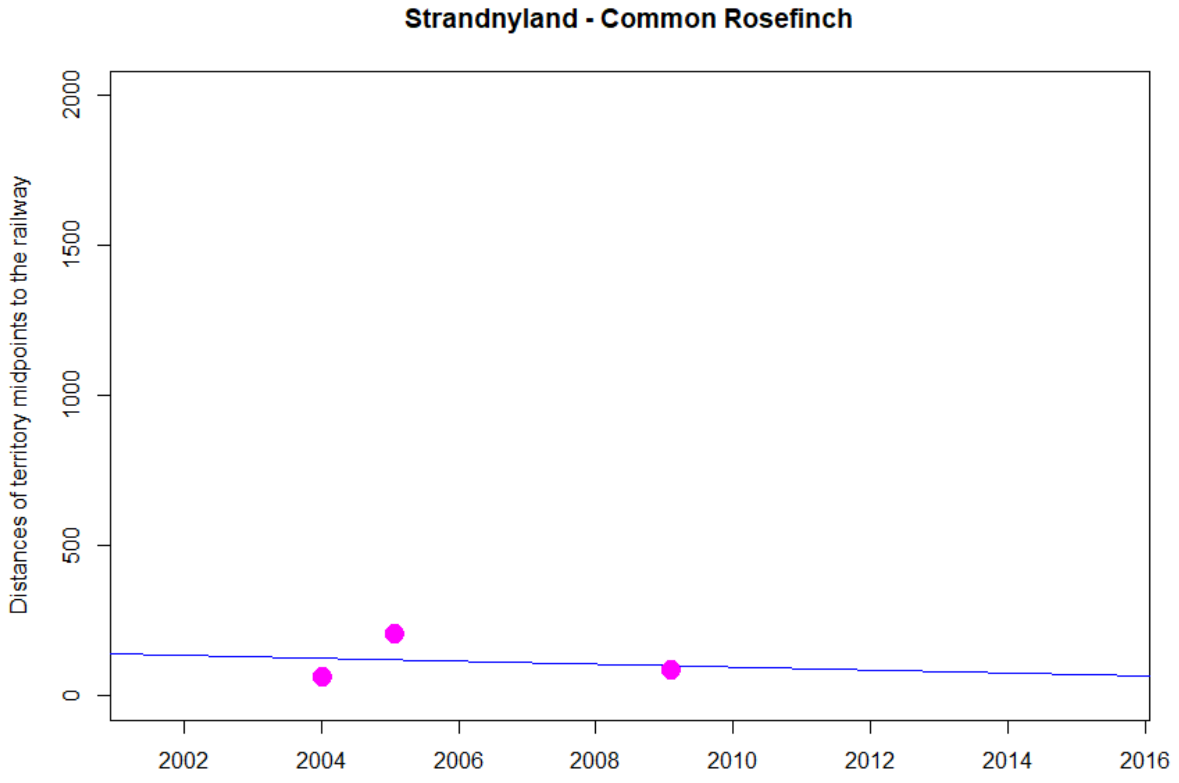

Strandnyland - Common Rosefinch
Distances of territory midpoints to the railway
0
500
1000
1500
2000
2002
2004
2006
2008
2010
2012
2014
2016


NA

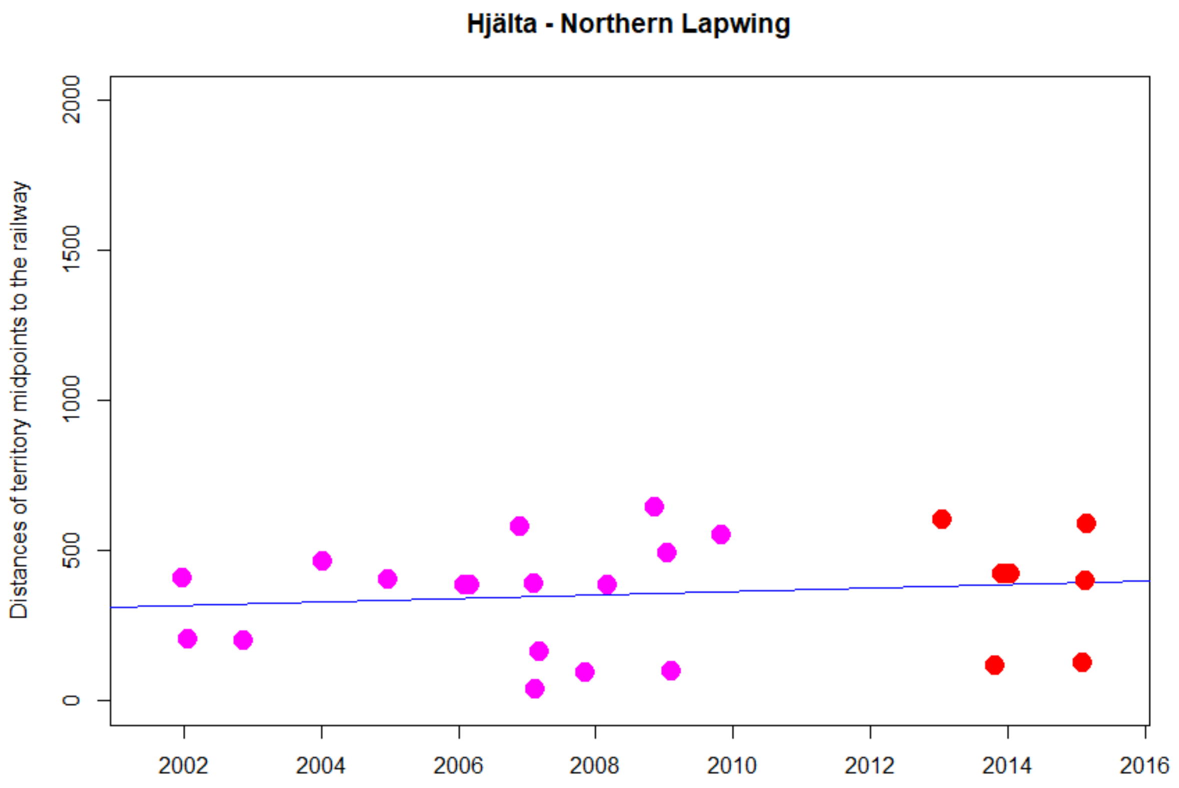


```
> cor.test(DNyland$Year,DNyland$NEAR_DIST_Botnia, method="kendall")

        Kendall's rank correlation tau

data:  DNyland$Year and DNyland$NEAR_DIST_Botnia
z = 0.69983, p-value = 0.484
alternative hypothesis: true tau is not equal to 0
sample estimates:
      tau
0.1053108
```

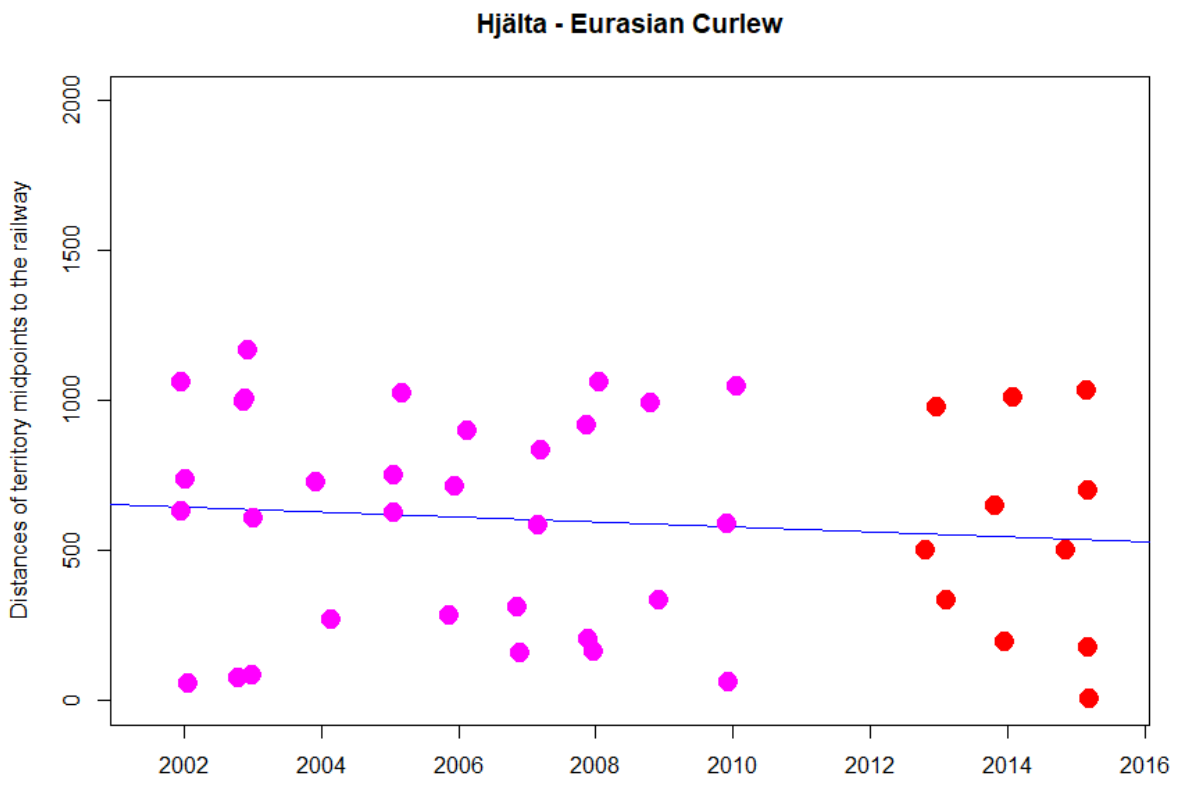


```
> cor.test(DNyland$Year,DNyland$NEAR_DIST_Botnia, method="kendall")

        Kendall's rank correlation tau

data:  DNyland$Year and DNyland$NEAR_DIST_Botnia
z = -0.64286, p-value = 0.5203
alternative hypothesis: true tau is not equal to 0
sample estimates:
        tau
-0.07104511
```

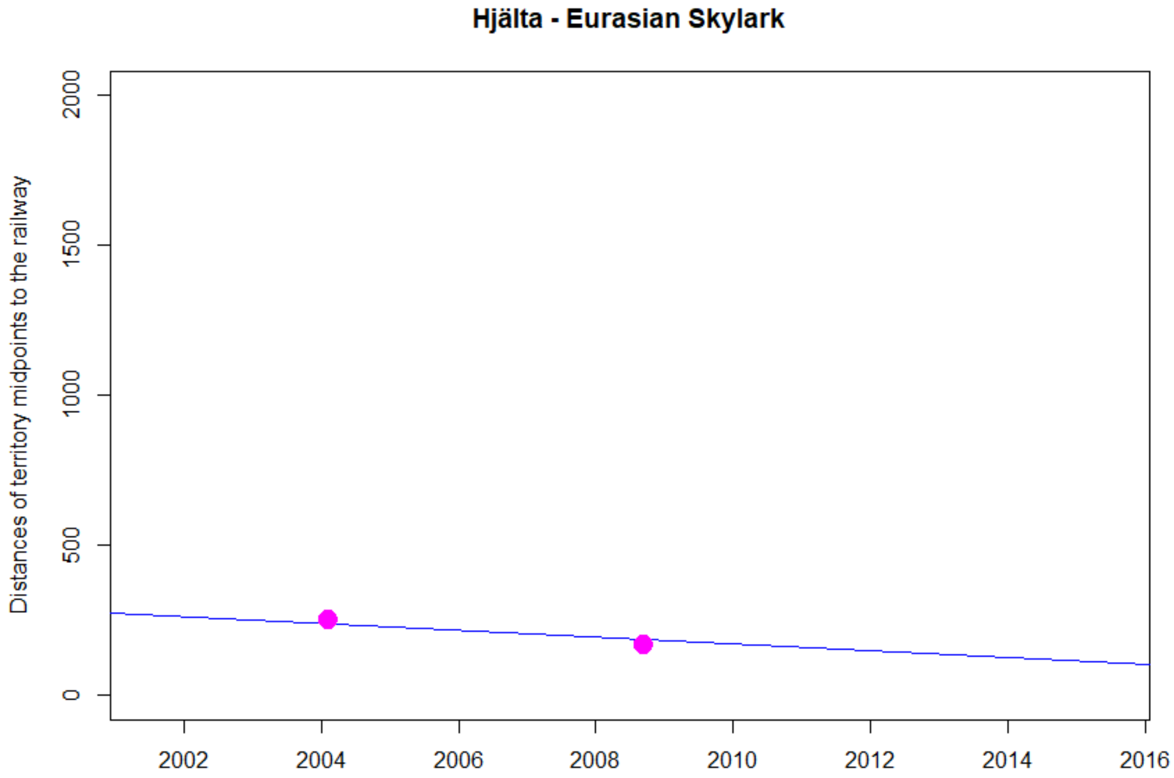
Hjälta - Eurasian Skylark
Distances of territory midpoints to the railway
0
500
1000
1500
2000
2002
2004
2006
2008
2010
2012
2014
2016

NA

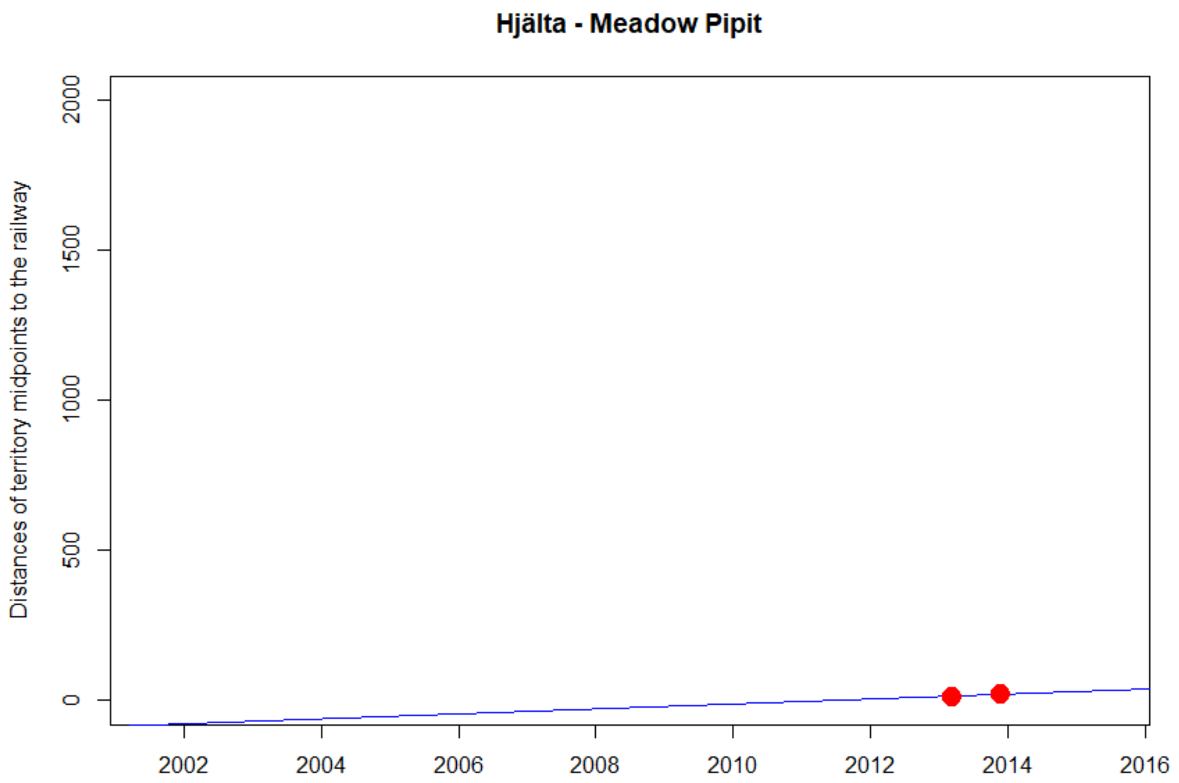
Hjälta - Meadow Pipit
Distances of territory midpoints to the railway
0
500
1000
1500
2000
2002
2004
2006
2008
2010
2012
2014
2016

NA

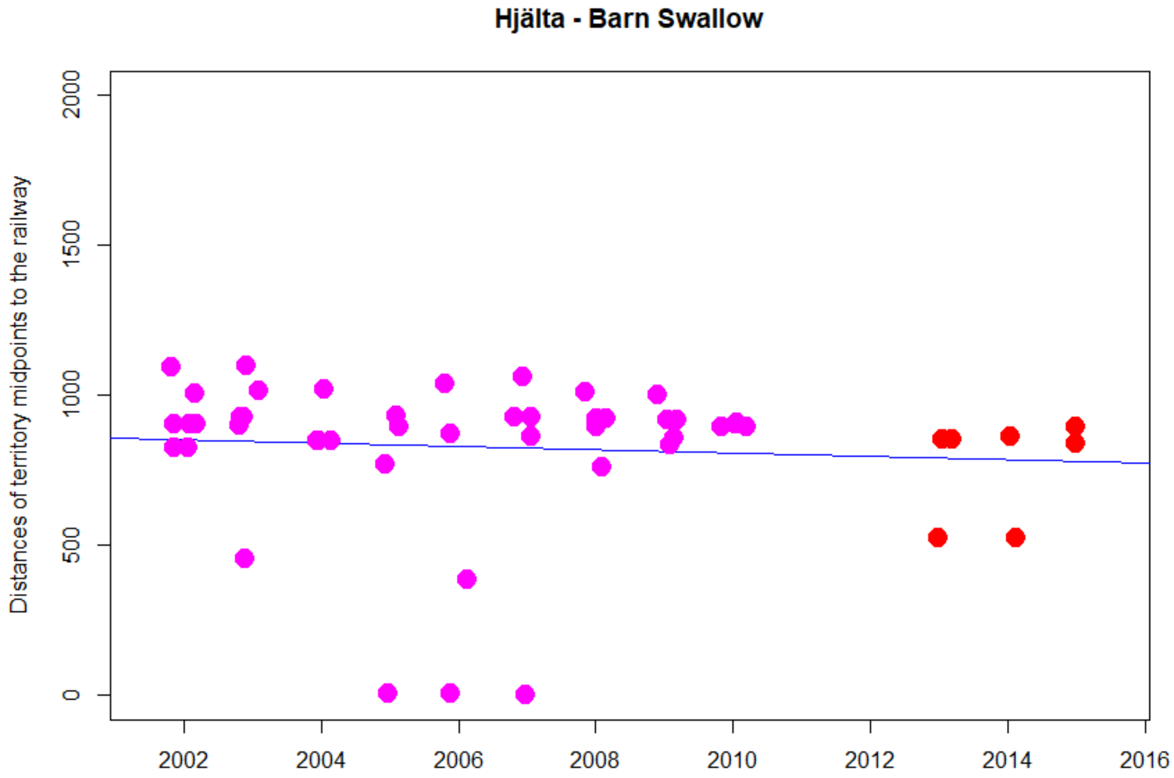


```
> cor.test(DNyland$Year,DNyland$NEAR_DIST_Botnia, method="kendall")

        Kendall's rank correlation tau

data:  DNyland$Year and DNyland$NEAR_DIST_Botnia
z = -1.6666, p-value = 0.09559
alternative hypothesis: true tau is not equal to 0
sample estimates:
       tau
-0.1671042
```

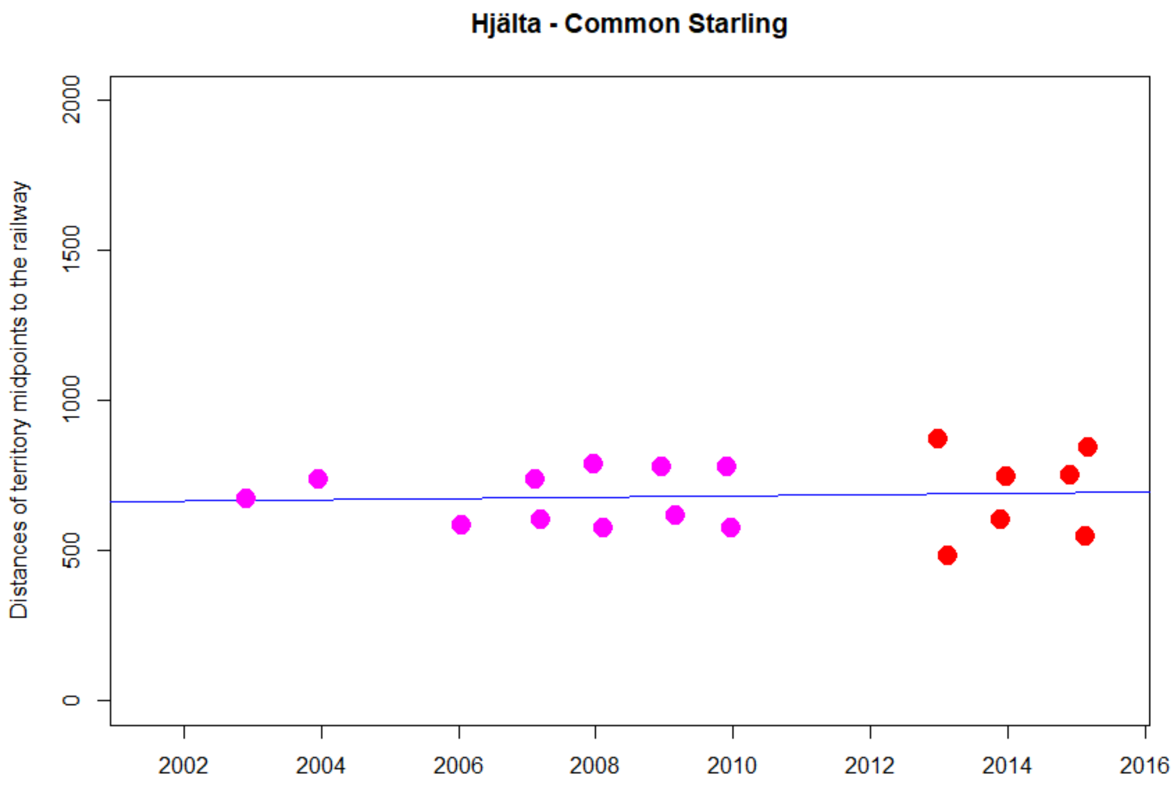


```
> cor.test(DNyland$Year,DNyland$NEAR_DIST_Botnia, method="kendall")

        Kendall's rank correlation tau

data:  DNyland$Year and DNyland$NEAR_DIST_Botnia
z = 0.45772, p-value = 0.6472
alternative hypothesis: true tau is not equal to 0
sample estimates:
       tau
0.08084521
```

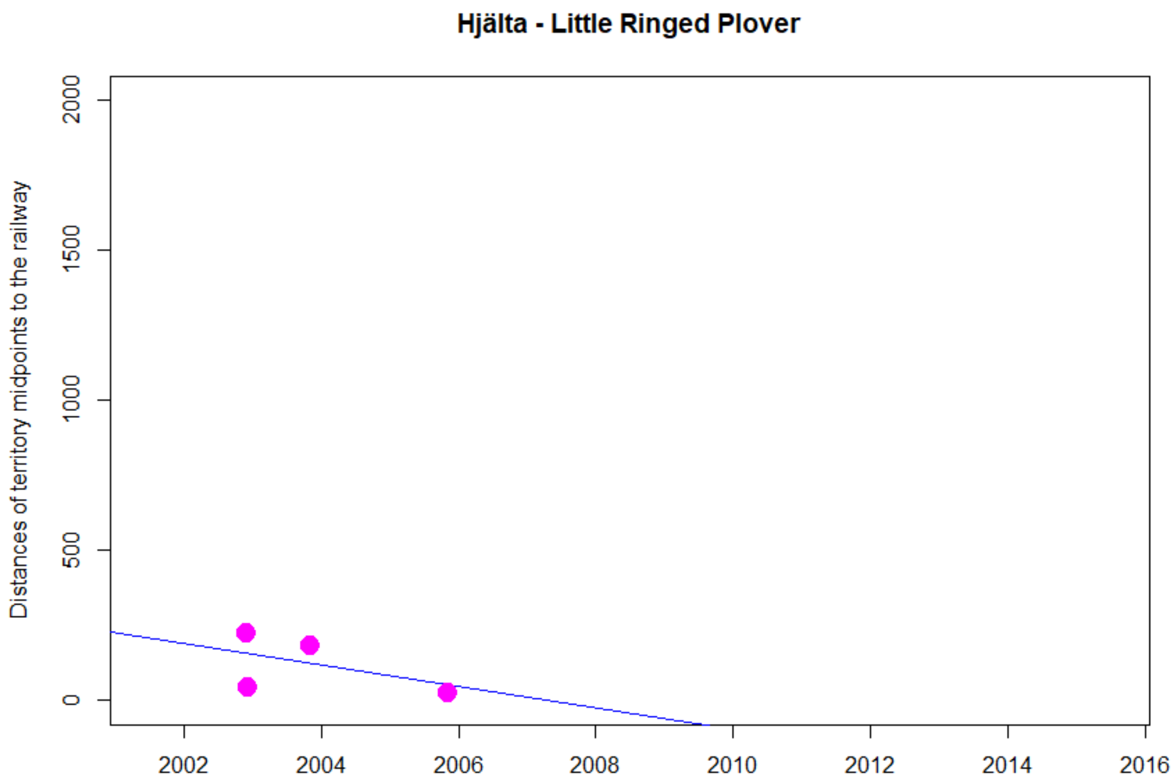


```
> cor.test(DNyland$Year,DNyland$NEAR_DIST_Botnia, method="kendall")

        Kendall's rank correlation tau

data:  DNyland$Year and DNyland$NEAR_DIST_Botnia
z = -1.0835, p-value = 0.2786
alternative hypothesis: true tau is not equal to 0
sample estimates:
       tau
-0.5477226
```

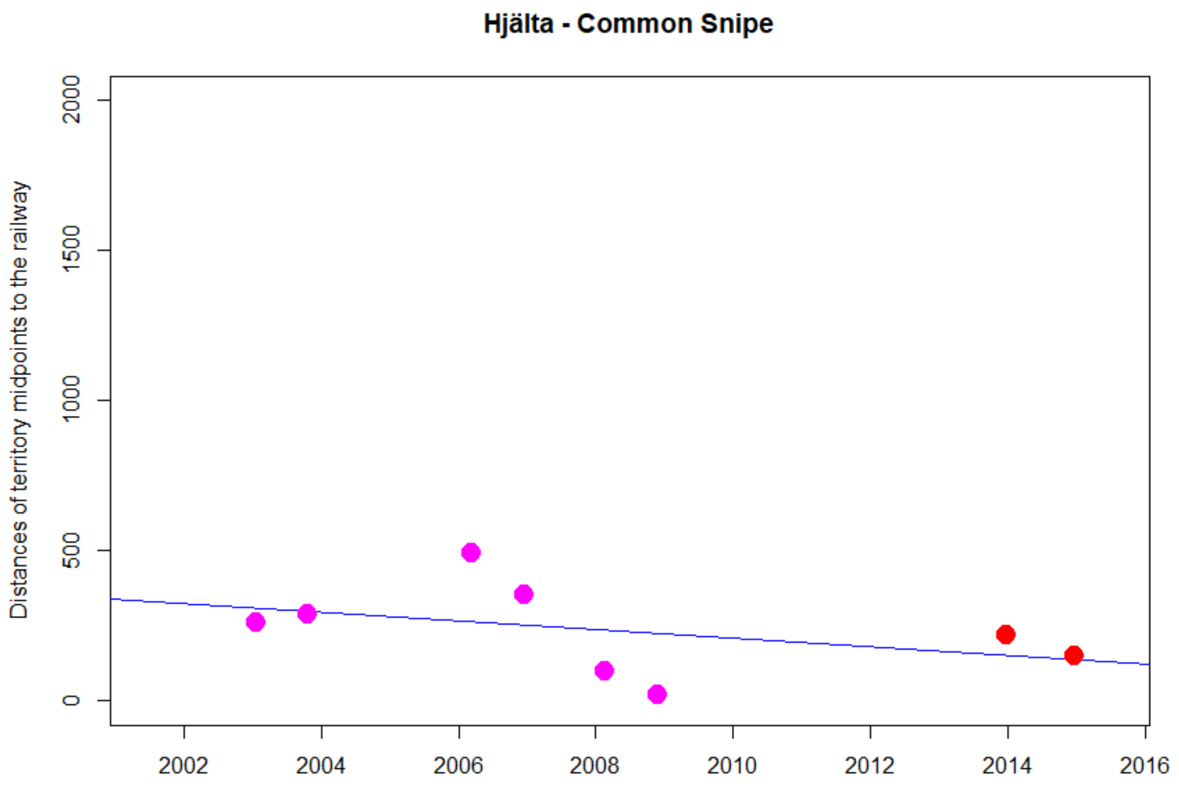


```
> cor.test(DNyland$Year,DNyland$NEAR_DIST_Botnia, method="kendall")

        Kendall's rank correlation tau

data:  DNyland$Year and DNyland$NEAR_DIST_Botnia
T = 9, p-value = 0.2751
alternative hypothesis: true tau is not equal to 0
sample estimates:
       tau
-0.3571429
```

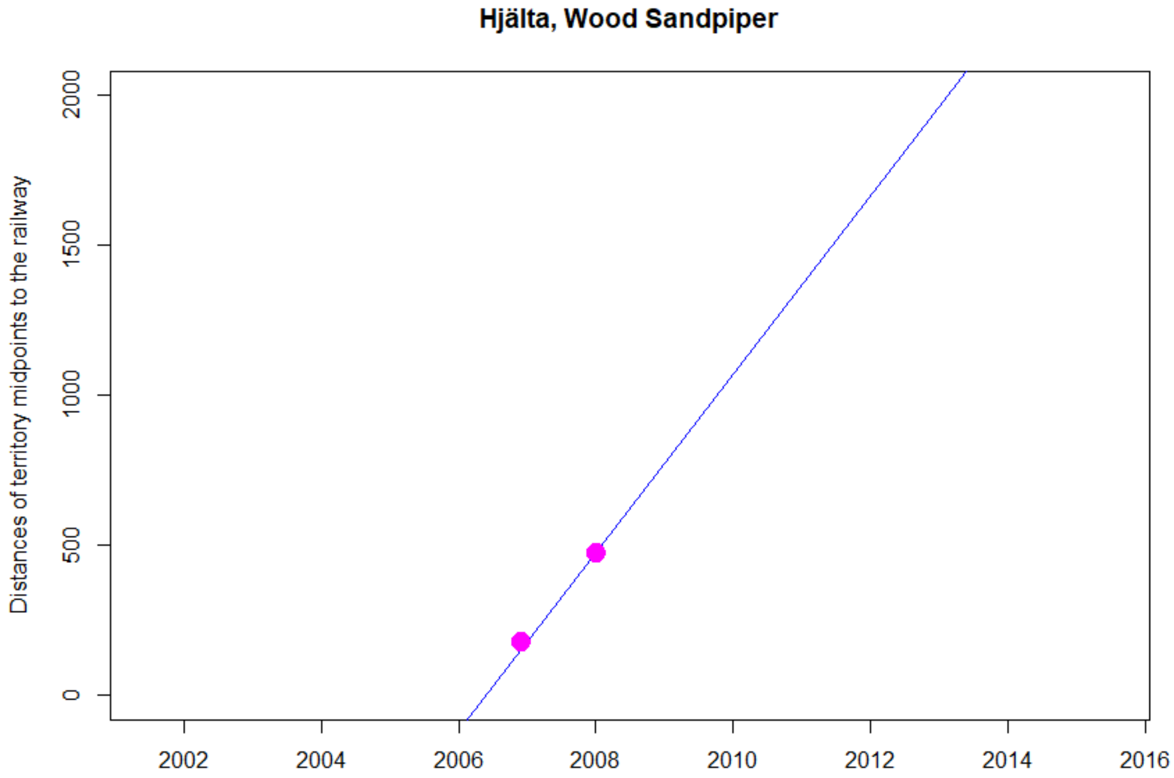
Hjälta, Wood Sandpiper
Distances of territory midpoints to the railway
0
500
1000
1500
2000
2002
2004
2006
2008
2010
2012
2014
2016

NA

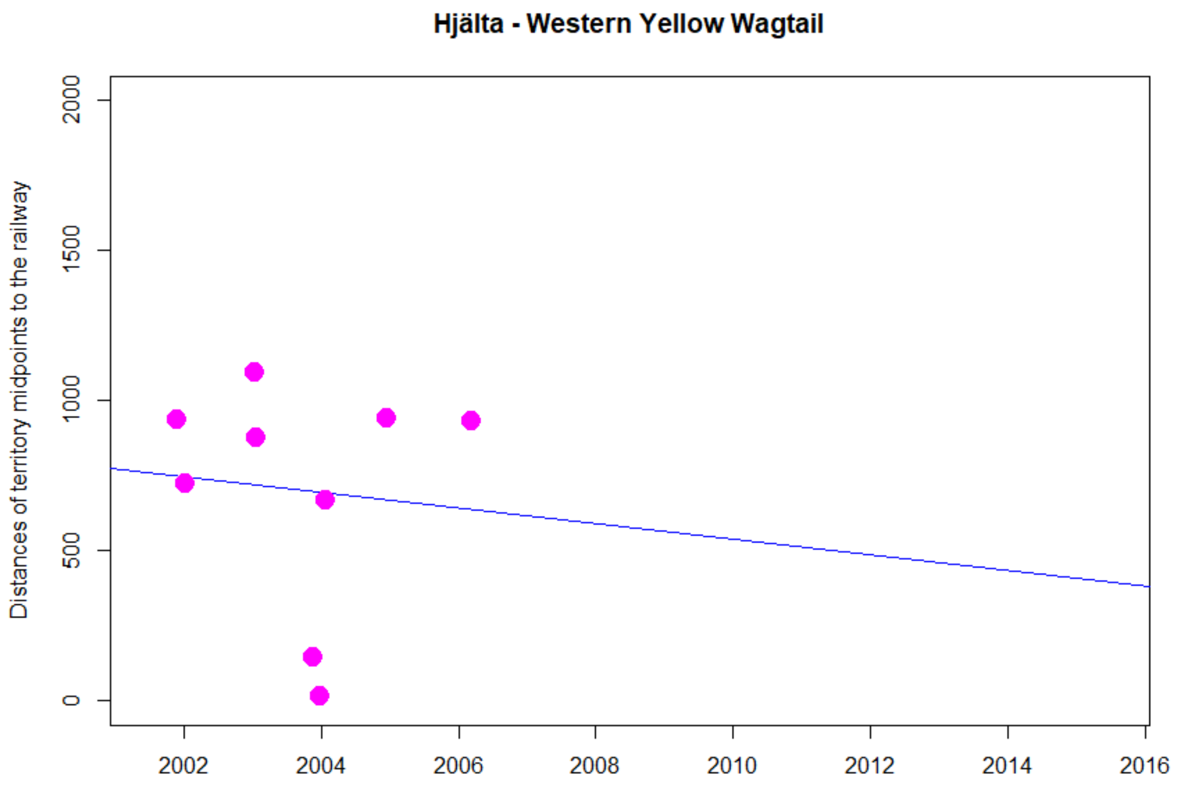


```
> cor.test(DNyland$Year,DNyland$NEAR_DIST_Botnia, method="kendall")

        Kendall's rank correlation tau

data:  DNyland$Year and DNyland$NEAR_DIST_Botnia
z = -0.32287, p-value = 0.7468
alternative hypothesis: true tau is not equal to 0
sample estimates:
        tau
-0.08980265
```

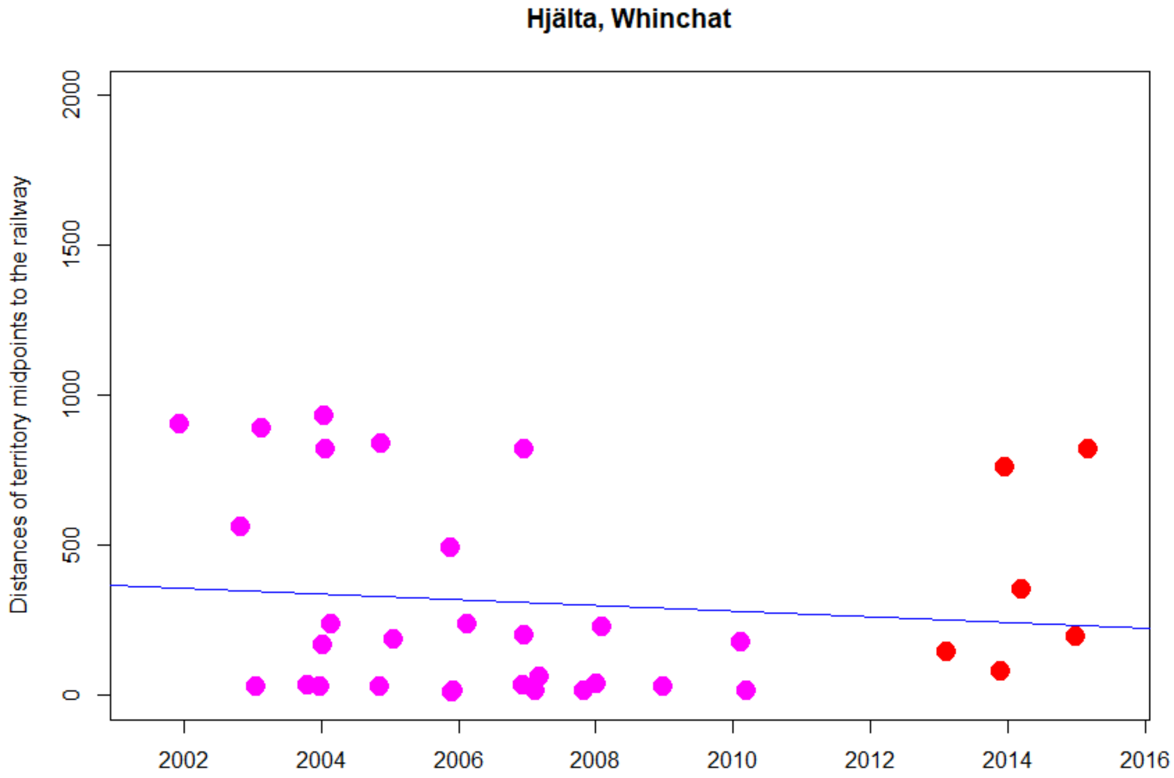


```
> cor.test(DNyland$Year,DNyland$NEAR_DIST_Botnia, method="kendall")

        Kendall's rank correlation tau

data:  DNyland$Year and DNyland$NEAR_DIST_Botnia
z = -1.0458, p-value = 0.2956
alternative hypothesis: true tau is not equal to 0
sample estimates:
       tau
-0.1301043
```

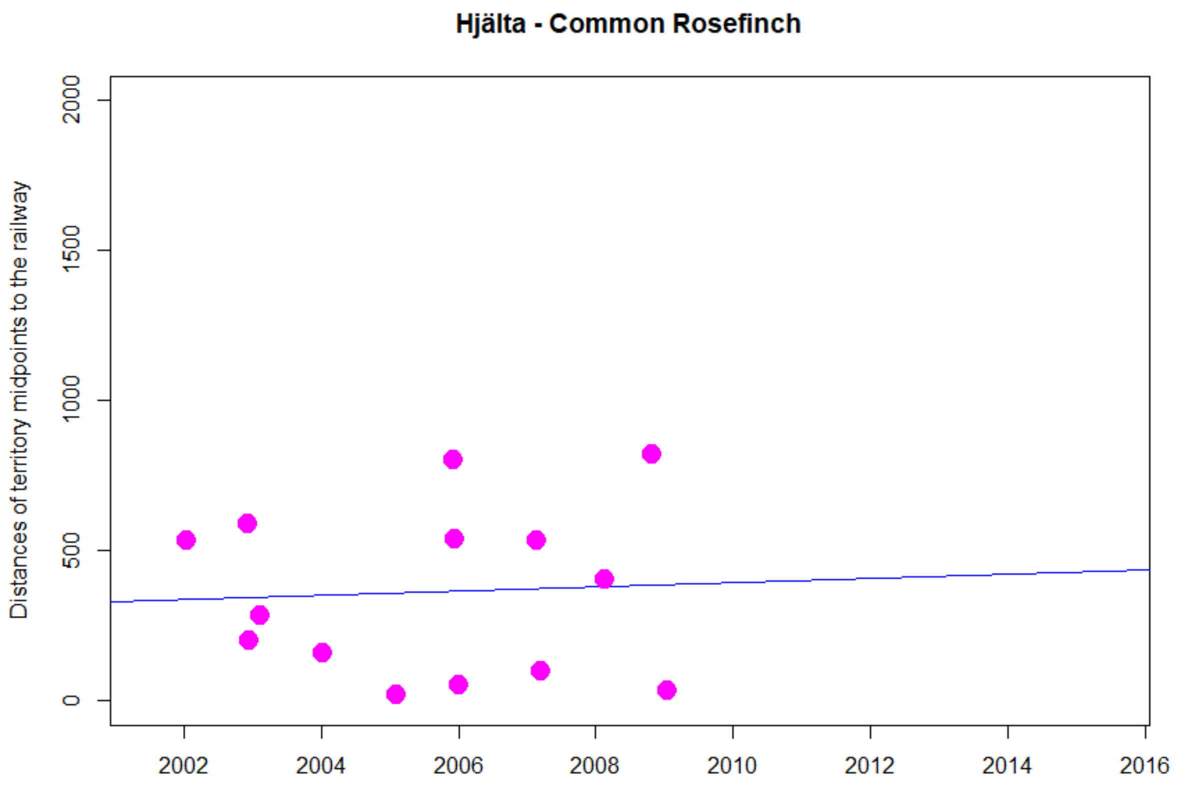


```
> cor.test(DNyland$Year,DNyland$NEAR_DIST_Botnia, method="kendall")

        Kendall's rank correlation tau

data:  DNyland$Year and DNyland$NEAR_DIST_Botnia
z = -0.27763, p-value = 0.7813
alternative hypothesis: true tau is not equal to 0
sample estimates:
       tau
-0.0575321
```

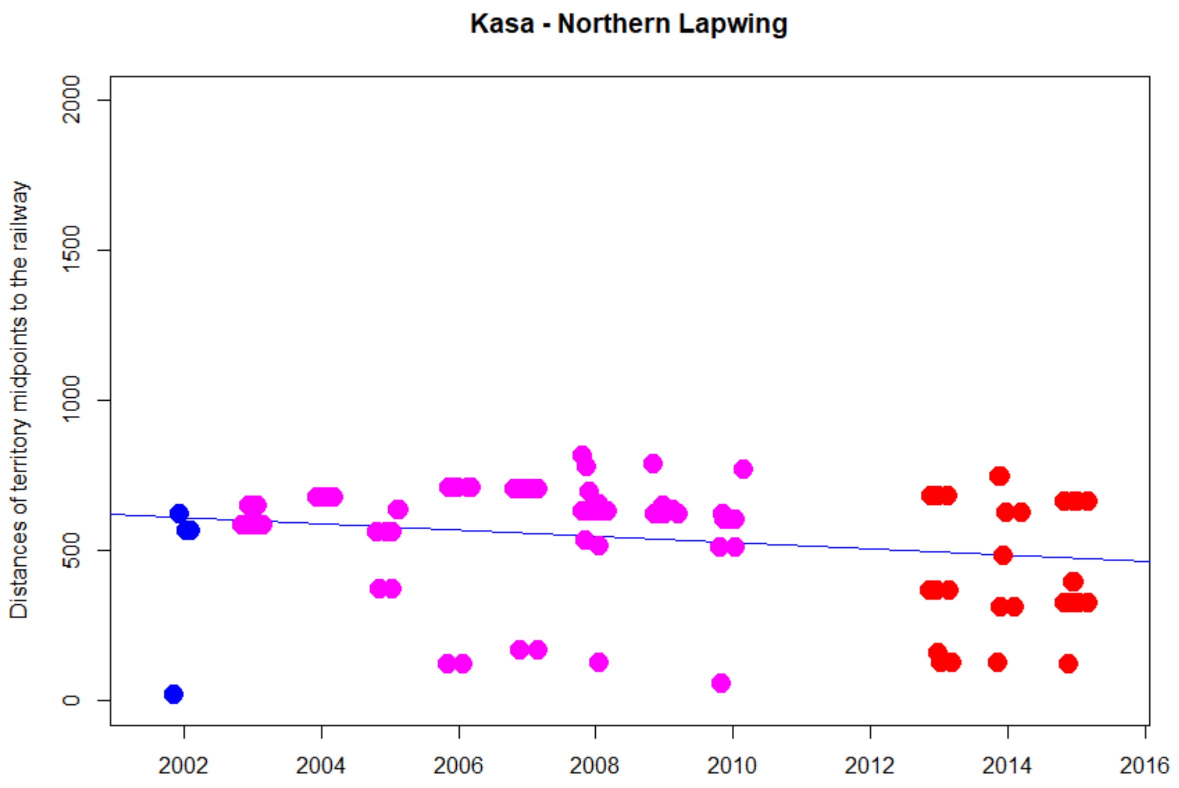


```
> cor.test(DNyland$Year,DNyland$NEAR_DIST_Botnia, method="kendall")

        Kendall's rank correlation tau

data:  DNyland$Year and DNyland$NEAR_DIST_Botnia
z = -1.6296, p-value = 0.1032
alternative hypothesis: true tau is not equal to 0
sample estimates:
       tau
-0.1122012
```

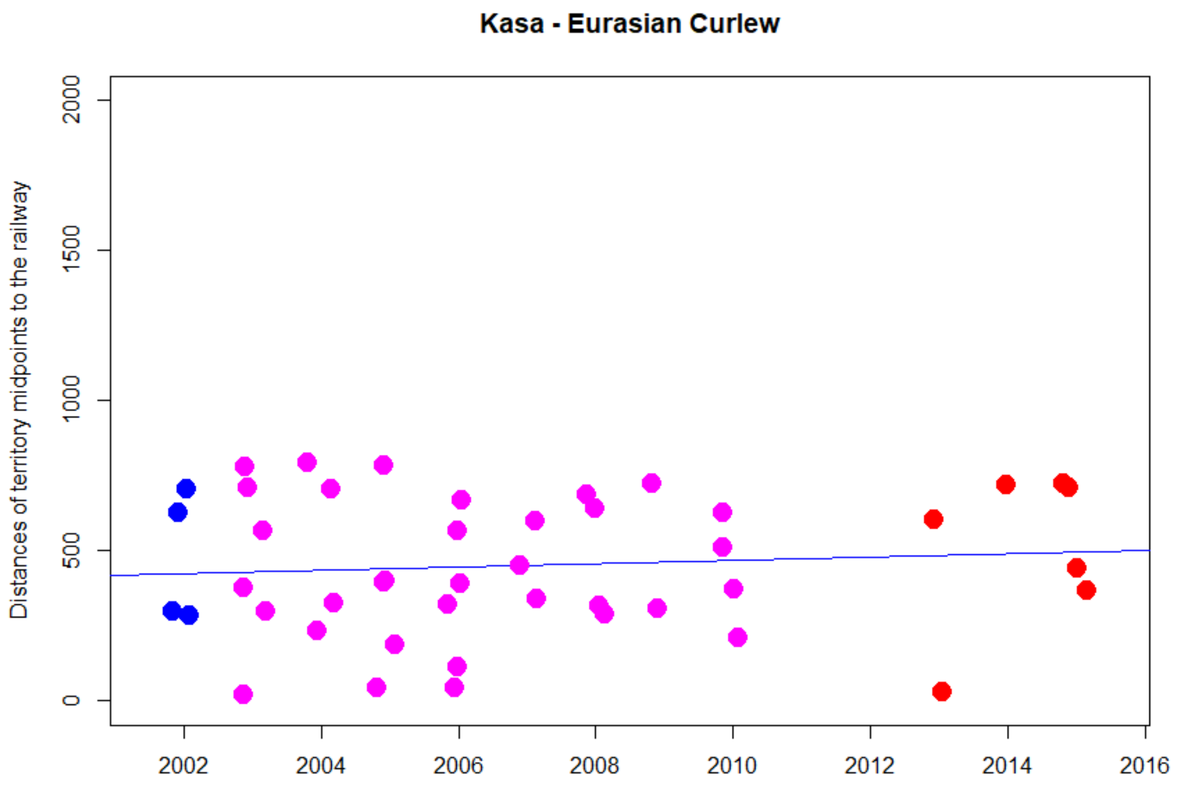


```
> cor.test(DNyland$Year,DNyland$NEAR_DIST_Botnia, method="kendall")

        Kendall's rank correlation tau

data:  DNyland$Year and DNyland$NEAR_DIST_Botnia
z = 0.61993, p-value = 0.5353
alternative hypothesis: true tau is not equal to 0
sample estimates:
       tau
0.06619305
```

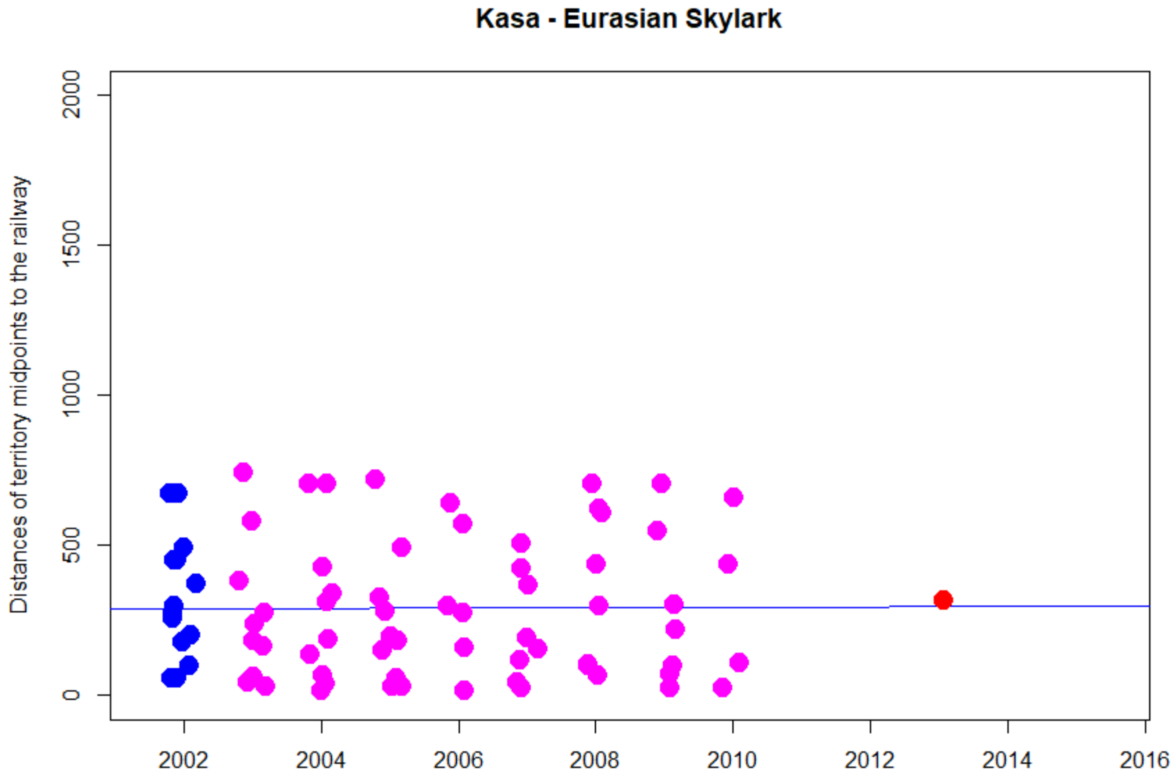


```
> cor.test(DNyland$Year,DNyland$NEAR_DIST_Botnia, method="kendall")

        Kendall's rank correlation tau

data:  DNyland$Year and DNyland$NEAR_DIST_Botnia
z = -0.23936, p-value = 0.8108
alternative hypothesis: true tau is not equal to 0
sample estimates:
        tau
-0.01939739
```

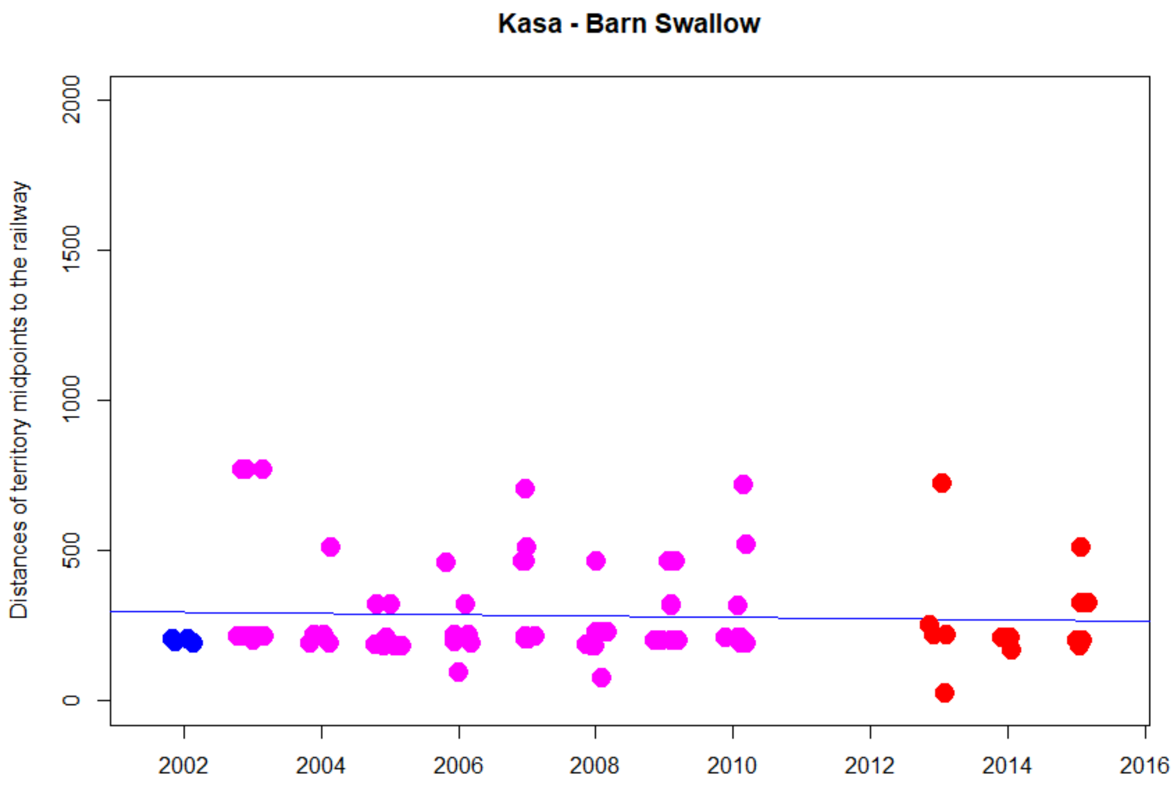


```
> cor.test(DNyland$Year,DNyland$NEAR_DIST_Botnia, method="kendall")

        Kendall's rank correlation tau

data:  DNyland$Year and DNyland$NEAR_DIST_Botnia
z = 0.25702, p-value = 0.7972
alternative hypothesis: true tau is not equal to 0
sample estimates:
       tau
0.02004661
```

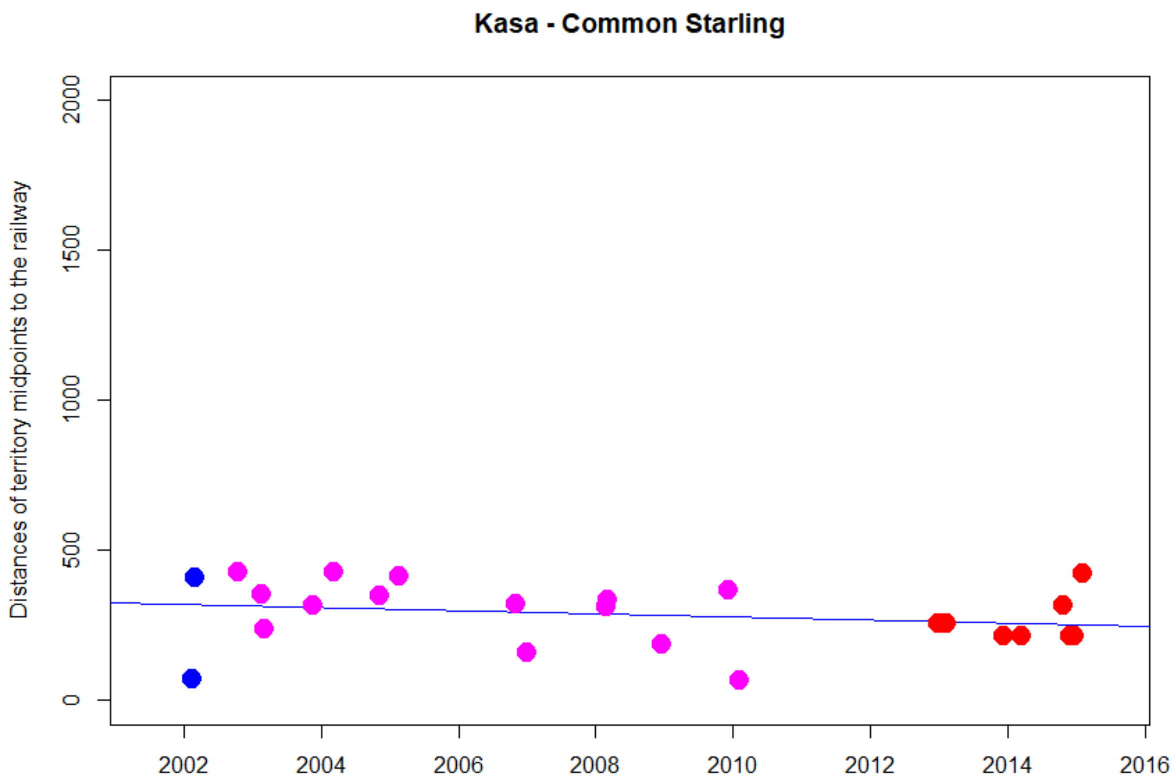


```
> cor.test(DNyland$Year,DNyland$NEAR_DIST_Botnia, method="kendall")

        Kendall's rank correlation tau

data:  DNyland$Year and DNyland$NEAR_DIST_Botnia
z = -1.3741, p-value = 0.1694
alternative hypothesis: true tau is not equal to 0
sample estimates:
       tau
-0.2068385
```

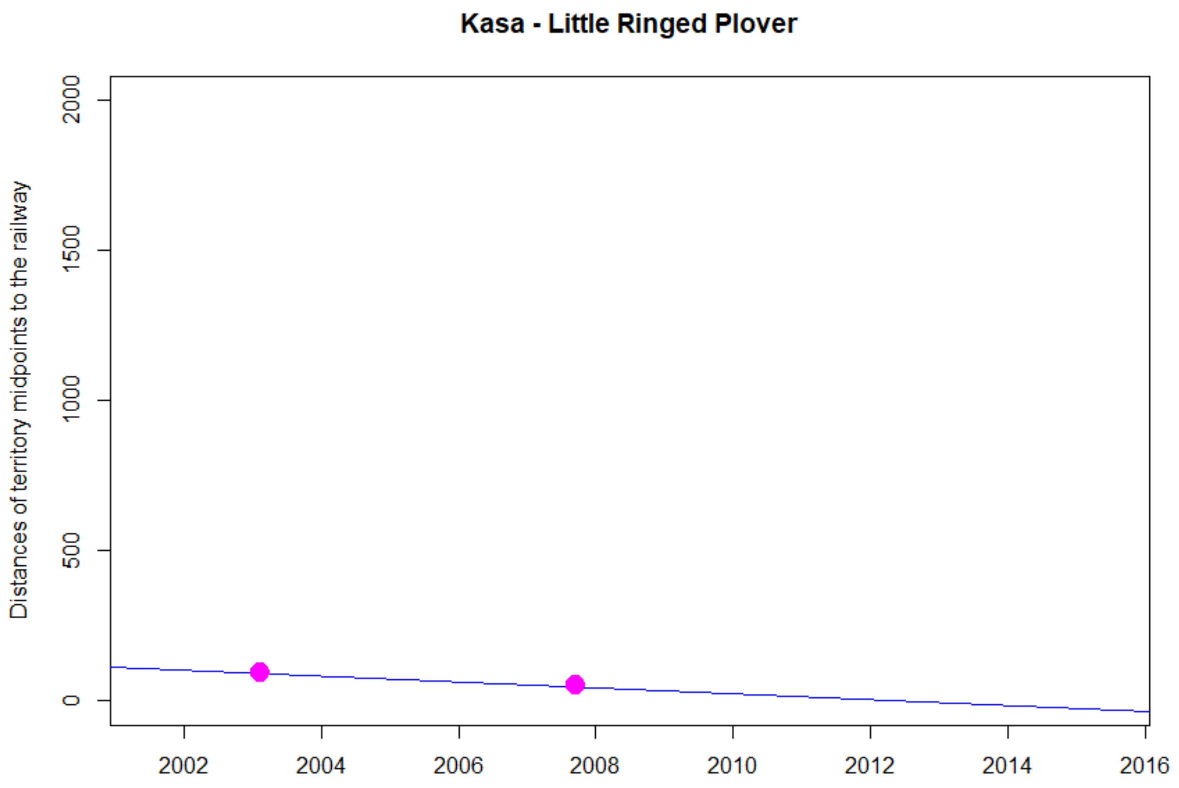
Kasa - Little Ringed Plover
Distances of territory midpoints to the railway
0
500
1000
1500
2000
2002
2004
2006
2008
2010
2012
2014
2016

NA

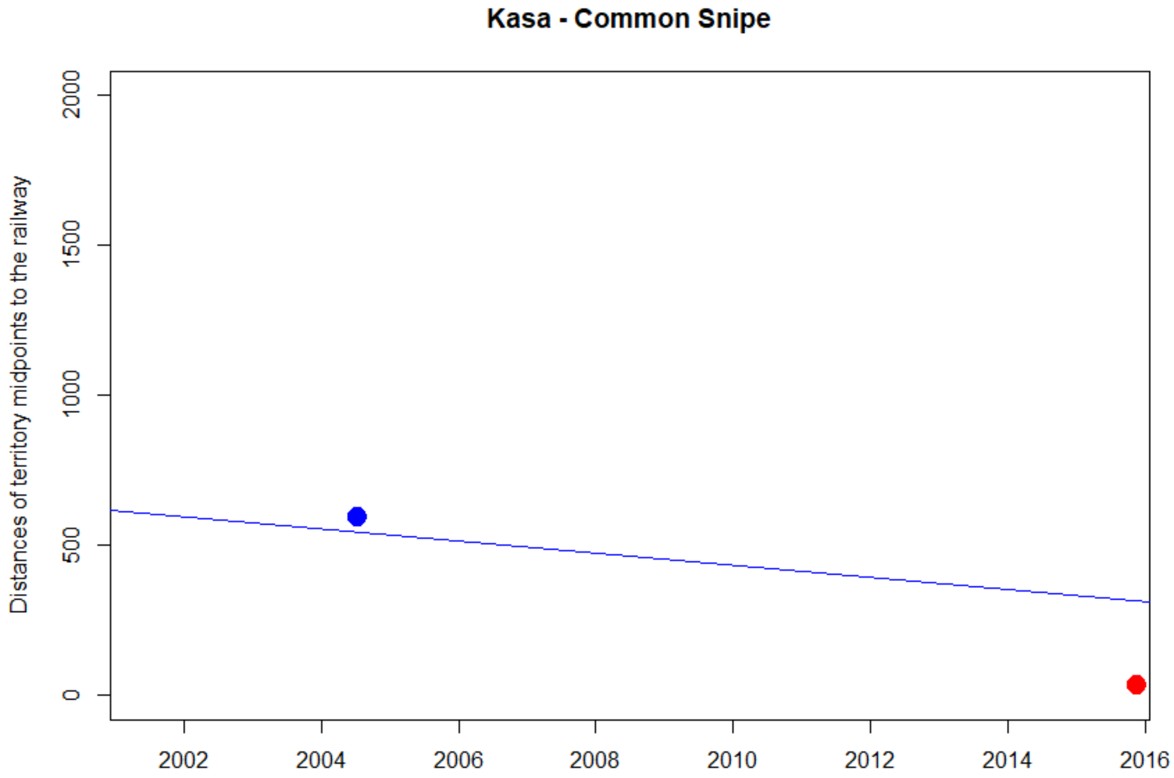
Kasa - Common Snipe
Distances of territory midpoints to the railway
0
500
1000
1500
2000
2002
2004
2006
2008
2010
2012
2014
2016

NA

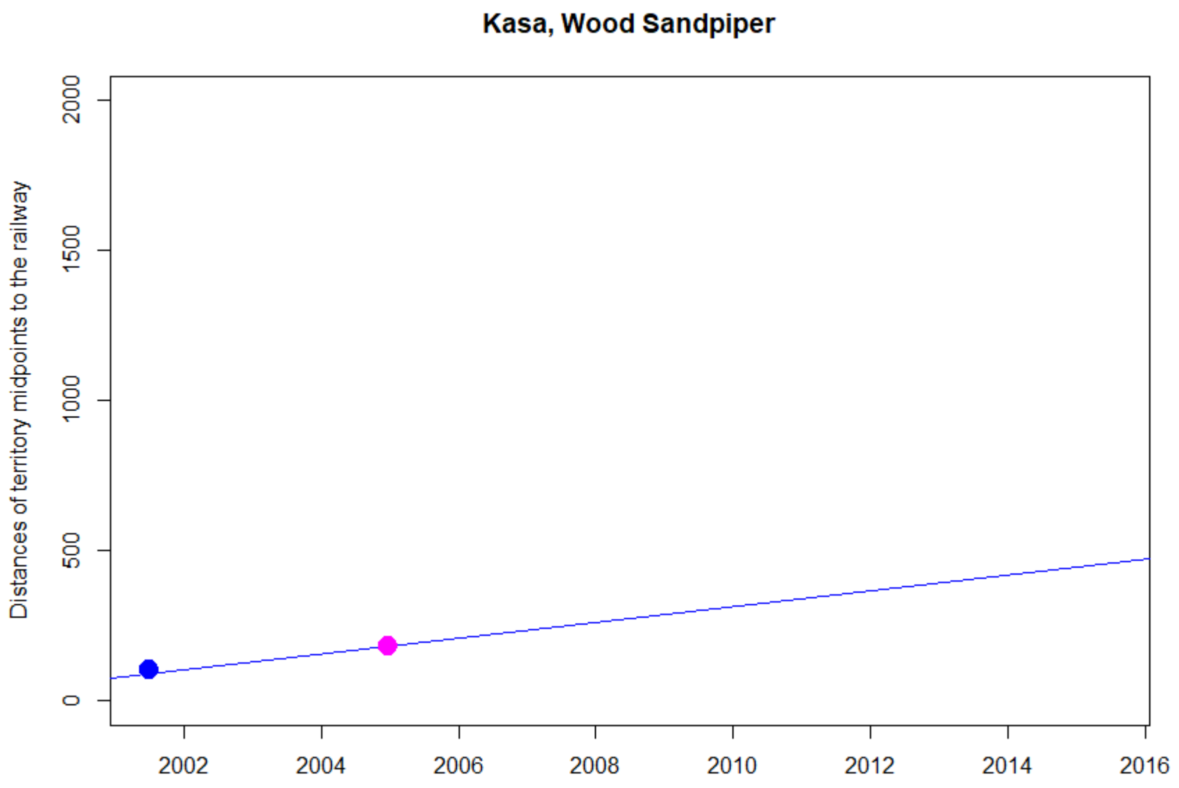
Kasa, Wood Sandpiper
Distances of territory midpoints to the railway
0
500
1000
1500
2000
2002
2004
2006
2008
2010
2012
2014
2016

NA

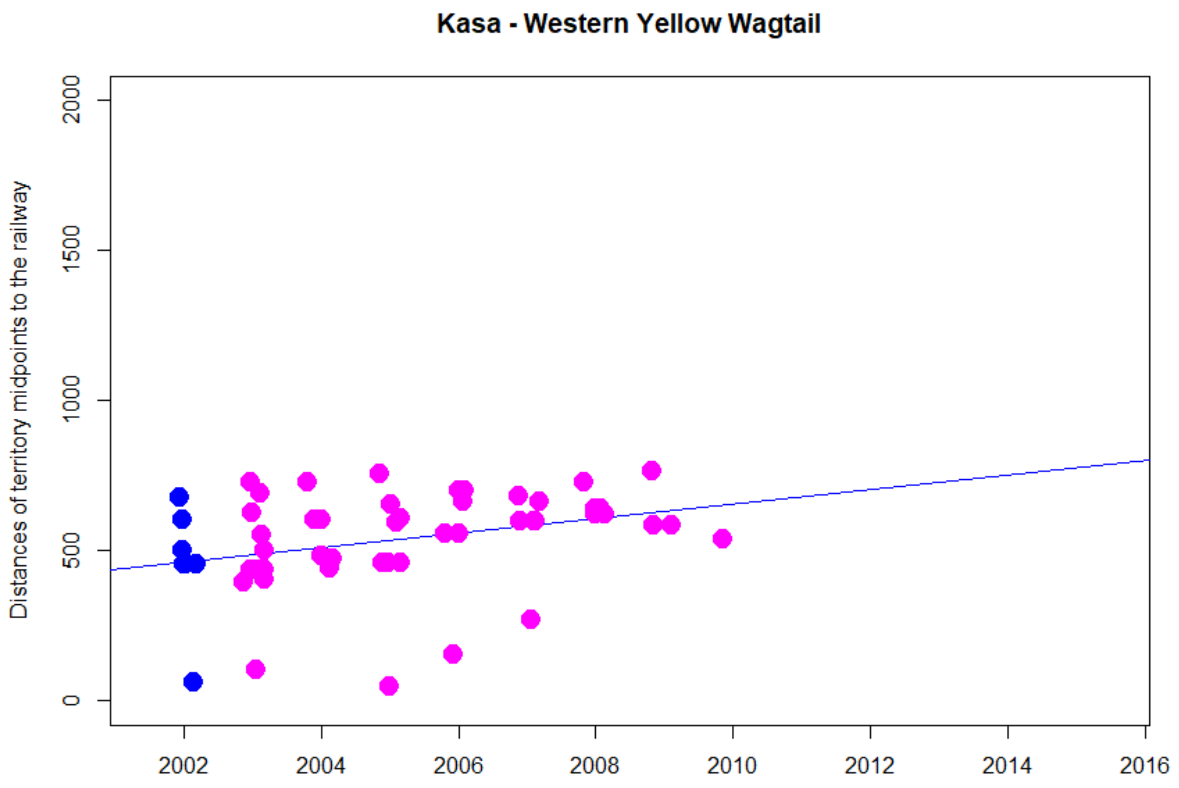


```
> cor.test(DNyland$Year,DNyland$NEAR_DIST_Botnia, method="kendall")

        Kendall's rank correlation tau

data:  DNyland$Year and DNyland$NEAR_DIST_Botnia
z = 2.8152, p-value = 0.004875
alternative hypothesis: true tau is not equal to 0
sample estimates:
      tau
0.2749943
```

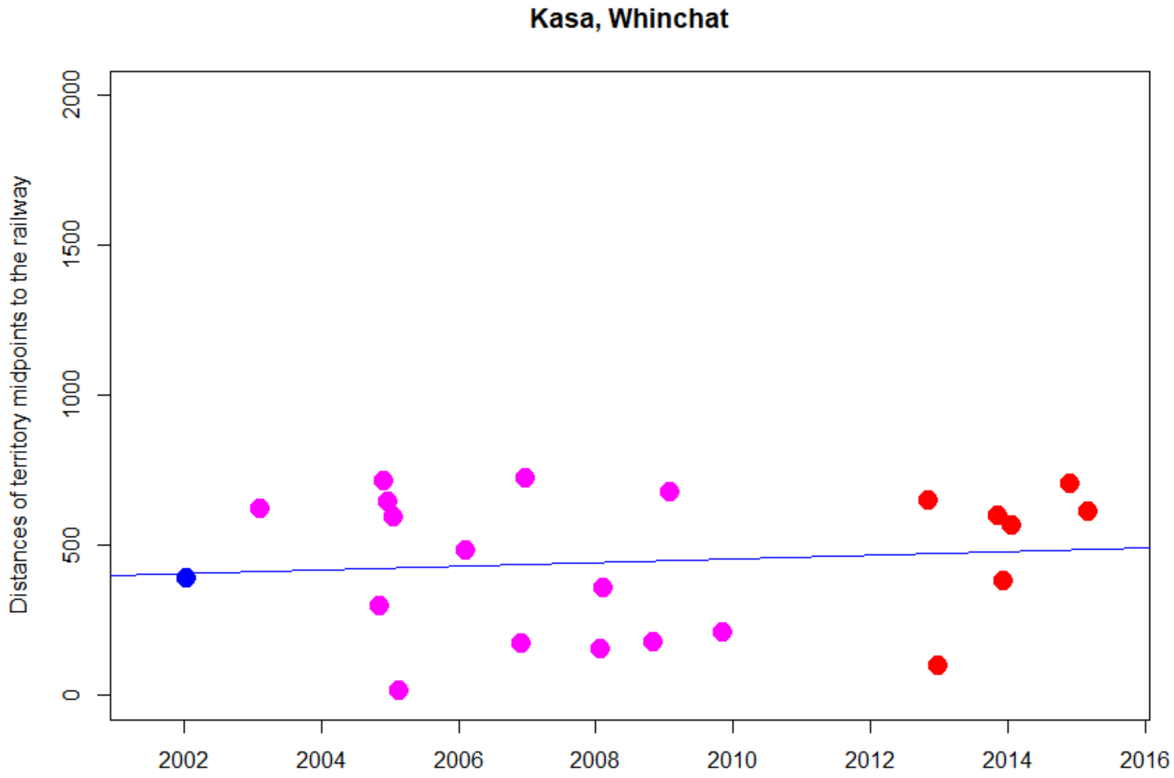


```
> cor.test(DNyland$Year,DNyland$NEAR_DIST_Botnia, method="kendall")

        Kendall's rank correlation tau

data:  DNyland$Year and DNyland$NEAR_DIST_Botnia
z = 0.37032, p-value = 0.7111
alternative hypothesis: true tau is not equal to 0
sample estimates:
       tau
0.05860674
```

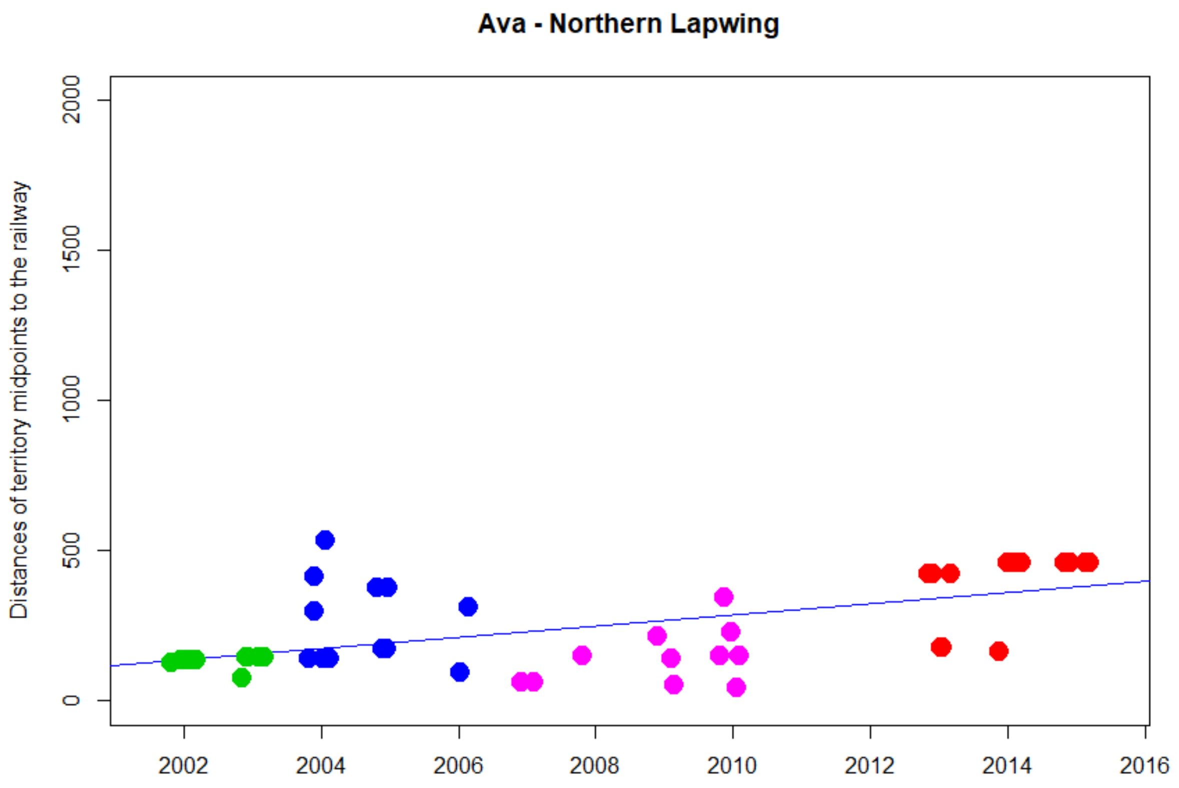


```
> cor.test(DNyland$Year,DNyland$NEAR_DIST_Botnia, method="kendall")

        Kendall's rank correlation tau

data:  DNyland$Year and DNyland$NEAR_DIST_Botnia
z = 4.7685, p-value = 1.856e-06
alternative hypothesis: true tau is not equal to 0
sample estimates:
      tau
0.4859241
```

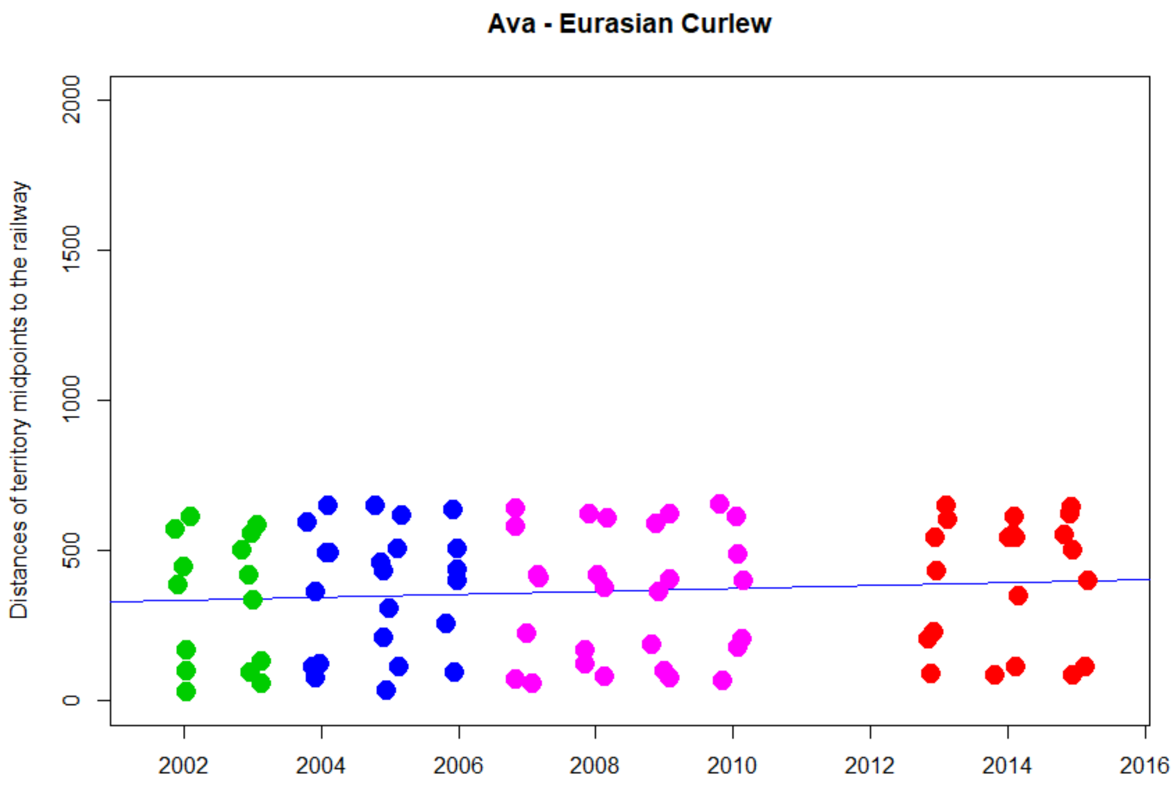


```
> cor.test(DNyland$Year,DNyland$NEAR_DIST_Botnia, method="kendall")

        Kendall's rank correlation tau

data:  DNyland$Year and DNyland$NEAR_DIST_Botnia
z = 0.89083, p-value = 0.373
alternative hypothesis: true tau is not equal to 0
sample estimates:
       tau
0.06722385
```

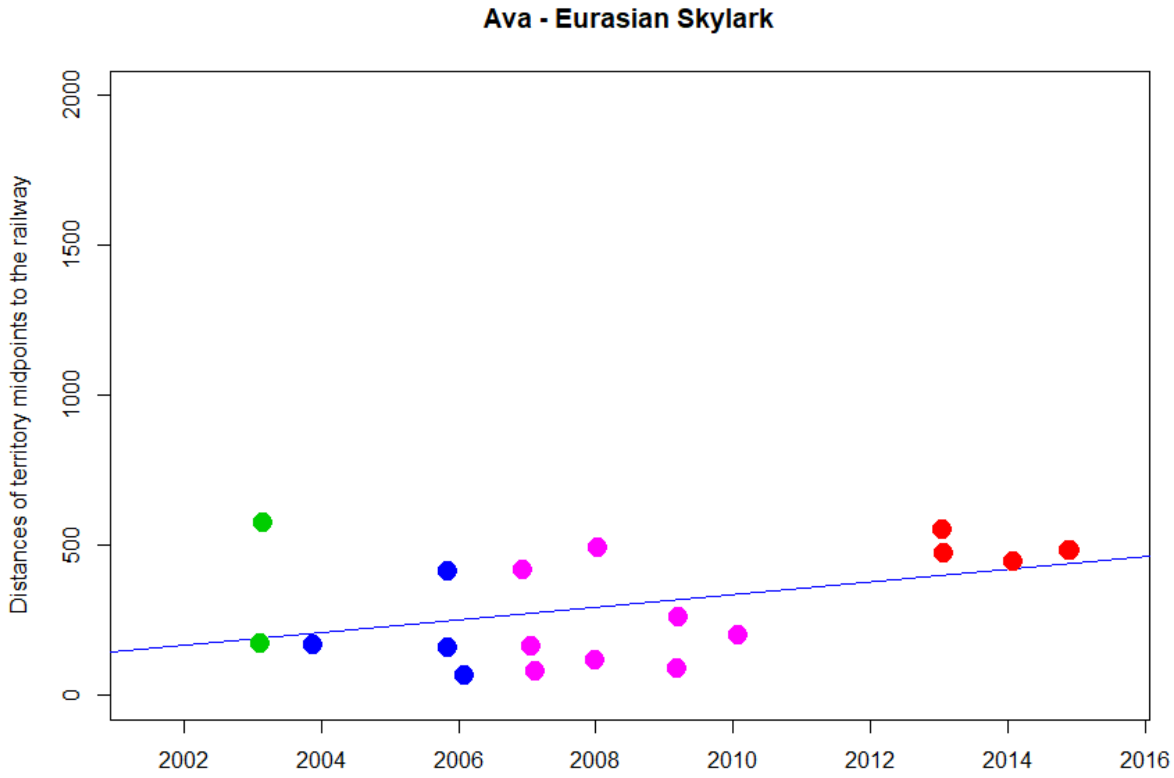


```
> cor.test(DNyland$Year,DNyland$NEAR_DIST_Botnia, method="kendall")

        Kendall's rank correlation tau

data:  DNyland$Year and DNyland$NEAR_DIST_Botnia
z = 1.552, p-value = 0.1207
alternative hypothesis: true tau is not equal to 0
sample estimates:
      tau
0.2667892
```

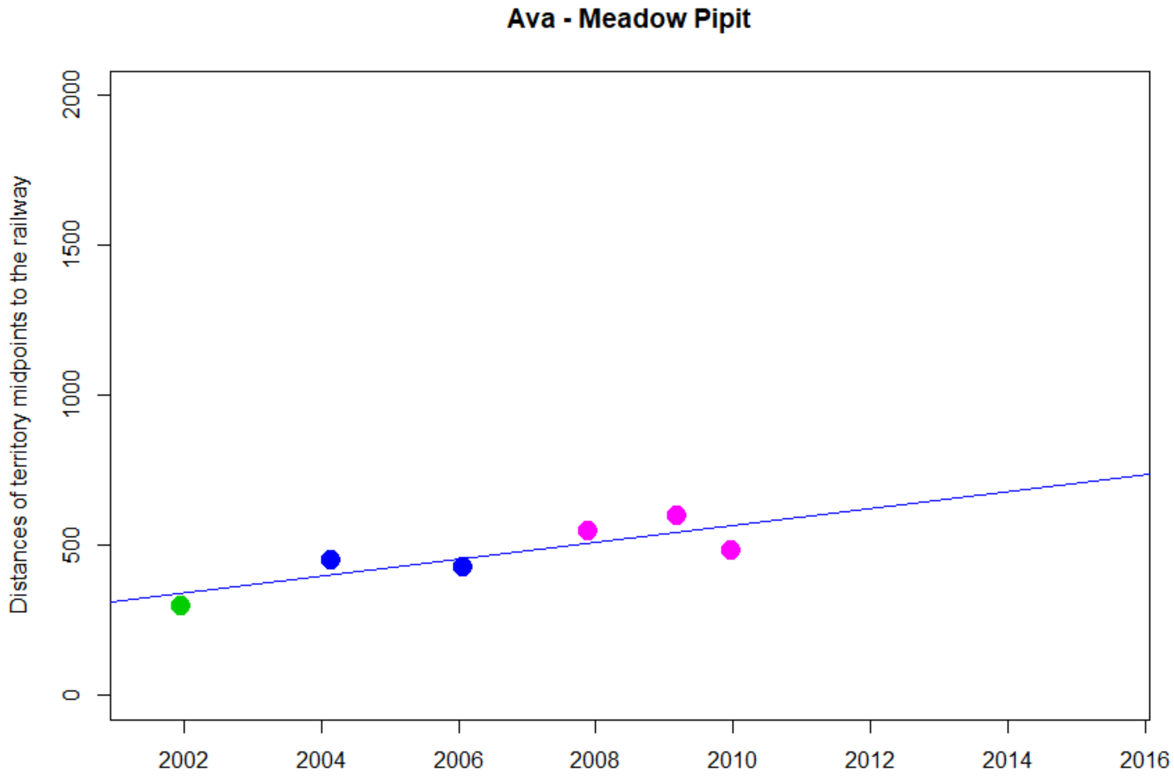


```
> cor.test(DNyland$Year,DNyland$NEAR_DIST_Botnia, method="kendall")

        Kendall's rank correlation tau

data:  DNyland$Year and DNyland$NEAR_DIST_Botnia
T = 12, p-value = 0.1361
alternative hypothesis: true tau is not equal to 0
sample estimates:
tau
0.6
```

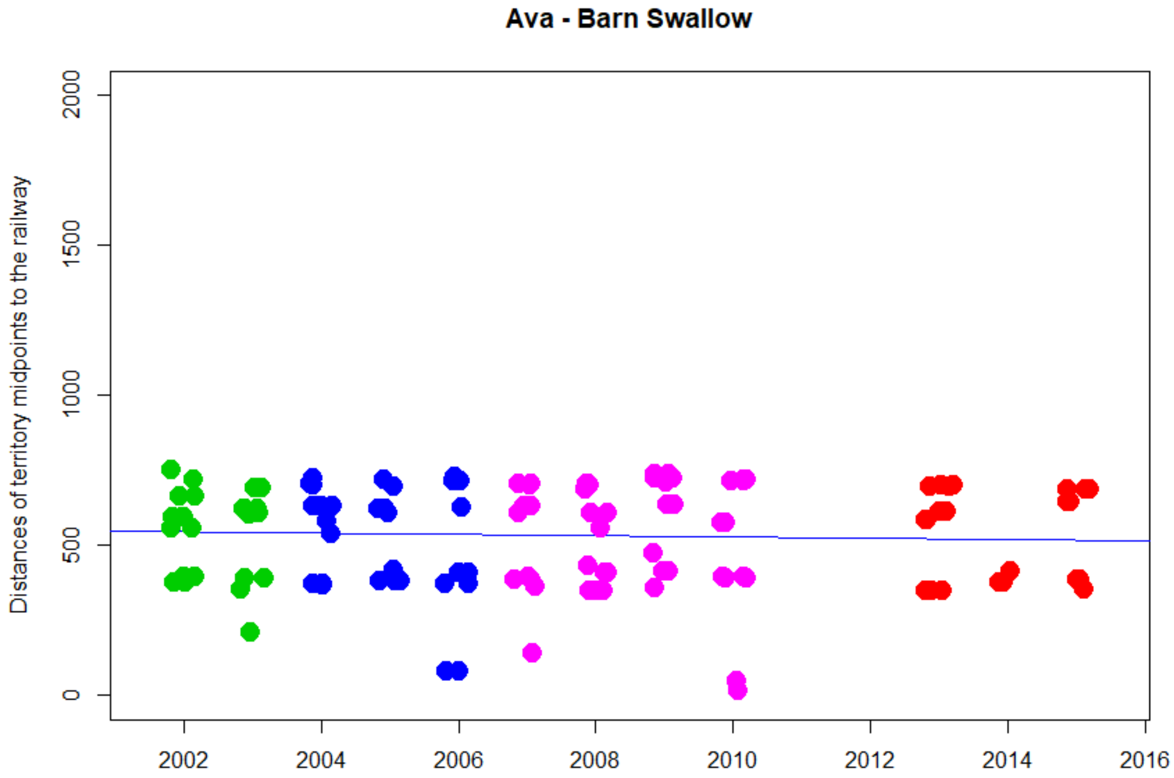


```
> cor.test(DNyland$Year,DNyland$NEAR_DIST_Botnia, method="kendall")

        Kendall's rank correlation tau

data:  DNyland$Year and DNyland$NEAR_DIST_Botnia
z = -0.28062, p-value = 0.779
alternative hypothesis: true tau is not equal to 0
sample estimates:
        tau
-0.01720921
```

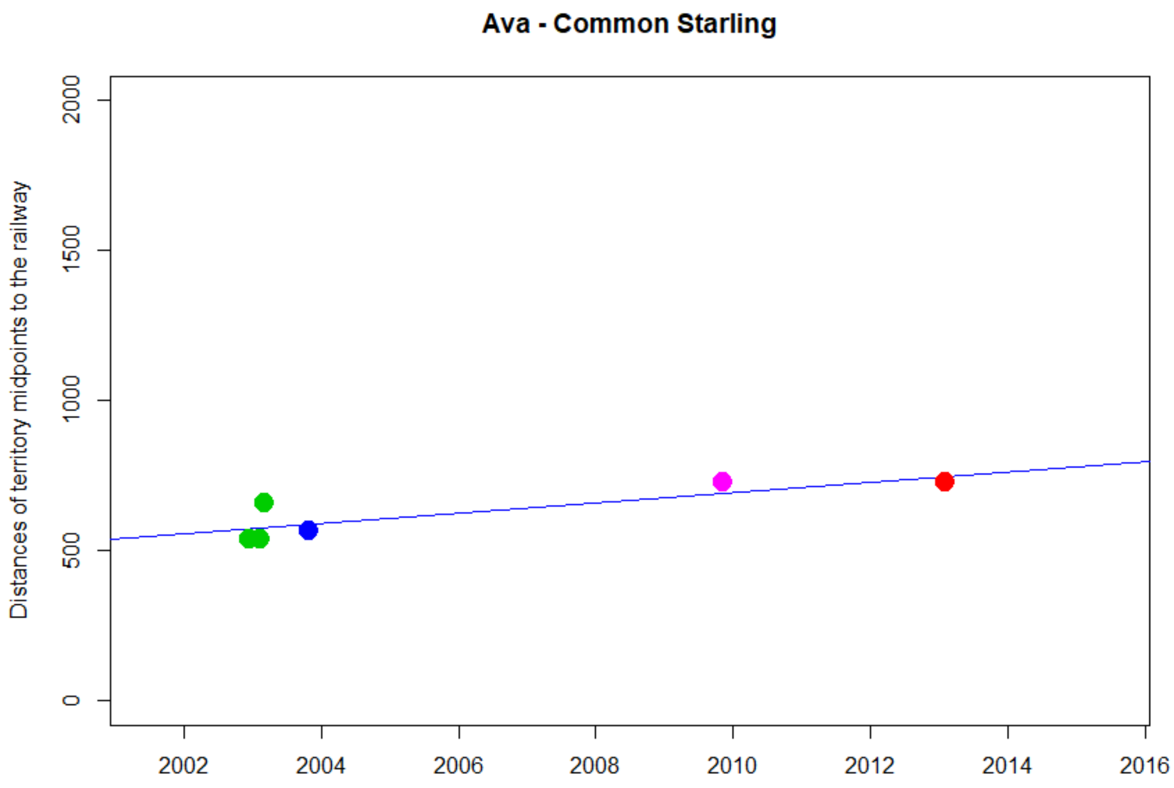


```
> cor.test(DNyland$Year,DNyland$NEAR_DIST_Botnia, method="kendall")

        Kendall's rank correlation tau

data:  DNyland$Year and DNyland$NEAR_DIST_Botnia
z = 1.6375, p-value = 0.1015
alternative hypothesis: true tau is not equal to 0
sample estimates:
      tau
0.6172134
```

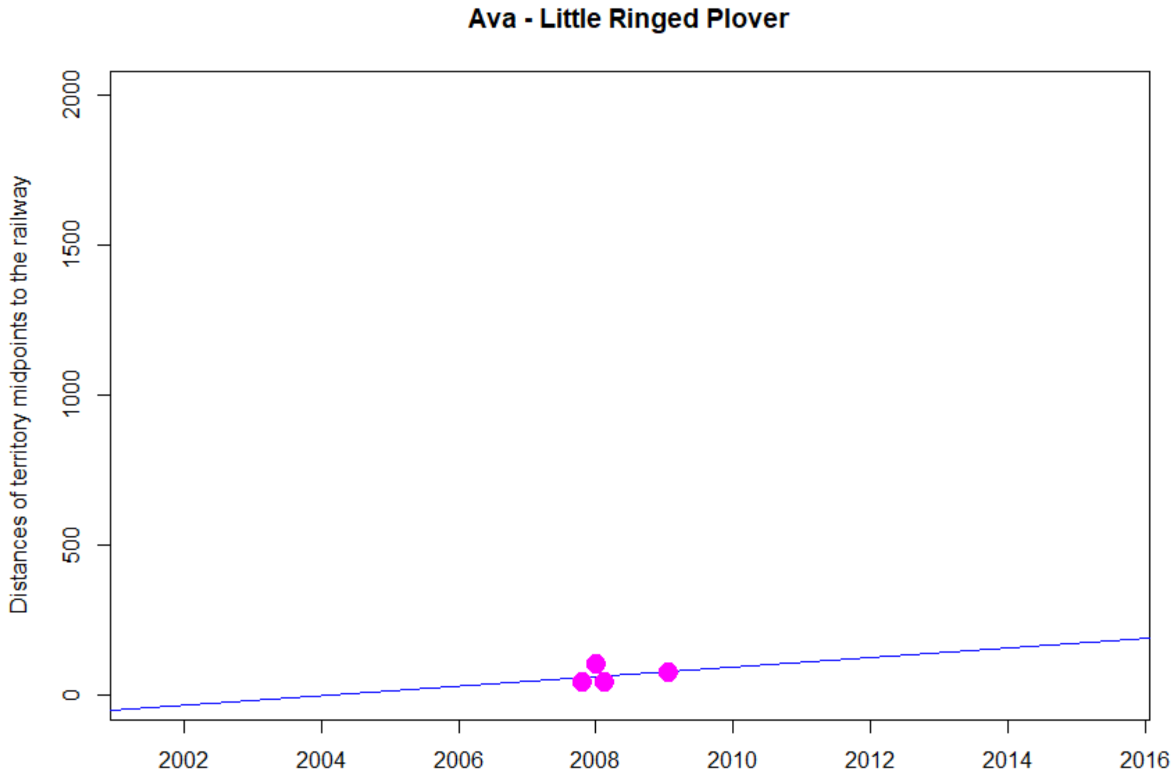


```
> cor.test(DNyland$Year,DNyland$NEAR_DIST_Botnia, method="kendall")

        Kendall's rank correlation tau

data:  DNyland$Year and DNyland$NEAR_DIST_Botnia
z = 0.4714, p-value = 0.6374
alternative hypothesis: true tau is not equal to 0
sample estimates:
      tau
0.2581989
```

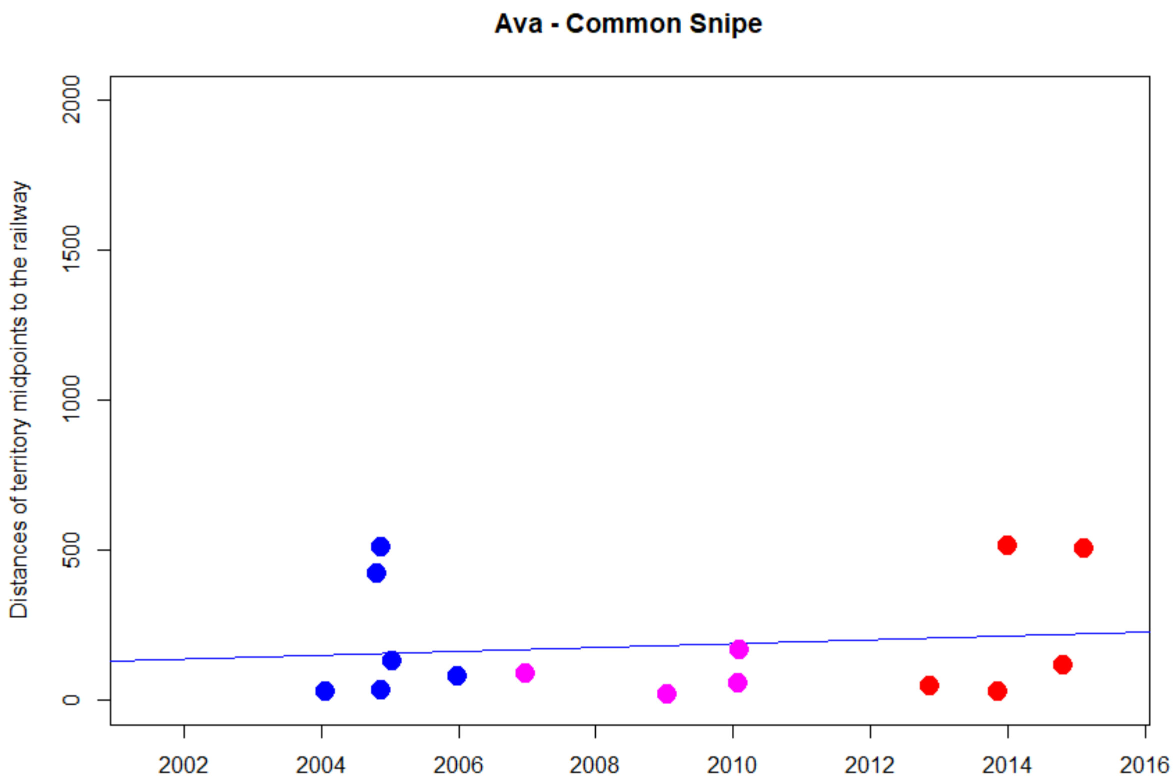


```
> cor.test(DNyland$Year,DNyland$NEAR_DIST_Botnia, method="kendall")

        Kendall's rank correlation tau

data:  DNyland$Year and DNyland$NEAR_DIST_Botnia
z = 0.5021, p-value = 0.6156
alternative hypothesis: true tau is not equal to 0
sample estimates:
       tau
0.09960238
```

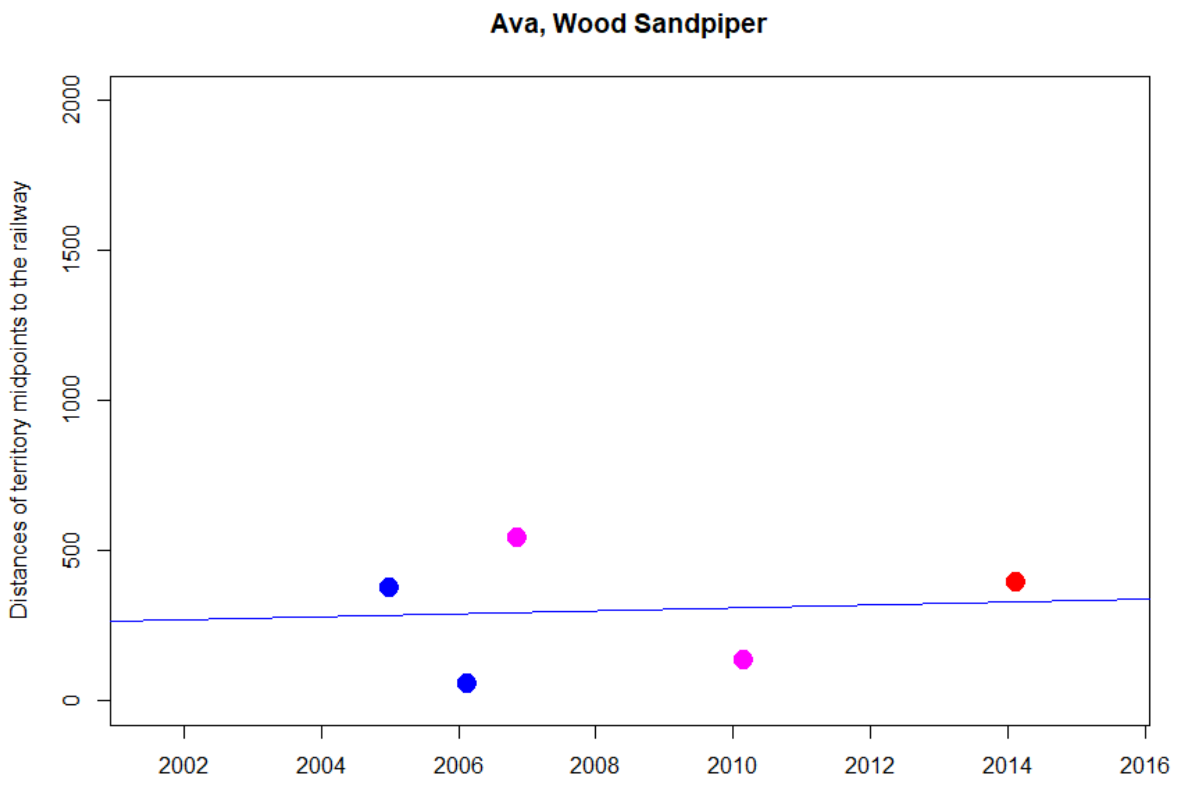


```
> cor.test(DNyland$Year,DNyland$NEAR_DIST_Botnia, method="kendall")

        Kendall's rank correlation tau

data:  DNyland$Year and DNyland$NEAR_DIST_Botnia
T = 6, p-value = 0.8167
alternative hypothesis: true tau is not equal to 0
sample estimates:
tau
0.2
```

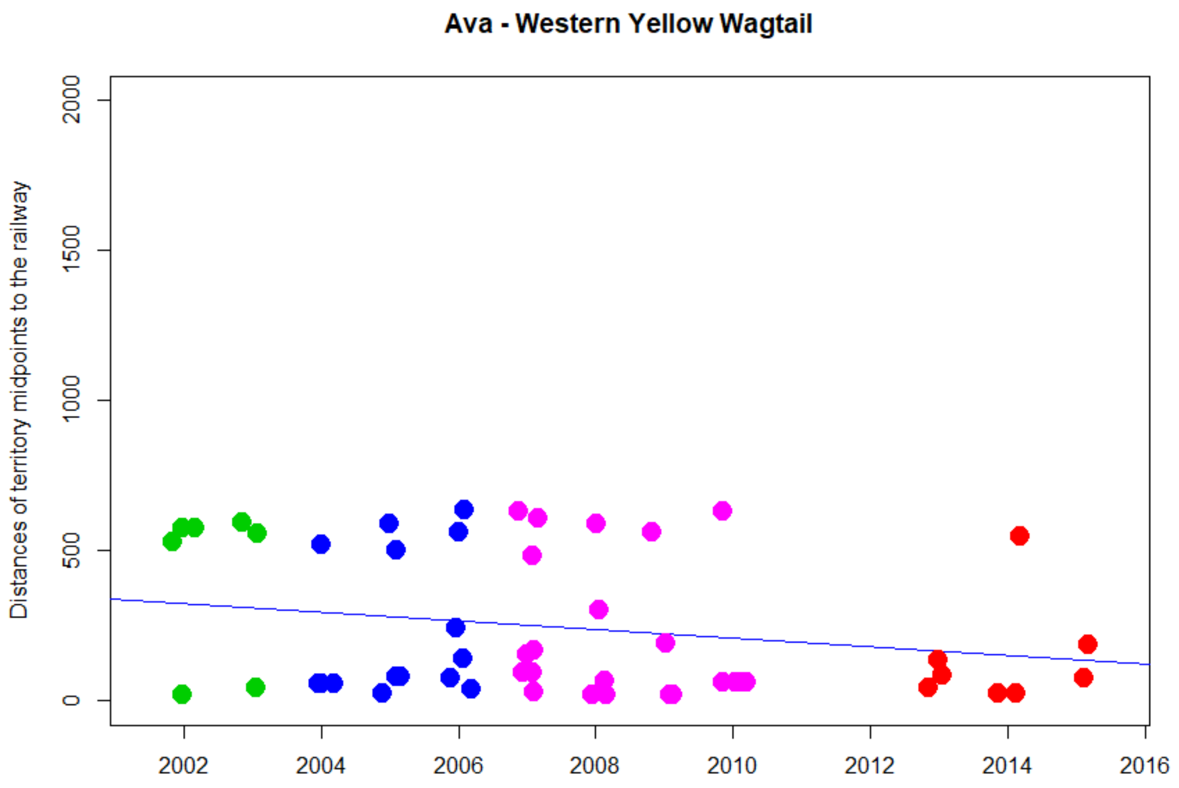


```
> cor.test(DNyland$Year,DNyland$NEAR_DIST_Botnia, method="kendall")

        Kendall's rank correlation tau

data:  DNyland$Year and DNyland$NEAR_DIST_Botnia
z = -1.1177, p-value = 0.2637
alternative hypothesis: true tau is not equal to 0
sample estimates:
       tau
-0.1080315
```

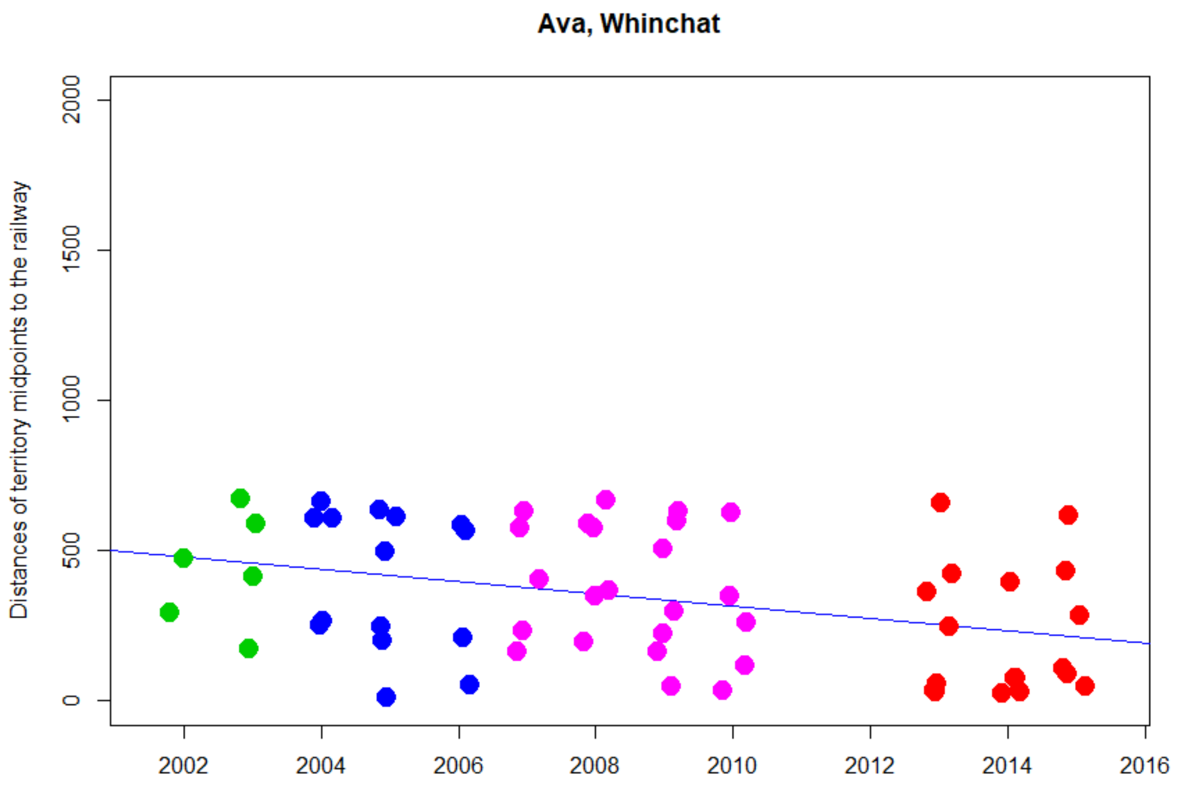


```
> cor.test(DNyland$Year,DNyland$NEAR_DIST_Botnia, method="kendall")

        Kendall's rank correlation tau

data:  DNyland$Year and DNyland$NEAR_DIST_Botnia
z = -2.7405, p-value = 0.006135
alternative hypothesis: true tau is not equal to 0
sample estimates:
       tau
-0.2468808
```

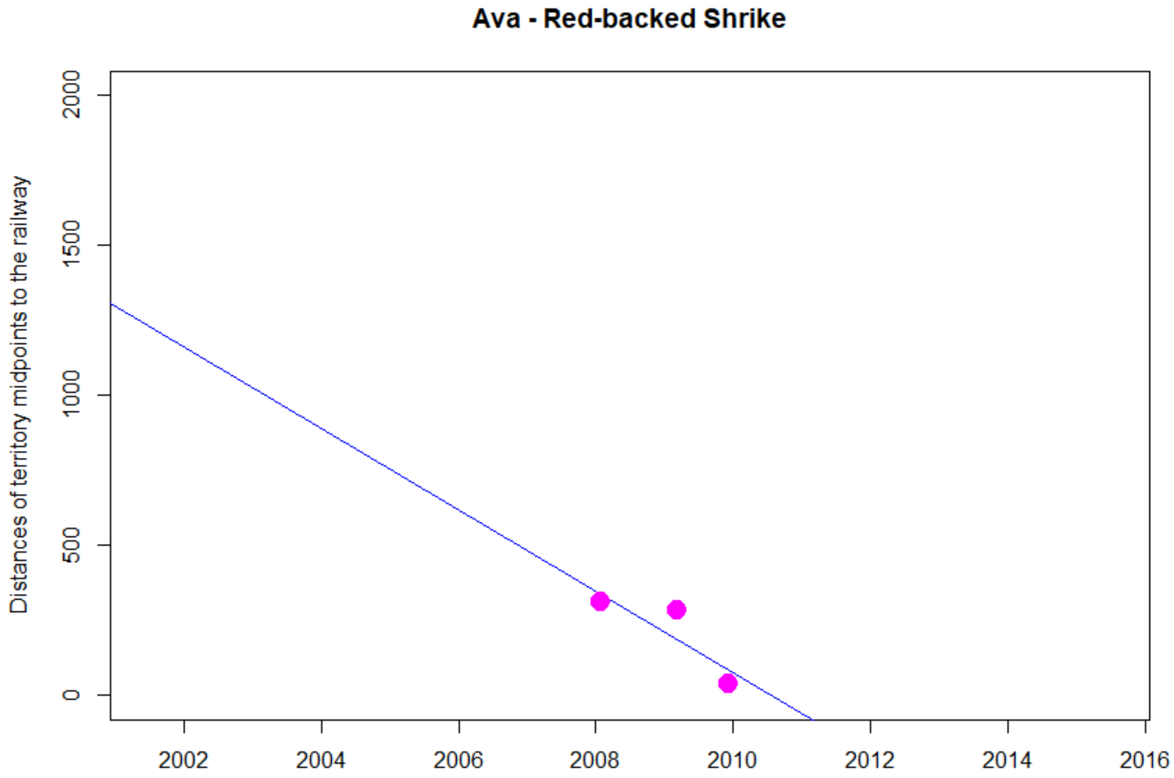
Ava - Red-backed Shrike
Distances of territory midpoints to the railway
0
500
1000
1500
2000
2002
2004
2006
2008
2010
2012
2014
2016

NA

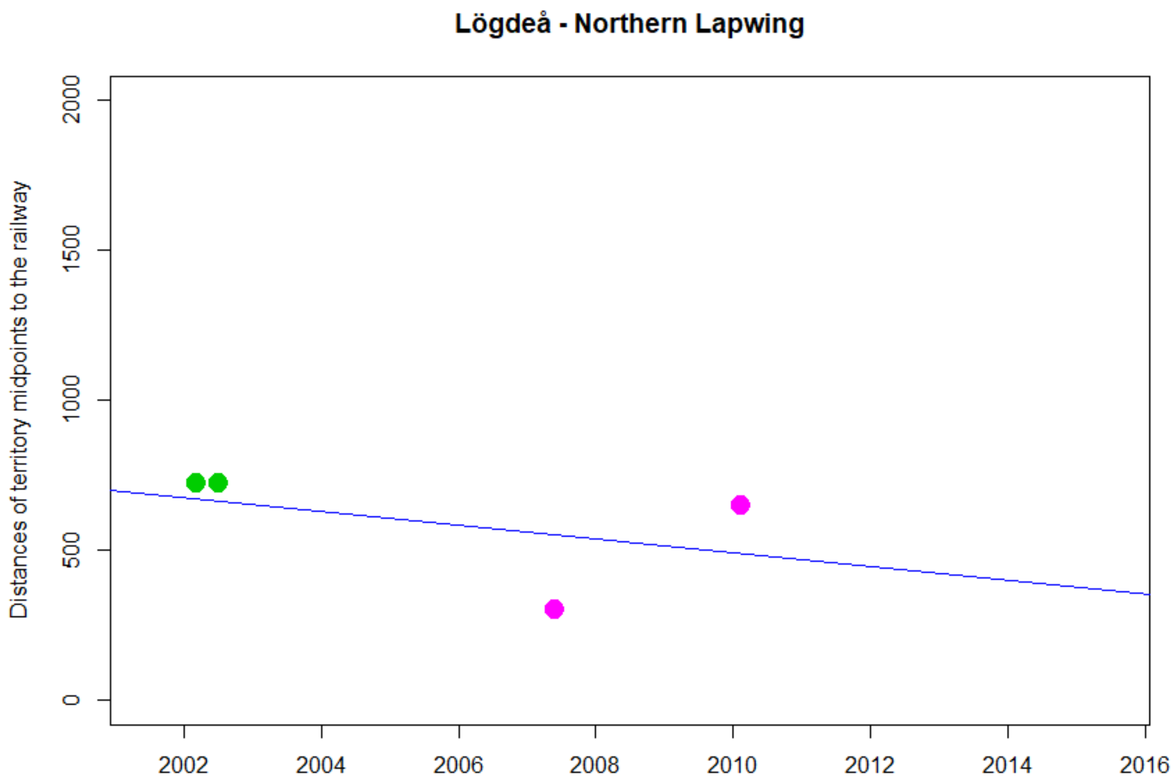


```
> cor.test(DNyland$Year,DNyland$NEAR_DIST_Botnia, method="kendall")

        Kendall's rank correlation tau

data:  DNyland$Year and DNyland$NEAR_DIST_Botnia
z = -1.1476, p-value = 0.2511
alternative hypothesis: true tau is not equal to 0
sample estimates:
 tau
-0.6
```

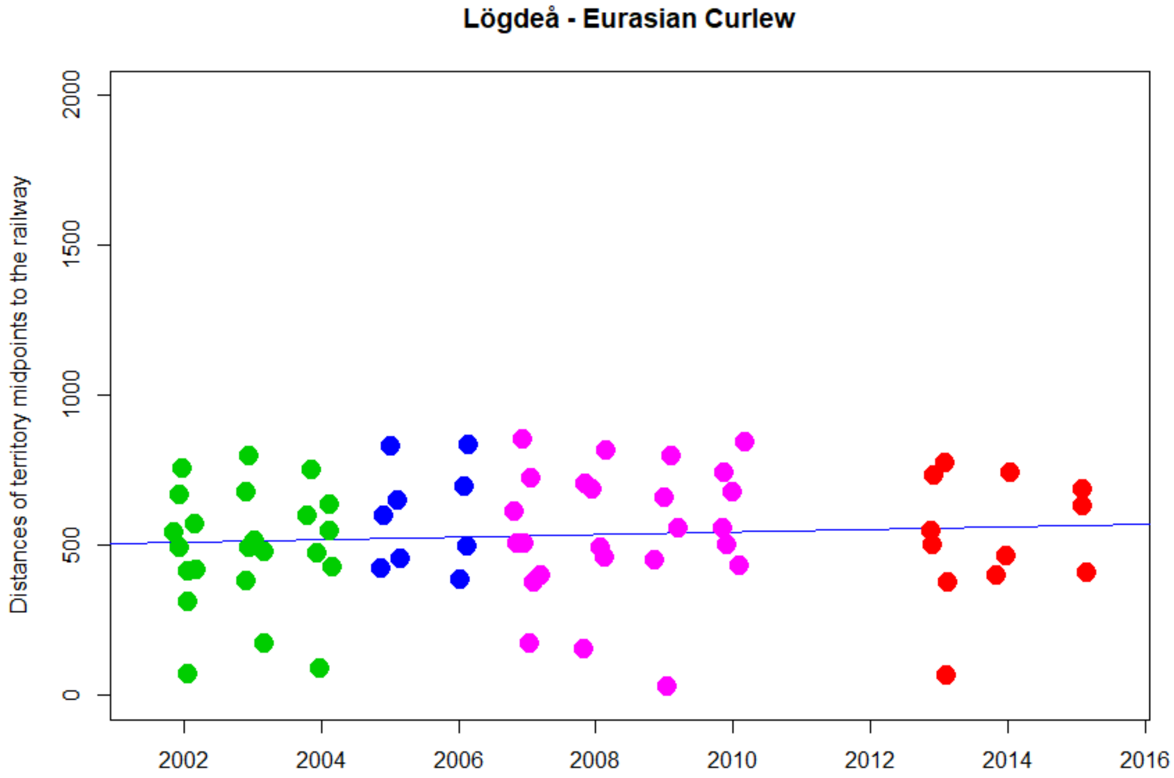


```
> cor.test(DNyland$Year,DNyland$NEAR_DIST_Botnia, method="kendall")

        Kendall's rank correlation tau

data:  DNyland$Year and DNyland$NEAR_DIST_Botnia
z = 0.92149, p-value = 0.3568
alternative hypothesis: true tau is not equal to 0
sample estimates:
       tau
0.07857439
```

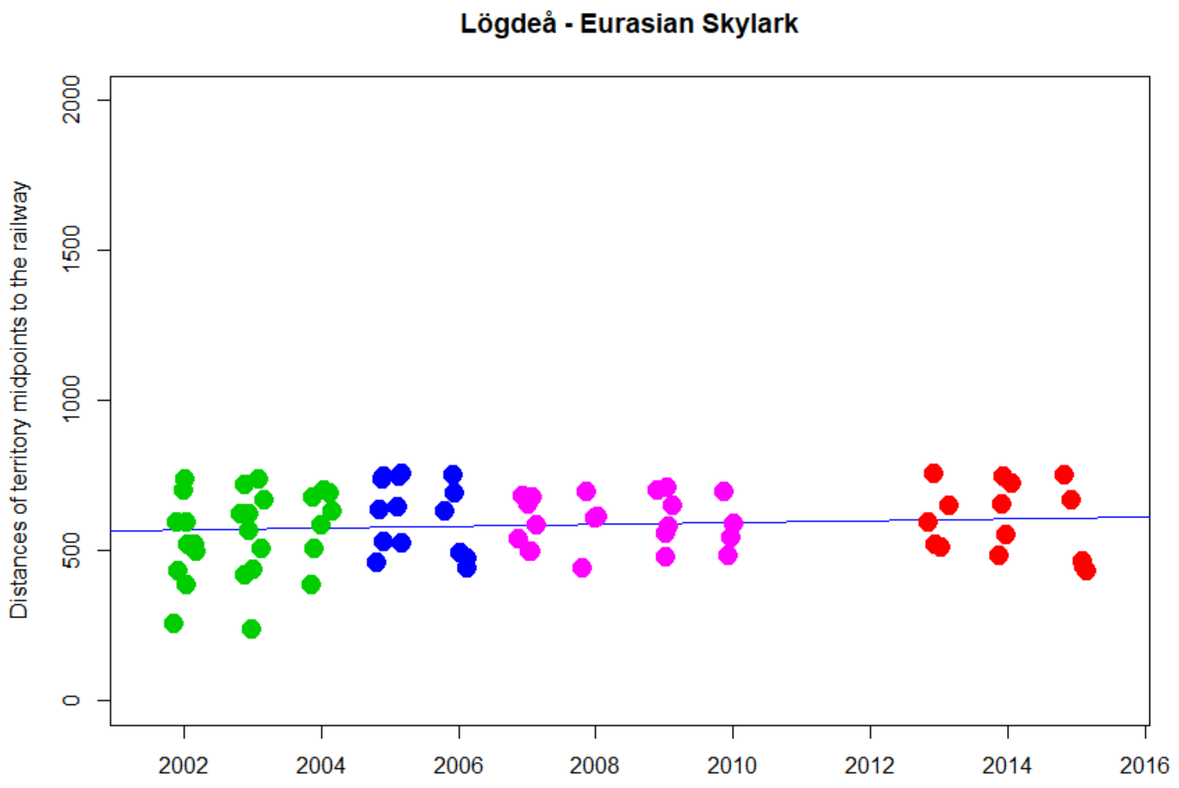


```
> cor.test(DNyland$Year,DNyland$NEAR_DIST_Botnia, method="kendall")

        Kendall's rank correlation tau

data:  DNyland$Year and DNyland$NEAR_DIST_Botnia
z = 0.9151, p-value = 0.3601
alternative hypothesis: true tau is not equal to 0
sample estimates:
       tau
0.07325113
```

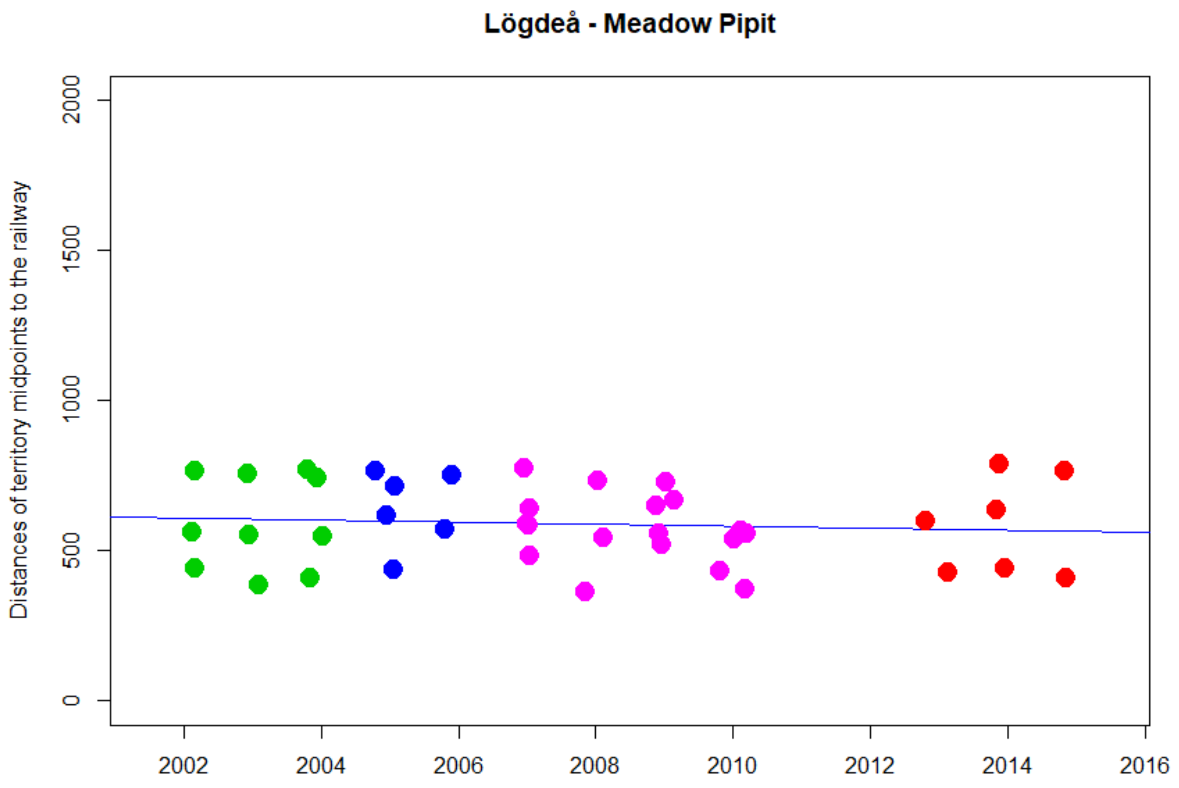


```
> cor.test(DNyland$Year,DNyland$NEAR_DIST_Botnia, method="kendall")

        Kendall's rank correlation tau

data:  DNyland$Year and DNyland$NEAR_DIST_Botnia
z = -0.80178, p-value = 0.4227
alternative hypothesis: true tau is not equal to 0
sample estimates:
       tau
-0.0897613
```

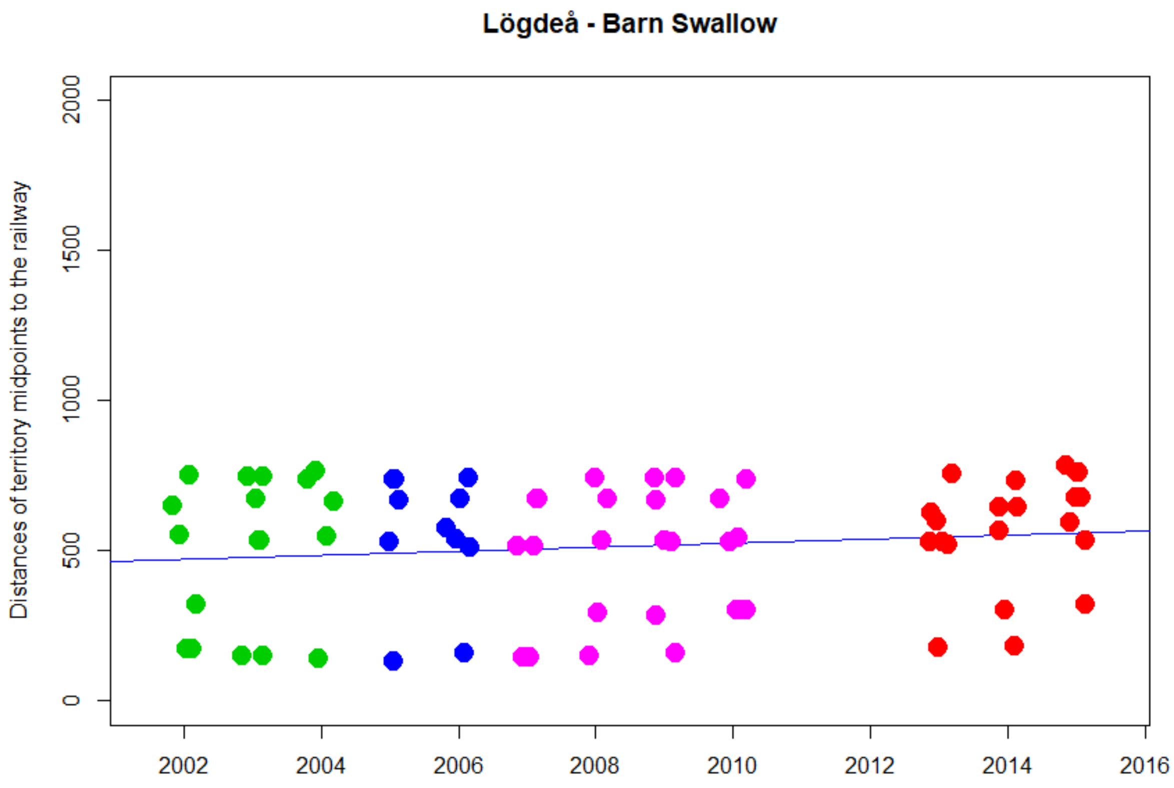


```
> cor.test(DNyland$Year,DNyland$NEAR_DIST_Botnia, method="kendall")

        Kendall's rank correlation tau

data:  DNyland$Year and DNyland$NEAR_DIST_Botnia
z = 1.0073, p-value = 0.3138
alternative hypothesis: true tau is not equal to 0
sample estimates:
       tau
0.08120214
```

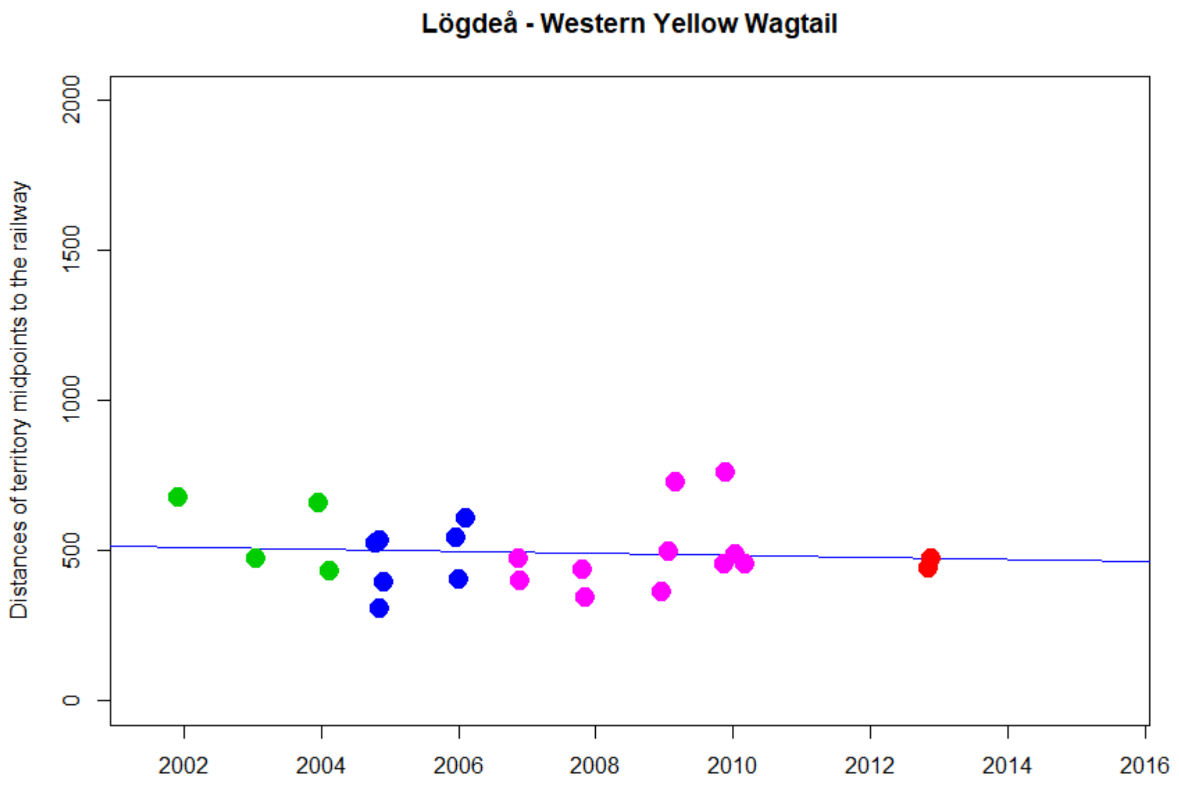


```
> cor.test(DNyland$Year,DNyland$NEAR_DIST_Botnia, method="kendall")

        Kendall's rank correlation tau

data:  DNyland$Year and DNyland$NEAR_DIST_Botnia
z = -0.3004, p-value = 0.7639
alternative hypothesis: true tau is not equal to 0
sample estimates:
        tau
-0.04540441
```

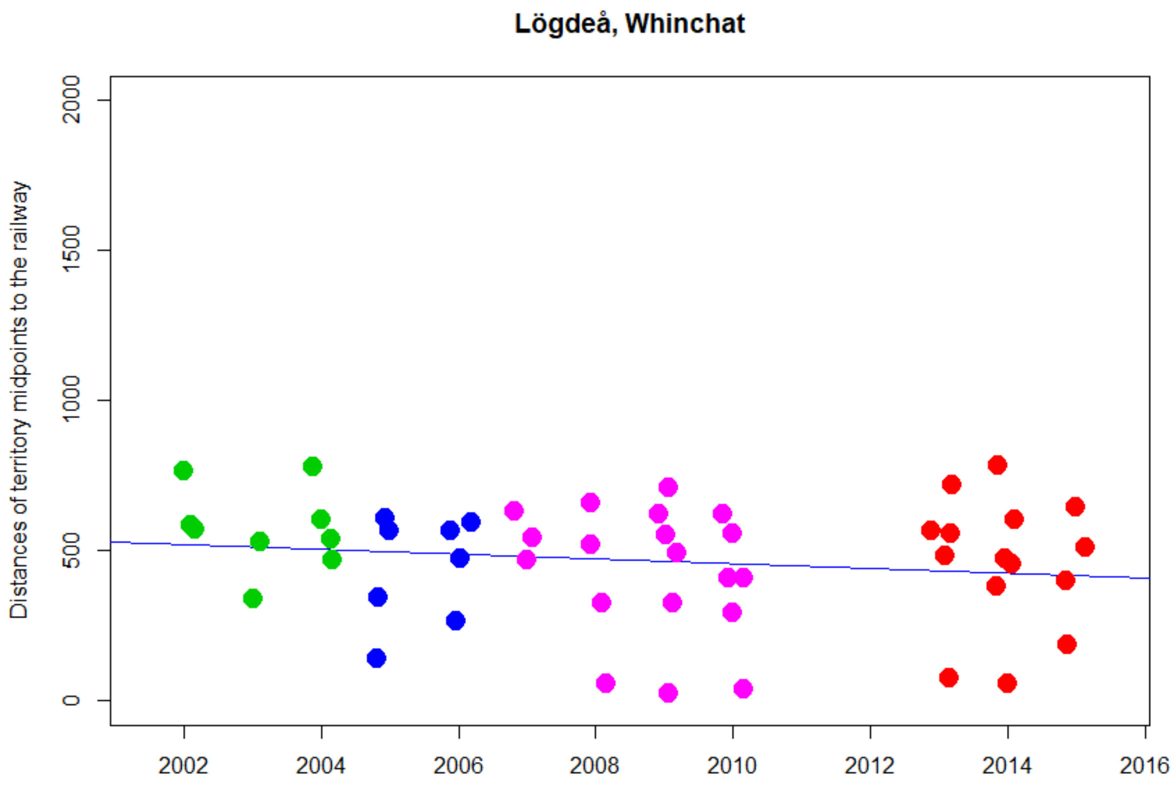


```
> cor.test(DNyland$Year,DNyland$NEAR_DIST_Botnia, method="kendall")

        Kendall's rank correlation tau

data:  DNyland$Year and DNyland$NEAR_DIST_Botnia
z = -1.134, p-value = 0.2568
alternative hypothesis: true tau is not equal to 0
sample estimates:
       tau
-0.1119271
```

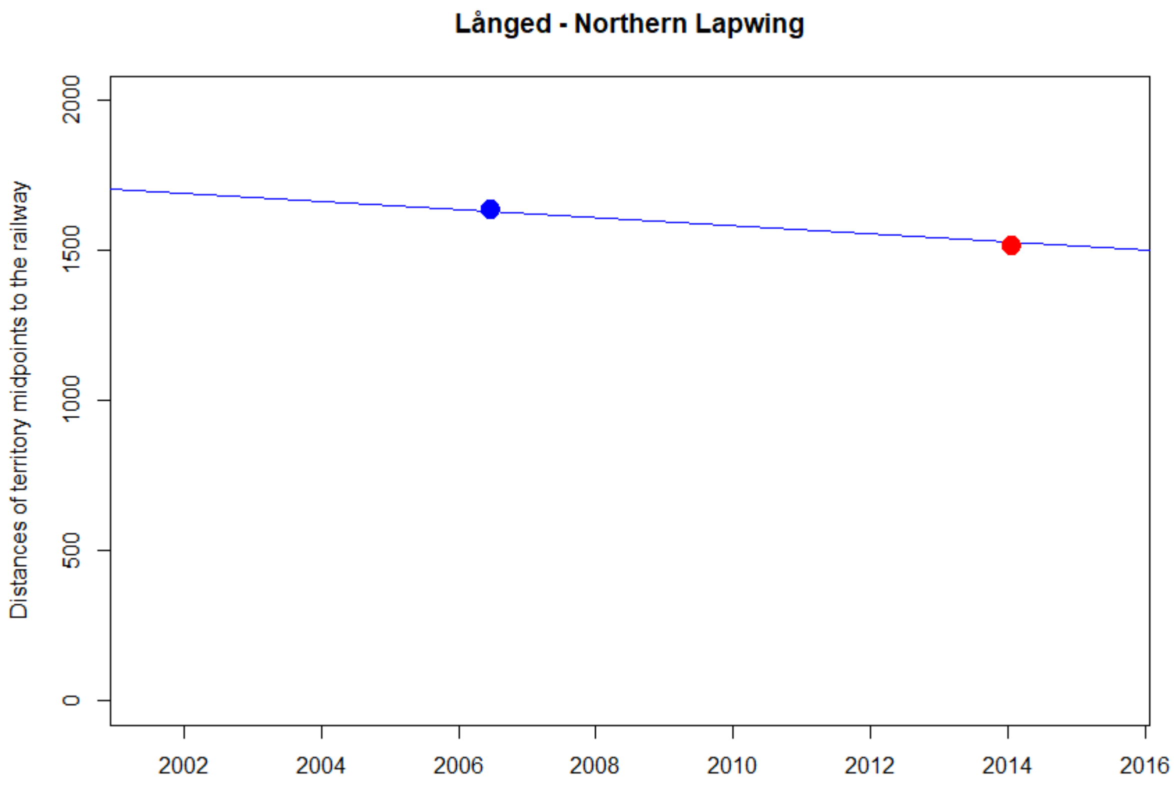
Långed - Northern Lapwing
Distances of territory midpoints to the railway
0
500
1000
1500
2000
2002
2004
2006
2008
2010
2012
2014
2016

NA

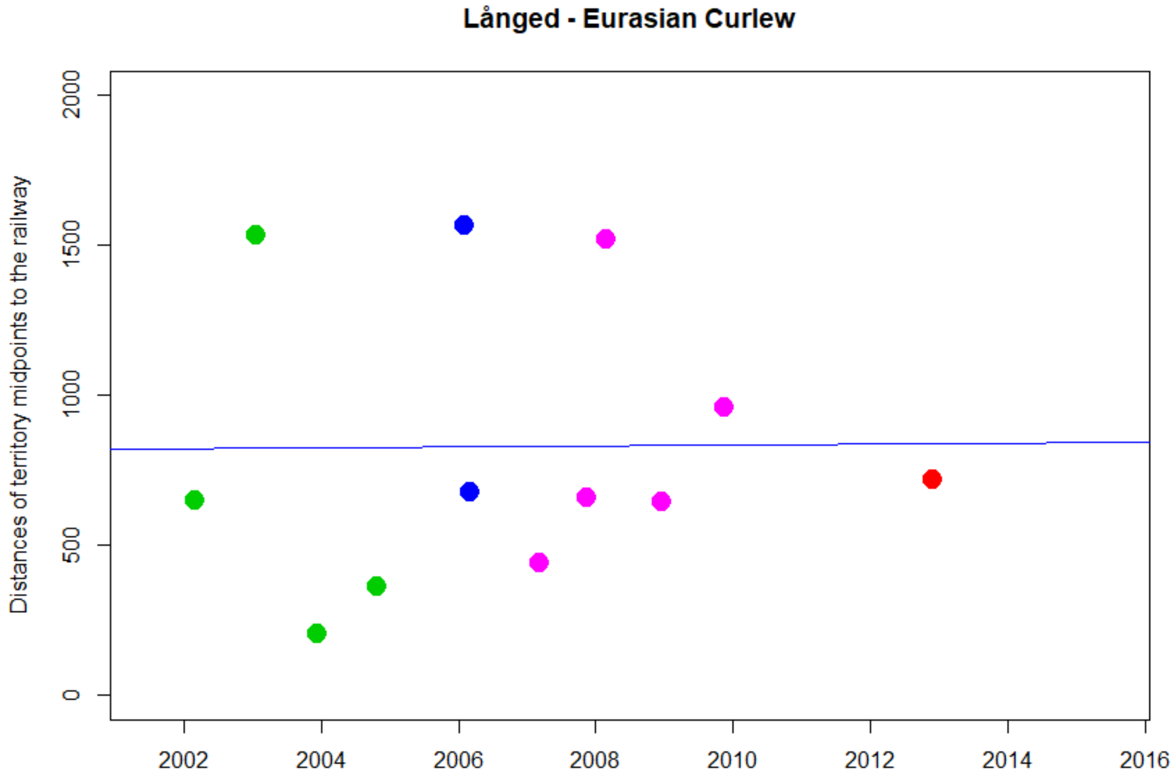


```
> cor.test(DNyland$Year,DNyland$NEAR_DIST_Botnia, method="kendall")

        Kendall's rank correlation tau

data:  DNyland$Year and DNyland$NEAR_DIST_Botnia
z = 0.68897, p-value = 0.4908
alternative hypothesis: true tau is not equal to 0
sample estimates:
      tau
0.1538644
```

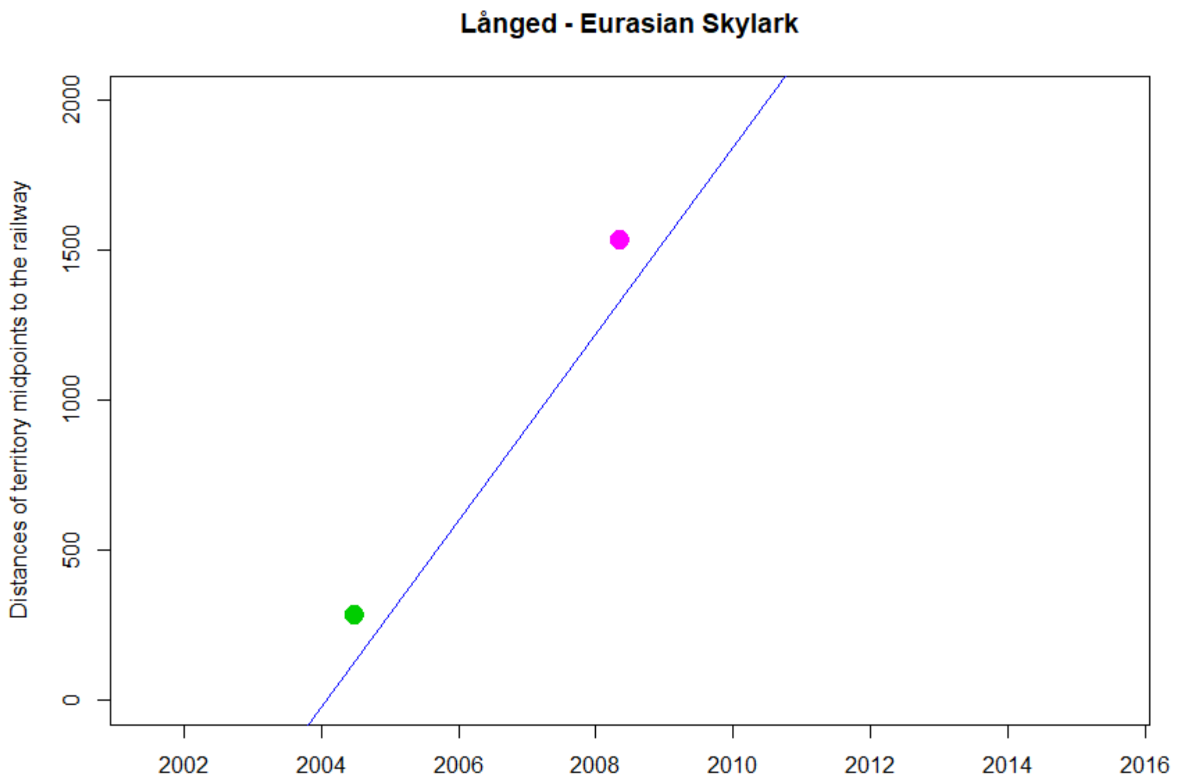
Långed - Eurasian Skylark
Distances of territory midpoints to the railway
0
500
1000
1500
2000
2002
2004
2006
2008
2010
2012
2014
2016

NA

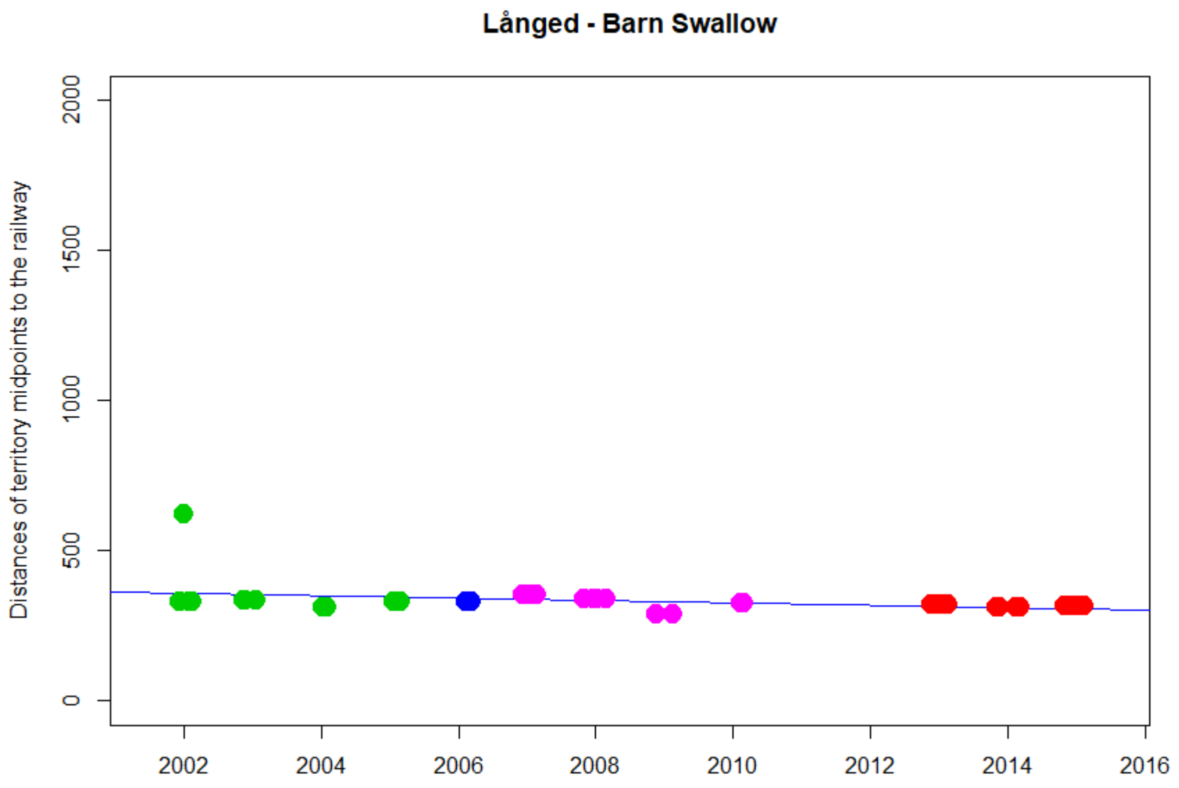


```
> cor.test(DNyland$Year,DNyland$NEAR_DIST_Botnia, method="kendall")

        Kendall's rank correlation tau

data:  DNyland$Year and DNyland$NEAR_DIST_Botnia
z = -3.7094, p-value = 0.0002077
alternative hypothesis: true tau is not equal to 0
sample estimates:
       tau
-0.4122664
```

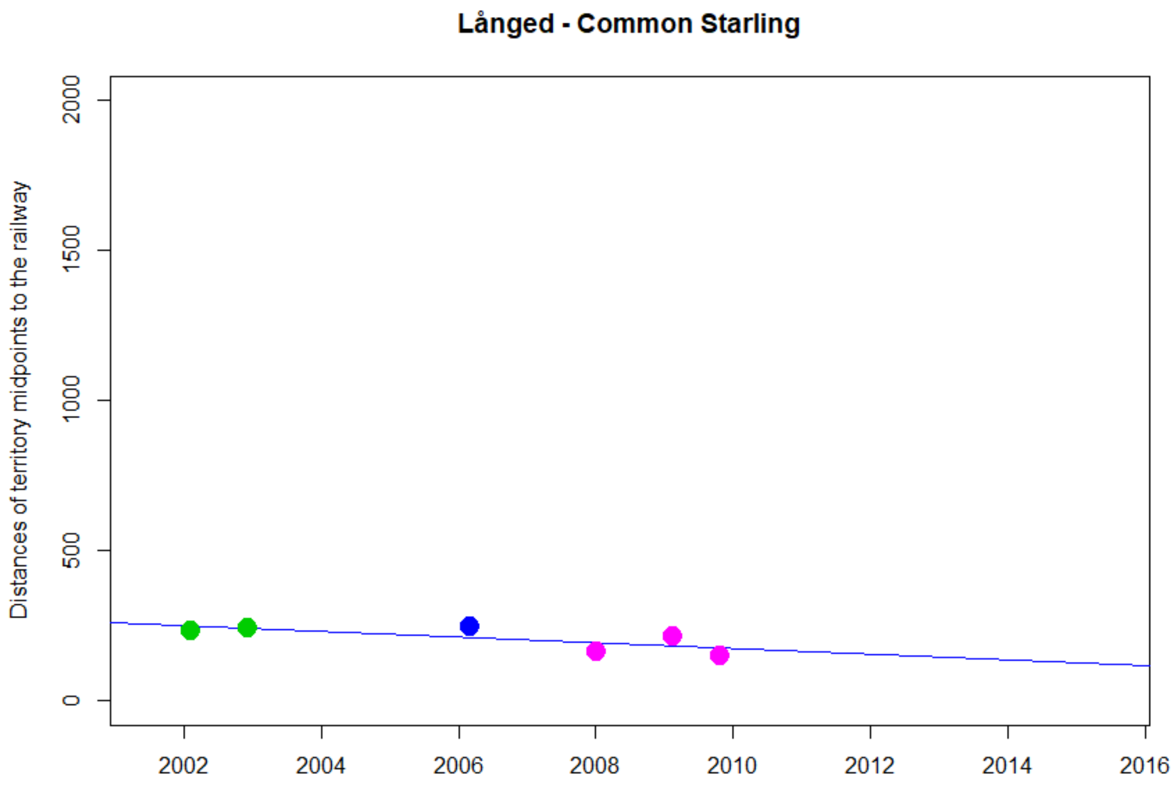


```
> cor.test(DNyland$Year,DNyland$NEAR_DIST_Botnia, method="kendall")

        Kendall's rank correlation tau

data:  DNyland$Year and DNyland$NEAR_DIST_Botnia
T = 4, p-value = 0.2722
alternative hypothesis: true tau is not equal to 0
sample estimates:
       tau 
-0.4666667 
```

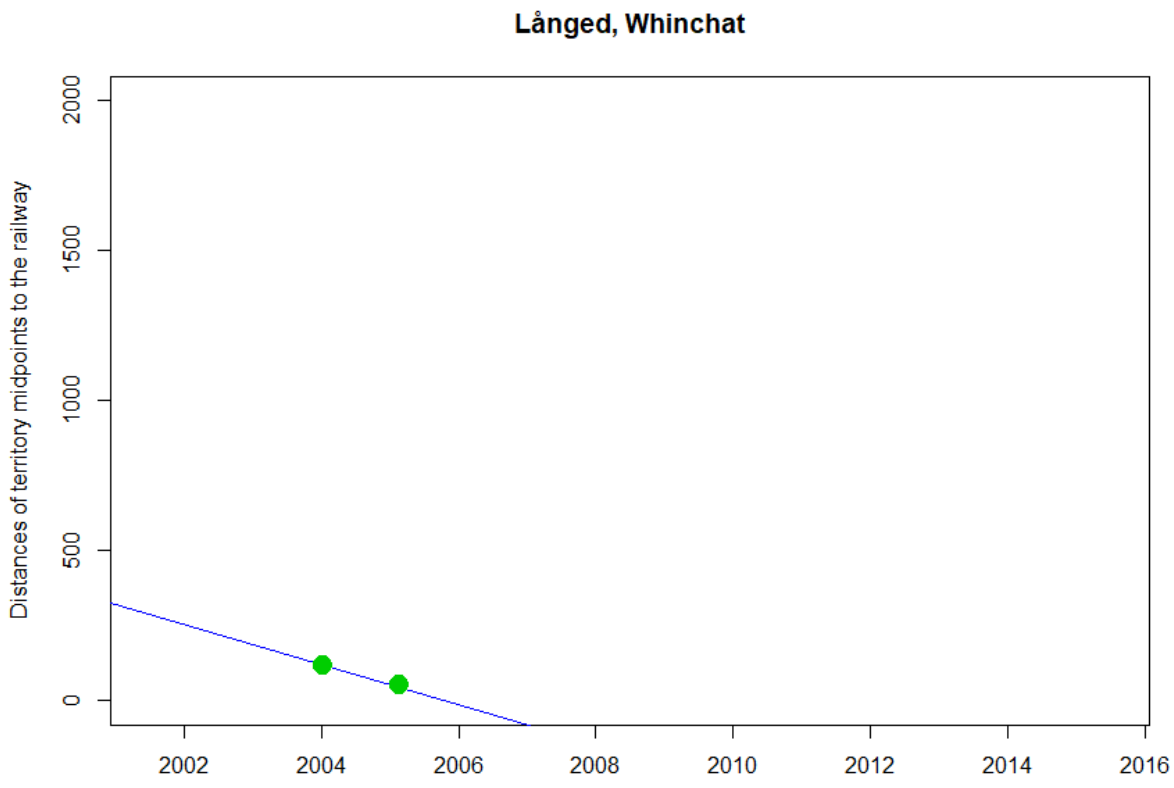
Långed, Whinchat
Distances of territory midpoints to the railway
0
500
1000
1500
2000
2002
2004
2006
2008
2010
2012
2014
2016

NA

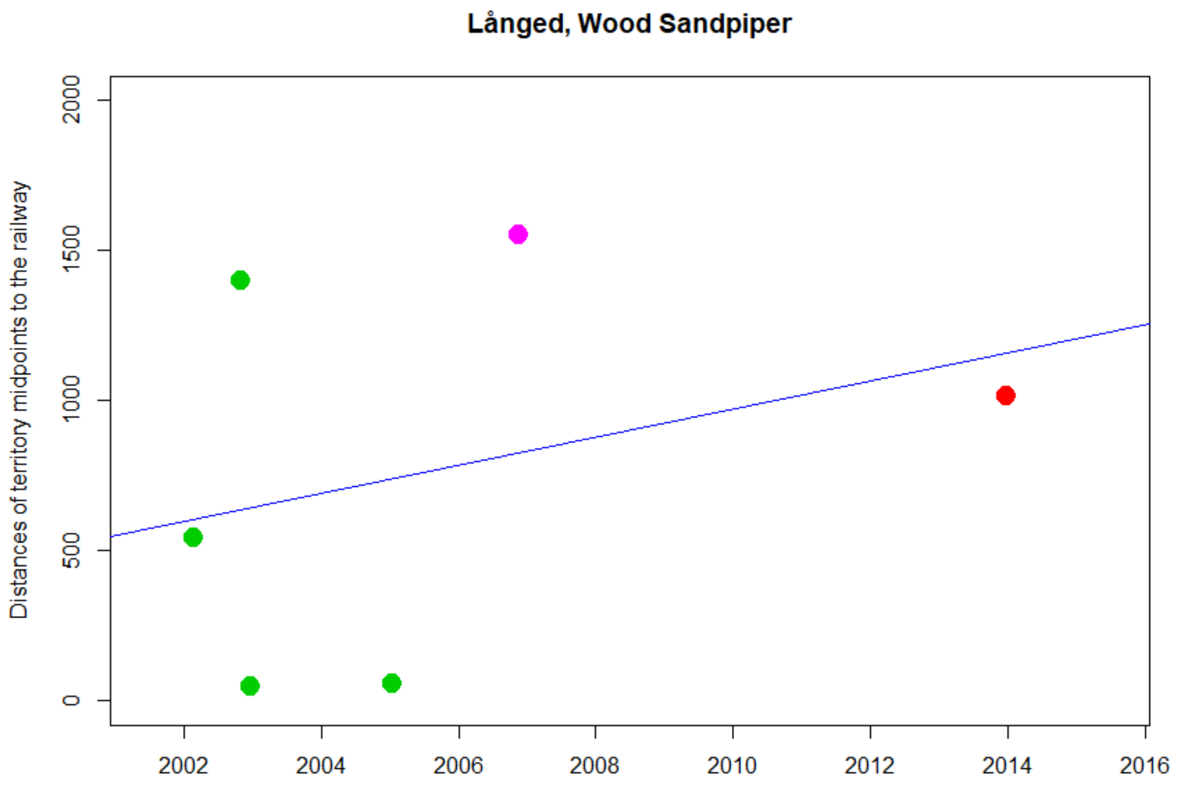


```
> cor.test(DNyland$Year,DNyland$NEAR_DIST_Botnia, method="kendall")

        Kendall's rank correlation tau

data:  DNyland$Year and DNyland$NEAR_DIST_Botnia
z = 0.76509, p-value = 0.4442
alternative hypothesis: true tau is not equal to 0
sample estimates:
      tau 
0.2760262 
```

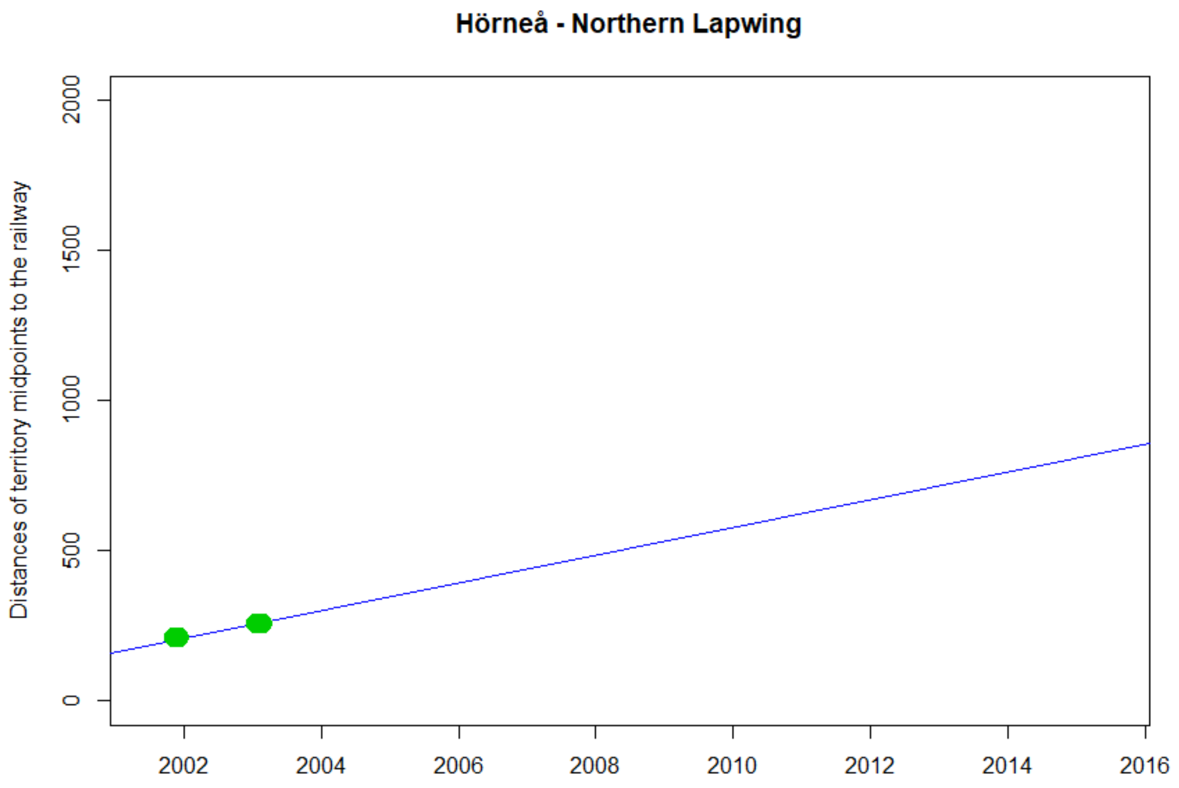
Hörneå - Northern Lapwing
Distances of territory midpoints to the railway
0
500
1000
1500
2000
2002
2004
2006
2008
2010
2012
2014
2016

NA

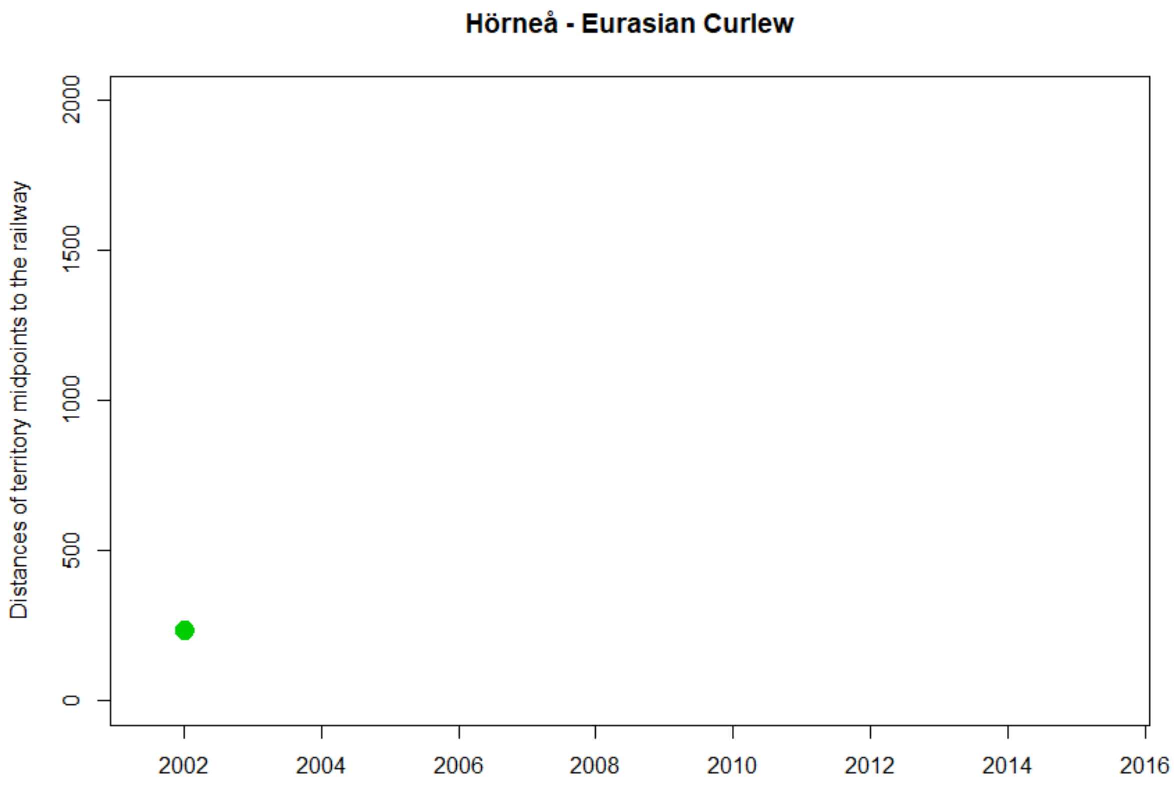
Hörneå - Eurasian Curlew
Distances of territory midpoints to the railway
0
500
1000
1500
2000
2002
2004
2006
2008
2010
2012
2014
2016

NA

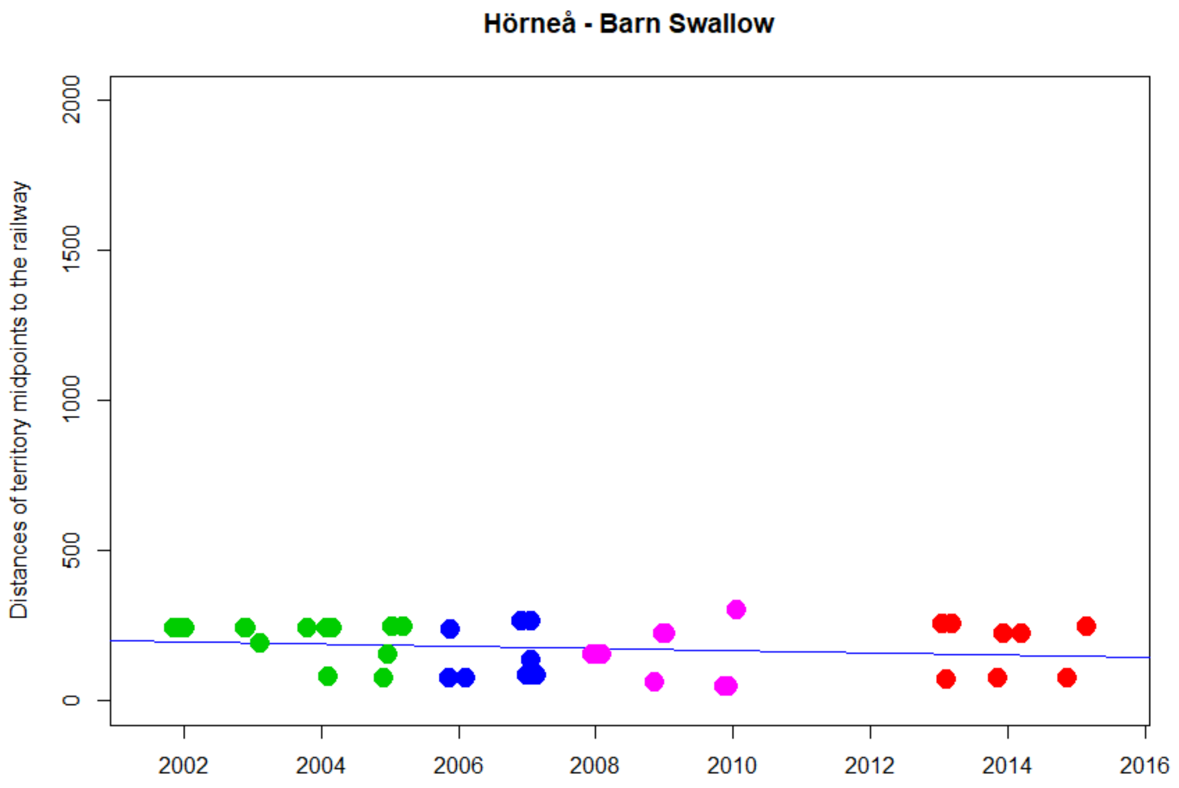


```
> cor.test(DNyland$Year,DNyland$NEAR_DIST_Botnia, method="kendall")

        Kendall's rank correlation tau

data:  DNyland$Year and DNyland$NEAR_DIST_Botnia
z = -1.2902, p-value = 0.197
alternative hypothesis: true tau is not equal to 0
sample estimates:
       tau
-0.1478837
```

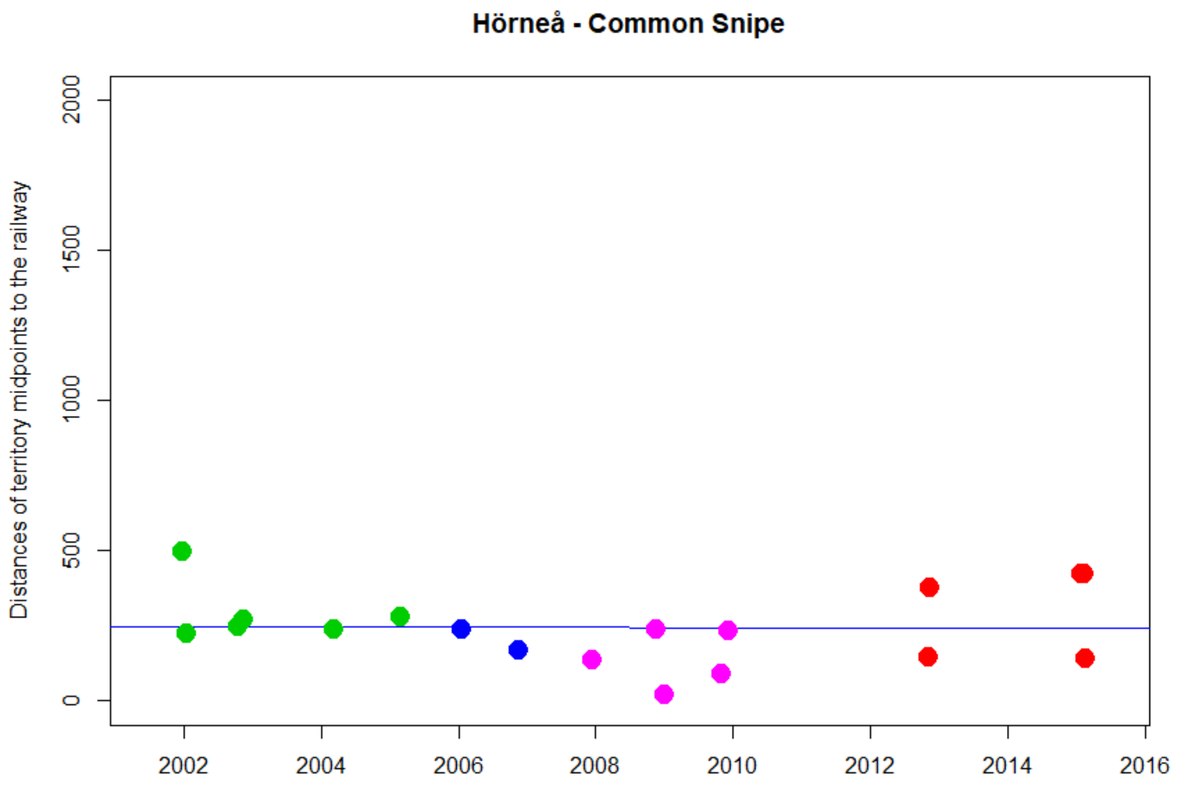


```
> cor.test(DNyland$Year,DNyland$NEAR_DIST_Botnia, method="kendall")

        Kendall's rank correlation tau

data:  DNyland$Year and DNyland$NEAR_DIST_Botnia
z = -0.64841, p-value = 0.5167
alternative hypothesis: true tau is not equal to 0
sample estimates:
       tau
-0.1145099
```

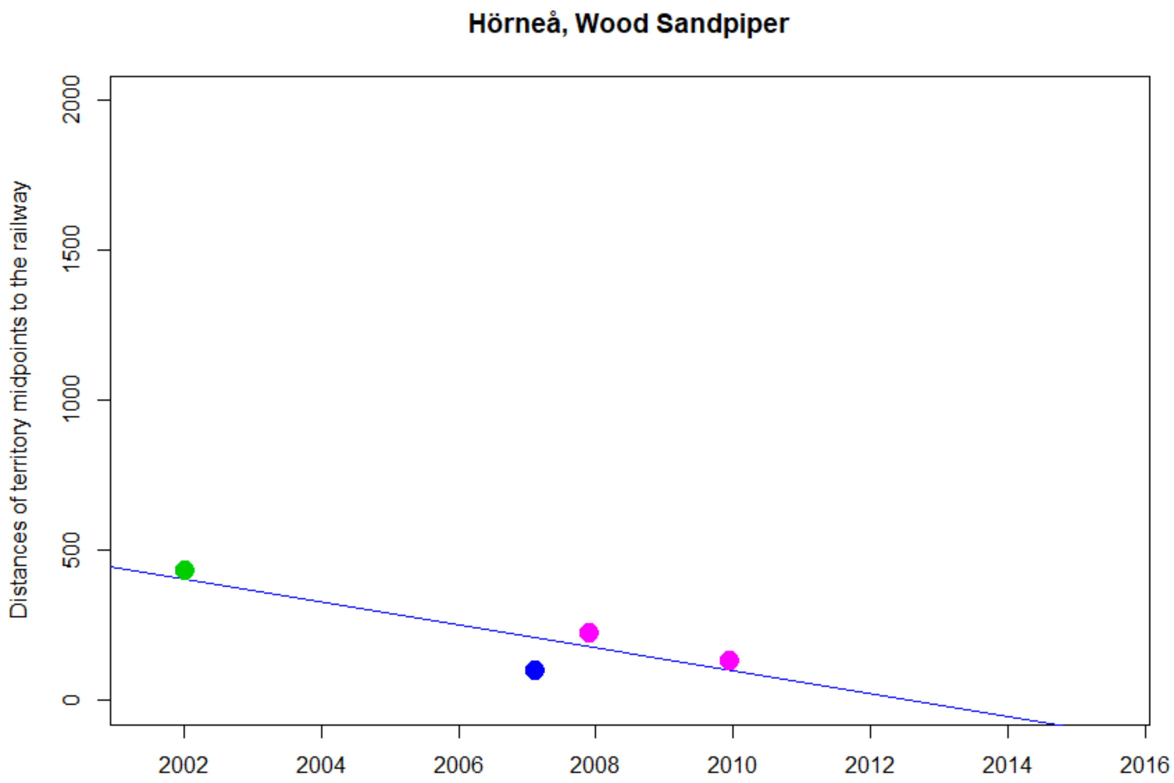


```
> cor.test(DNyland$Year,DNyland$NEAR_DIST_Botnia, method="kendall")

        Kendall's rank correlation tau

data:  DNyland$Year and DNyland$NEAR_DIST_Botnia
T = 2, p-value = 0.75
alternative hypothesis: true tau is not equal to 0
sample estimates:
       tau
-0.3333333
```

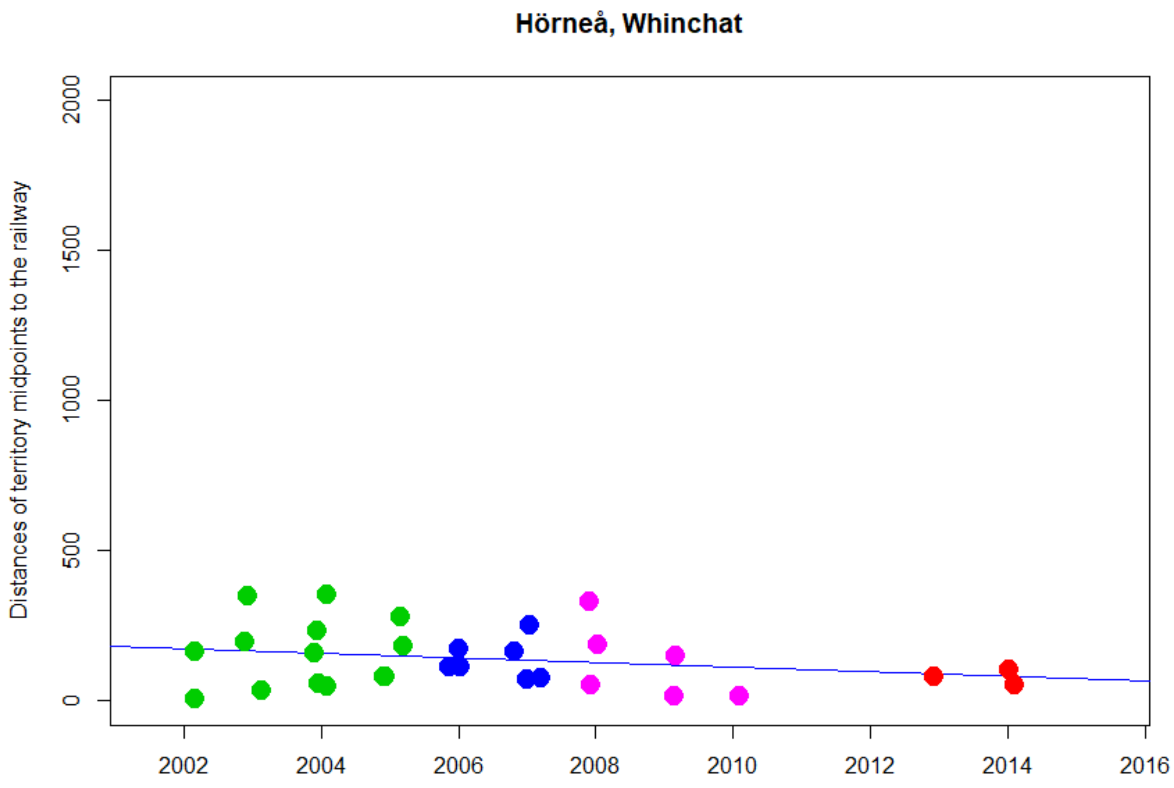


```
> cor.test(DNyland$Year,DNyland$NEAR_DIST_Botnia, method="kendall")

        Kendall's rank correlation tau

data:  DNyland$Year and DNyland$NEAR_DIST_Botnia
z = -0.98899, p-value = 0.3227
alternative hypothesis: true tau is not equal to 0
sample estimates:
       tau
-0.1318395
```

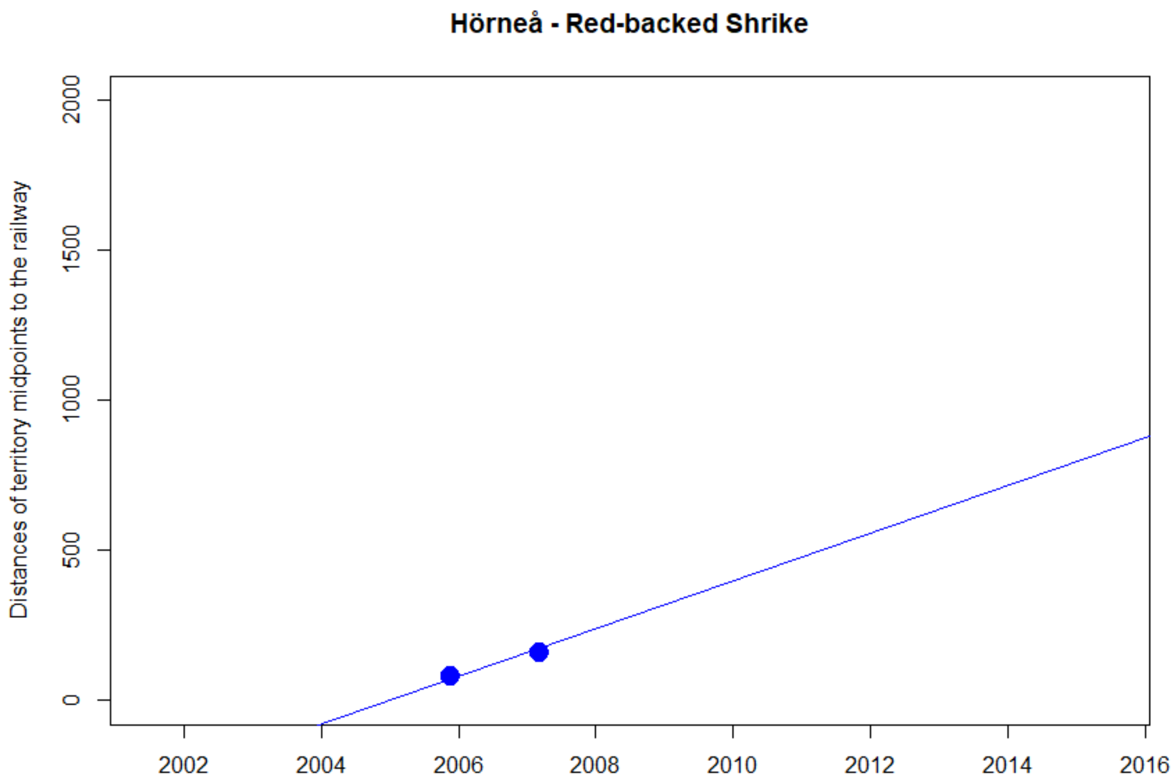
Hörneå - Red-backed Shrike
Distances of territory midpoints to the railway
0
500
1000
1500
2000
2002
2004
2006
2008
2010
2012
2014
2016

NA

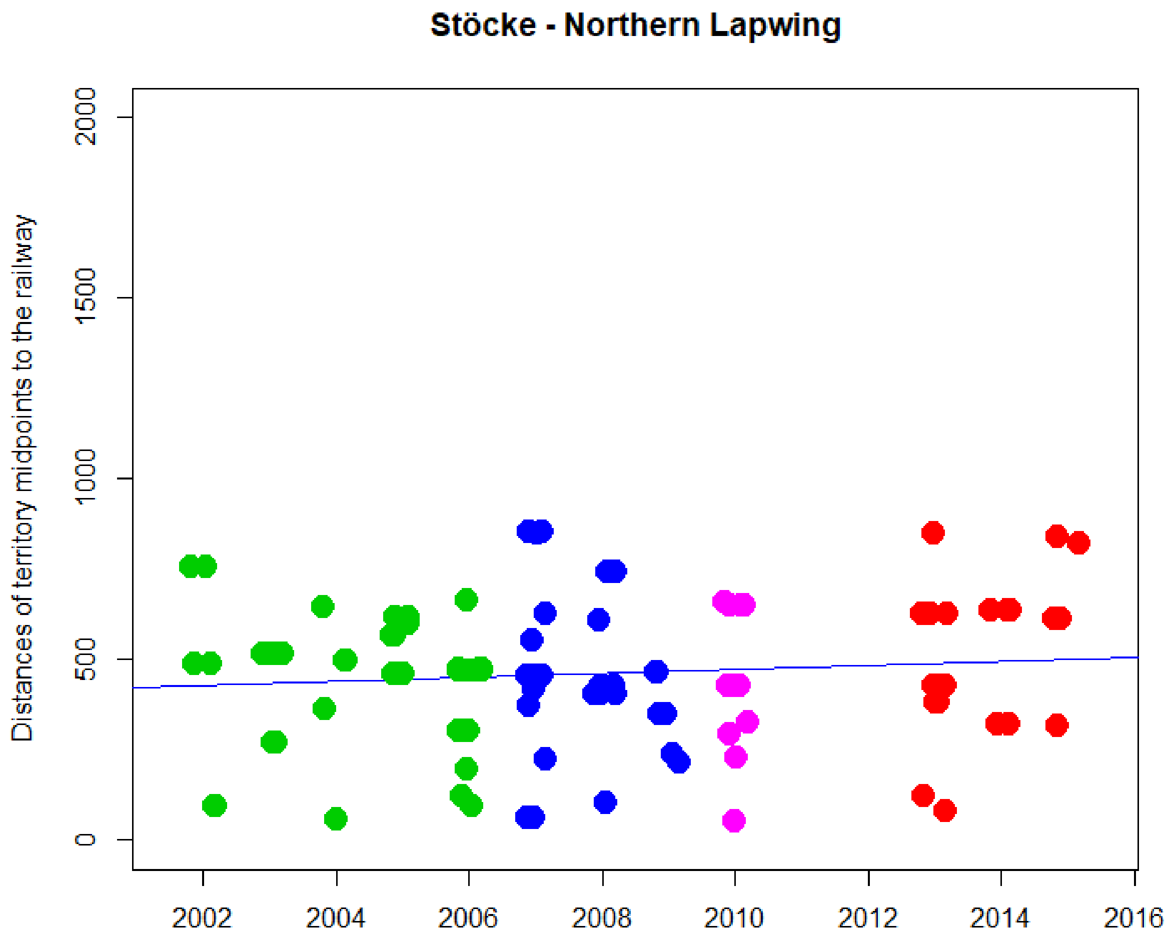


```
> cor.test(DNyland$Year,DNyland$NEAR_DIST_Botnia, method="kendall")

        Kendall's rank correlation tau

data:  DNyland$Year and DNyland$NEAR_DIST_Botnia
z = -0.25381, p-value = 0.7996
alternative hypothesis: true tau is not equal to 0
sample estimates:
        tau
-0.01689091
```

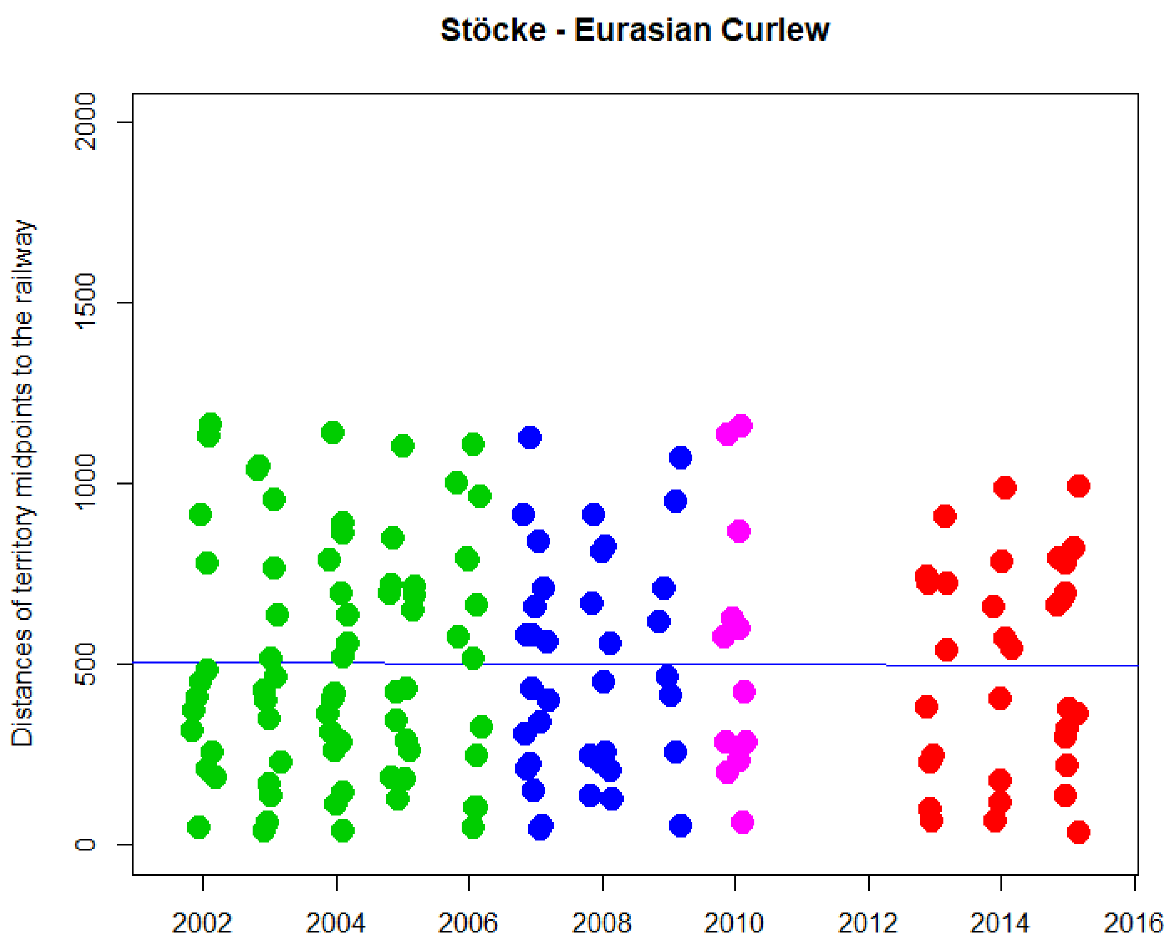


```
> cor.test(DNyland$Year,DNyland$NEAR_DIST_Botnia, method="kendall")

        Kendall's rank correlation tau

data:  DNyland$Year and DNyland$NEAR_DIST_Botnia
z = 0.094788, p-value = 0.9245
alternative hypothesis: true tau is not equal to 0
sample estimates:
        tau
0.005332744
```

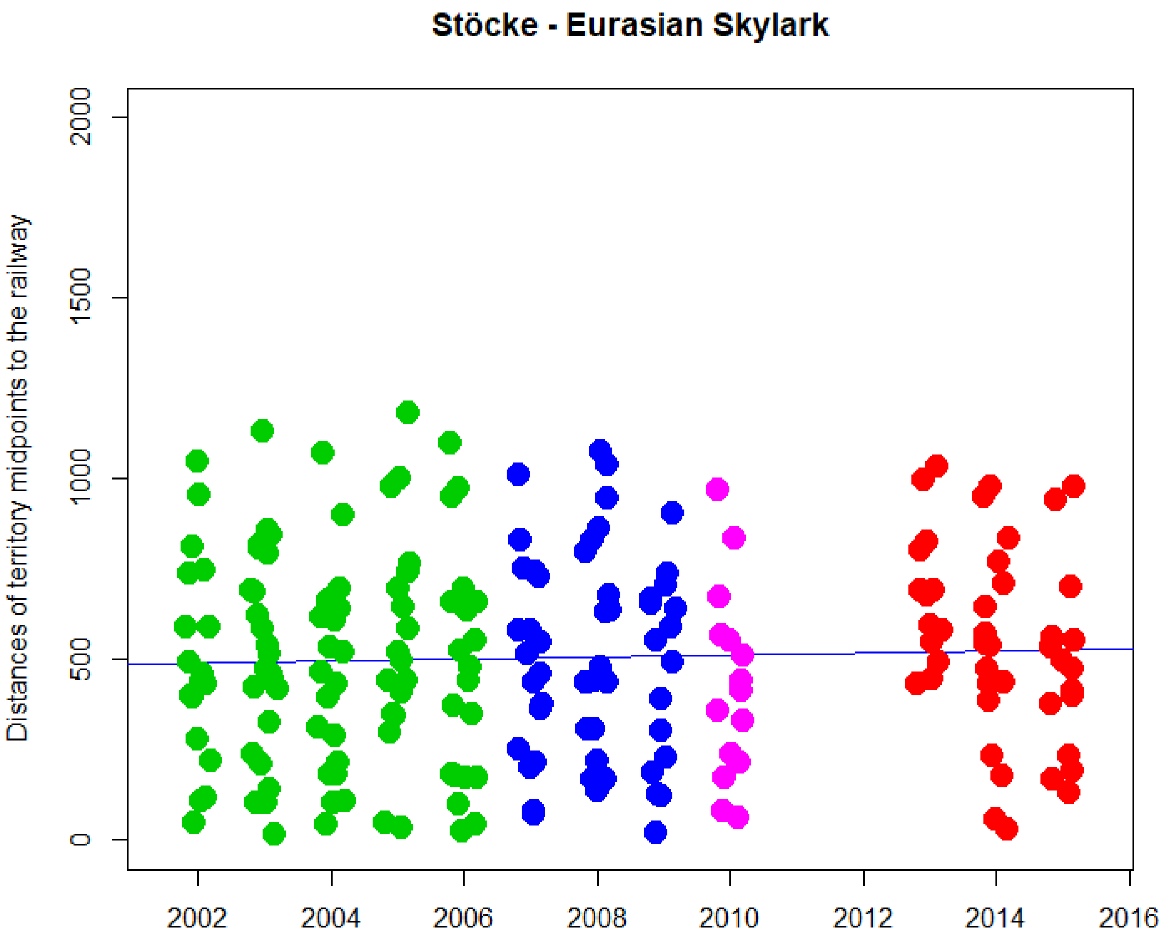


```
> cor.test(DNyland$Year,DNyland$NEAR_DIST_Botnia, method="kendall")

        Kendall's rank correlation tau

data:  DNyland$Year and DNyland$NEAR_DIST_Botnia
z = 0.43258, p-value = 0.6653
alternative hypothesis: true tau is not equal to 0
sample estimates:
       tau
0.02042237
```

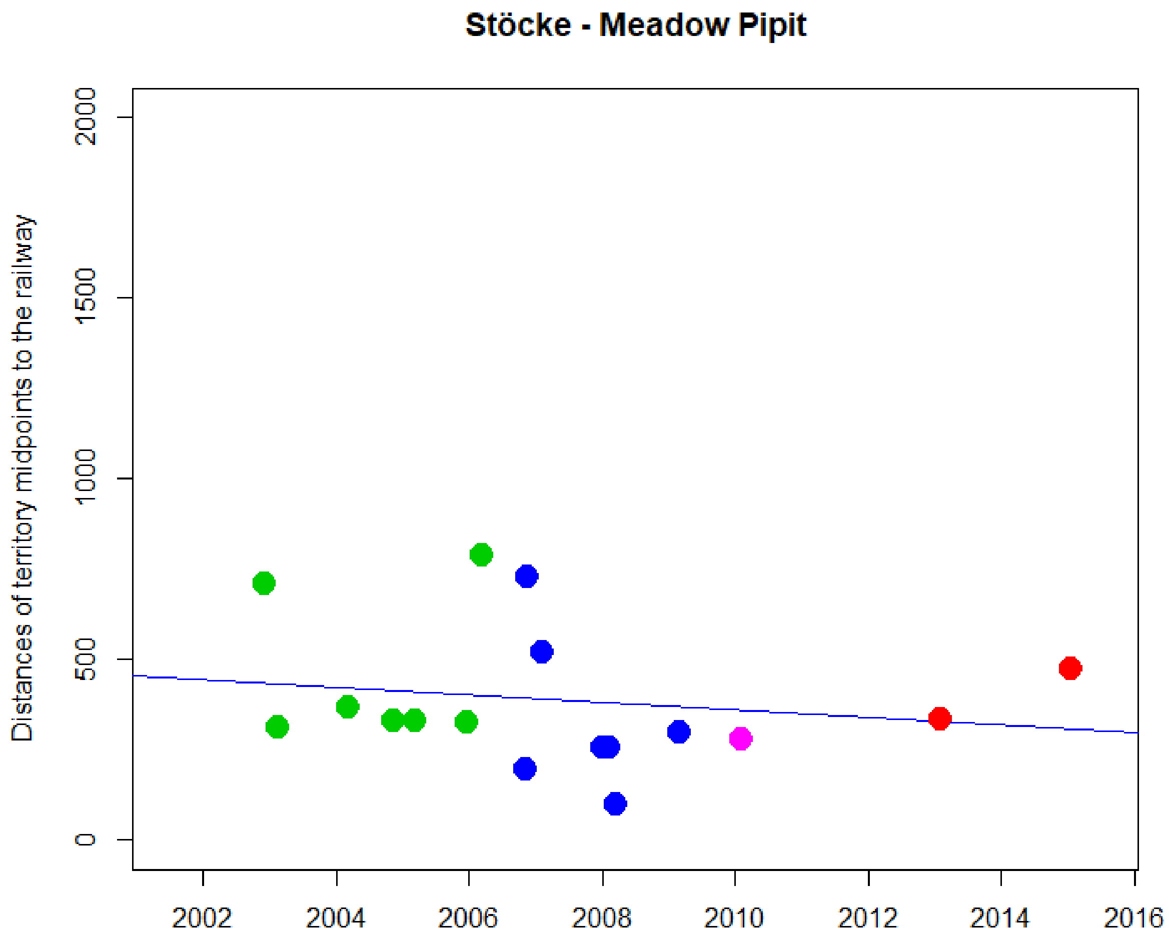


```
> cor.test(DNyland$Year,DNyland$NEAR_DIST_Botnia, method="kendall")

        Kendall's rank correlation tau

data:  DNyland$Year and DNyland$NEAR_DIST_Botnia
z = -0.95739, p-value = 0.3384
alternative hypothesis: true tau is not equal to 0
sample estimates:
       tau
-0.1763086
```

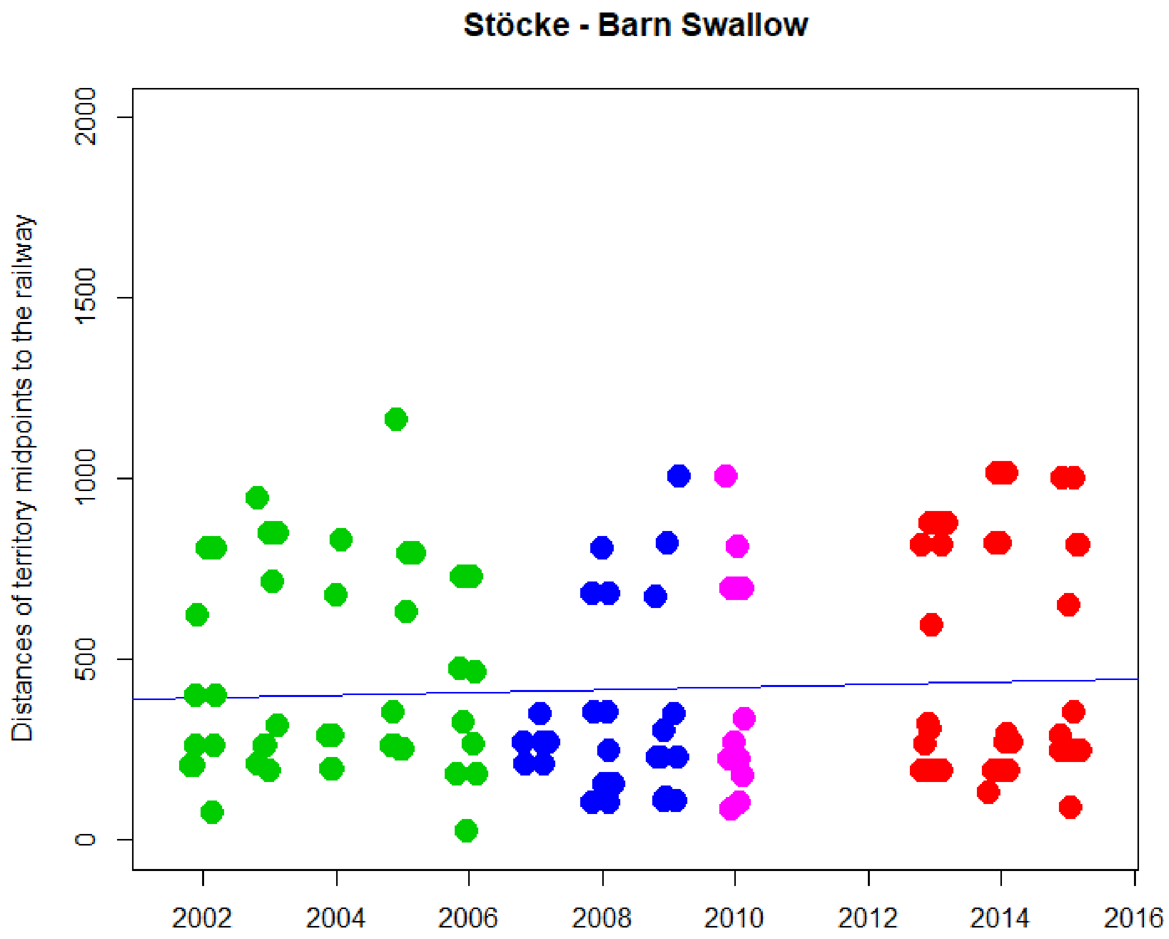


```
> cor.test(DNyland$Year,DNyland$NEAR_DIST_Botnia, method="kendall")

        Kendall's rank correlation tau

data:  DNyland$Year and DNyland$NEAR_DIST_Botnia
z = 0.1583, p-value = 0.8742
alternative hypothesis: true tau is not equal to 0
sample estimates:
        tau
0.009680589
```

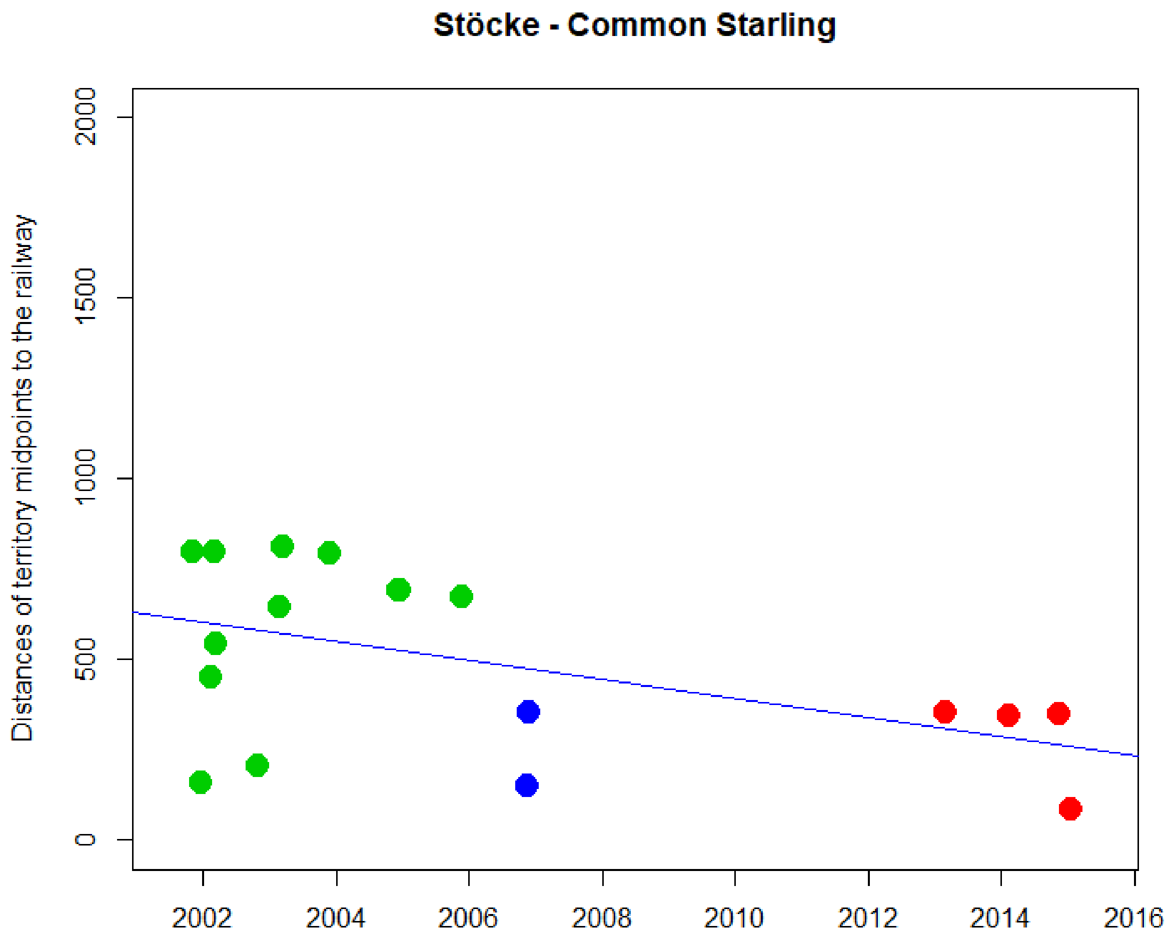


```
> cor.test(DNyland$Year,DNyland$NEAR_DIST_Botnia, method="kendall")

        Kendall's rank correlation tau

data:  DNyland$Year and DNyland$NEAR_DIST_Botnia
z = -2.0447, p-value = 0.04088
alternative hypothesis: true tau is not equal to 0
sample estimates:
       tau
-0.3684912
```

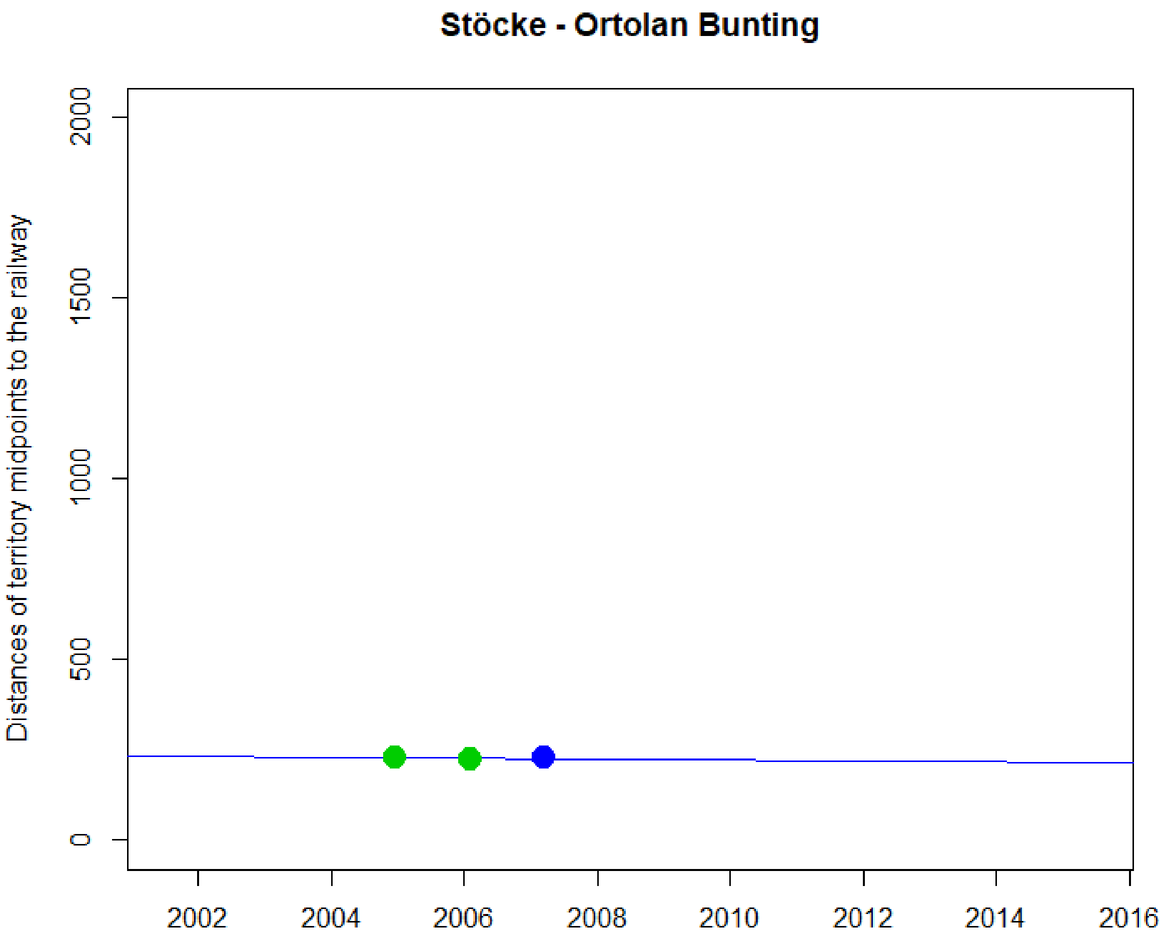

Stöcke - Ortolan Bunting
Distances of territory midpoints to the railway
0
500
1000
1500
2000
2002
2004
2006
2008
2010
2012
2014
2016


NA

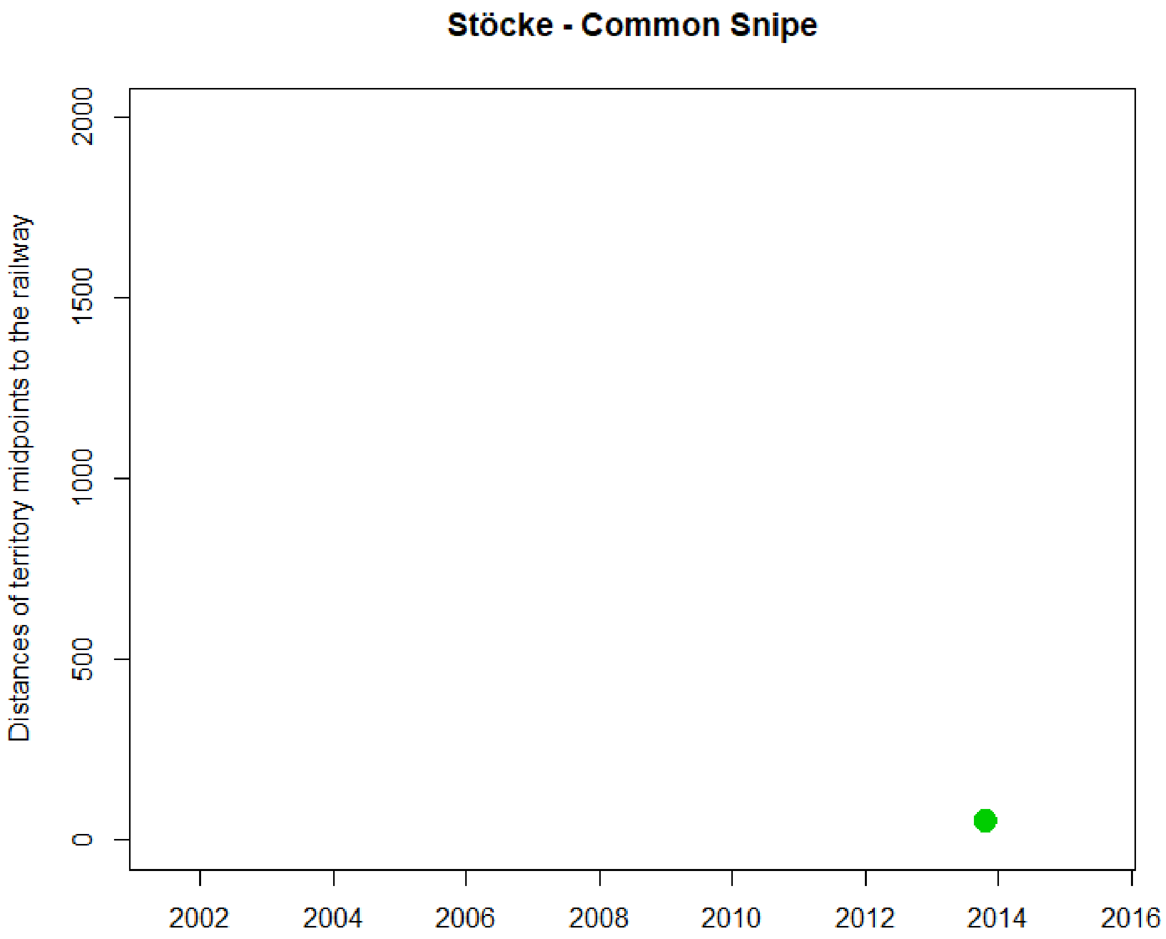
Stöcke - Common Snipe
Distances of territory midpoints to the railway
0
500
1000
1500
2000
2002
2004
2006
2008
2010
2012
2014
2016

NA

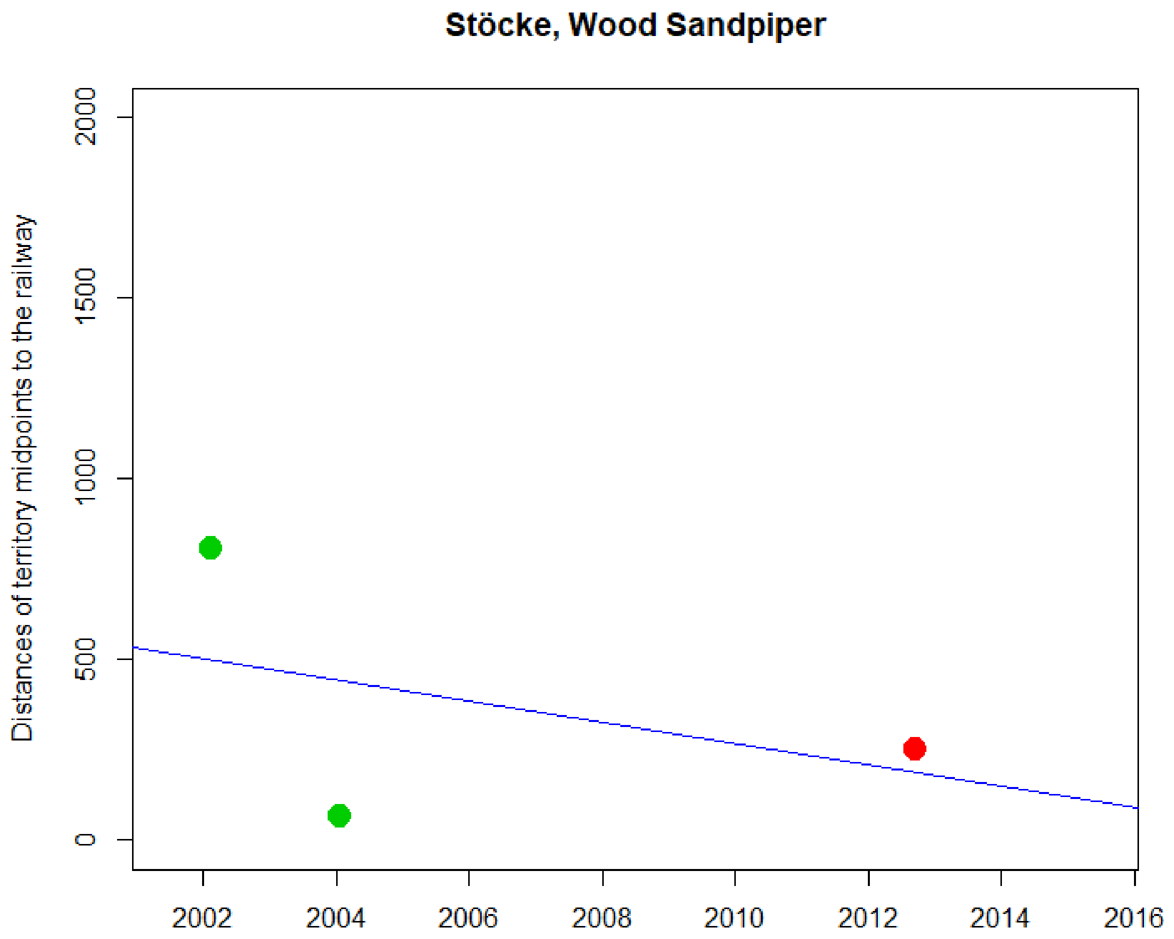
Stöcke, Wood Sandpiper
Distances of territory midpoints to the railway
0
500
1000
1500
2000
2002
2004
2006
2008
2010
2012
2014
2016

NA

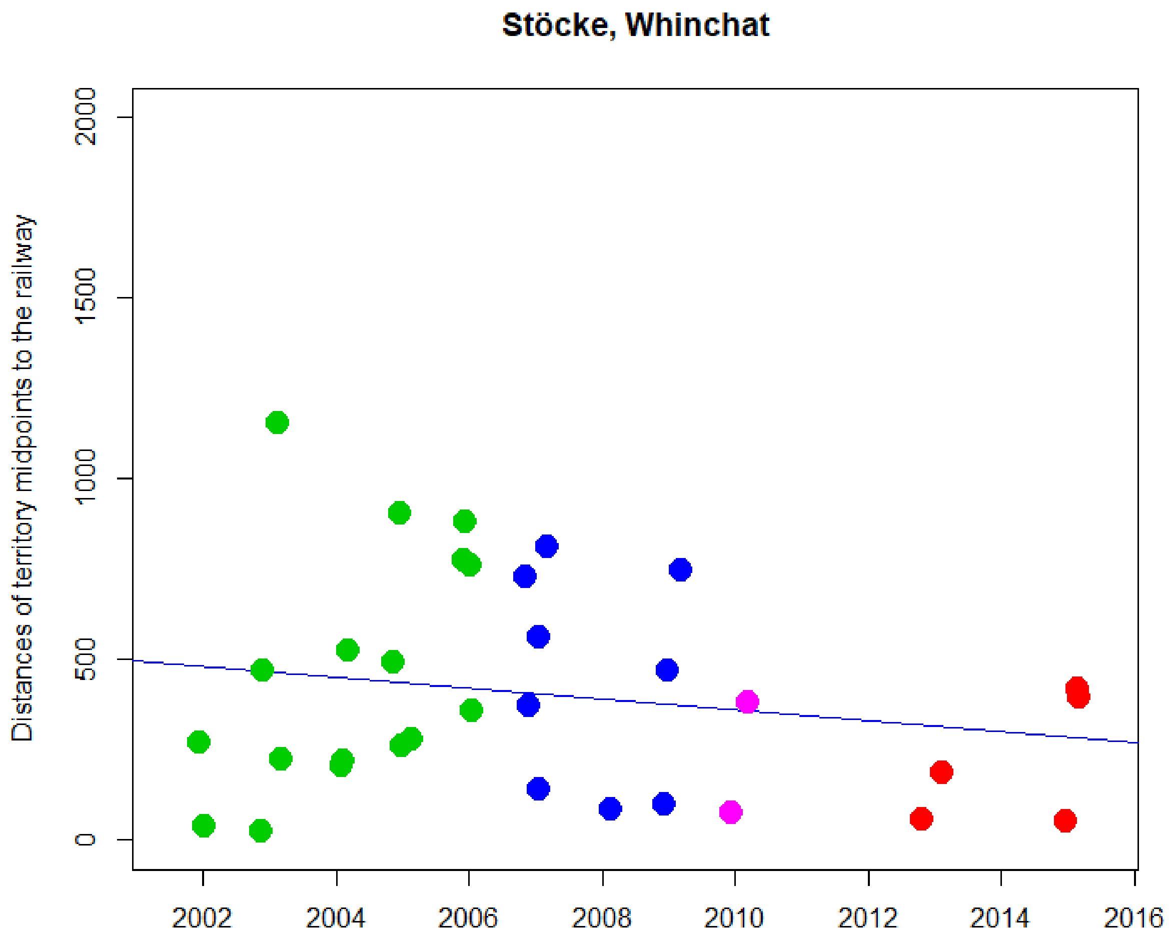


```
> cor.test(DNyland$Year,DNyland$NEAR_DIST_Botnia, method="kendall")

        Kendall's rank correlation tau

data:  DNyland$Year and DNyland$NEAR_DIST_Botnia
z = -0.62404, p-value = 0.5326
alternative hypothesis: true tau is not equal to 0
sample estimates:
        tau
-0.07880125
```

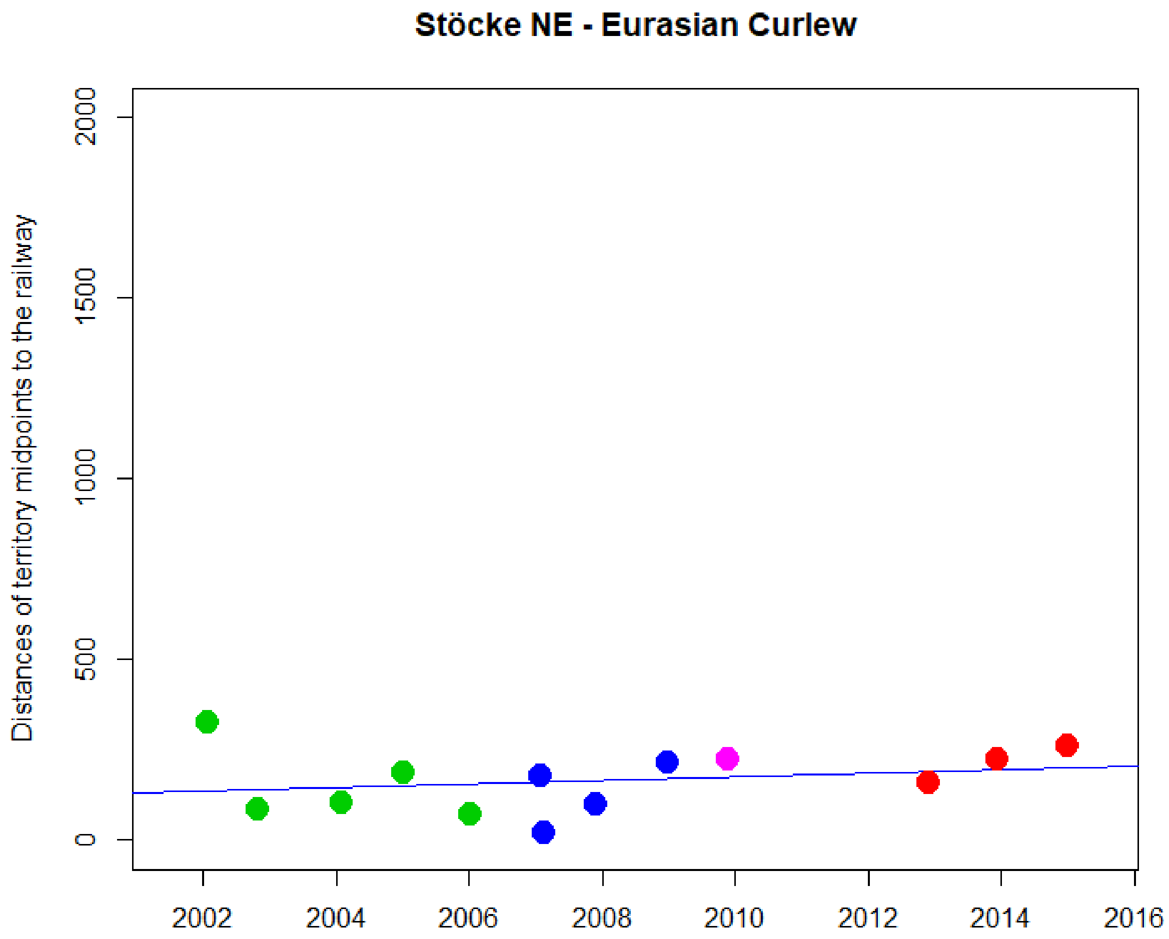


```
> cor.test(DNyland$Year,DNyland$NEAR_DIST_Botnia, method="kendall")

        Kendall's rank correlation tau

data:  DNyland$Year and DNyland$NEAR_DIST_Botnia
z = 1.2836, p-value = 0.1993
alternative hypothesis: true tau is not equal to 0
sample estimates:
      tau
0.2709734
```

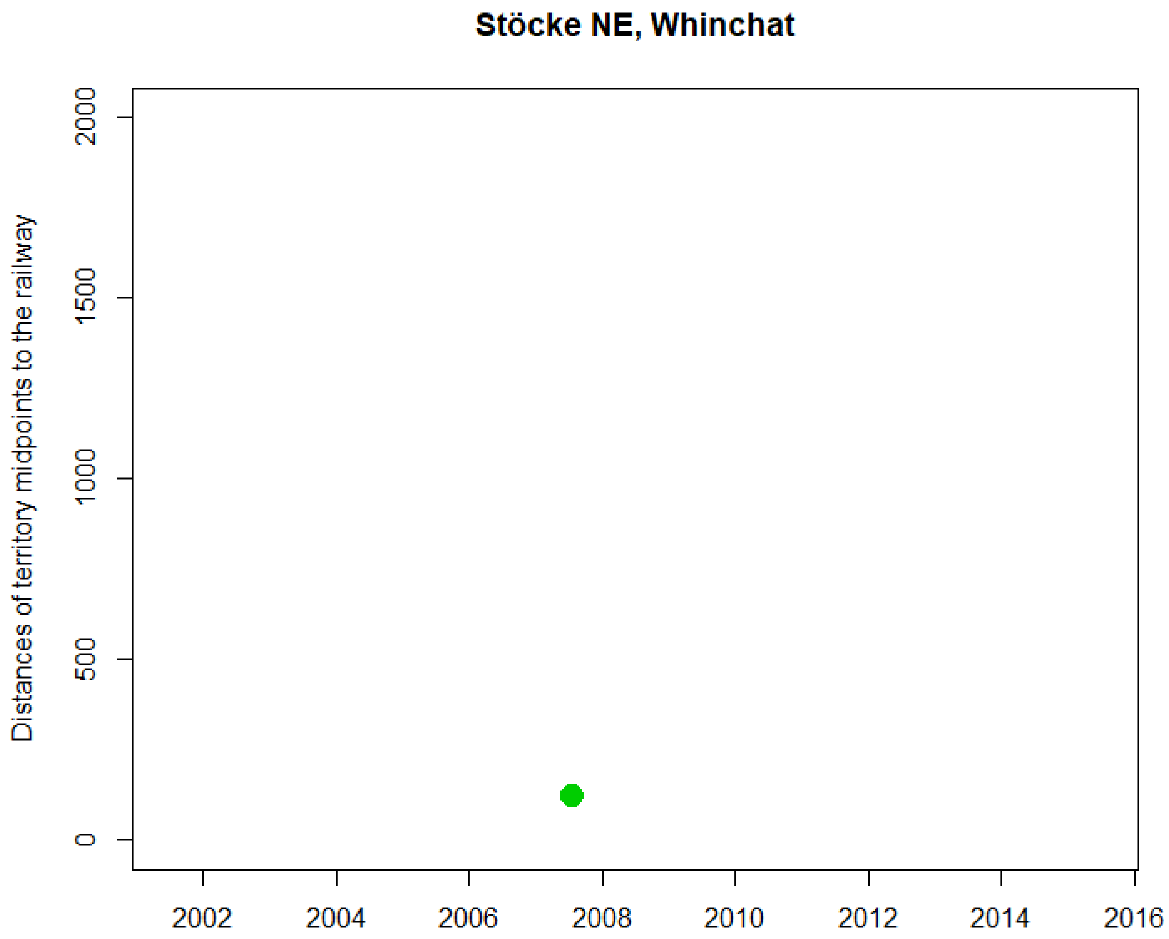
Stöcke NE, Whinchat
Distances of territory midpoints to the railway
0
500
1000
1500
2000
2002
2004
2006
2008
2010
2012
2014
2016

NA

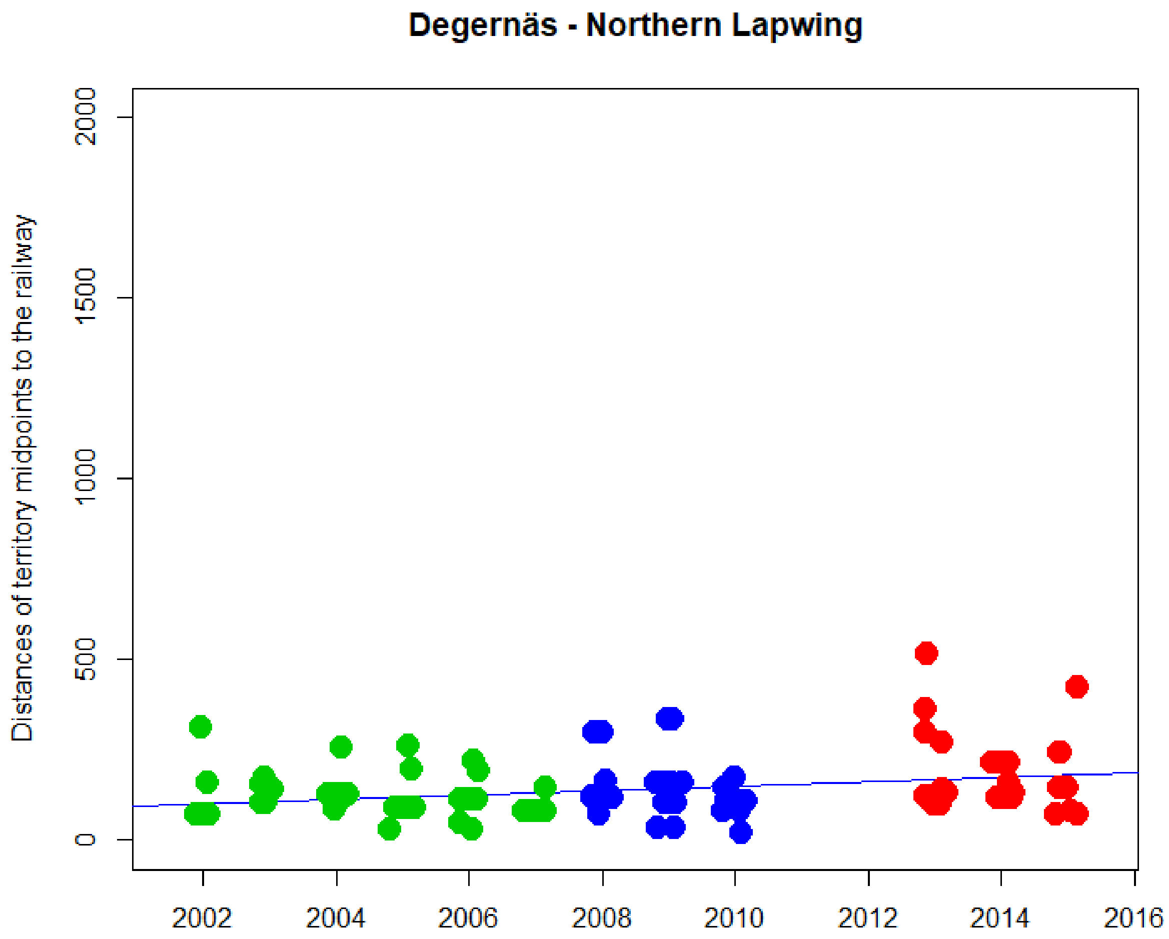


```
> cor.test(DNyland$Year,DNyland$NEAR_DIST_Botnia, method="kendall")

        Kendall's rank correlation tau

data:  DNyland$Year and DNyland$NEAR_DIST_Botnia
z = 2.5621, p-value = 0.0104
alternative hypothesis: true tau is not equal to 0
sample estimates:
      tau
0.1658561
```

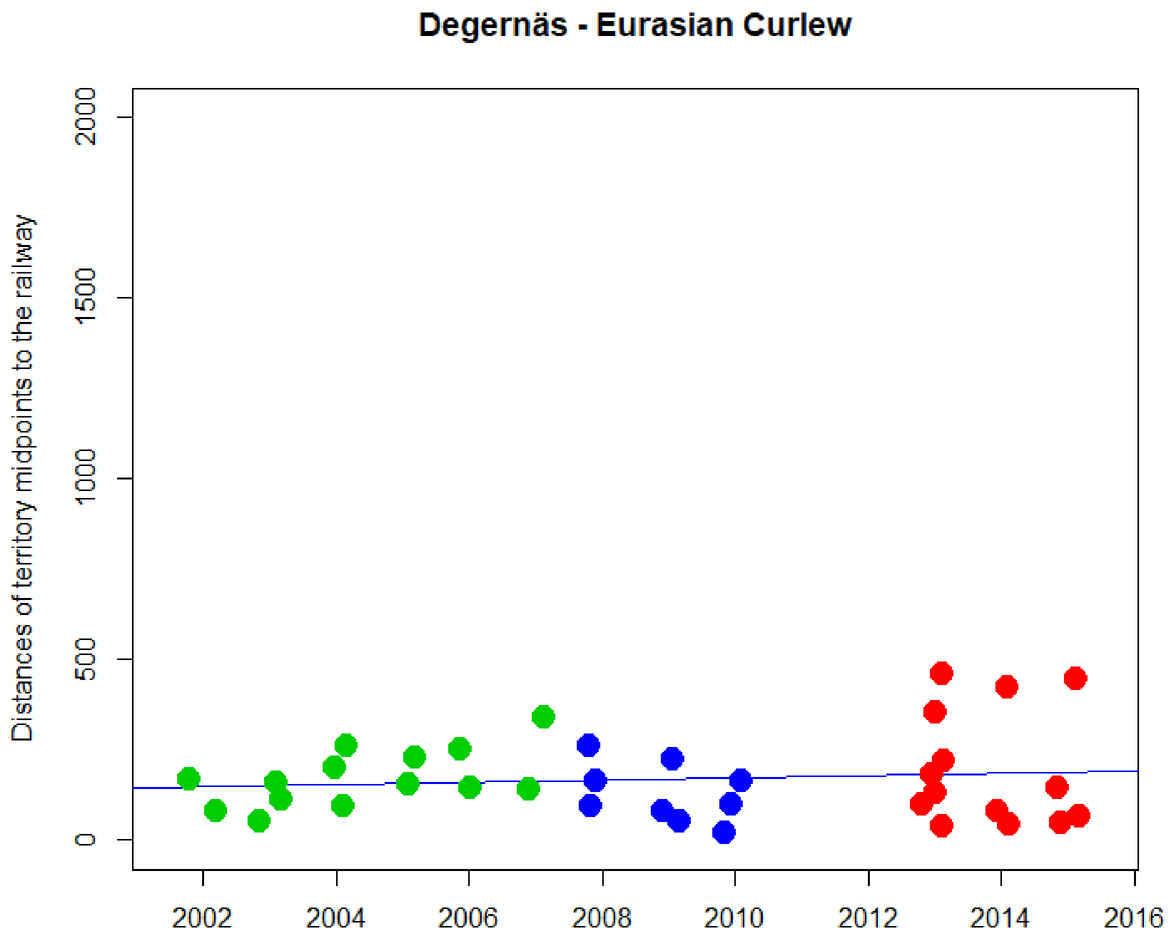


```
> cor.test(DNyland$Year,DNyland$NEAR_DIST_Botnia, method="kendall")

        Kendall's rank correlation tau

data:  DNyland$Year and DNyland$NEAR_DIST_Botnia
z = -0.3292, p-value = 0.742
alternative hypothesis: true tau is not equal to 0
sample estimates:
        tau
-0.03899961
```

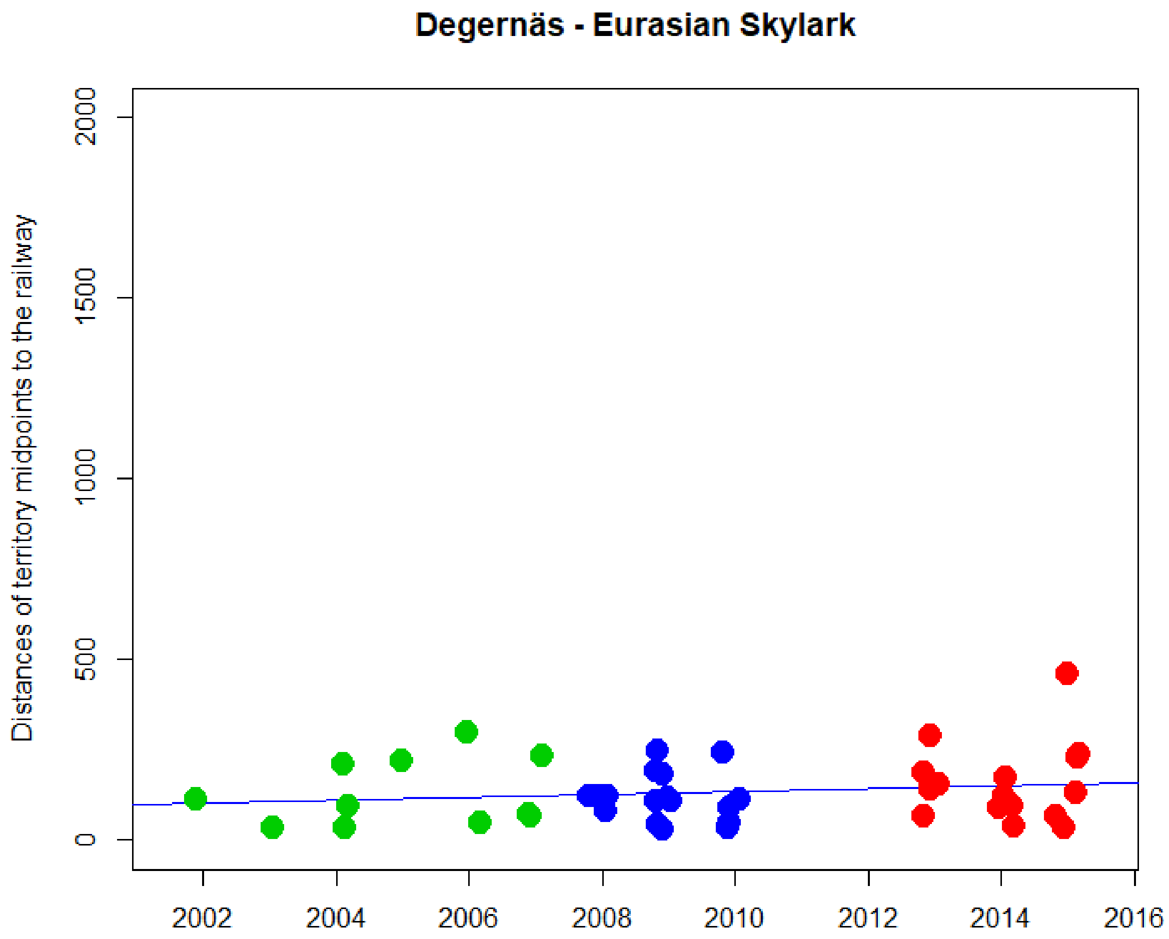


```
> cor.test(DNyland$Year,DNyland$NEAR_DIST_Botnia, method="kendall")

        Kendall's rank correlation tau

data:  DNyland$Year and DNyland$NEAR_DIST_Botnia
z = 0.89777, p-value = 0.3693
alternative hypothesis: true tau is not equal to 0
sample estimates:
       tau
0.09660575
```

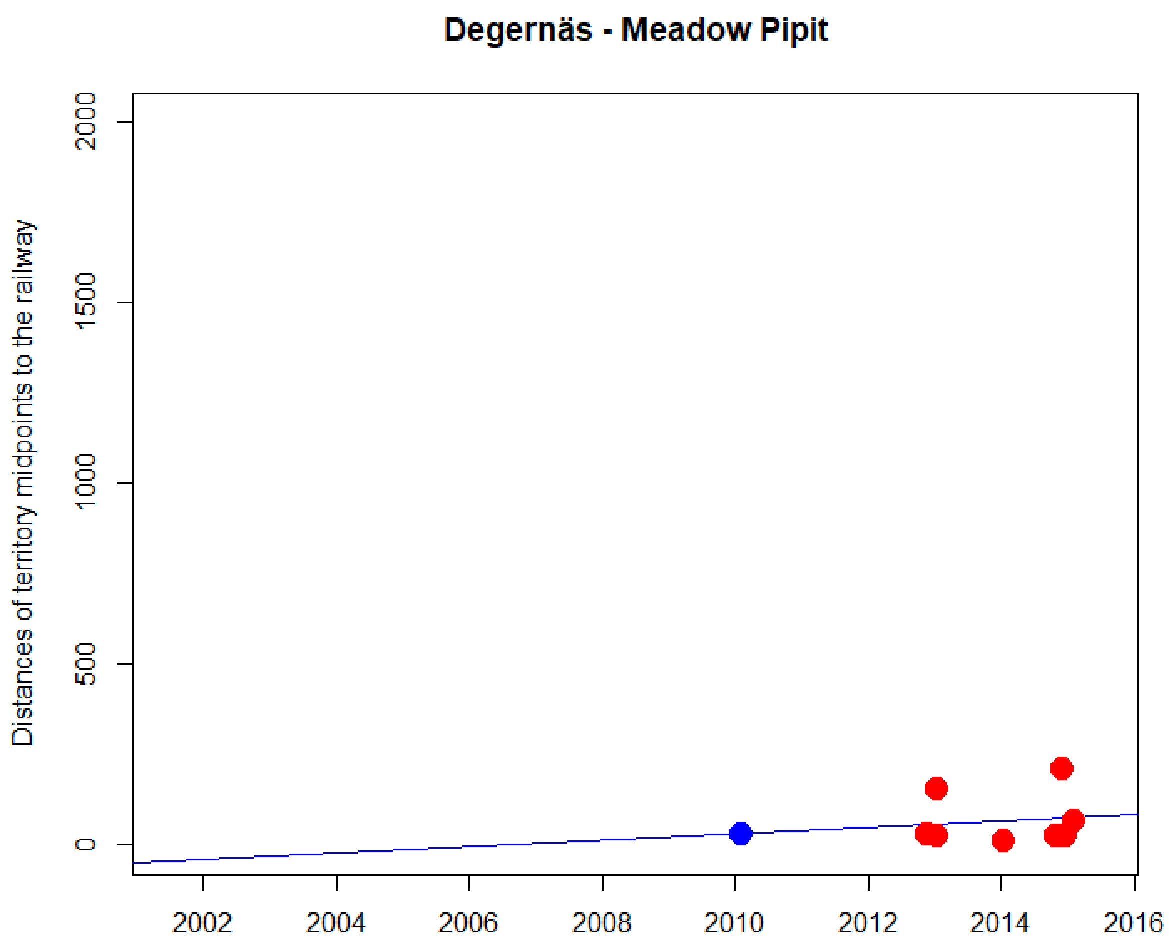


```
> cor.test(DNyland$Year,DNyland$NEAR_DIST_Botnia, method="kendall")

        Kendall's rank correlation tau

data:  DNyland$Year and DNyland$NEAR_DIST_Botnia
z = 0.11204, p-value = 0.9108
alternative hypothesis: true tau is not equal to 0
sample estimates:
       tau
0.03207501
```

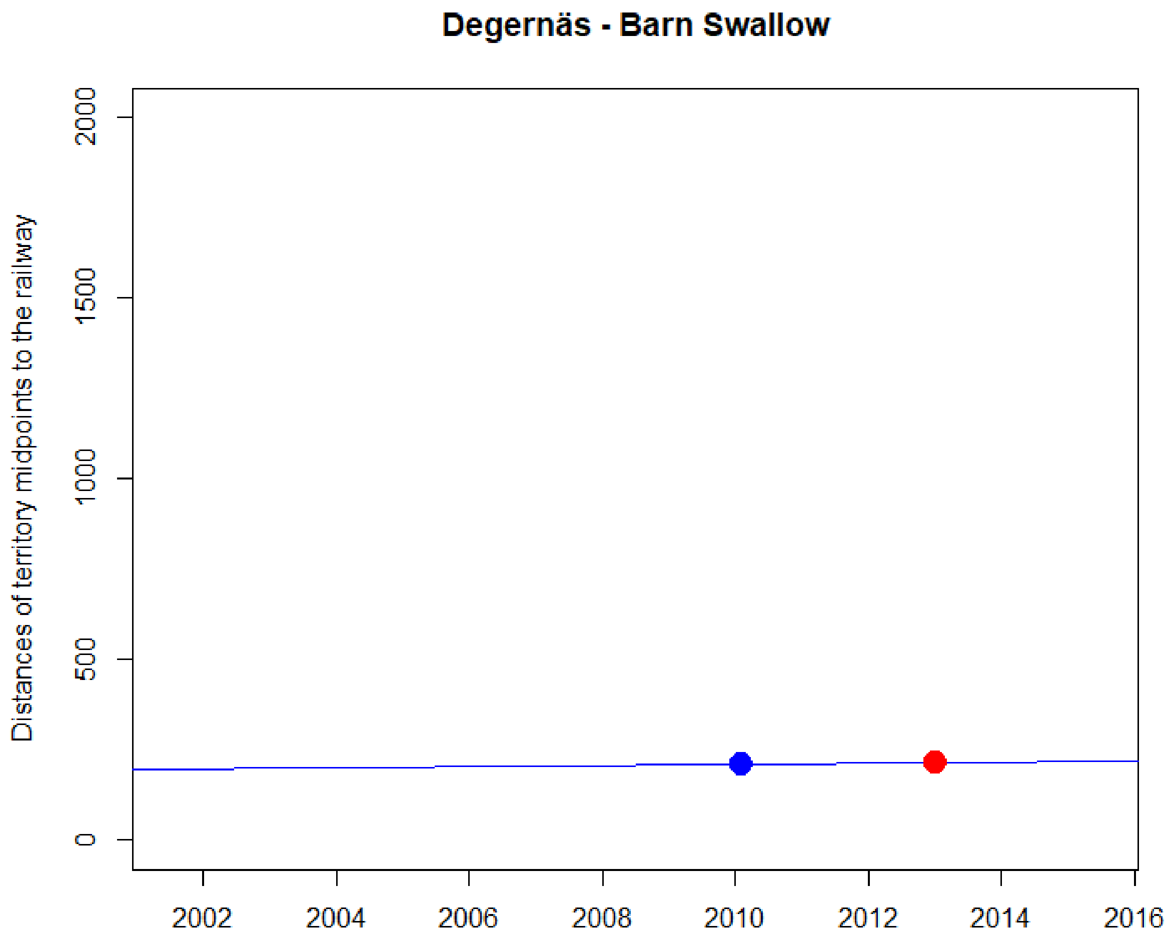
Degernäs - Barn Swallow
Distances of territory midpoints to the railway
0
500
1000
1500
2000
2002
2004
2006
2008
2010
2012
2014
2016

NA

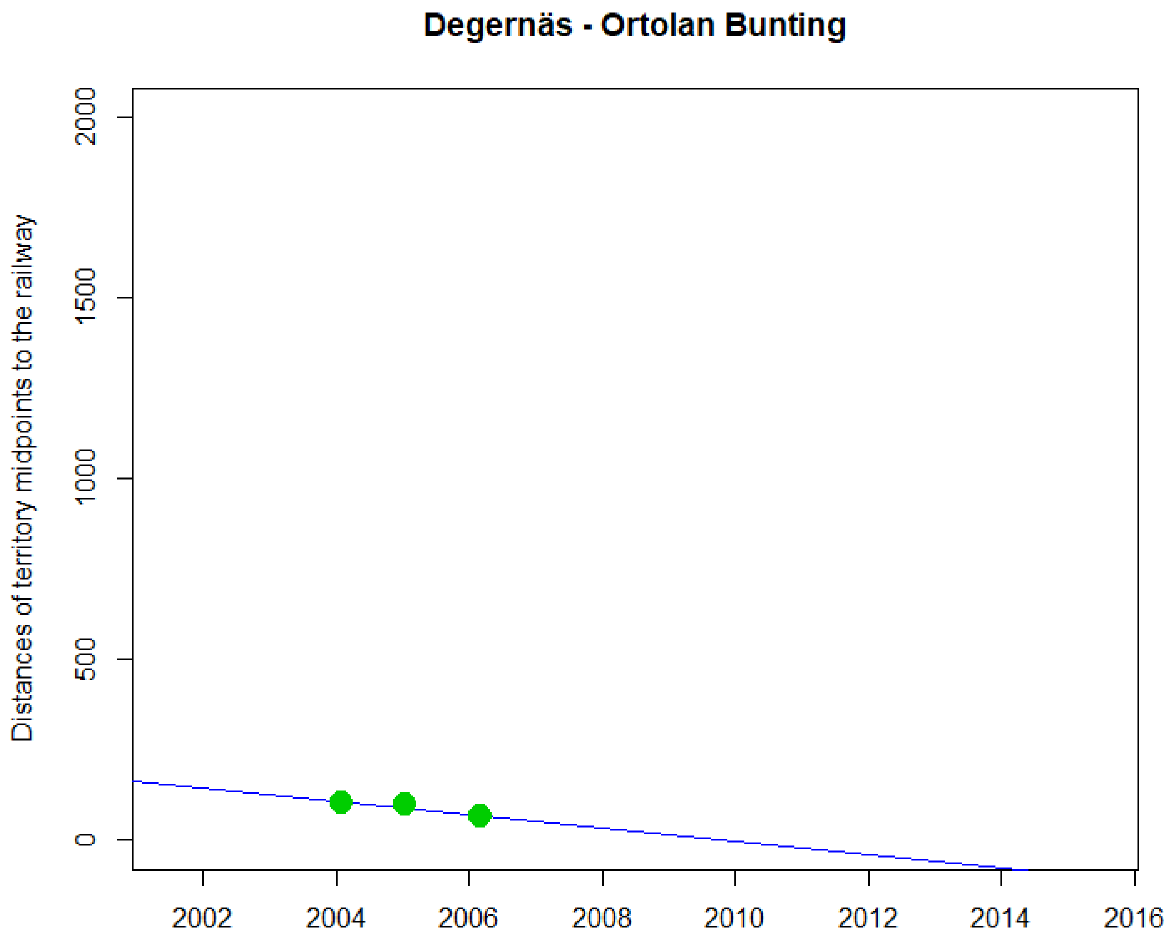
Degernäs - Ortolan Bunting
Distances of territory midpoints to the railway
0
500
1000
1500
2000
2002
2004
2006
2008
2010
2012
2014
2016

NA

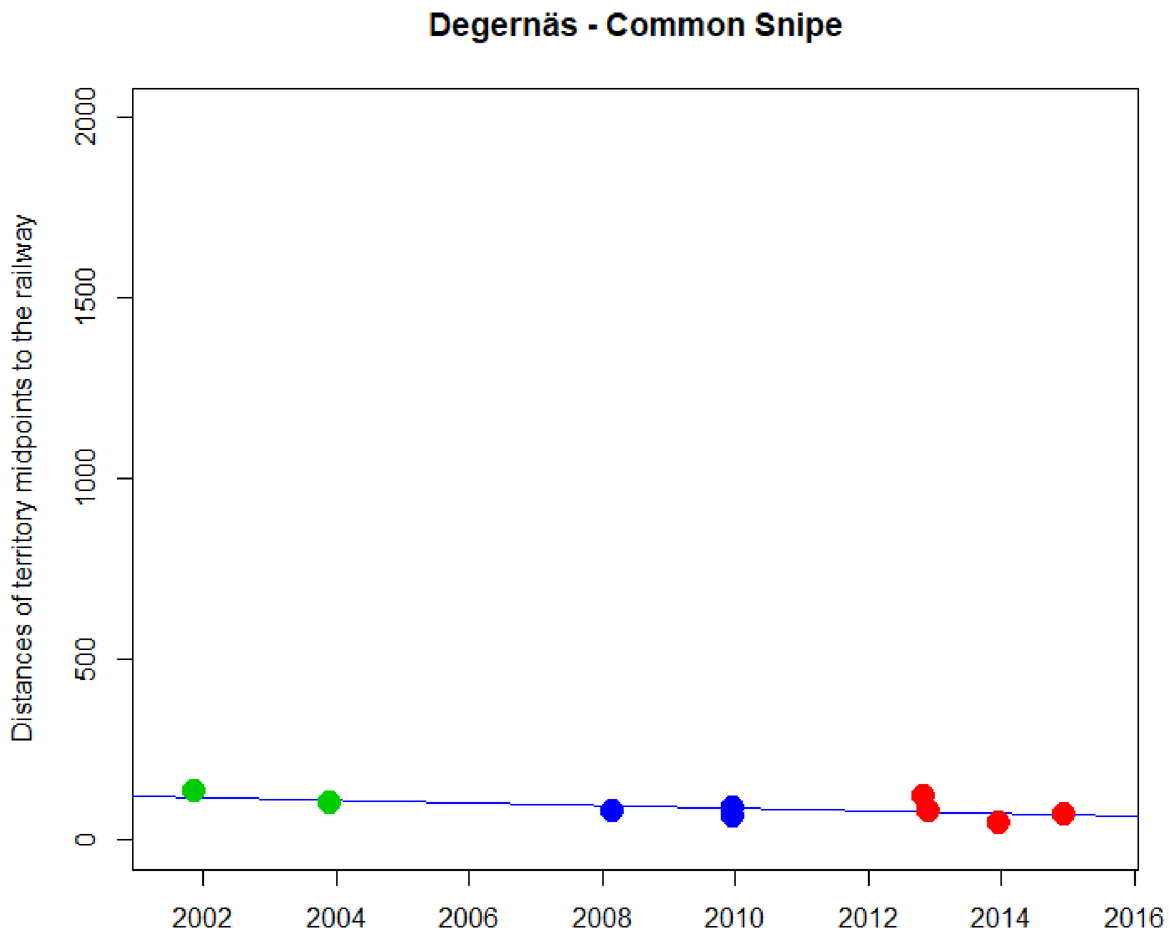


```
> cor.test(DNyland$Year,DNyland$NEAR_DIST_Botnia, method="kendall")

        Kendall's rank correlation tau

data:  DNyland$Year and DNyland$NEAR_DIST_Botnia
z = -1.8974, p-value = 0.05778
alternative hypothesis: true tau is not equal to 0
sample estimates:
       tau
-0.5144958
```

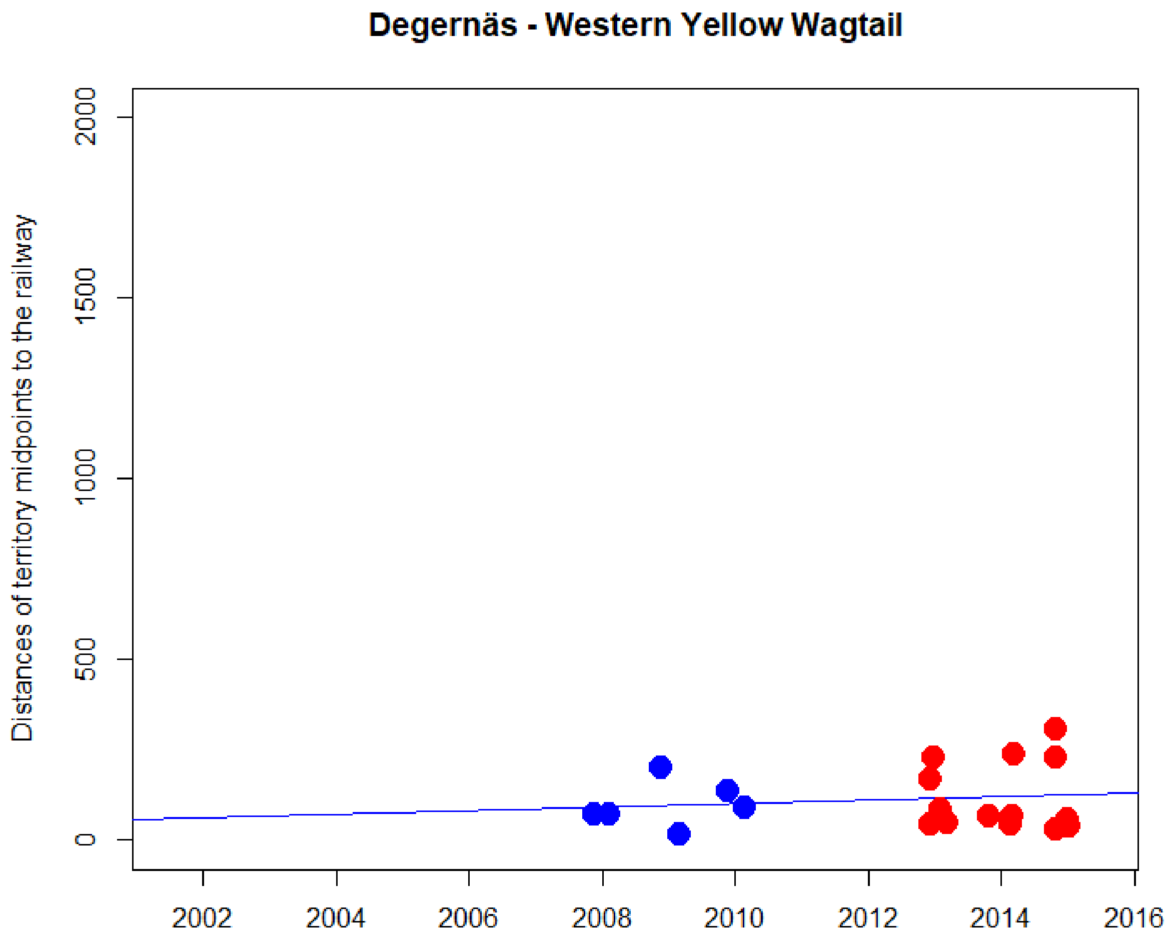


```
> cor.test(DNyland$Year,DNyland$NEAR_DIST_Botnia, method="kendall")

        Kendall's rank correlation tau

data:  DNyland$Year and DNyland$NEAR_DIST_Botnia
z = -0.23291, p-value = 0.8158
alternative hypothesis: true tau is not equal to 0
sample estimates:
       tau
-0.0402352
```

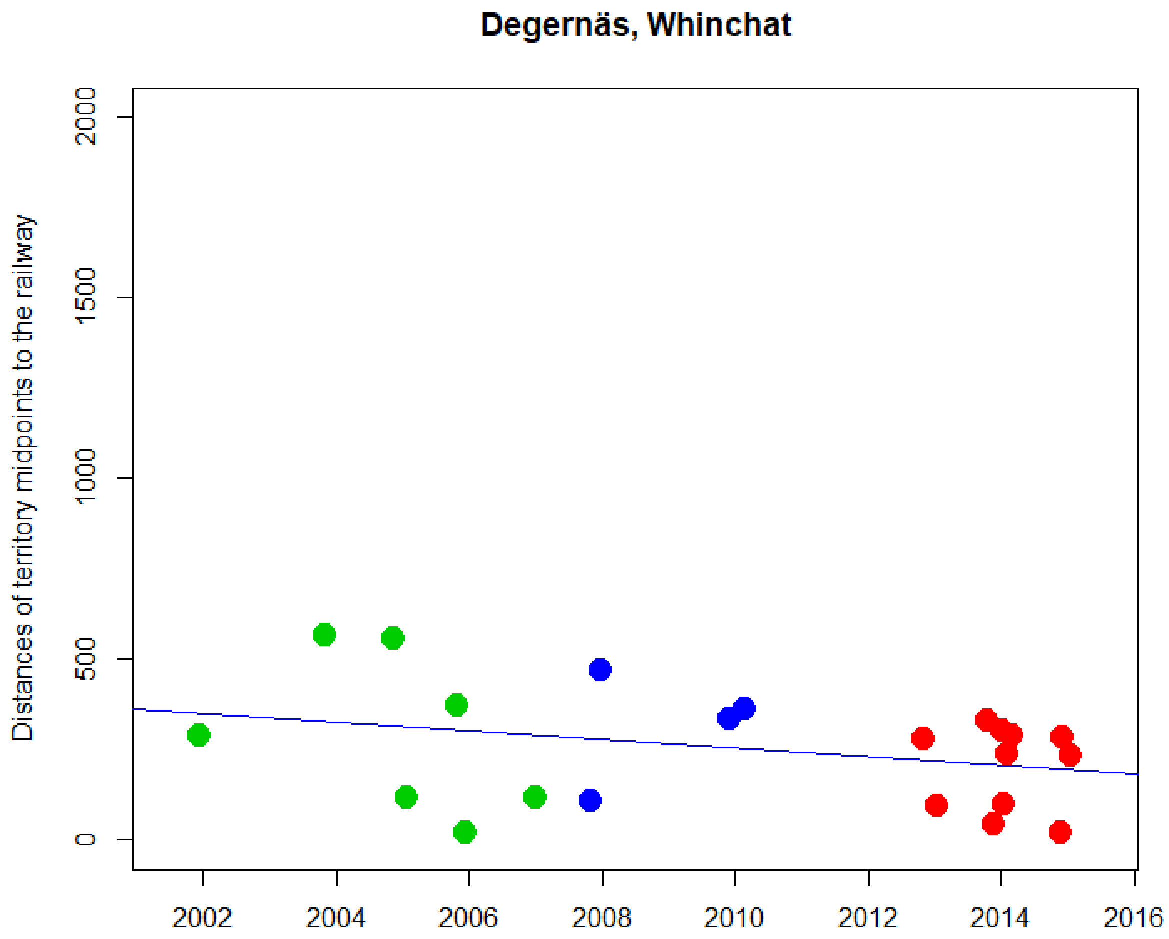


```
> cor.test(DNyland$Year,DNyland$NEAR_DIST_Botnia, method="kendall")

        Kendall's rank correlation tau

data:  DNyland$Year and DNyland$NEAR_DIST_Botnia
z = -1.8318, p-value = 0.06698
alternative hypothesis: true tau is not equal to 0
sample estimates:
       tau
-0.2919727
```

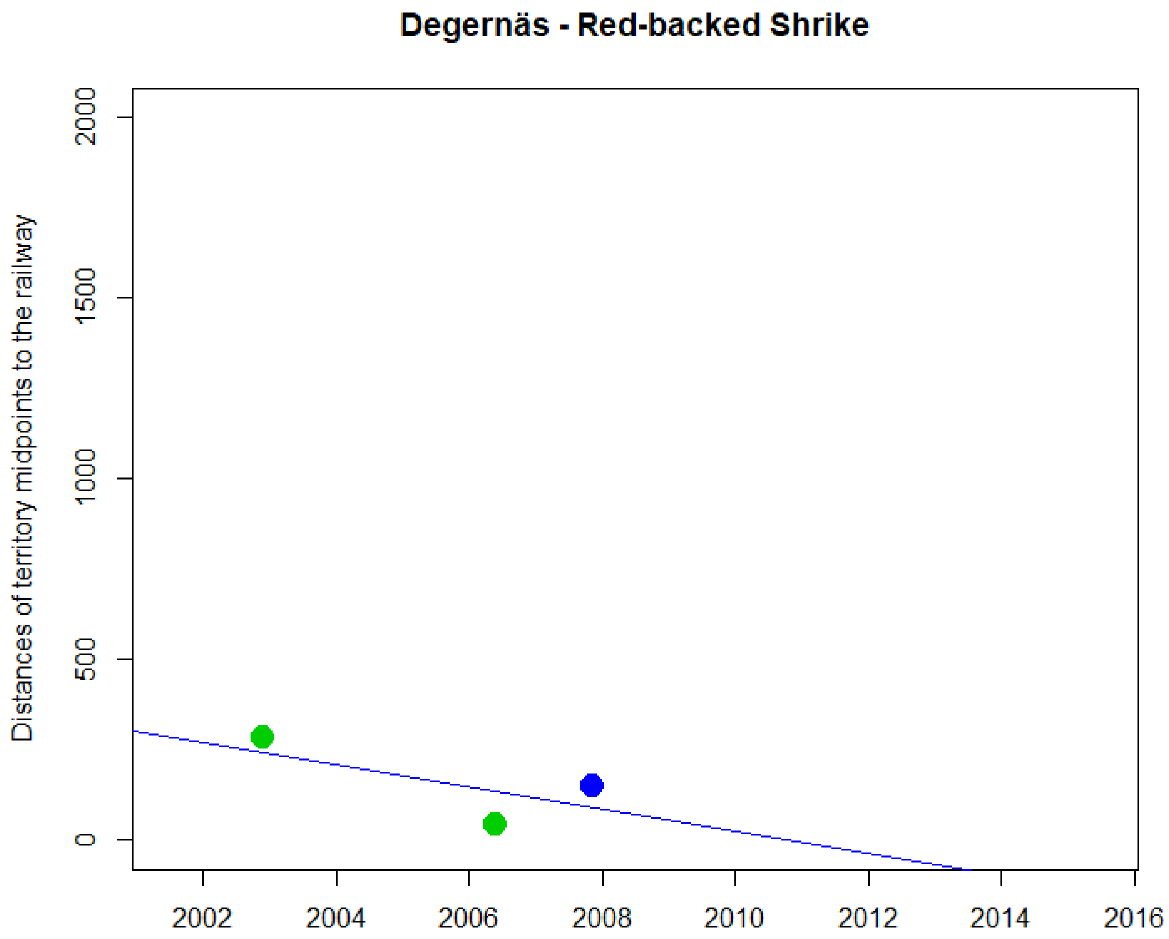
Degernäs - Red-backed Shrike
Distances of territory midpoints to the railway
0
500
1000
1500
2000
2002
2004
2006
2008
2010
2012
2014
2016

NA
