## Supplementary material for "Are railways detrimental to bird populations? A BDACI study on the construction of the Bothnia Line Railway": Mixed effects model output for TM to BLR distances

Supplementary material **Dist5**

### Mixed effect modelling for distances between territory midpoints and the Bothnia Line Railway across phases of railway construction for all species combined and for individual species

Output from R-script presented *as is*.

Lme models with formula NEAR_DIST_Botnia ~ 1 , random = ~ 1|Site (the "empty" model) and NEAR_DIST_Botnia ~ Status , random = ~ 1|Site, respectively, with Status levels *Before*, *Construction, Ready* and *Traffic*. The addition of the Status variable to the empty models resulted in significant improvement expressed in lower AICc values.

Negative estimates (column "Value") indicate that the distances to the railway decreased in relation to conditions *Before* railway construction (the Intercept in the models).

Significant (*P*<0.05) coefficients are marked with yellow background.

#### All species

```
> Dlme00<-lme(NEAR_DIST_Botnia ~ 1 , random = ~ 1|Site, data=BotDist)
> summary(Dlme00)
Linear mixed-effects model fit by REML
 Data: BotDist
       AIC      BIC    logLik
  44425.02 44443.28 -22209.51

Random effects:
 Formula: ~1 | Site
        (Intercept) Residual
StdDev:    146.5067 225.4836

Fixed effects: NEAR_DIST_Botnia ~ 1
               Value Std.Error   DF  t-value p-value
(Intercept) 343.0013  41.12983 3232 8.339478       0

Standardized Within-Group Residuals:
        Min          Q1         Med          Q3         Max
-2.37238798 -0.67322057 -0.05567101  0.71788882  5.10782262

Number of Observations: 3245
Number of Groups: 13
> AIC(Dlme00)
[1] 44425.02
> Dlme01<-lme(NEAR_DIST_Botnia ~ Status , random = ~ 1|Site, data=BotDist)
> summary(Dlme01)
Linear mixed-effects model fit by REML
 Data: BotDist
       AIC      BIC    logLik
  44407.17 44443.68 -22197.59

Random effects:
 Formula: ~1 | Site
        (Intercept) Residual
StdDev:     147.188 225.4462

Fixed effects: NEAR_DIST_Botnia ~ 1 + Status
                      Value Std.Error   DF   t-value p-value
(Intercept)        355.3200  42.17652 3229  8.424594  0.0000
StatusConstruction -19.1241  12.67410 3229 -1.508912  0.1314
StatusReady        -20.0267  12.73645 3229 -1.572396  0.1160
StatusTrafic        -4.7586  12.38332 3229 -0.384271  0.7008

Correlation:
                   (Intr) SttsCn SttsRd
StatusConstruction -0.150
StatusReady        -0.184  0.529
StatusTrafic       -0.162  0.501  0.588

Standardized Within-Group Residuals:
        Min          Q1         Med          Q3         Max
-2.40350153 -0.67648078 -0.05647363  0.70426553  5.15123100

Number of Observations: 3245
Number of Groups: 13
> AIC(Dlme01)
[1] 44407.17
> AICc(Dlme00)
[1] 44425.03
> AICc(Dlme01)
[1] 44407.2
```

#### Northern Lapwing

```
VvDist<-BotDist[BotDist$Species=="Vv",]
> ## Empty model
> Dlme00<-lme(fixed = NEAR_DIST_Botnia ~ 1 , random = ~ 1|Site, data=VvDis
t)
> summary(Dlme00)
Linear mixed-effects model fit by REML
 Data: VvDist
       AIC      BIC    logLik
  8430.111 8443.547 -4212.056

Random effects:
 Formula: ~1 | Site
        (Intercept) Residual
StdDev:    359.6532 149.1158

Fixed effects: NEAR_DIST_Botnia ~ 1
               Value Std.Error  DF t-value p-value
(Intercept) 423.8729  101.6526 639 4.16982       0

Standardized Within-Group Residuals:
        Min          Q1         Med          Q3         Max
-3.49344722 -0.55881648 -0.02441984  0.59981100  2.62421042

Number of Observations: 652
Number of Groups: 13
> AICc(Dlme00)
[1] 8430.148
> ## Status included
> Dlme01<-lme(fixed = NEAR_DIST_Botnia ~ Status , random = ~ 1|Site, data=
VvDist)
> summary(Dlme01)
Linear mixed-effects model fit by REML
 Data: VvDist
      AIC      BIC    logLik
  8412.39 8439.233 -4200.195

Random effects:
 Formula: ~1 | Site
        (Intercept) Residual
StdDev:    358.3085 149.3044

Fixed effects: NEAR_DIST_Botnia ~ Status
                      Value Std.Error  DF  t-value p-value
(Intercept)        409.4100 102.01788 636 4.013120  0.0001
StatusConstruction  19.4936  18.46794 636 1.055537  0.2916
StatusReady         18.6960  19.33524 636 0.966939  0.3339
StatusTrafic        18.0698  18.40048 636 0.982030  0.3265
 Correlation:
                   (Intr) SttsCn SttsRd
StatusConstruction -0.090
StatusReady        -0.106  0.505
StatusTrafic       -0.100  0.499  0.611

Standardized Within-Group Residuals:
        Min          Q1         Med          Q3         Max
-3.49533279 -0.56326683 -0.02280604  0.59743512  2.57011186

Number of Observations: 652
Number of Groups: 13
> AICc(Dlme01)
[1] 8412.52
```

### Eurasian Curlew

```
> NaDist<-BotDist[BotDist$Species=="Na",]
> ## Empty model
> Dlme00<-lme(fixed = NEAR_DIST_Botnia ~ 1 , random = ~ 1|Site, data=NaDis
t)
> summary(Dlme00)
Linear mixed-effects model fit by REML
 Data: NaDist
       AIC      BIC    logLik
  7162.642 7175.351 -3578.321

Random effects:
 Formula: ~1 | Site
        (Intercept) Residual
StdDev:    187.3792 256.6562

Fixed effects: NEAR_DIST_Botnia ~ 1
               Value Std.Error  DF  t-value p-value
(Intercept) 385.3768  55.78788 499 6.907895       0

Standardized Within-Group Residuals:
        Min          Q1         Med          Q3         Max
-2.26238875 -0.62724388 -0.02884387  0.68119110  3.10725140

Number of Observations: 512
Number of Groups: 13
> AICc(Dlme00)
[1] 7162.689
> ## Status included
> Dlme01<-lme(fixed = NEAR_DIST_Botnia ~ Status , random = ~ 1|Site, data=
NaDist)
> summary(Dlme01)
Linear mixed-effects model fit by REML
 Data: NaDist
       AIC      BIC    logLik
  7142.171 7167.554 -3565.085

Random effects:
 Formula: ~1 | Site
        (Intercept) Residual
StdDev:    186.9849 257.3497

Fixed effects: NEAR_DIST_Botnia ~ Status
                      Value Std.Error  DF  t-value p-value
(Intercept)        372.7429  59.74608 496 6.238785  0.0000
StatusConstruction  13.1606  34.21712 496 0.384620  0.7007
StatusReady         19.6057  35.18869 496 0.557159  0.5777
StatusTrafic        14.3106  33.68114 496 0.424886  0.6711
 Correlation:
                   (Intr) SttsCn SttsRd
StatusConstruction -0.260
StatusReady        -0.327  0.451
StatusTrafic       -0.272  0.439  0.510

Standardized Within-Group Residuals:
       Min         Q1        Med         Q3        Max
-2.2417162 -0.6425096 -0.0203046  0.6600936  3.0951515

Number of Observations: 512
Number of Groups: 13
> AICc(Dlme01)
[1] 7142.337
```

### Eurasian Skylark

```
> AaDist<-BotDist[BotDist$Species=="Aa",]
> ## Empty model
> Dlme00<-lme(fixed = NEAR_DIST_Botnia ~ 1 , random = ~ 1|Site, data=AaDis
t)
> summary(Dlme00)
Linear mixed-effects model fit by REML
 Data: AaDist
       AIC      BIC    logLik
  6627.217 6639.751 -3310.609

Random effects:
 Formula: ~1 | Site
        (Intercept) Residual
StdDev:    197.6241 225.3438

Fixed effects: NEAR_DIST_Botnia ~ 1
               Value Std.Error  DF  t-value p-value
(Intercept) 341.0679  65.82334 472 5.181565       0

Standardized Within-Group Residuals:
        Min          Q1         Med          Q3         Max
-2.16854670 -0.58414097 -0.08547115  0.61214514  3.76949041

Number of Observations: 483
Number of Groups: 11
> AICc(Dlme00)
[1] 6627.267
> ## Status included
> Dlme01<-lme(fixed = NEAR_DIST_Botnia ~ Status , random = ~ 1|Site, data=
AaDist)
> summary(Dlme01)
Linear mixed-effects model fit by REML
 Data: AaDist
      AIC     BIC   logLik
  6604.02 6629.05 -3296.01

Random effects:
 Formula: ~1 | Site
        (Intercept) Residual
StdDev:    201.4352 225.1646

Fixed effects: NEAR_DIST_Botnia ~ Status
                      Value Std.Error  DF   t-value p-value
(Intercept)        335.1788  69.74759 469  4.805597  0.0000
StatusConstruction  13.1301  28.92666 469  0.453911  0.6501
StatusReady        -14.5305  35.20255 469 -0.412768  0.6800
StatusTrafic        47.0578  30.93694 469  1.521088  0.1289
 Correlation:
                   (Intr) SttsCn SttsRd
StatusConstruction -0.209
StatusReady        -0.253  0.459
StatusTrafic       -0.183  0.413  0.368

Standardized Within-Group Residuals:
        Min          Q1         Med          Q3         Max
-2.26867283 -0.59660509 -0.07413254  0.60045788  3.80328891

Number of Observations: 483
Number of Groups: 11
> AICc(Dlme01)
[1] 6604.196
```

### Meadow Pipit

```
## Meadow Pipit
> ApDist<-BotDist[BotDist$Species=="Ap",]
> ## Empty model
> Dlme00<-lme(fixed = NEAR_DIST_Botnia ~ 1 , random = ~ 1|Site, data=ApDis
t)
> summary(Dlme00)
Linear mixed-effects model fit by REML
 Data: ApDist
      AIC      BIC   logLik
  979.526 986.4785 -486.763

Random effects:
 Formula: ~1 | Site
        (Intercept) Residual
StdDev:     224.879  141.135

Fixed effects: NEAR_DIST_Botnia ~ 1
               Value Std.Error DF  t-value p-value
(Intercept) 298.7797  96.61891 70 3.092352  0.0029

Standardized Within-Group Residuals:
       Min         Q1        Med         Q3        Max
-2.0512394 -0.4793522 -0.2090073  0.6265341  2.8623878

Number of Observations: 76
Number of Groups: 6
> AICc(Dlme00)
[1] 979.8594
> ## Status included
> Dlme01<-lme(fixed = NEAR_DIST_Botnia ~ Status , random = ~ 1|Site, data=
ApDist)
> summary(Dlme01)
Linear mixed-effects model fit by REML
 Data: ApDist
       AIC      BIC    logLik
  956.5174 970.1774 -472.2587

Random effects:
 Formula: ~1 | Site
        (Intercept) Residual
StdDev:    221.2077 143.8918

Fixed effects: NEAR_DIST_Botnia ~ Status
                      Value Std.Error DF   t-value p-value
(Intercept)        320.7227 100.23008 67  3.199865  0.0021
StatusConstruction -25.0075  49.73438 67 -0.502820  0.6167
StatusReady        -24.6691  47.97726 67 -0.514182  0.6088
StatusTrafic       -33.9048  56.34159 67 -0.601772  0.5494
 Correlation:
                   (Intr) SttsCn SttsRd
StatusConstruction -0.216
StatusReady        -0.218  0.460
StatusTrafic       -0.281  0.427  0.447

Standardized Within-Group Residuals:
       Min         Q1        Med         Q3        Max
-1.9475411 -0.5353112 -0.2384340  0.6702618  2.6981555

Number of Observations: 76
Number of Groups: 6
> AICc(Dlme01)
[1] 957.7348
```

### Barn Swallow

```
## Barn Swallow
> HrDist<-BotDist[BotDist$Species=="Hr",]
> ## Empty model
> Dlme00<-lme(fixed = NEAR_DIST_Botnia ~ 1 , random = ~ 1|Site, data=HrDis
t)
> summary(Dlme00)
Linear mixed-effects model fit by REML
 Data: HrDist
       AIC      BIC    logLik
  9603.494 9617.211 -4798.747

Random effects:
 Formula: ~1 | Site
        (Intercept) Residual
StdDev:    200.8513 192.2223

Fixed effects: NEAR_DIST_Botnia ~ 1
               Value Std.Error  DF  t-value p-value
(Intercept) 390.8909  59.36308 704 6.584747       0

Standardized Within-Group Residuals:
        Min          Q1         Med          Q3         Max
-4.23744594 -0.61748458 -0.07179096  0.54566881  3.87025076

Number of Observations: 716
Number of Groups: 12
> AICc(Dlme00)
[1] 9603.528
> ## Status included
> Dlme01<-lme(fixed = NEAR_DIST_Botnia ~ Status , random = ~ 1|Site, data=
HrDist)
> summary(Dlme01)
Linear mixed-effects model fit by REML
 Data: HrDist
       AIC      BIC    logLik
  9581.747 9609.155 -4784.873

Random effects:
 Formula: ~1 | Site
        (Intercept) Residual
StdDev:    200.2101 192.0551

Fixed effects: NEAR_DIST_Botnia ~ Status
                      Value Std.Error  DF   t-value p-value
(Intercept)        408.5715  61.49267 701  6.644232  0.0000
StatusConstruction -44.0155  24.48887 701 -1.797369  0.0727
StatusReady        -22.0607  22.55510 701 -0.978081  0.3284
StatusTrafic        -3.4209  22.76057 701 -0.150299  0.8806
 Correlation:
                   (Intr) SttsCn SttsRd
StatusConstruction -0.204
StatusReady        -0.249  0.556
StatusTrafic       -0.227  0.521  0.624

Standardized Within-Group Residuals:
        Min          Q1         Med          Q3         Max
-4.22746232 -0.62015928 -0.08705626  0.58276291  3.80030880

Number of Observations: 716
Number of Groups: 12
> AICc(Dlme01)
[1] 9581.865
```

### Common Starling

```
## Common Starling
> SvDist<-BotDist[BotDist$Species=="Sv",]
> ## Empty model
> Dlme00<-lme(fixed = NEAR_DIST_Botnia ~ 1 , random = ~ 1|Site, data=SvDis
t)
> summary(Dlme00)
Linear mixed-effects model fit by REML
 Data: SvDist
      AIC     BIC    logLik
  1083.51 1090.73 -538.7549

Random effects:
 Formula: ~1 | Site
        (Intercept) Residual
StdDev:    223.4093 149.3514

Fixed effects: NEAR_DIST_Botnia ~ 1
               Value Std.Error DF  t-value p-value
(Intercept) 445.6389  82.97094 75 5.371024       0

Standardized Within-Group Residuals:
        Min          Q1         Med          Q3         Max
-2.72146859 -0.51081577  0.05805498  0.59874374  2.14282943

Number of Observations: 83
Number of Groups: 8
> AICc(Dlme00)
[1] 1083.814
> ## Status included
> Dlme01<-lme(fixed = NEAR_DIST_Botnia ~ Status , random = ~ 1|Site, data=
SvDist)
> summary(Dlme01)
Linear mixed-effects model fit by REML
 Data: SvDist
       AIC      BIC    logLik
  1052.033 1066.249 -520.0163

Random effects:
 Formula: ~1 | Site
        (Intercept) Residual
StdDev:    230.2754 143.4722

Fixed effects: NEAR_DIST_Botnia ~ Status
                       Value Std.Error DF   t-value p-value
(Intercept)         523.9500  91.47060 72  5.728069  0.0000
StatusConstruction -135.8122  59.90977 72 -2.266945  0.0264
StatusReady         -89.7697  56.07866 72 -1.600783  0.1138
StatusTrafic       -141.7822  54.59826 72 -2.596827  0.0114
 Correlation:
                   (Intr) SttsCn SttsRd
StatusConstruction -0.247
StatusReady        -0.347  0.447
StatusTrafic       -0.306  0.415  0.716

Standardized Within-Group Residuals:
         Min           Q1          Med           Q3          Max
-2.671752773 -0.584040660 -0.001807768  0.708975930  1.900124243

Number of Observations: 83
Number of Groups: 8
> AICc(Dlme01)
[1] 1053.138
```

**Ortolan Bunting**
**Sample size too small. See Supplementary material Dist4 for details.**

```
## Ortolan Bunting
> EhDist<-BotDist[BotDist$Species=="Eh",]
> ## Empty model
> Dlme00<-lme(fixed = NEAR_DIST_Botnia ~ 1 , random = ~ 1|Site, data=EhDis
t)
> summary(Dlme00)
Linear mixed-effects model fit by REML
 Data: EhDist
       AIC      BIC    logLik
  53.43052 52.25883 -23.71526

Random effects:
 Formula: ~1 | Site
        (Intercept) Residual
StdDev:    97.37233  14.1305

Fixed effects: NEAR_DIST_Botnia ~ 1
               Value Std.Error DF  t-value p-value
(Intercept) 158.4462  69.09388  4 2.293202  0.0836

Standardized Within-Group Residuals:
       Min         Q1        Med         Q3        Max
-1.6223035 -0.1485051  0.1616524  0.4645309  0.9853992

Number of Observations: 6
Number of Groups: 2
> AICc(Dlme00)
[1] 65.43052
> ## Status included
> Dlme01<-lme(fixed = NEAR_DIST_Botnia ~ Status , random = ~ 1|Site, data=
EhDist)
> summary(Dlme01)
Linear mixed-effects model fit by REML
 Data: EhDist
       AIC      BIC    logLik
  47.74858 45.29376 -19.87429

Random effects:
 Formula: ~1 | Site
        (Intercept) Residual
StdDev:    96.96875 16.31271

Fixed effects: NEAR_DIST_Botnia ~ Status
                       Value Std.Error DF   t-value p-value
(Intercept)        158.16170  68.97015  3 2.2931904  0.1056
StatusConstruction   1.70731  19.95561  3 0.0855554  0.9372
 Correlation:
                   (Intr)
StatusConstruction -0.048

Standardized Within-Group Residuals:
       Min         Q1        Med         Q3        Max
-1.4151757 -0.1100233  0.1324779  0.4061637  0.8436846

Number of Observations: 6
Number of Groups: 2
> AICc(Dlme01)
[1] 87.74858
```

### Little Ringed Plover
### Too few cases (only two status classes)

```
## Little Ringed Plover
> CdDist<-BotDist[BotDist$Species=="Cd",]
> ## Empty model
> Dlme00<-lme(fixed = NEAR_DIST_Botnia ~ 1 , random = ~ 1|Site, data=CdDis
t)
> summary(Dlme00)
Linear mixed-effects model fit by REML
 Data: CdDist
       AIC      BIC    logLik
  203.8035 206.3031 -98.90173

Random effects:
 Formula: ~1 | Site
        (Intercept) Residual
StdDev:    6.872718 74.47058

Fixed effects: NEAR_DIST_Botnia ~ 1
               Value Std.Error DF  t-value p-value
(Intercept) 113.1398  17.86439 13 6.333258       0

Standardized Within-Group Residuals:
       Min         Q1        Med         Q3        Max
-1.1746761 -0.9082618 -0.2726545  0.8856288  1.5731738

Number of Observations: 18
Number of Groups: 5
> AICc(Dlme00)
[1] 205.5177
> ## Status included
> Dlme01<-lme(fixed = NEAR_DIST_Botnia ~ Status , random = ~ 1|Site, data=
CdDist)
> summary(Dlme01)
Linear mixed-effects model fit by REML
 Data: CdDist
       AIC     BIC    logLik
  192.5207 195.611 -92.26033

Random effects:
 Formula: ~1 | Site
        (Intercept) Residual
StdDev: 0.006632374 70.72466

Fixed effects: NEAR_DIST_Botnia ~ Status
                Value Std.Error DF   t-value p-value
(Intercept)  231.7276  70.72466 12  3.276475  0.0066  = Construction
StatusReady -125.5840  72.77508 12 -1.725646  0.1100
 Correlation:
            (Intr)
StatusReady -0.972

Standardized Within-Group Residuals:
       Min         Q1        Med         Q3        Max
-1.1333882 -0.8575461 -0.1886927  0.6599714  1.7314529

Number of Observations: 18
Number of Groups: 5
> AICc(Dlme01)
[1] 195.5976
```

### Common Snipe

```
## Common Snipe
> GgDist<-BotDist[BotDist$Species=="Gg",]
> ## Empty model
> Dlme00<-lme(fixed = NEAR_DIST_Botnia ~ 1 , random = ~ 1|Site, data=GgDis
t)
> summary(Dlme00)
Linear mixed-effects model fit by REML
 Data: GgDist
       AIC      BIC    logLik
  942.5448 949.3328 -468.2724

Random effects:
 Formula: ~1 | Site
        (Intercept) Residual
StdDev:    286.7027 142.7737

Fixed effects: NEAR_DIST_Botnia ~ 1
               Value Std.Error DF  t-value p-value
(Intercept) 278.1736  94.52238 62 2.942939  0.0046

Standardized Within-Group Residuals:
        Min          Q1         Med          Q3         Max
-2.63777657 -0.56938926 -0.09197647  0.31958313  2.31161347

Number of Observations: 72
Number of Groups: 10
> AICc(Dlme00)
[1] 942.8977
> ## Status included
> Dlme01<-lme(fixed = NEAR_DIST_Botnia ~ Status , random = ~ 1|Site, data=
GgDist)
> summary(Dlme01)
Linear mixed-effects model fit by REML
 Data: GgDist
      AIC   BIC    logLik
  915.083 928.4 -451.5415

Random effects:
 Formula: ~1 | Site
        (Intercept) Residual
StdDev:     274.707 141.5627

Fixed effects: NEAR_DIST_Botnia ~ Status
                       Value Std.Error DF   t-value p-value
(Intercept)         320.5403 100.69189 59  3.183378  0.0023
StatusConstruction  -36.9006  62.47469 59 -0.590648  0.5570
StatusReady        -101.3117  59.05335 59 -1.715596  0.0915
StatusTrafic         -9.5377  56.49742 59 -0.168816  0.8665
 Correlation:
                   (Intr) SttsCn SttsRd
StatusConstruction -0.358
StatusReady        -0.387  0.626
StatusTrafic       -0.390  0.670  0.681

Standardized Within-Group Residuals:
       Min         Q1        Med         Q3        Max
-2.7334297 -0.5206156 -0.1528158  0.3676452  2.2509204

Number of Observations: 72
Number of Groups: 10
> AICc(Dlme01)
[1] 916.3753
```

### Green Sandpiper

```
## Green Sandpiper
> ToDist<-BotDist[BotDist$Species=="To",]
> ## Empty model
> Dlme00<-lme(fixed = NEAR_DIST_Botnia ~ 1 , random = ~ 1|Site, data=ToDis
t)
> summary(Dlme00)
Linear mixed-effects model fit by REML
 Data: ToDist
       AIC      BIC  logLik
  408.4199 412.3074 -201.21

Random effects:
 Formula: ~1 | Site
        (Intercept) Residual
StdDev:    158.6572 367.2928

Fixed effects: NEAR_DIST_Botnia ~ 1
               Value Std.Error DF  t-value p-value
(Intercept) 323.8639  88.71014 18 3.650811  0.0018

Standardized Within-Group Residuals:
       Min         Q1        Med         Q3        Max
-1.3928304 -0.6039837 -0.2513598  0.2724166  2.7058288

Number of Observations: 28
Number of Groups: 10
> AICc(Dlme00)
[1] 409.4199
> ## Status included
> Dlme01<-lme(fixed = NEAR_DIST_Botnia ~ Status , random = ~ 1|Site, data=
ToDist)
> summary(Dlme01)
Linear mixed-effects model fit by REML
 Data: ToDist
      AIC      BIC    logLik
  375.749 382.8174 -181.8745

Random effects:
 Formula: ~1 | Site
        (Intercept) Residual
StdDev:    154.4894 381.7217

Fixed effects: NEAR_DIST_Botnia ~ Status
                       Value Std.Error DF    t-value p-value
(Intercept)         329.7558  157.5653 15  2.0928193  0.0538
StatusConstruction -181.9476  239.6288 15 -0.7592893  0.4594
StatusReady          54.6304  195.2328 15  0.2798218  0.7834
StatusTrafic         57.4462  241.3724 15  0.2379984  0.8151
 Correlation:
                   (Intr) SttsCn SttsRd
StatusConstruction -0.631
StatusReady        -0.751  0.532
StatusTrafic       -0.547  0.380  0.451

Standardized Within-Group Residuals:
       Min         Q1        Med         Q3        Max
-1.2857380 -0.5824287 -0.1451811  0.2976837  2.5148779

Number of Observations: 28
Number of Groups: 10
> AICc(Dlme01)
[1] 379.749
```

### Western Yellow Wagtail

```
## Western Yellow Wagtail
> MfDist<-BotDist[BotDist$Species=="Mf",]
> ## Empty model
> Dlme00<-lme(fixed = NEAR_DIST_Botnia ~ 1 , random = ~ 1|Site, data=MfDis
t)
> summary(Dlme00)
Linear mixed-effects model fit by REML
 Data: MfDist
       AIC      BIC    logLik
  2727.318 2737.242 -1360.659

Random effects:
 Formula: ~1 | Site
        (Intercept) Residual
StdDev:    204.8601 191.7257

Fixed effects: NEAR_DIST_Botnia ~ 1
               Value Std.Error  DF  t-value p-value
(Intercept) 384.0458  79.56773 196 4.826652       0

Standardized Within-Group Residuals:
       Min         Q1        Med         Q3        Max
-3.4487774 -0.5922780 -0.1990709  0.6936595  2.1816253

Number of Observations: 203
Number of Groups: 7
> AICc(Dlme00)
[1] 2727.438
> ## Status included
> Dlme01<-lme(fixed = NEAR_DIST_Botnia ~ Status , random = ~ 1|Site, data=
MfDist)
> summary(Dlme01)
Linear mixed-effects model fit by REML
 Data: MfDist
       AIC      BIC    logLik
  2697.552 2717.311 -1342.776

Random effects:
 Formula: ~1 | Site
        (Intercept) Residual
StdDev:    206.1031  189.639

Fixed effects: NEAR_DIST_Botnia ~ Status
                       Value Std.Error  DF   t-value p-value
(Intercept)         527.7769  97.20022 193  5.429791  0.0000
StatusConstruction -167.3500  63.71434 193 -2.626568  0.0093
StatusReady        -153.3327  61.35567 193 -2.499080  0.0133
StatusTrafic       -142.3549  70.46590 193 -2.020196  0.0447
 Correlation: 
                   (Intr) SttsCn SttsRd
StatusConstruction -0.510              
StatusReady        -0.555  0.819       
StatusTrafic       -0.480  0.736  0.742

Standardized Within-Group Residuals:
       Min         Q1        Med         Q3        Max
-3.4869436 -0.6255617 -0.1671672  0.6424730  2.2304907

Number of Observations: 203
Number of Groups: 7
> AICc(Dlme01)
[1] 2697.98
```

### Whinchat

```
## Whinchat
> SrDist<-BotDist[BotDist$Species=="Sr",]
> ## Empty model
> Dlme00<-lme(fixed = NEAR_DIST_Botnia ~ 1 , random = ~ 1|Site, data=SrDis
t)
> summary(Dlme00)
Linear mixed-effects model fit by REML
 Data: SrDist
       AIC      BIC    logLik
  4620.225 4631.694 -2307.113

Random effects:
 Formula: ~1 | Site
        (Intercept) Residual
StdDev:    107.2664 214.1919

Fixed effects: NEAR_DIST_Botnia ~ 1
               Value Std.Error  DF  t-value p-value
(Intercept) 308.9952  33.92609 326 9.107891       0

Standardized Within-Group Residuals:
       Min         Q1        Med         Q3        Max
-2.0145305 -0.6553652 -0.0591114  0.6095372  3.5395786

Number of Observations: 339
Number of Groups: 13
> AICc(Dlme00)
[1] 4620.297
> ## Status included
> Dlme01<-lme(fixed = NEAR_DIST_Botnia ~ Status , random = ~ 1|Site, data=
SrDist)
> summary(Dlme01)
Linear mixed-effects model fit by REML
 Data: SrDist
       AIC      BIC    logLik
  4593.794 4616.678 -2290.897

Random effects:
 Formula: ~1 | Site
        (Intercept) Residual
StdDev:    115.3373  212.807

Fixed effects: NEAR_DIST_Botnia ~ Status
                      Value Std.Error  DF   t-value p-value
(Intercept)        371.1135  45.80975 323  8.101191  0.0000
StatusConstruction -41.5062  41.92362 323 -0.990044  0.3229
StatusReady        -88.2531  39.88467 323 -2.212707  0.0276
StatusTrafic       -85.8842  39.78627 323 -2.158639  0.0316
 Correlation:
                   (Intr) SttsCn SttsRd
StatusConstruction -0.463
StatusReady        -0.574  0.595
StatusTrafic       -0.552  0.574  0.709

Standardized Within-Group Residuals:
        Min          Q1         Med          Q3         Max
-1.92533419 -0.68213361 -0.05000712  0.56528227  3.40230808

Number of Observations: 339
Number of Groups: 13
> AICc(Dlme01)
[1] 4594.047
```

### Red-backed Shrike
### Too few?

```
## Red-backed Shrike
> LcDist<-BotDist[BotDist$Species=="Lc",]
> ## Empty model
> Dlme00<-lme(fixed = NEAR_DIST_Botnia ~ 1 , random = ~ 1|Site, data=LcDis
t)
> summary(Dlme00)
Linear mixed-effects model fit by REML
 Data: LcDist 
       AIC      BIC    logLik
  185.3265 187.2437 -89.66327

Random effects:
 Formula: ~1 | Site
        (Intercept) Residual
StdDev:    200.2162  92.4556

Fixed effects: NEAR_DIST_Botnia ~ 1 
              Value Std.Error DF  t-value p-value
(Intercept) 233.289  93.75113 10 2.488386  0.0321

Standardized Within-Group Residuals:
       Min         Q1        Med         Q3        Max 
-1.8939965 -0.4571189  0.1689810  0.5399027  1.2900244 

Number of Observations: 15
Number of Groups: 5 
> AICc(Dlme00)
[1] 187.5084
> ## Status included
> Dlme01<-lme(fixed = NEAR_DIST_Botnia ~ Status , random = ~ 1|Site, data=
LcDist)
> summary(Dlme01)
Linear mixed-effects model fit by REML
 Data: LcDist 
       AIC     BIC    logLik
  148.2666 150.654 -68.13331

Random effects:
 Formula: ~1 | Site
        (Intercept) Residual
StdDev:    52.91367  93.7105

Fixed effects: NEAR_DIST_Botnia ~ Status 
                      Value Std.Error DF   t-value p-value
(Intercept)        158.7274  82.29878  9  1.928672  0.0859
StatusConstruction -27.3465  95.20723  9 -0.287231  0.7804
StatusReady        -17.9130  96.00584  2 -0.186582  0.8692
StatusTrafic       478.3265 135.47917  2  3.530627  0.0717
 Correlation: 
                   (Intr) SttsCn SttsRd
StatusConstruction -0.728              
StatusReady        -0.857  0.624       
StatusTrafic       -0.607  0.442  0.521

Standardized Within-Group Residuals:
        Min          Q1         Med          Q3         Max 
-1.46994284 -0.50252489  0.02826636  0.25410301  1.43704329 

Number of Observations: 15
Number of Groups: 5 
> AICc(Dlme01)
[1] 158.7666
```

#### Common Rosefinch

#### Too few?

```
## Common Rosefinch
> CeDist<-BotDist[BotDist$Species=="Ce",]
> ## Empty model
> Dlme00<-lme(fixed = NEAR_DIST_Botnia ~ 1 , random = ~ 1|Site, data=CeDis
t)
> summary(Dlme00)
Linear mixed-effects model fit by REML
 Data: CeDist
       AIC      BIC   logLik
  564.1861 569.3268 -279.093

Random effects:
 Formula: ~1 | Site
        (Intercept) Residual
StdDev:  0.03972304 209.0613

Fixed effects: NEAR_DIST_Botnia ~ 1
               Value Std.Error DF  t-value p-value
(Intercept) 304.9774  32.25887 37 9.454062       0

Standardized Within-Group Residuals:
       Min         Q1        Med         Q3        Max
-1.3734490 -0.6905885 -0.1365209  0.6626389  2.4642323

Number of Observations: 42
Number of Groups: 5
> AICc(Dlme00)
[1] 564.8177
> ## Status included
> Dlme01<-lme(fixed = NEAR_DIST_Botnia ~ Status , random = ~ 1|Site, data=
CeDist)
> summary(Dlme01)
Linear mixed-effects model fit by REML
 Data: CeDist
       AIC     BIC    logLik
  534.5985 544.424 -261.2992

Random effects:
 Formula: ~1 | Site
        (Intercept) Residual
StdDev:  0.02128036 213.7959

Fixed effects: NEAR_DIST_Botnia ~ Status
                       Value Std.Error DF    t-value p-value
(Intercept)         341.4154  151.1765 34  2.2583889  0.0305
StatusConstruction  -67.3675  174.5636 34 -0.3859192  0.7020
StatusReady         -21.6469  155.9770 34 -0.1387827  0.8904
StatusTrafic       -151.7127  195.1681 34 -0.7773438  0.4423
 Correlation: 
                   (Intr) SttsCn SttsRd
StatusConstruction -0.866              
StatusReady        -0.969  0.839       
StatusTrafic       -0.775  0.671  0.751

Standardized Within-Group Residuals:
       Min         Q1        Med         Q3        Max
-1.4056783 -0.6797631 -0.1111570  0.6586806  2.3404778

Number of Observations: 42
Number of Groups: 5
> AICc(Dlme01)
[1] 536.9985
```
