## Supplementary material for "Are railways detrimental to bird populations? A BDACI study on the construction of the Bothnia Line Railway": Trends of numbers of territories for Control, Impact and all sites

### Supplementary material **Terr1**

### Temporal trends for numbers of territories of individual study species in Impact sites (red), Control sites (blue) and all sites combined (black)

Most correlation tests generated warning messages due to ties (data points with the same Y-value). The warning texts were omitted from the output text for readability, but the tau value estimates marked with a * character. For models with ties, exact P-values could not be calculated and the estimated P-values should be treated with caution.

Jitter along the x-axis added to separate overlapping points if needed.

*Please note that the numbers of territories are very low for some species and correlations and trends for those species should be interpreted with care.*

National trend data 1996-2015 from the “Monitoring population changes of birds in Sweden” programme are included for each species. Source: Green, M., Haas, F. & Lindström, Å. 2016. Övervakning av fåglarnas populationsutveckling. Årsrapport för 2015. Rapport, Biologiska institutionen, Lunds universitet. 88 pp.

Contents


| Species | page |
|---|---|
| Barn Swallow | 2 |
| Eurasian Curlew | 3 |
| Eurasian Skylark | 4 |
| Common Rosefinch | 5 |
| Common Snipe | 6 |
| Common Starling | 7 |
| Green Sandpiper | 8 |
| Little Ringed Plover | 9 |
| Meadow Pipit | 10 |
| Northern Lapwing | 11 |
| Ortolan Bunting | 12 |
| Red-backed Shrike | 13 |
| Western Yellow Wagtail | 14 |
| Whinchat | 15 |

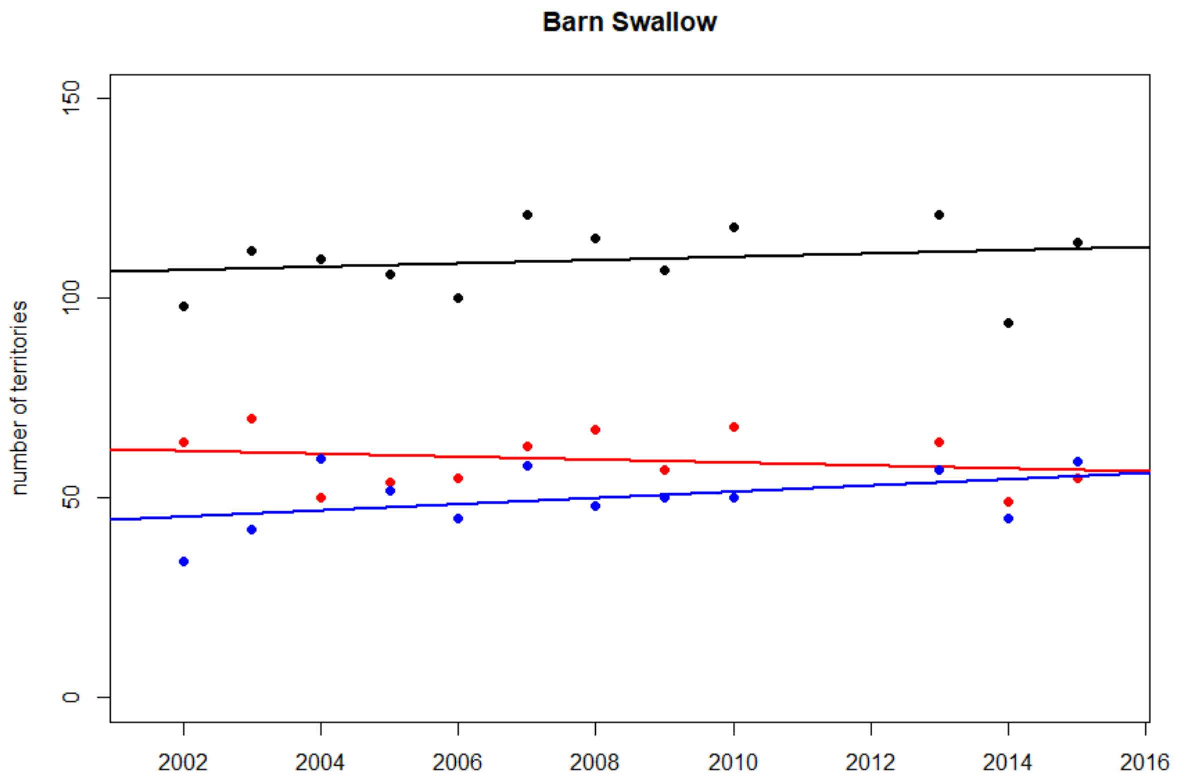


Total yearly numbers of territories of Barn Swallow in Impact sites (red), in Control sites (blue) and in all sites (black)

**Swedish national trend 1996-2015: +1.5 % per year (***)**

**Impact sites**
Kendall's rank correlation tau = -0.062
z = -0.276, p-value = 0.783

**Control sites**
Kendall's rank correlation tau = 0.277
z = 1.240, p-value = 0.215

**All sites**
Kendall's rank correlation tau = 0.198
z = 0.894, p-value = 0.372

**Difference between Impact and Control sites**
Kendall's rank correlation tau = -0.290
z = -1.306, p-value = 0.1916

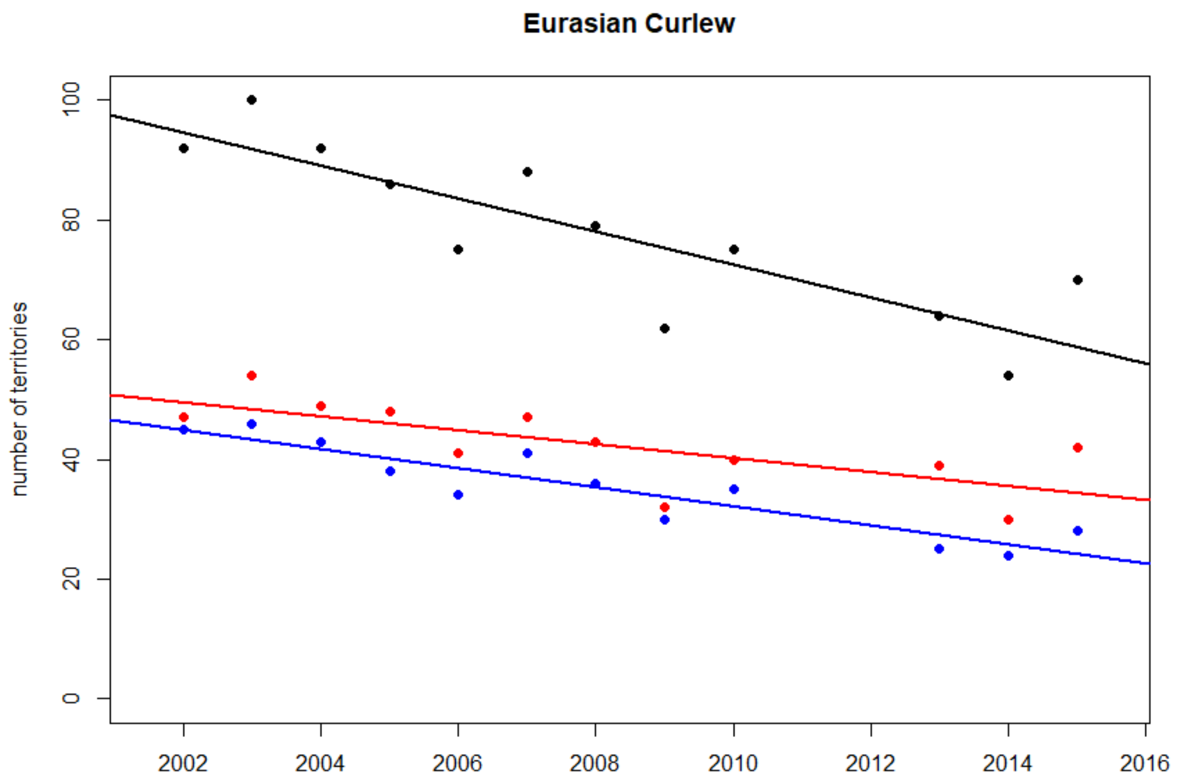


Total yearly numbers of territories of Eurasian Curlew in Impact sites (red), in Control sites (blue) and in all sites (black)

**Swedish national trend 1996-2015: -2.0 % per year (***)**

**Impact sites**
Kendall's rank correlation tau = -0.626
z = -2.818, p-value = 0.005

**Control sites**
Kendall's rank correlation tau = -0.758
T = 8, p-value < 0.001

**All sites**
Kendall's rank correlation tau = -0.708
z = -3.169, p-value = 0.0015

**Difference between Impact and Control sites**
Kendall's rank correlation tau = 0.159
z = 0.697, p-value = 0.486

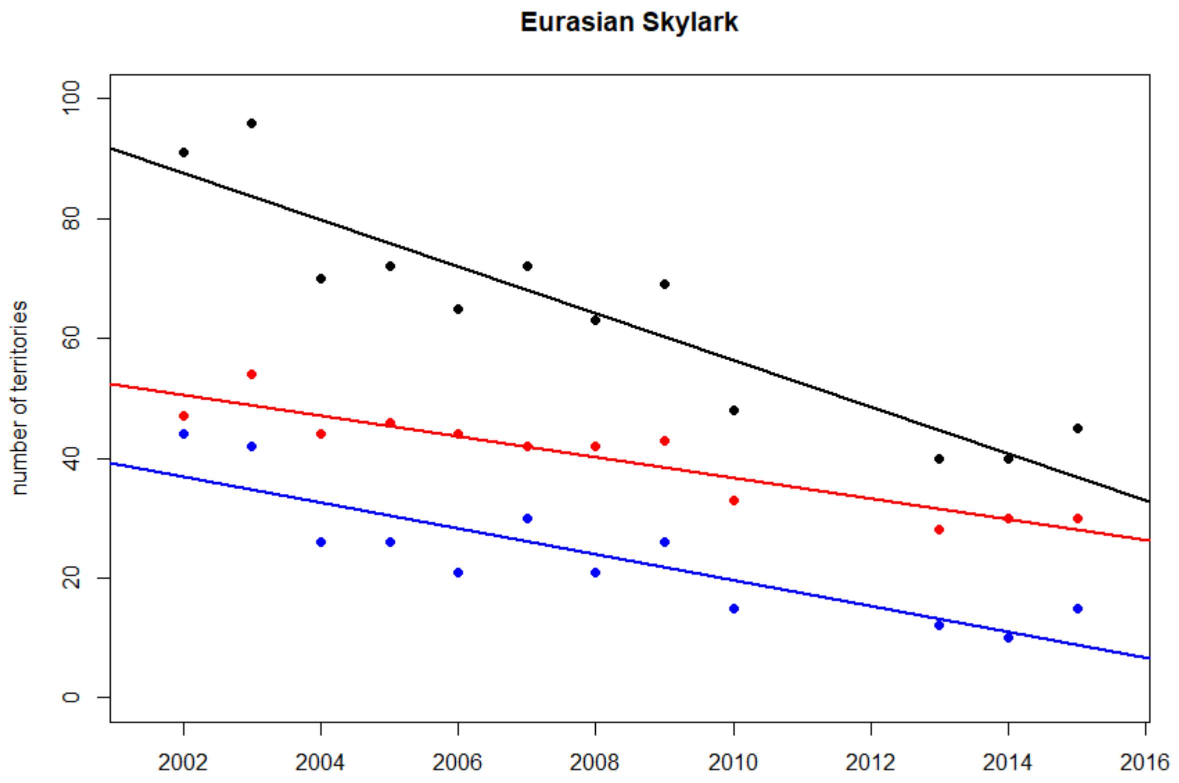


Total yearly numbers of territories of Eurasian Skylark in Impact sites (red), in Control sites (blue) and in all sites (black)

**Swedish national trend 1996-2015: -1.7 % per year (***)**

**Impact sites**
Kendall's rank correlation tau = -0.791
z = -3.522, p-value < 0.001***

**Control sites**
Kendall's rank correlation tau = -0.741
z = -3.267, p-value = 0.0011**

**All sites**
Kendall's rank correlation tau = -0.739
z = -3.307, p-value < 0.001***

**Difference between Impact and Control sites**
Kendall's rank correlation tau = 0.140
z = 0.622, p-value = 0.534

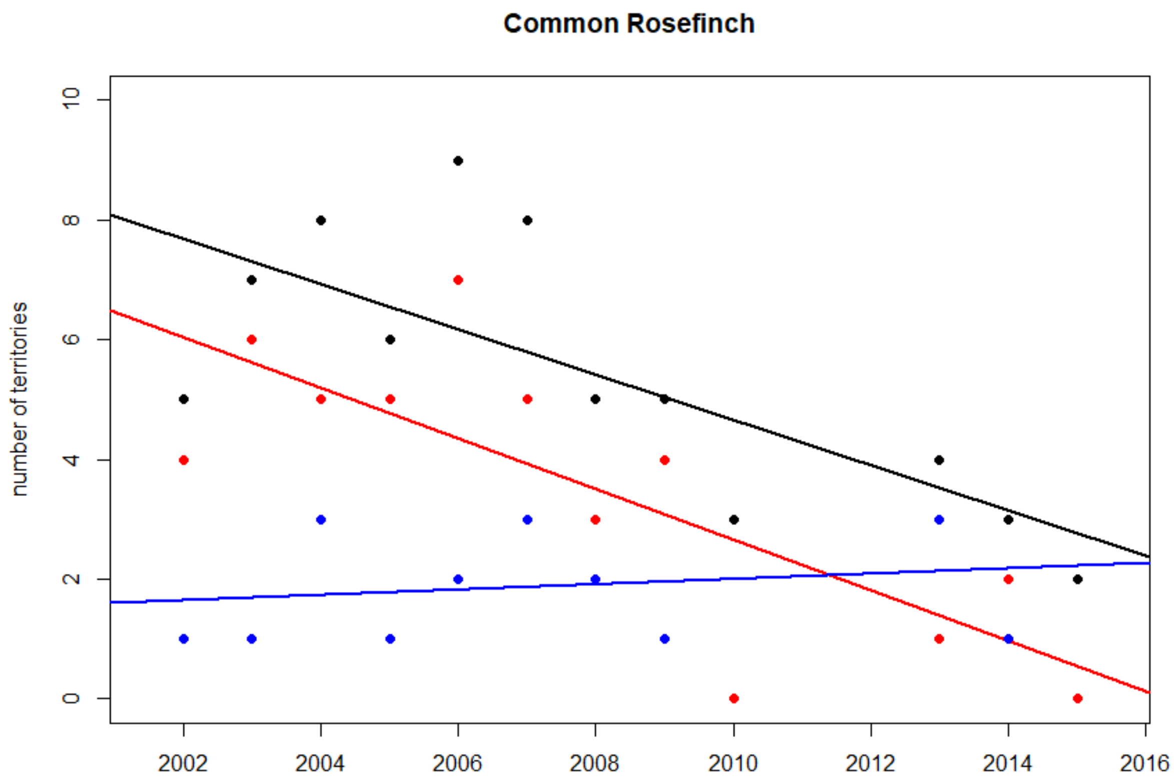


Total yearly numbers of territories of Common Rosefinch in Impact sites (red), in Control sites (blue) and in all sites (black)

**Impact sites**
Kendall's rank correlation tau = -0.583
z = -2.572, p-value = 0.0101*

**Control sites**
Kendall's rank correlation tau = 0.198
z = 0.812, p-value = 0.417

**All sites**
Kendall's rank correlation tau = -0.583
z = -2.572, p-value = 0.0101*

**Difference between Impact and Control sites**
Kendall's rank correlation tau = -0.552
z = -2.429, p-value = 0.015*

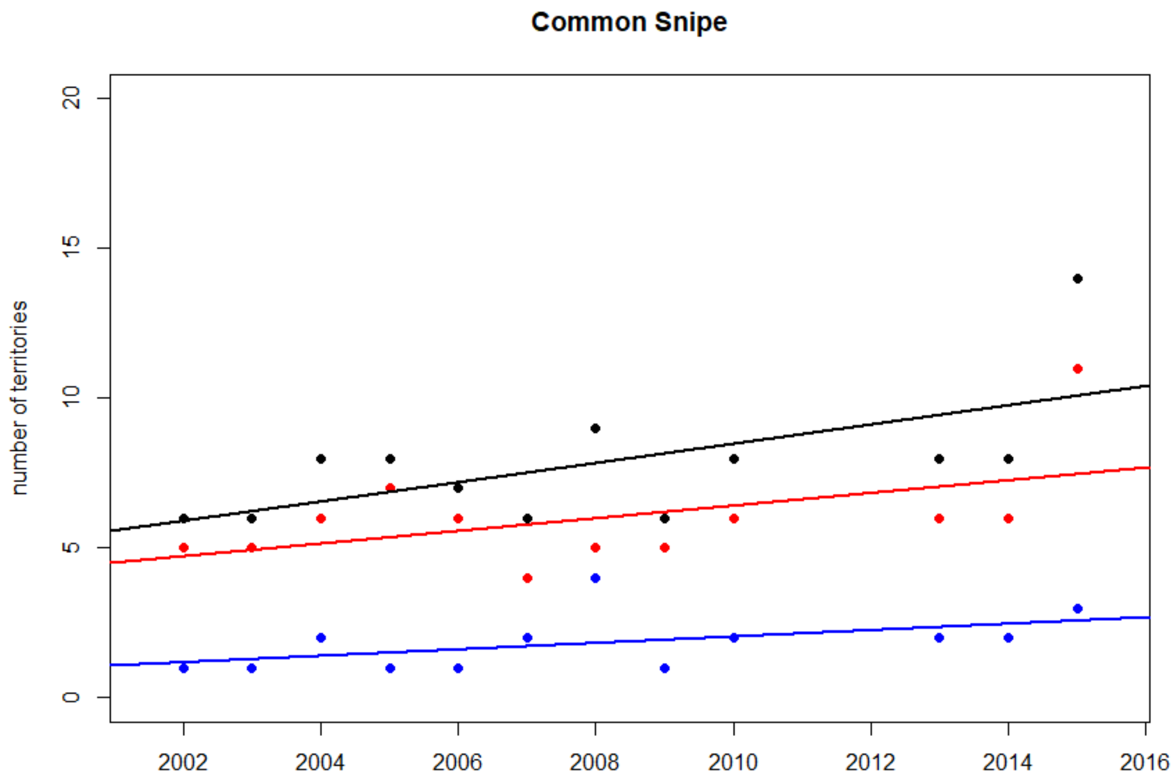


Total yearly numbers of territories of Common Snipe in Impact sites (red), in Control sites (blue) and in all sites (black)

**Swedish national trend 1996-2015: +1.1 % per year (**)**

**Impact sites**
Kendall's rank correlation tau = 0.348
z = 1.461, p-value = 0.144

**Control sites**
Kendall's rank correlation tau = 0.508
z = 2.091, p-value = 0.037*

**All sites**
Kendall's rank correlation tau = 0.453
z = 1.900, p-value = 0.058

**Difference between Impact and Control sites**
Kendall's rank correlation tau = 0.092
z = 0.385, p-value = 0.700

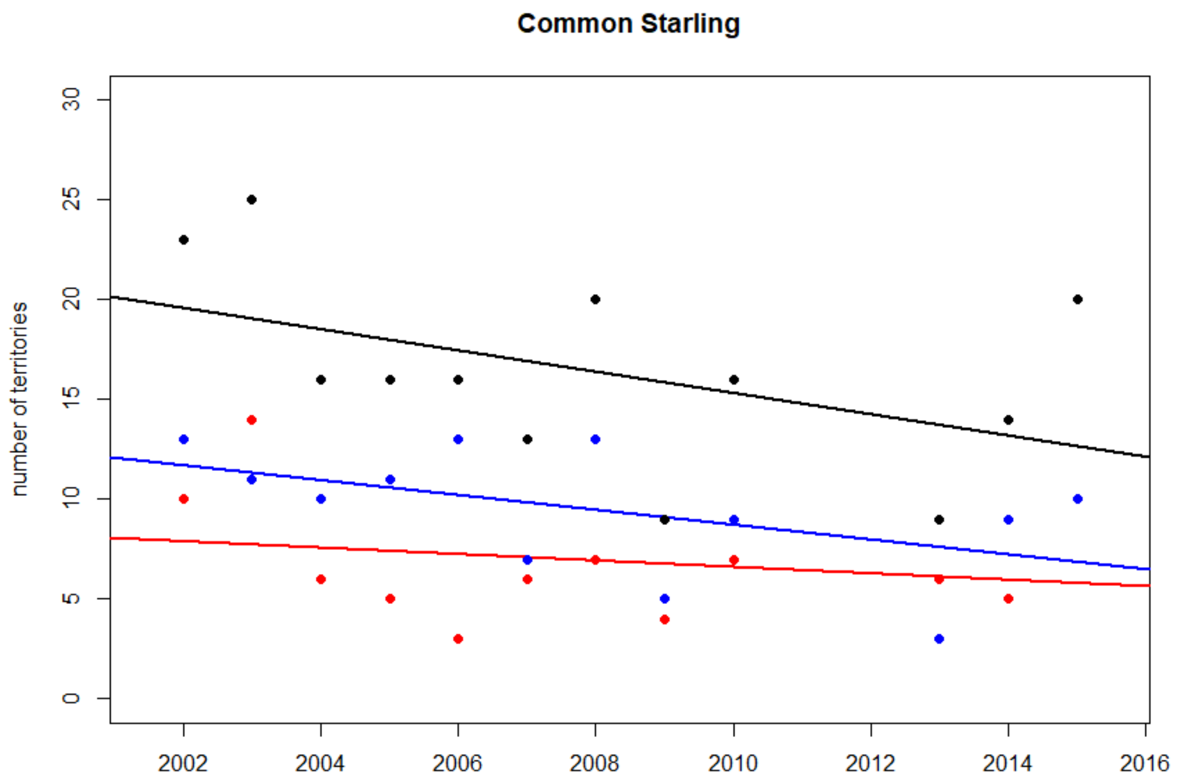


Total yearly numbers of territories of Common Starling in Impact sites (red), in Control sites (blue) and in all sites (black)

**Swedish national trend 1996-2015: -3.3 % per year (***)**

**Impact sites**
Kendall's rank correlation tau = -0.095
z = -0.418, p-value = 0.676

**Control sites**
Kendall's rank correlation tau = -0.381
z = -1.672, p-value = 0.095

**All sites**
Kendall's rank correlation tau = -0.356
z = -1.548, p-value = 0.122

**Difference between Impact and Control sites**
Kendall's rank correlation tau = 0.156
z = 0.692 p-value = 0.489

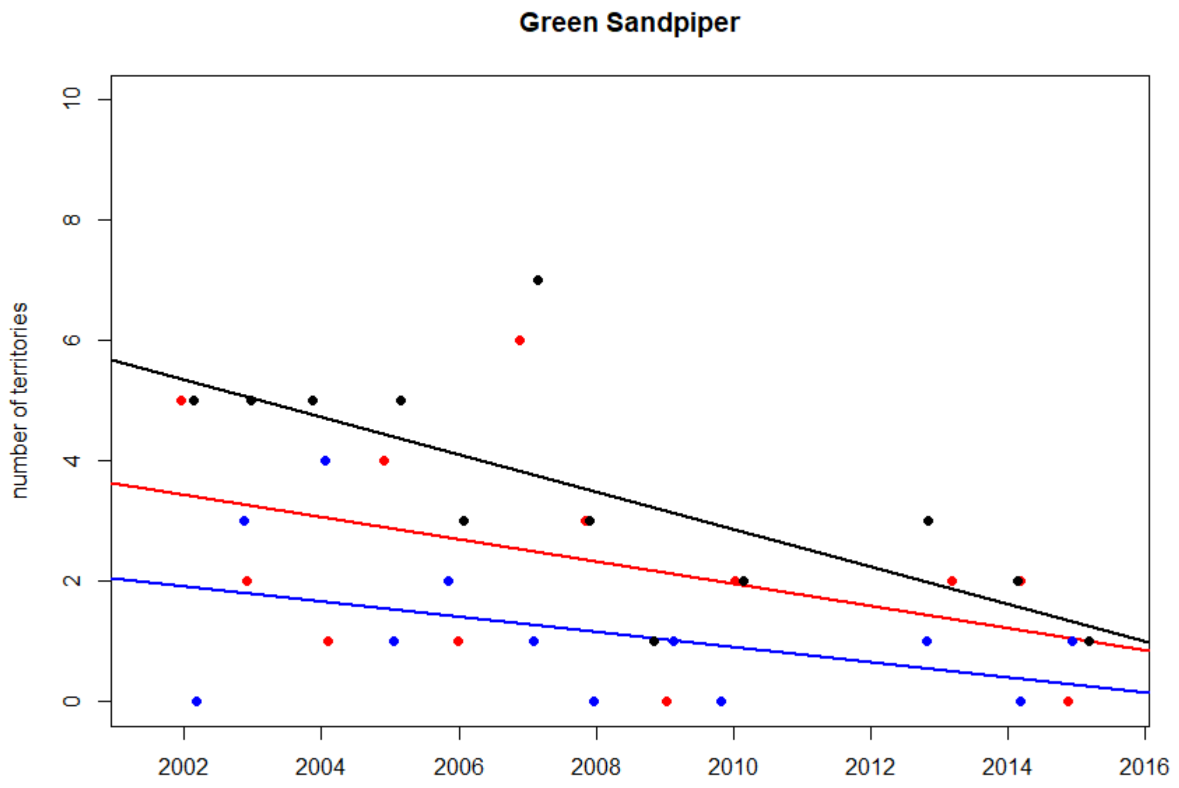


Total yearly numbers of territories of Green Sandpiper in Impact sites (red), in Control sites (blue) and in all sites (black)

**Swedish national trend 1996-2015: 3.8 % per year (***)**

**Impact sites**
Kendall's rank correlation tau = -0.323
z = -1.407, p-value = 0.159

**Control sites**
Kendall's rank correlation tau = -0.313
z = -1.315, p-value = 0.189

**All sites**
Kendall's rank correlation tau = -0.614
z = -2.627, p-value = 0.009**

**Difference between Impact and Control sites**
Kendall's rank correlation tau = -0.114
z = -0.494, p-value = 0.622

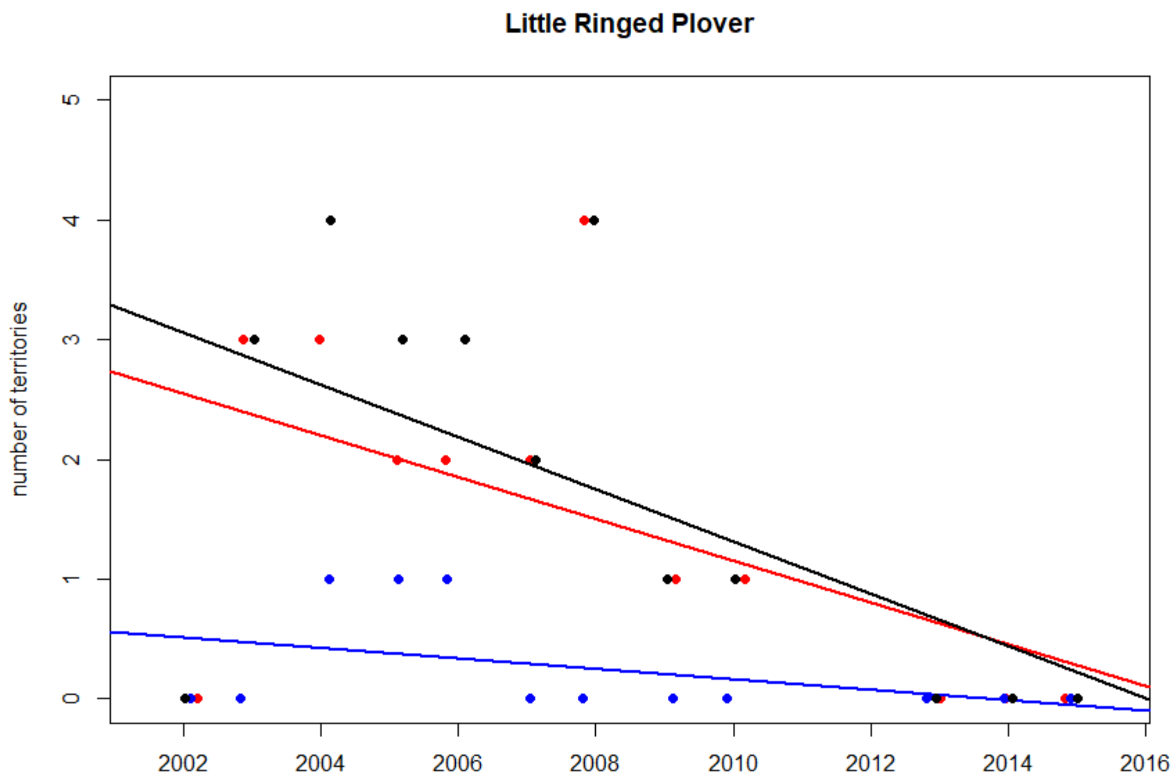


Total yearly numbers of territories of Little Ringed Plover in Impact sites (red), in Control sites (blue) and in all sites (black)

**Swedish national trend 1996-2015: +1.6 % per year (NS)**

**Impact sites**
Kendall's rank correlation tau = -0.481
z = -2.059, p-value = 0.039*

**Control sites**
Kendall's rank correlation tau = -0.355
z = -1.387, p-value = 0.1655

**All sites**
Kendall's rank correlation tau = -0.481
z = -2.059, p-value = 0.039*

**Difference between Impact and Control sites**
Kendall's rank correlation tau = -0.389
z = -1.650, p-value = 0.099

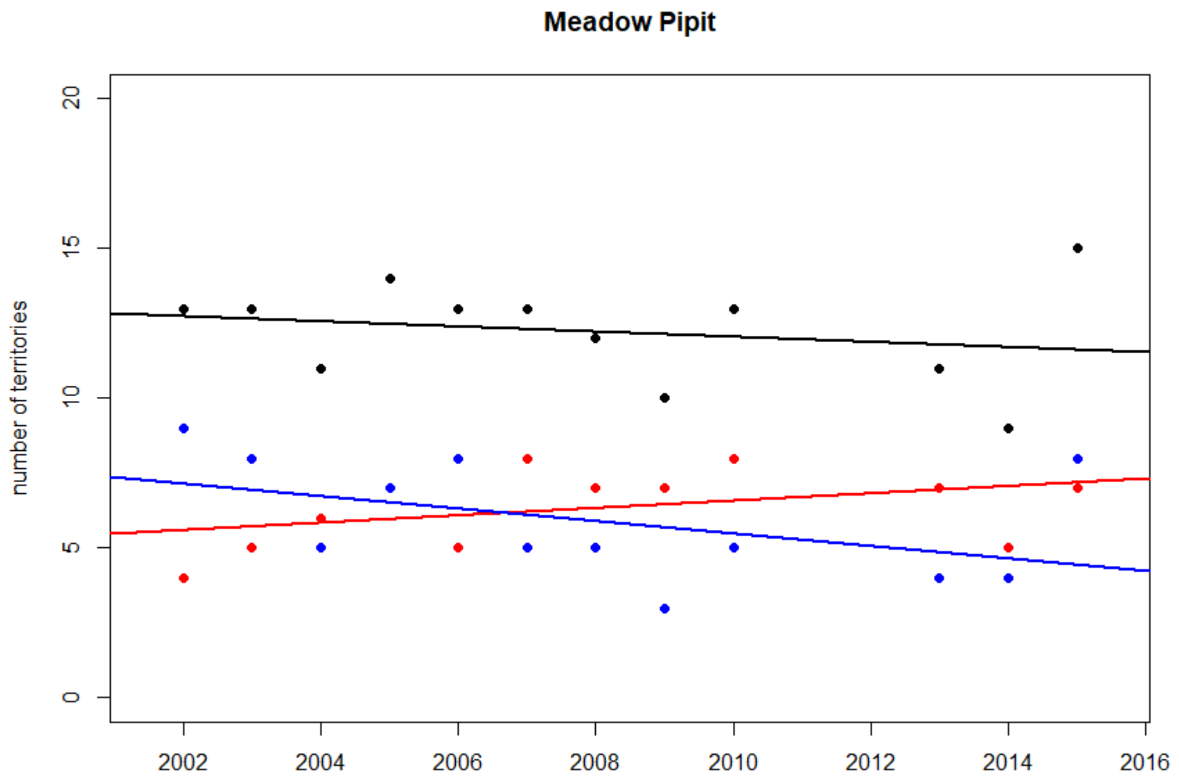


Total yearly numbers of territories of Meadow Pipit in Impact sites (red), in Control sites (blue) and in all sites (black)

**Swedish national trend 1996-2015: -1.2 % per year (***)**

**Impact sites**
Kendall's rank correlation tau = 0.376
z = 1.591, p-value = 0.112

**Control sites**
Kendall's rank correlation tau = -0.461
z = -1.983, p-value = 0.047*

**All sites**
Kendall's rank correlation tau = -0.216
z = -0.931, p-value = 0.352

**Difference between Impact and Control sites**
Kendall's rank correlation tau = 0.362
z = 1.599, p-value = 0.110

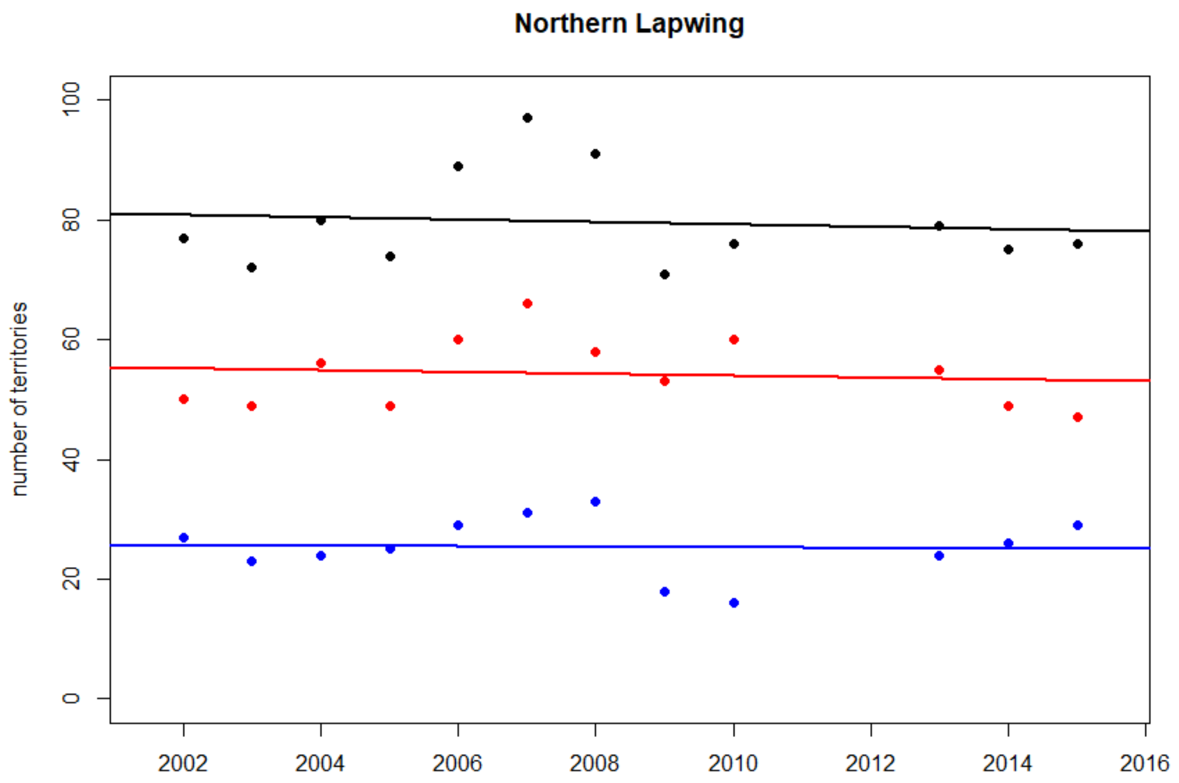


Total yearly numbers of territories of Northern Lapwing in Impact sites (red), in Control sites (blue) and in all sites (black)

**Swedish national trend 1996-2015: -0.7 % per year (NS)**

**Impact sites**
Kendall's rank correlation tau = -0.094
z = -0.416, p-value = 0.677
**Control sites**
Kendall's rank correlation tau = 0.092
z = 0.413, p-value = 0.679
**All sites**
Kendall's rank correlation tau = -0.015
z = -0.069, p-value = 0.945
**Difference between Impact and Control sites**
Kendall's rank correlation tau = 0.016
z = 0.069, p-value = 0.945

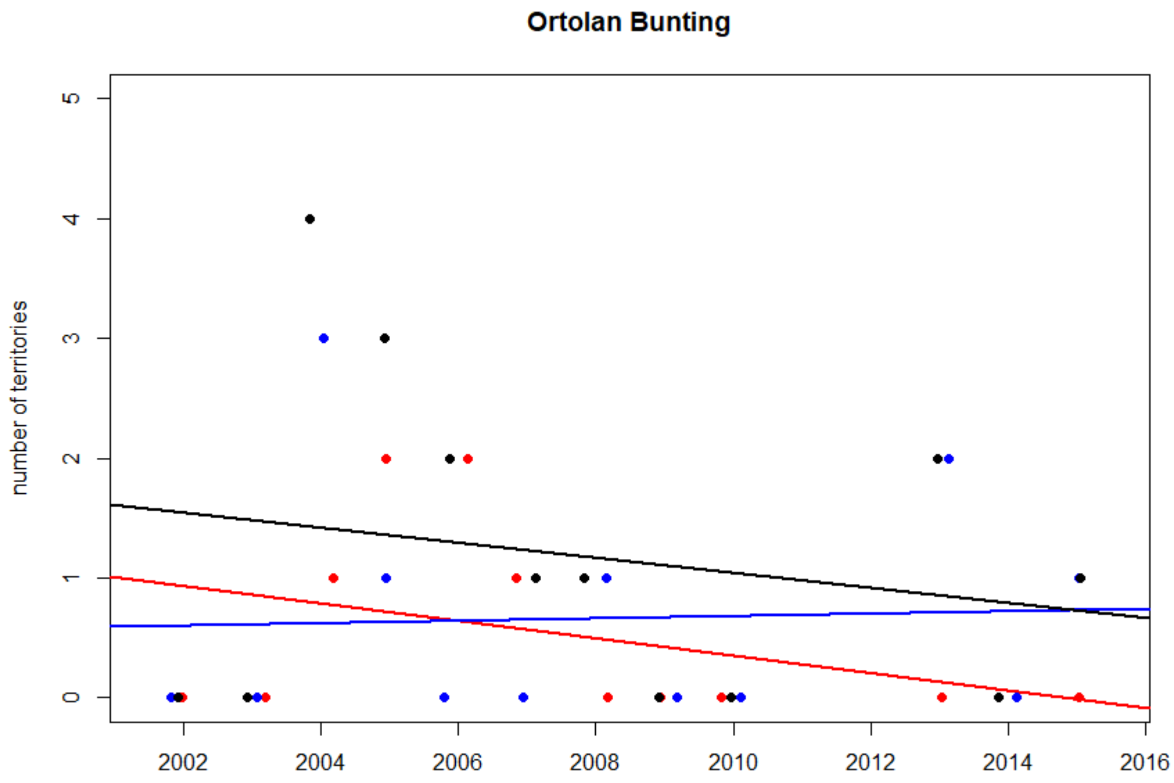


Total yearly numbers of territories of Ortolan Bunting in Impact sites (red), in Control sites (blue) and in all sites (black)

**Swedish national trend 1996-2015: -7.1 % per year (***)**

**Impact sites**
Kendall's rank correlation tau = -0.328
z = -1.327, p-value = 0.184

**Control sites**
Kendall's rank correlation tau = 0.076
z = 0.312, p-value = 0.755

**All sites**
Kendall's rank correlation tau = -0.171
z = -0.723, p-value = 0.470

**Difference between Impact and Control sites**
Kendall's rank correlation tau = -0.220
z = -0.936, p-value = 0.349

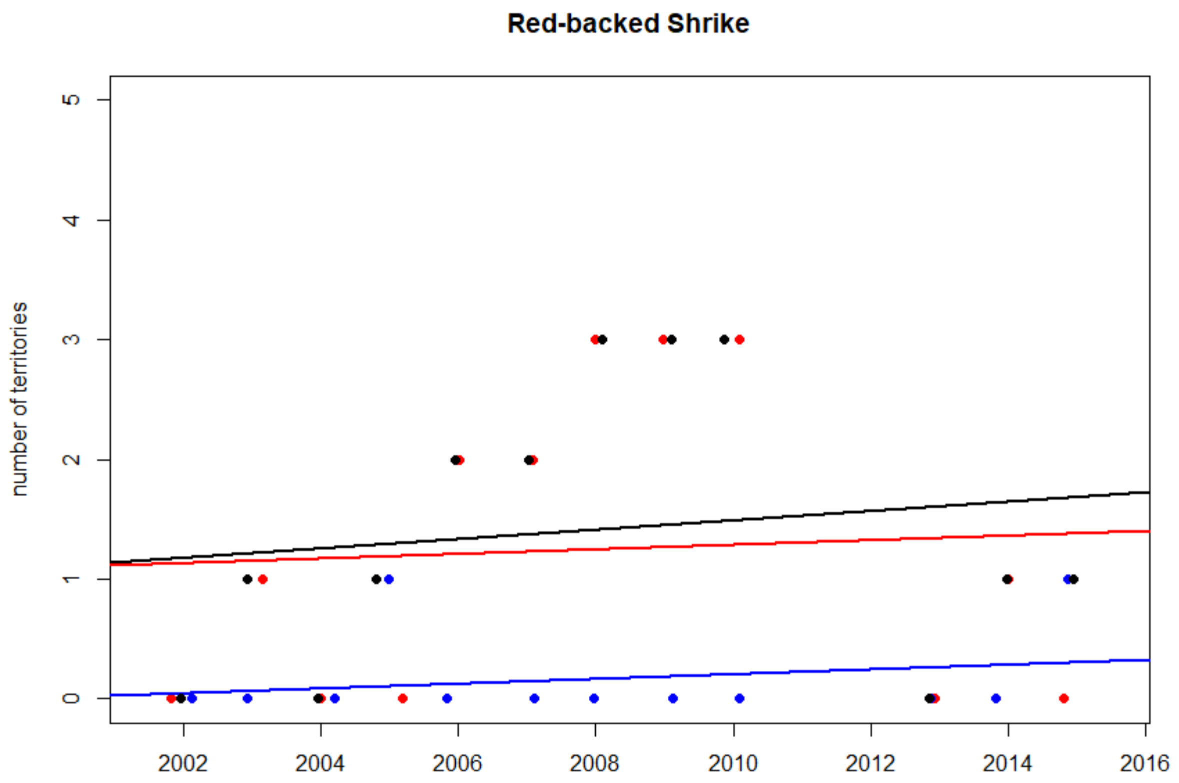


Total yearly numbers of territories of Red-backed Shrike in Impact sites (red), in Control sites (blue) and in all sites (black)

**Swedish national trend 1996-2015: -4.3 % per year (***)**

**Impact sites**
Kendall's rank correlation tau = 0.190
z = 0.797, p-value = 0.425

**Control sites**
Kendall's rank correlation tau = 0.165
z = 0.645, p-value = 0.519

**All sites**
Kendall's rank correlation tau = 0.287
z = 1.215, p-value = 0.224

**Difference between Impact and Control sites**
Kendall's rank correlation tau = 0.114
z = 0.492, p-value = 0.623

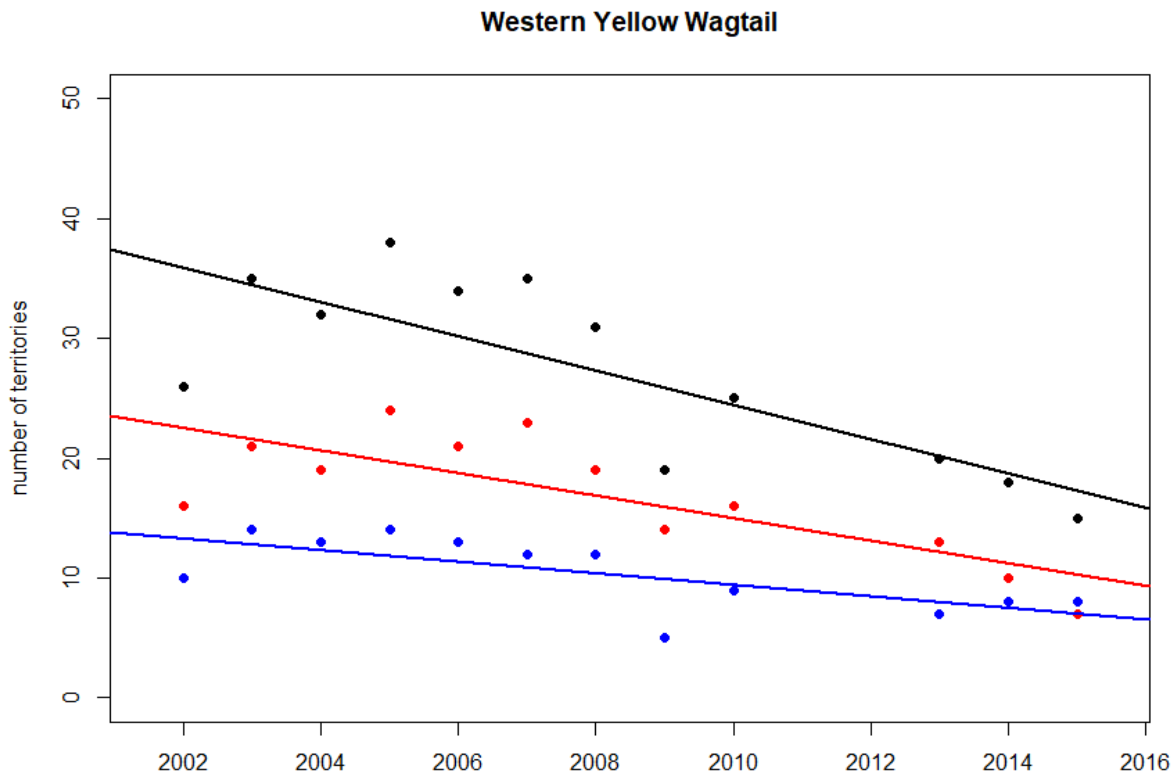


Total yearly numbers of territories of Western Yellow Wagtail in Impact sites (red), in Control sites (blue) and in all sites (black)

**Swedish national trend 1996-2015: -0.5 % per year (NS) for subspecies *thunbergi* and +5.0 % per year (***) for subspecies *flava*.**

**Impact sites**
Kendall's rank correlation tau = -0.574
z = -2.555, p-value = 0.011*

**Control sites**
Kendall's rank correlation tau = -0.563
z = -2.492, p-value = 0.013*

**All sites**
Kendall's rank correlation tau = -0.595
z = -2.681, p-value = 0.007**

**Difference between Impact and Control sites**
Kendall's rank correlation tau = -0.286
z = -1.256, p-value = 0.209

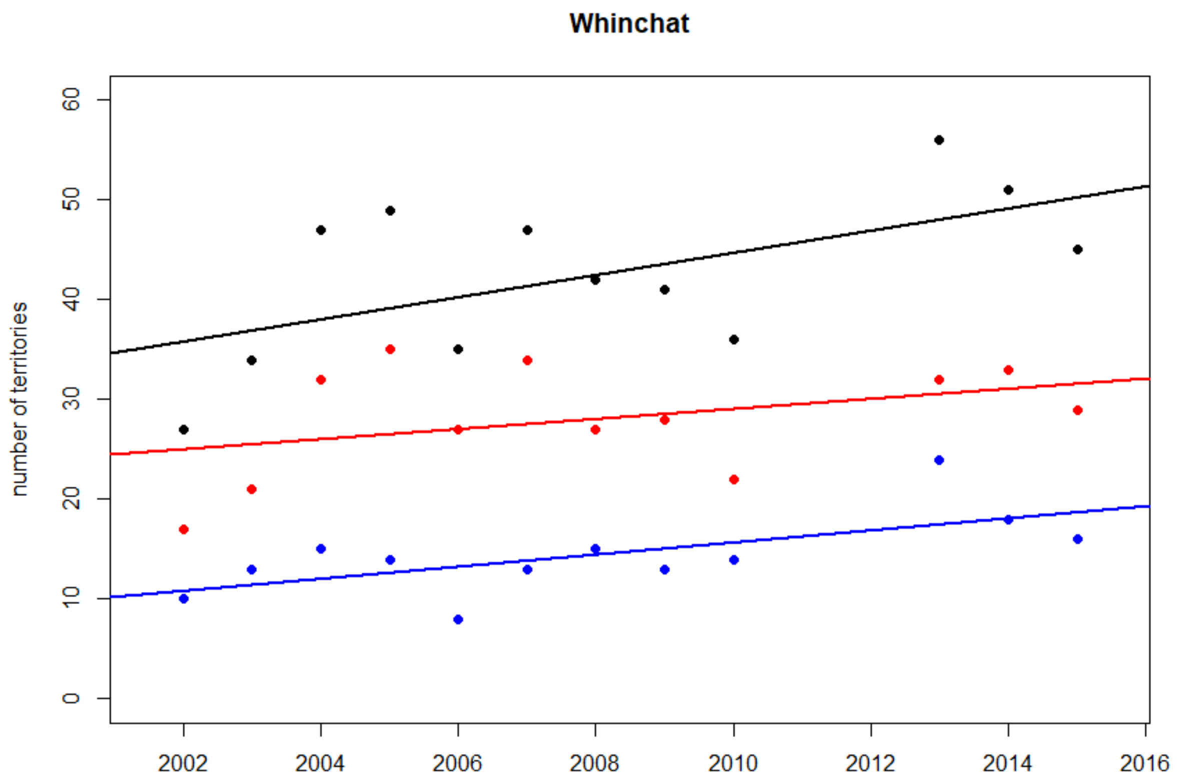


Total yearly numbers of territories of Whinchat in Impact sites (red), in Control sites (blue) and in all sites (black)

**Swedish national trend 1996-2015: -1.5 % per year (***)**

**Impact sites**
Kendall's rank correlation tau = 0.246
z = 1.102, p-value = 0.270

**Control sites**
Kendall's rank correlation tau = 0.489
z = 2.155, p-value = 0.032*

**All sites**
Kendall's rank correlation tau = 0.351
z = 1.581, p-value = 0.114

**Difference between Impact and Control sites**
Kendall's rank correlation tau = -0.016
z = -0.070, p-value = 0.945
