## Supplementary material for "Are railways detrimental to bird populations? A BDACI study on the construction of the Bothnia Line Railway": Trends of territory numbers per site och species

### Supplementary file **Terr2**

### Temporal trends per study site for numbers of territories of individual species and all species combined

Plots and Kendall’s rank correlation tests for all species combined, but only plots for individual species (zero-only sites omitted).

Sites in alphabetical order

Significant level indicators:
* = P < 0.05, ** = P < 0.01, *** = P < 0.001.

Values for individual years are colour-coded in accordance of the phases of the railway construction process: *Before* (green), *Construction* (blue), *Ready* (purple) and *Traffic* (red). Control sites are coded in black for all years. A simple linear regression line added for visualization.

Contents page


| Contents | page |
|---|---|
| All species combined | 2-18 |
| Barn Swallow | 19-27 |
| Common Rosefinch | 28-32 |
| Common Snipe | 33-39 |
| Common Starling | 40-46 |
| Eurasian Curlew | 47-56 |
| Eurasian Golden Plover | 57 |
| Eurasian Skylark | 58-66 |
| Green sandpiper | 67-74 |
| Little Ringed Plover | 75-77 |
| Meadow Pipit | 78-82 |
| Northern Lapwing | 83-92 |
| Ortolan Bunting | 93-95 |
| Red-backed Shrike | 96-98 |
| Western Yellow Wagtail | 99-104 |
| Whinchat | 105-114 |

### All species combined

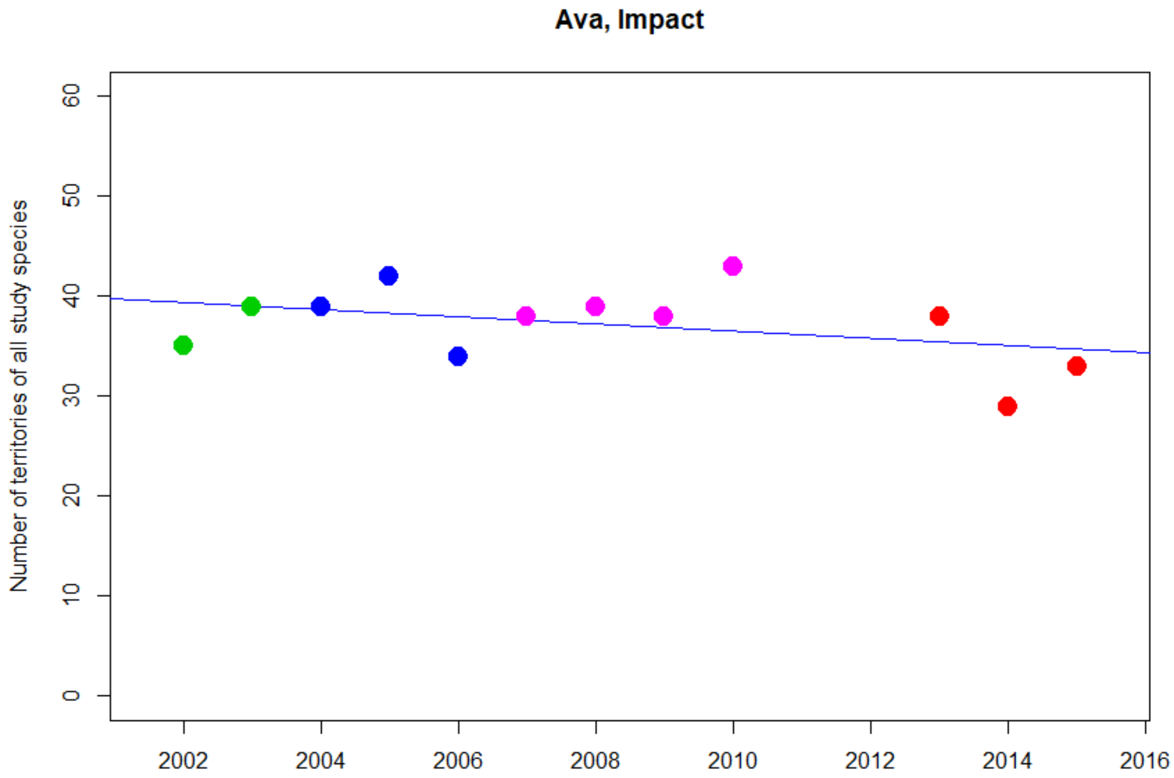


Kendall's rank correlation tau = -0.222
z = -0.977, p-value = 0.329

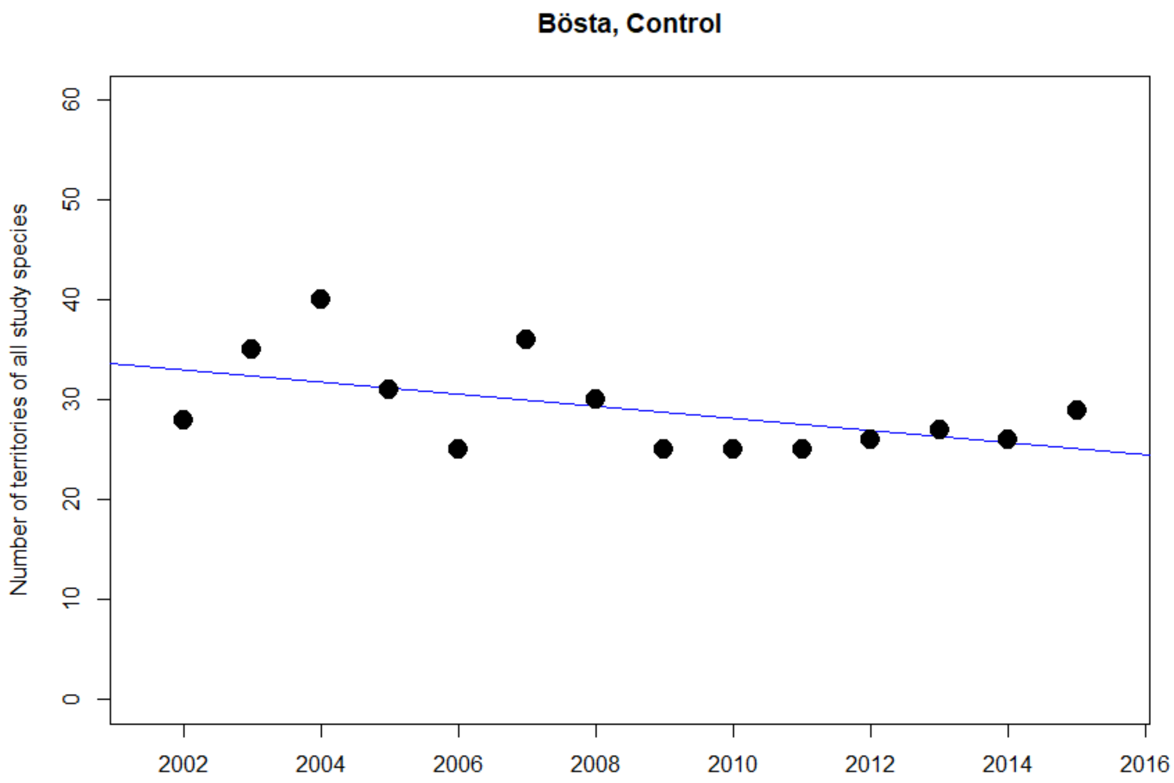


Kendall's rank correlation tau = -0.252
z = -1.222, p-value = 0.222

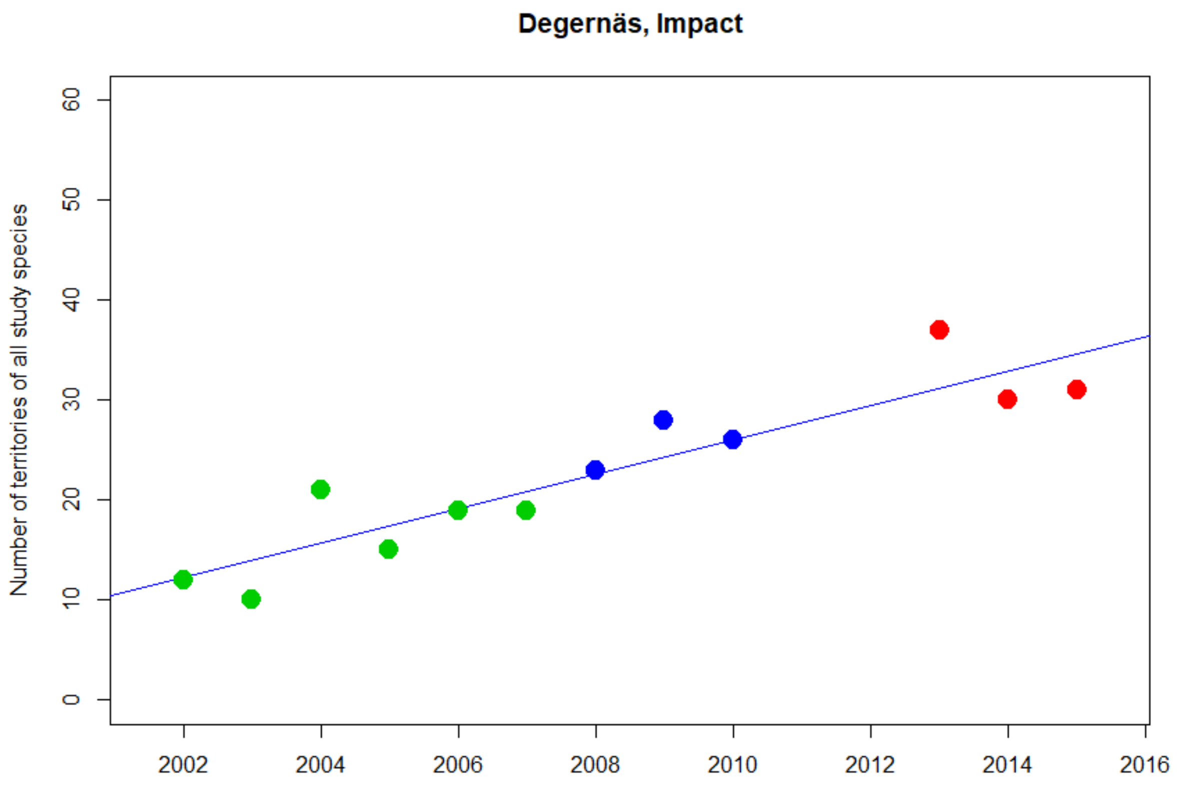


Kendall's rank correlation tau = 0.779
z = 3.506, p-value < 0.001***

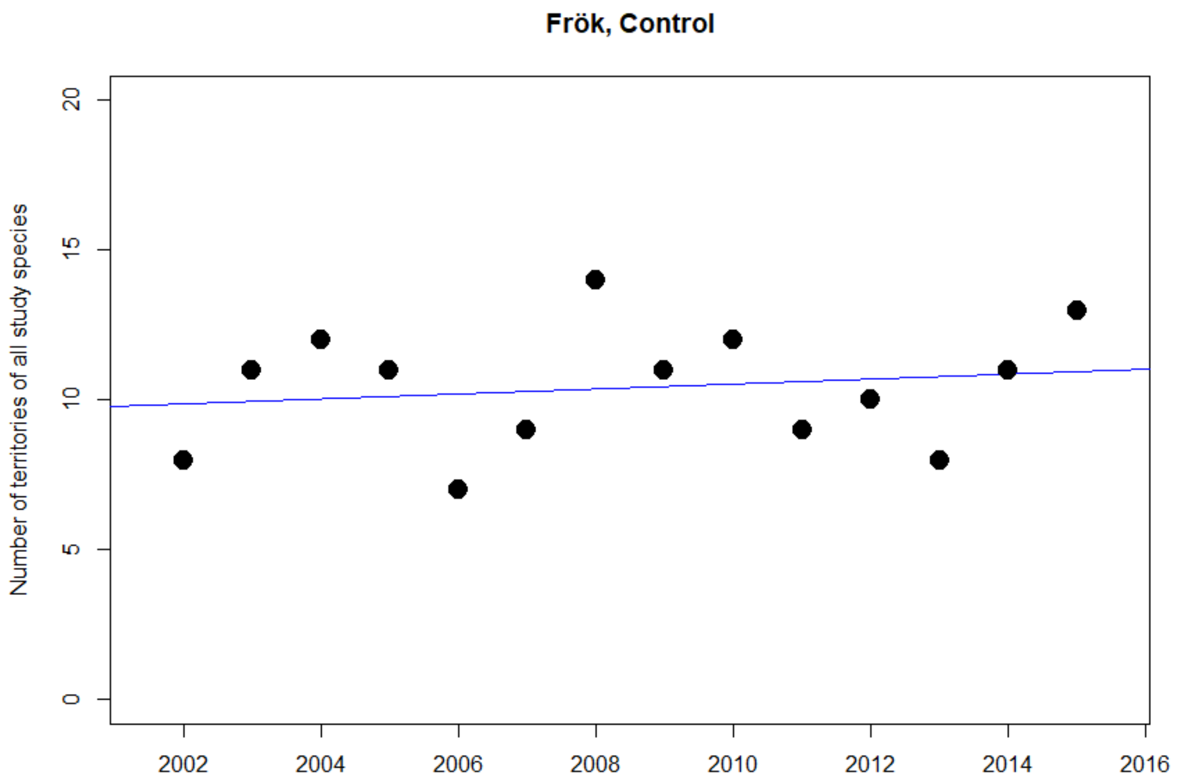


Kendall's rank correlation tau = 0.116
z = 0.557, p-value = 0.577

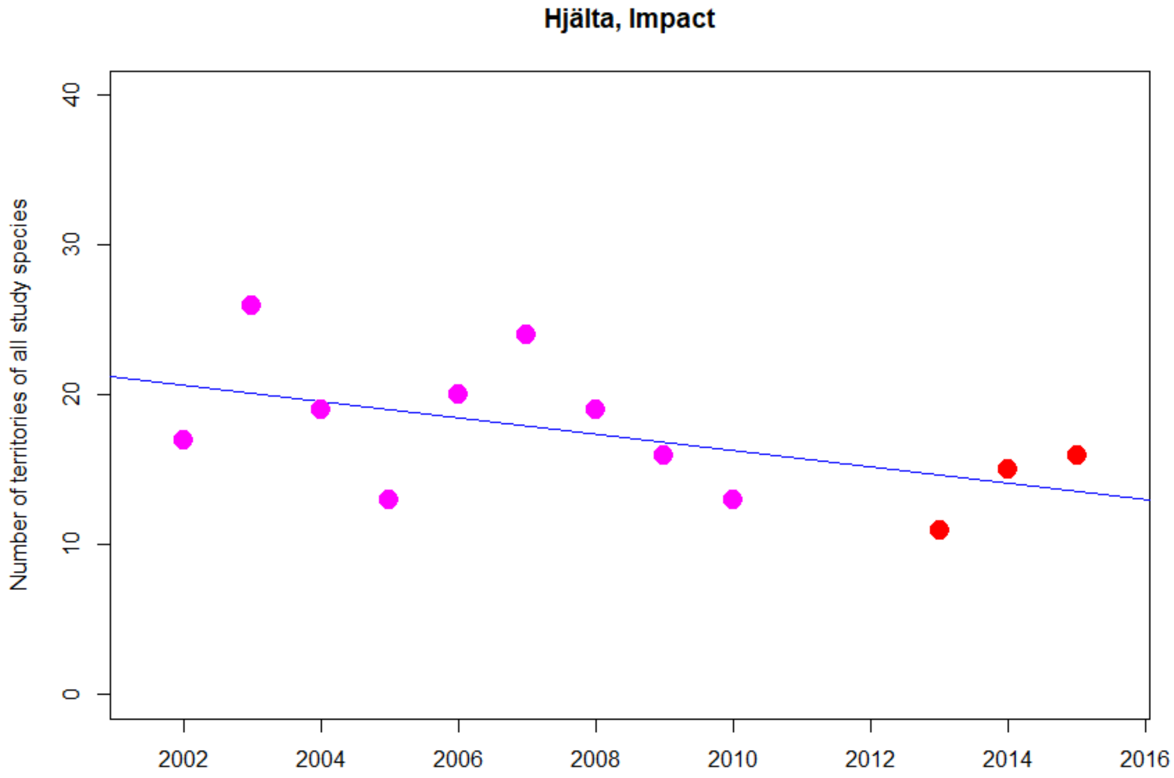


Kendall's rank correlation tau = -0.388
z = -1.727, p-value = 0.084

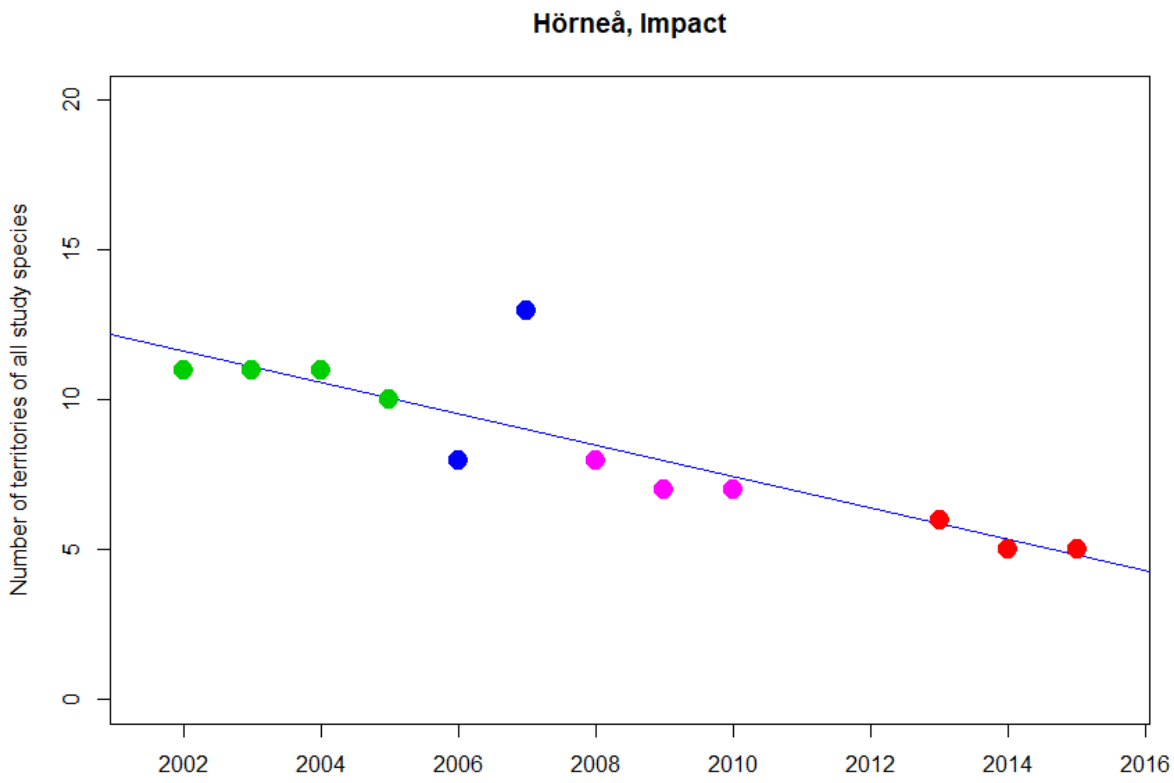


Kendall's rank correlation tau = -0.795
z = -3.484, p-value < 0.001***

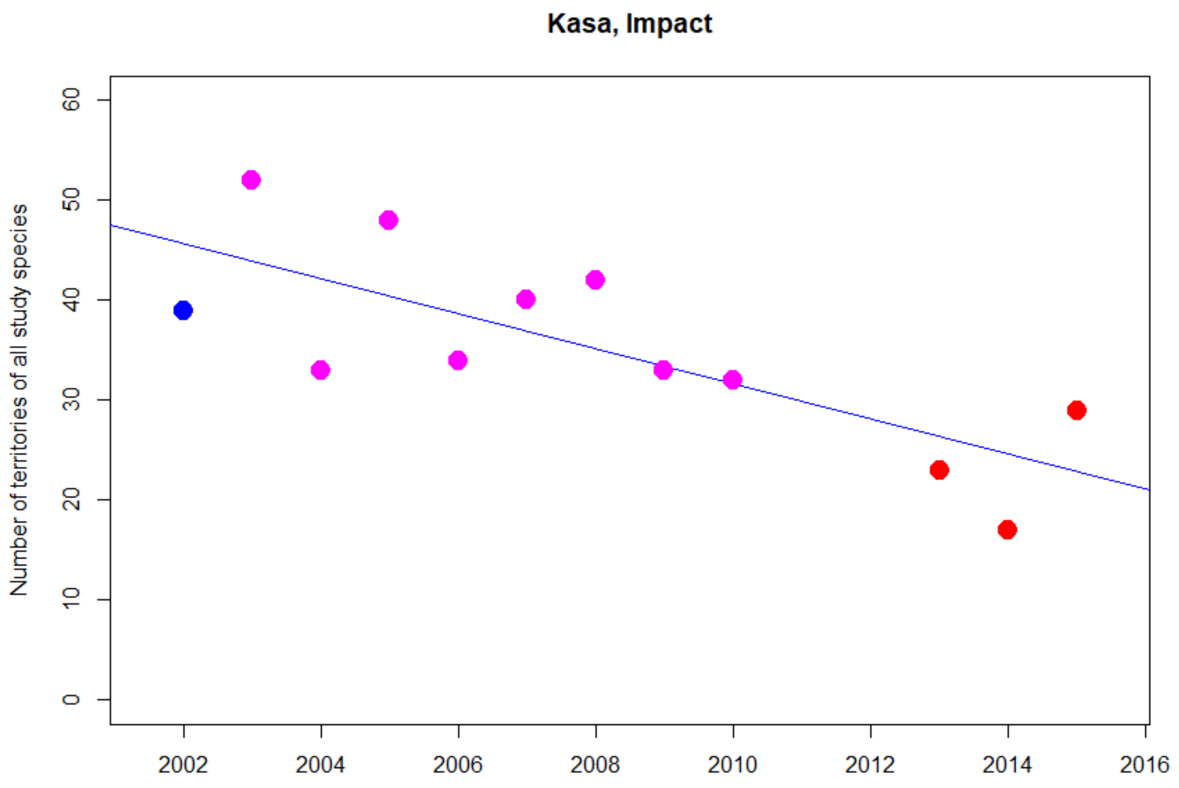


Kendall's rank correlation tau = -0.595
z = -2.681, p-value = 0.007**

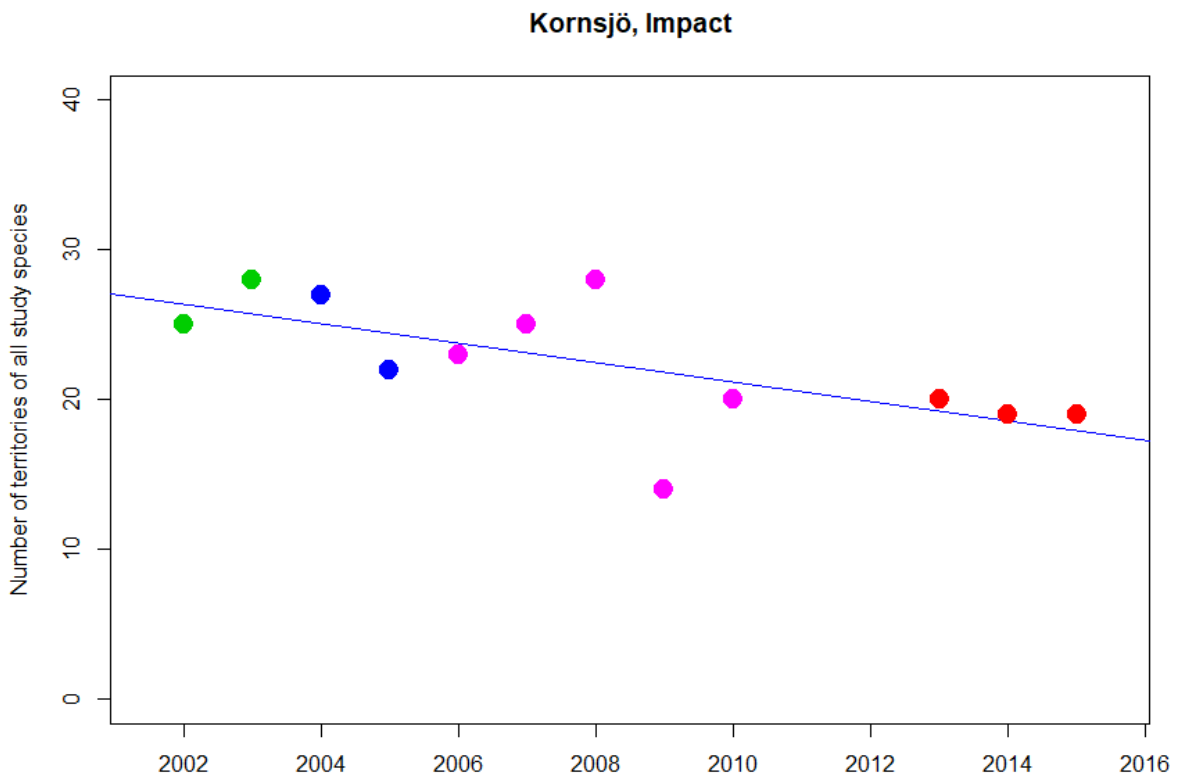


Kendall's rank correlation tau = -0.532
z = -2.354, p-value = 0.019*

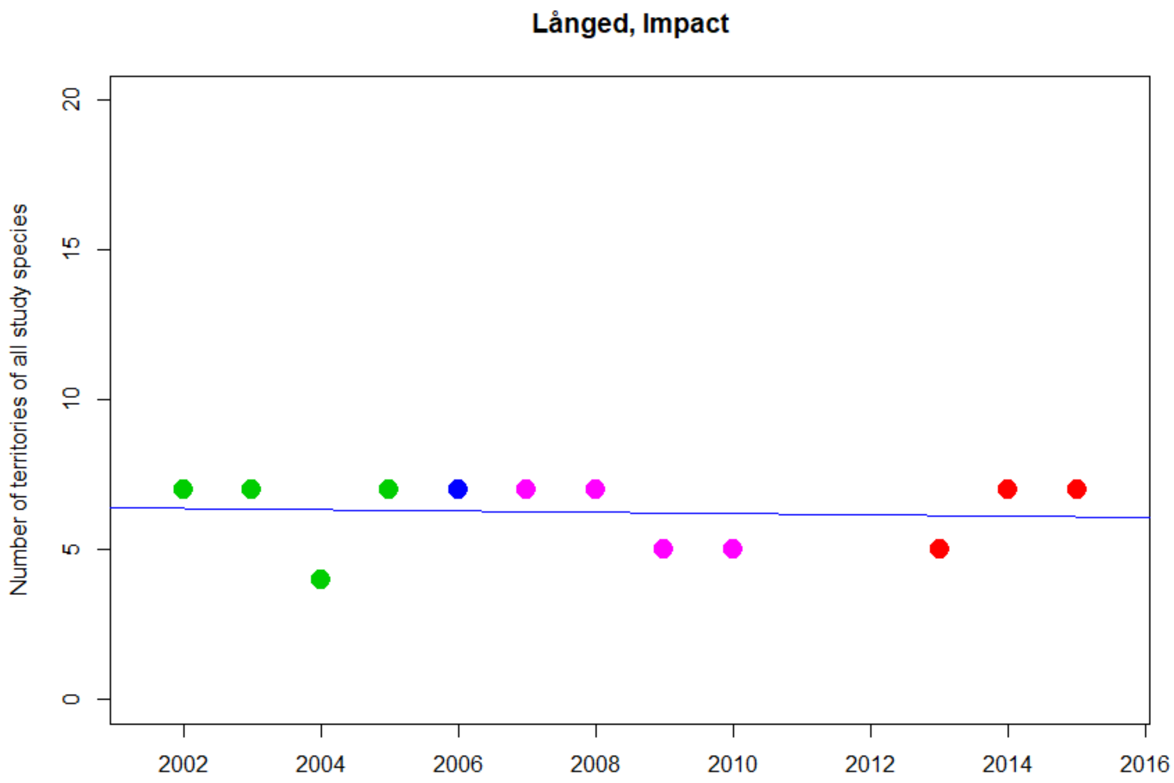


Kendall's rank correlation tau = -0.104
z = -0.417, p-value = 0.677

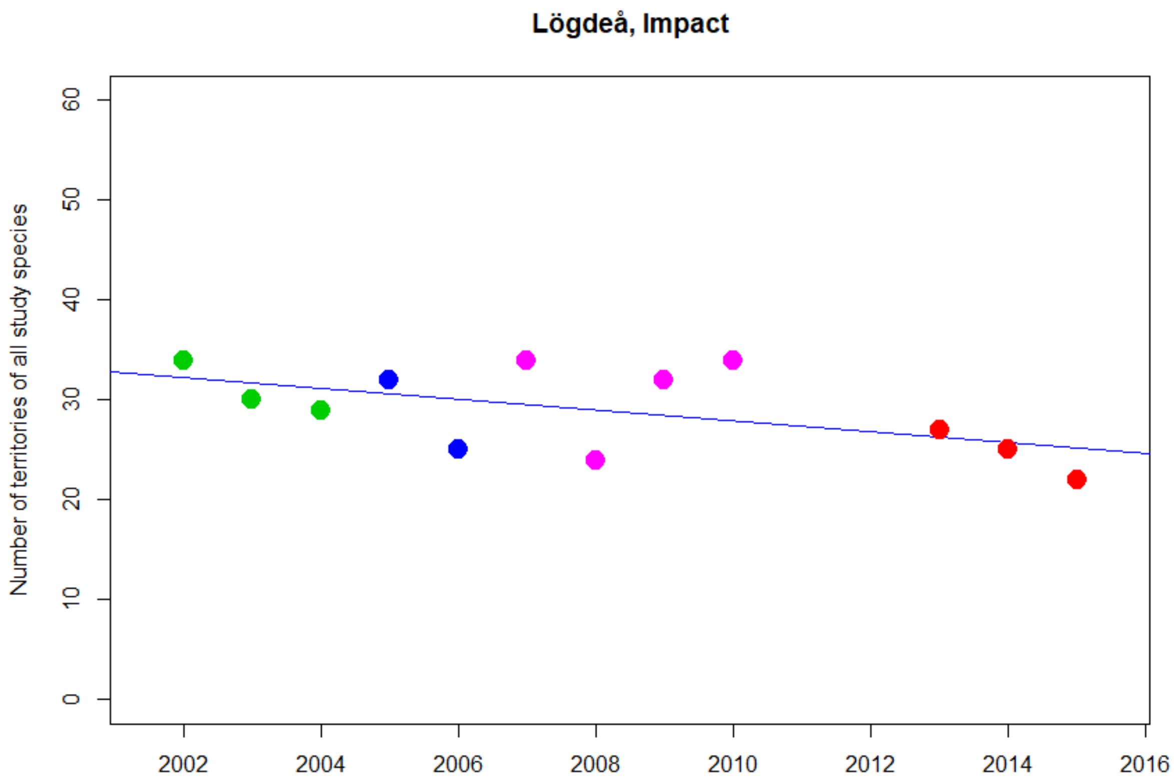


Kendall's rank correlation tau = -0.362
z = -1.599, p-value = 0.110

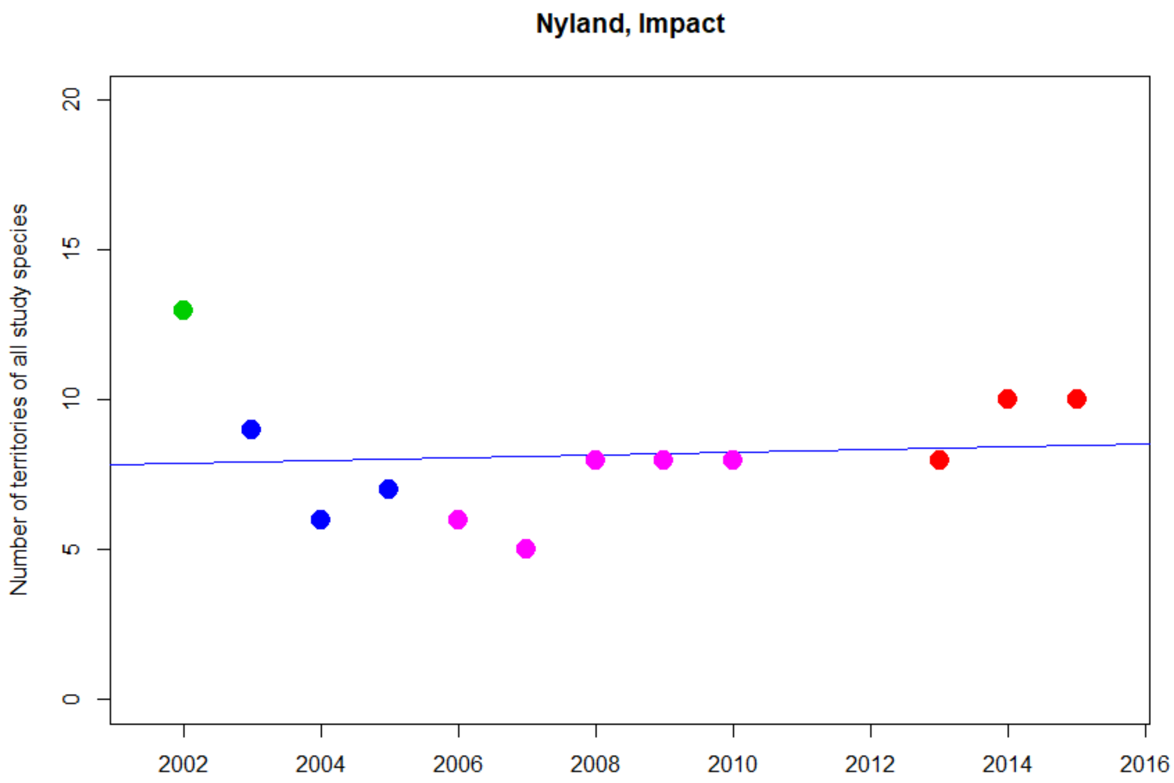


Kendall's rank correlation tau = 0.194
z = 0.844, p-value = 0.399

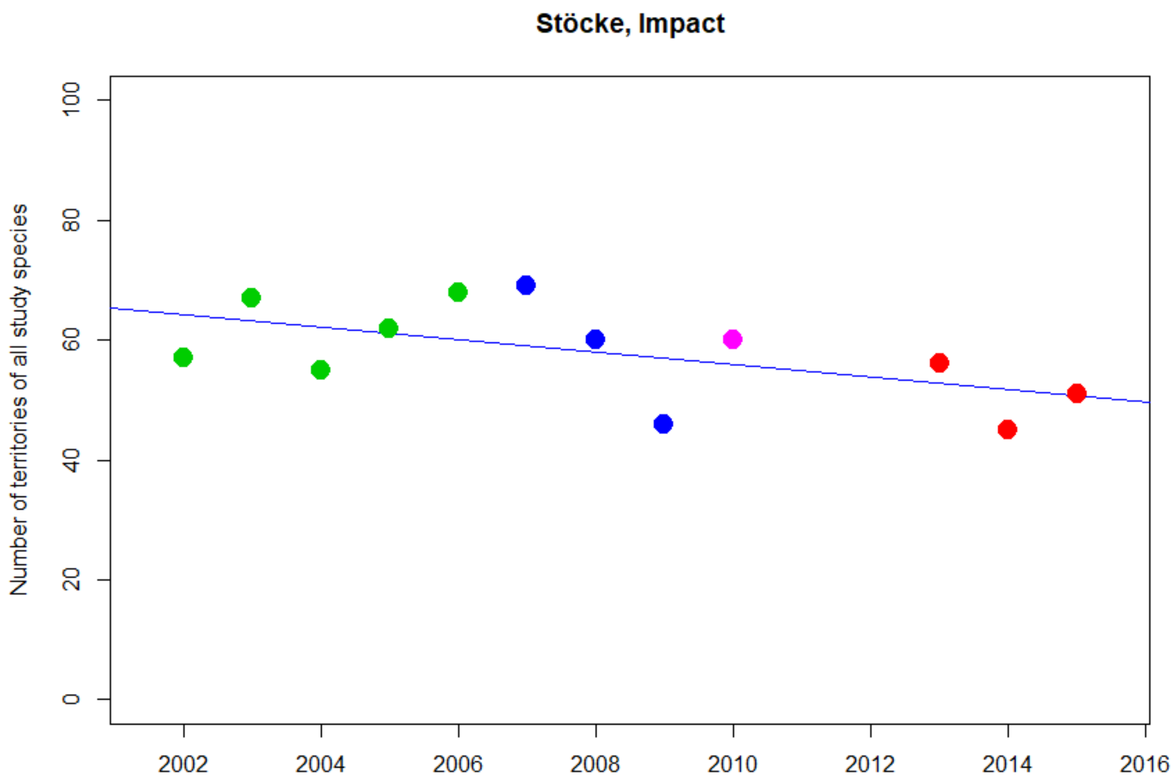


Kendall's rank correlation tau = -0.351
z = -1.581, p-value = 0.114

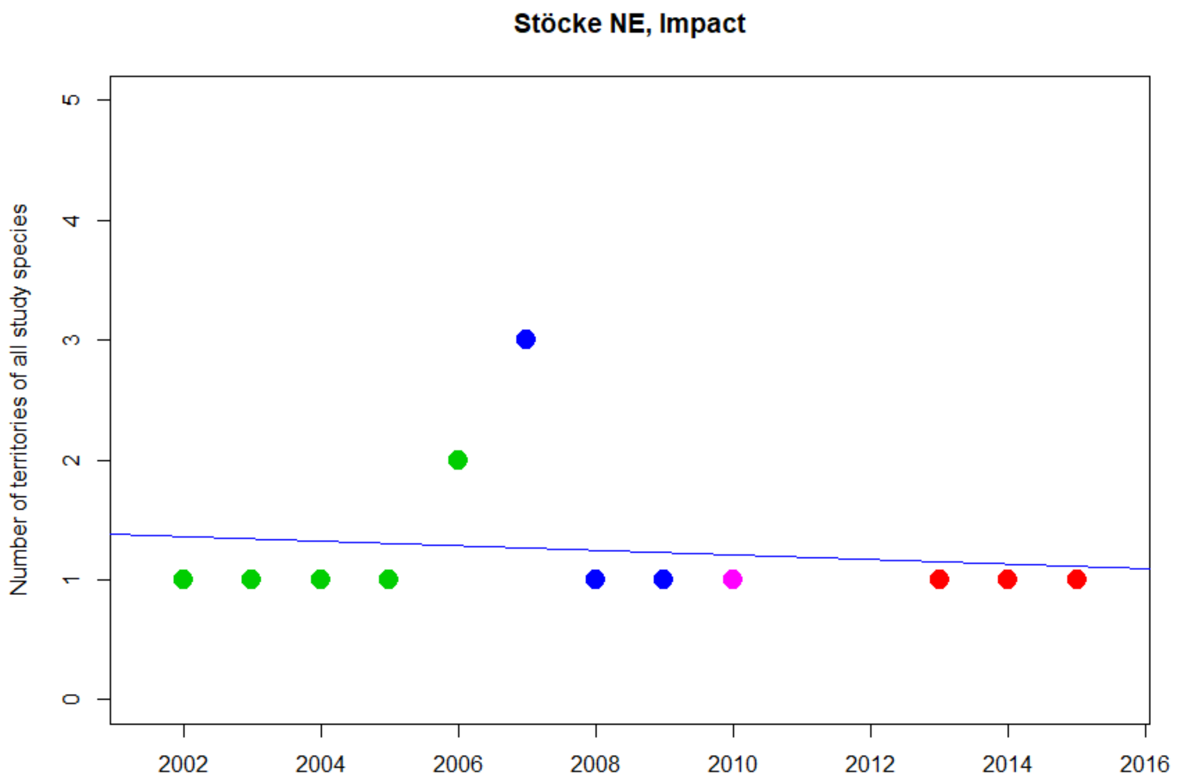


Kendall's rank correlation tau = -0.081
z = -0.320, p-value = 0.749

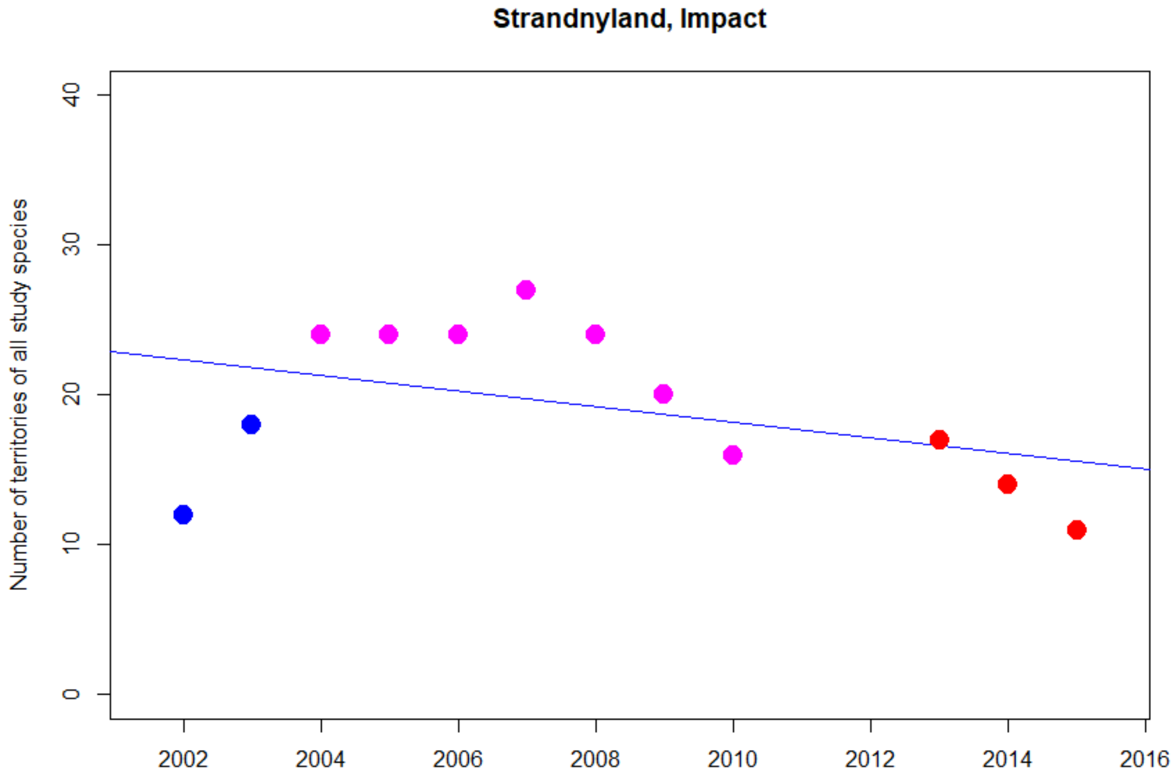


Kendall's rank correlation tau = -0.318
z = -1.400, p-value = 0.161

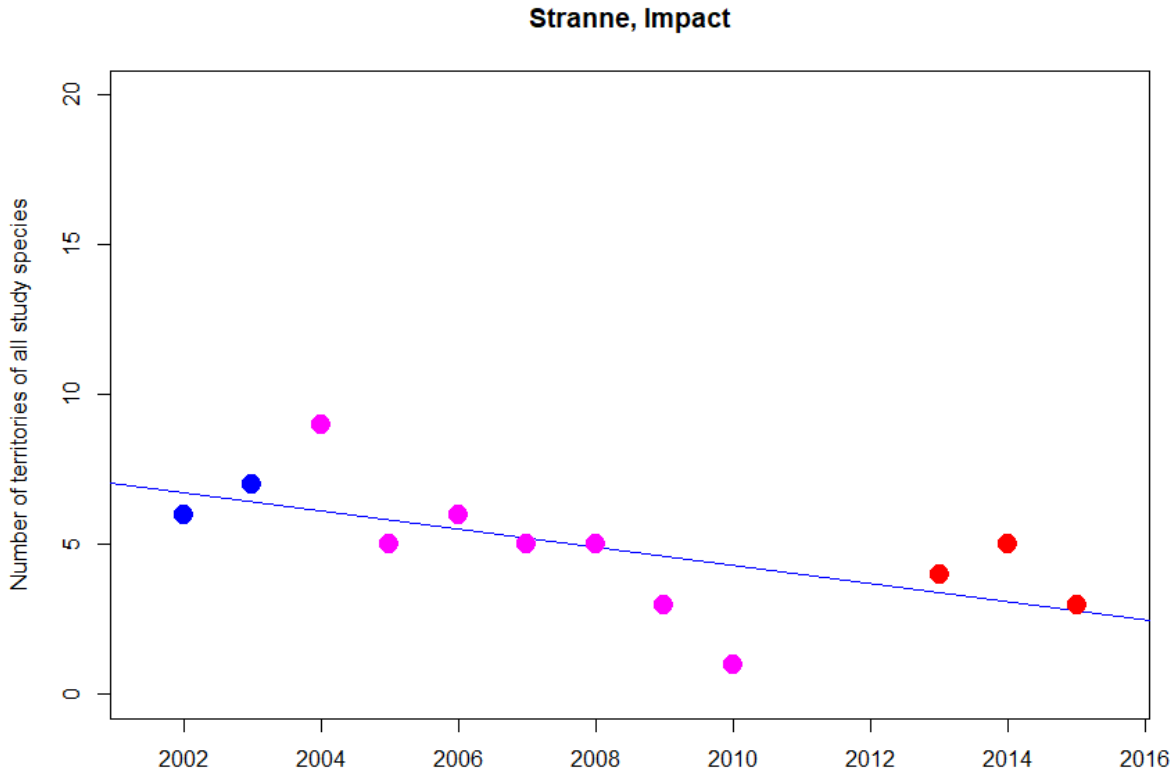


Kendall's rank correlation tau = -0.6141828
z = -2.6737, p-value = 0.008**

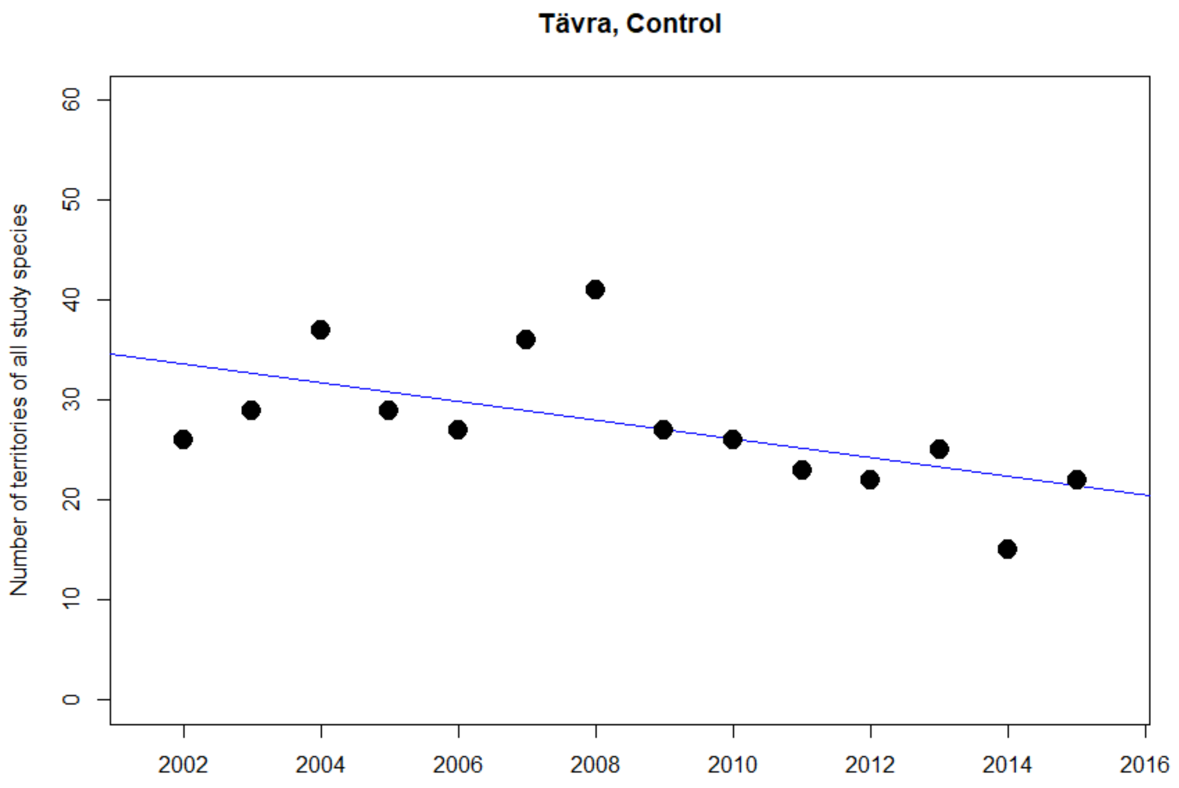


Kendall's rank correlation tau = -0.551
z = -2.699, p-value = 0.007**

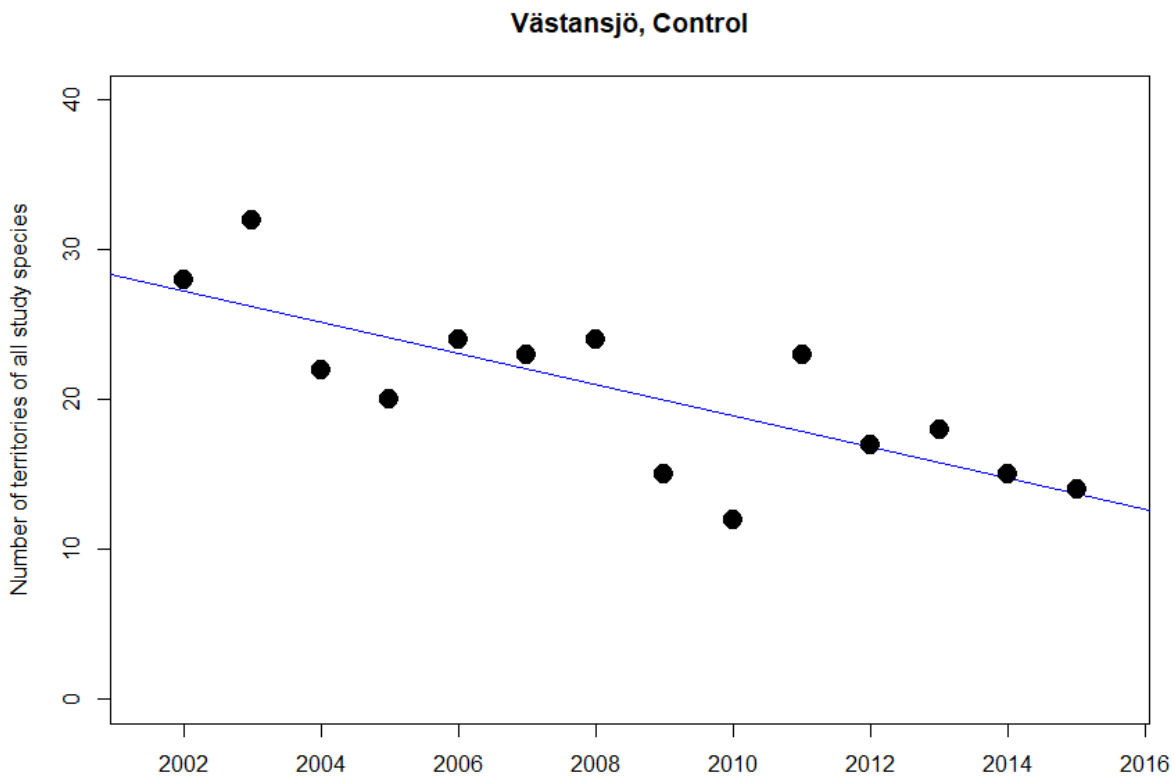


Kendall's rank correlation tau = -0.559
z = -2.750, p-value = 0.006**

### Barn Swallow

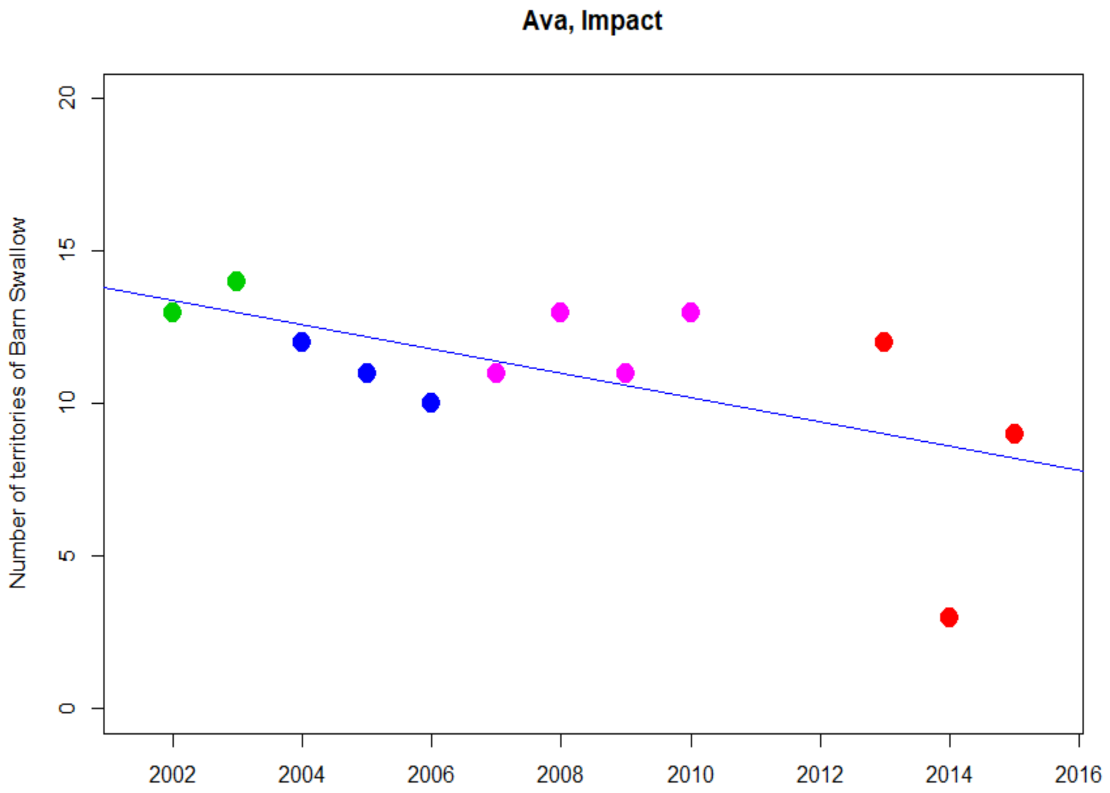


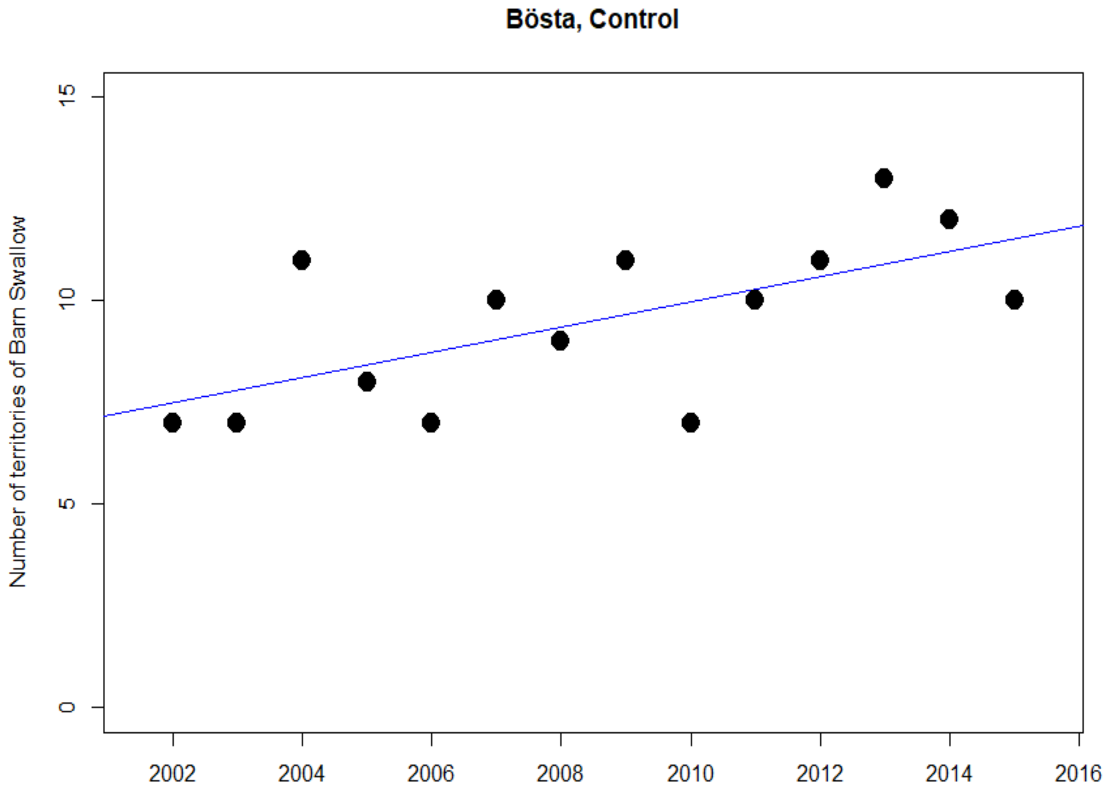

**Degernäs, Impact**

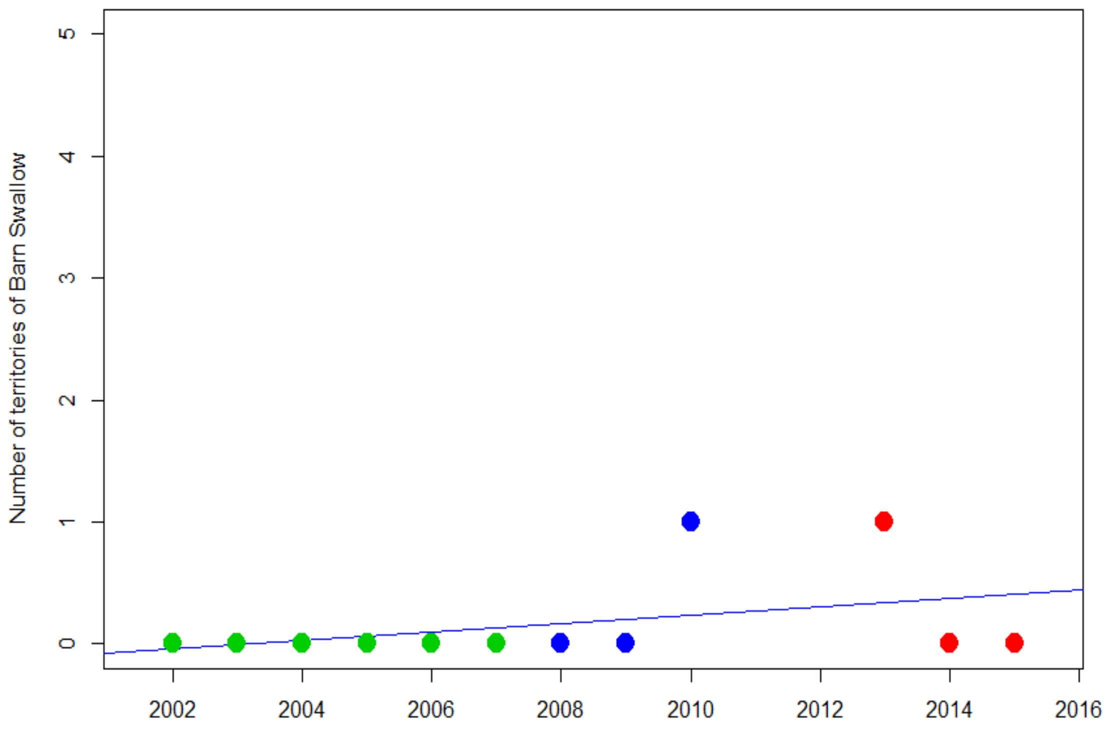


**Frök, Control**

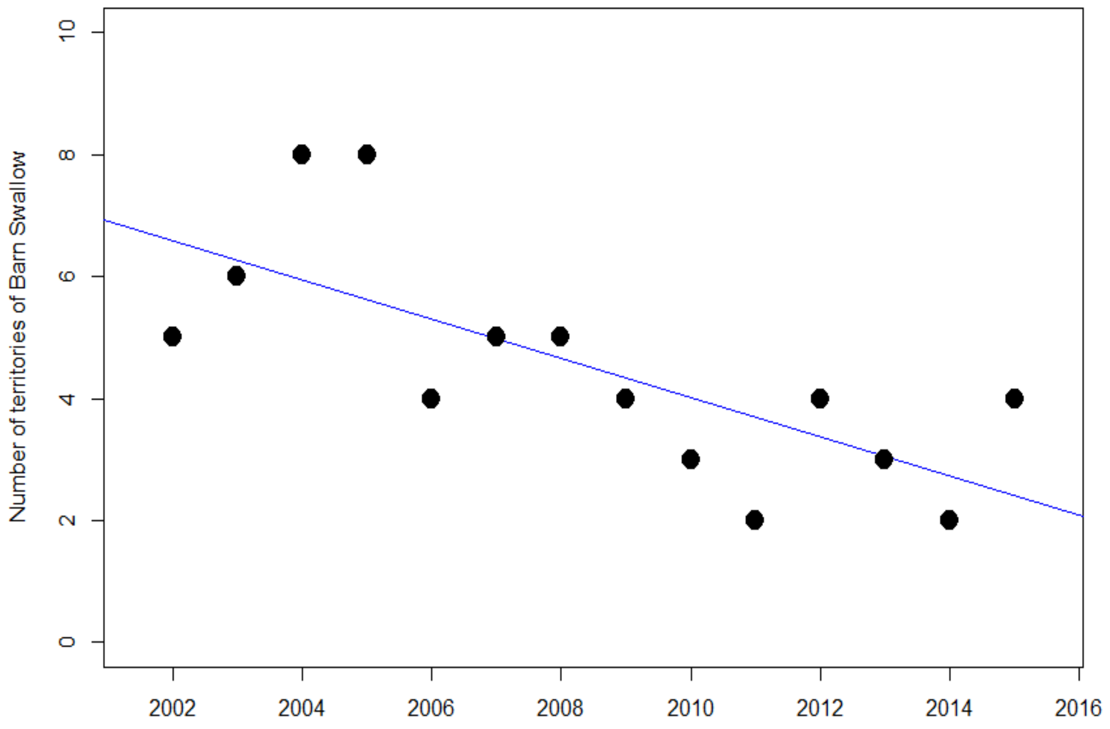

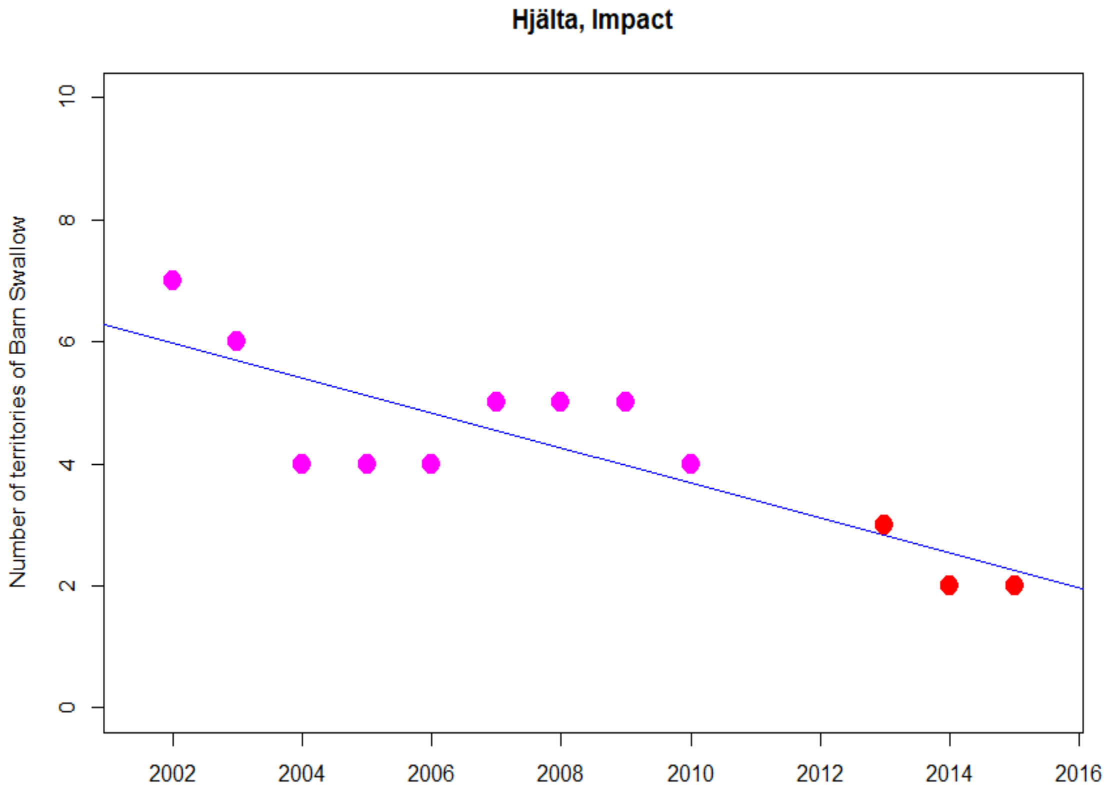
Hjälta, Impact
Number of territories of Barn Swallow
0
2
4
6
8
10
2002
2004
2006
2008
2010
2012
2014
2016

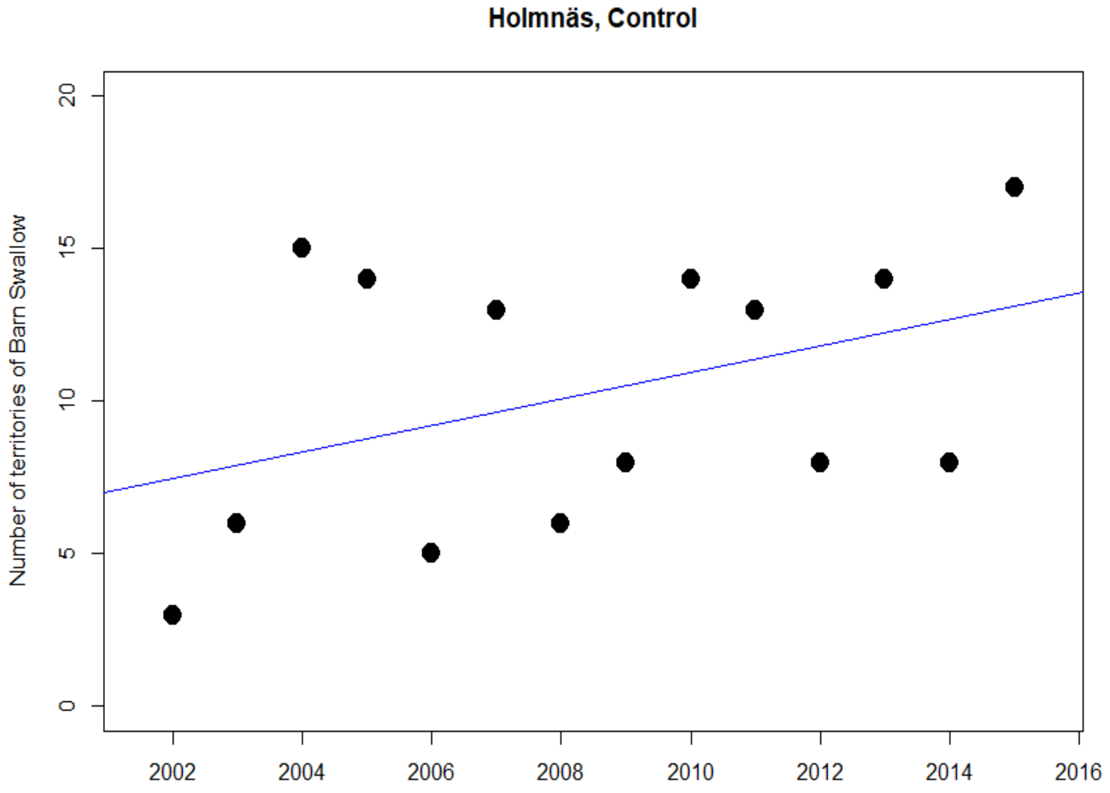
Holmnäs, Control
Number of territories of Barn Swallow
0
5
10
15
20
2002
2004
2006
2008
2010
2012
2014
2016

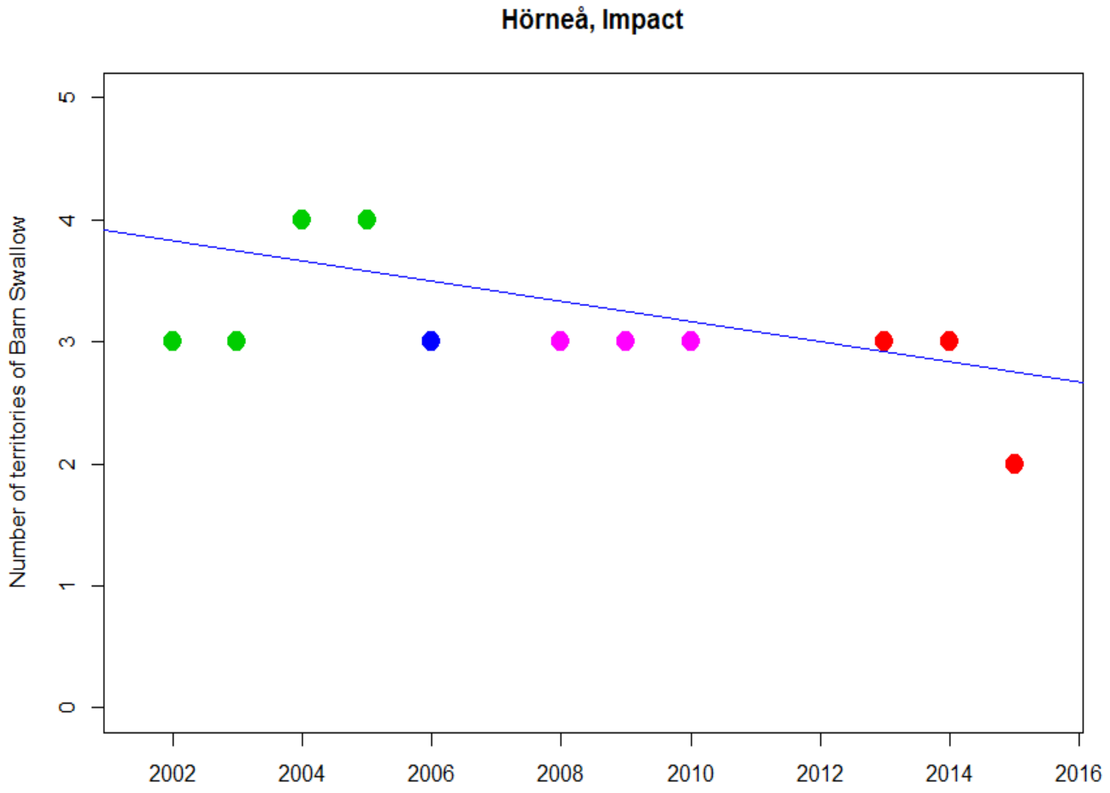
Hörneå, Impact
Number of territories of Barn Swallow
0
1
2
3
4
5
2002
2004
2006
2008
2010
2012
2014
2016

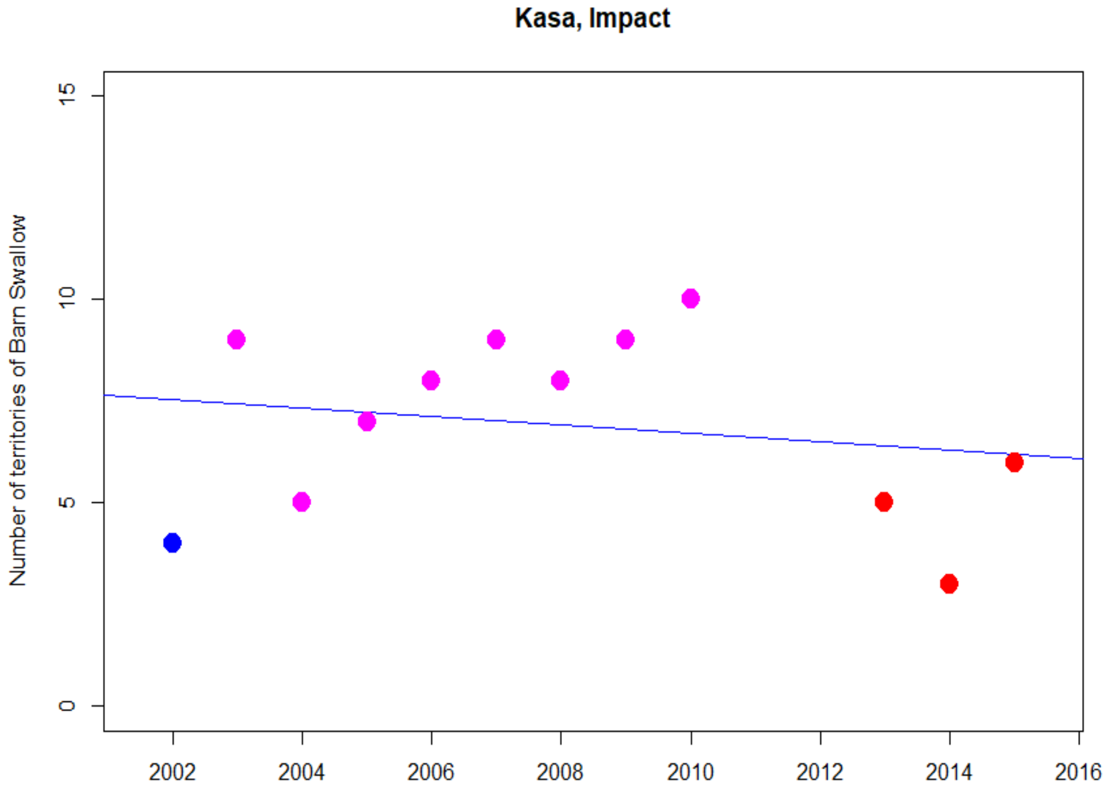
Kasa, Impact
Number of territories of Barn Swallow
0
5
10
15
2002
2004
2006
2008
2010
2012
2014
2016

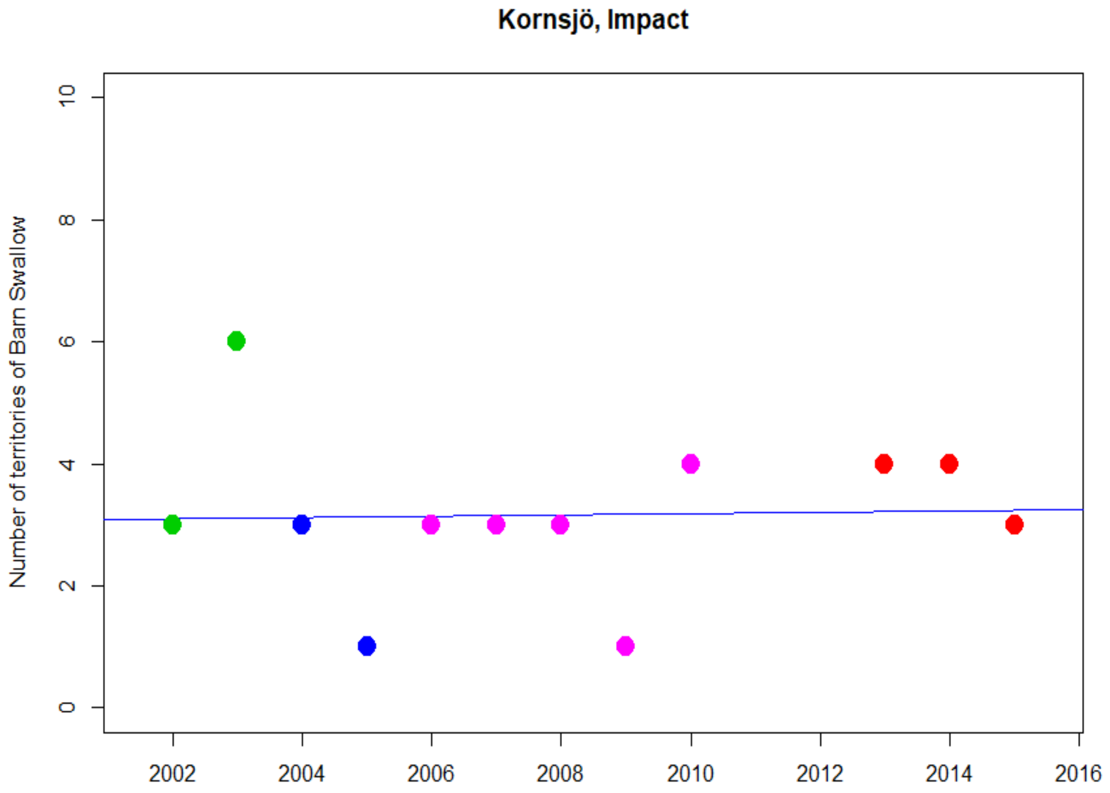
Kornsjö, Impact
Number of territories of Barn Swallow
0
2
4
6
8
10
2002
2004
2006
2008
2010
2012
2014
2016

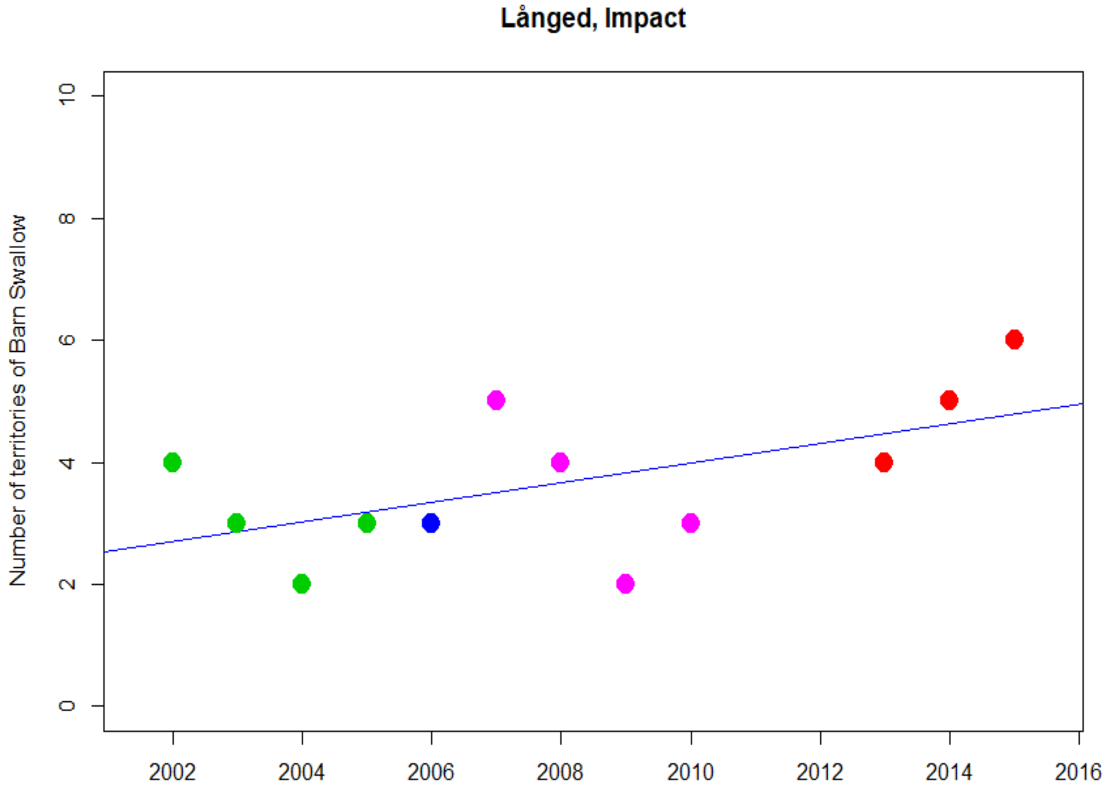
Långed, Impact
Number of territories of Barn Swallow
0
2
4
6
8
10
2002
2004
2006
2008
2010
2012
2014
2016

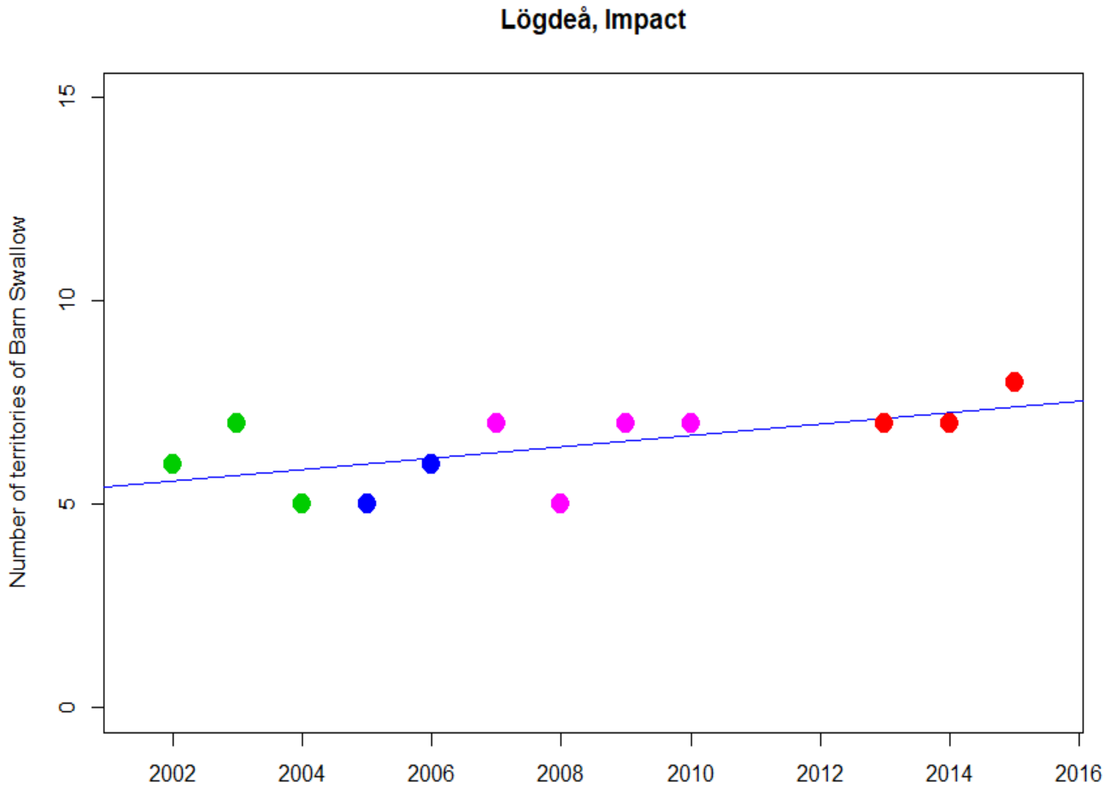
Lögdeå, Impact
Number of territories of Barn Swallow
0
5
10
15
2002
2004
2006
2008
2010
2012
2014
2016

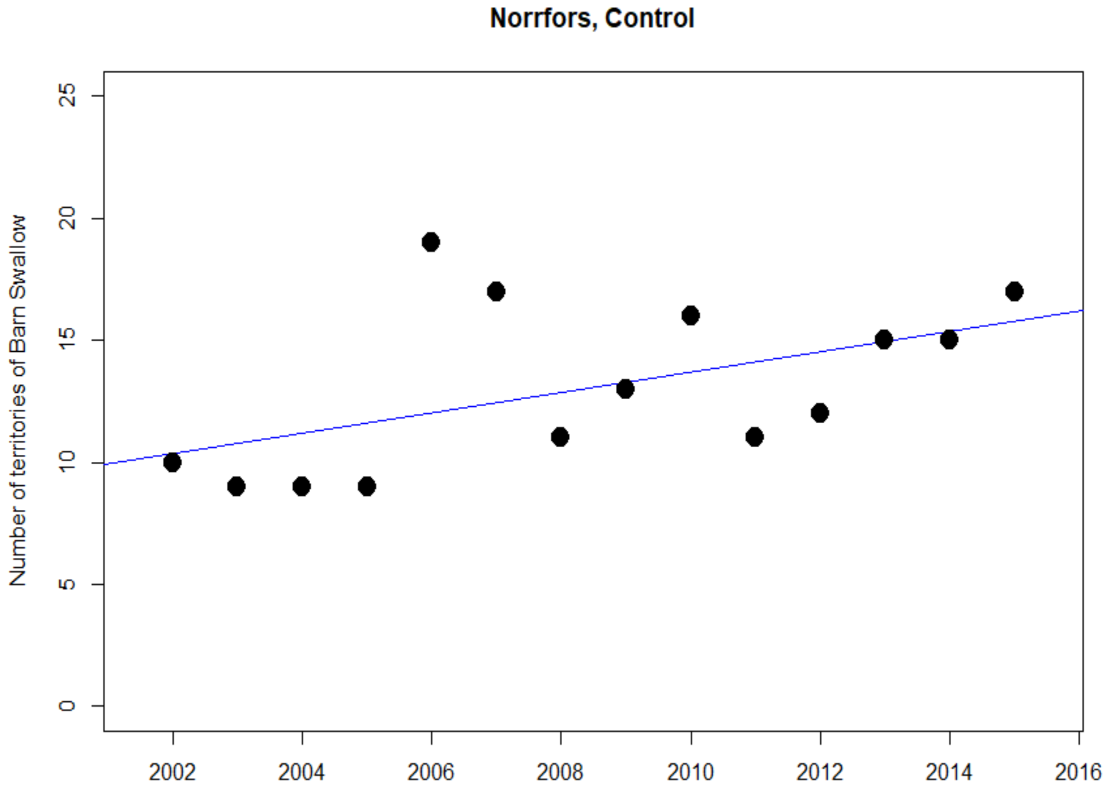
Norrfors, Control
Number of territories of Barn Swallow
0
5
10
15
20
25
2002
2004
2006
2008
2010
2012
2014
2016

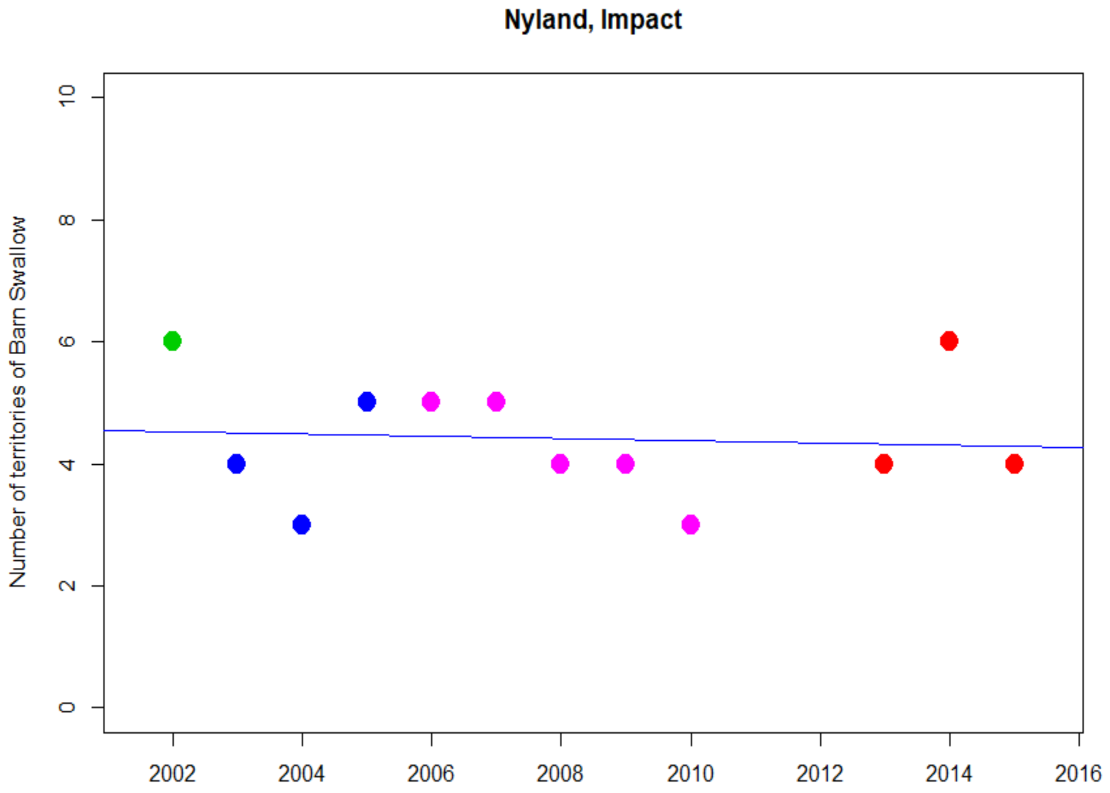
Nyland, Impact
Number of territories of Barn Swallow
0
2
4
6
8
10
2002
2004
2006
2008
2010
2012
2014
2016

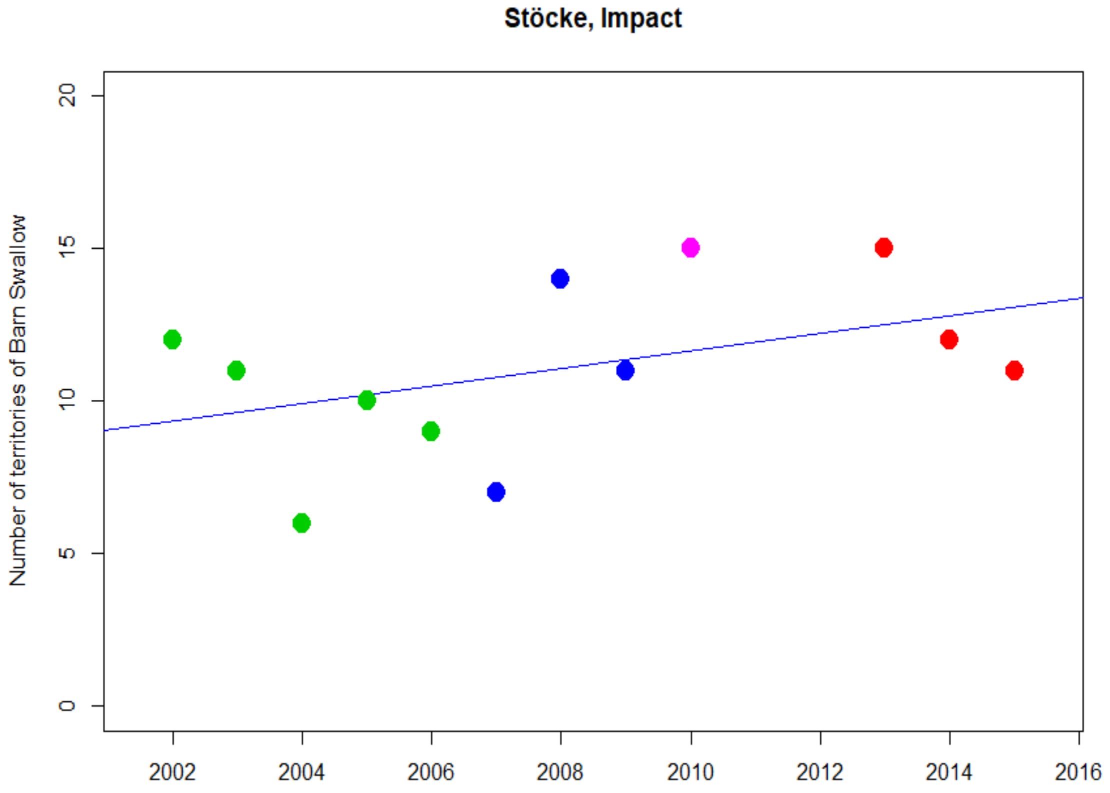
Stöcke, Impact
Number of territories of Barn Swallow
0
5
10
15
20
2002
2004
2006
2008
2010
2012
2014
2016

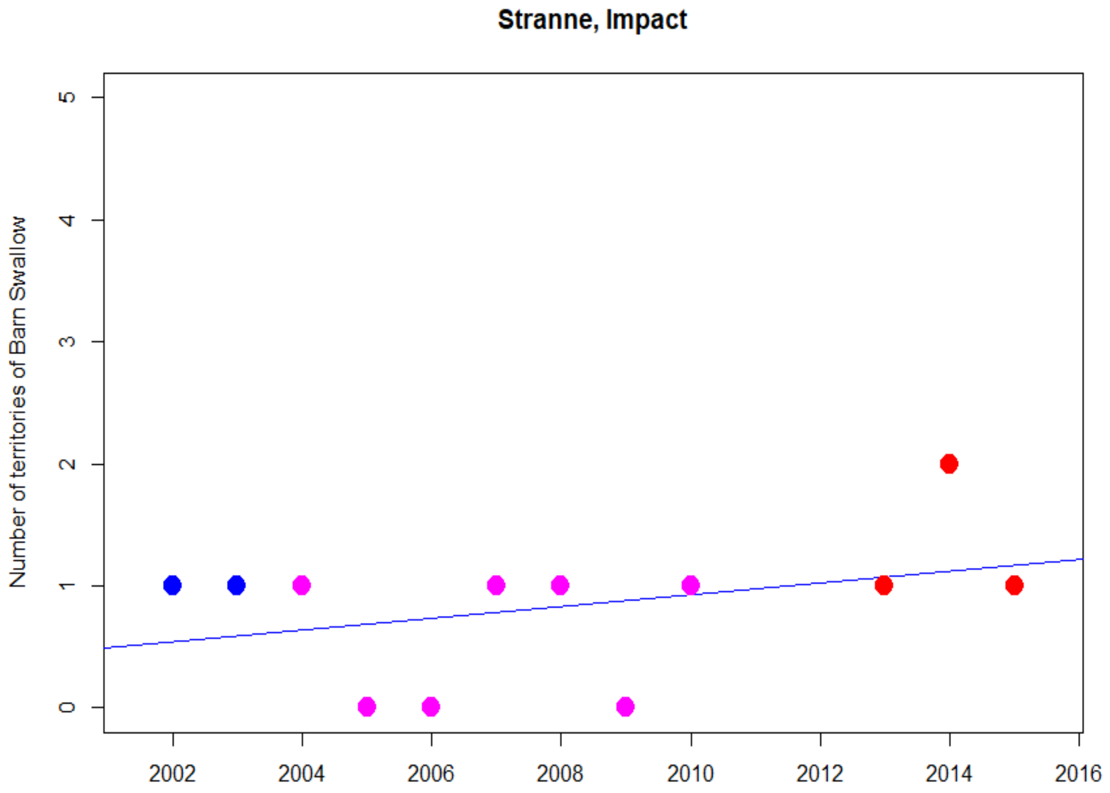
Stranne, Impact
Number of territories of Barn Swallow
0
1
2
3
4
5
2002
2004
2006
2008
2010
2012
2014
2016

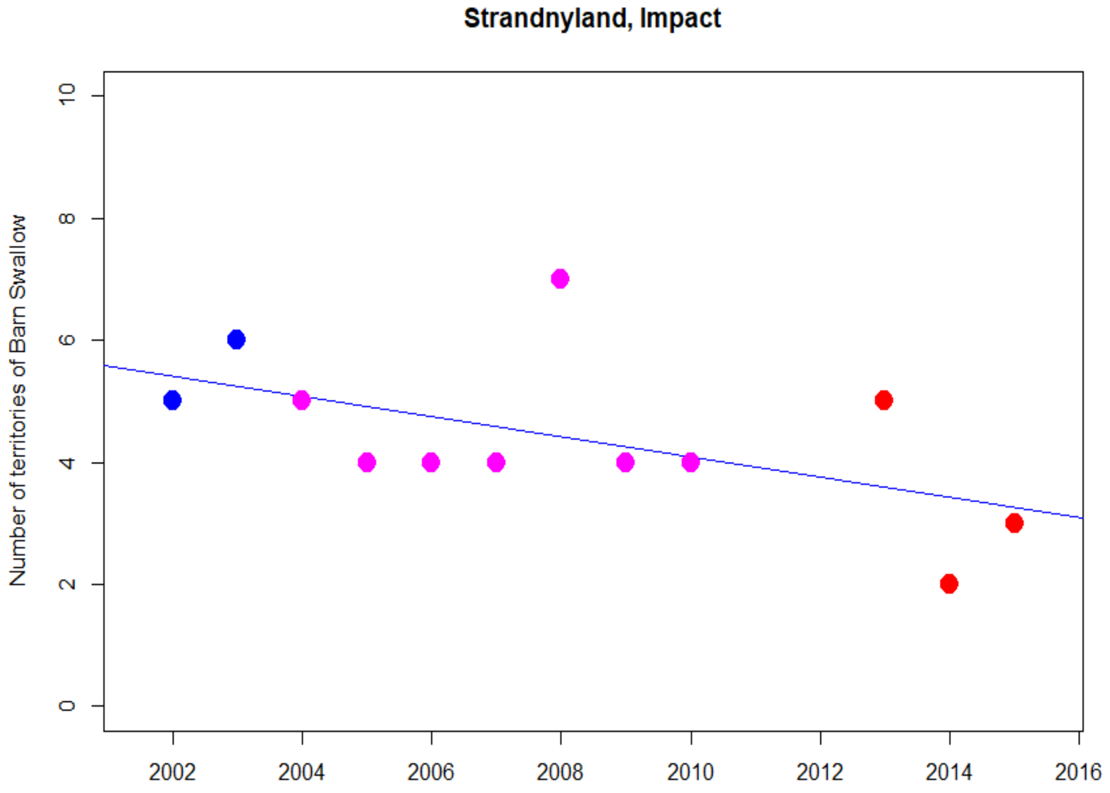
Strandnyland, Impact
Number of territories of Barn Swallow
0
2
4
6
8
10
2002
2004
2006
2008
2010
2012
2014
2016

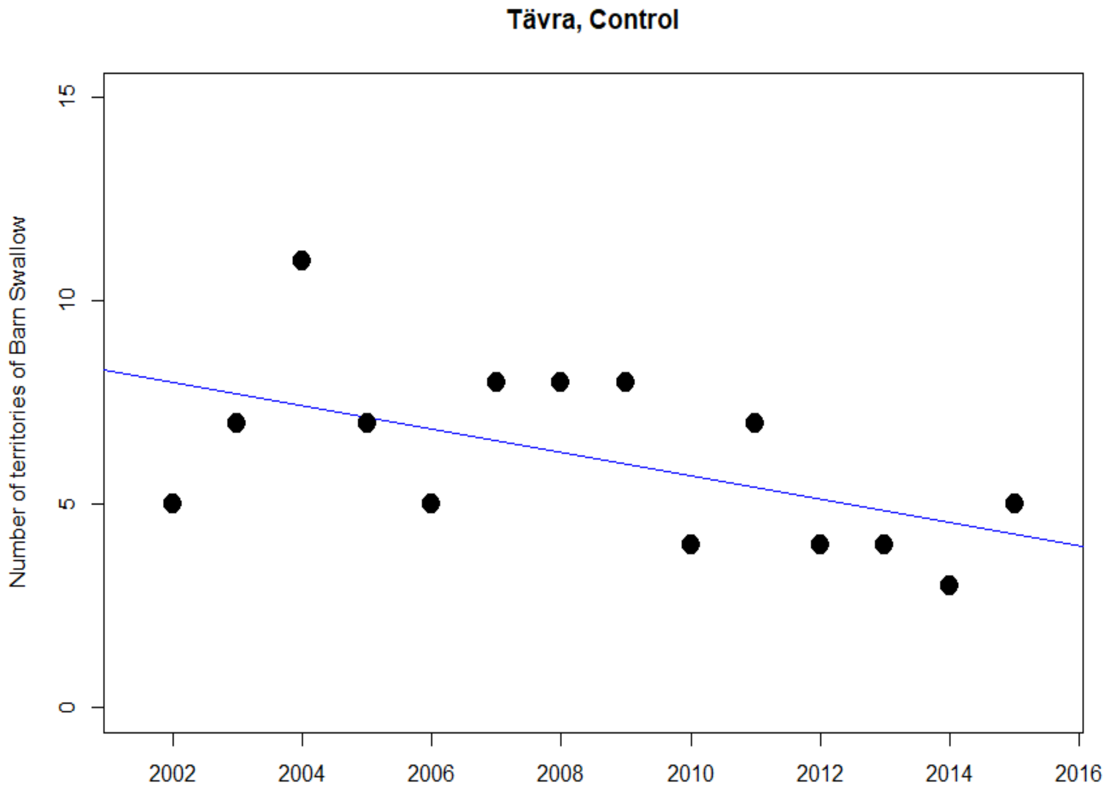
Tävra, Control
Number of territories of Barn Swallow
0
5
10
15
2002
2004
2006
2008
2010
2012
2014
2016

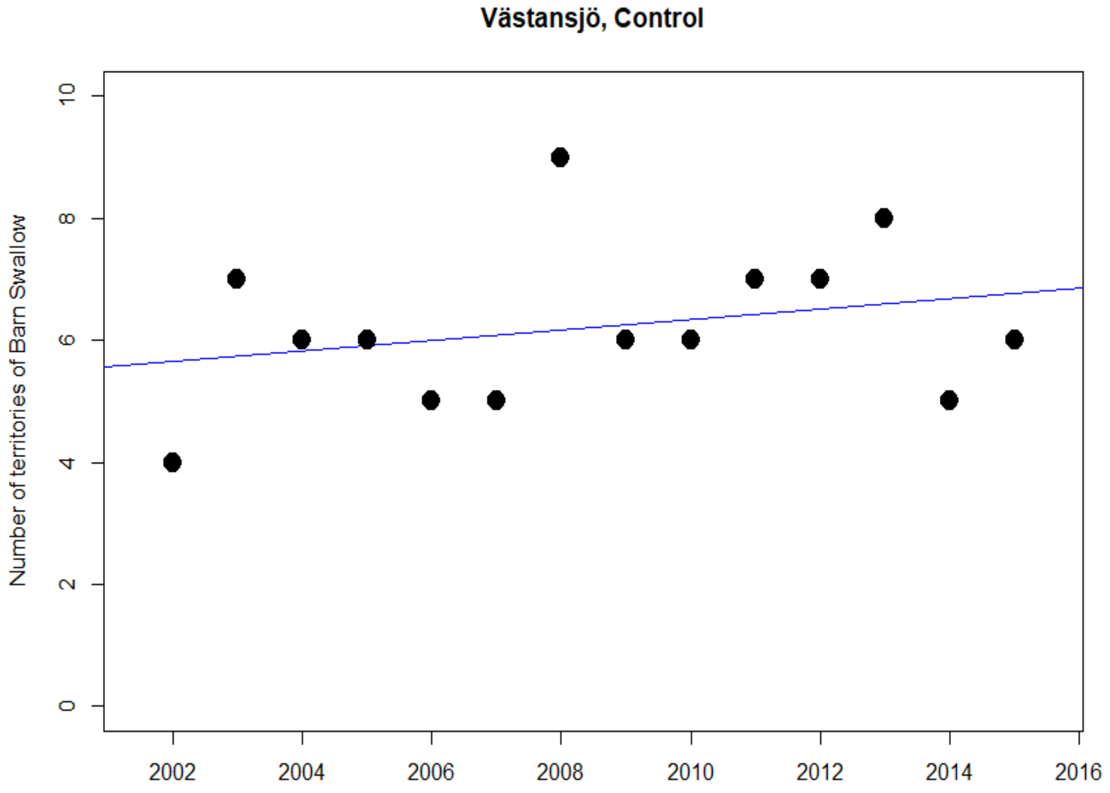
Västansjö, Control
Number of territories of Barn Swallow
0
2
4
6
8
10
2002
2004
2006
2008
2010
2012
2014
2016

### Common Rosefinch

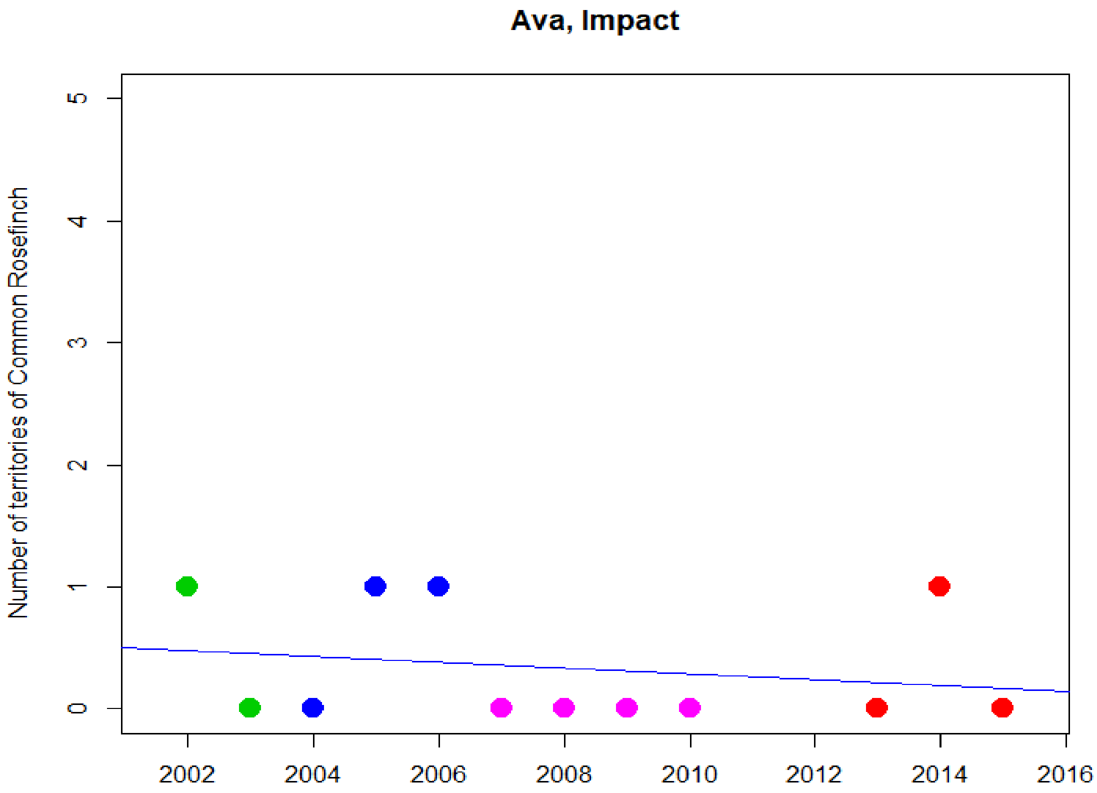


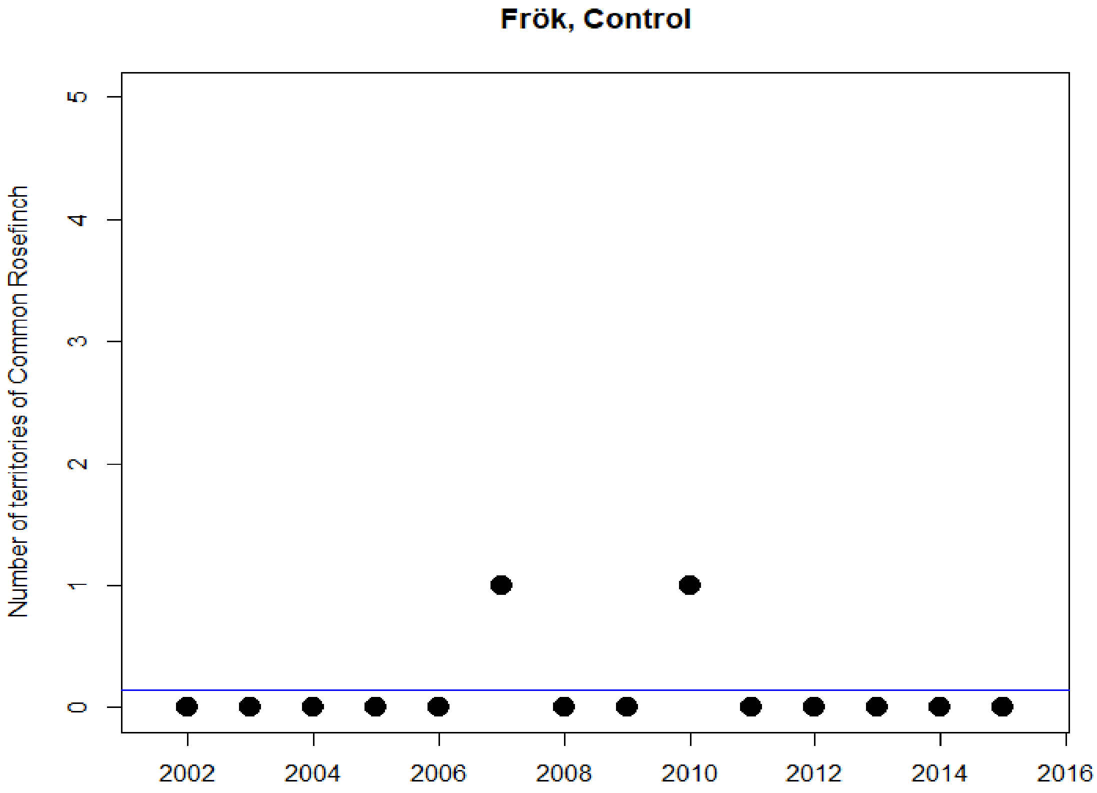

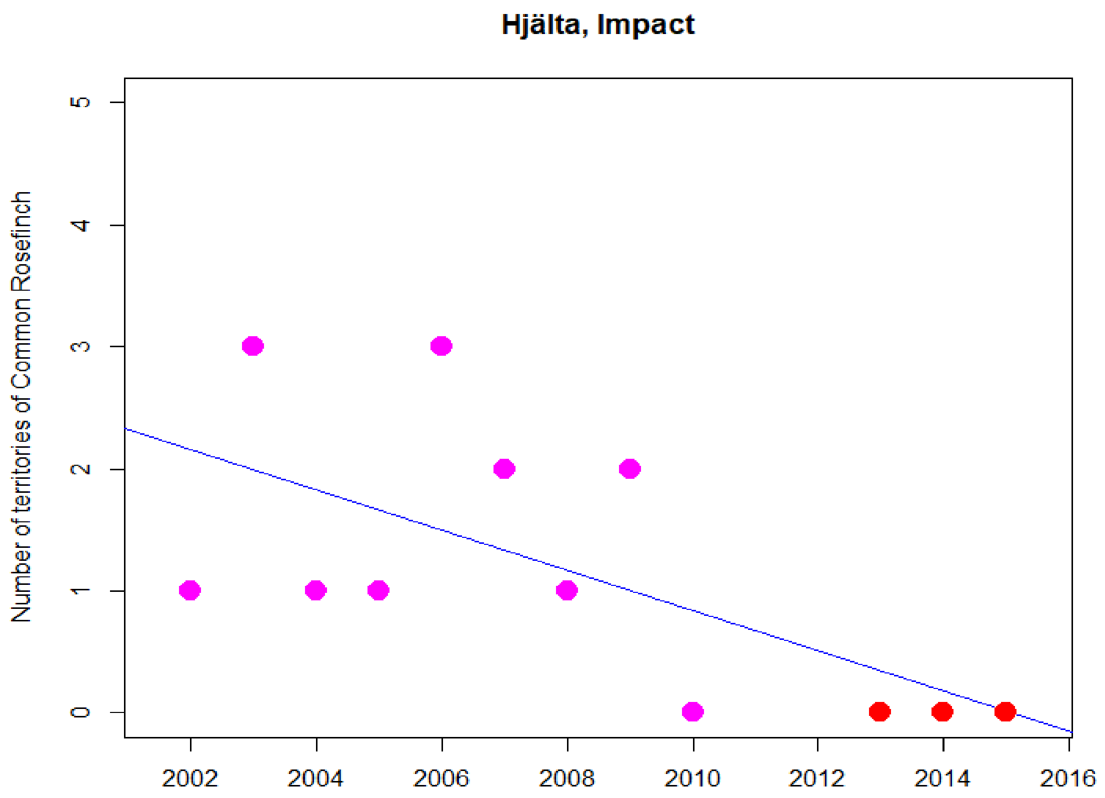
Hjälta, Impact
Number of territories of Common Rosefinch
0
1
2
3
4
5
2002
2004
2006
2008
2010
2012
2014
2016

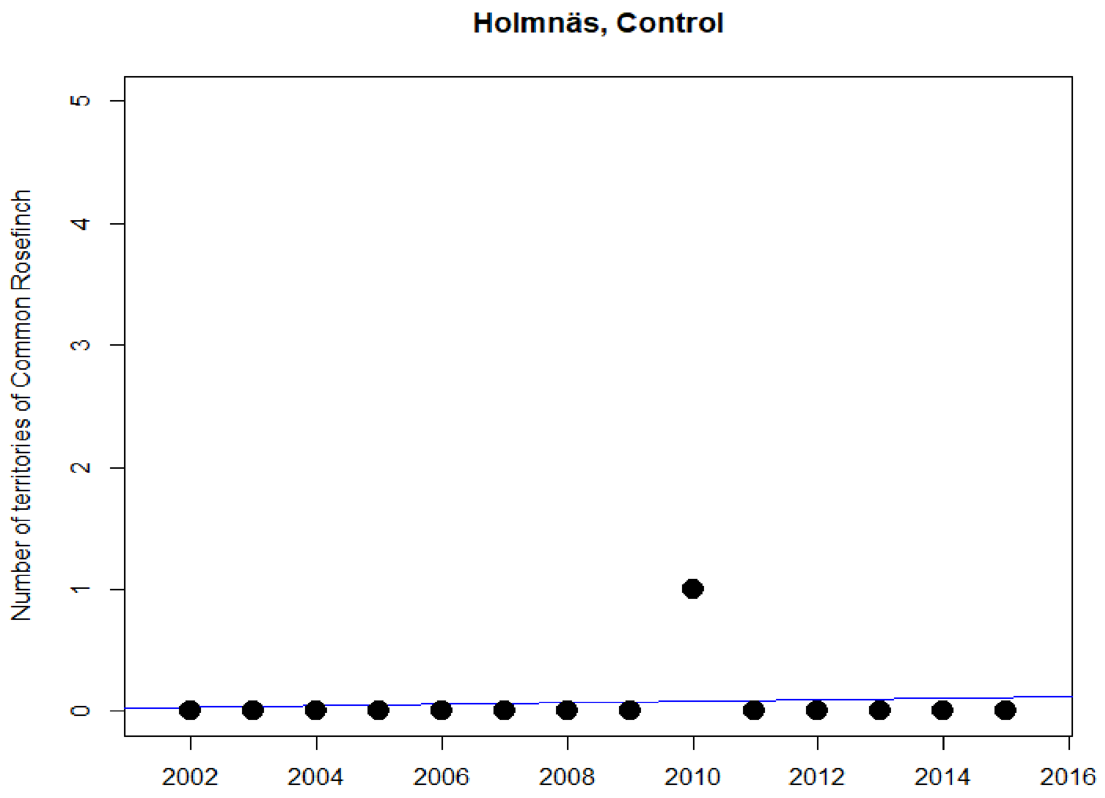
Holmnäs, Control
Number of territories of Common Rosefinch
0
1
2
3
4
5
2002
2004
2006
2008
2010
2012
2014
2016

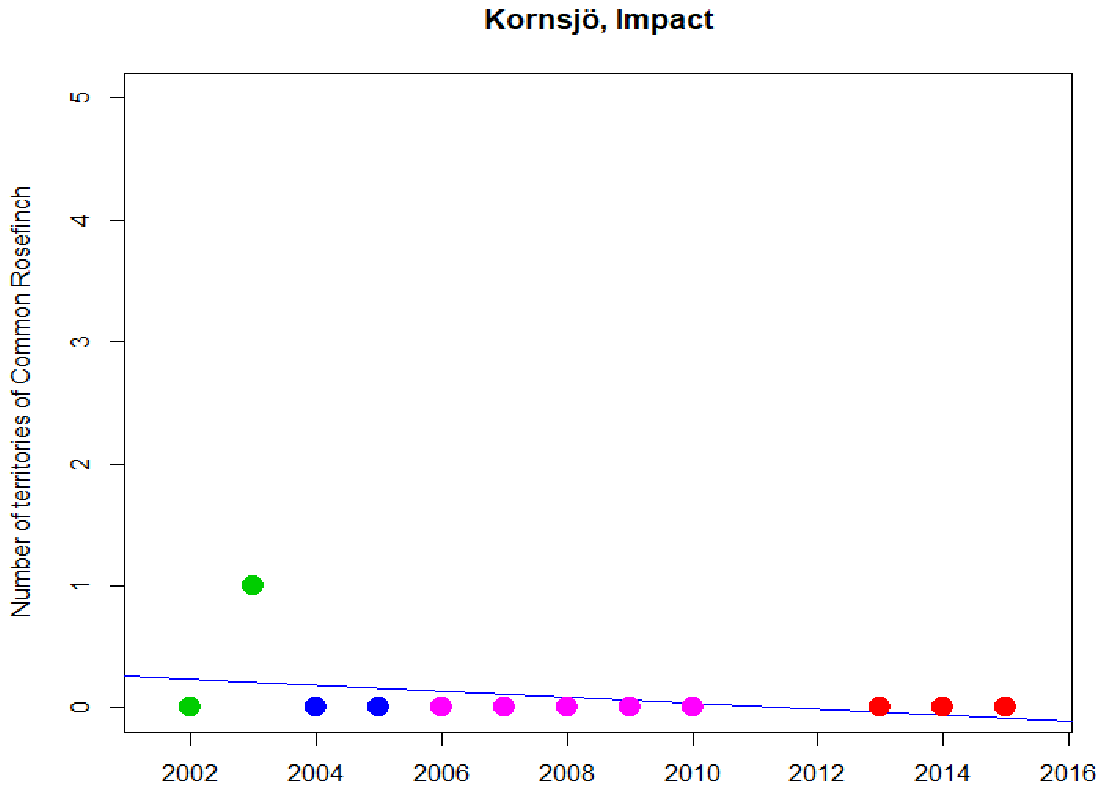
Kornsjö, Impact
Number of territories of Common Rosefinch
0
1
2
3
4
5
2002
2004
2006
2008
2010
2012
2014
2016

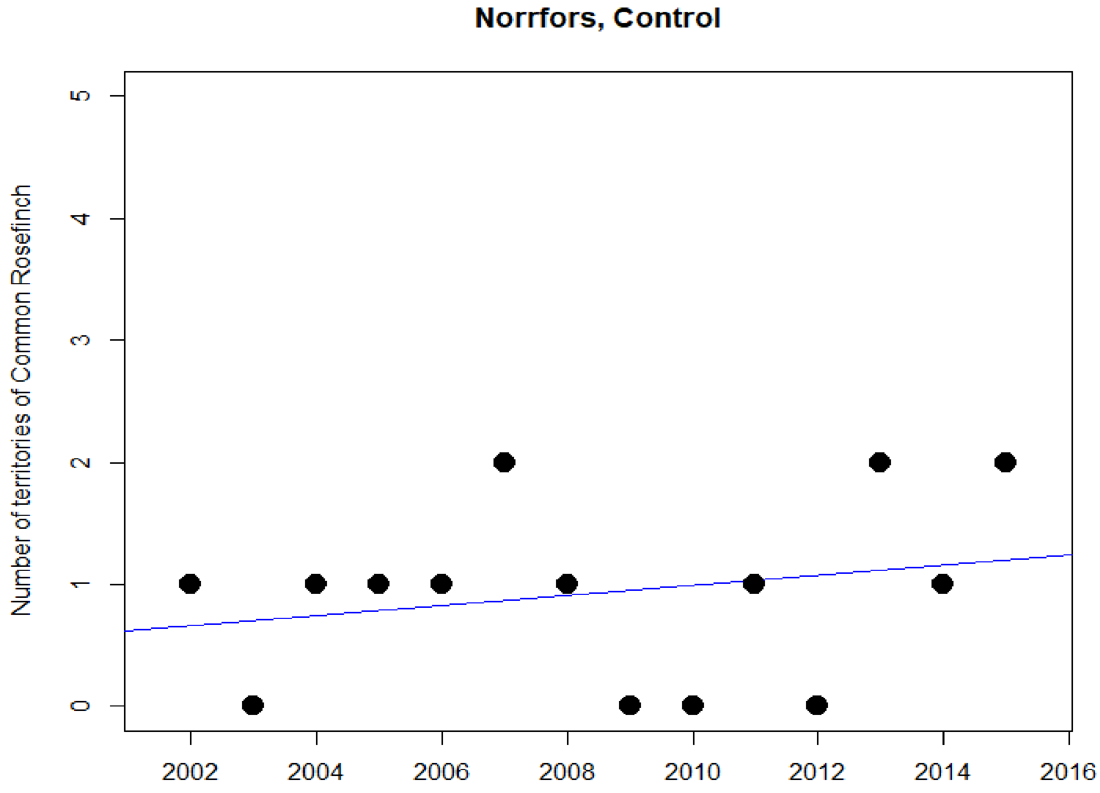
Norrfors, Control
Number of territories of Common Rosefinch
0
1
2
3
4
5
2002
2004
2006
2008
2010
2012
2014
2016

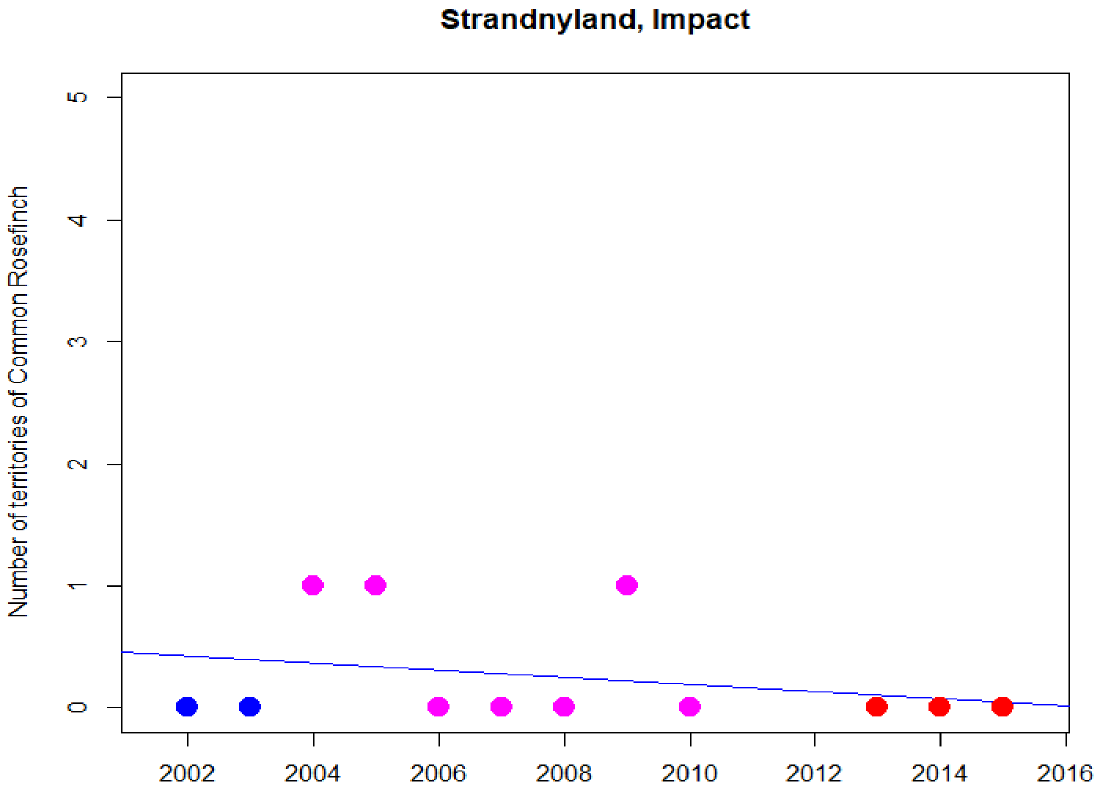
Strandnyland, Impact
Number of territories of Common Rosefinch
0
1
2
3
4
5
2002
2004
2006
2008
2010
2012
2014
2016

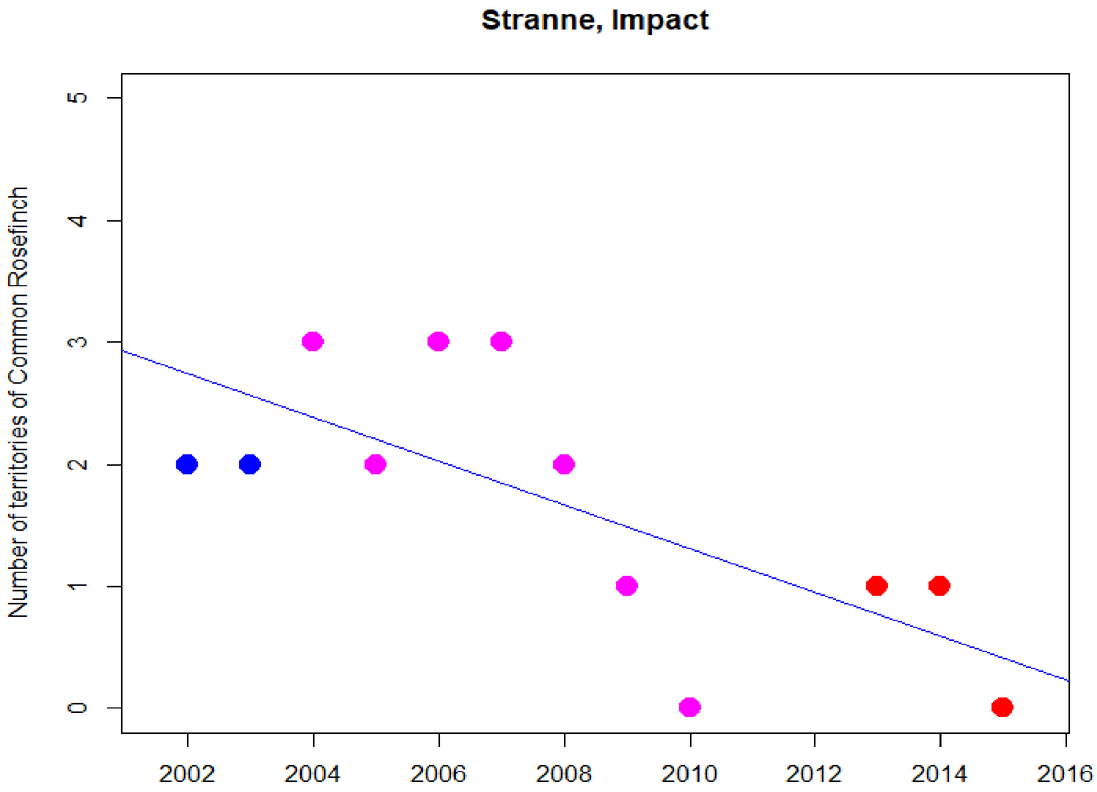
Stranne, Impact
Number of territories of Common Rosefinch
0
1
2
3
4
5
2002
2004
2006
2008
2010
2012
2014
2016

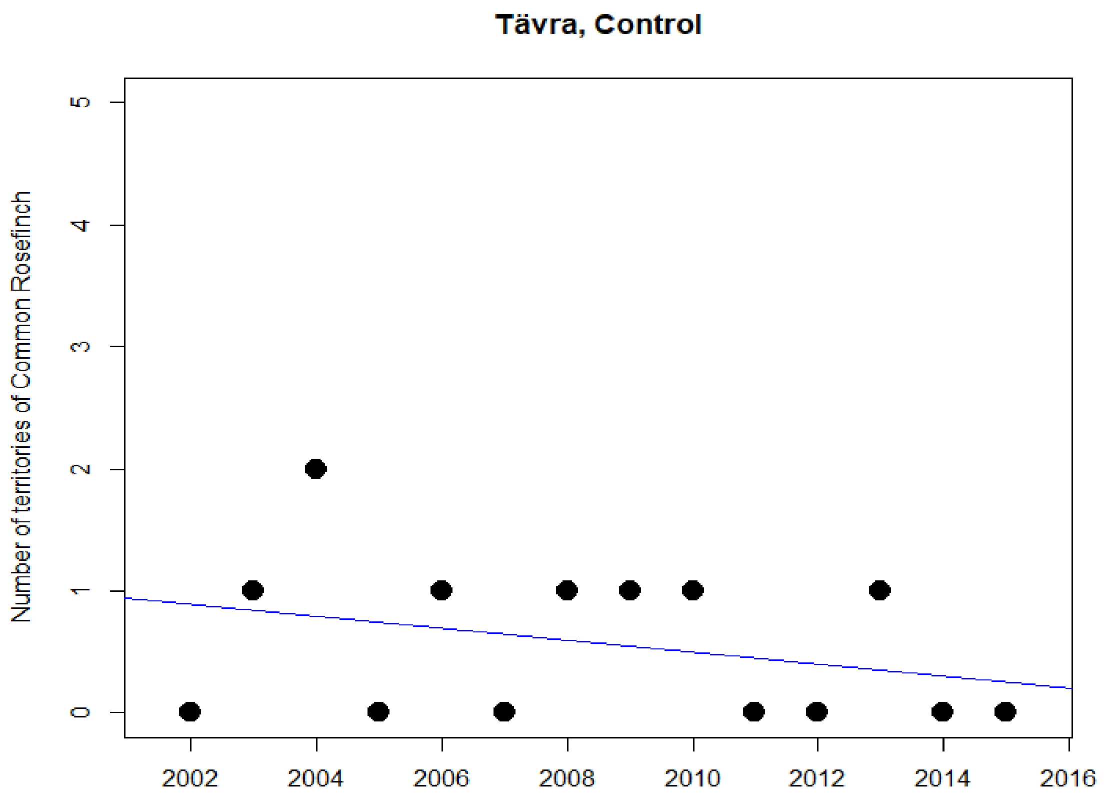
Tävra, Control
Number of territories of Common Rosefinch
0
1
2
3
4
5
2002
2004
2006
2008
2010
2012
2014
2016

### Common Snipe

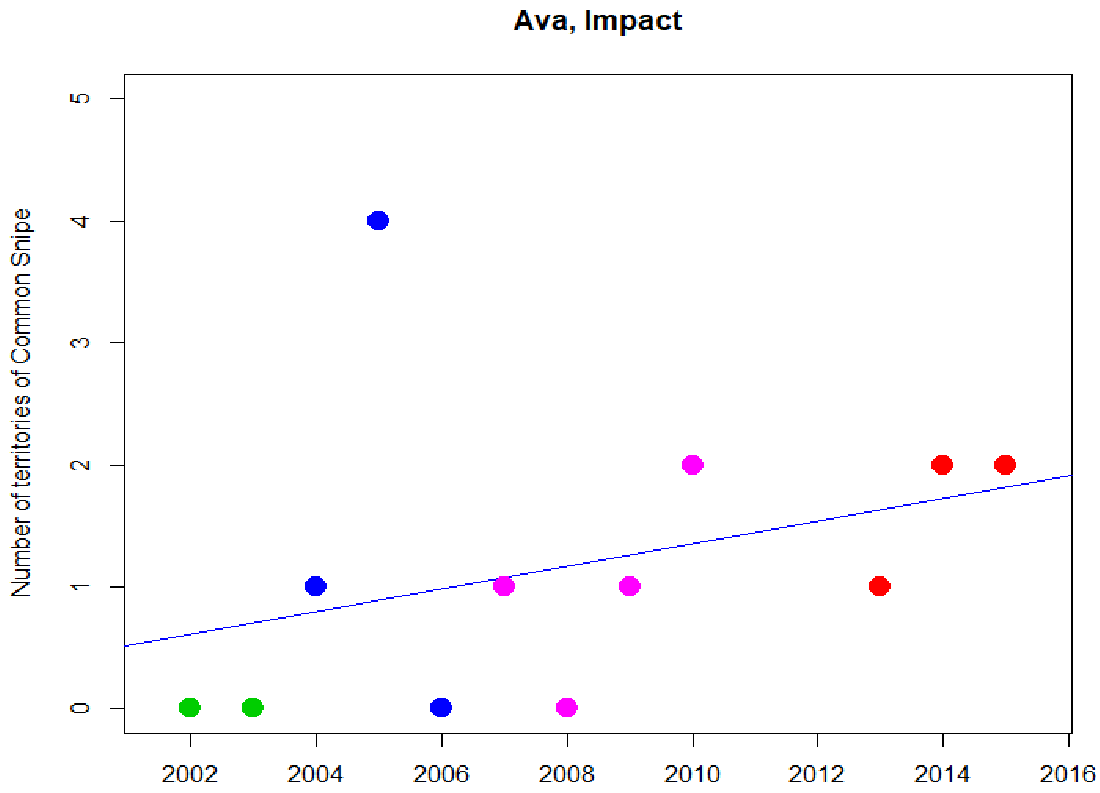


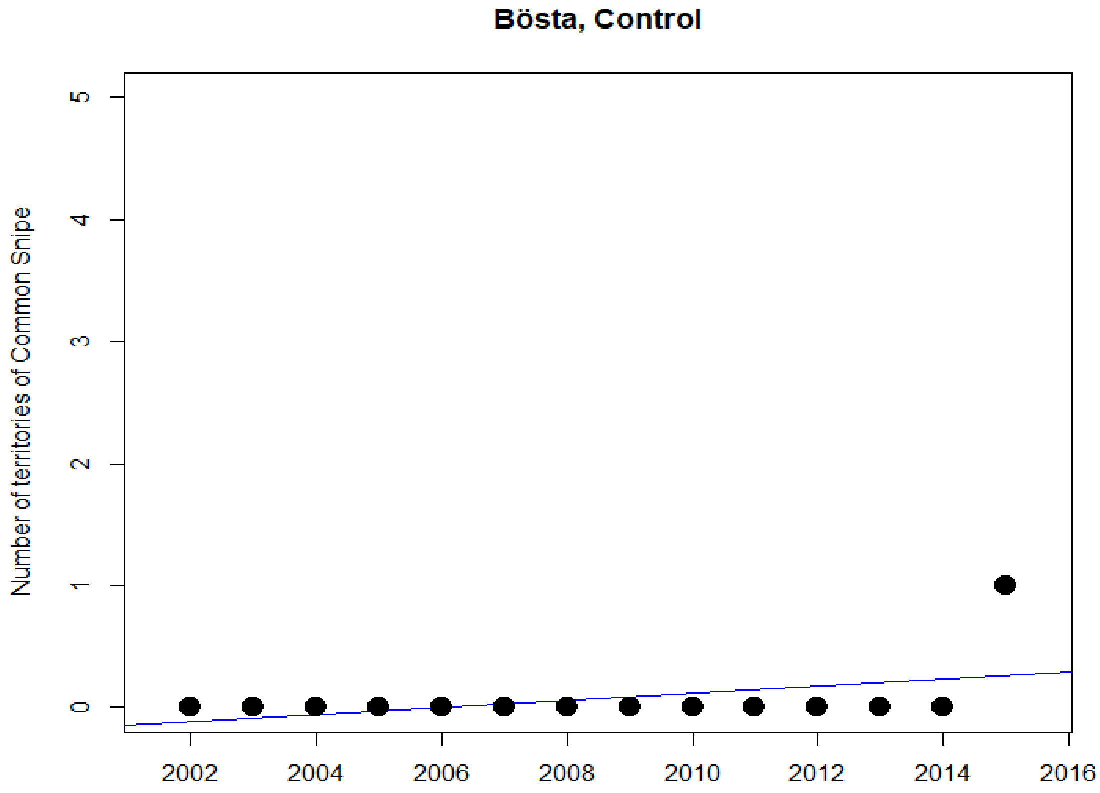

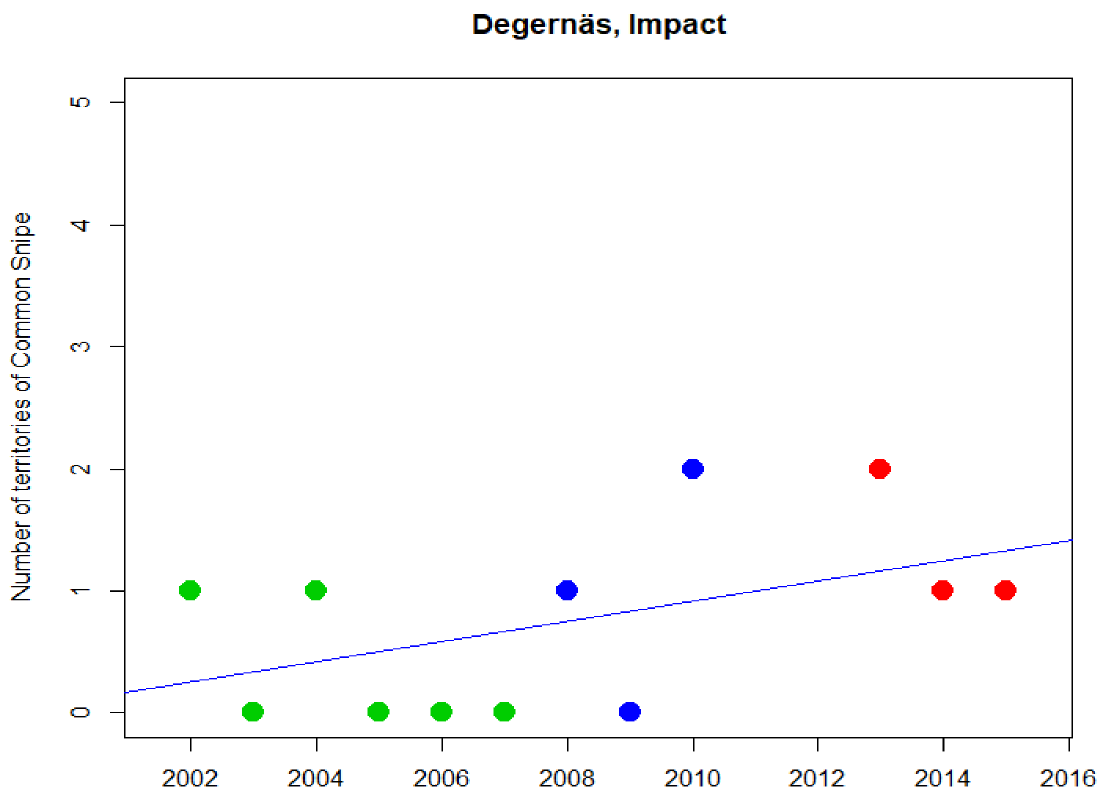

Degernäs, Impact
Number of territories of Common Snipe
0
1
2
3
4
5
2002
2004
2006
2008
2010
2012
2014
2016


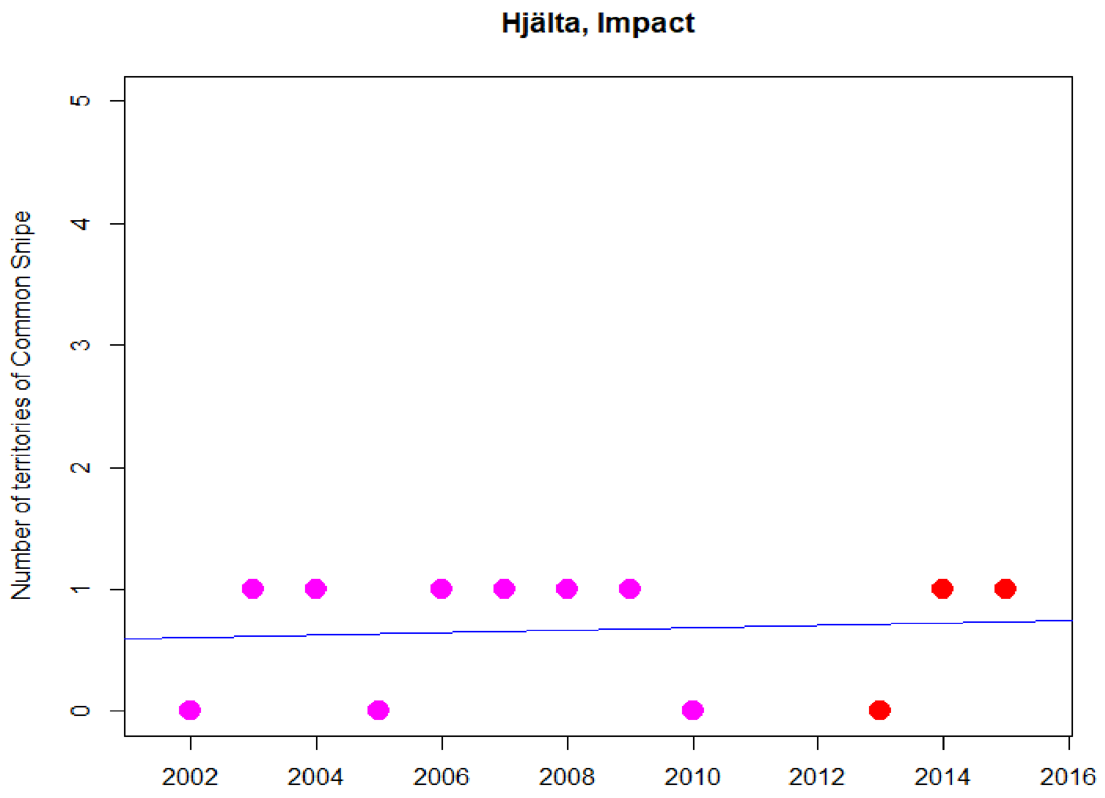

Hjälta, Impact
Number of territories of Common Snipe
0
1
2
3
4
5
2002
2004
2006
2008
2010
2012
2014
2016

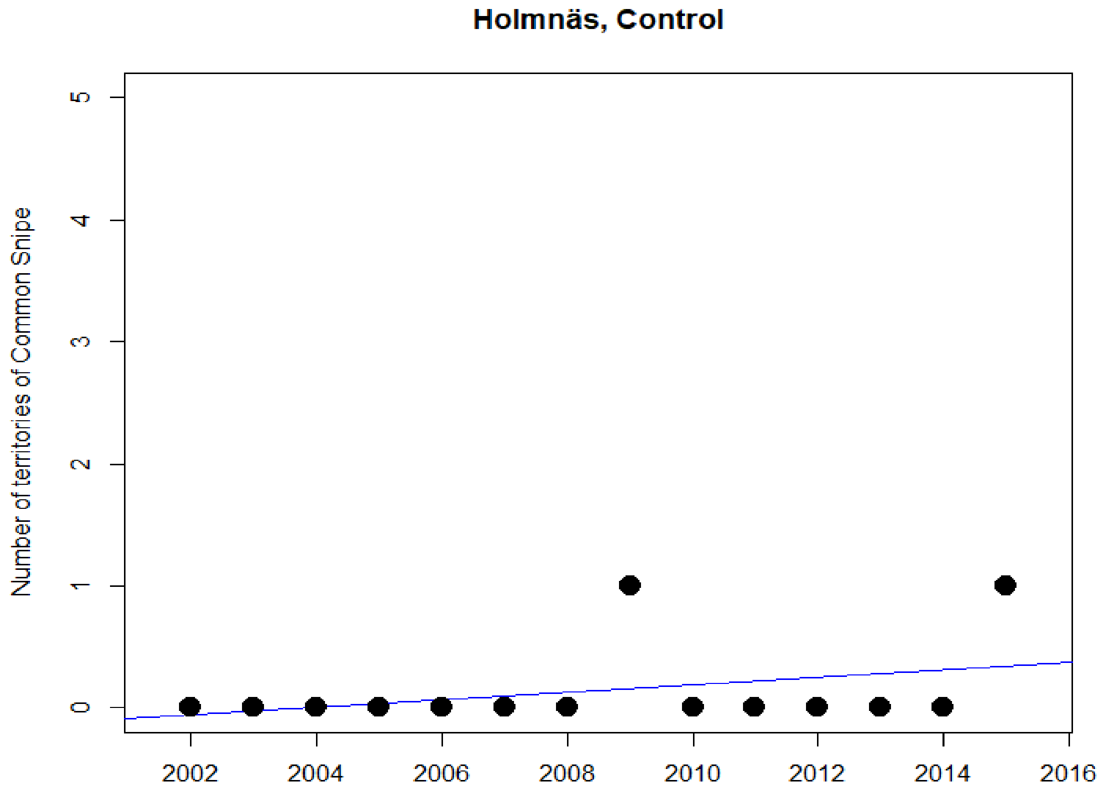
Holmnäs, Control
Number of territories of Common Snipe
0
1
2
3
4
5
2002
2004
2006
2008
2010
2012
2014
2016

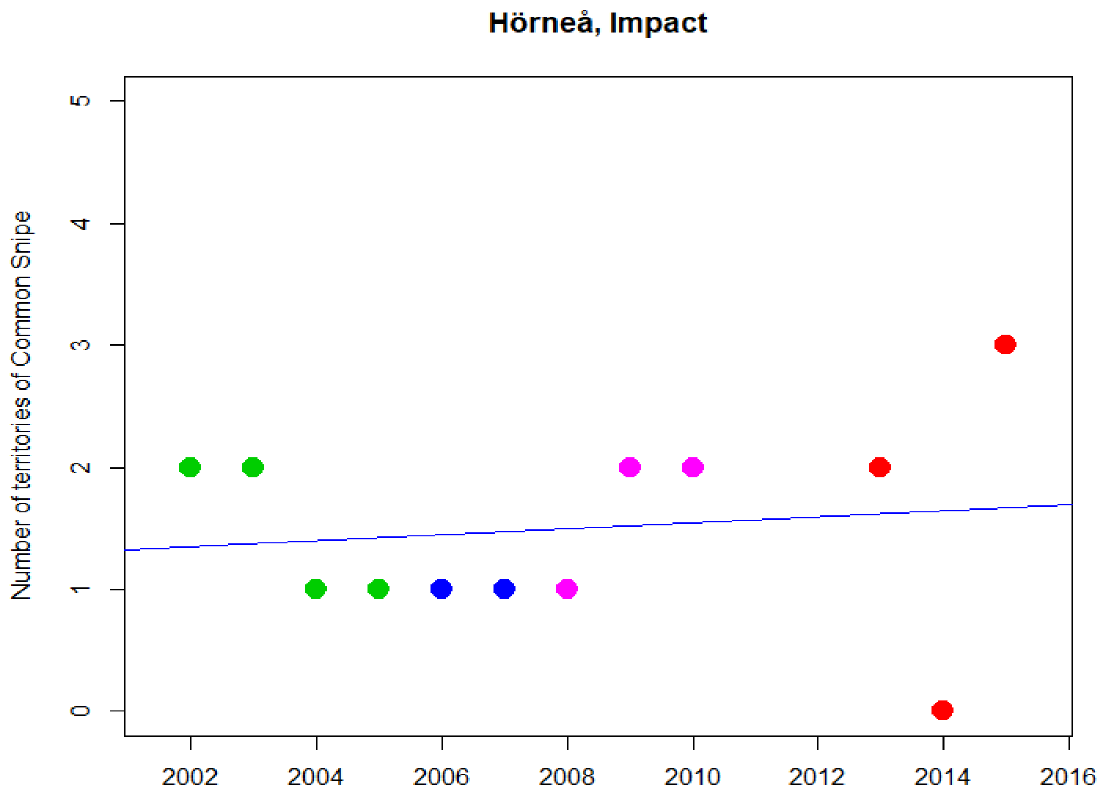
Hörneå, Impact
Number of territories of Common Snipe
0
1
2
3
4
5
2002
2004
2006
2008
2010
2012
2014
2016

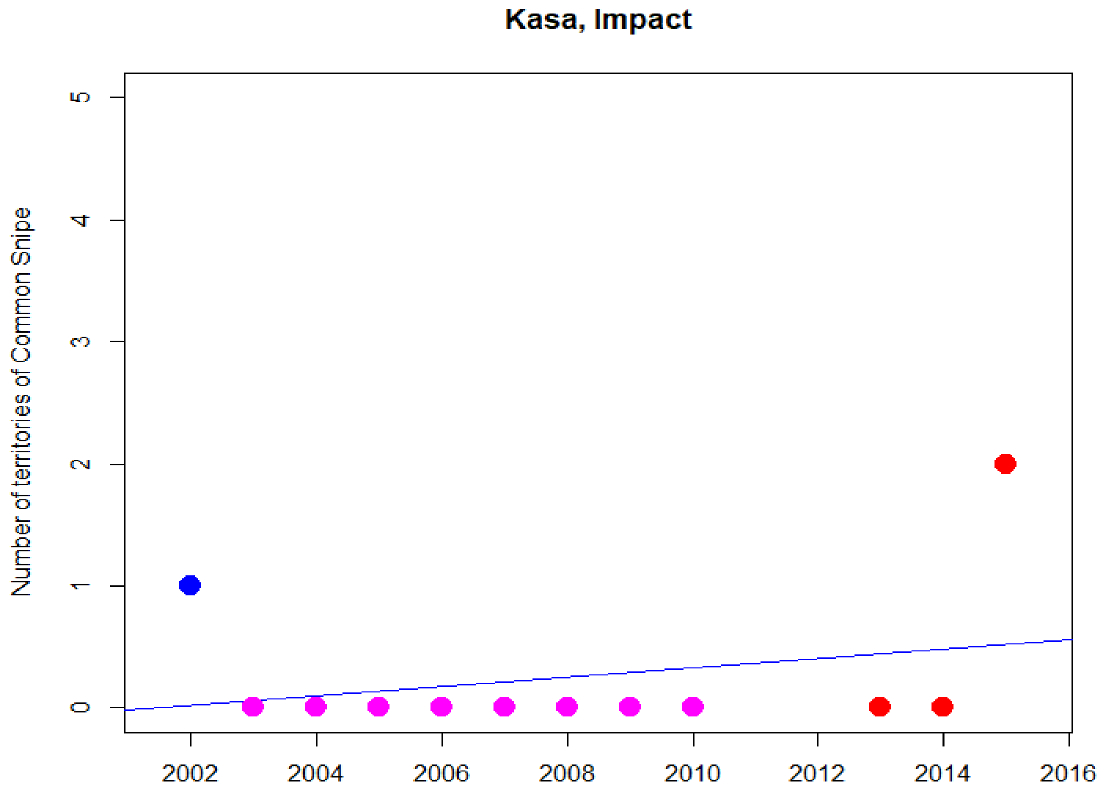
Kasa, Impact
Number of territories of Common Snipe
0
1
2
3
4
5
2002
2004
2006
2008
2010
2012
2014
2016

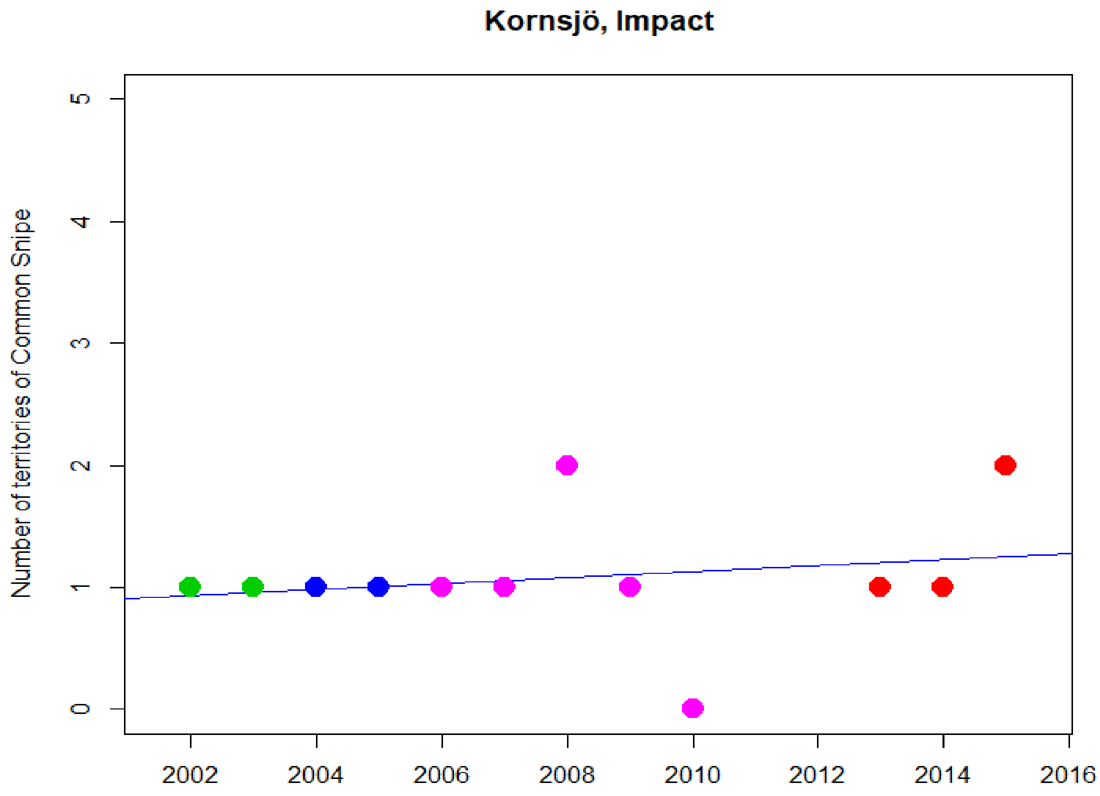
Kornsjö, Impact
Number of territories of Common Snipe
0
1
2
3
4
5
2002
2004
2006
2008
2010
2012
2014
2016

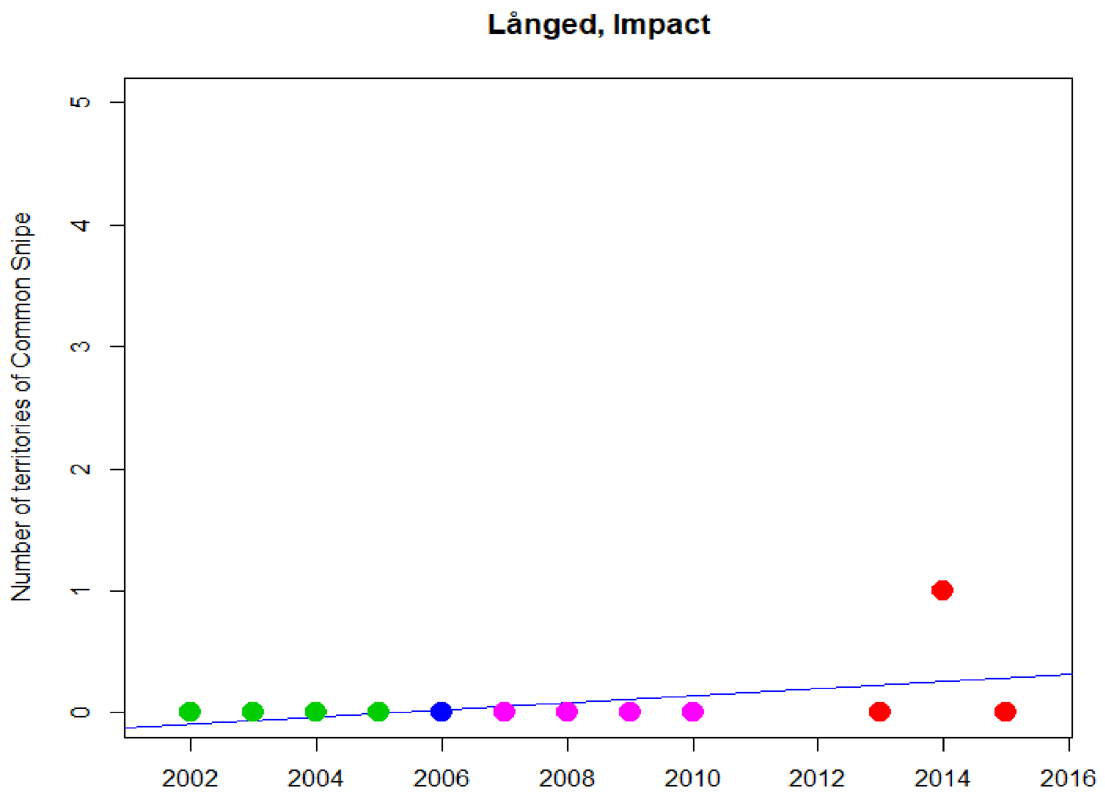
Långed, Impact
Number of territories of Common Snipe
0
1
2
3
4
5
2002
2004
2006
2008
2010
2012
2014
2016

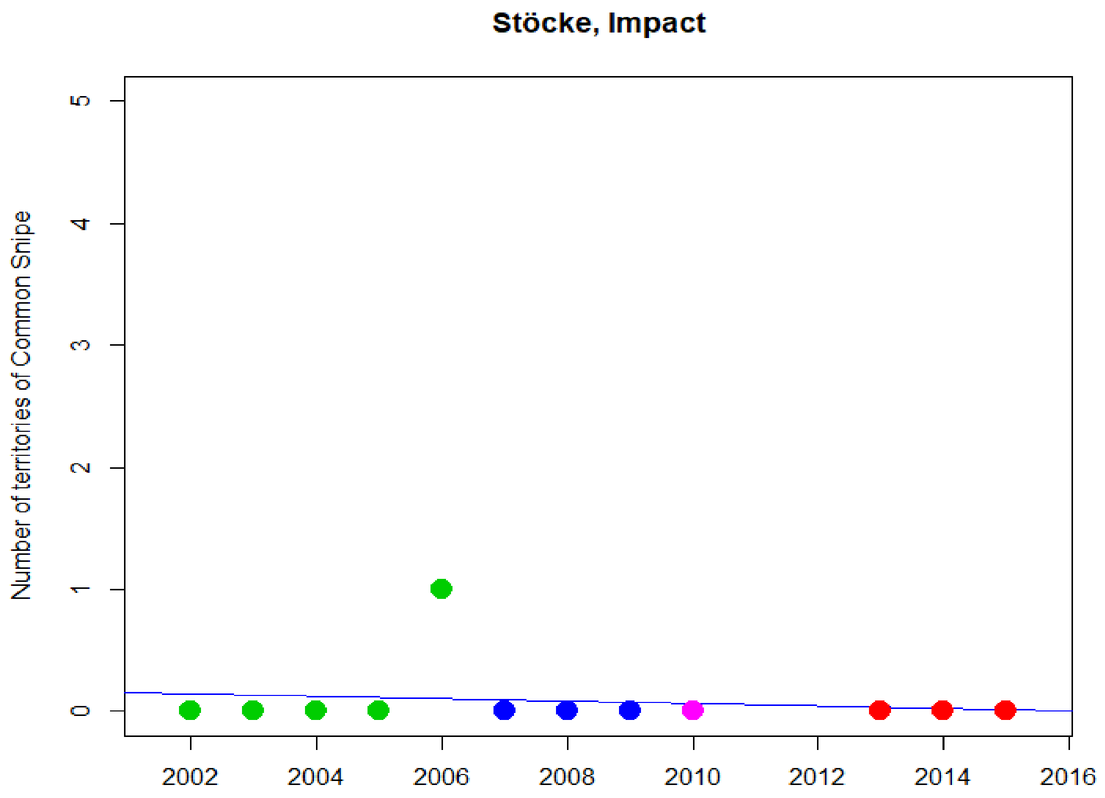
Stöcke, Impact
Number of territories of Common Snipe
0
1
2
3
4
5
2002
2004
2006
2008
2010
2012
2014
2016

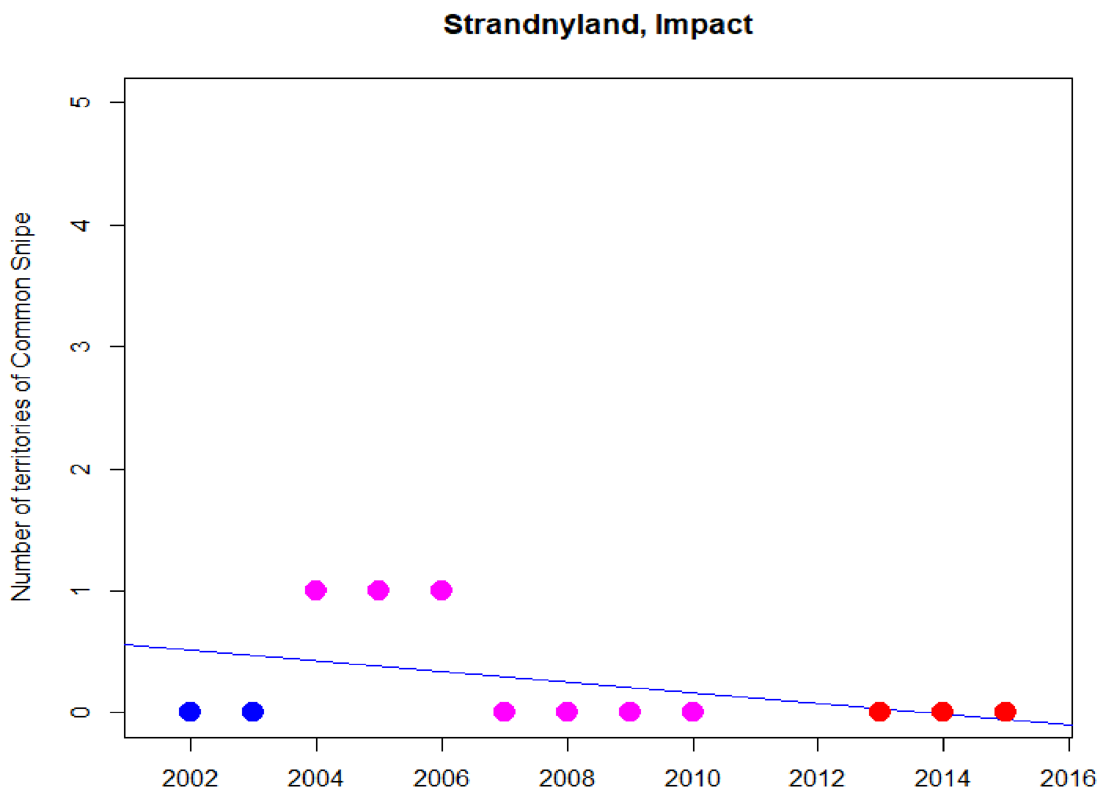
Strandnyland, Impact
Number of territories of Common Snipe
0
1
2
3
4
5
2002
2004
2006
2008
2010
2012
2014
2016

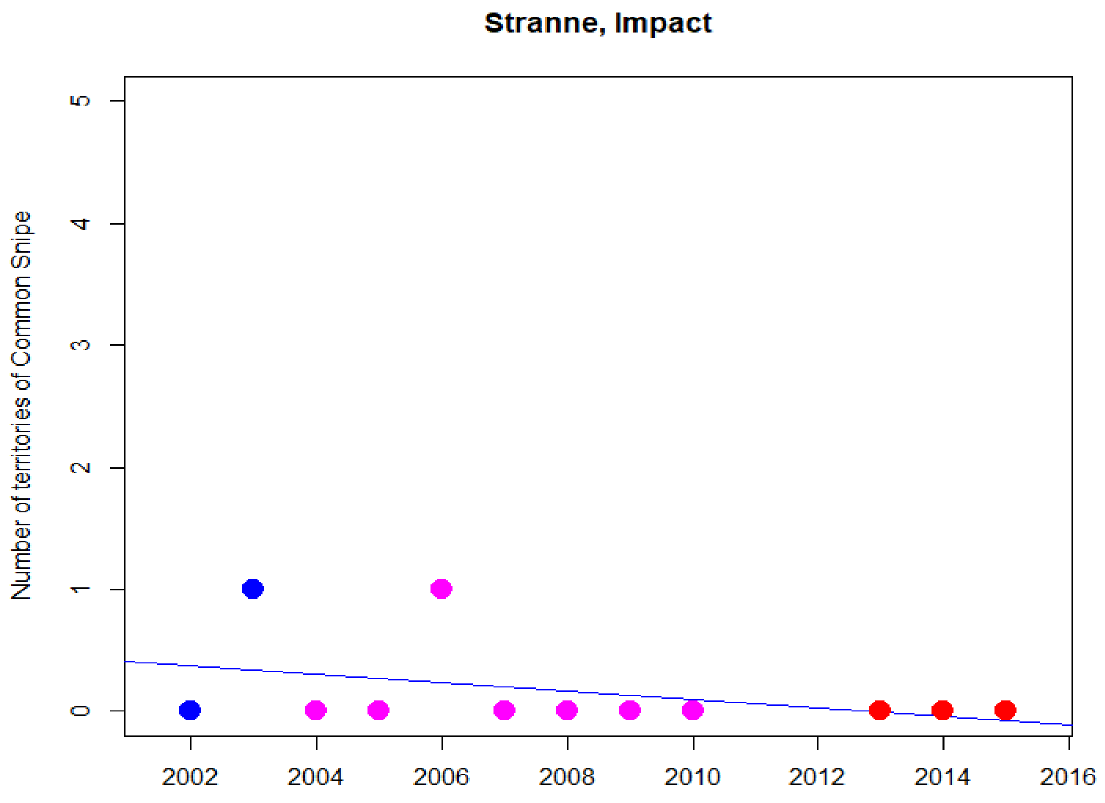
Stranne, Impact
Number of territories of Common Snipe
0
1
2
3
4
5
2002
2004
2006
2008
2010
2012
2014
2016

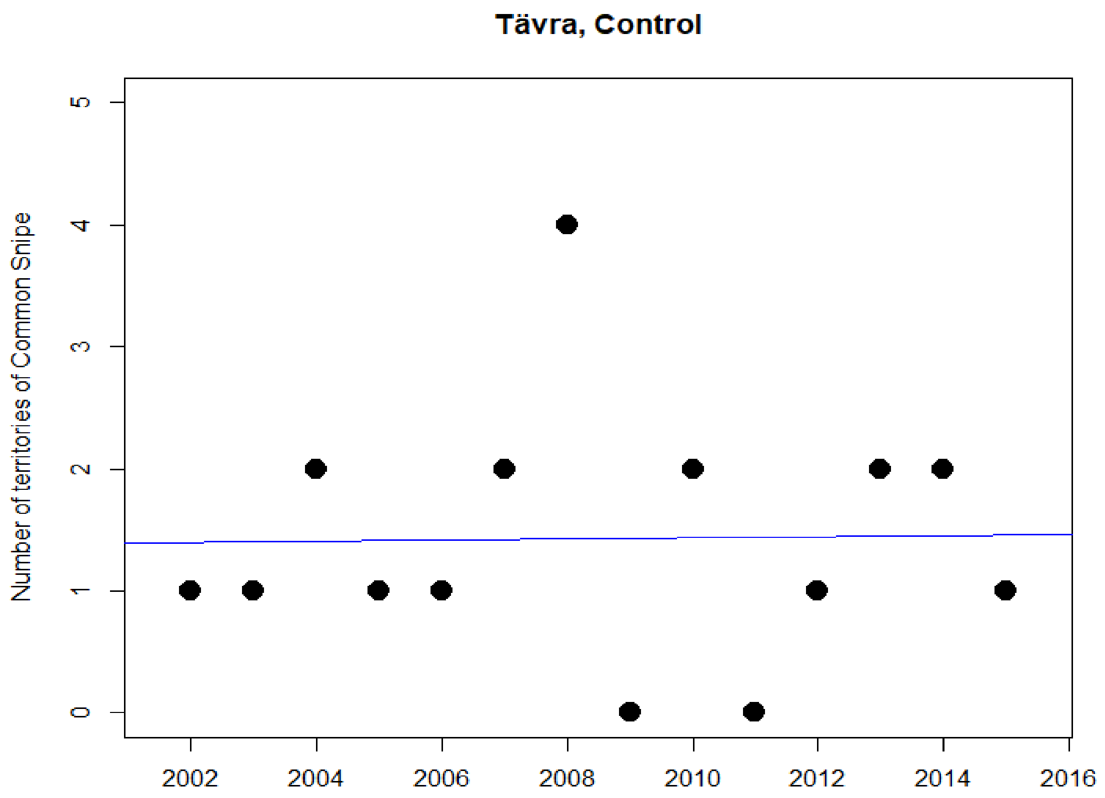
Tävra, Control
Number of territories of Common Snipe
0
1
2
3
4
5
2002
2004
2006
2008
2010
2012
2014
2016

### Common Starling

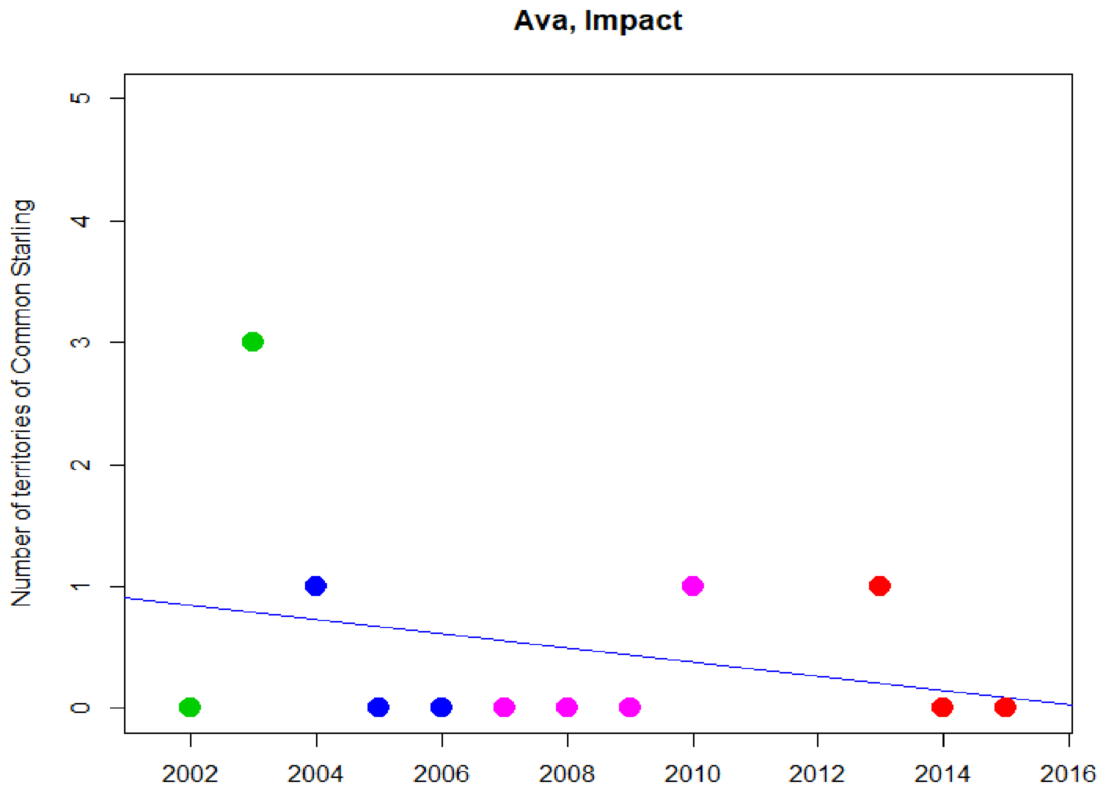


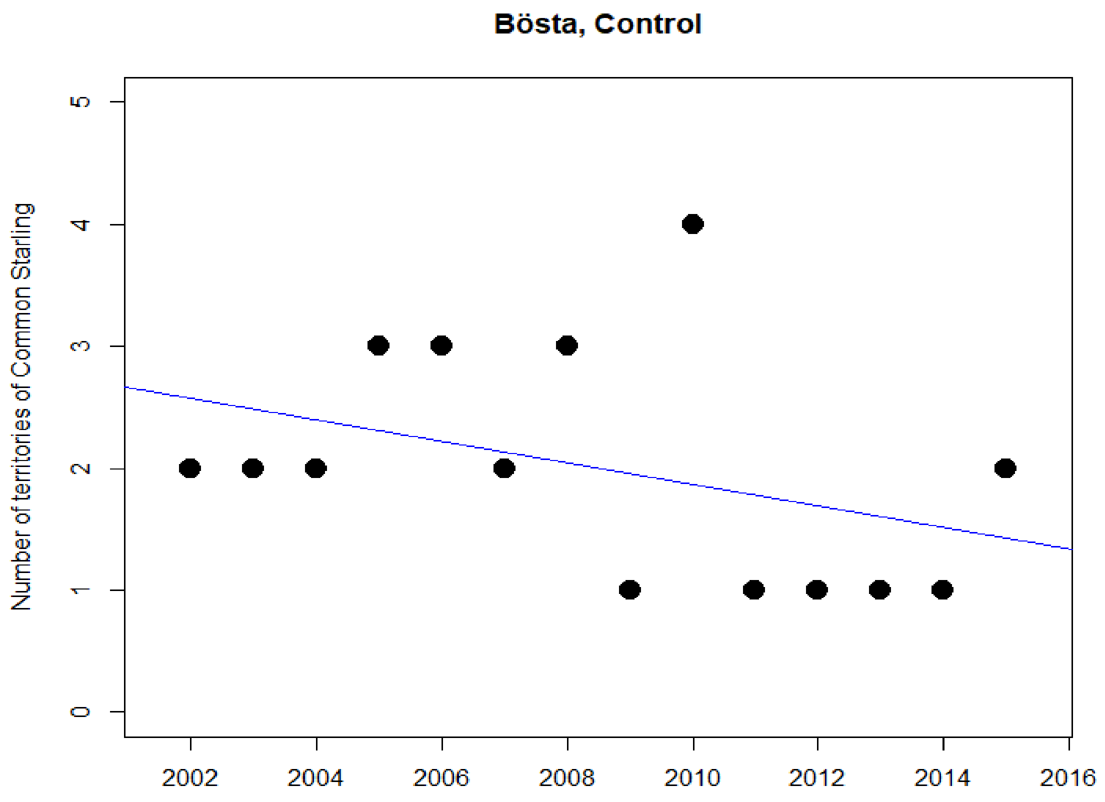

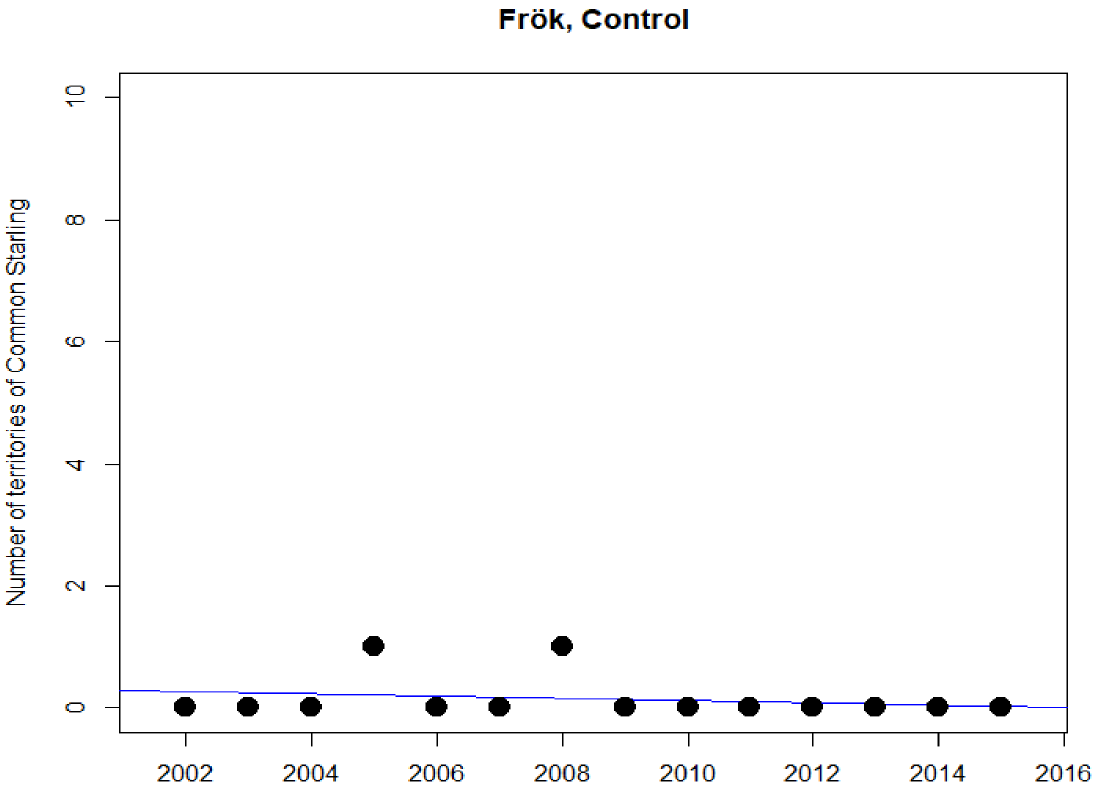

Frök, Control
Number of territories of Common Starling
0
2
4
6
8
10
2002
2004
2006
2008
2010
2012
2014
2016


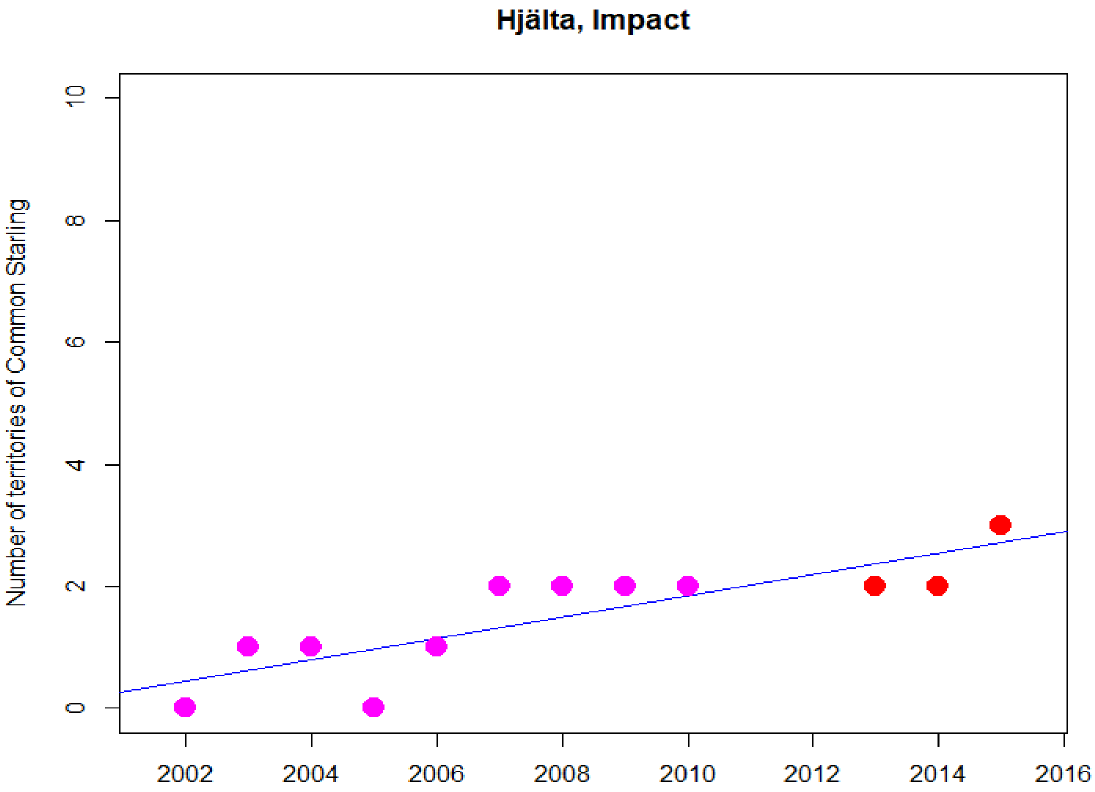

Hjälta, Impact
Number of territories of Common Starling
0
2
4
6
8
10
2002
2004
2006
2008
2010
2012
2014
2016

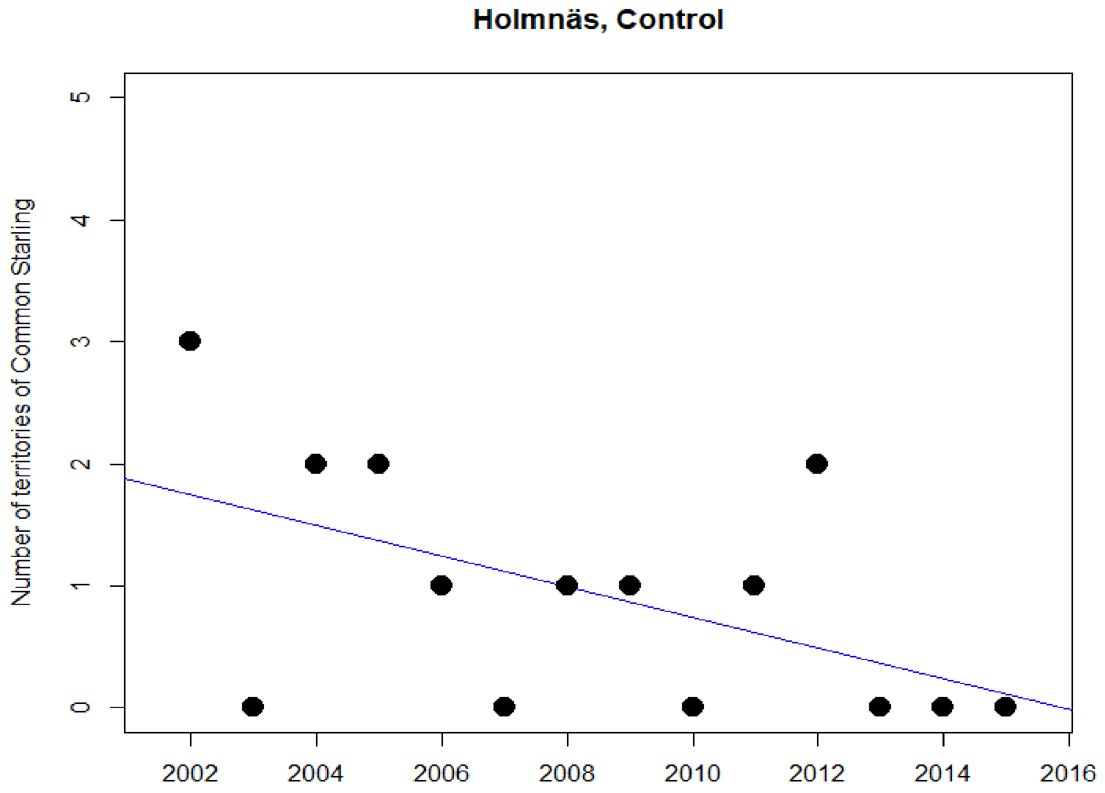
Holmnäs, Control
Number of territories of Common Starling
0
1
2
3
4
5
2002
2004
2006
2008
2010
2012
2014
2016

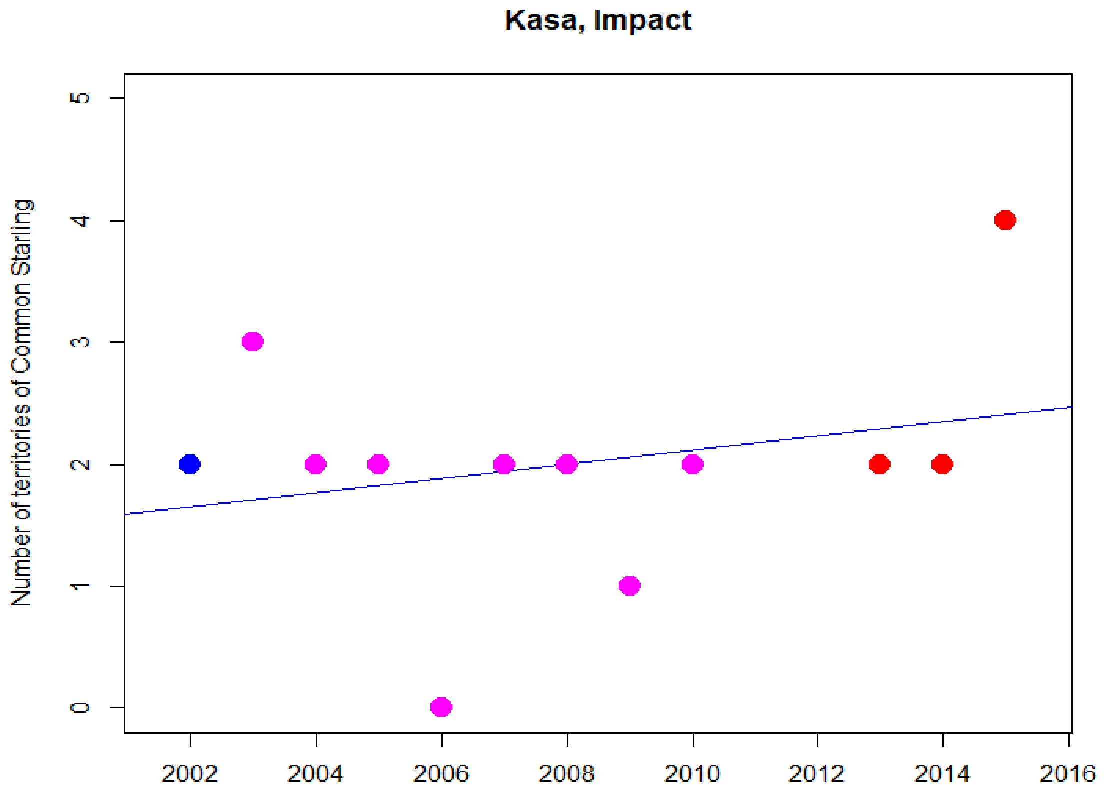
Kasa, Impact
Number of territories of Common Starling
0
1
2
3
4
5
2002
2004
2006
2008
2010
2012
2014
2016

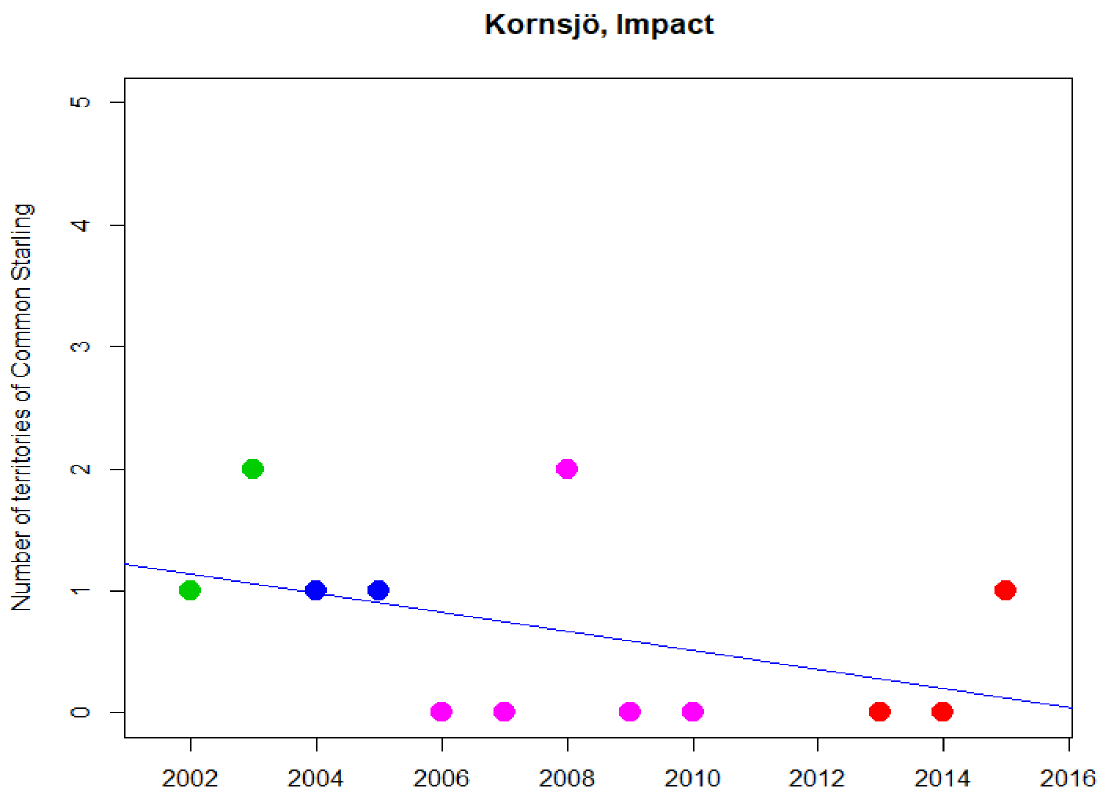
Kornsjö, Impact
Number of territories of Common Starling
0
1
2
3
4
5
2002
2004
2006
2008
2010
2012
2014
2016

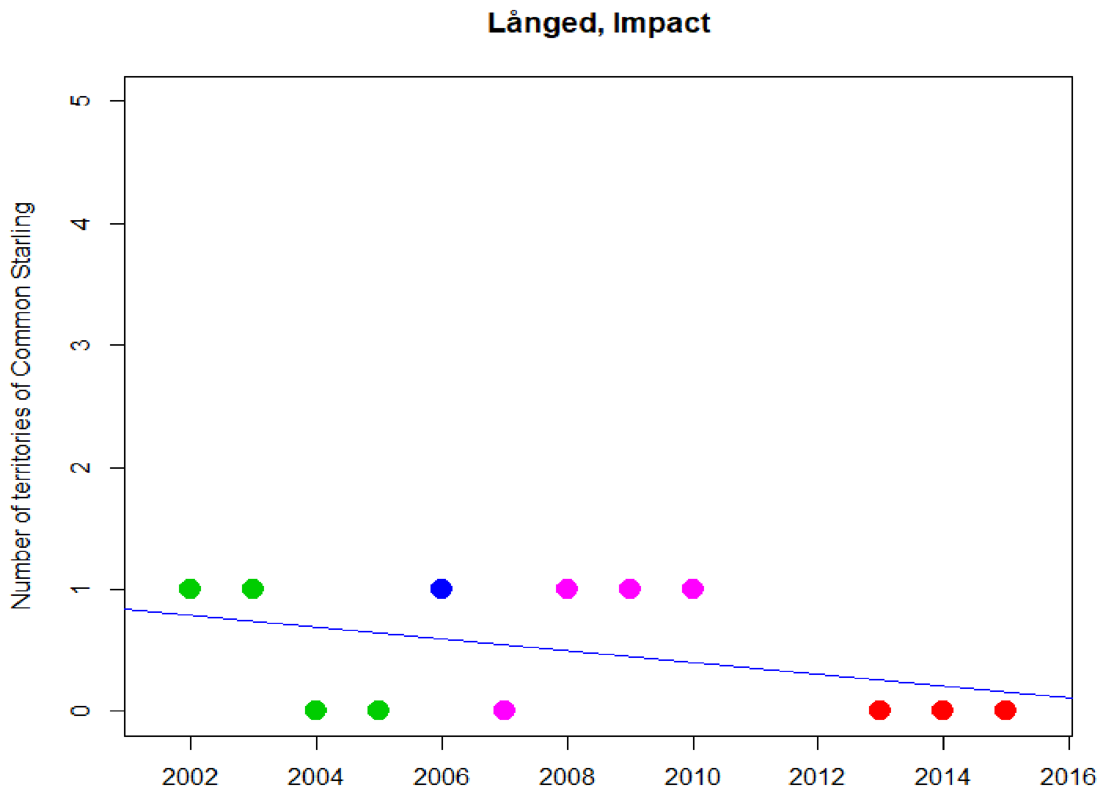
Långed, Impact
Number of territories of Common Starling
0
1
2
3
4
5
2002
2004
2006
2008
2010
2012
2014
2016

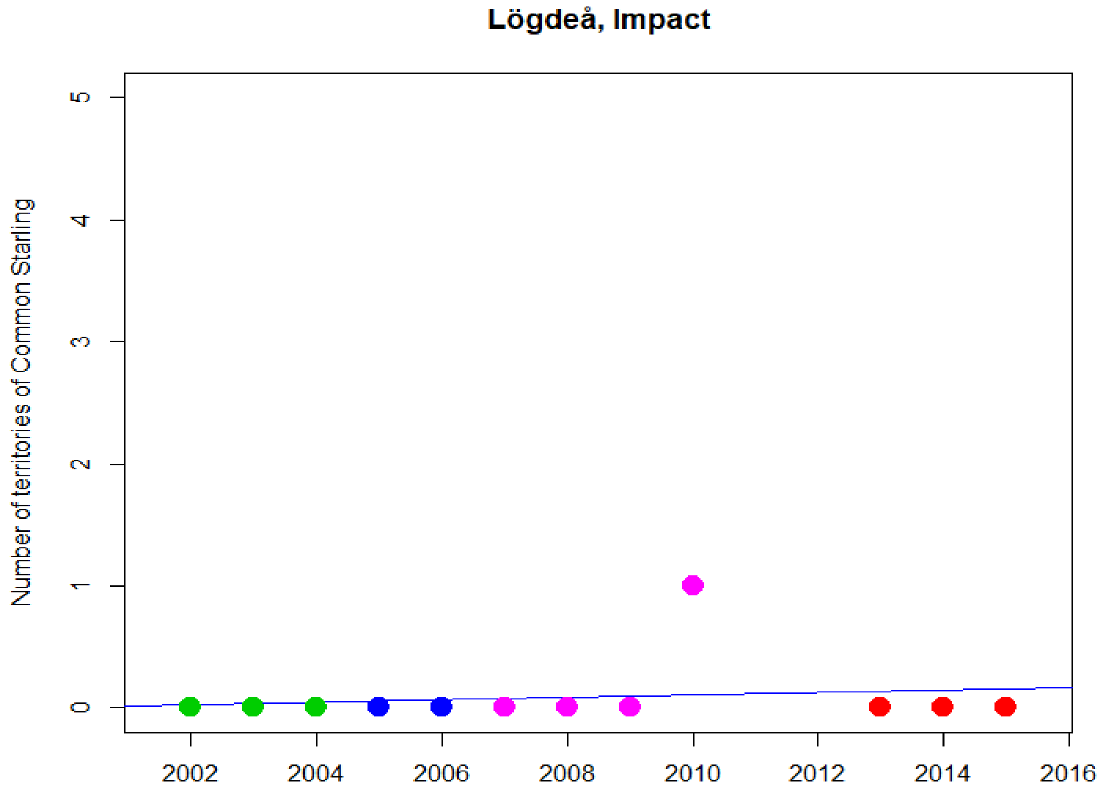
Lögdeå, Impact
Number of territories of Common Starling
0
1
2
3
4
5
2002
2004
2006
2008
2010
2012
2014
2016

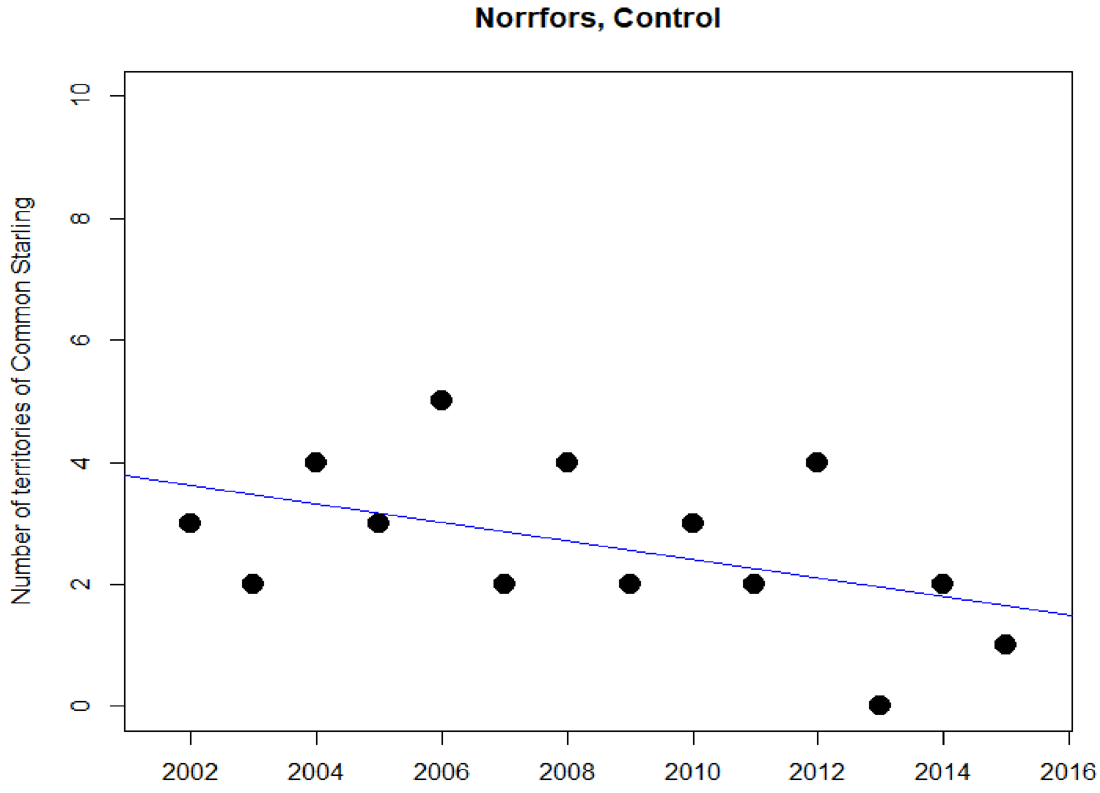
Norrfors, Control
Number of territories of Common Starling
0
2
4
6
8
10
2002
2004
2006
2008
2010
2012
2014
2016

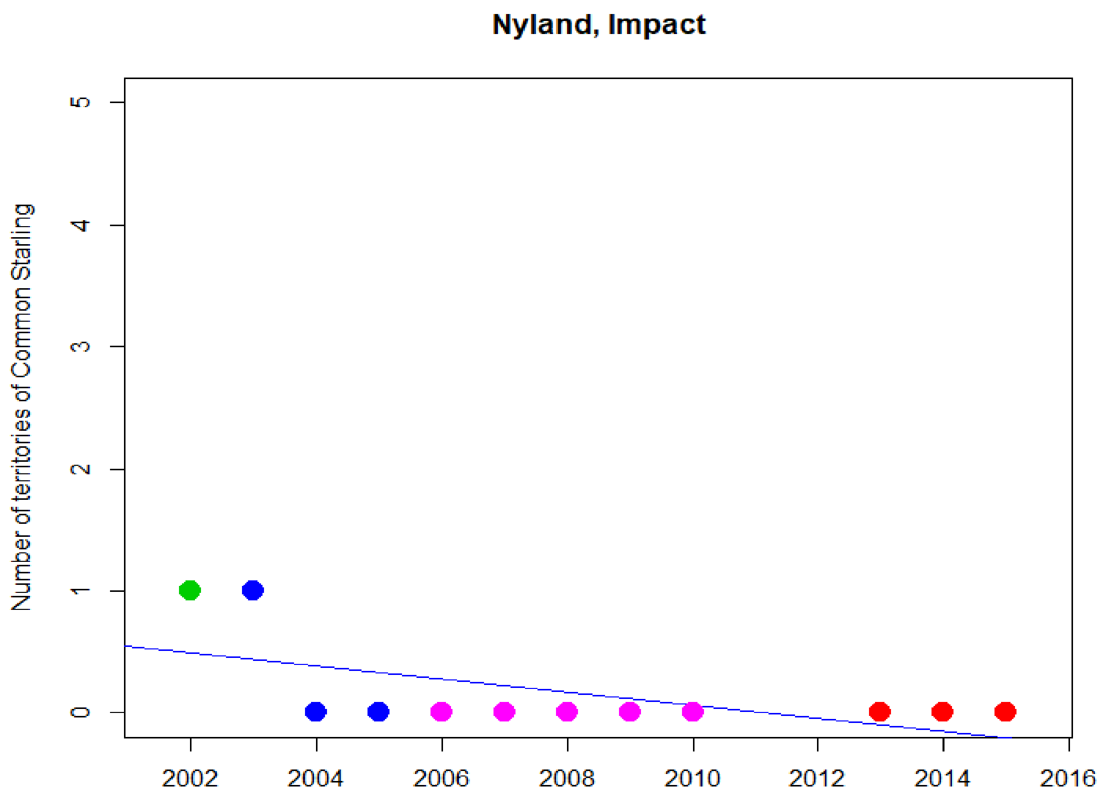
Nyland, Impact
Number of territories of Common Starling
0
1
2
3
4
5
2002
2004
2006
2008
2010
2012
2014
2016

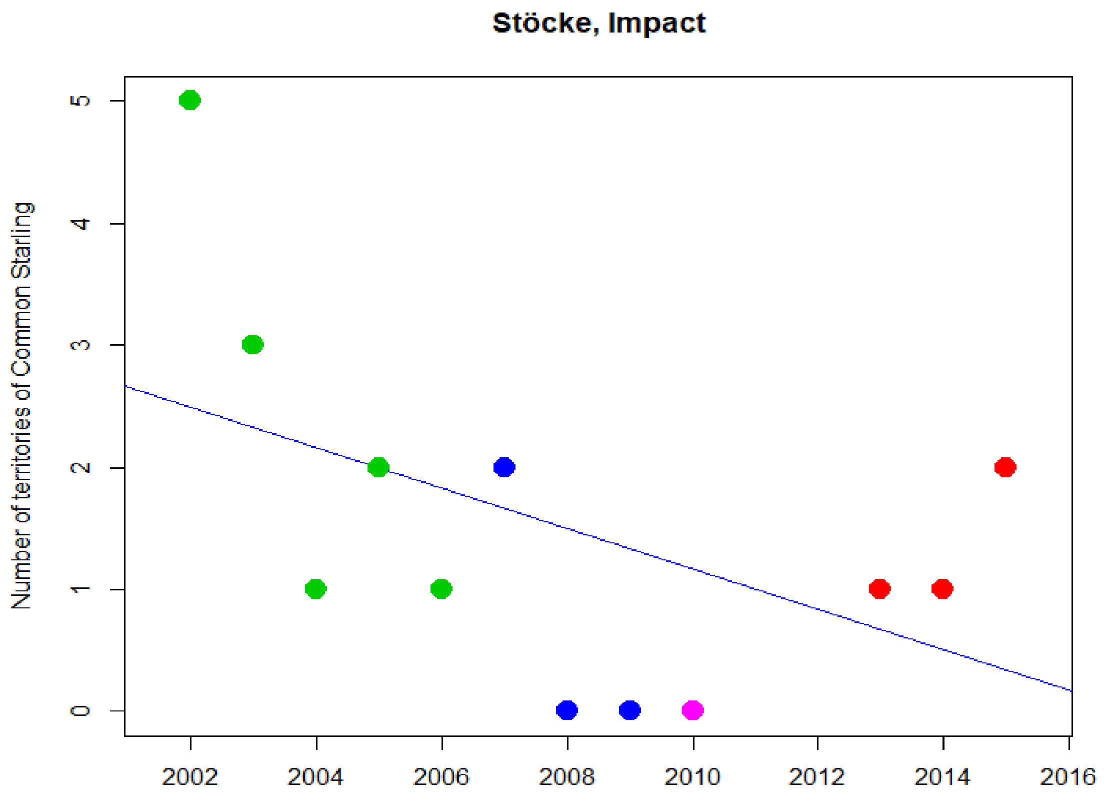
Stöcke, Impact
Number of territories of Common Starling
0
1
2
3
4
5
2002
2004
2006
2008
2010
2012
2014
2016

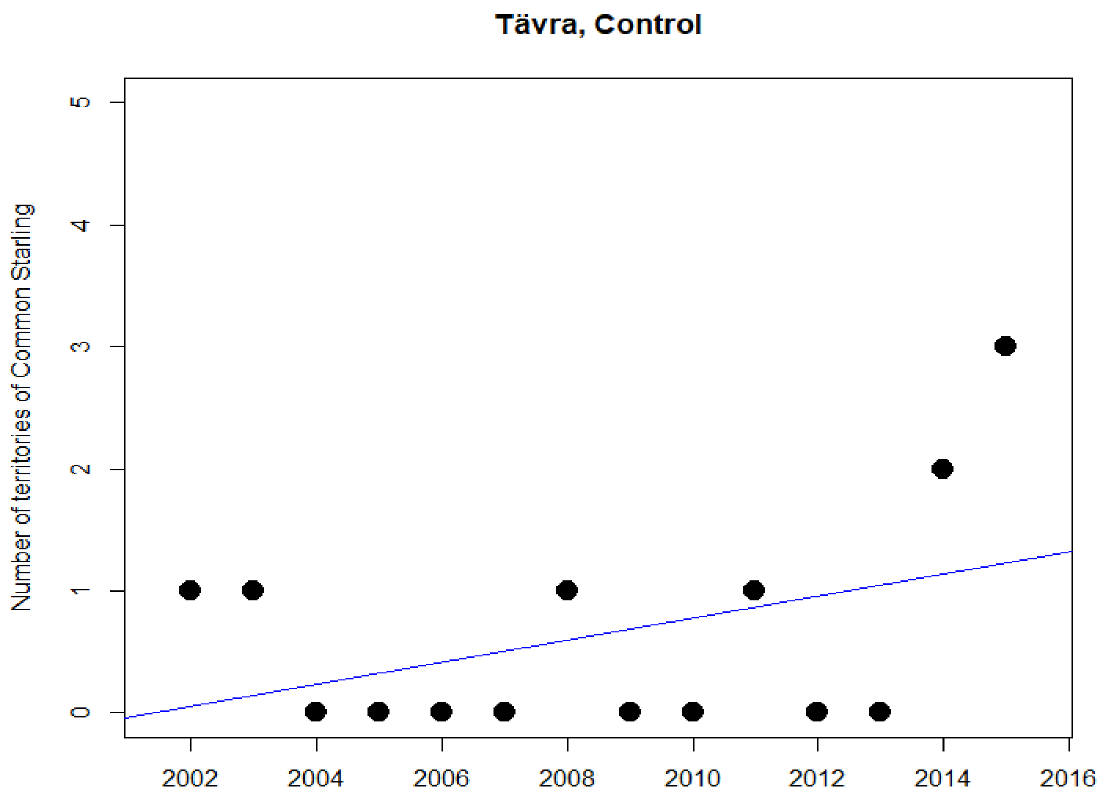

Tävra, Control
Number of territories of Common Starling
0
1
2
3
4
5
2002
2004
2006
2008
2010
2012
2014
2016


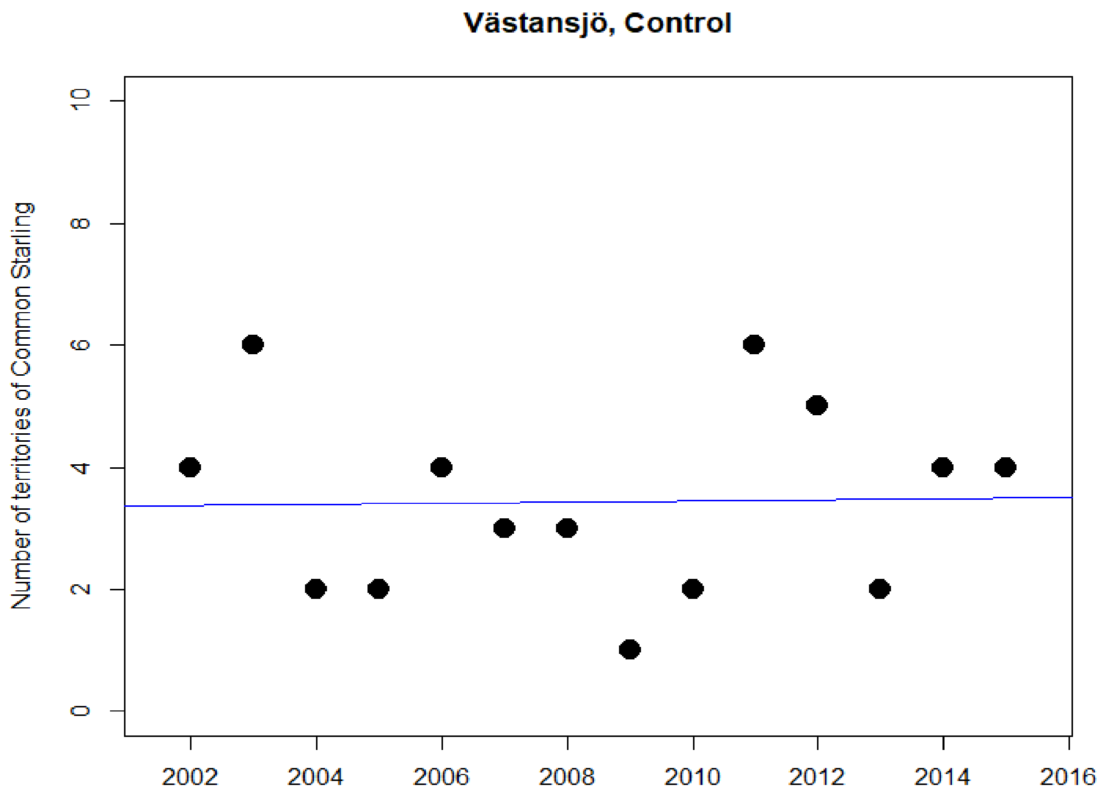

Västansjö, Control
Number of territories of Common Starling
0
2
4
6
8
10
2002
2004
2006
2008
2010
2012
2014
2016

### Eurasian Curlew

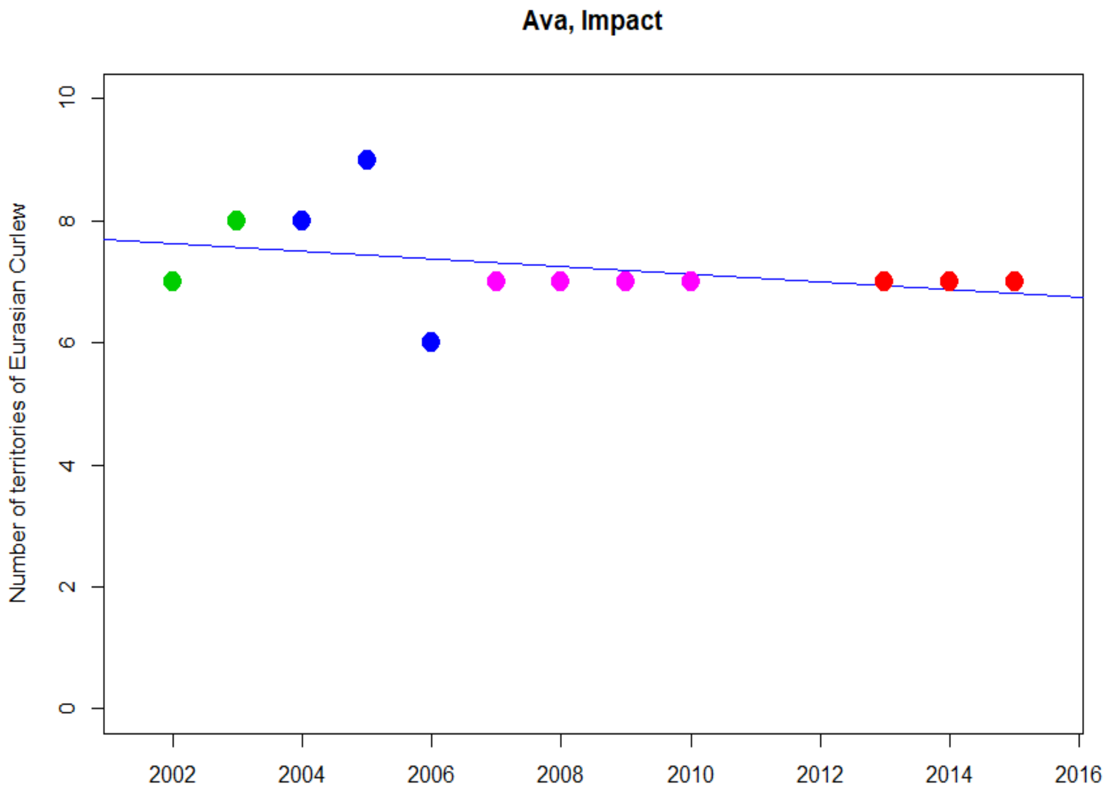


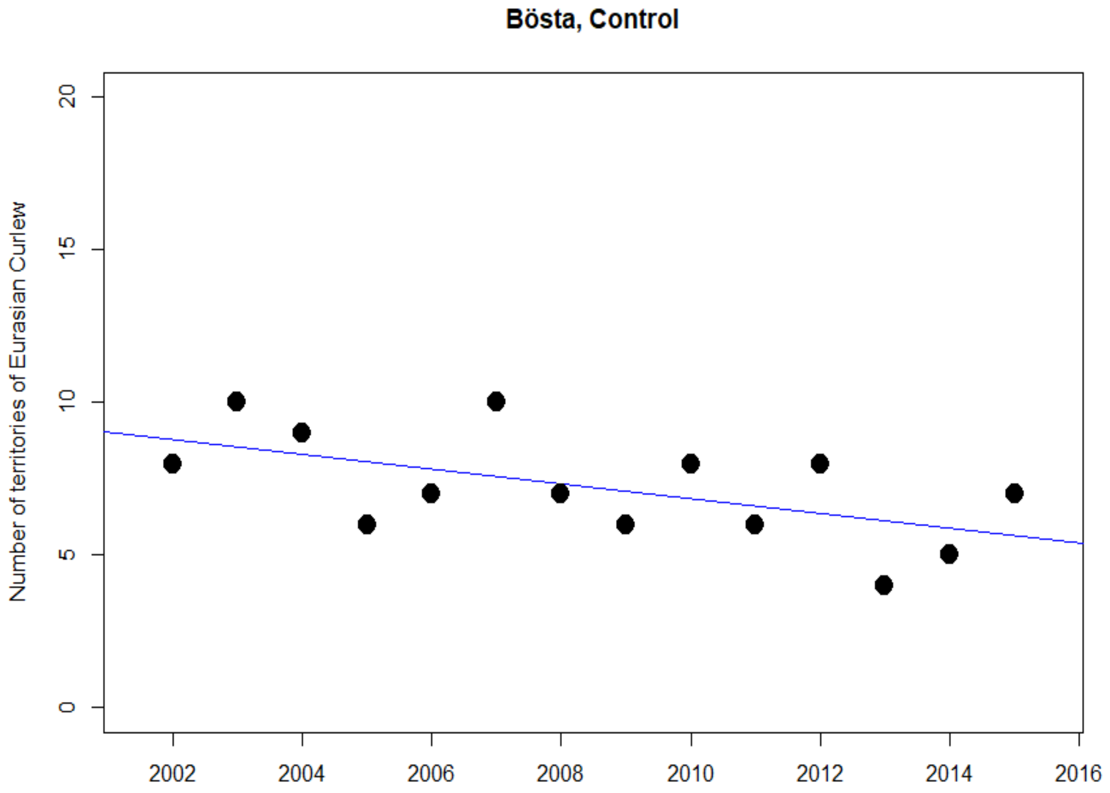

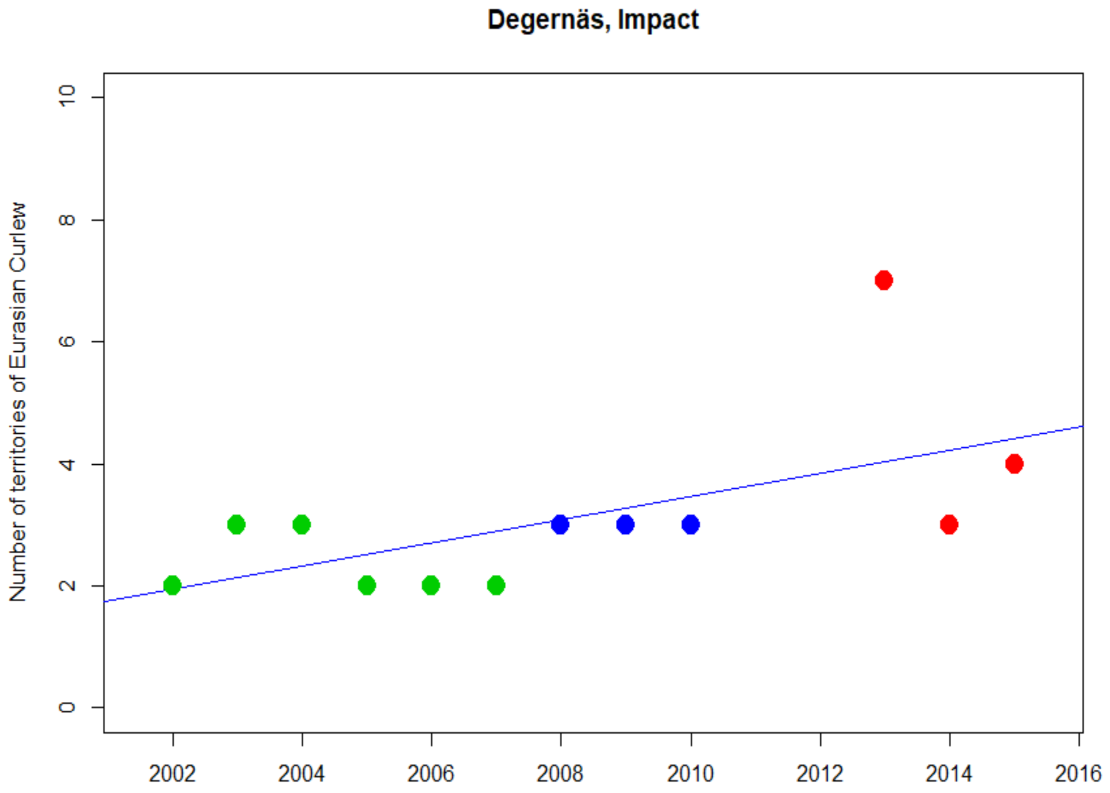
Degernäs, Impact
Number of territories of Eurasian Curlew
0
2
4
6
8
10
2002
2004
2006
2008
2010
2012
2014
2016

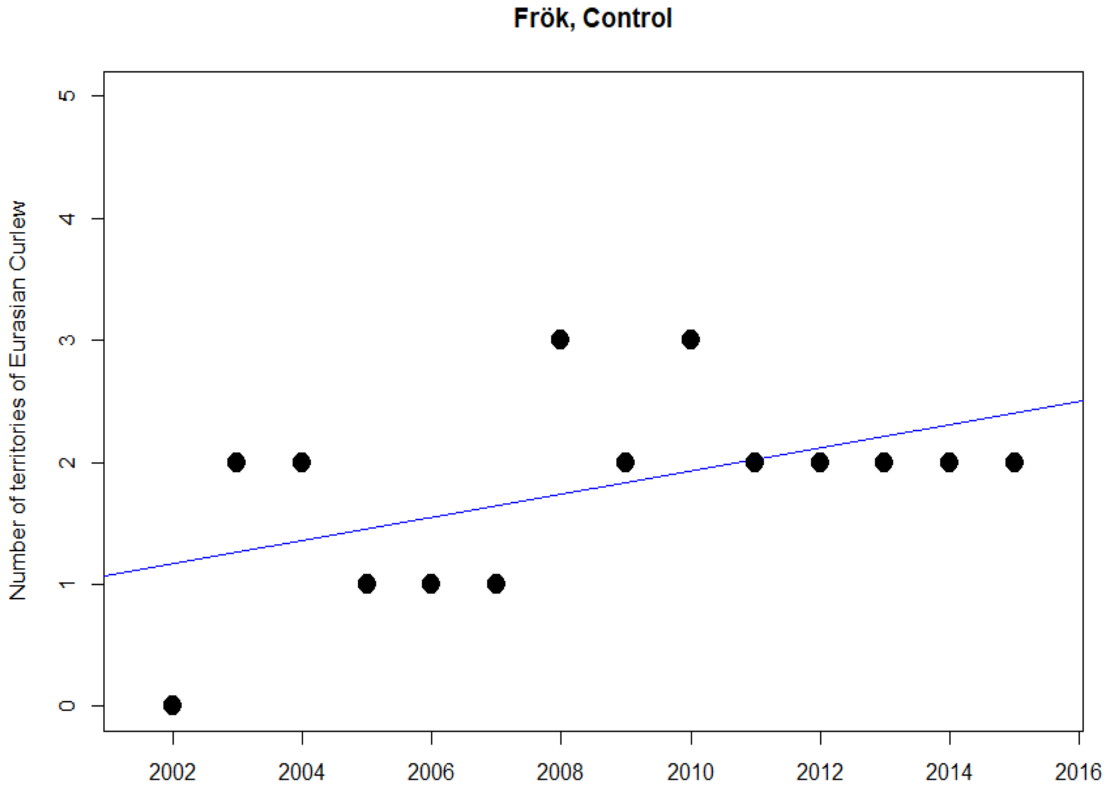
Frök, Control
Number of territories of Eurasian Curlew
0
1
2
3
4
5
2002
2004
2006
2008
2010
2012
2014
2016

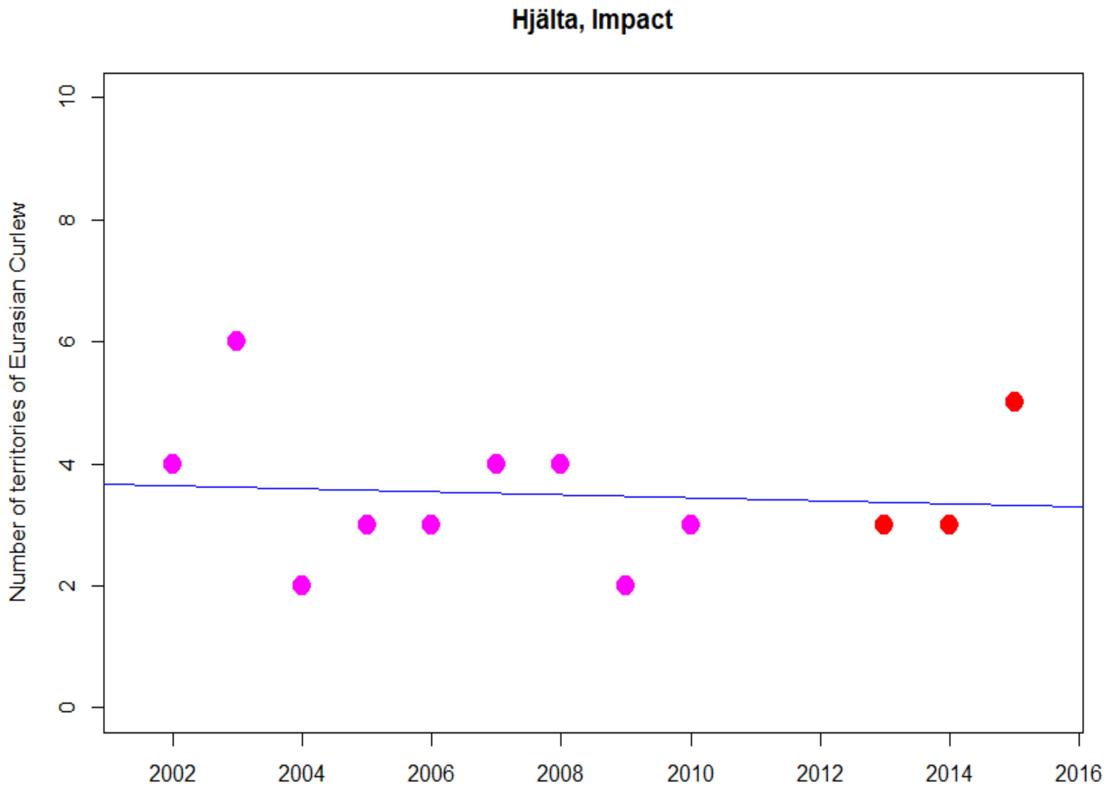
Hjälta, Impact
Number of territories of Eurasian Curlew
0
2
4
6
8
10
2002
2004
2006
2008
2010
2012
2014
2016

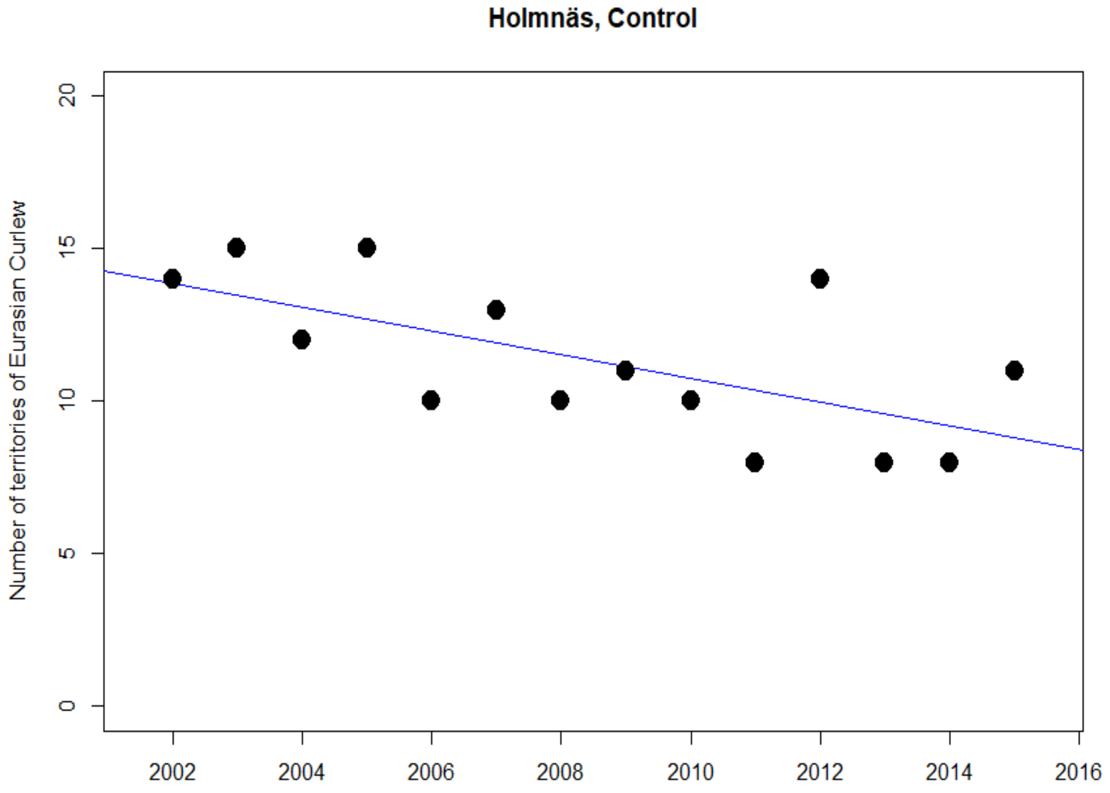
Holmnäs, Control
Number of territories of Eurasian Curlew
0
5
10
15
20
2002
2004
2006
2008
2010
2012
2014
2016

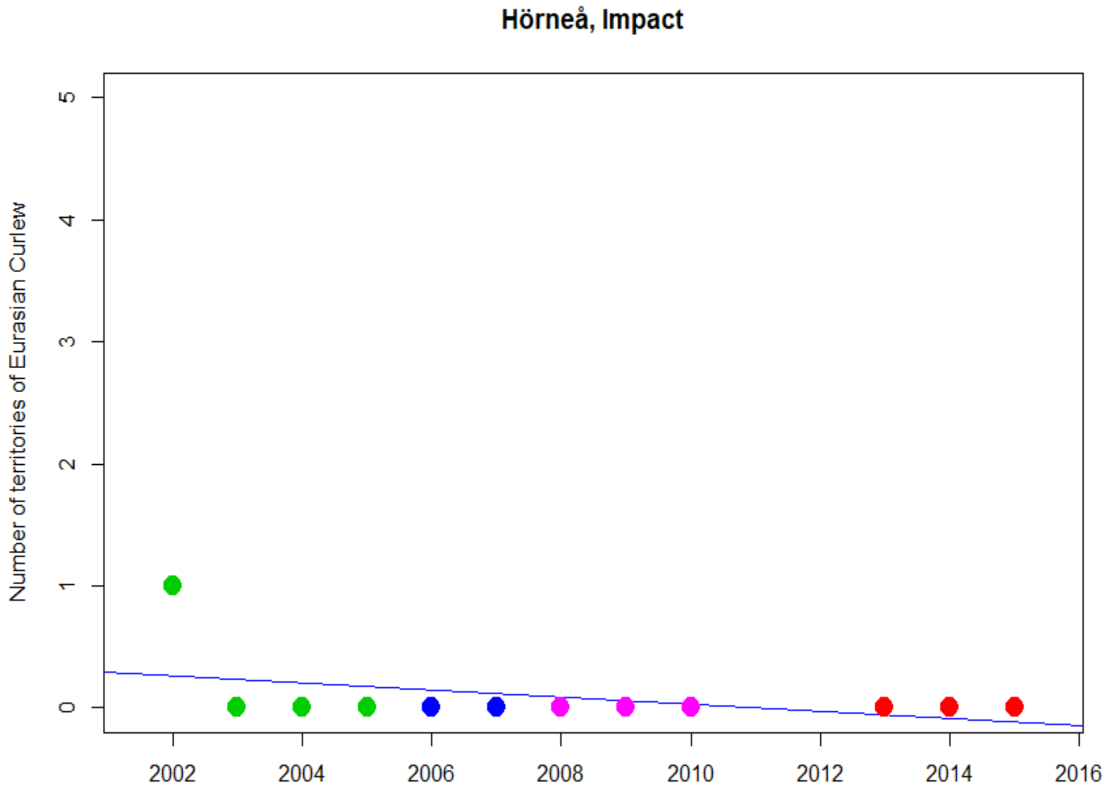

Hörneå, Impact
Number of territories of Eurasian Curlew
0
1
2
3
4
5
2002
2004
2006
2008
2010
2012
2014
2016


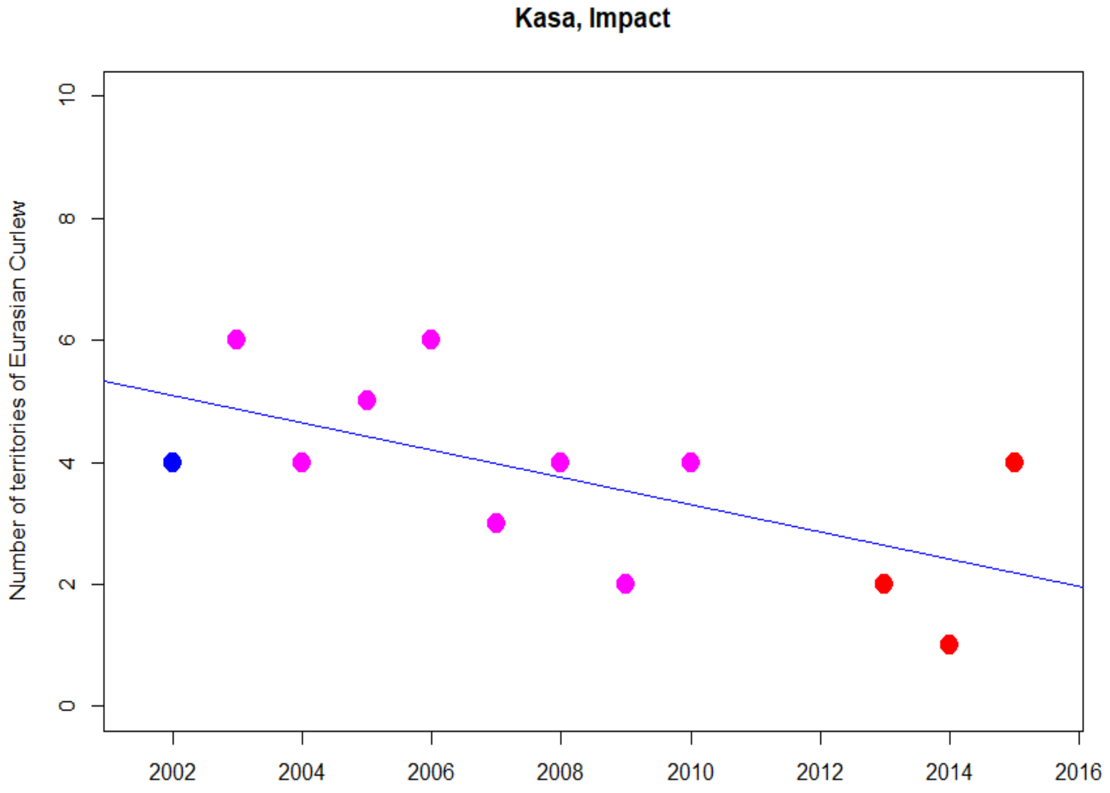

Kasa, Impact
Number of territories of Eurasian Curlew
0
2
4
6
8
10
2002
2004
2006
2008
2010
2012
2014
2016

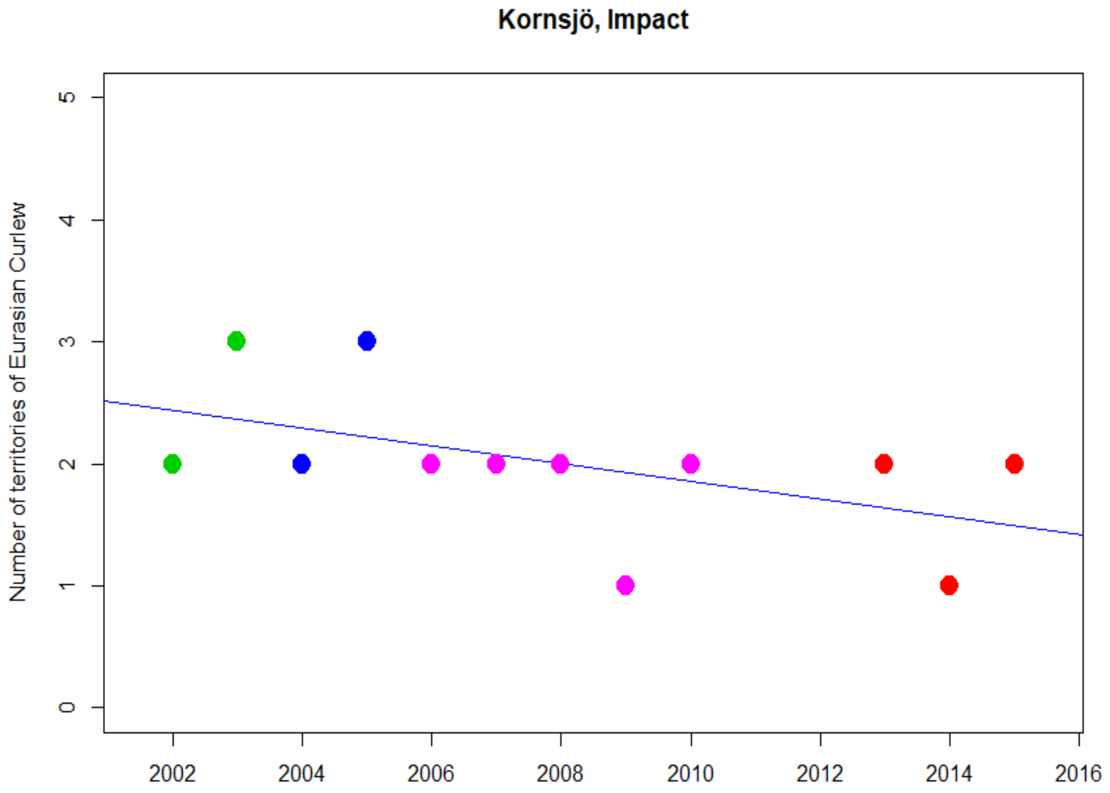
Kornsjö, Impact
Number of territories of Eurasian Curlew
0
1
2
3
4
5
2002
2004
2006
2008
2010
2012
2014
2016

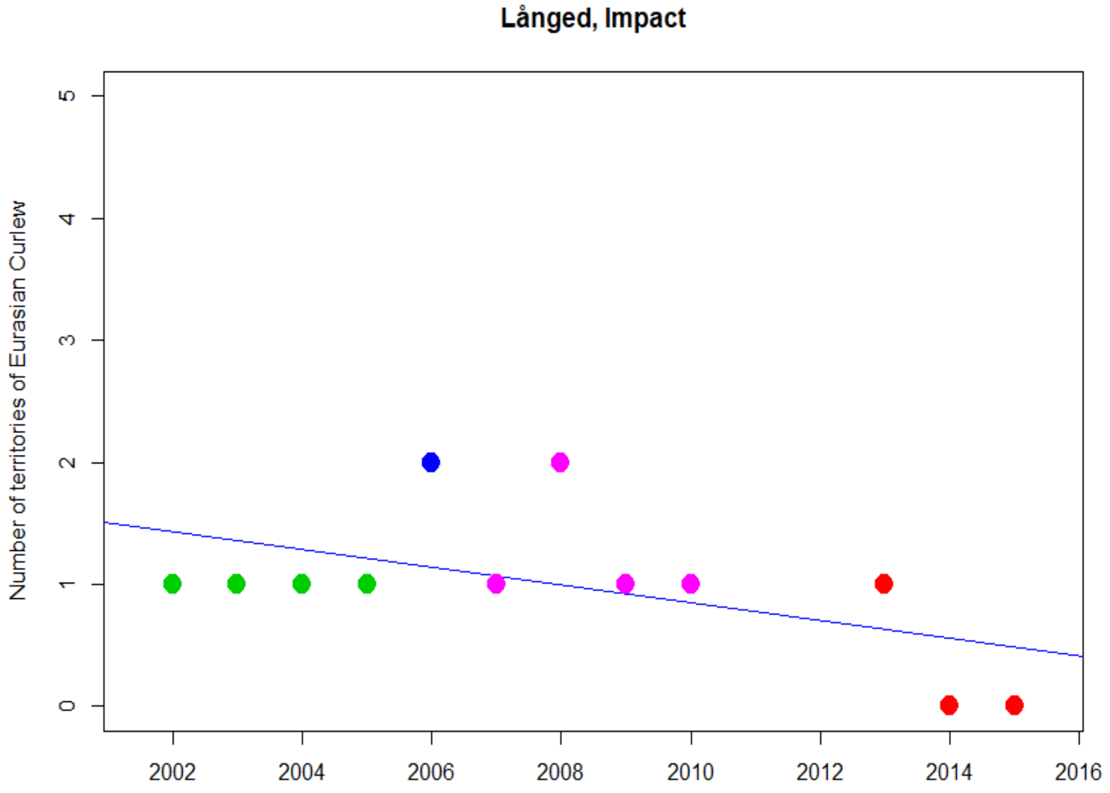
Långed, Impact
Number of territories of Eurasian Curlew
0
1
2
3
4
5
2002
2004
2006
2008
2010
2012
2014
2016

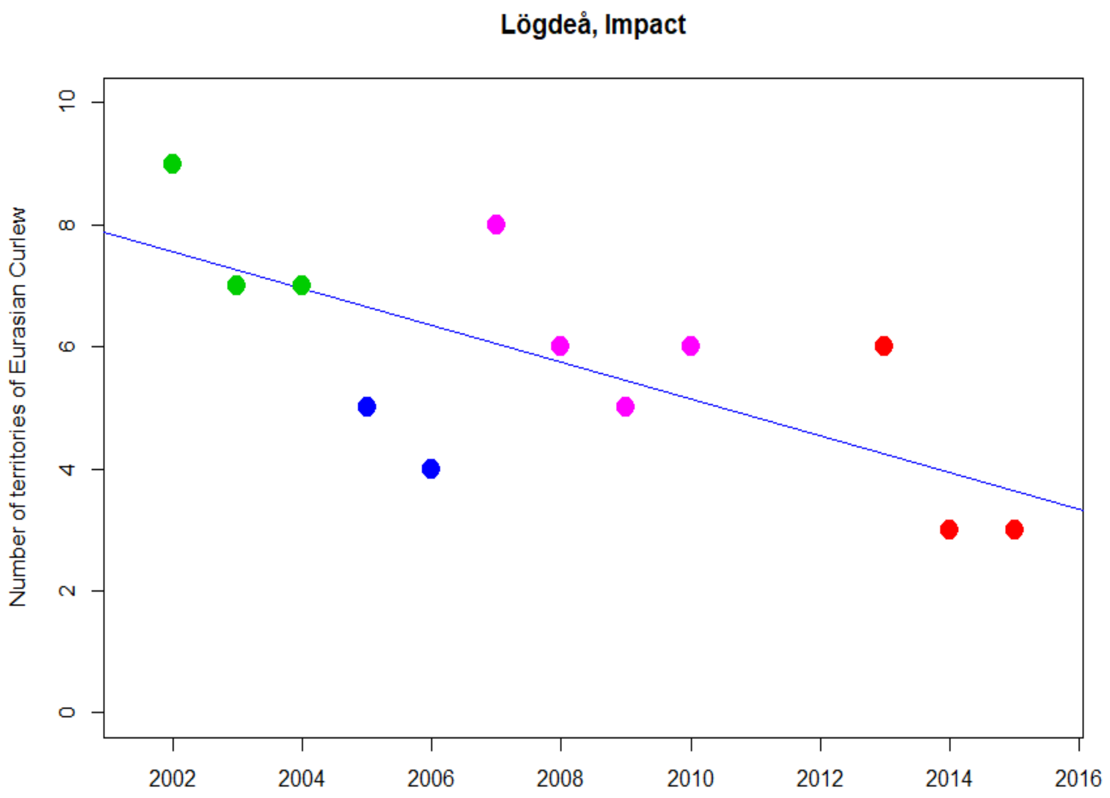
Lögdeå, Impact
Number of territories of Eurasian Curlew
0
2
4
6
8
10
2002
2004
2006
2008
2010
2012
2014
2016

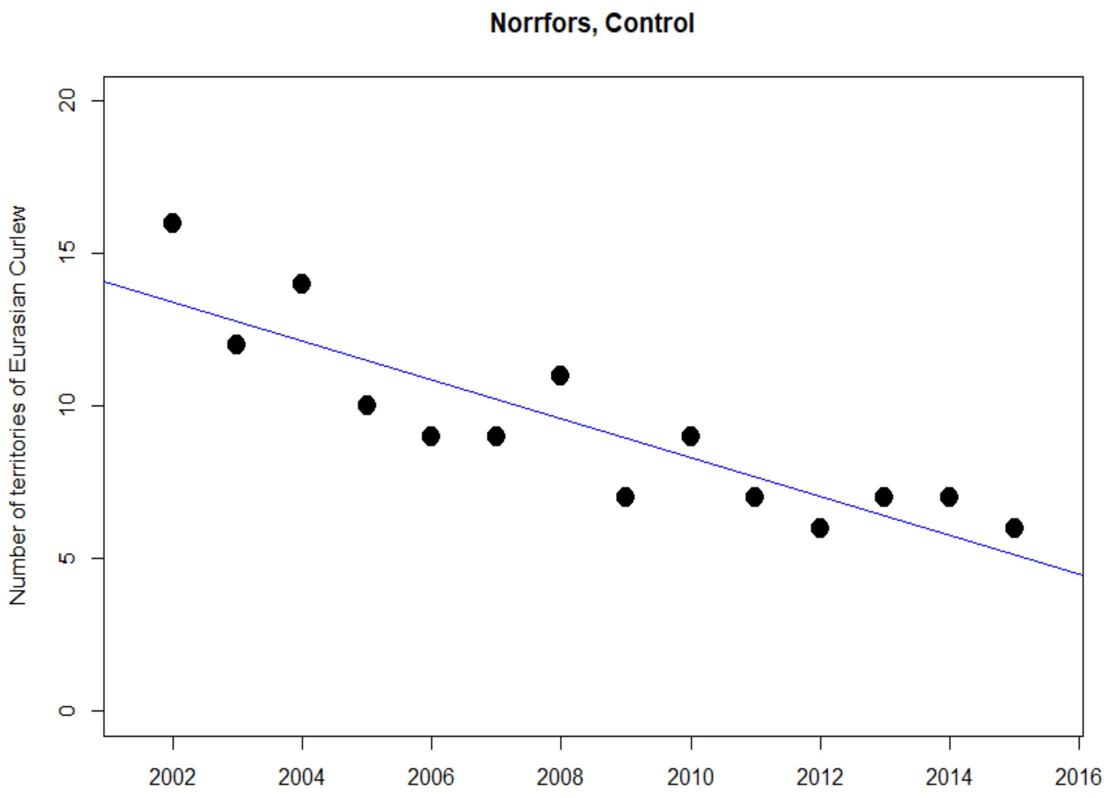
Norrfors, Control
Number of territories of Eurasian Curlew
0
5
10
15
20
2002
2004
2006
2008
2010
2012
2014
2016

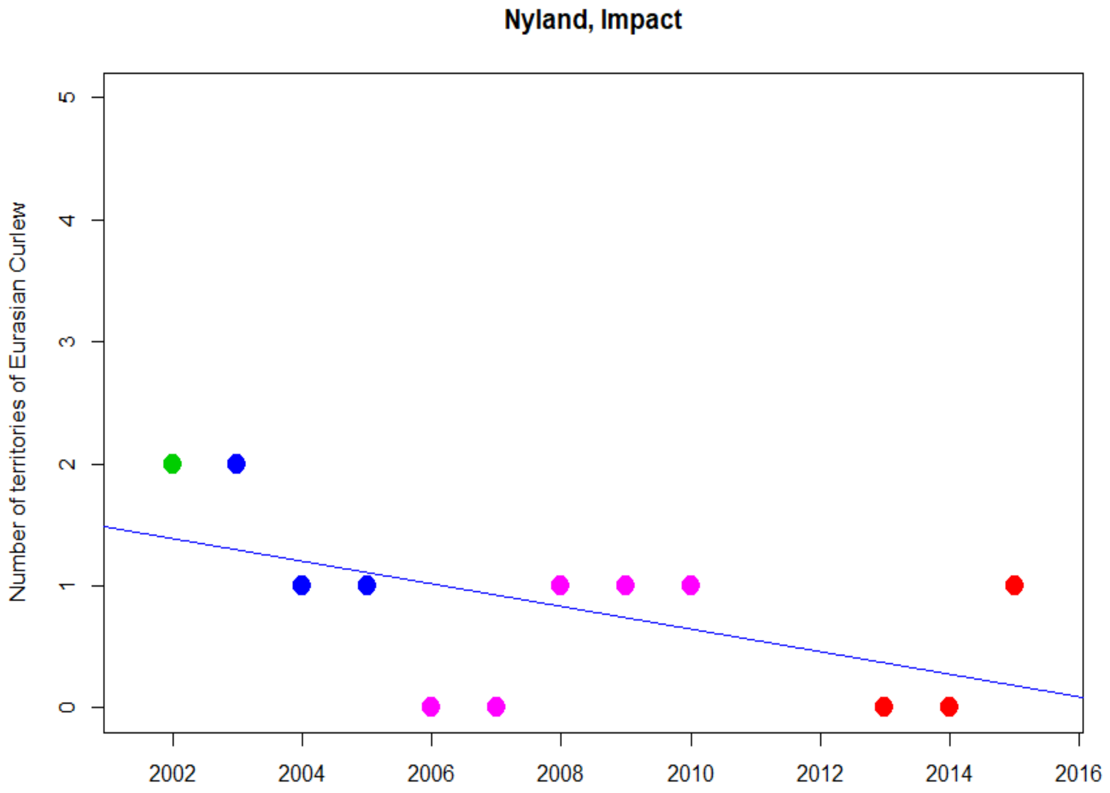
Nyland, Impact
Number of territories of Eurasian Curlew
0
1
2
3
4
5
2002
2004
2006
2008
2010
2012
2014
2016

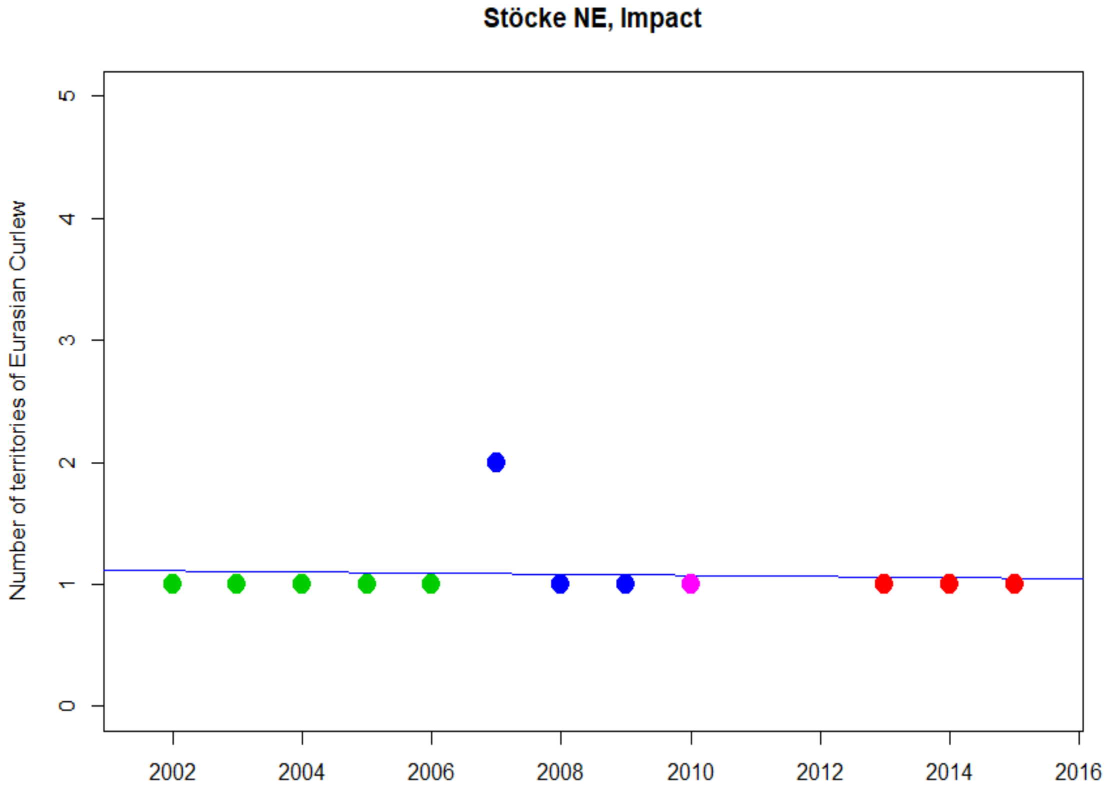
Stöcke NE, Impact
Number of territories of Eurasian Curlew
0
1
2
3
4
5
2002
2004
2006
2008
2010
2012
2014
2016

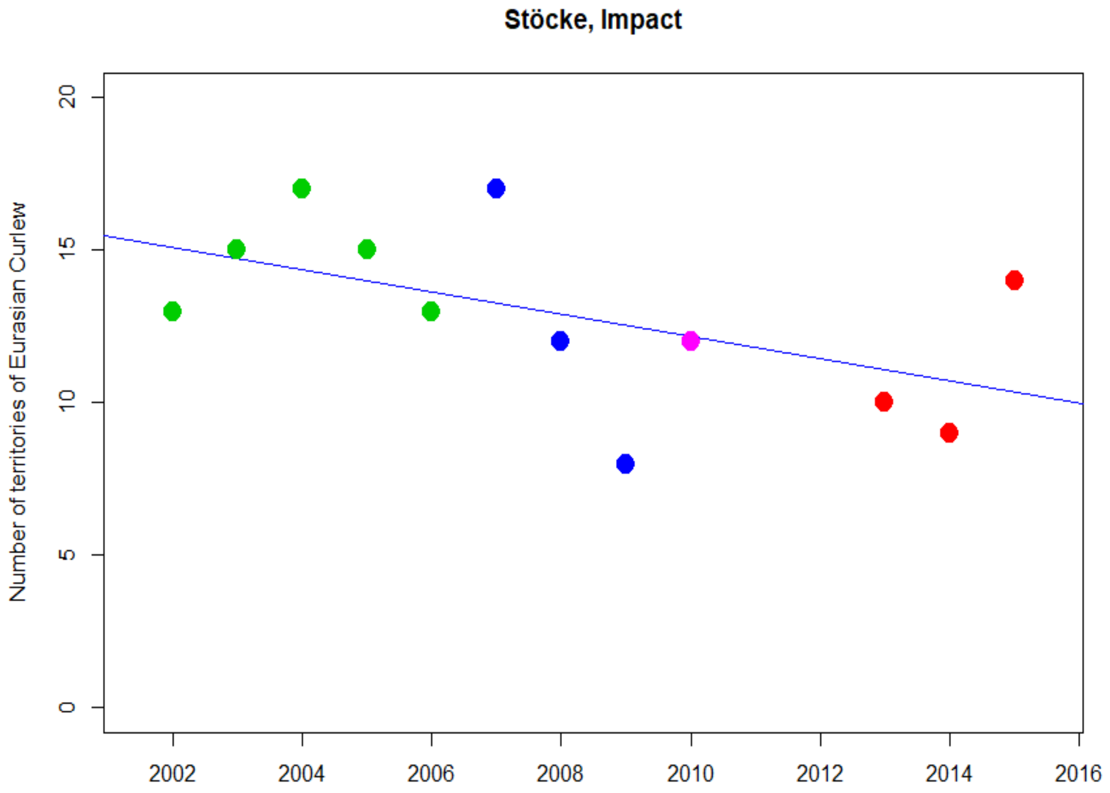
Stöcke, Impact
Number of territories of Eurasian Curlew
0
5
10
15
20
2002
2004
2006
2008
2010
2012
2014
2016

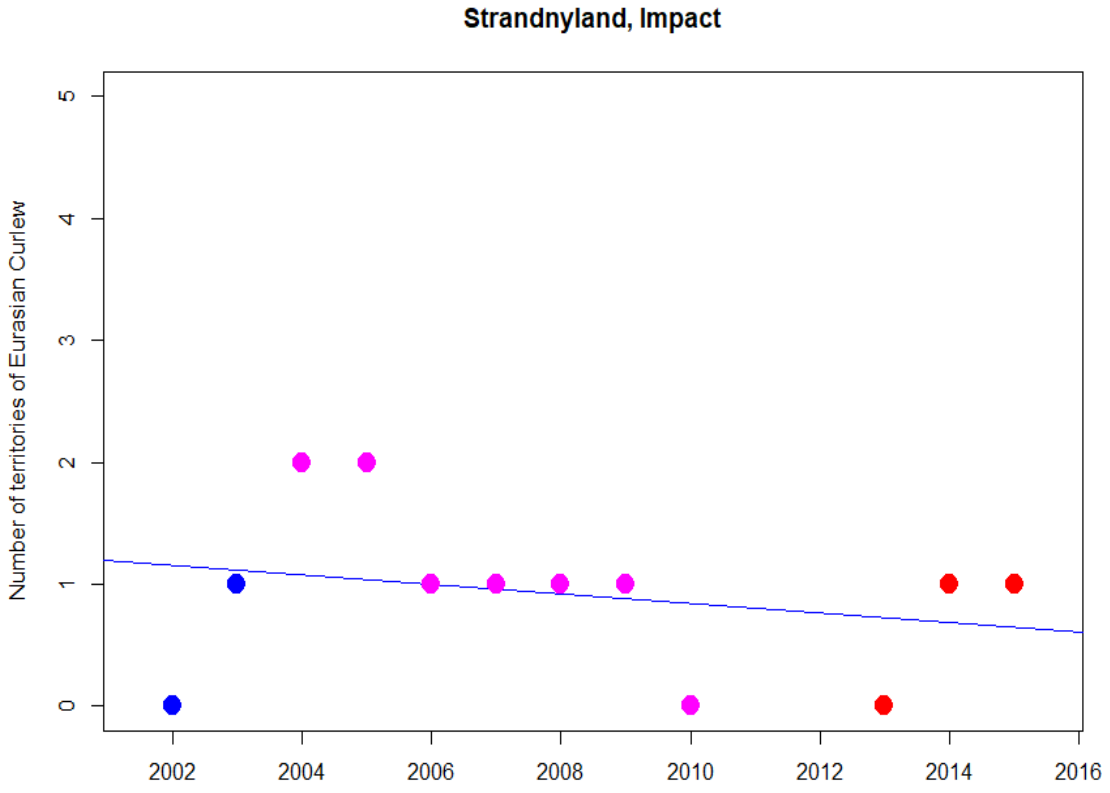
Strandnyland, Impact
Number of territories of Eurasian Curlew
0
1
2
3
4
5
2002
2004
2006
2008
2010
2012
2014
2016

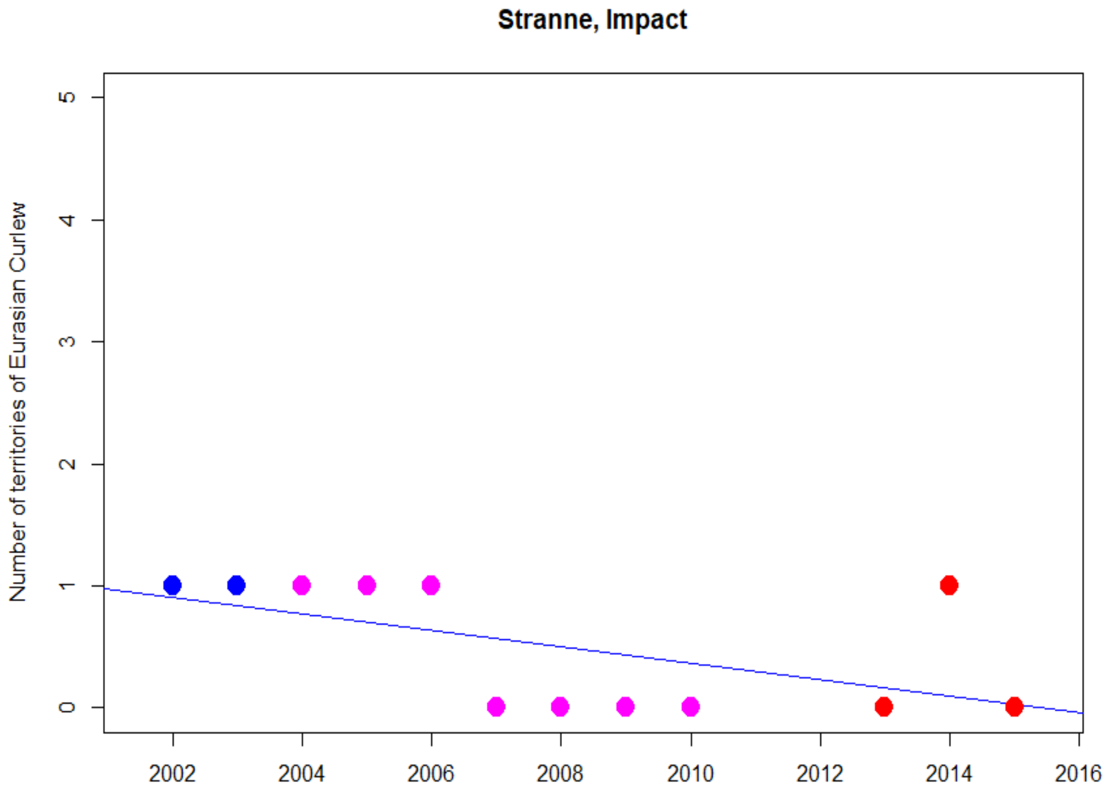
Stranne, Impact
Number of territories of Eurasian Curlew
0
1
2
3
4
5
2002
2004
2006
2008
2010
2012
2014
2016

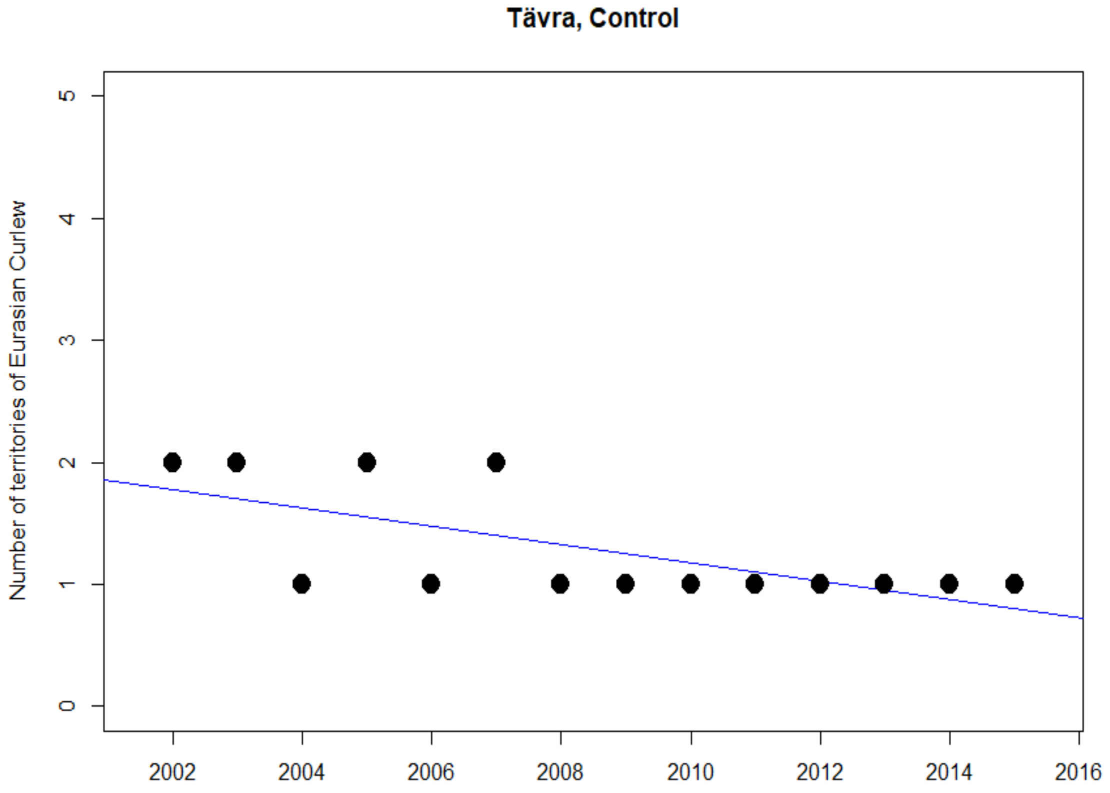
Tävra, Control
Number of territories of Eurasian Curlew
0
1
2
3
4
5
2002
2004
2006
2008
2010
2012
2014
2016

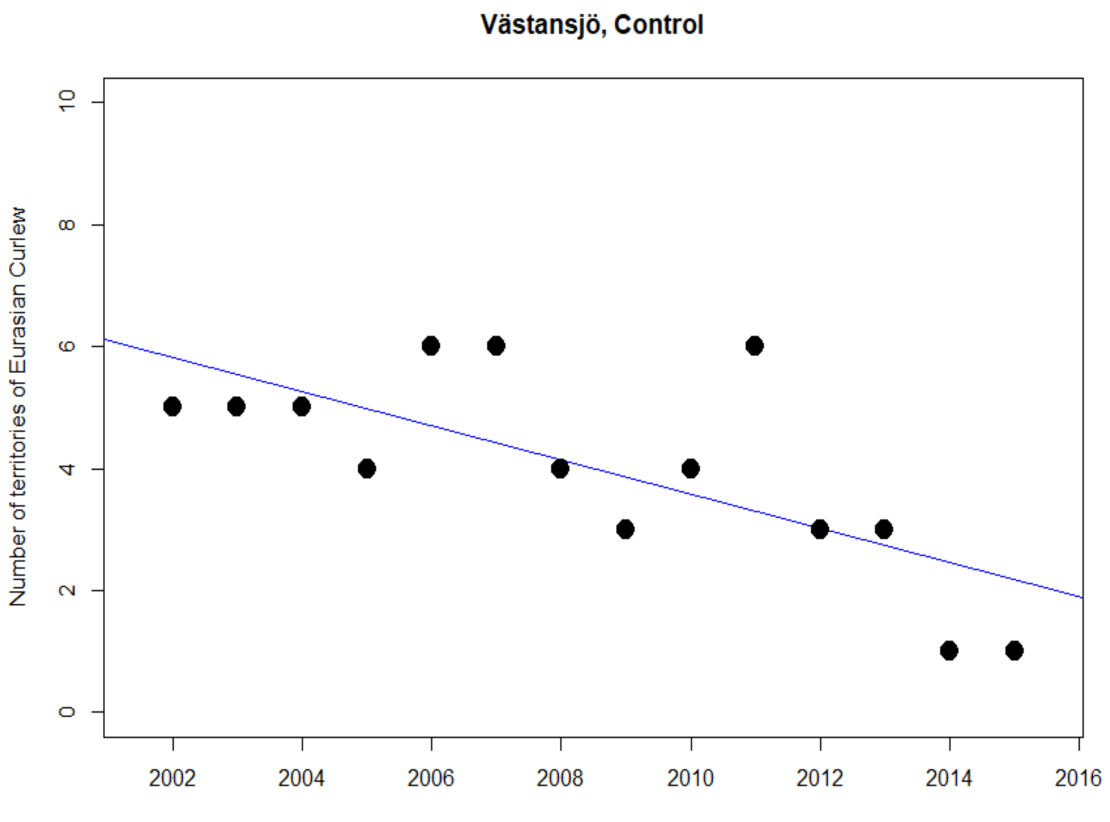
Västansjö, Control
Number of territories of Eurasian Curlew
0
2
4
6
8
10
2002
2004
2006
2008
2010
2012
2014
2016

### Eurasian Golden Plover

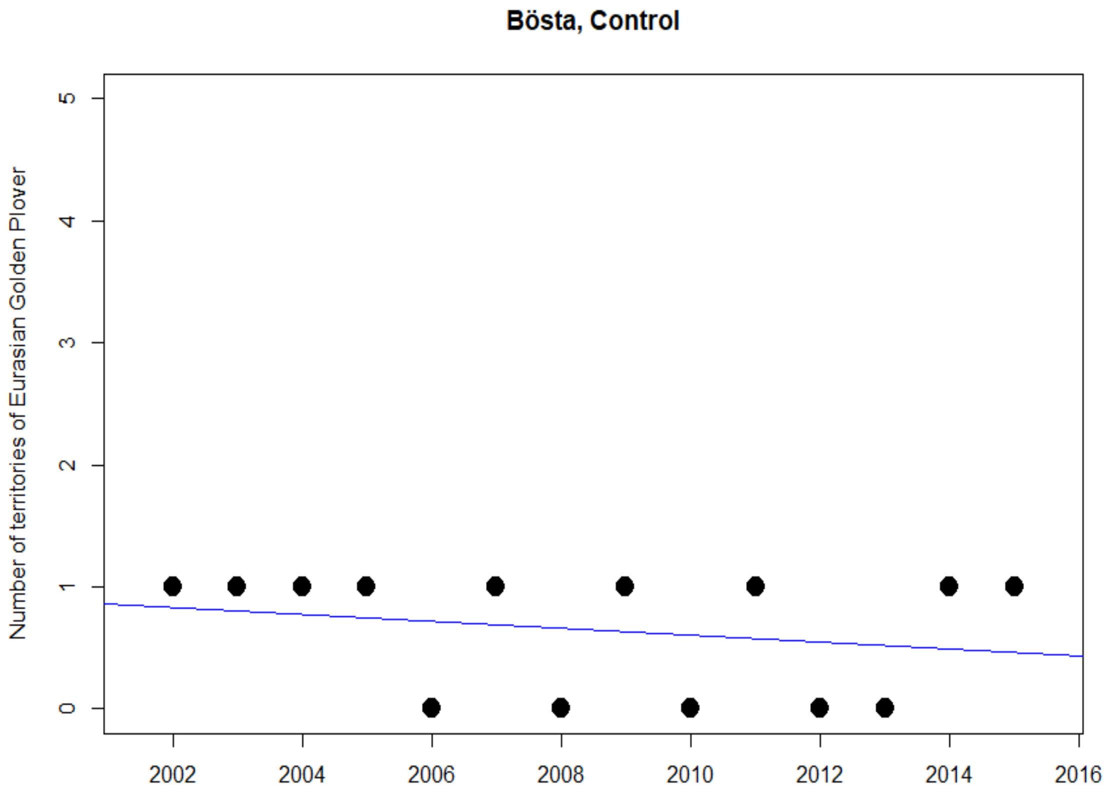

### Eurasian Skylark

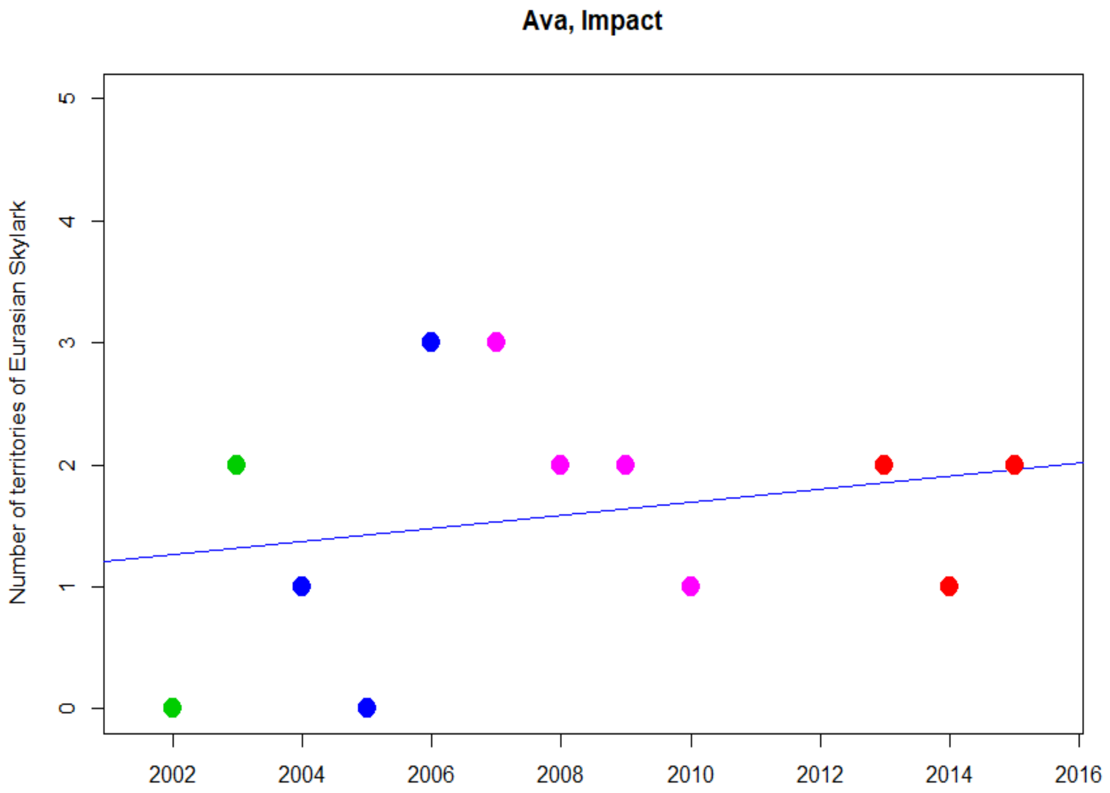


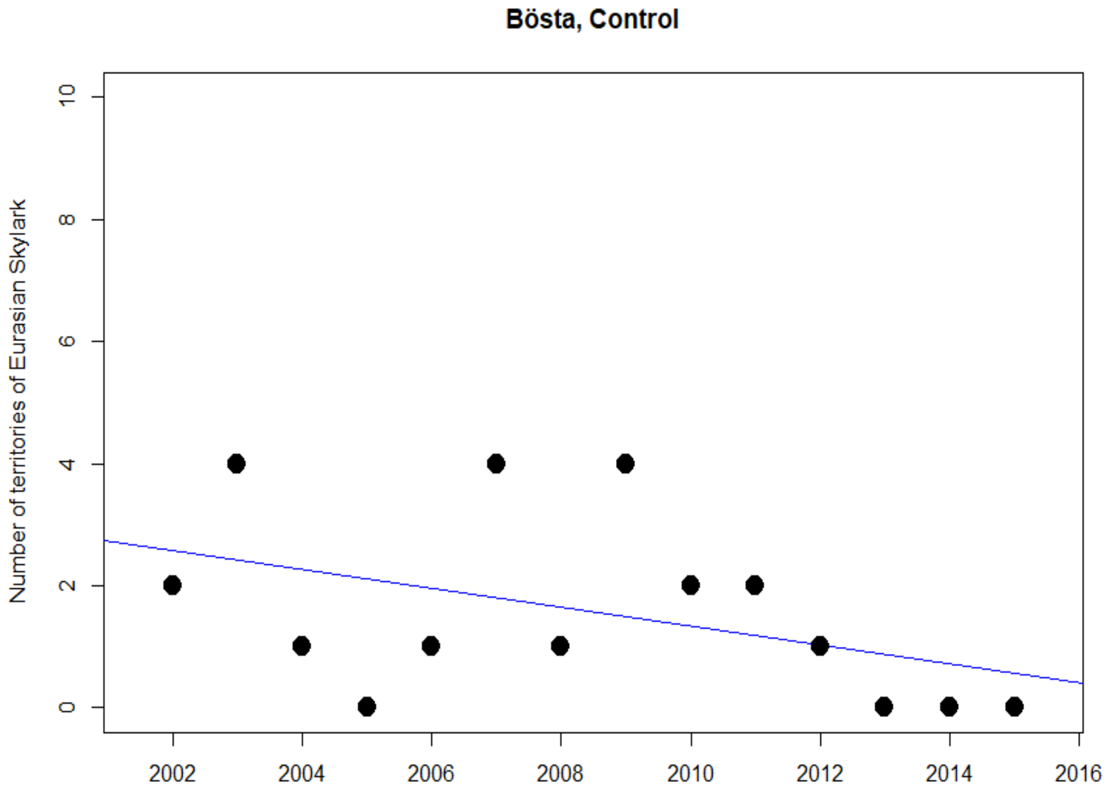

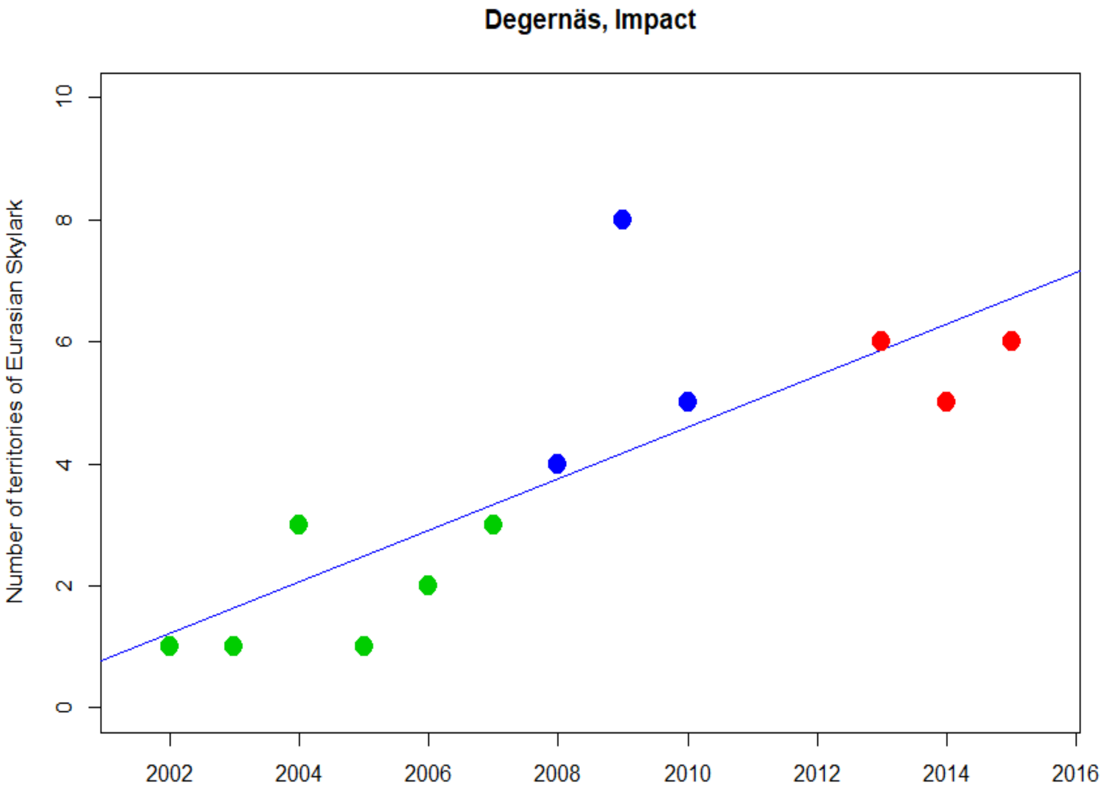
Degernäs, Impact
Number of territories of Eurasian Skylark
0
2
4
6
8
10
2002
2004
2006
2008
2010
2012
2014
2016

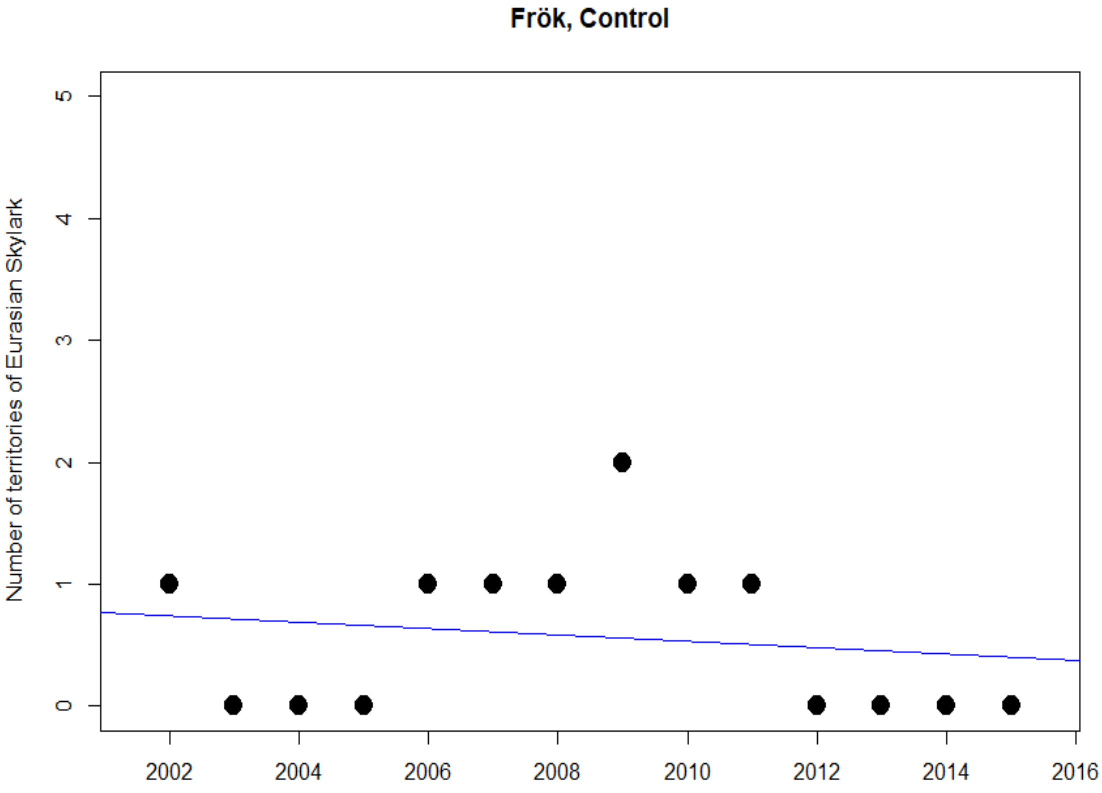
Frök, Control
Number of territories of Eurasian Skylark
0
1
2
3
4
5
2002
2004
2006
2008
2010
2012
2014
2016

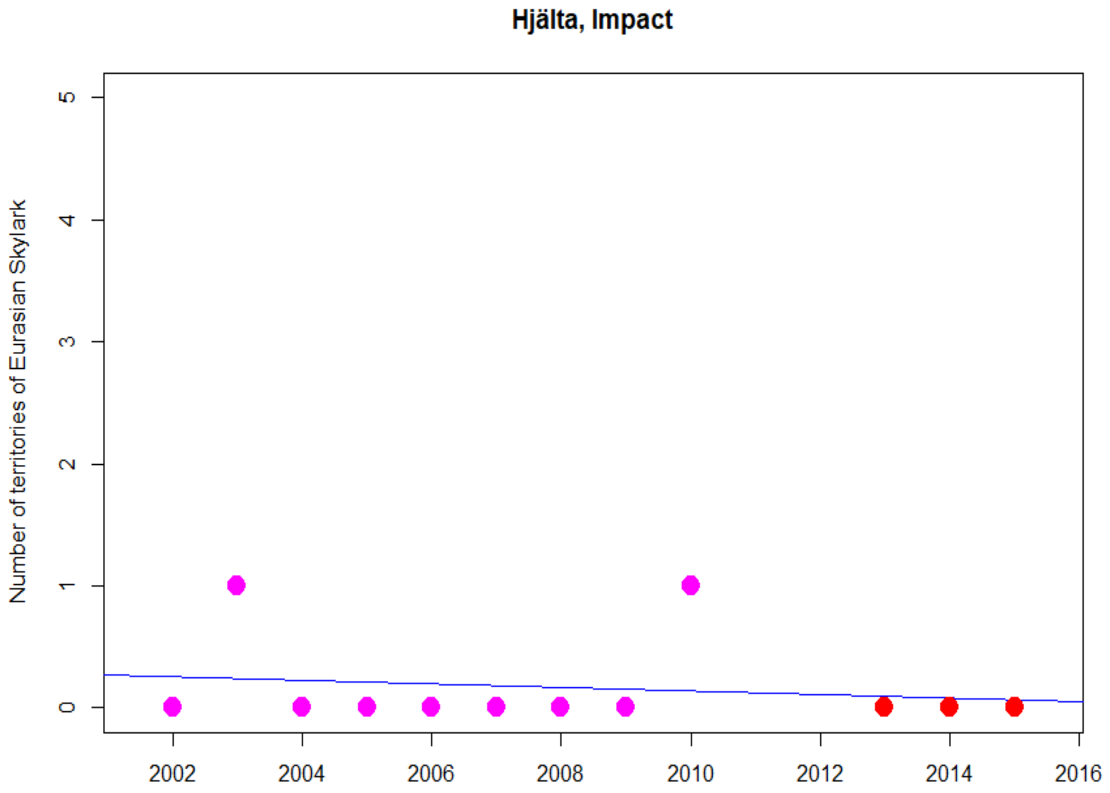
Hjälta, Impact
Number of territories of Eurasian Skylark
0
1
2
3
4
5
2002
2004
2006
2008
2010
2012
2014
2016

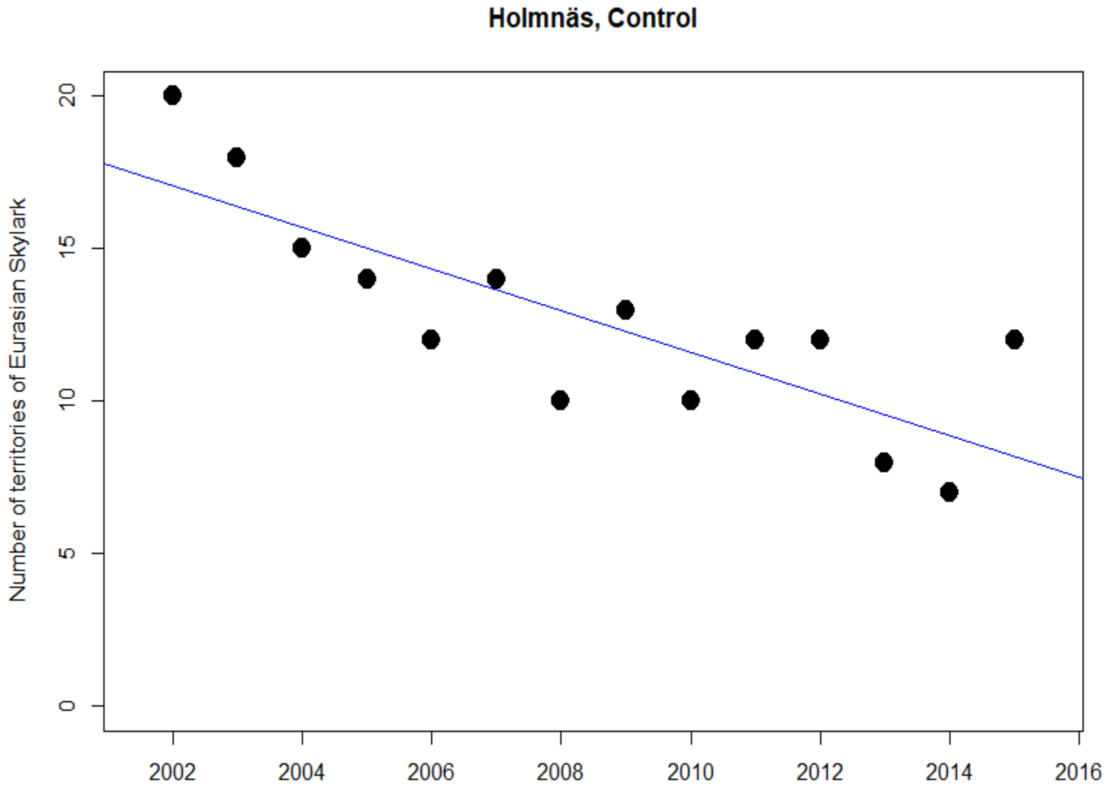
Holmnäs, Control
Number of territories of Eurasian Skylark
0
5
10
15
20
2002
2004
2006
2008
2010
2012
2014
2016

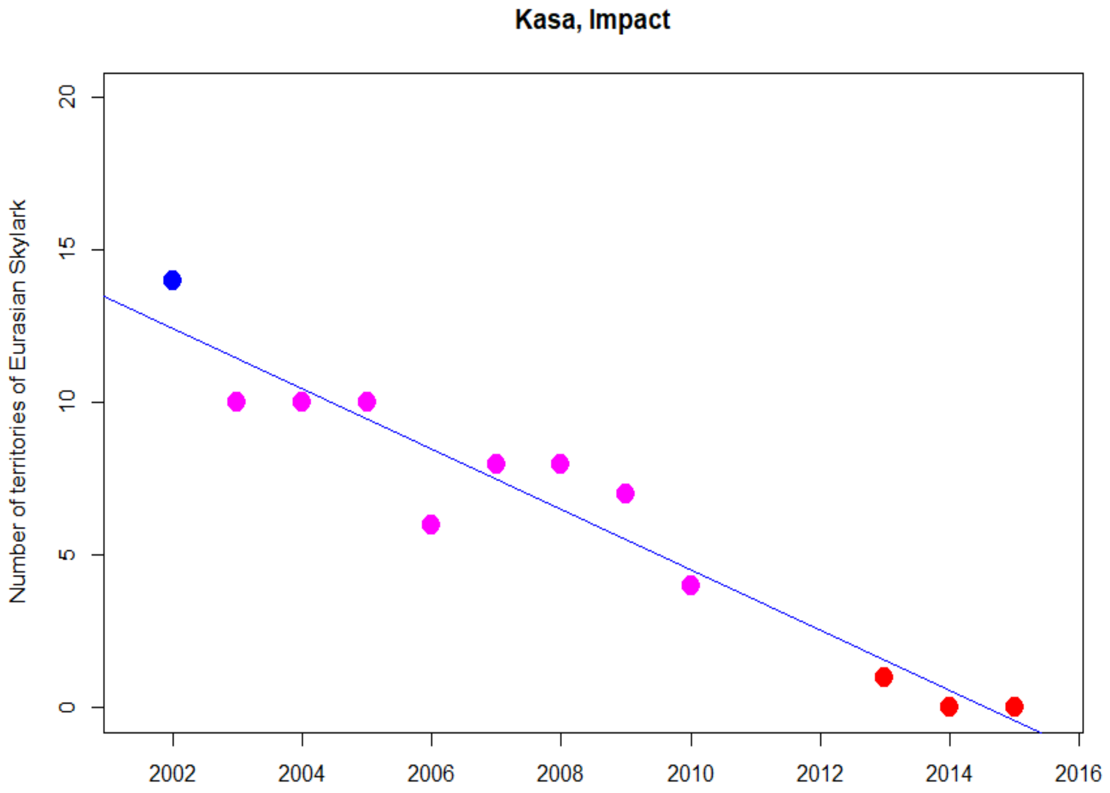
Kasa, Impact
Number of territories of Eurasian Skylark
0
5
10
15
20
2002
2004
2006
2008
2010
2012
2014
2016

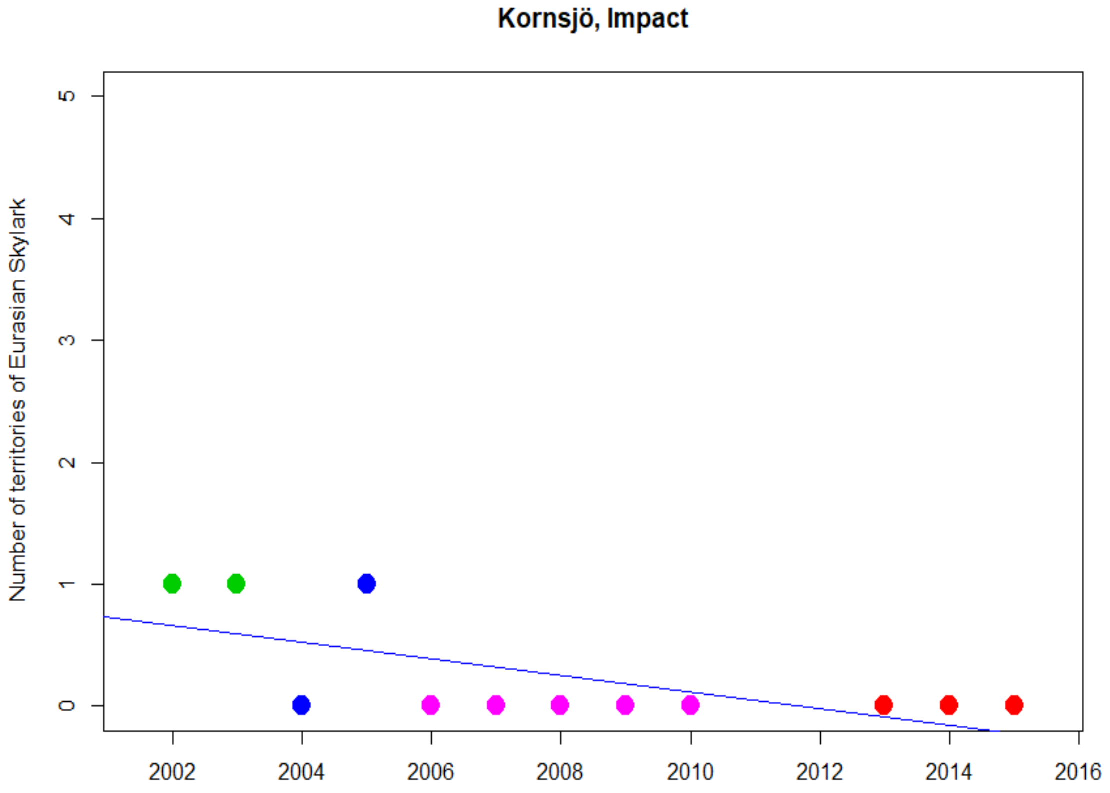
Kornsjö, Impact
Number of territories of Eurasian Skylark
0
1
2
3
4
5
2002
2004
2006
2008
2010
2012
2014
2016

**Långed, Impact**

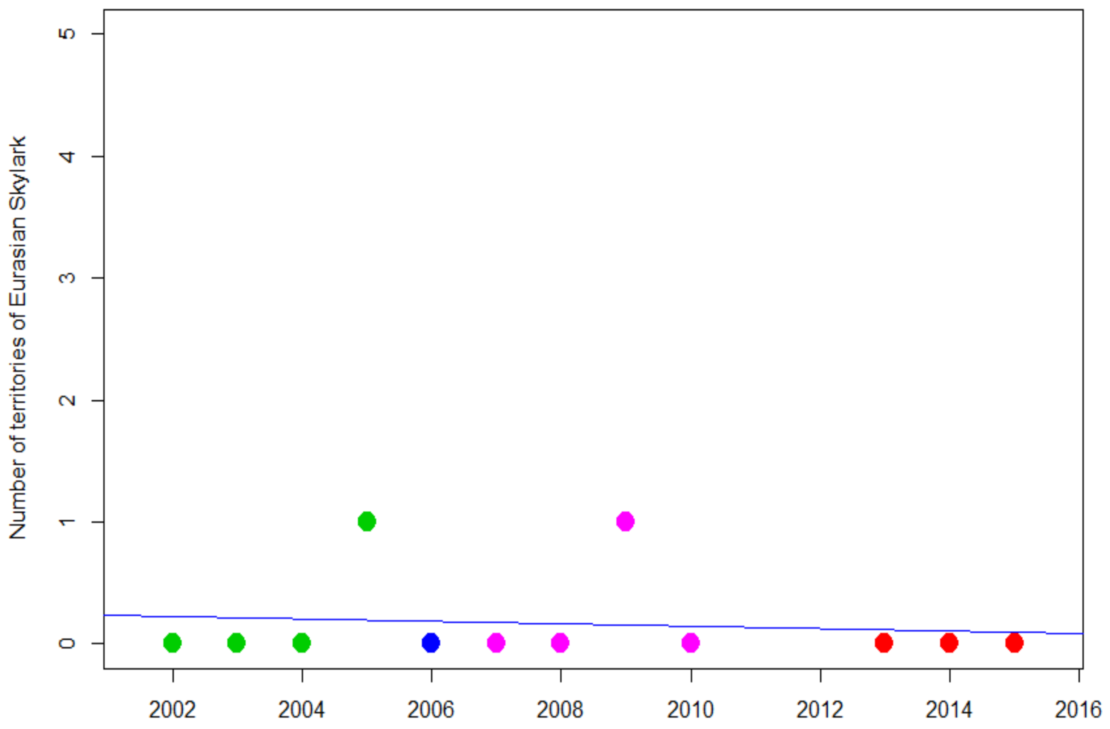


**Lögdeå, Impact**

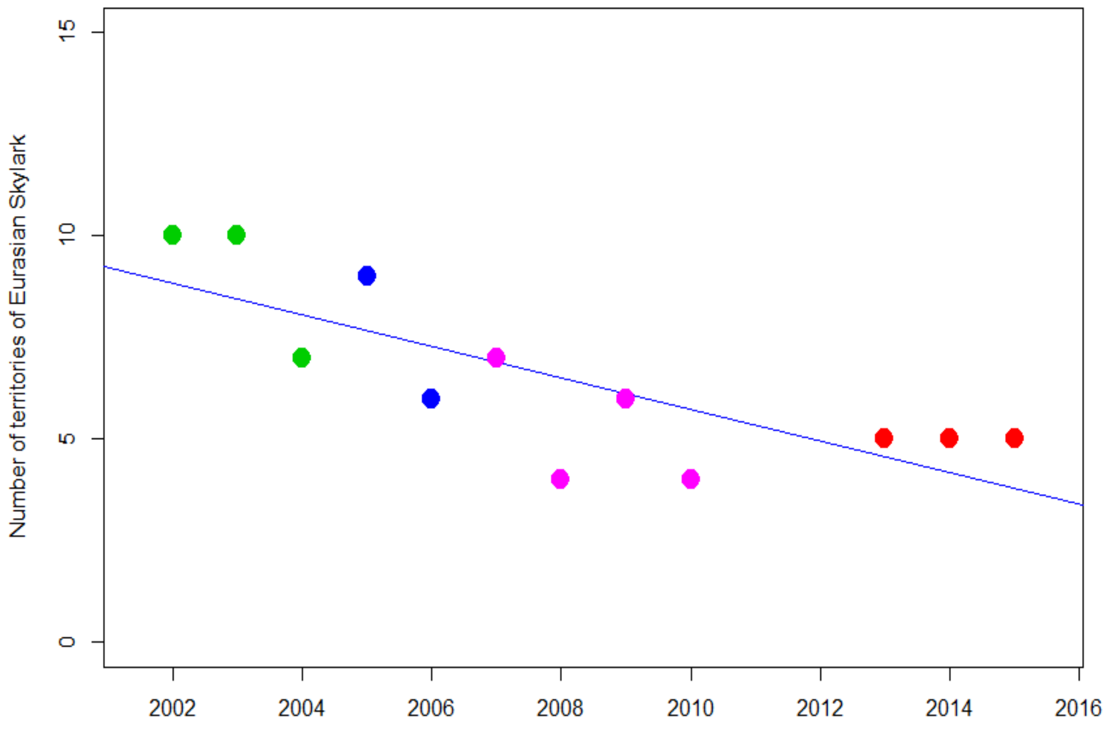

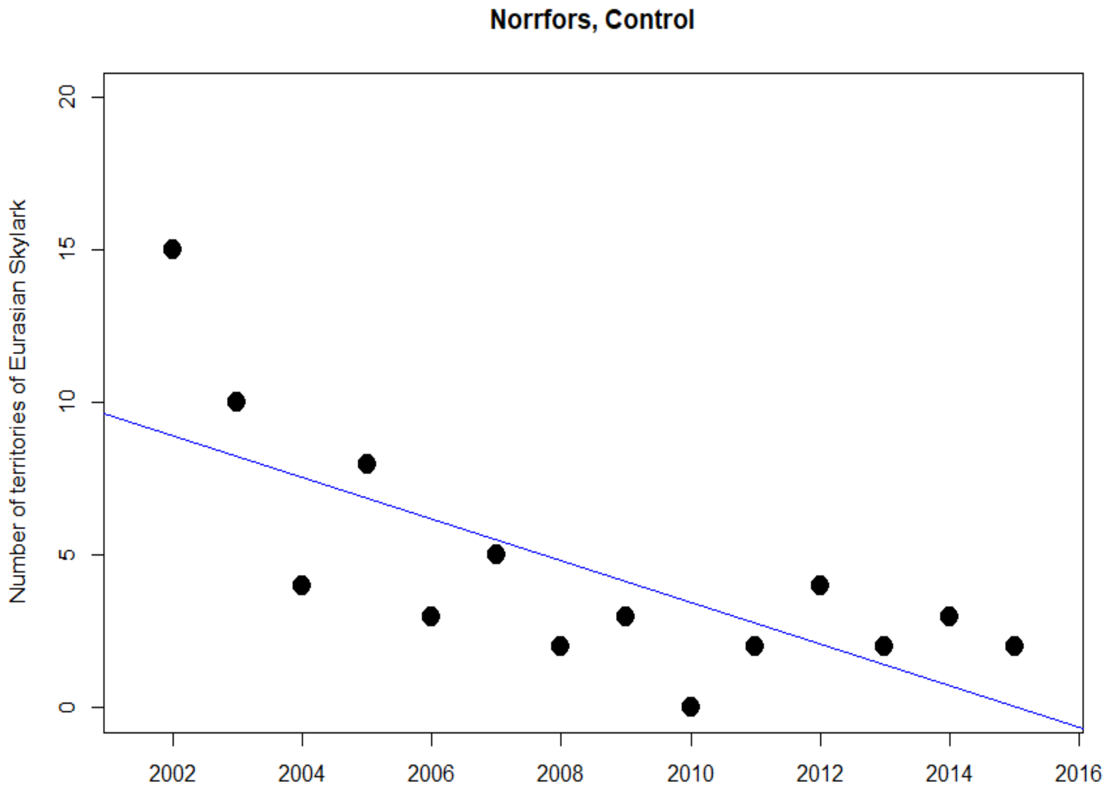
Norrfors, Control
Number of territories of Eurasian Skylark
0
5
10
15
20
2002
2004
2006
2008
2010
2012
2014
2016

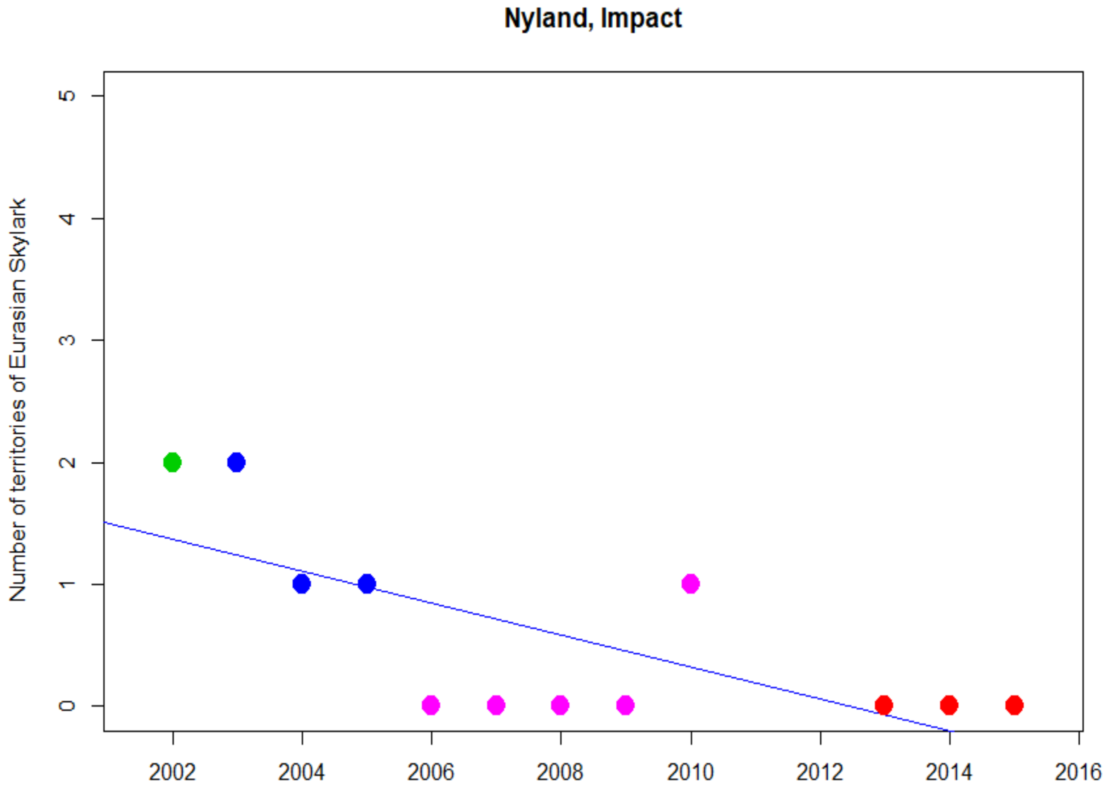
Nyland, Impact
Number of territories of Eurasian Skylark
0
1
2
3
4
5
2002
2004
2006
2008
2010
2012
2014
2016

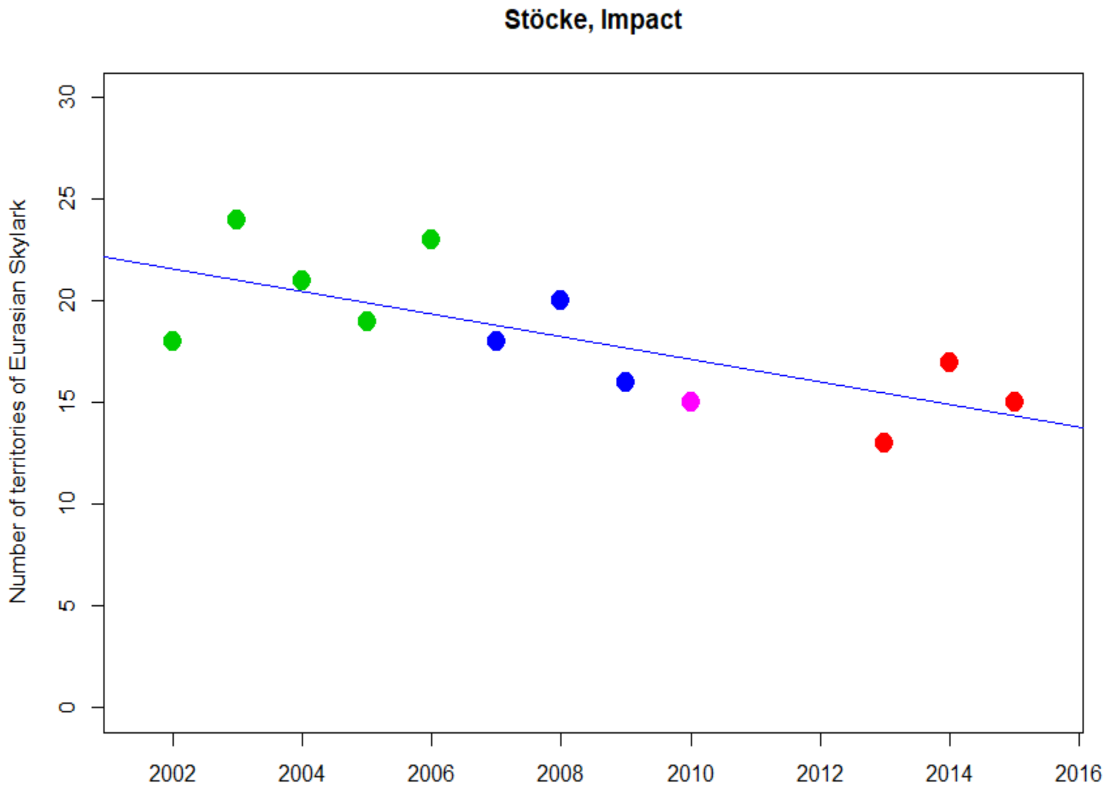
Stöcke, Impact
Number of territories of Eurasian Skylark
0
5
10
15
20
25
30
2002
2004
2006
2008
2010
2012
2014
2016

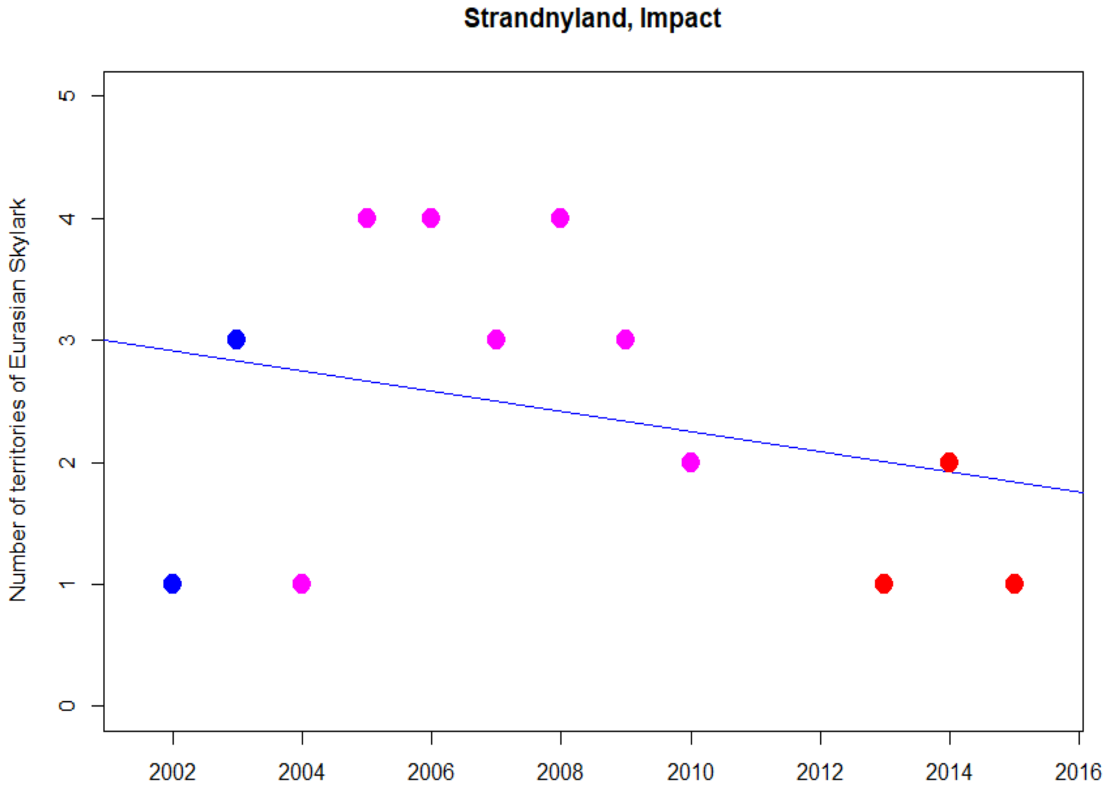
Strandnyland, Impact
Number of territories of Eurasian Skylark
0
1
2
3
4
5
2002
2004
2006
2008
2010
2012
2014
2016

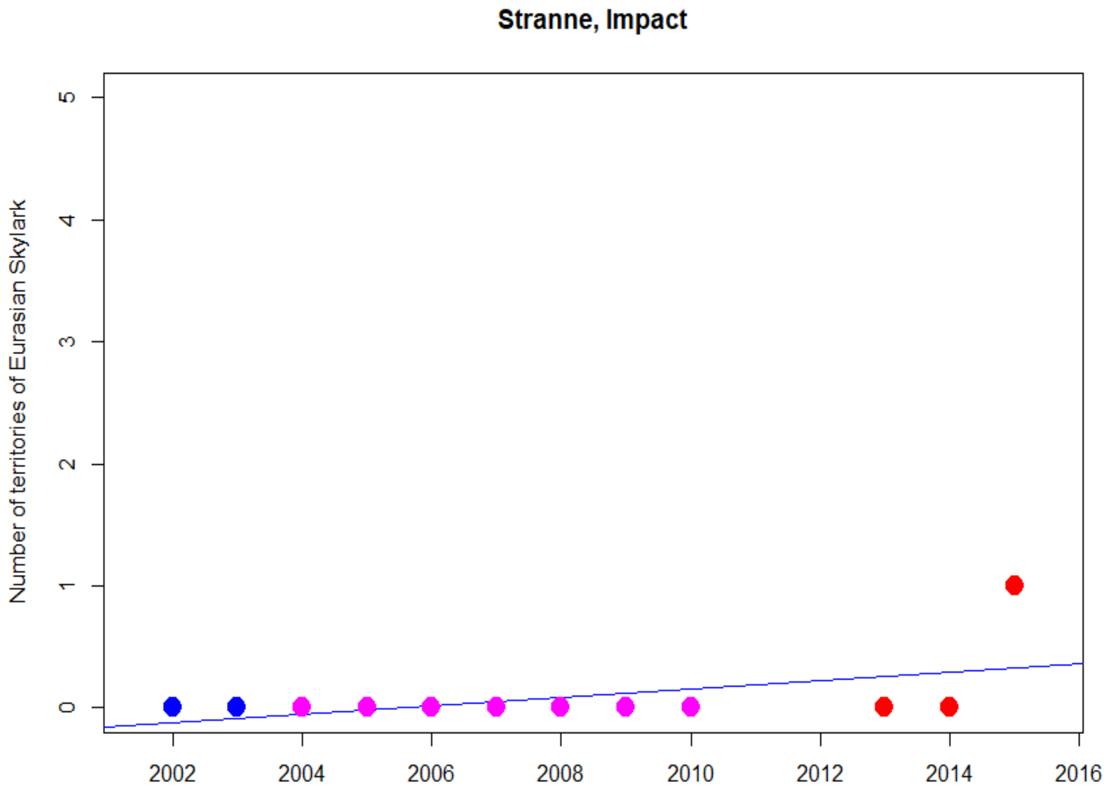

Stranne, Impact
Number of territories of Eurasian Skylark
0
1
2
3
4
5
2002
2004
2006
2008
2010
2012
2014
2016


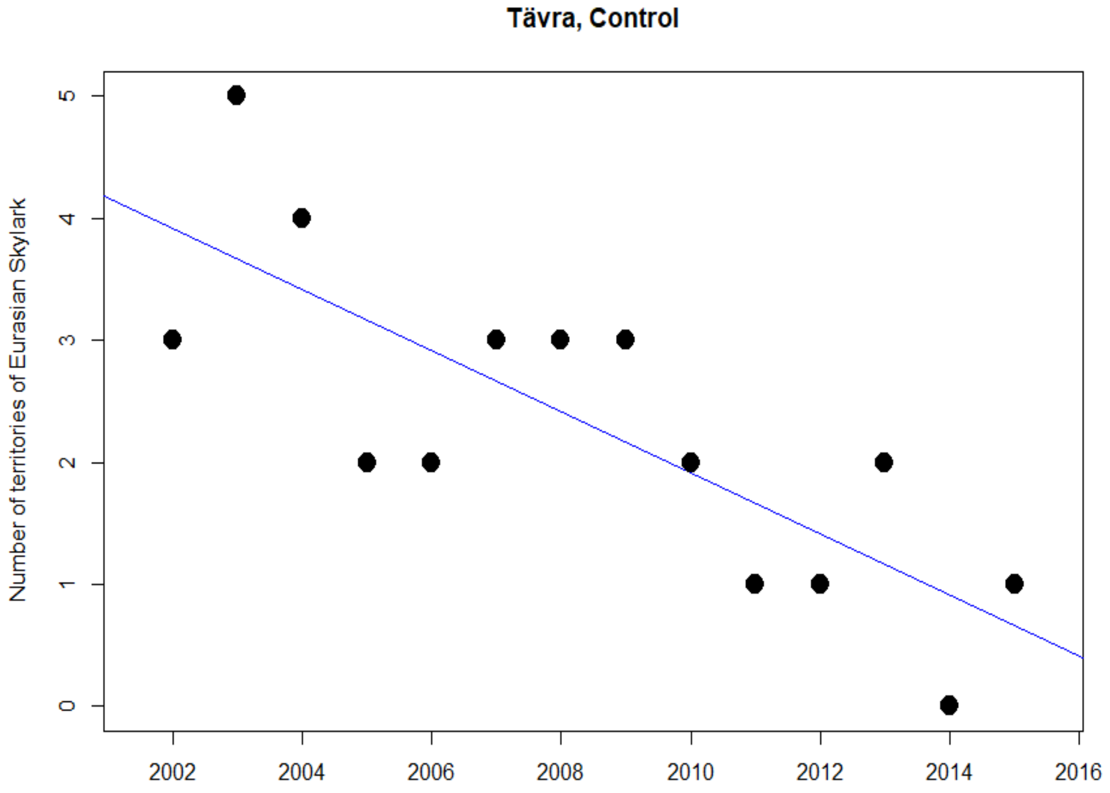

Tävra, Control
Number of territories of Eurasian Skylark
0
1
2
3
4
5
2002
2004
2006
2008
2010
2012
2014
2016

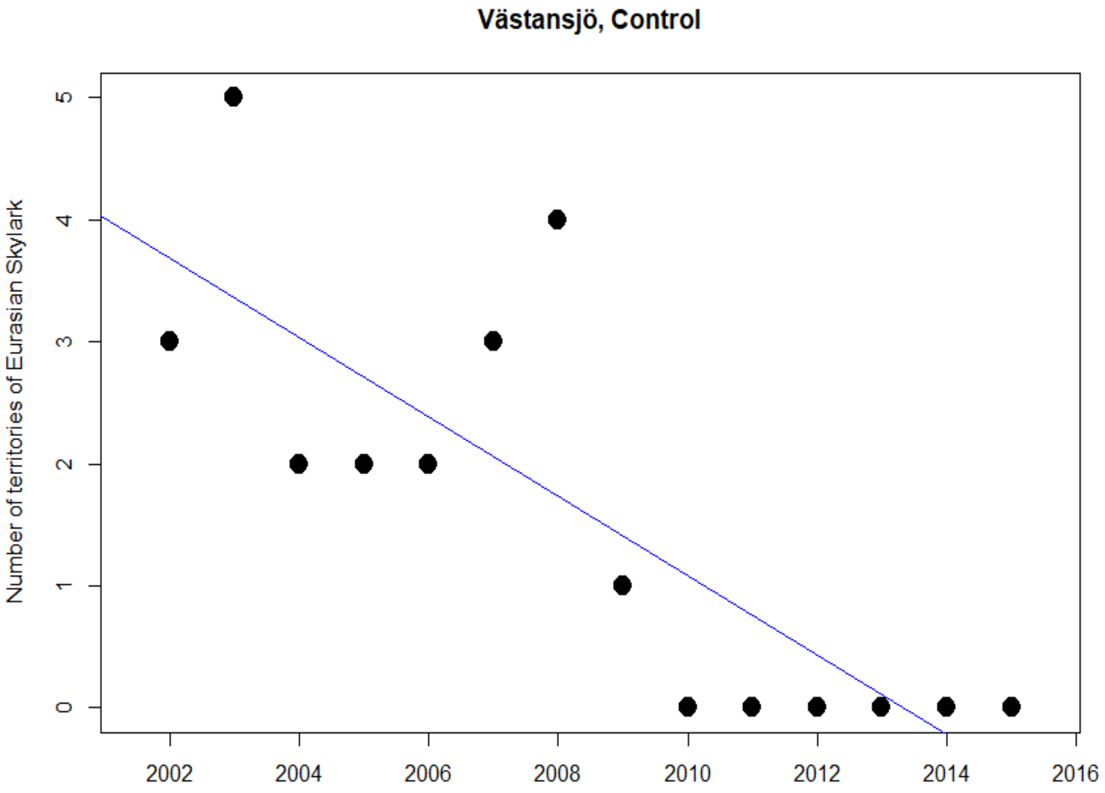
Västansjö, Control
Number of territories of Eurasian Skylark
0
1
2
3
4
5
2002
2004
2006
2008
2010
2012
2014
2016

### Green Sandpiper

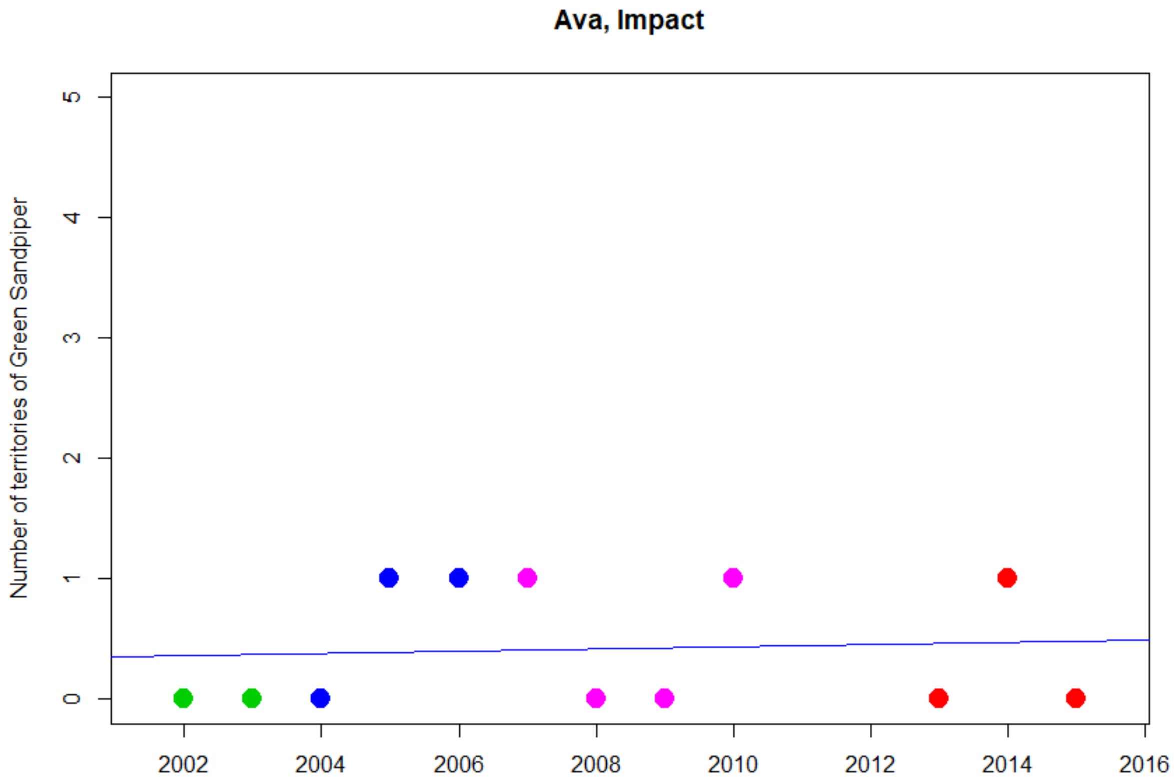


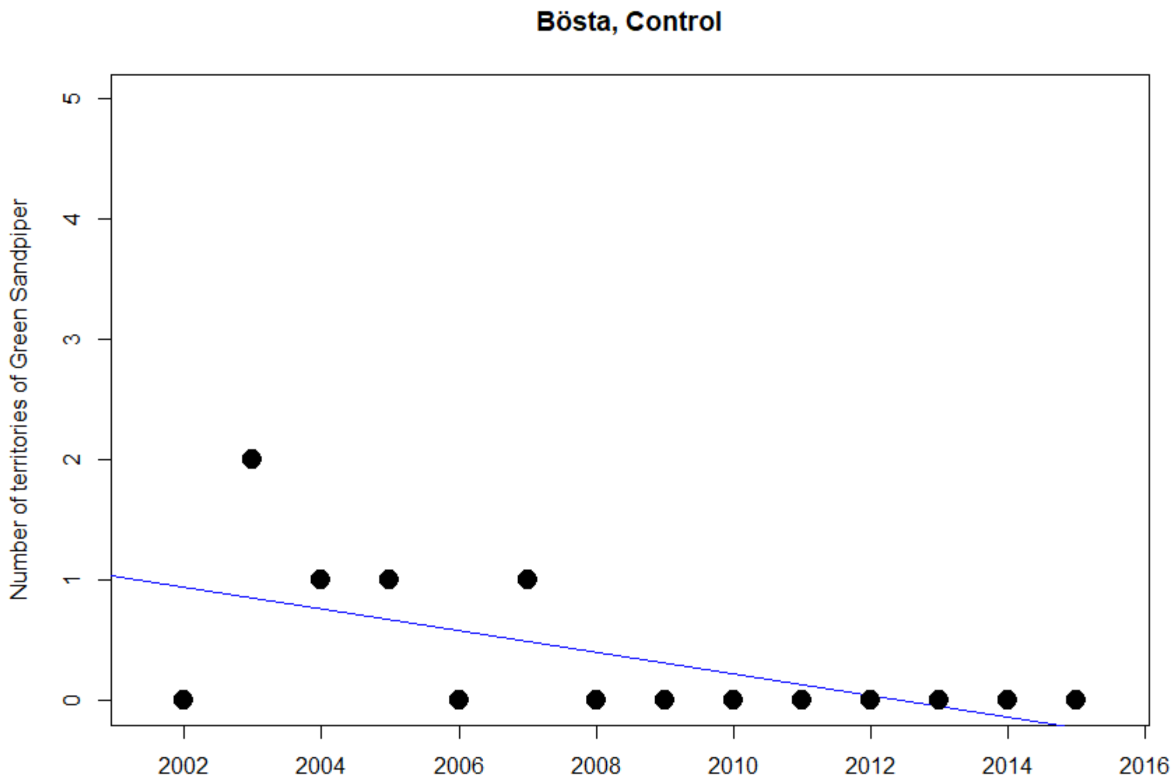

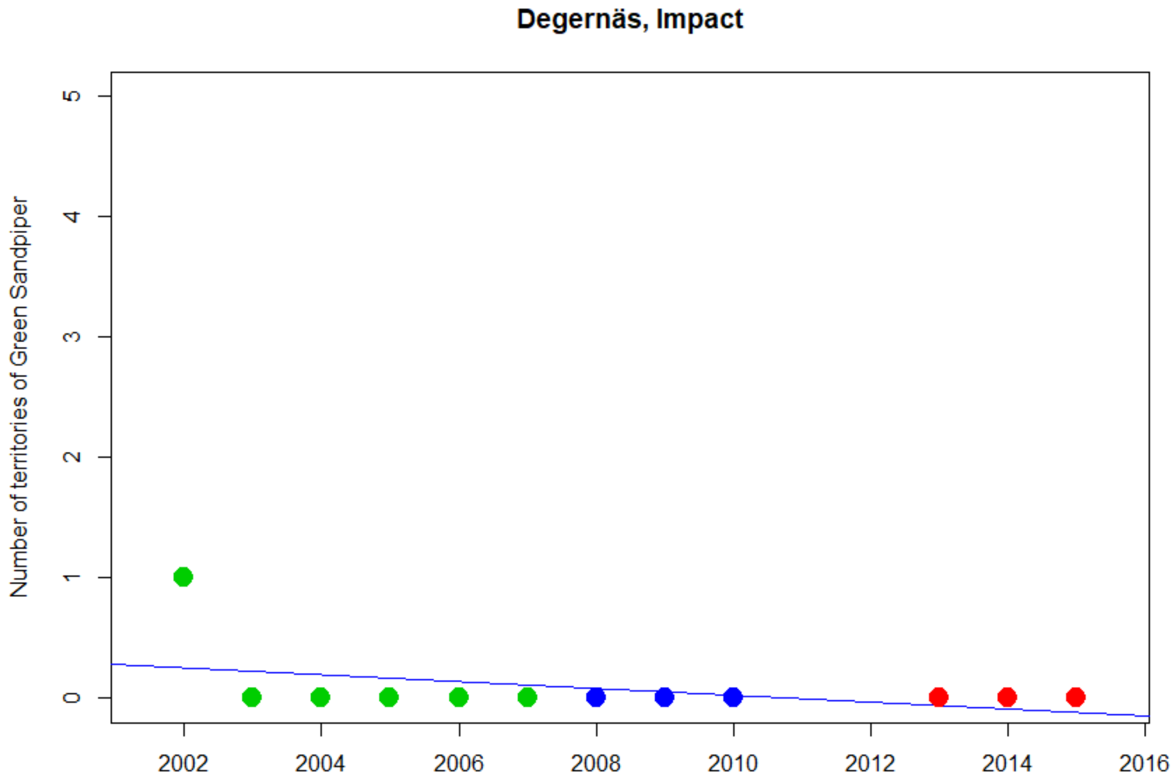
Degernäs, Impact
Number of territories of Green Sandpiper
0
1
2
3
4
5
2002
2004
2006
2008
2010
2012
2014
2016

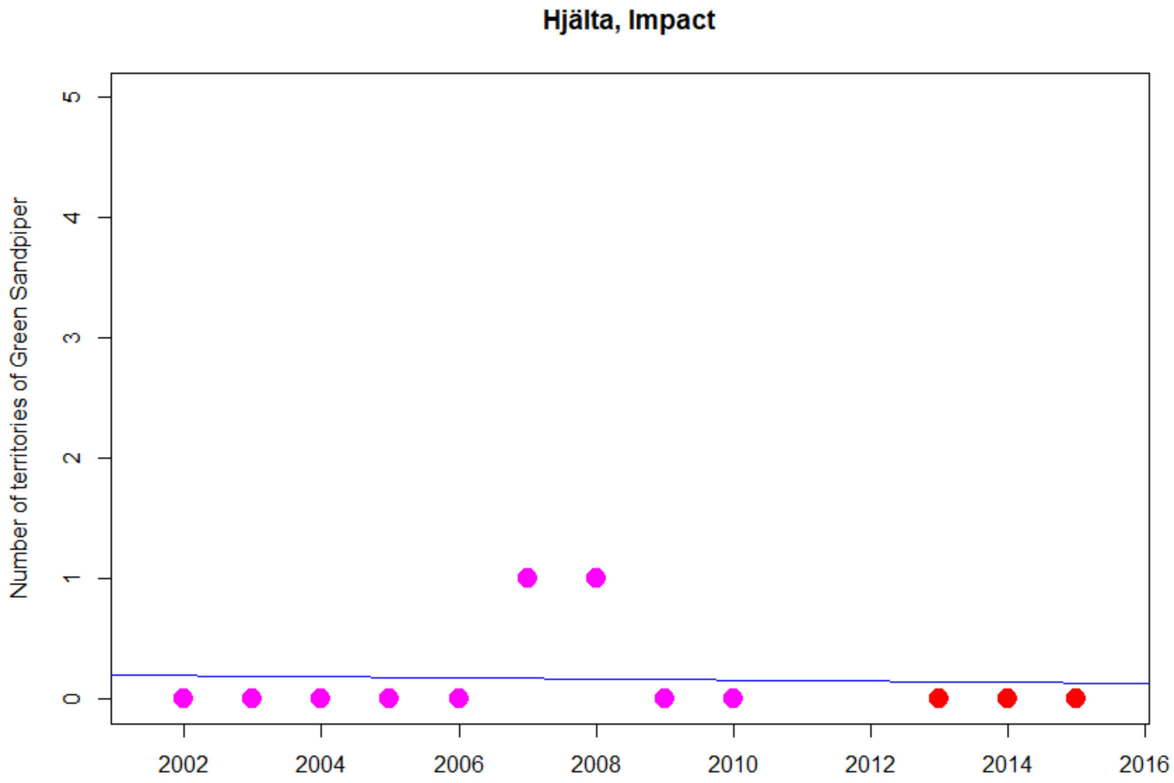
Hjälta, Impact
Number of territories of Green Sandpiper
0
1
2
3
4
5
2002
2004
2006
2008
2010
2012
2014
2016

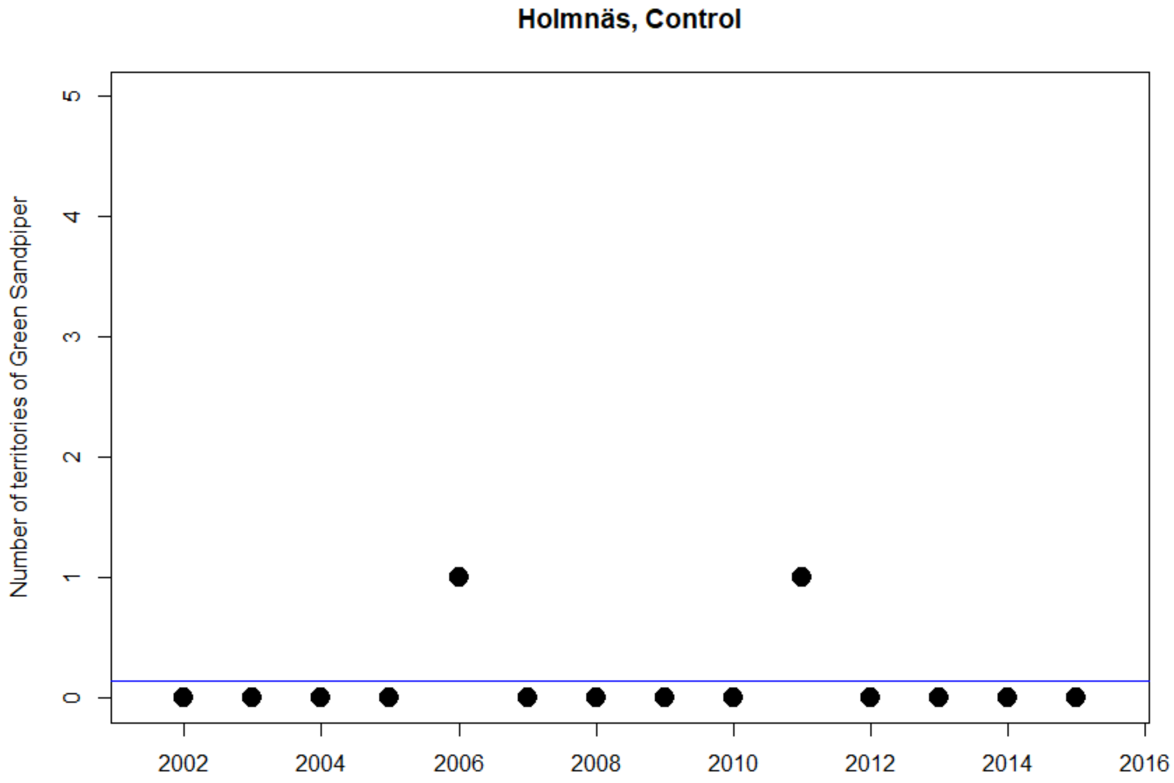
Holmnäs, Control
Number of territories of Green Sandpiper
0
1
2
3
4
5
2002
2004
2006
2008
2010
2012
2014
2016

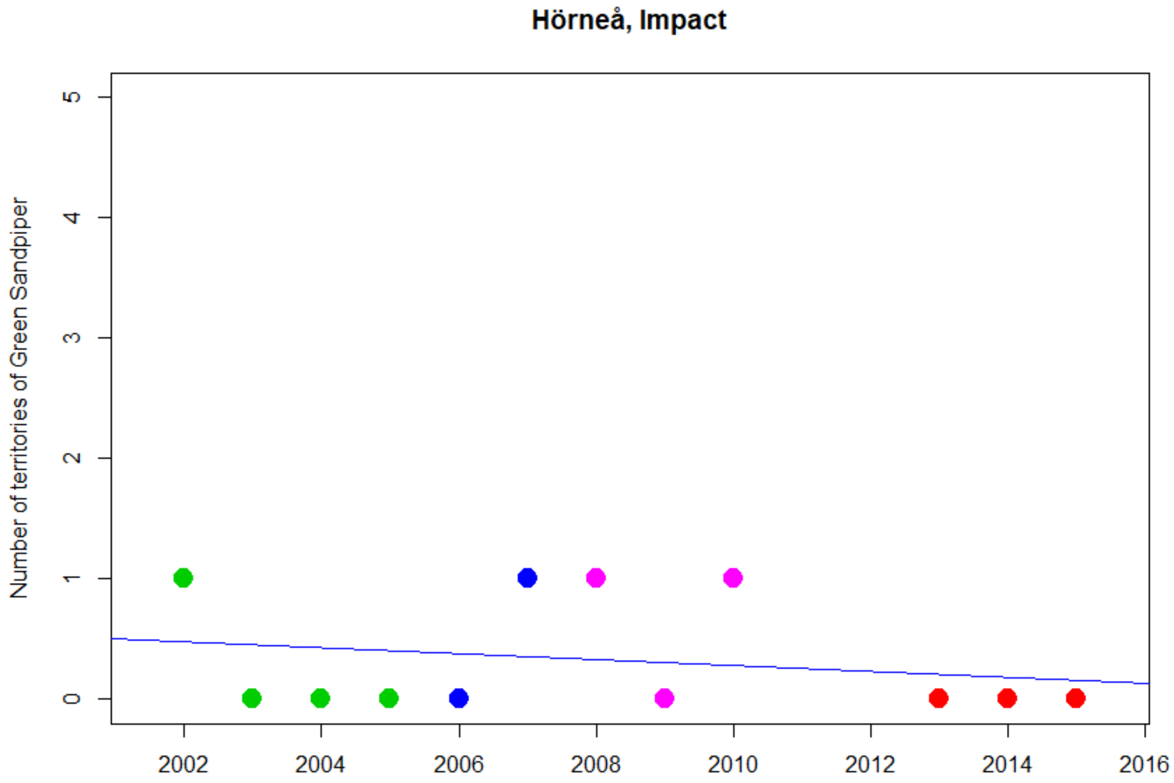
Hörneå, Impact
Number of territories of Green Sandpiper
0
1
2
3
4
5
2002
2004
2006
2008
2010
2012
2014
2016

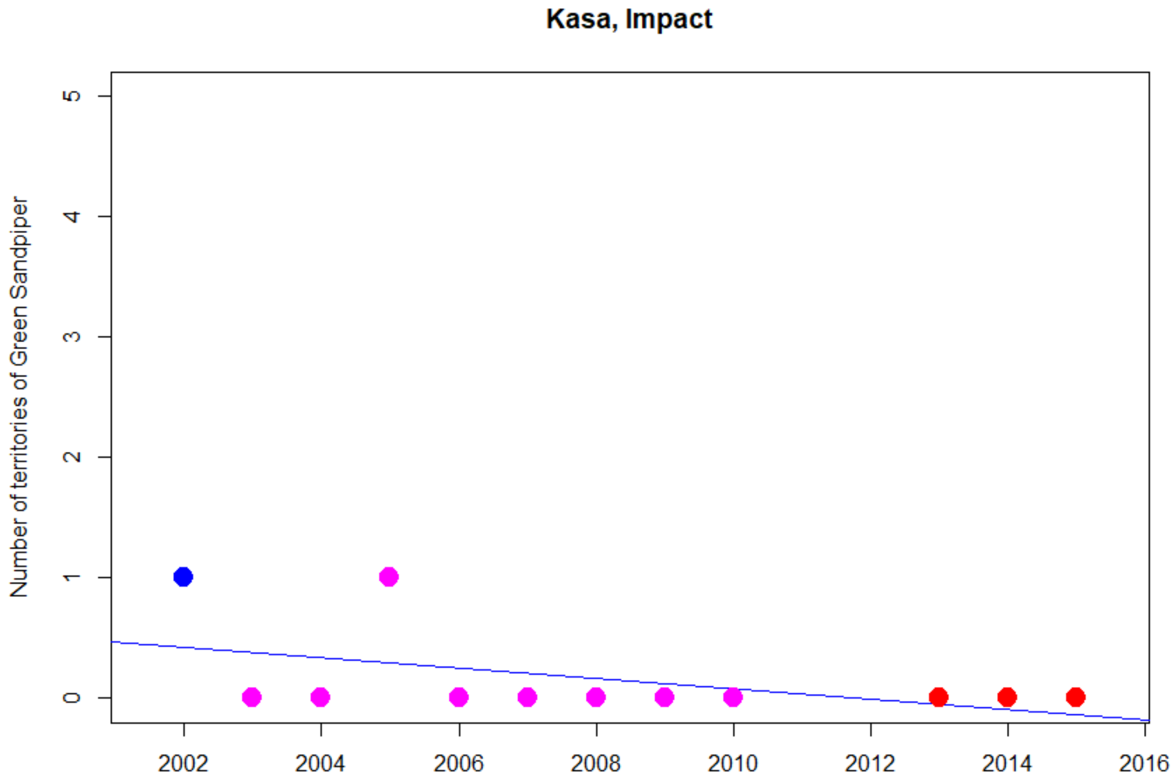
Kasa, Impact
Number of territories of Green Sandpiper
0
1
2
3
4
5
2002
2004
2006
2008
2010
2012
2014
2016

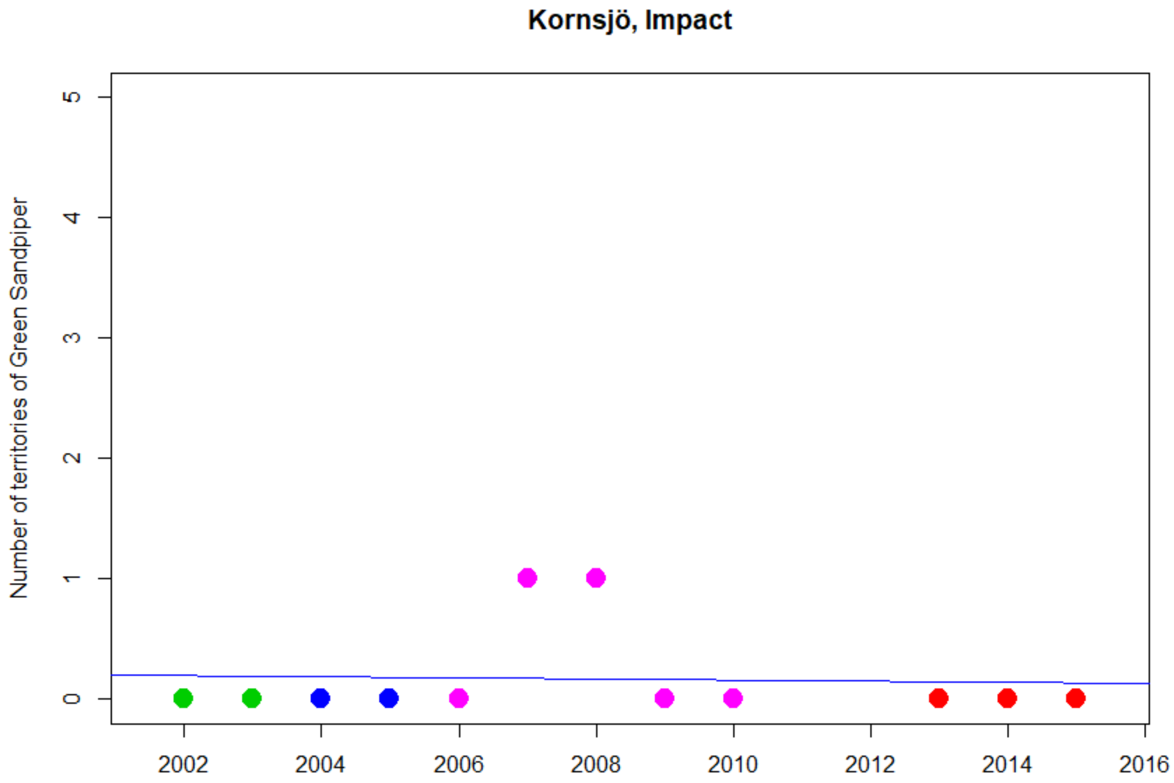
Kornsjö, Impact
Number of territories of Green Sandpiper
0
1
2
3
4
5
2002
2004
2006
2008
2010
2012
2014
2016

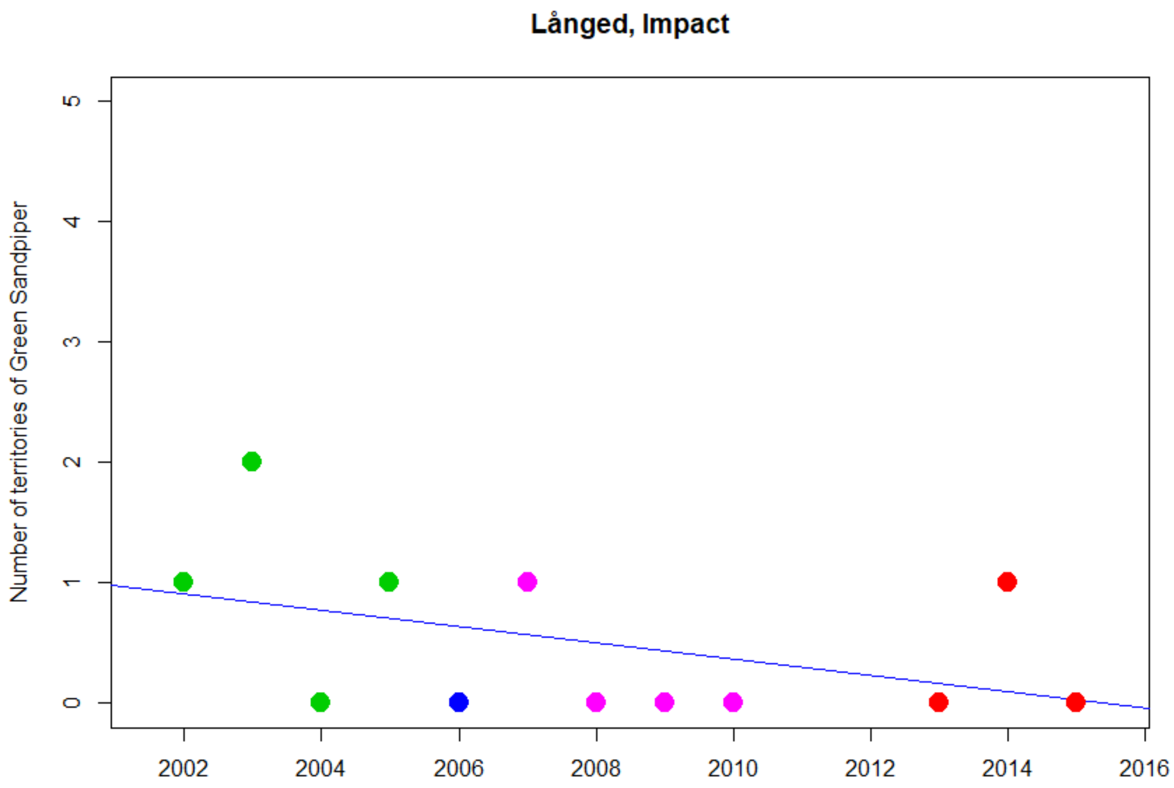
Långed, Impact
Number of territories of Green Sandpiper
0
1
2
3
4
5
2002
2004
2006
2008
2010
2012
2014
2016

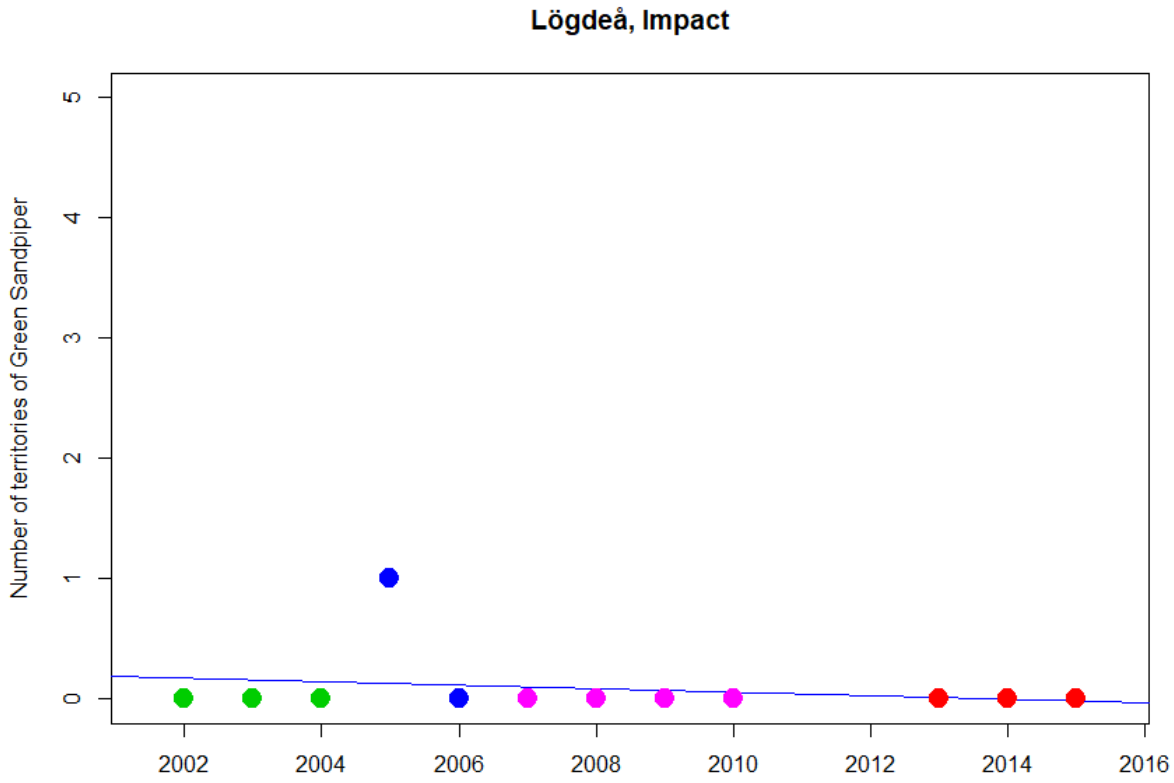
Lögdeå, Impact
Number of territories of Green Sandpiper
0
1
2
3
4
5
2002
2004
2006
2008
2010
2012
2014
2016

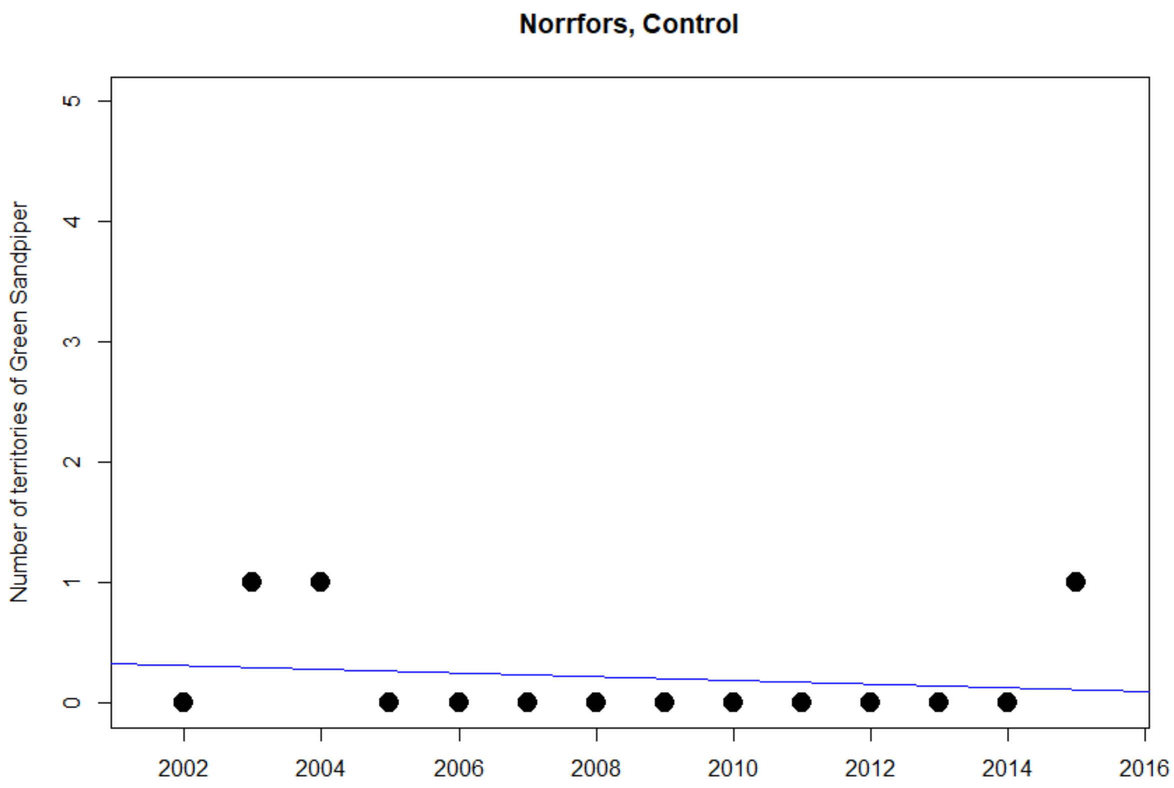
Norrfors, Control
Number of territories of Green Sandpiper
0
1
2
3
4
5
2002
2004
2006
2008
2010
2012
2014
2016

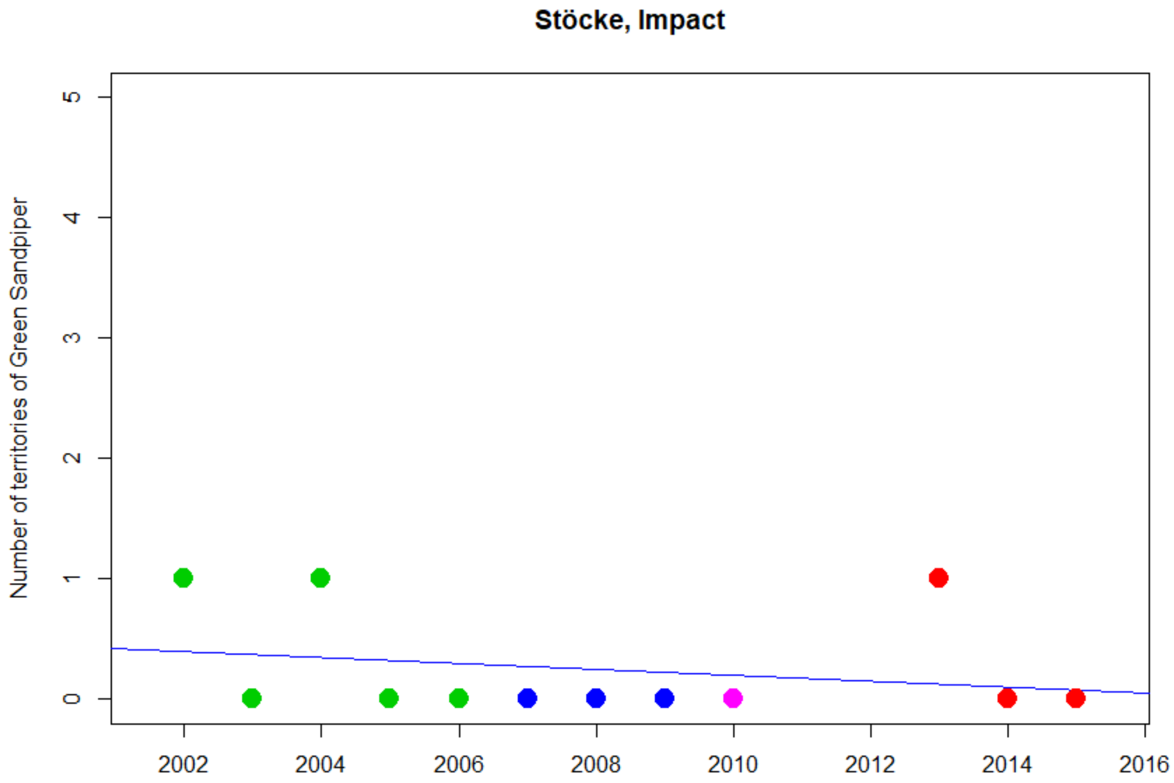
Stöcke, Impact
Number of territories of Green Sandpiper
0
1
2
3
4
5
2002
2004
2006
2008
2010
2012
2014
2016

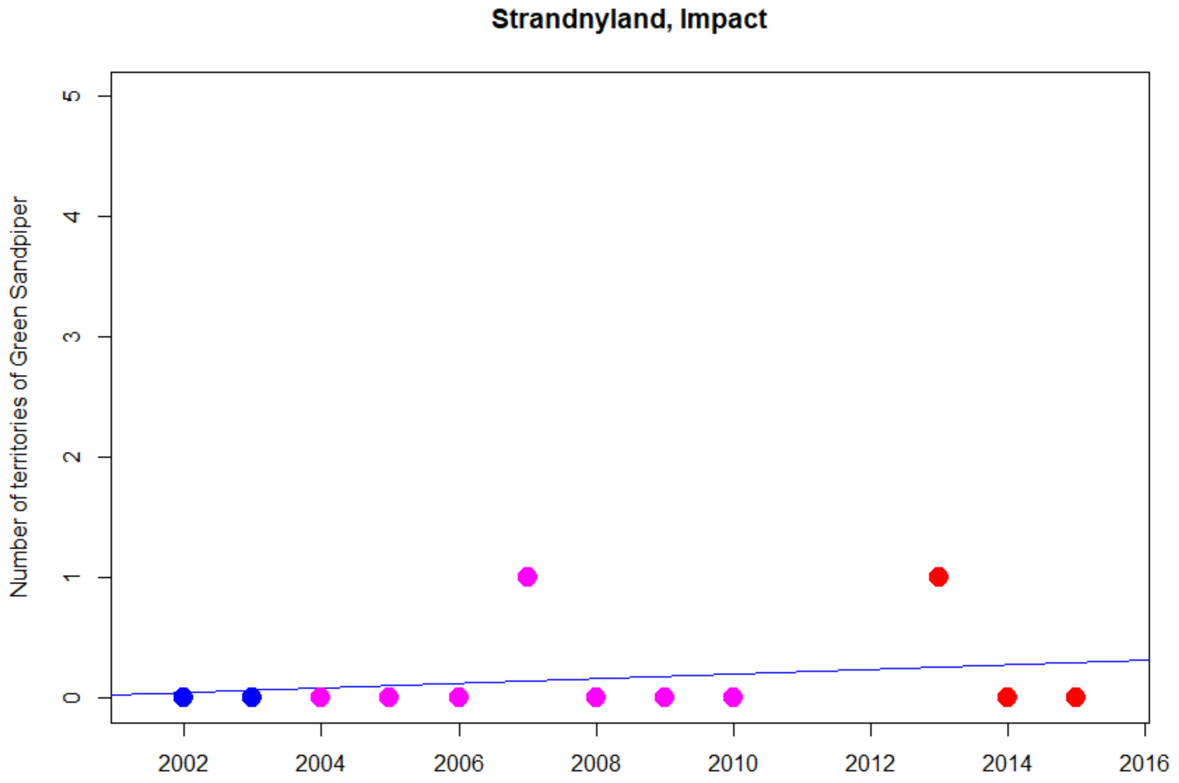
Strandnyland, Impact
Number of territories of Green Sandpiper
0
1
2
3
4
5
2002
2004
2006
2008
2010
2012
2014
2016

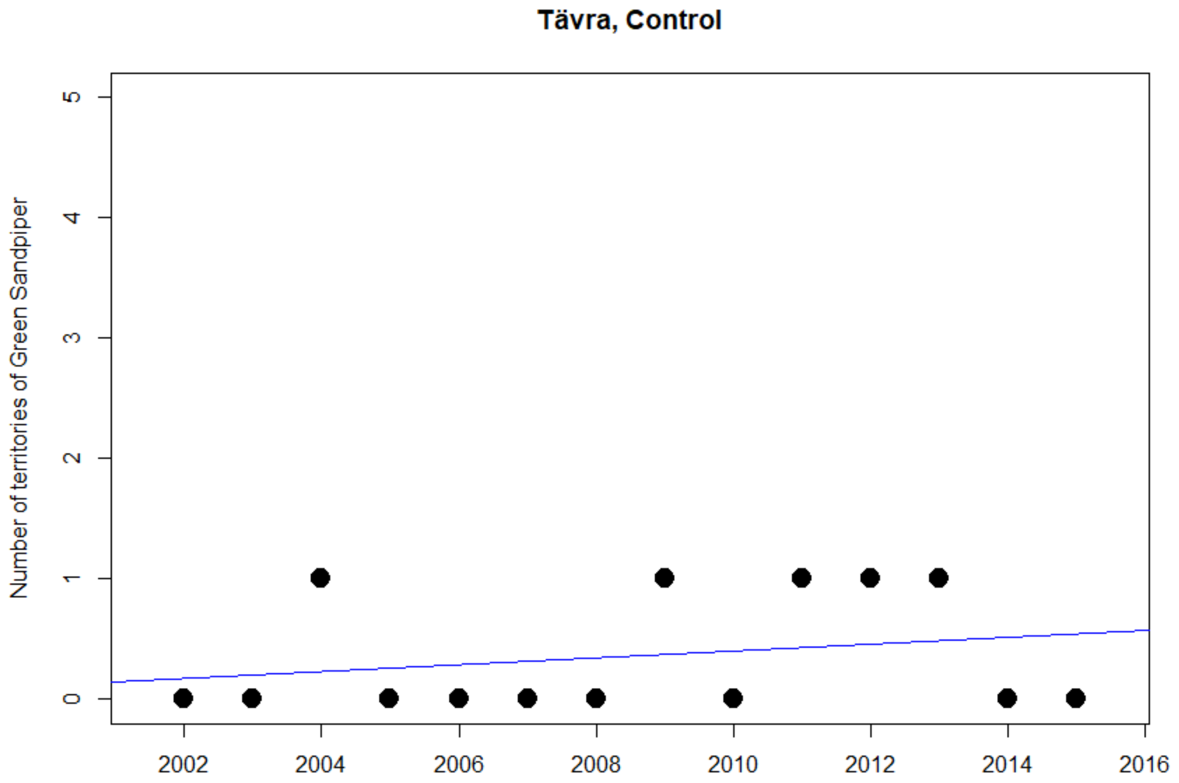
Tävra, Control
Number of territories of Green Sandpiper
0
1
2
3
4
5
2002
2004
2006
2008
2010
2012
2014
2016

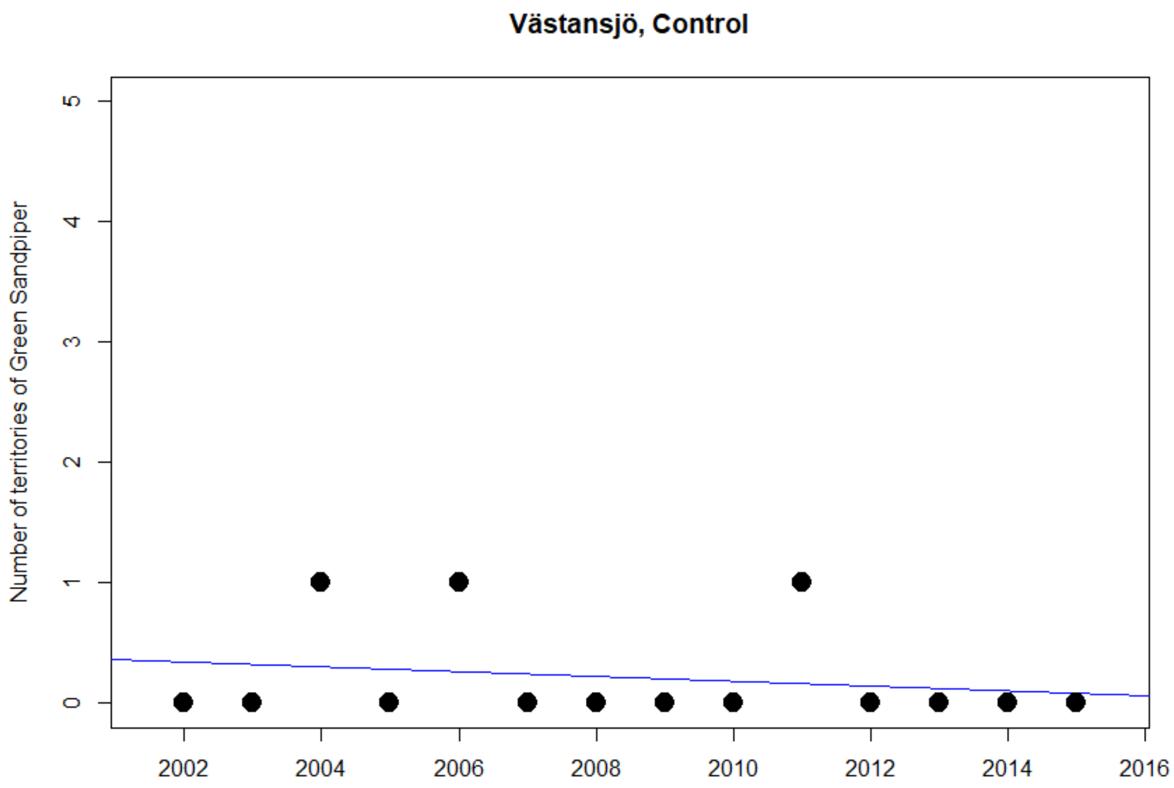

Västansjö, Control
Number of territories of Green Sandpiper
0
1
2
3
4
5
2002
2004
2006
2008
2010
2012
2014
2016

### Little Ringed Plover

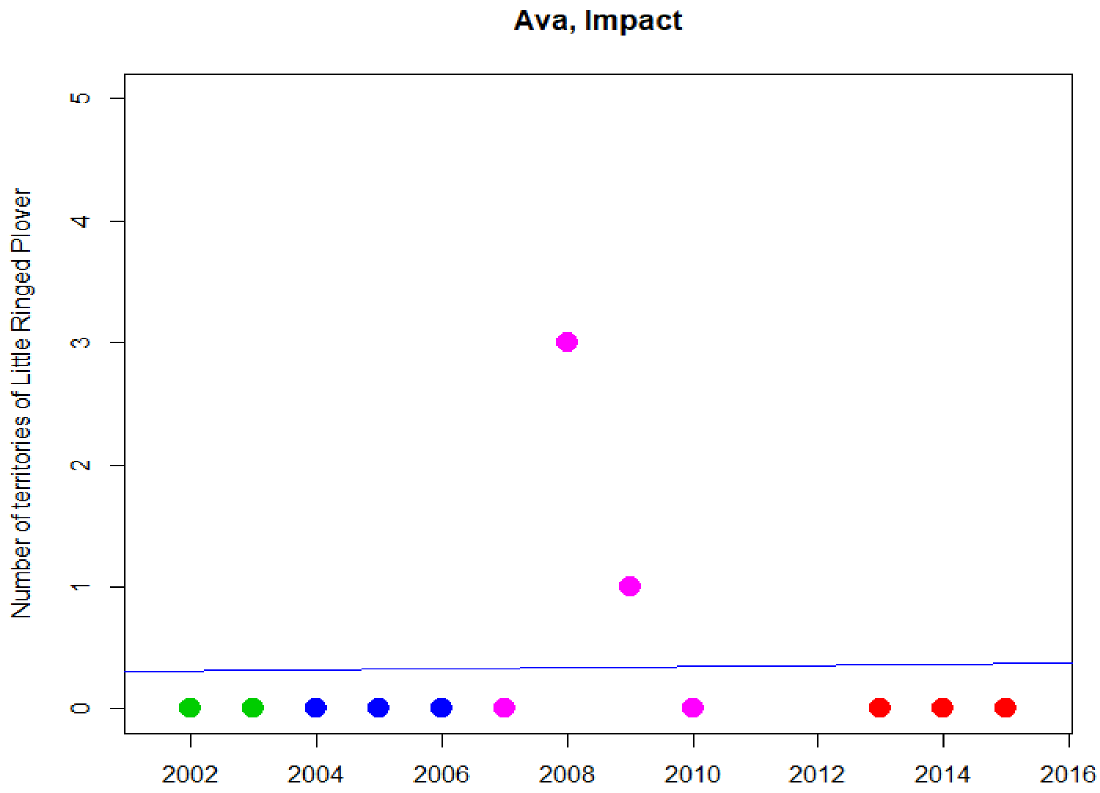


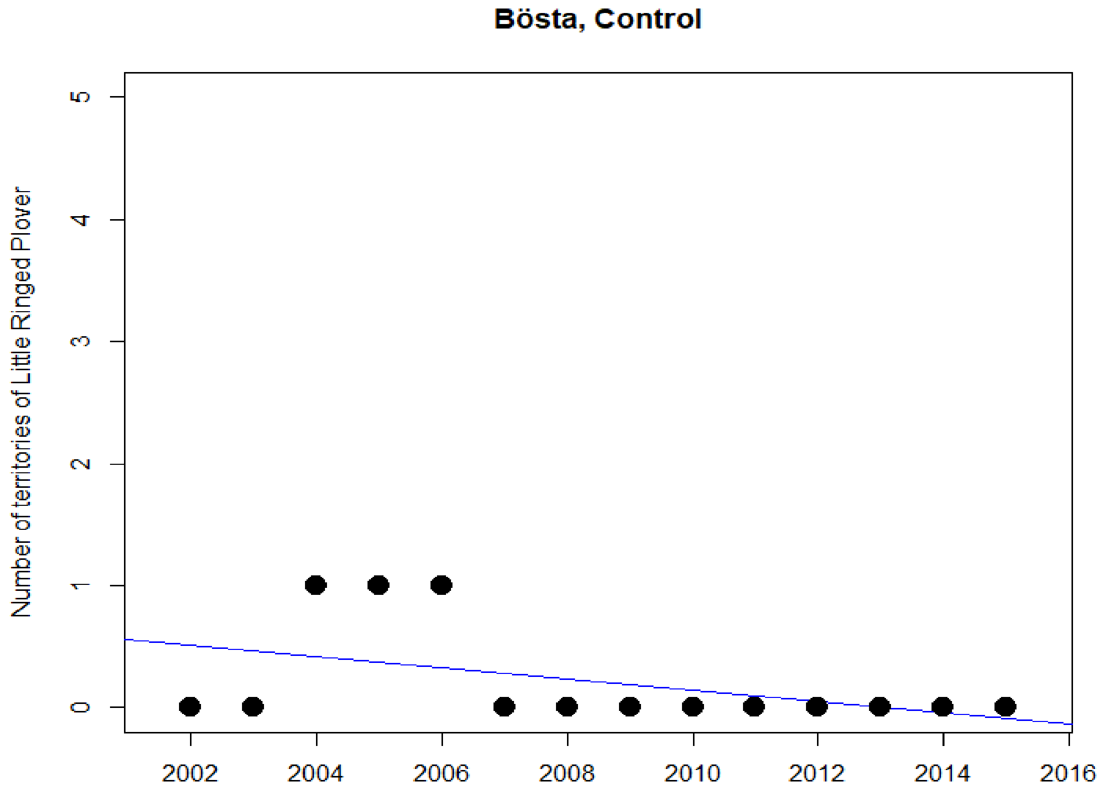

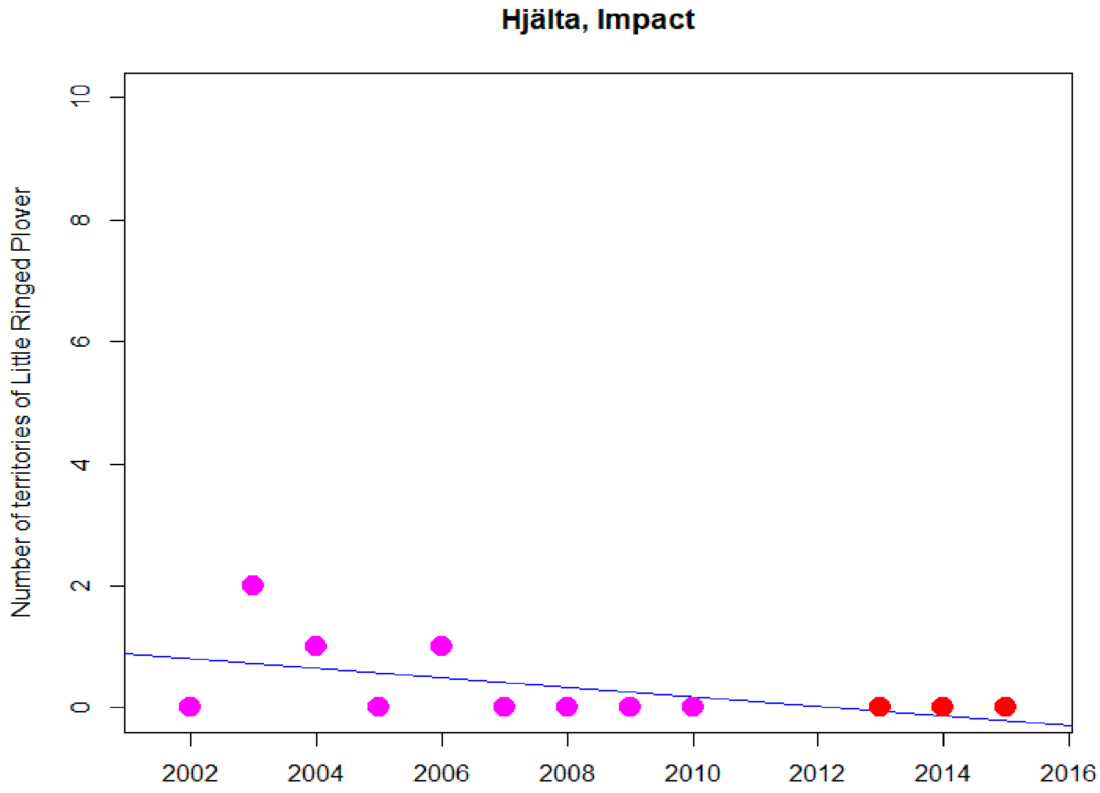
Hjälta, Impact
Number of territories of Little Ringed Plover
0
2
4
6
8
10
2002
2004
2006
2008
2010
2012
2014
2016

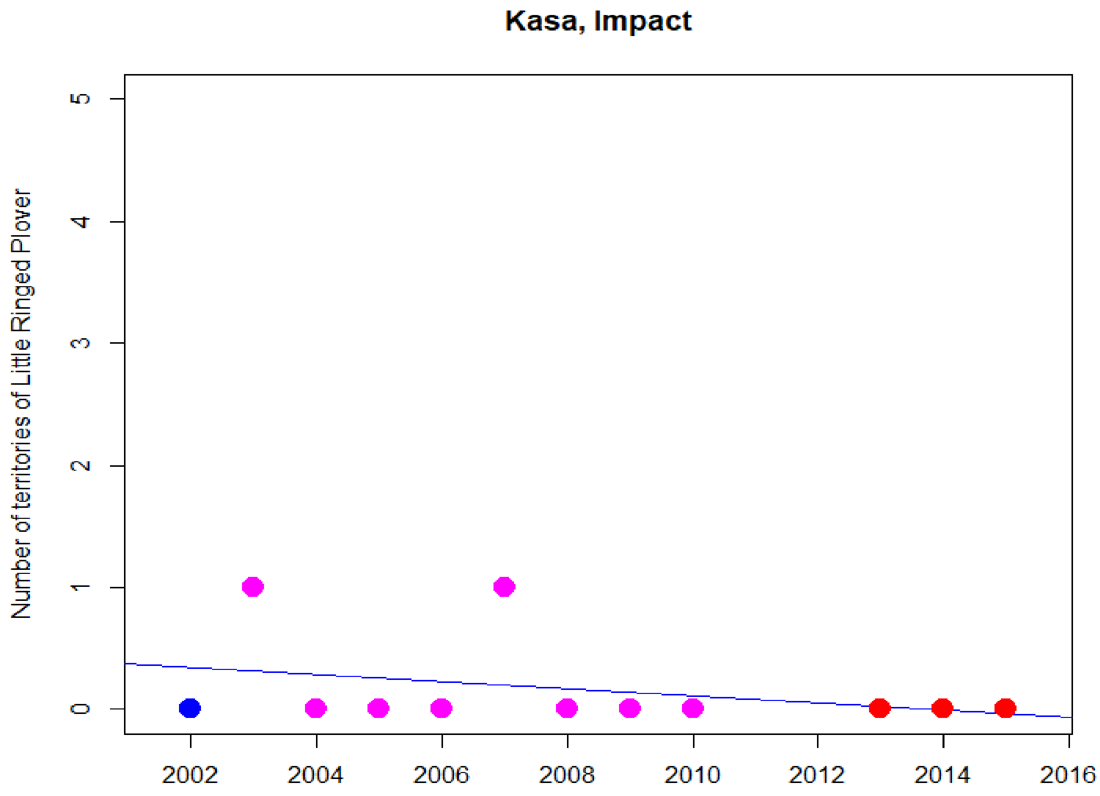
Kasa, Impact
Number of territories of Little Ringed Plover
0
1
2
3
4
5
2002
2004
2006
2008
2010
2012
2014
2016

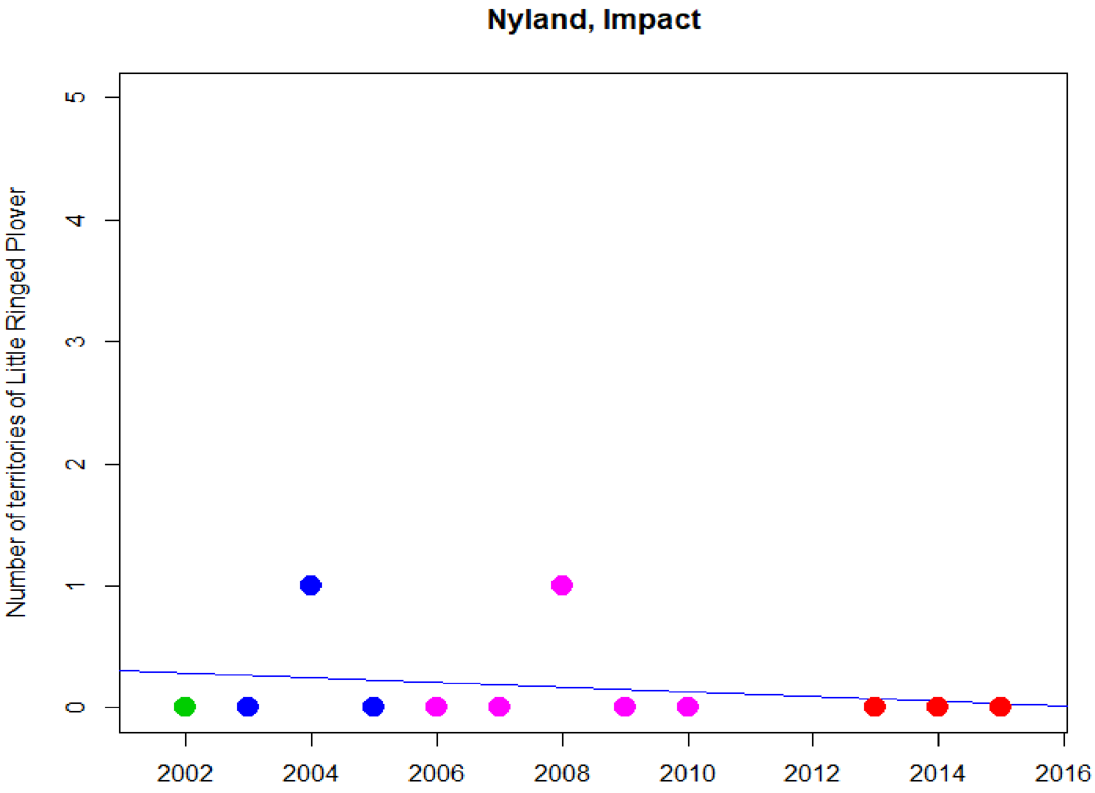

Nyland, Impact
Number of territories of Little Ringed Plover
0
1
2
3
4
5
2002
2004
2006
2008
2010
2012
2014
2016


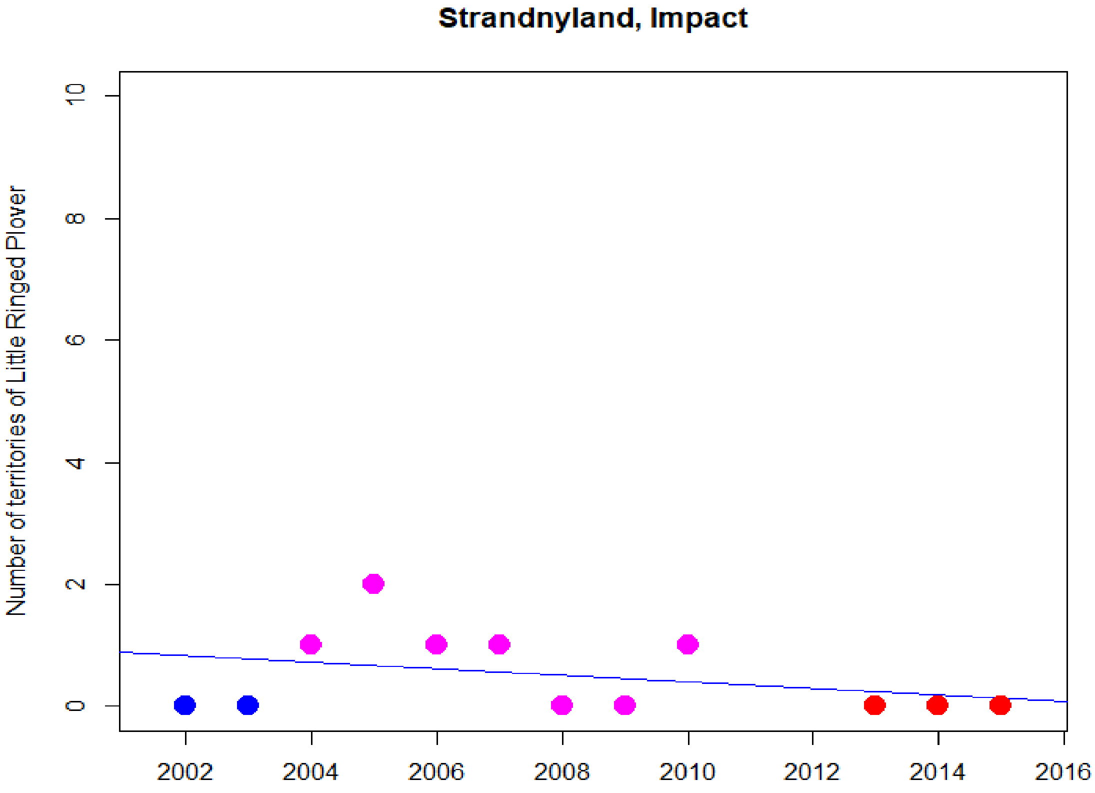

Strandnyland, Impact
Number of territories of Little Ringed Plover
0
2
4
6
8
10
2002
2004
2006
2008
2010
2012
2014
2016

### Meadow Pipit

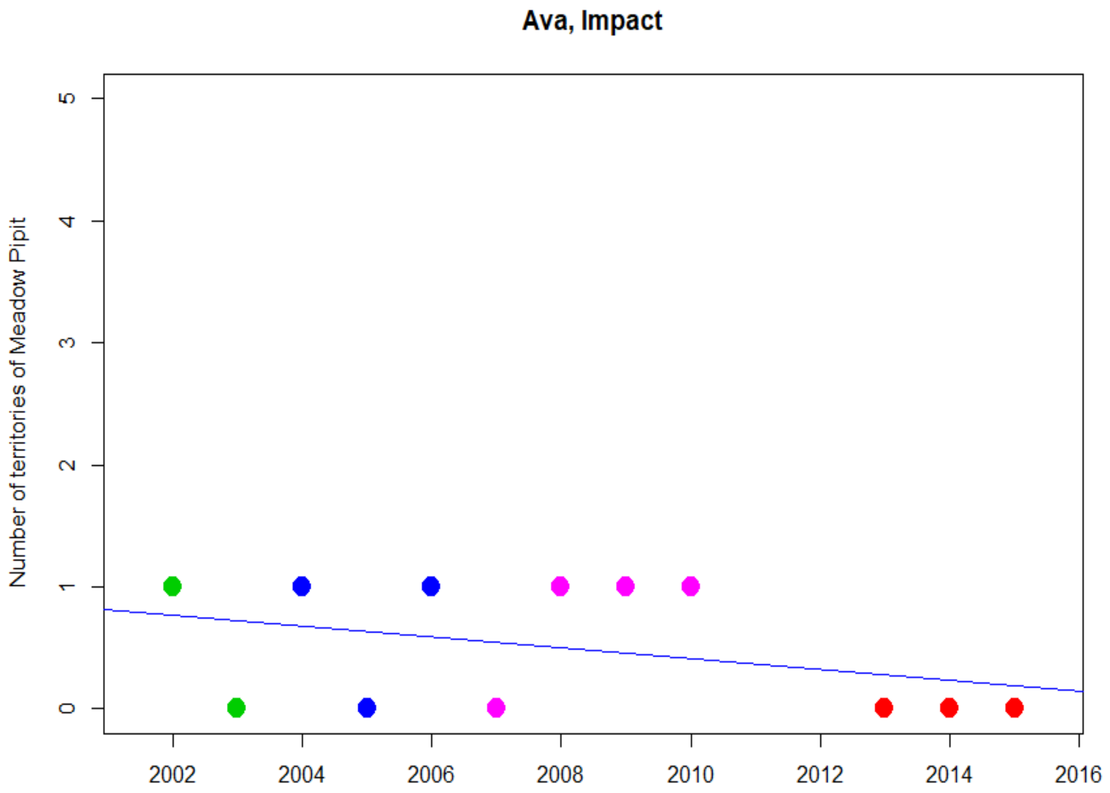


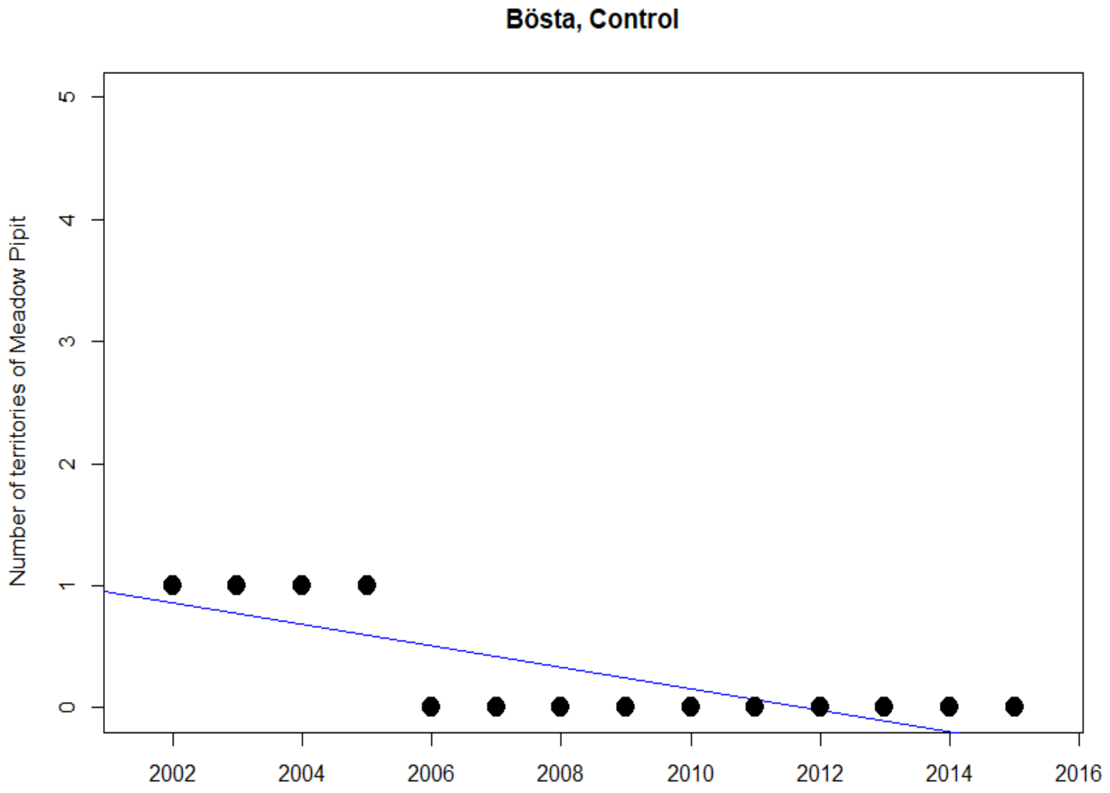

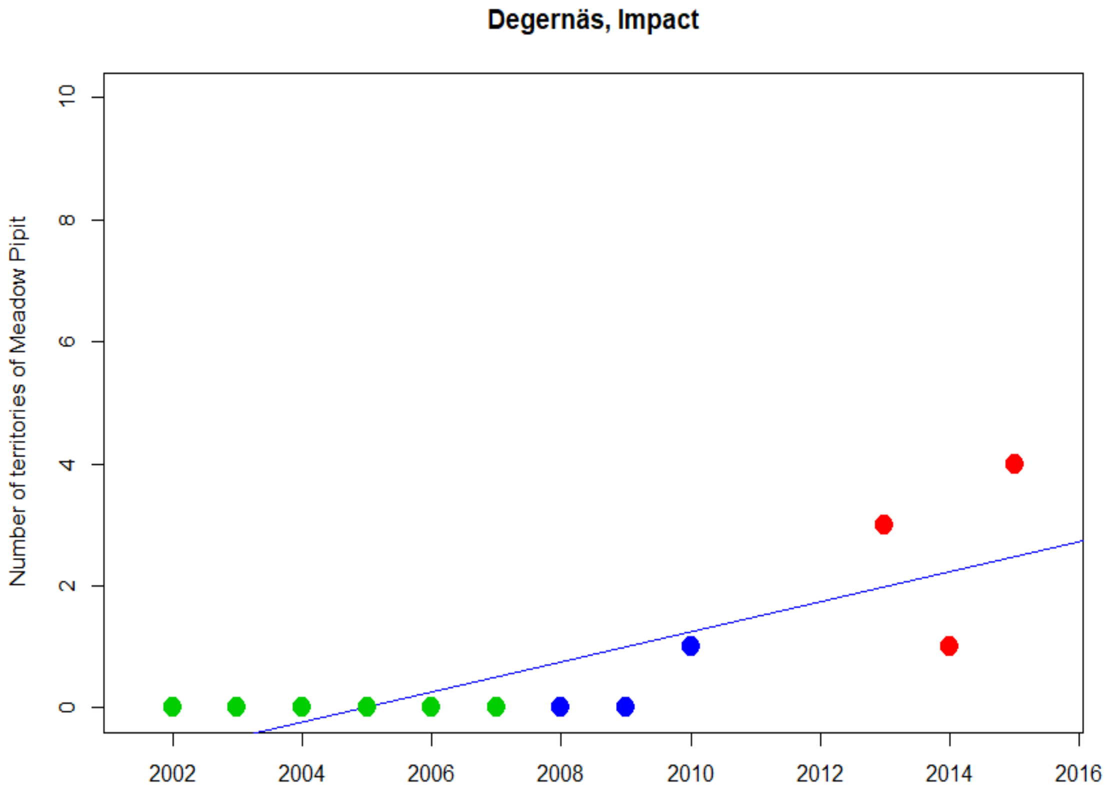
Degernäs, Impact
Number of territories of Meadow Pipit
0
2
4
6
8
10
2002
2004
2006
2008
2010
2012
2014
2016

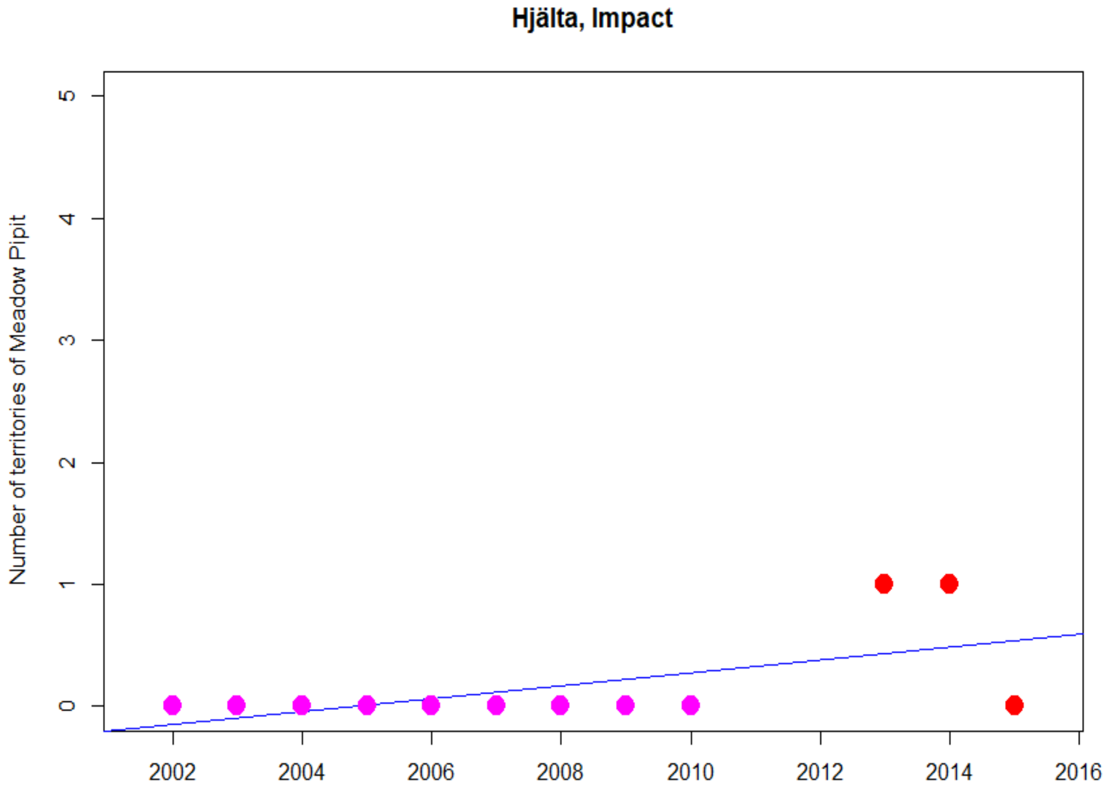
Hjälta, Impact
Number of territories of Meadow Pipit
0
1
2
3
4
5
2002
2004
2006
2008
2010
2012
2014
2016

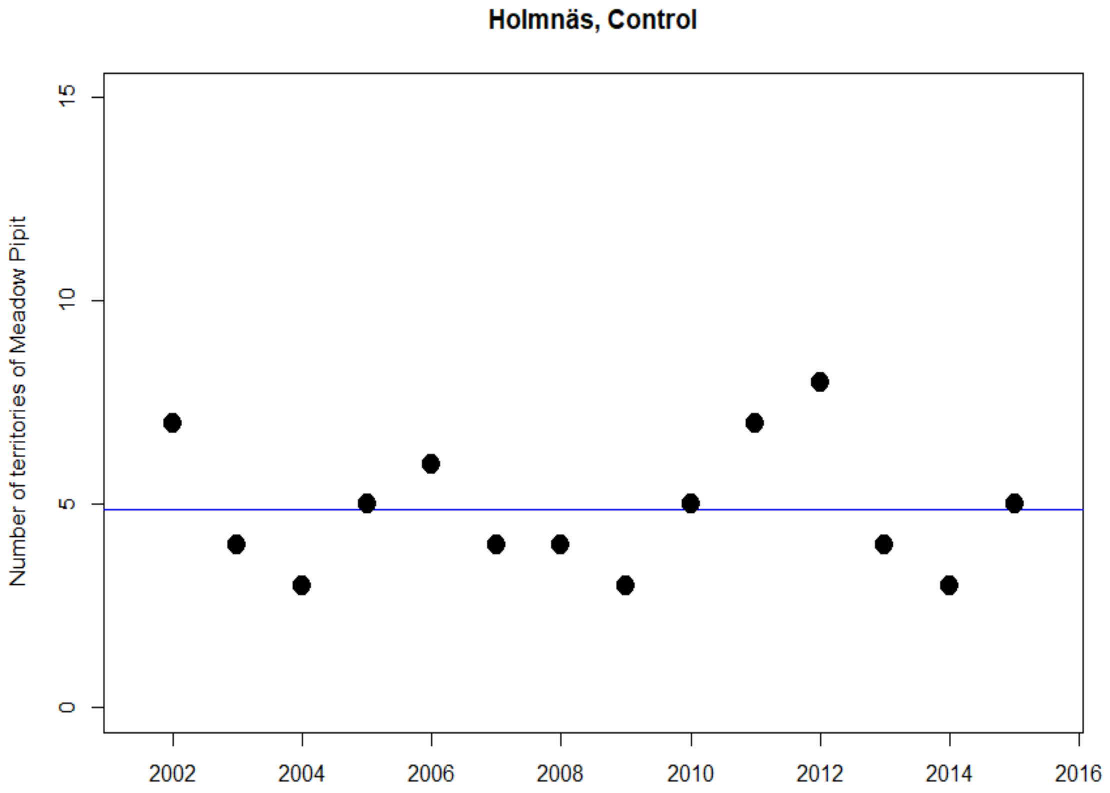
Holmnäs, Control
Number of territories of Meadow Pipit
0
5
10
15
2002
2004
2006
2008
2010
2012
2014
2016

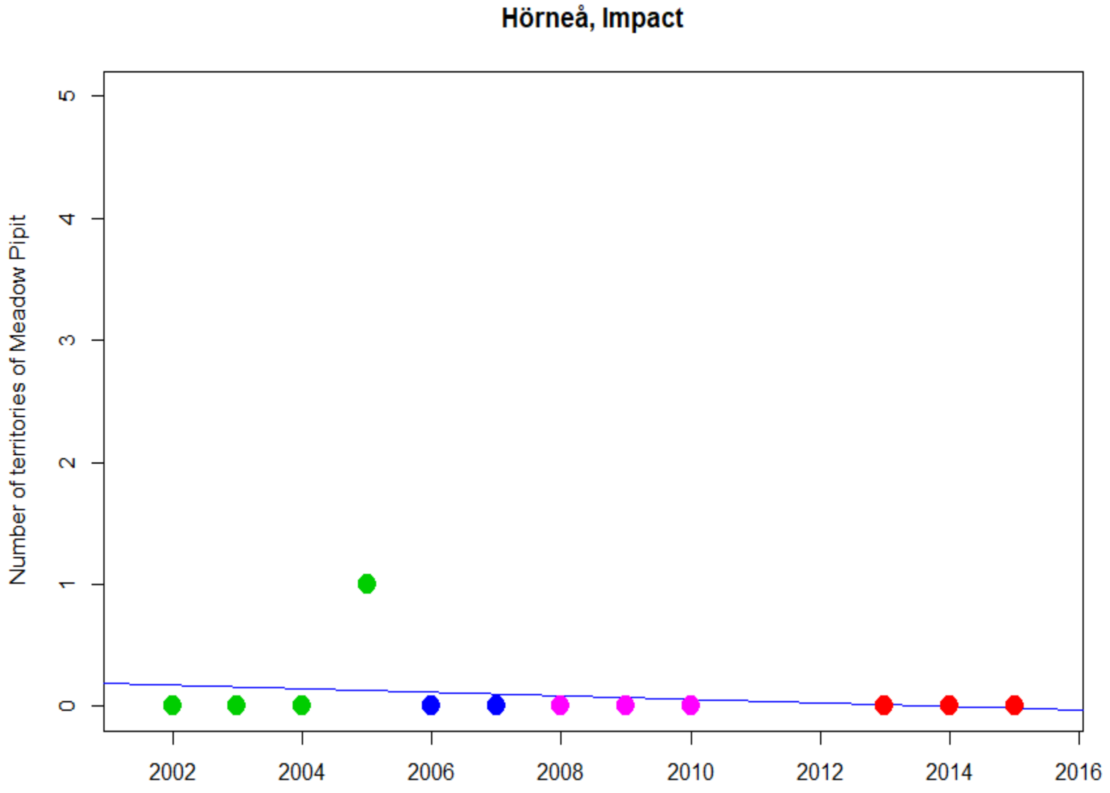
Hörneå, Impact
Number of territories of Meadow Pipit
0
1
2
3
4
5
2002
2004
2006
2008
2010
2012
2014
2016

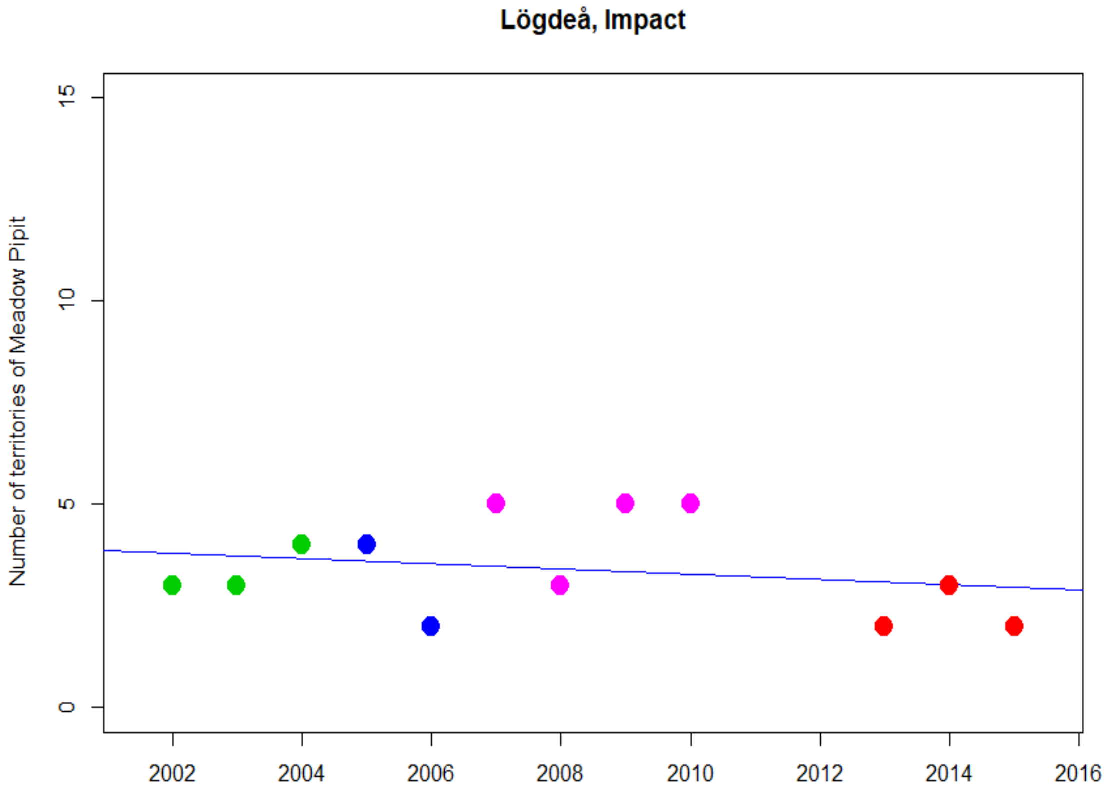
Lögdeå, Impact
Number of territories of Meadow Pipit
0
5
10
15
2002
2004
2006
2008
2010
2012
2014
2016

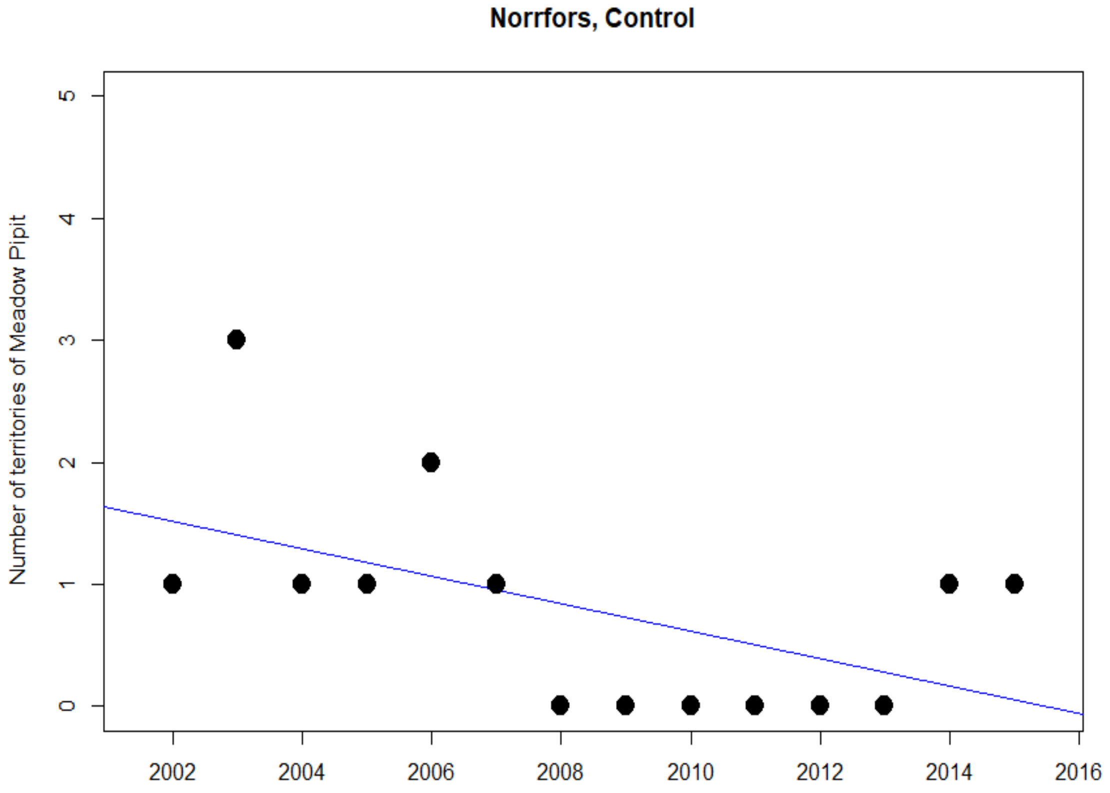
Norrfors, Control
Number of territories of Meadow Pipit
0
1
2
3
4
5
2002
2004
2006
2008
2010
2012
2014
2016

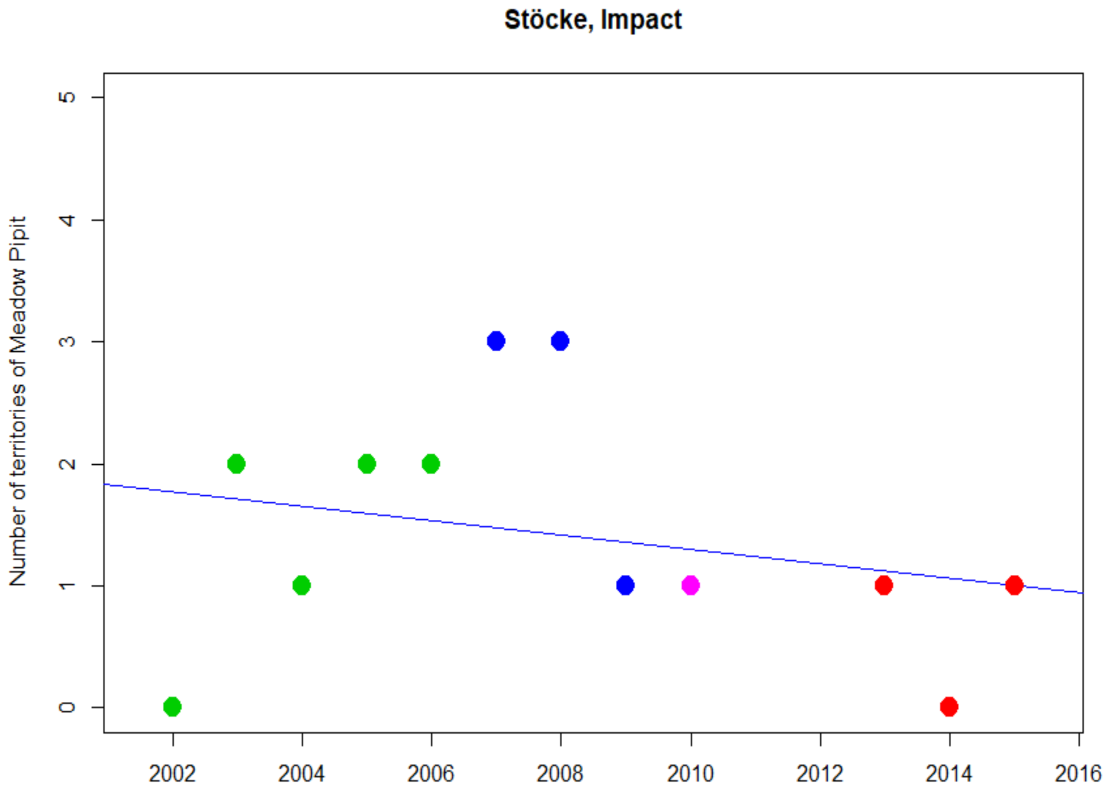
Stöcke, Impact
Number of territories of Meadow Pipit
0
1
2
3
4
5
2002
2004
2006
2008
2010
2012
2014
2016

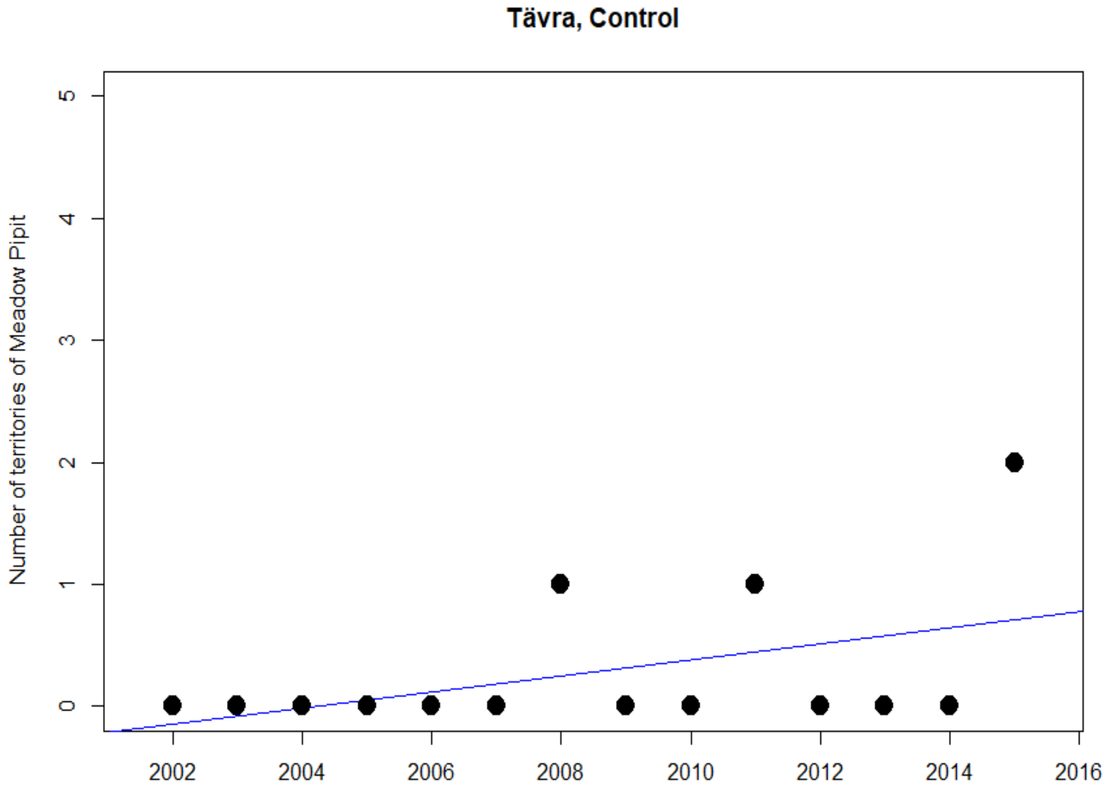
Tävra, Control
Number of territories of Meadow Pipit
0
1
2
3
4
5
2002
2004
2006
2008
2010
2012
2014
2016

### Northern lapwing

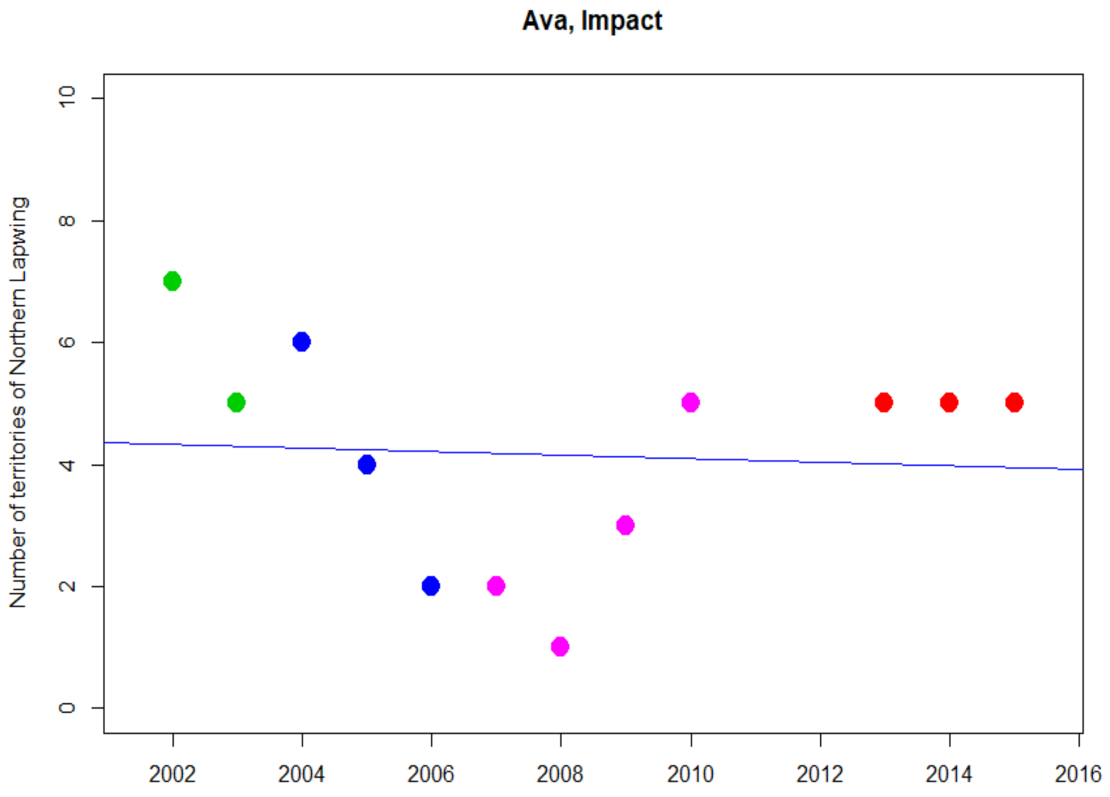


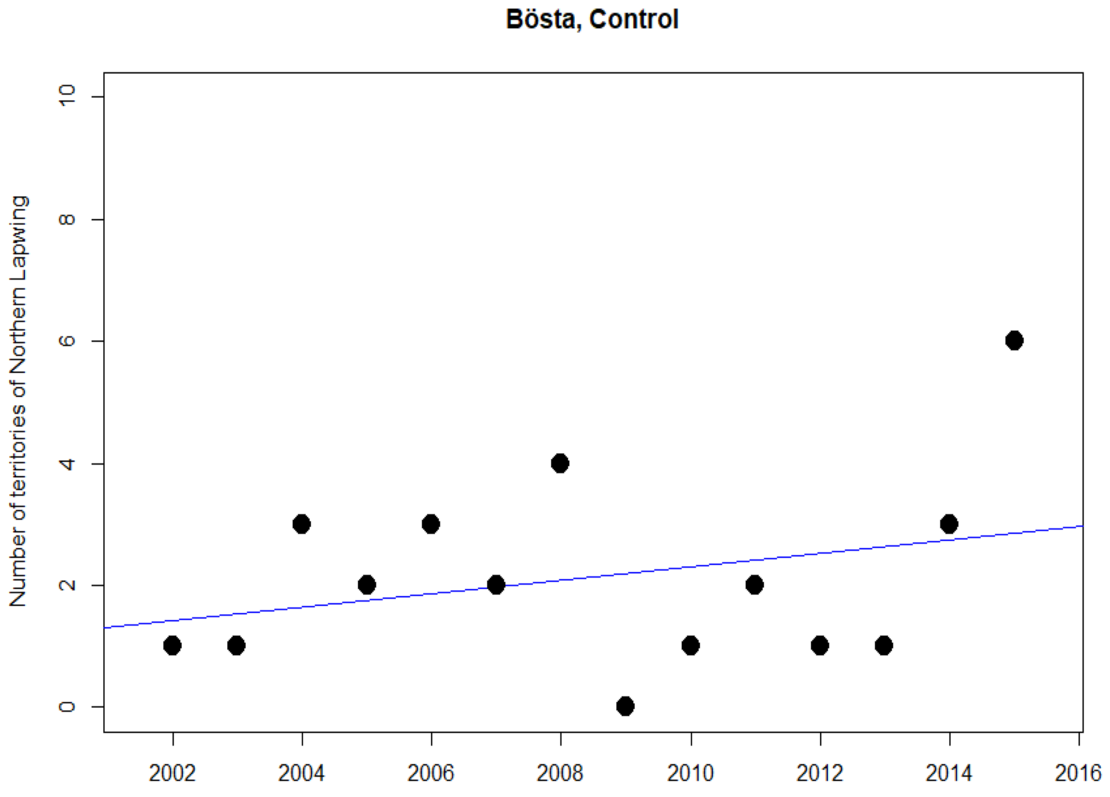

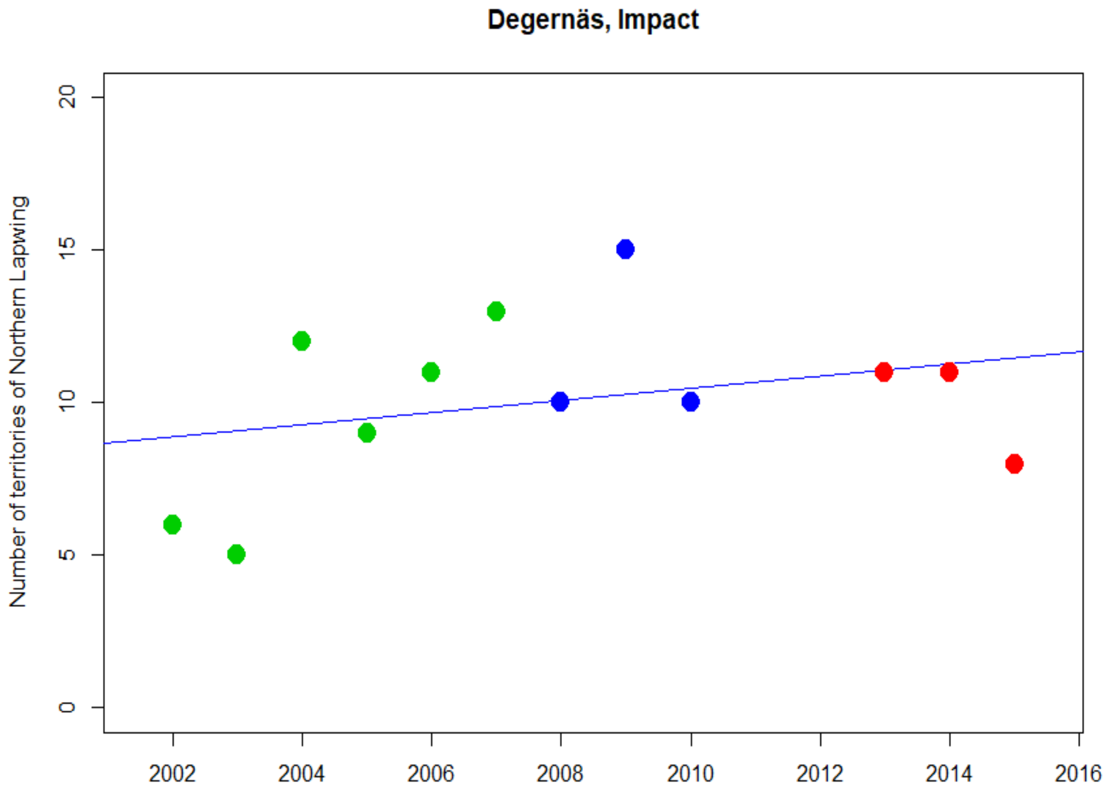
Degernäs, Impact
Number of territories of Northern Lapwing
0
5
10
15
20
2002
2004
2006
2008
2010
2012
2014
2016

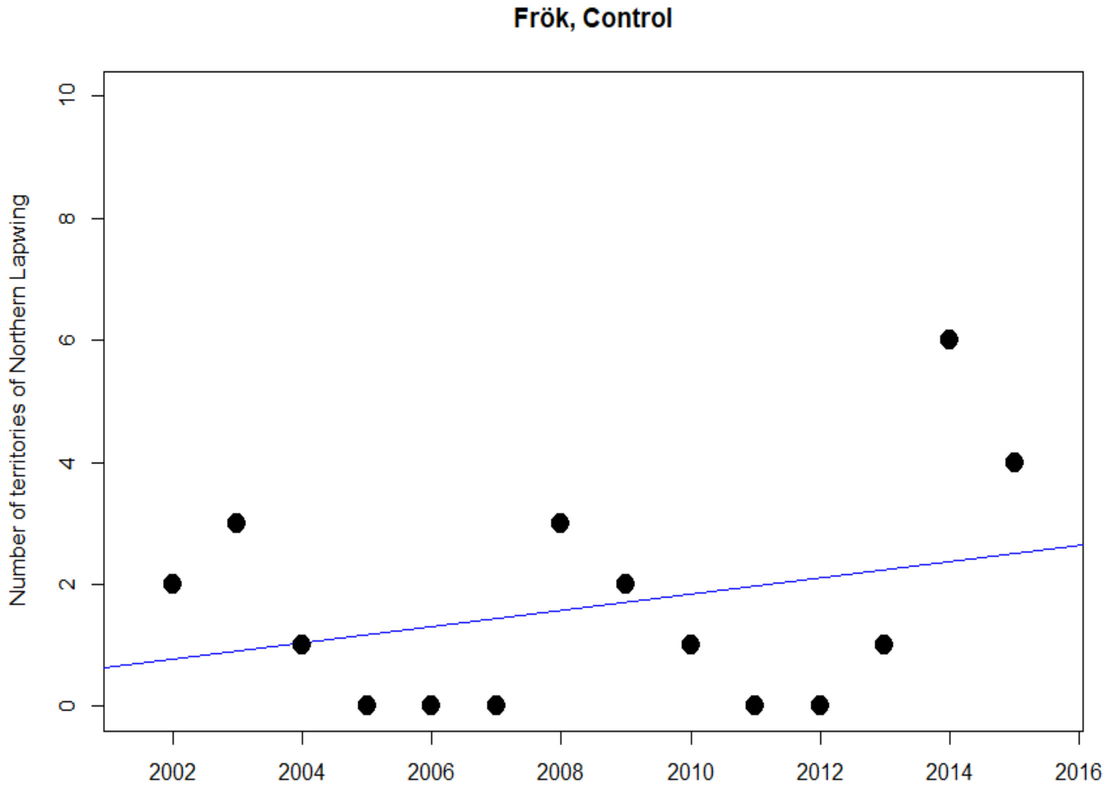
Frök, Control
Number of territories of Northern Lapwing
0
2
4
6
8
10
2002
2004
2006
2008
2010
2012
2014
2016

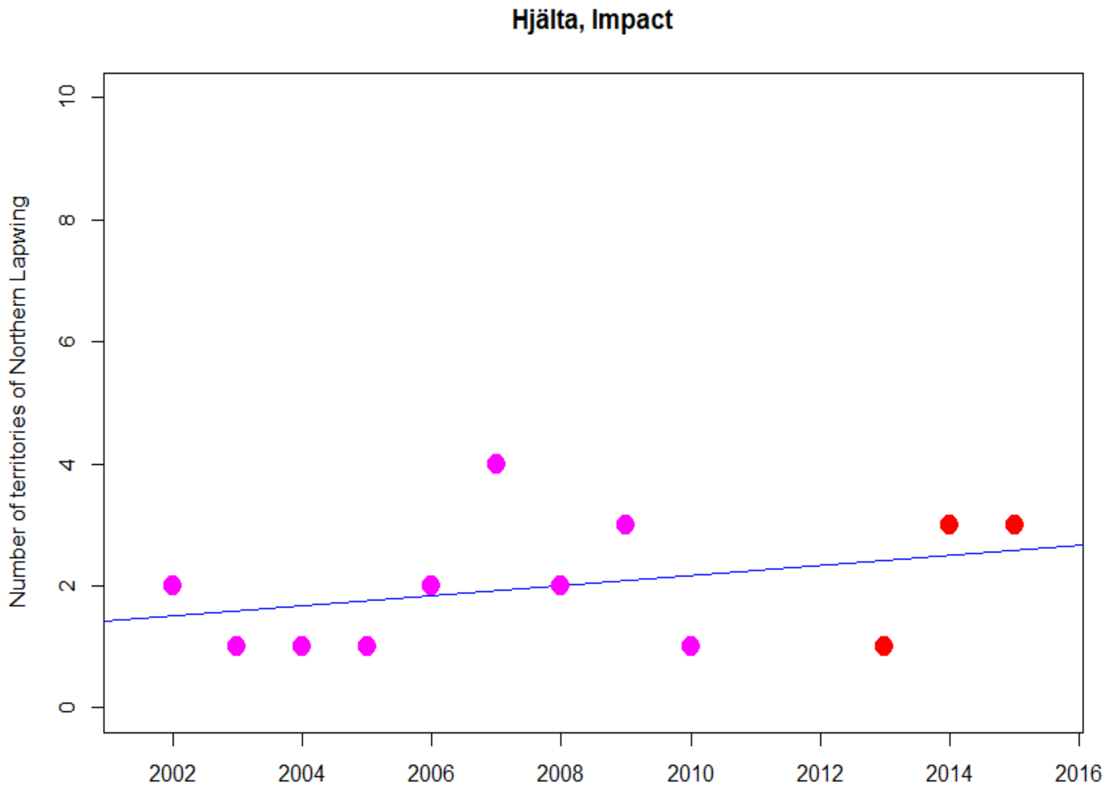
Hjälta, Impact
Number of territories of Northern Lapwing
0
2
4
6
8
10
2002
2004
2006
2008
2010
2012
2014
2016

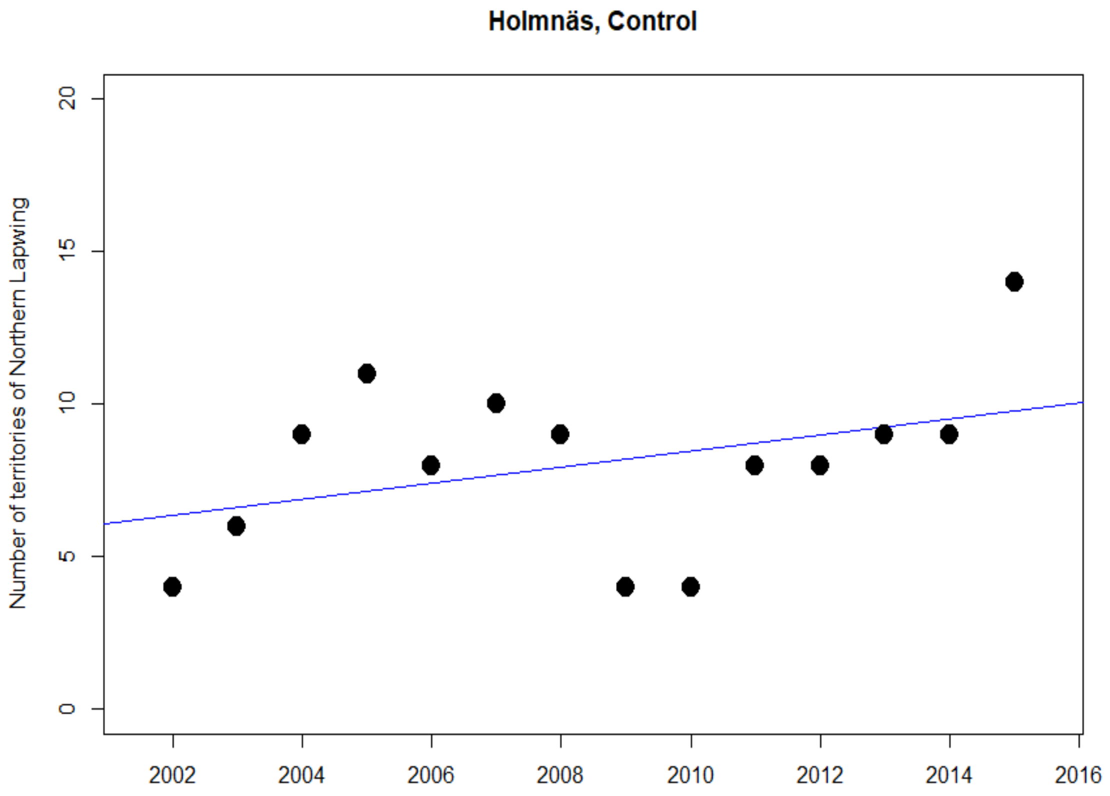
Holmnäs, Control
Number of territories of Northern Lapwing
0
5
10
15
20
2002
2004
2006
2008
2010
2012
2014
2016

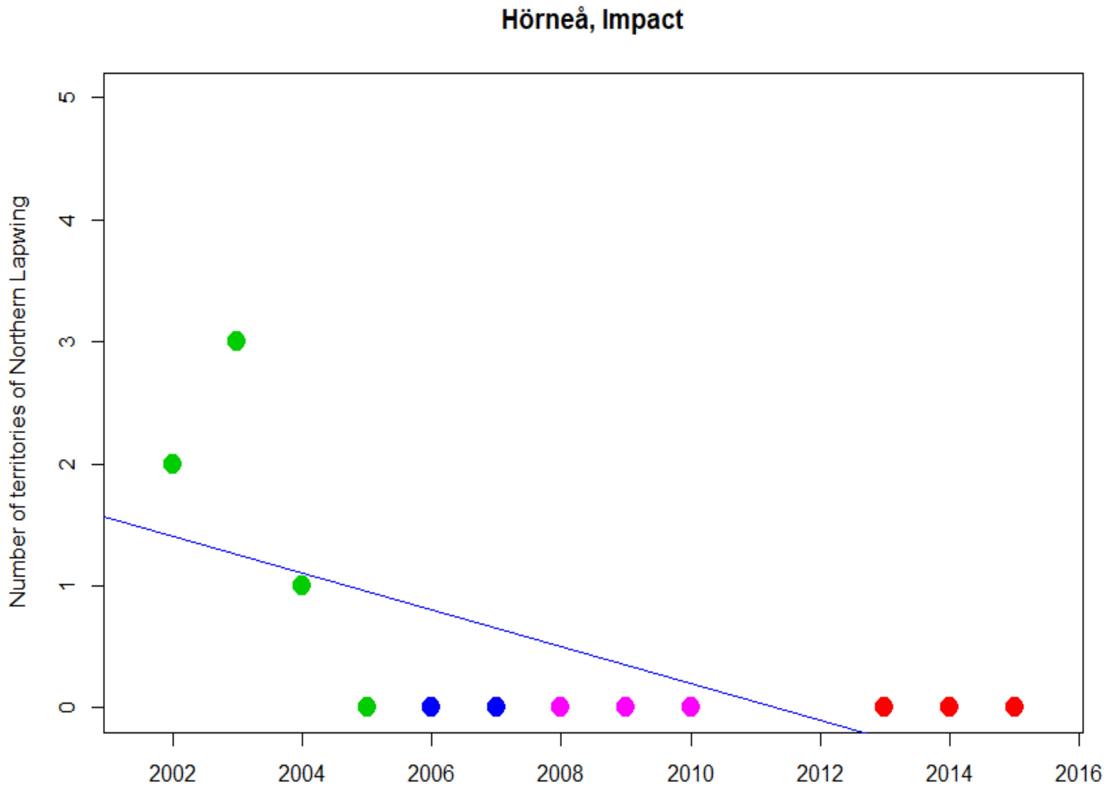
Hörneå, Impact
Number of territories of Northern Lapwing
0
1
2
3
4
5
2002
2004
2006
2008
2010
2012
2014
2016

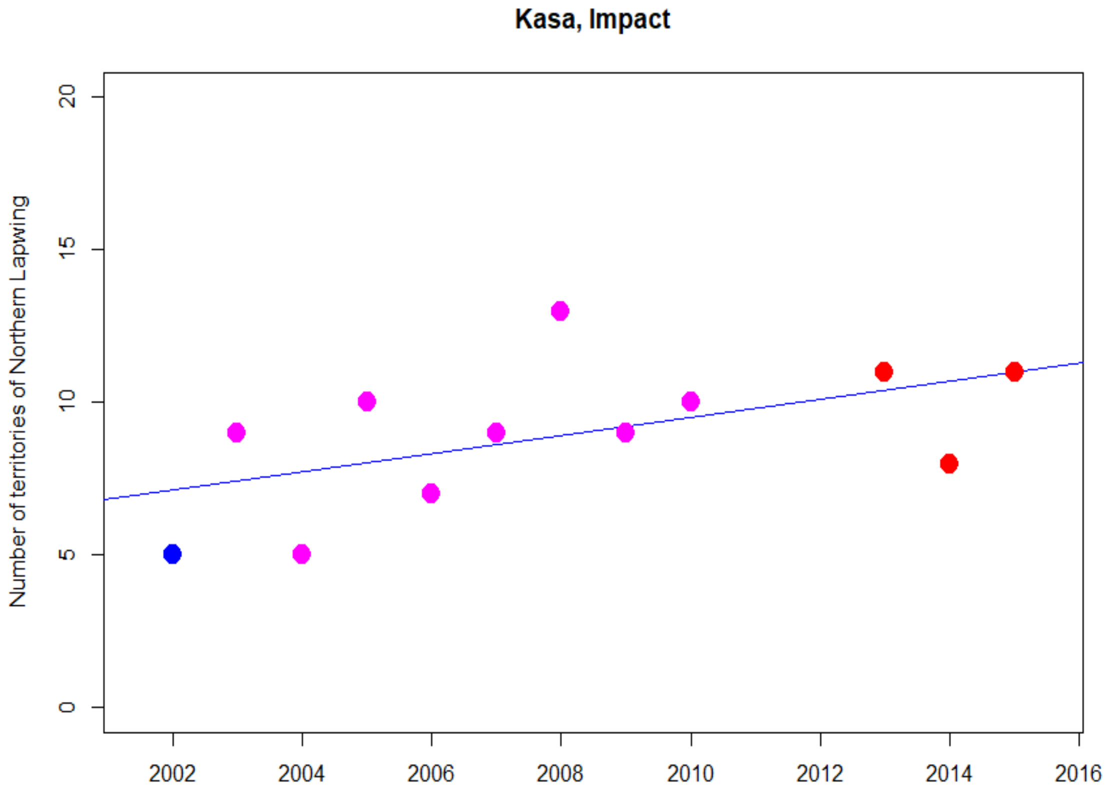
Kasa, Impact
Number of territories of Northern Lapwing
0
5
10
15
20
2002
2004
2006
2008
2010
2012
2014
2016

**Kornsjö, Impact**

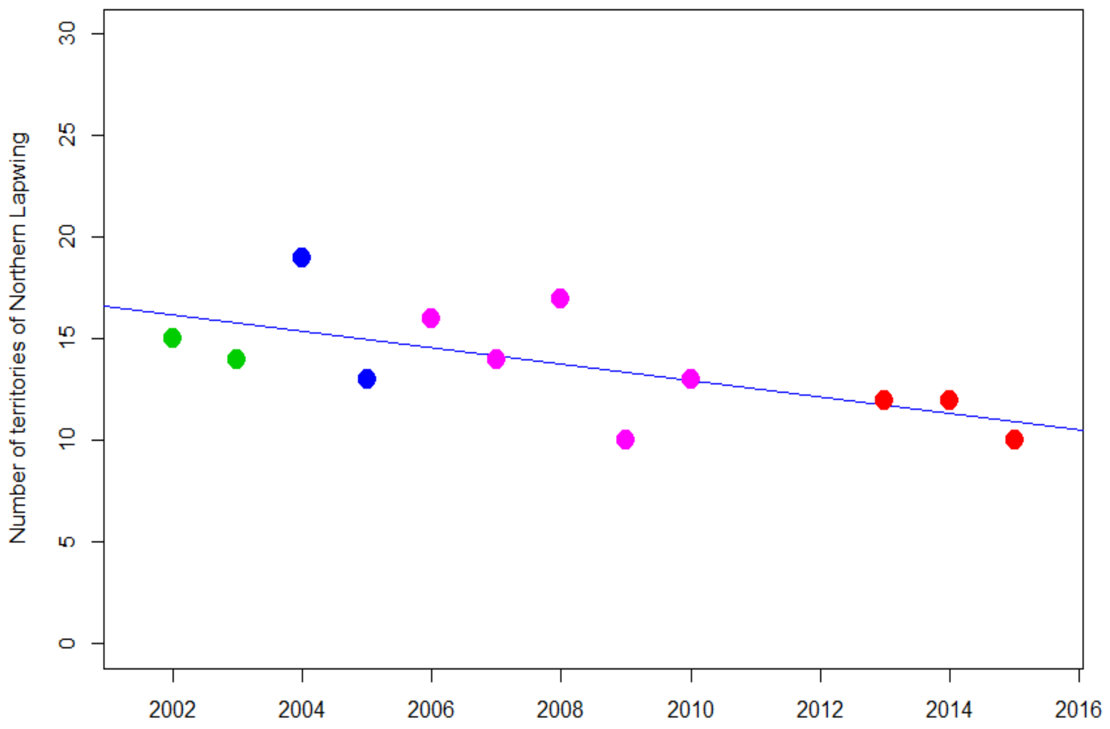


**Långed, Impact**

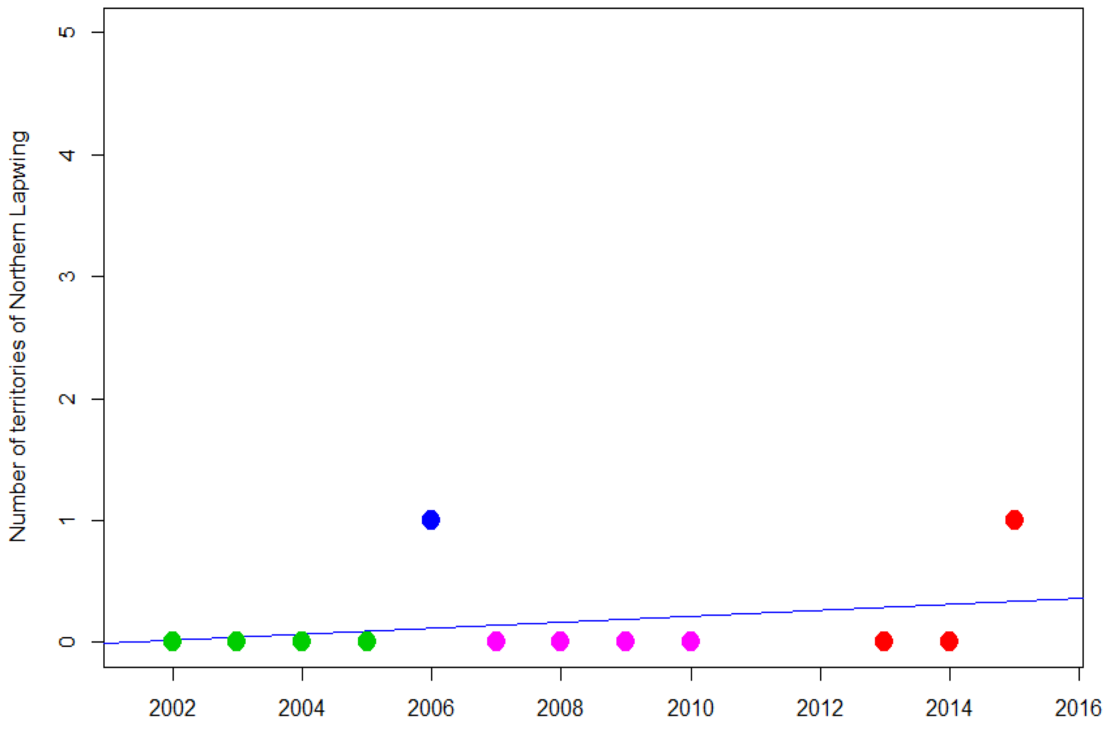

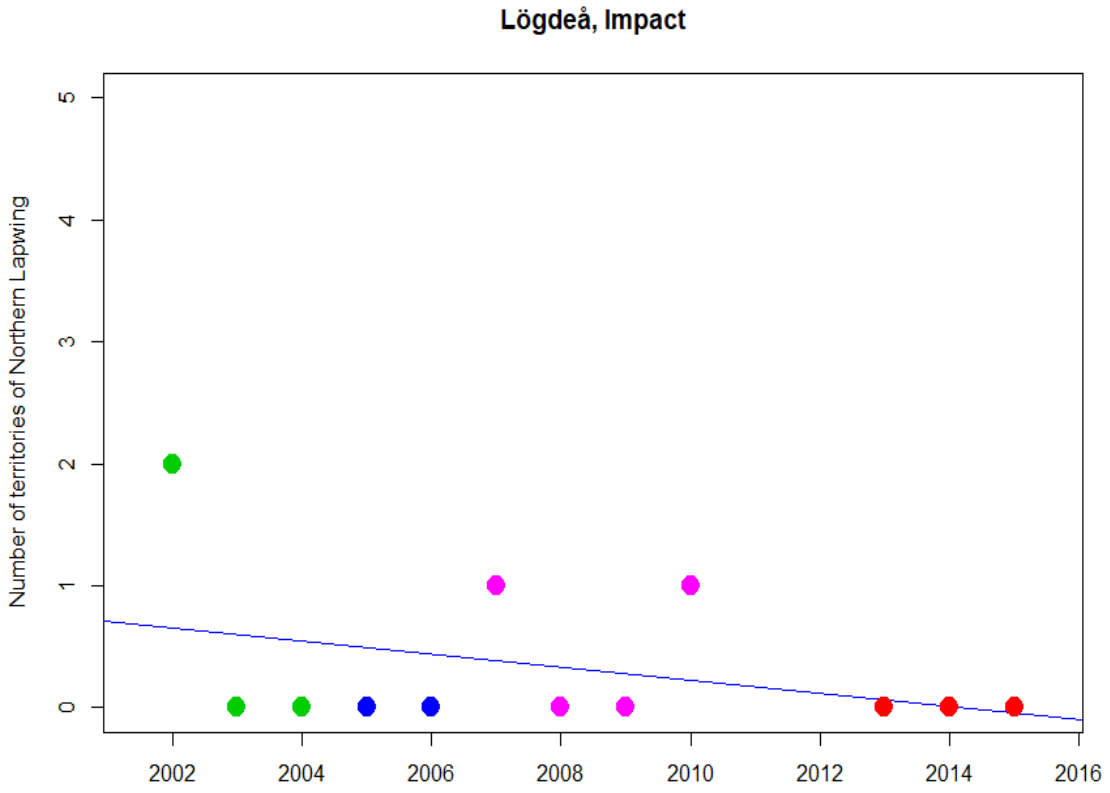
Lögdeå, Impact
Number of territories of Northern Lapwing
0
1
2
3
4
5
2002
2004
2006
2008
2010
2012
2014
2016

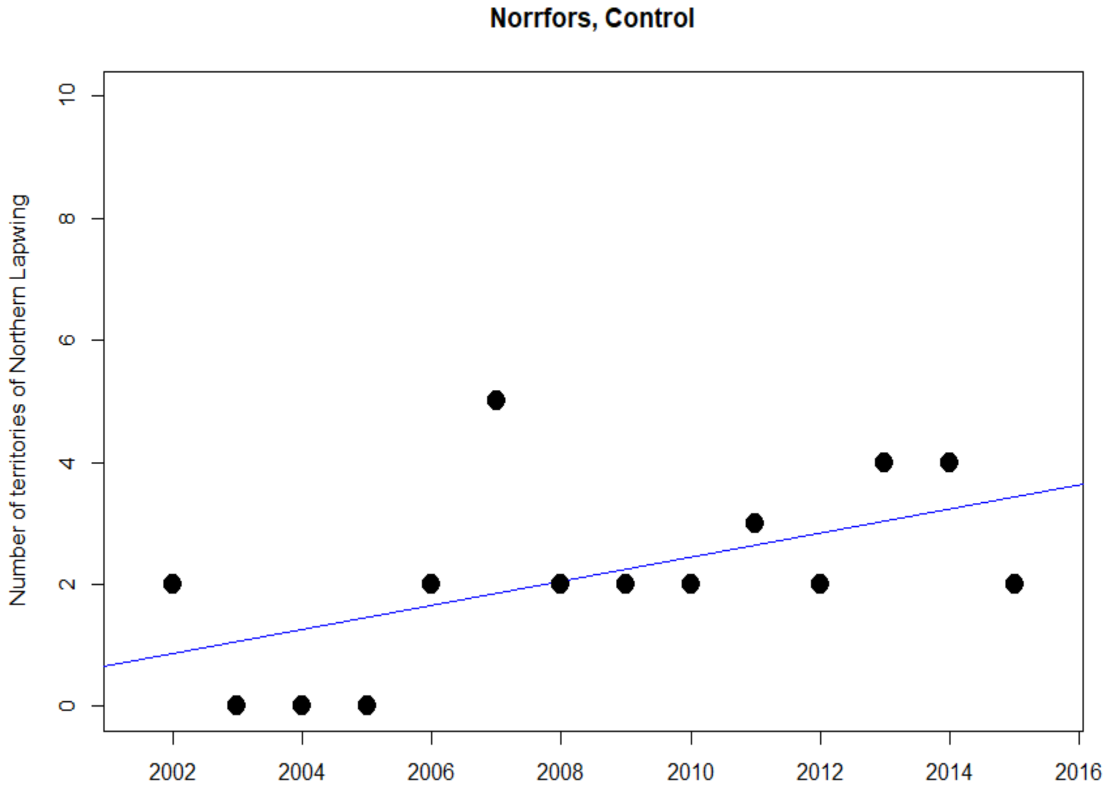
Norrfors, Control
Number of territories of Northern Lapwing
0
2
4
6
8
10
2002
2004
2006
2008
2010
2012
2014
2016

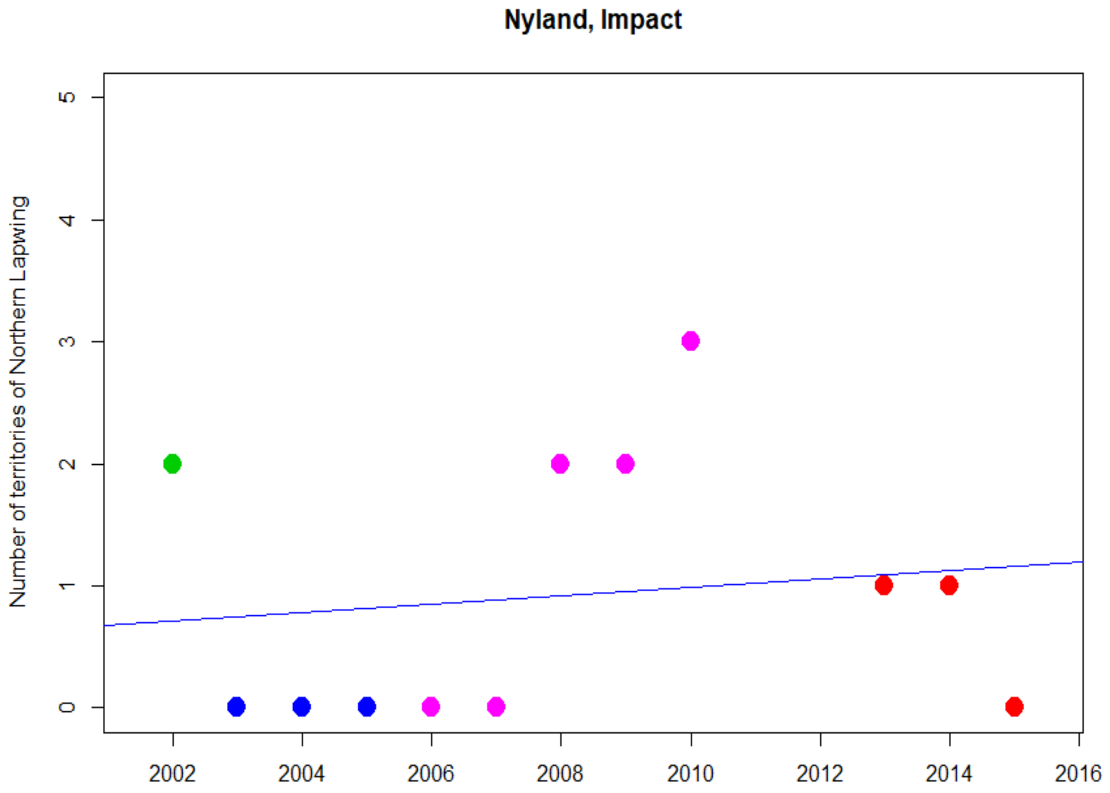
Nyland, Impact
Number of territories of Northern Lapwing
0
1
2
3
4
5
2002
2004
2006
2008
2010
2012
2014
2016

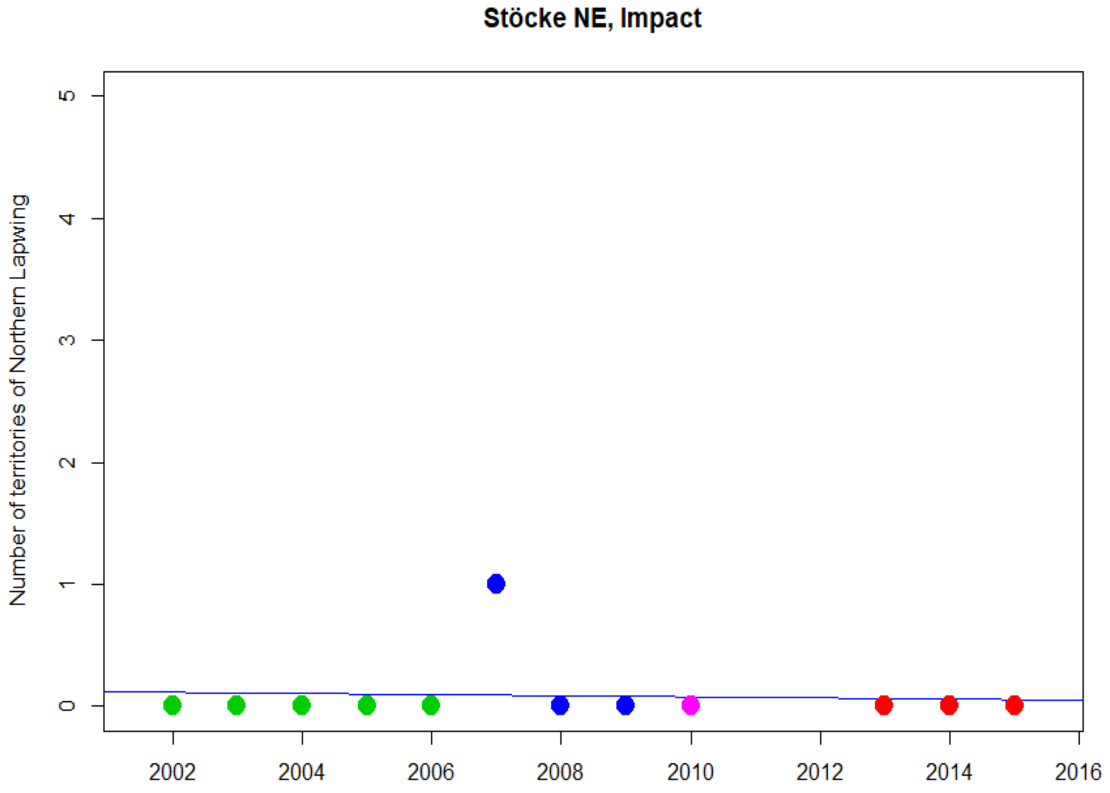
Stöcke NE, Impact
Number of territories of Northern Lapwing
0
1
2
3
4
5
2002
2004
2006
2008
2010
2012
2014
2016

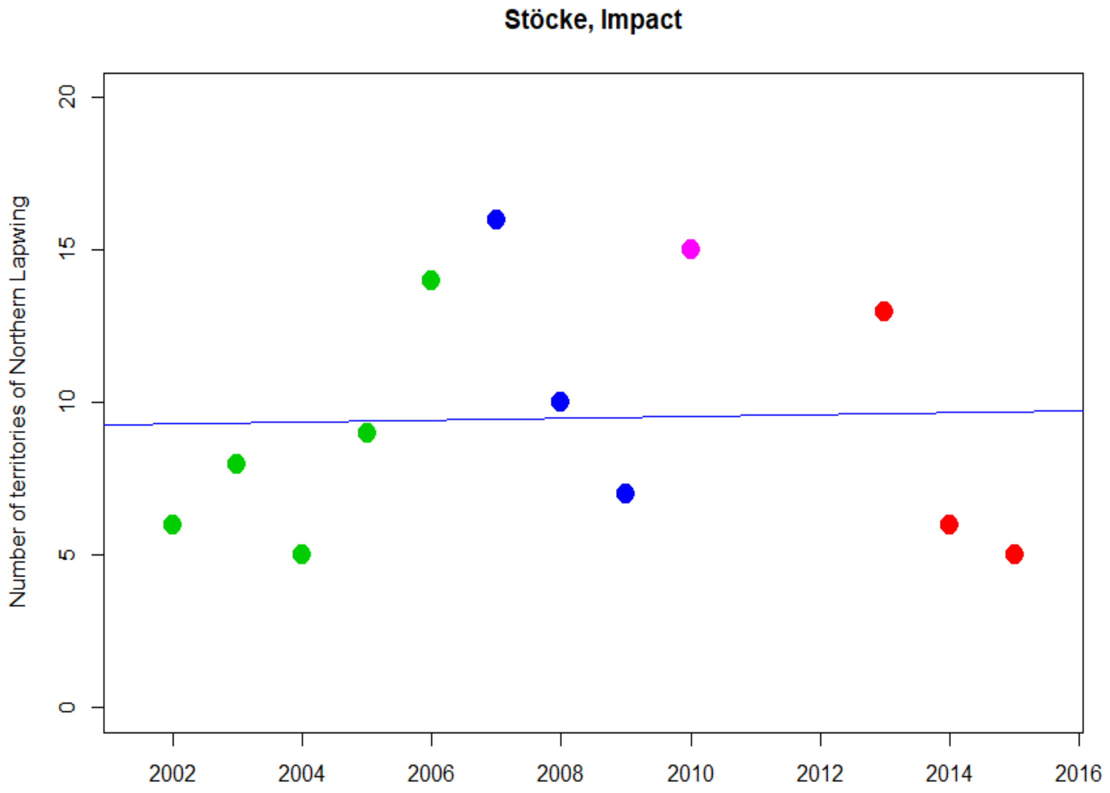
Stöcke, Impact
Number of territories of Northern Lapwing
0
5
10
15
20
2002
2004
2006
2008
2010
2012
2014
2016

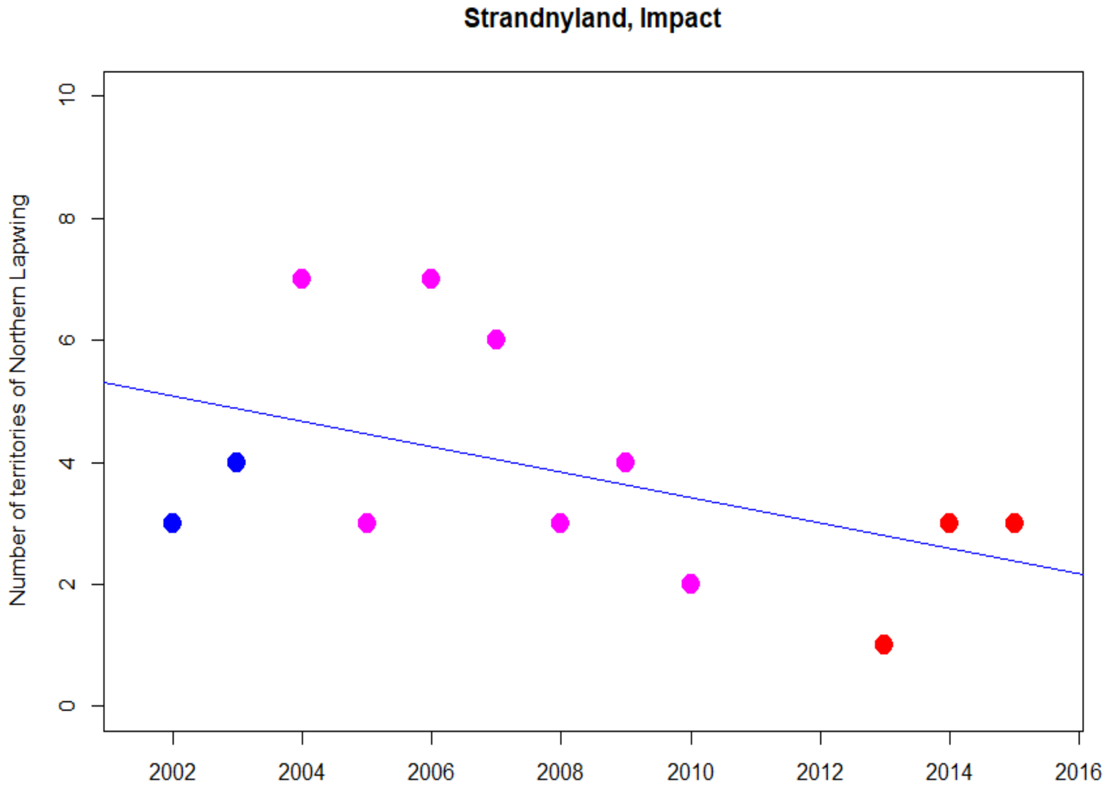
Strandnyland, Impact
Number of territories of Northern Lapwing
0
2
4
6
8
10
2002
2004
2006
2008
2010
2012
2014
2016

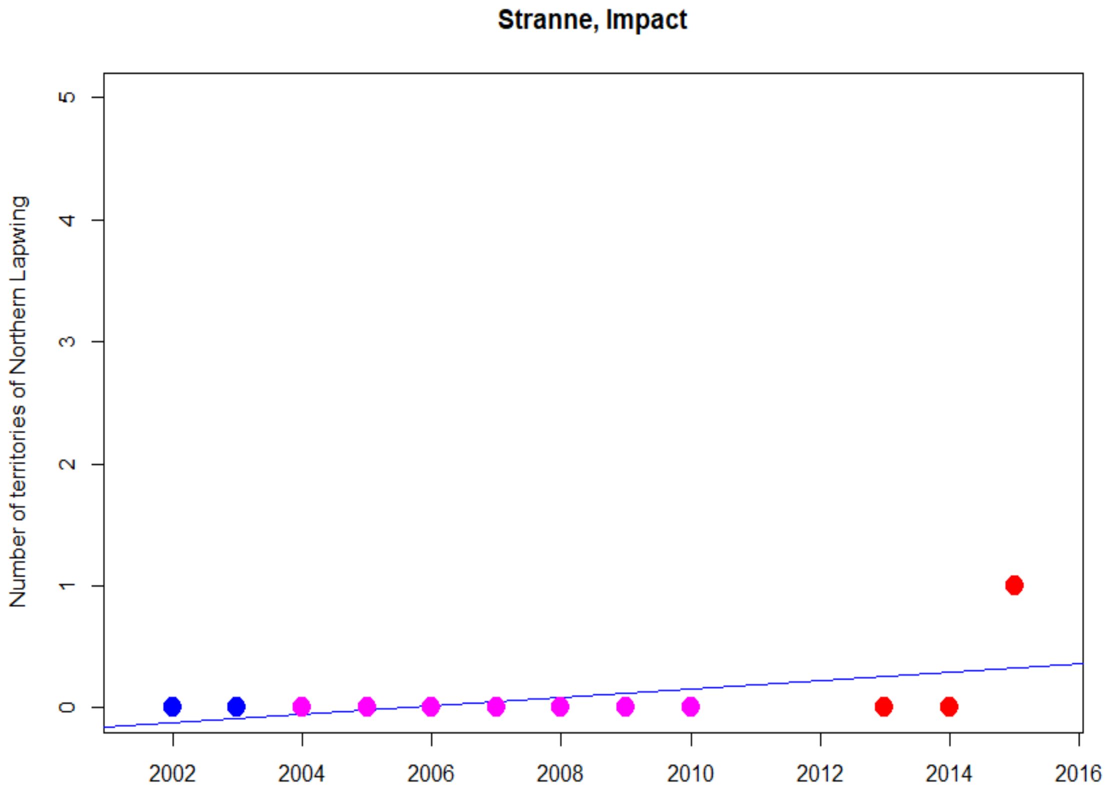
Stranne, Impact
Number of territories of Northern Lapwing
0
1
2
3
4
5
2002
2004
2006
2008
2010
2012
2014
2016

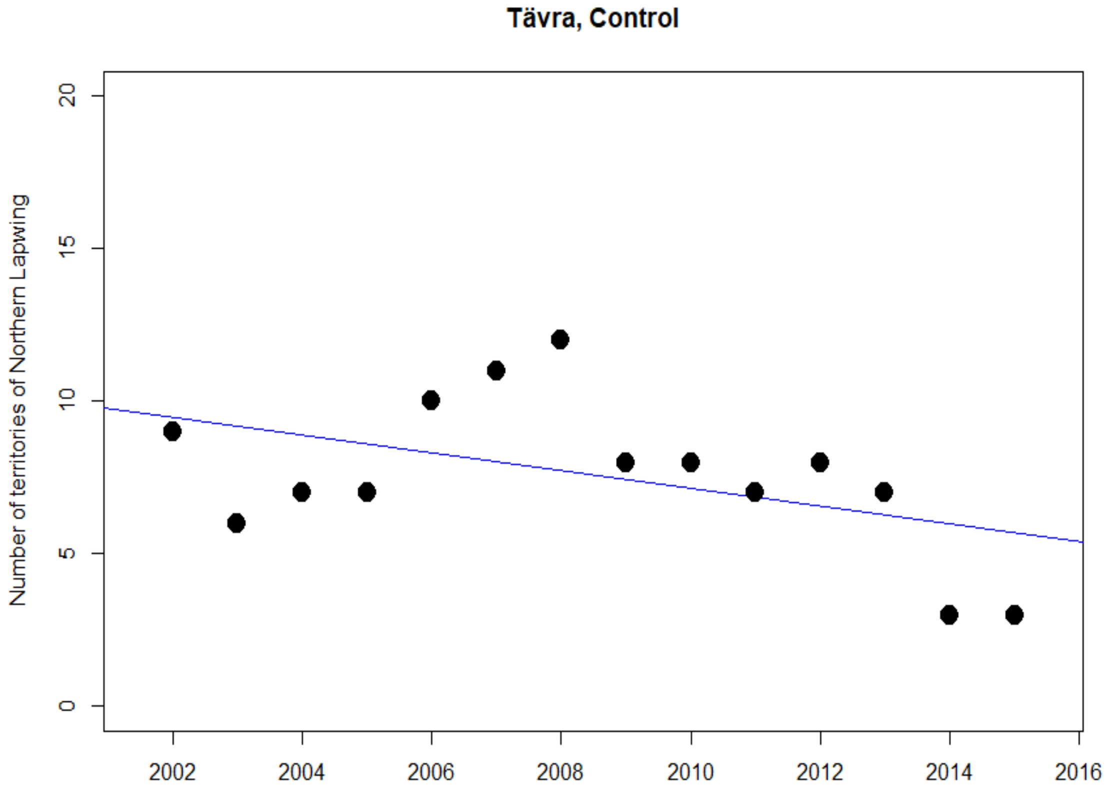
Tävra, Control
Number of territories of Northern Lapwing
0
5
10
15
20
2002
2004
2006
2008
2010
2012
2014
2016

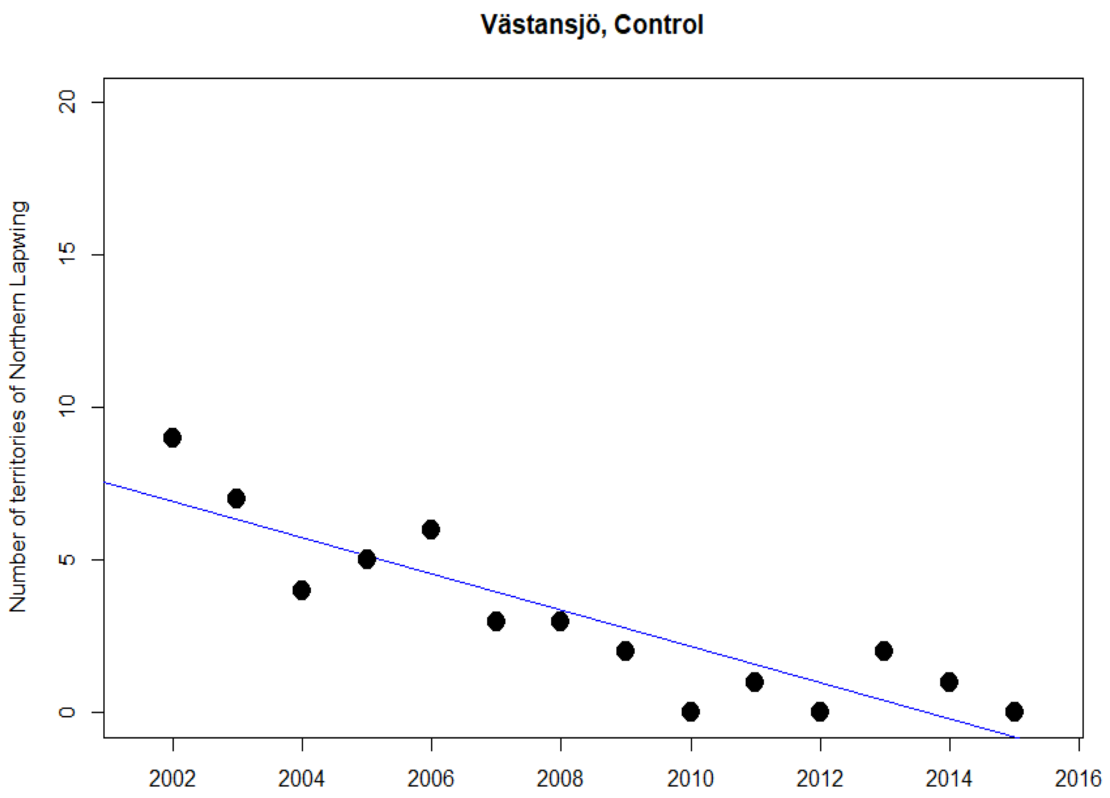
Västansjö, Control
Number of territories of Northern Lapwing
0
5
10
15
20
2002
2004
2006
2008
2010
2012
2014
2016

### Ortolan Bunting

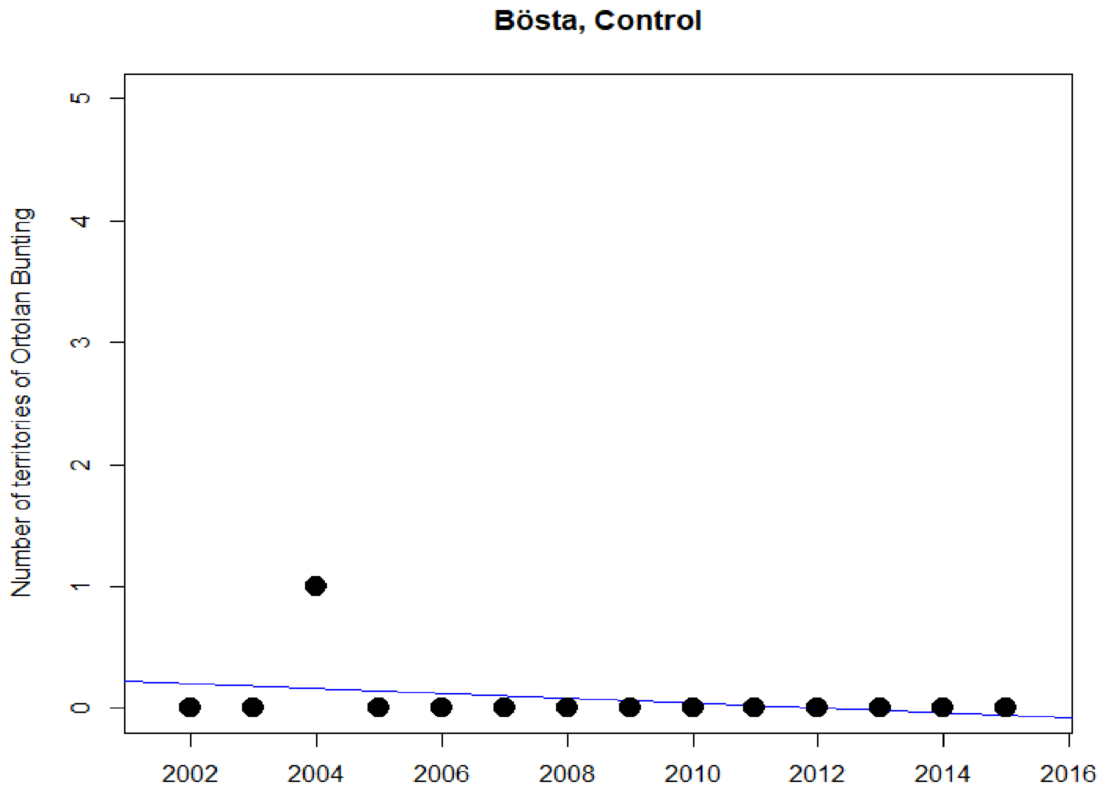


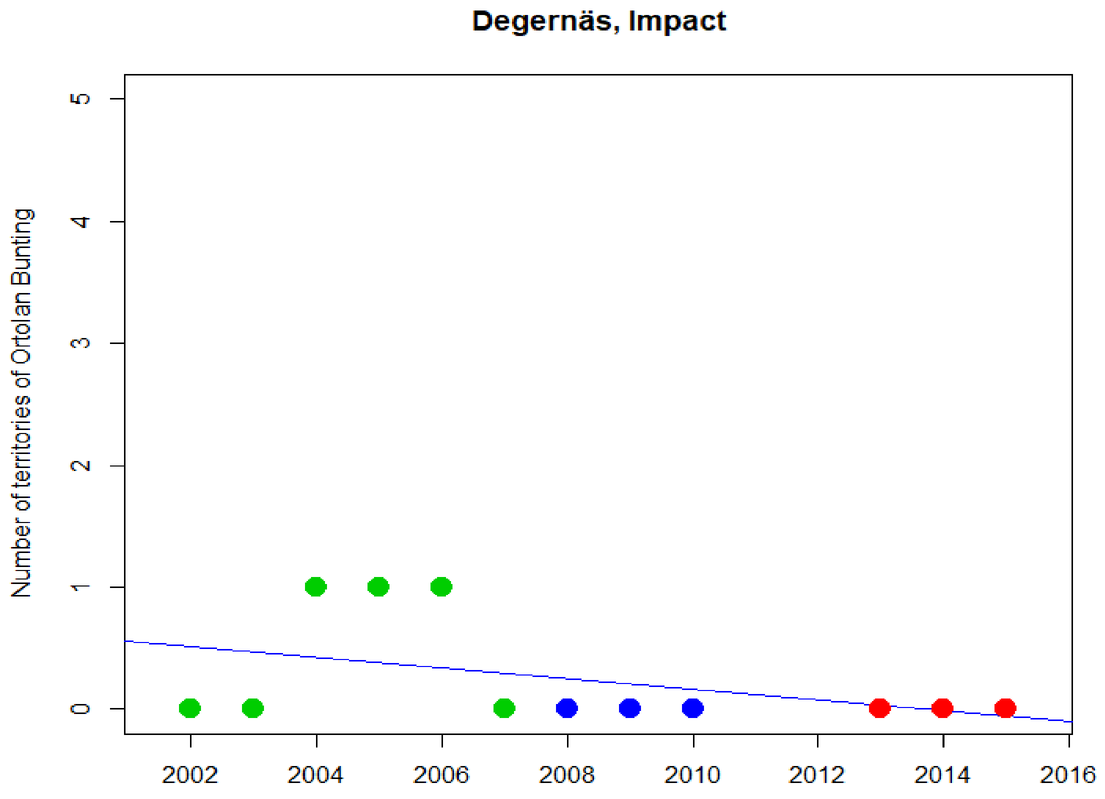

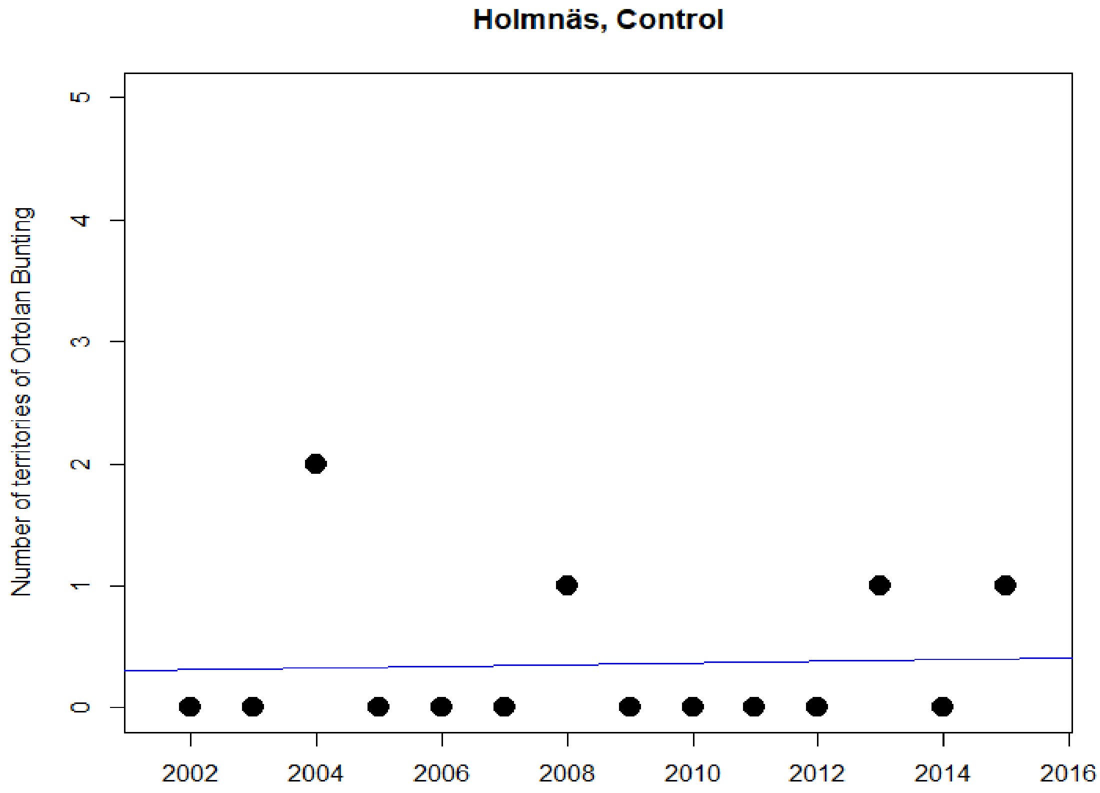
Holmnäs, Control
Number of territories of Ortolan Bunting
0
1
2
3
4
5
2002
2004
2006
2008
2010
2012
2014
2016

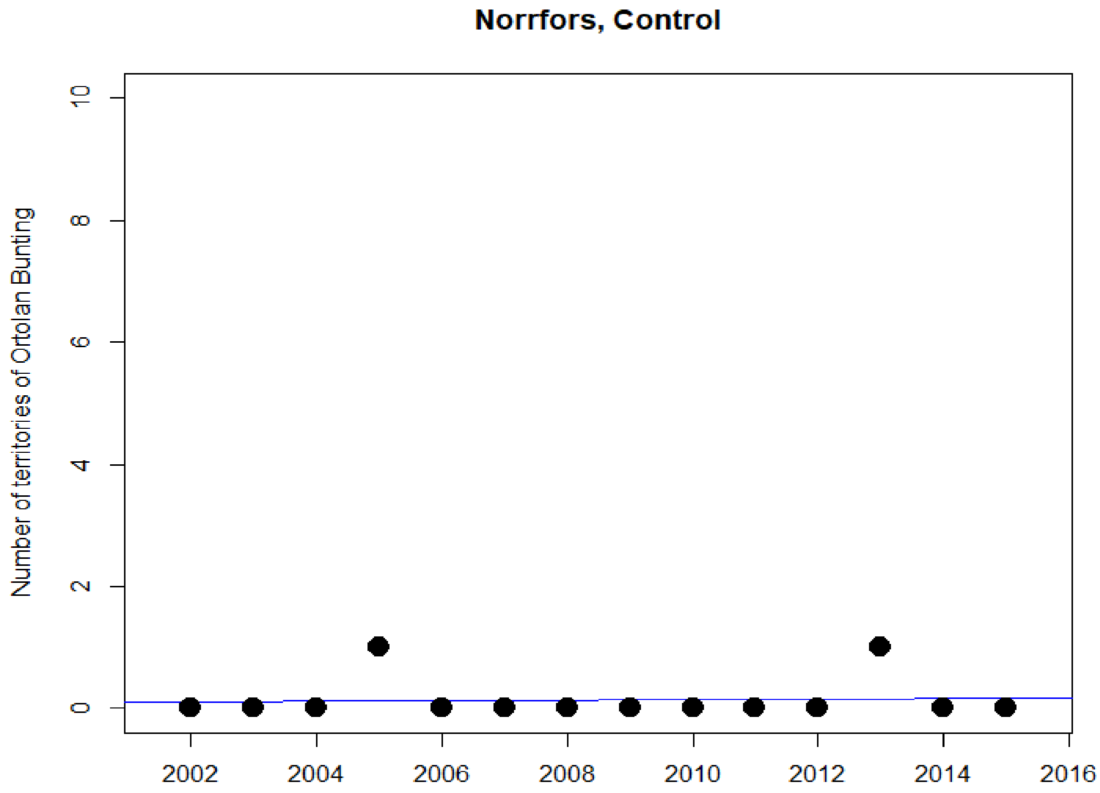
Norrfors, Control
Number of territories of Ortolan Bunting
0
2
4
6
8
10
2002
2004
2006
2008
2010
2012
2014
2016

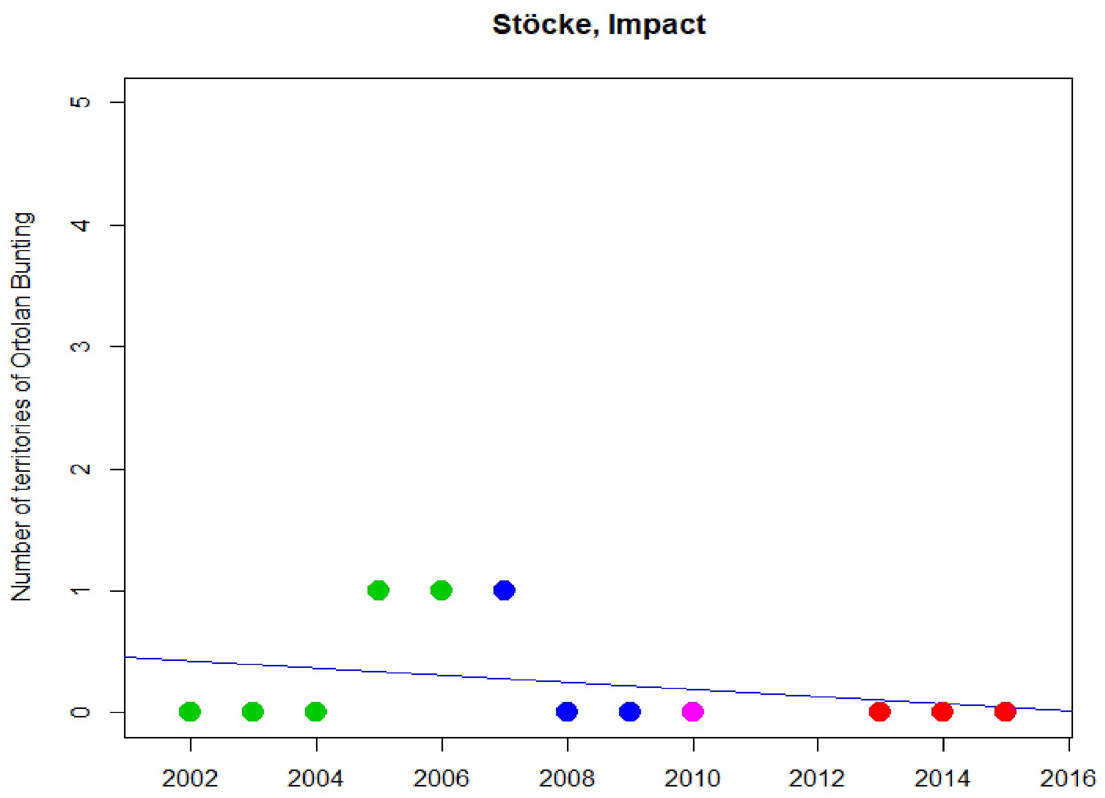

Stöcke, Impact
Number of territories of Ortolan Bunting
0
1
2
3
4
5
2002
2004
2006
2008
2010
2012
2014
2016

### Red-backed Shrike

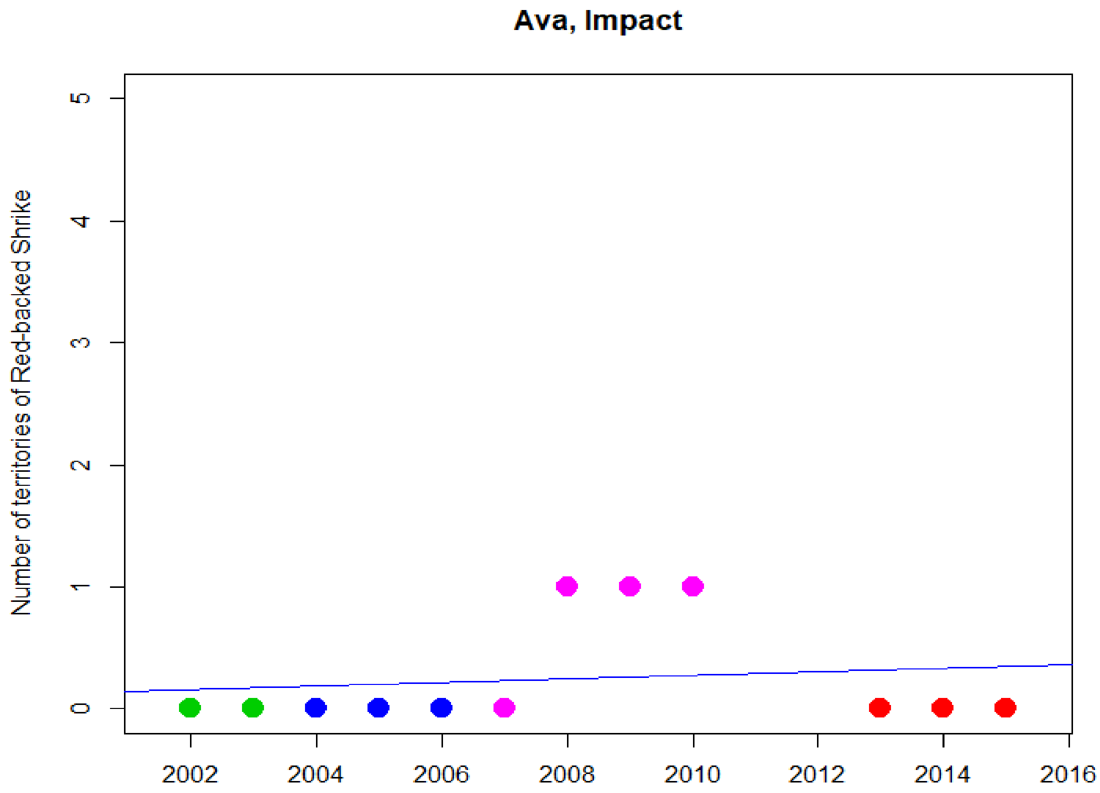


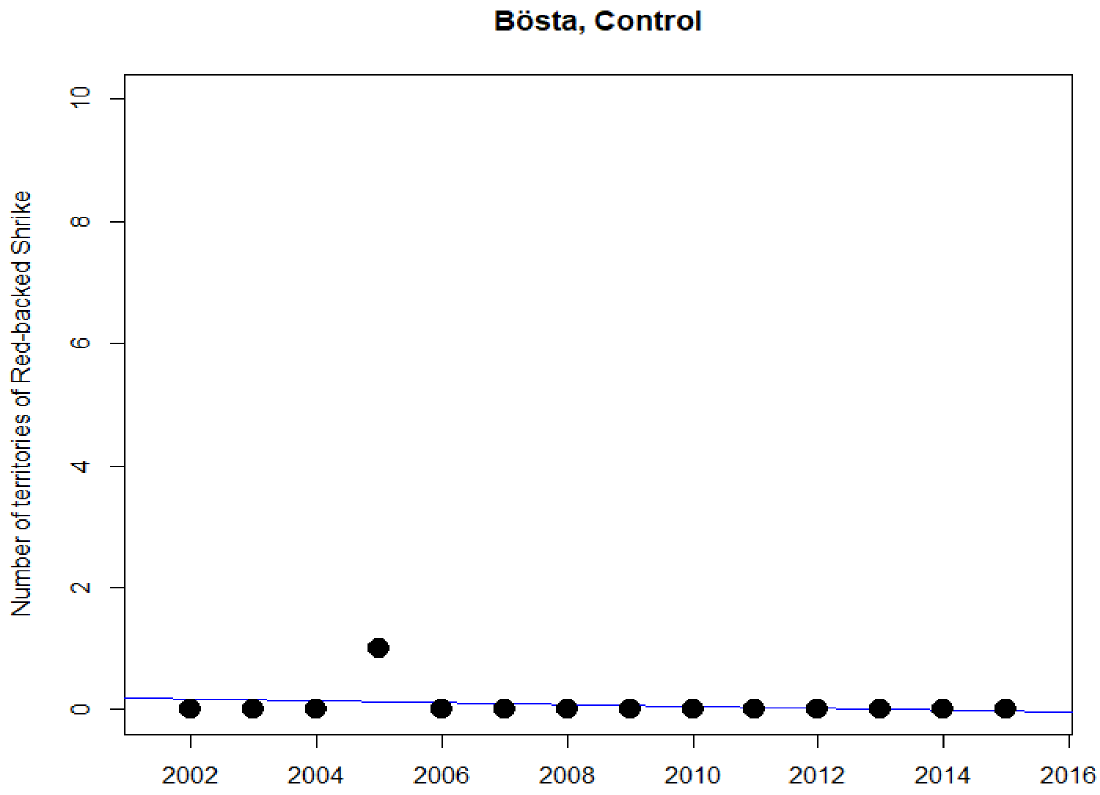

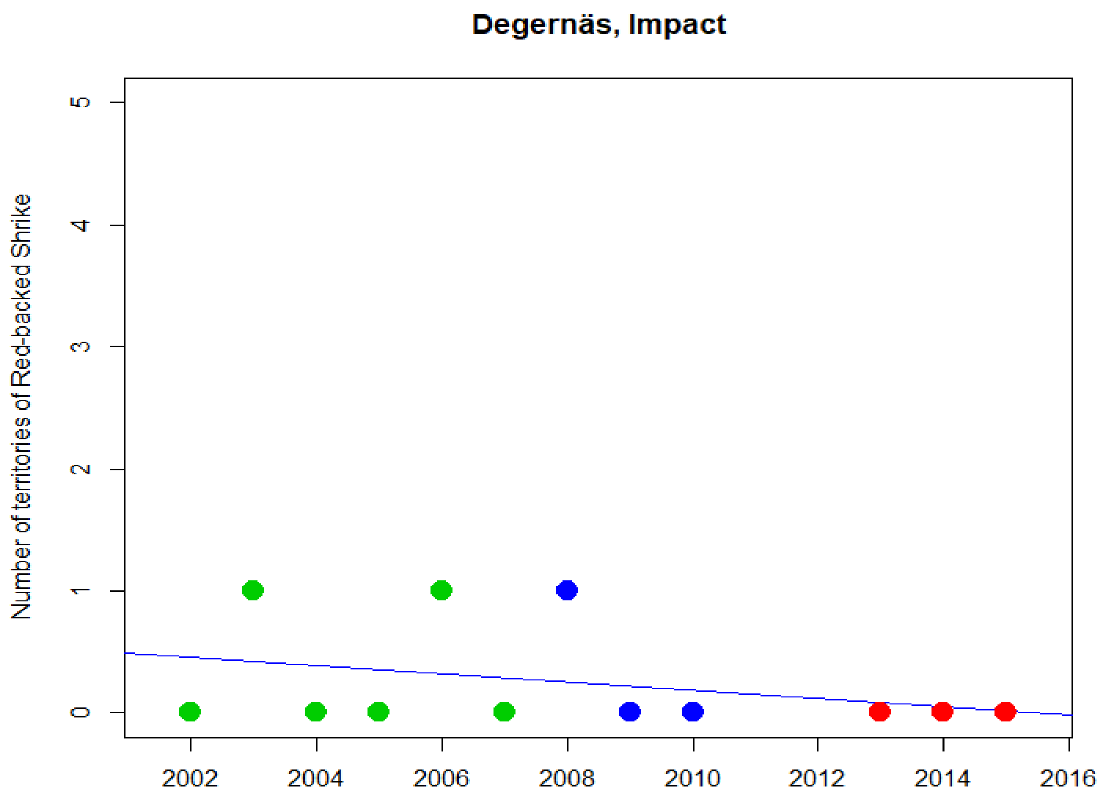
Degernäs, Impact
Number of territories of Red-backed Shrike
0
1
2
3
4
5
2002
2004
2006
2008
2010
2012
2014
2016

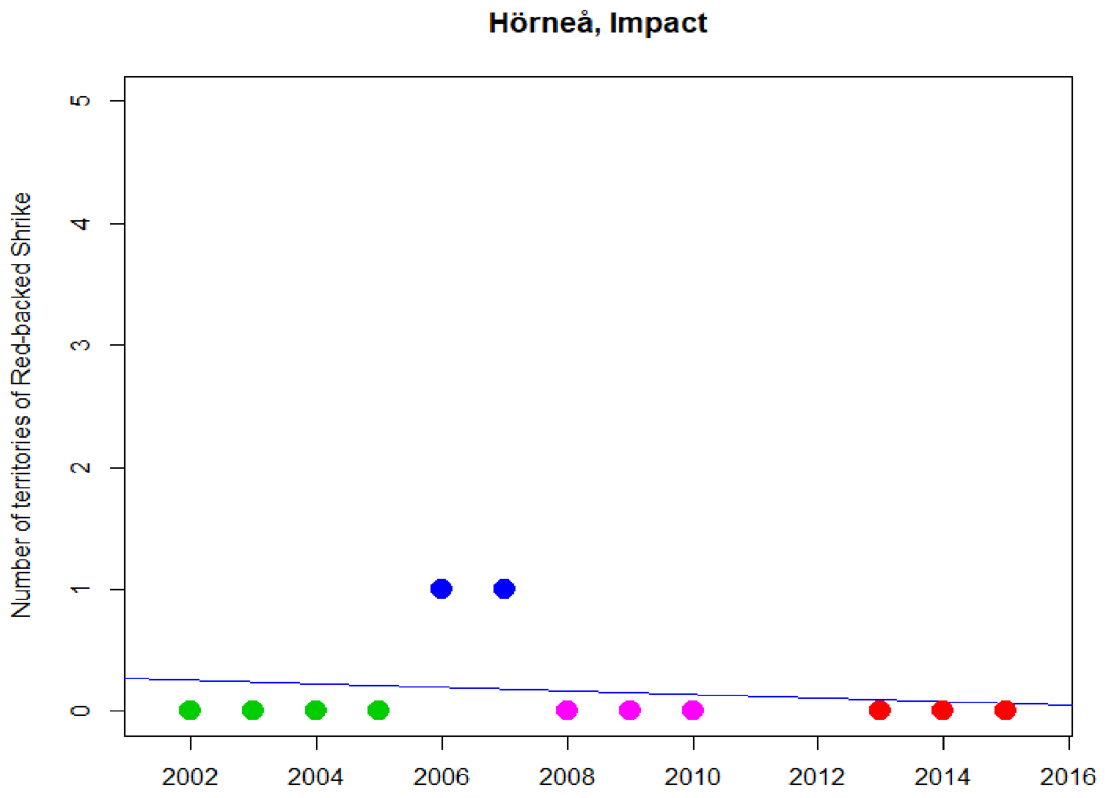
Hörneå, Impact
Number of territories of Red-backed Shrike
0
1
2
3
4
5
2002
2004
2006
2008
2010
2012
2014
2016

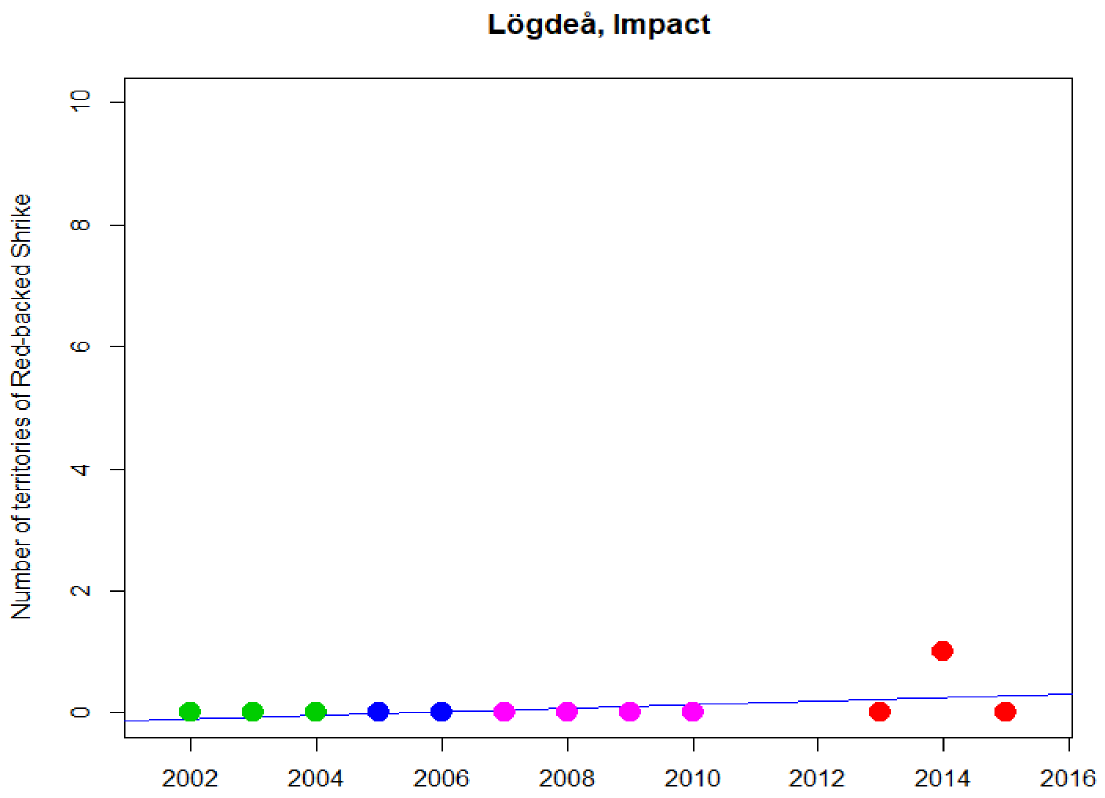
Lögdeå, Impact
Number of territories of Red-backed Shrike
0
2
4
6
8
10
2002
2004
2006
2008
2010
2012
2014
2016

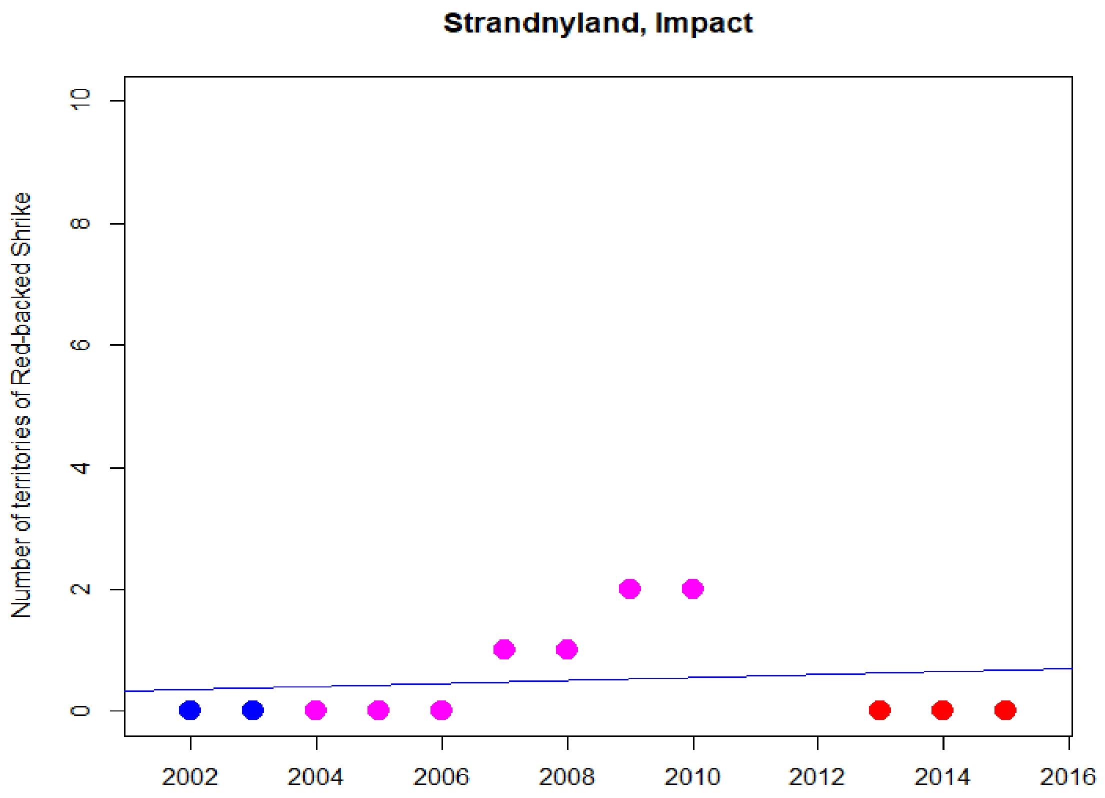
Strandnyland, Impact
Number of territories of Red-backed Shrike
0
2
4
6
8
10
2002
2004
2006
2008
2010
2012
2014
2016

### Western Yellow Wagtail

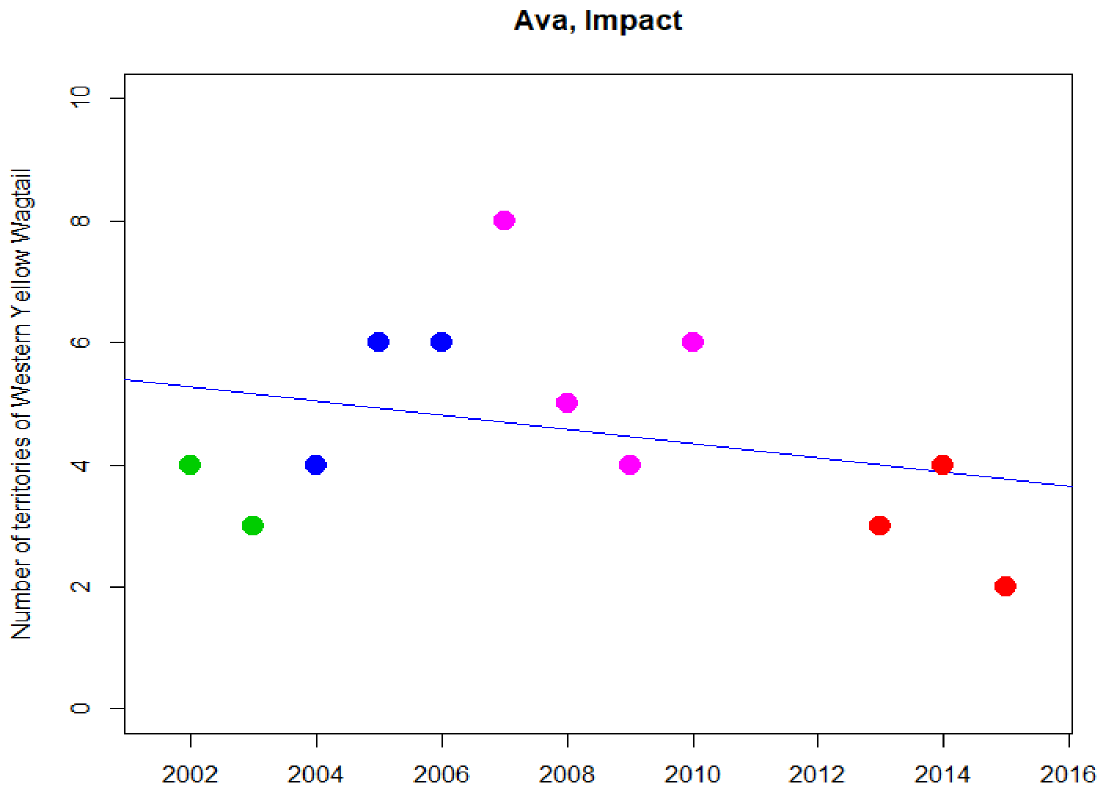


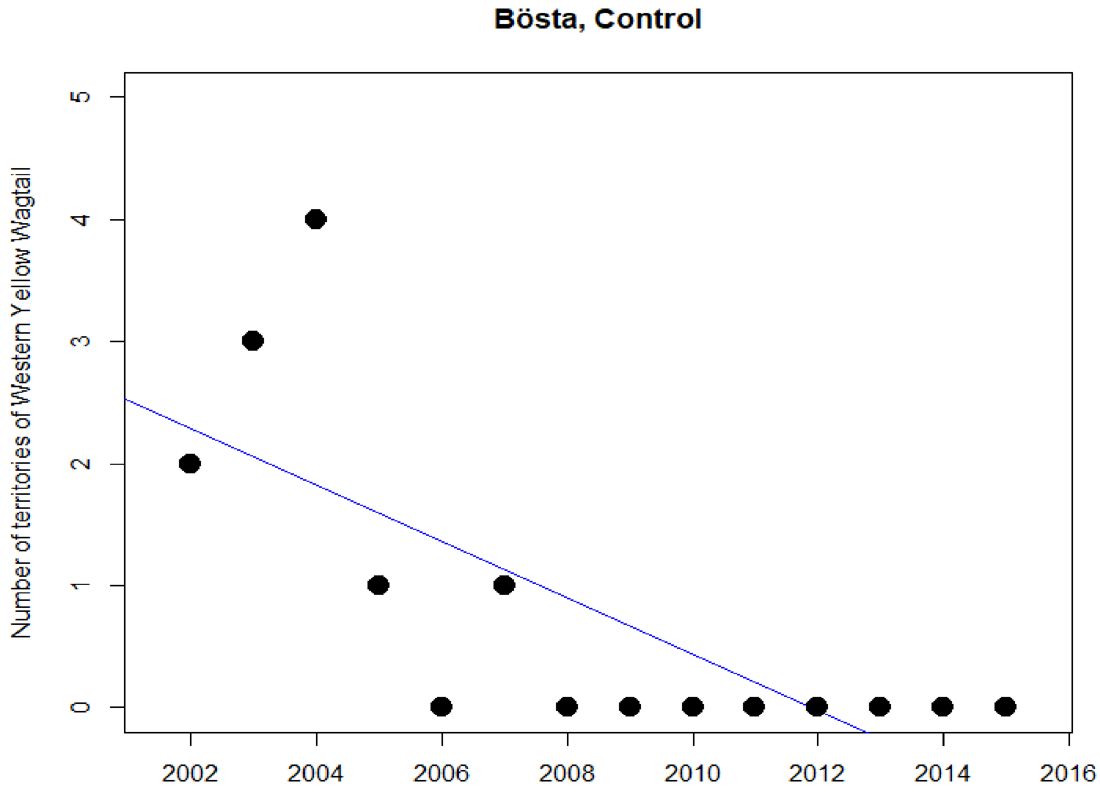

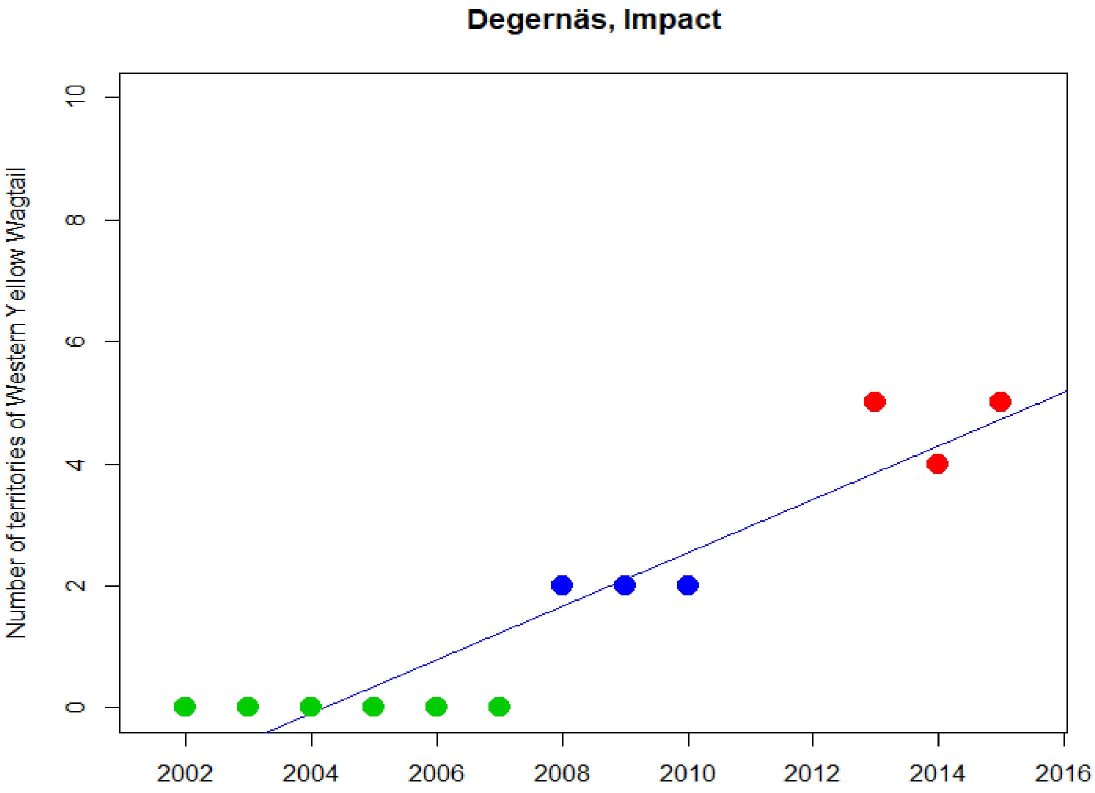
Degernäs, Impact
Number of territories of Western Yellow Wagtail
0
2
4
6
8
10
2002
2004
2006
2008
2010
2012
2014
2016

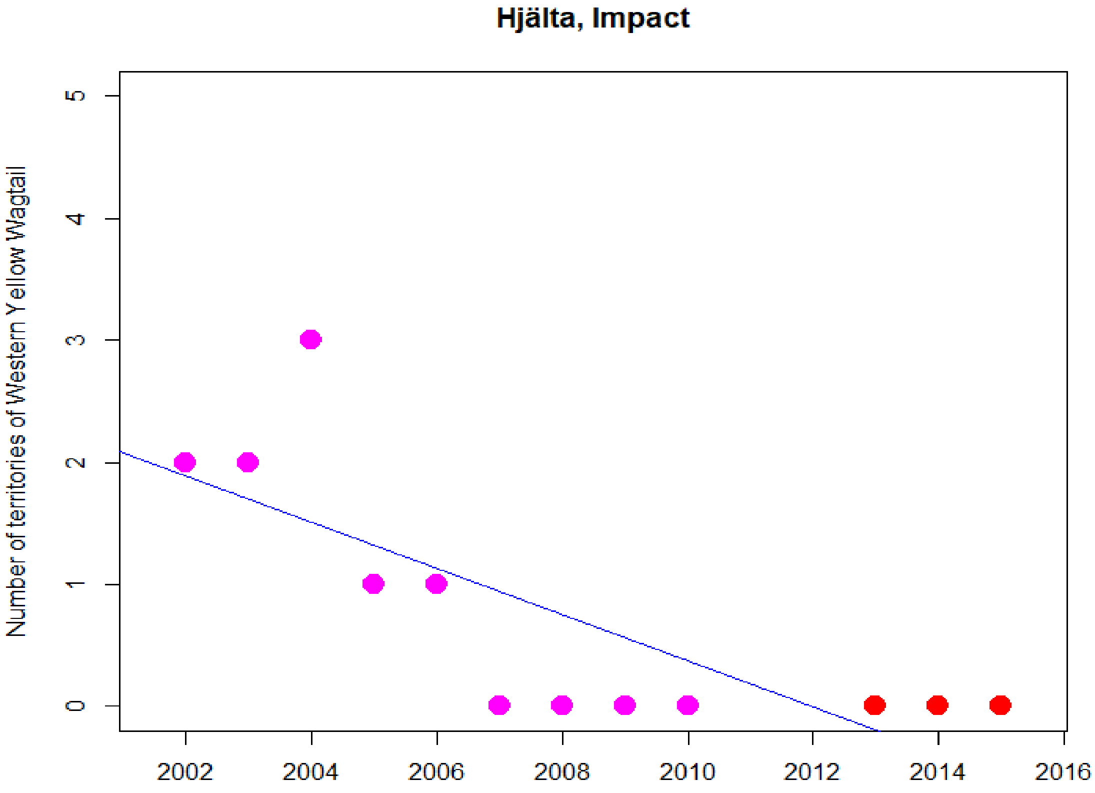
Hjälta, Impact
Number of territories of Western Yellow Wagtail
0
1
2
3
4
5
2002
2004
2006
2008
2010
2012
2014
2016

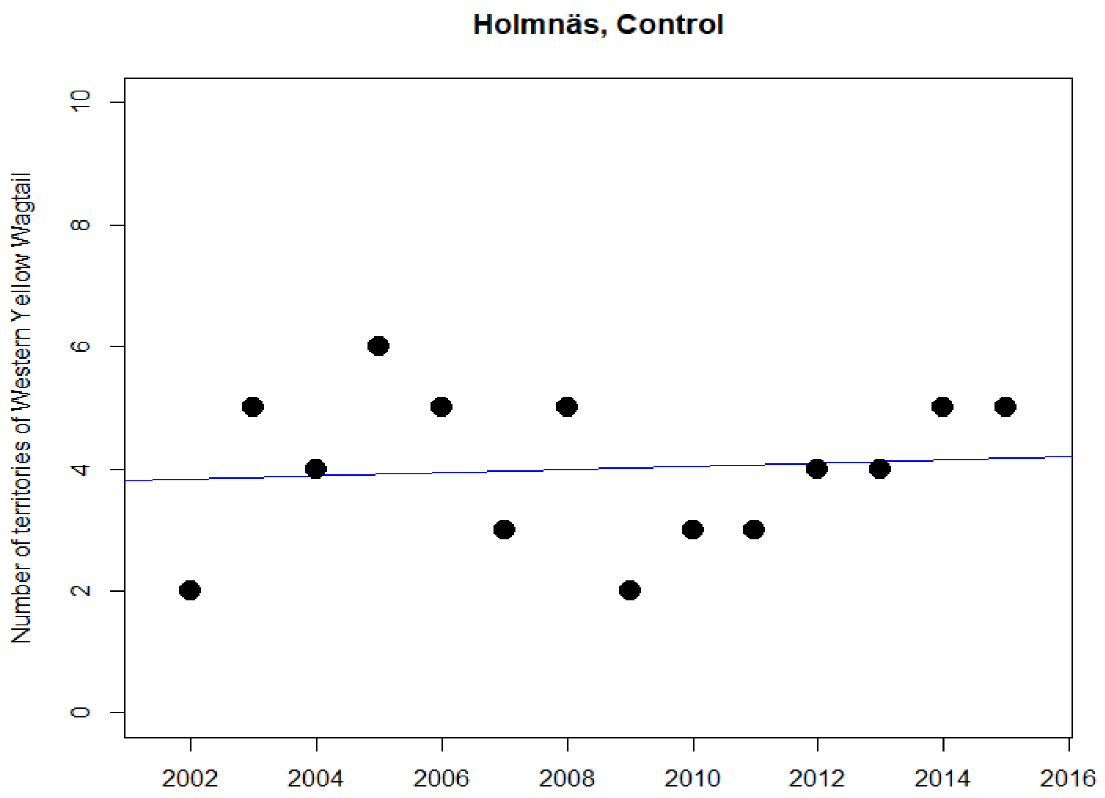
Holmnäs, Control
Number of territories of Western Yellow Wagtail
0
2
4
6
8
10
2002
2004
2006
2008
2010
2012
2014
2016

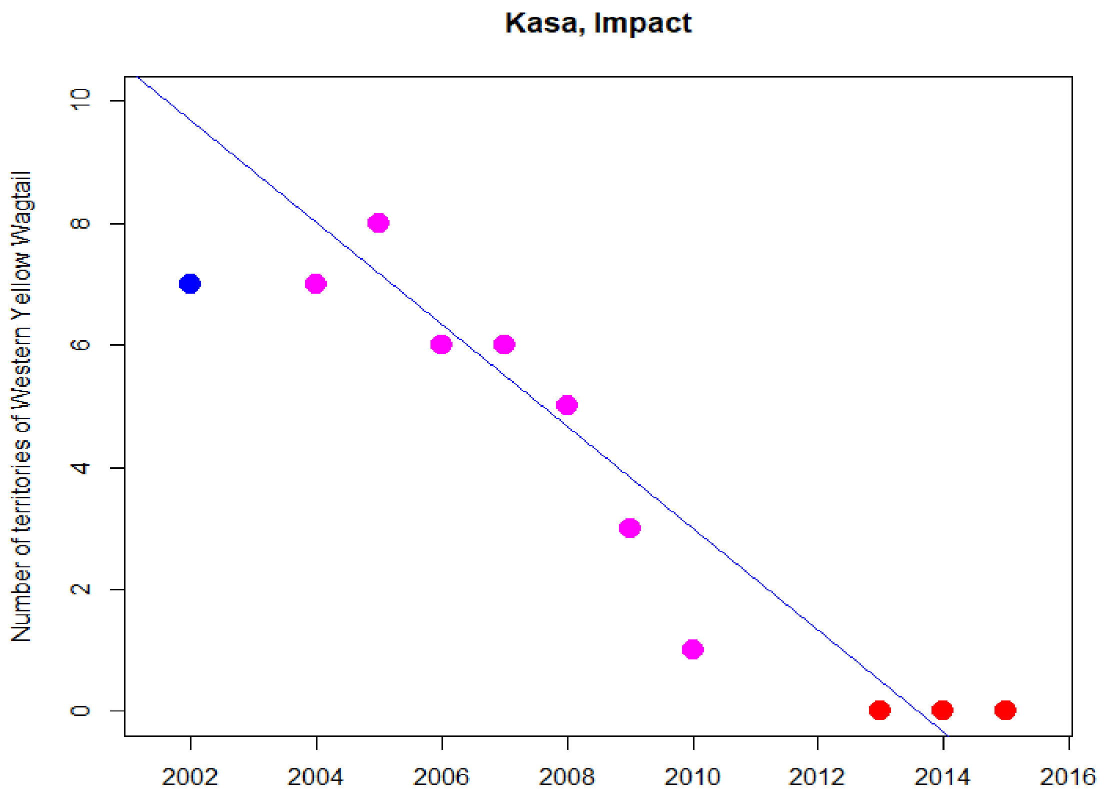
Kasa, Impact
Number of territories of Western Yellow Wagtail
0
2
4
6
8
10
2002
2004
2006
2008
2010
2012
2014
2016

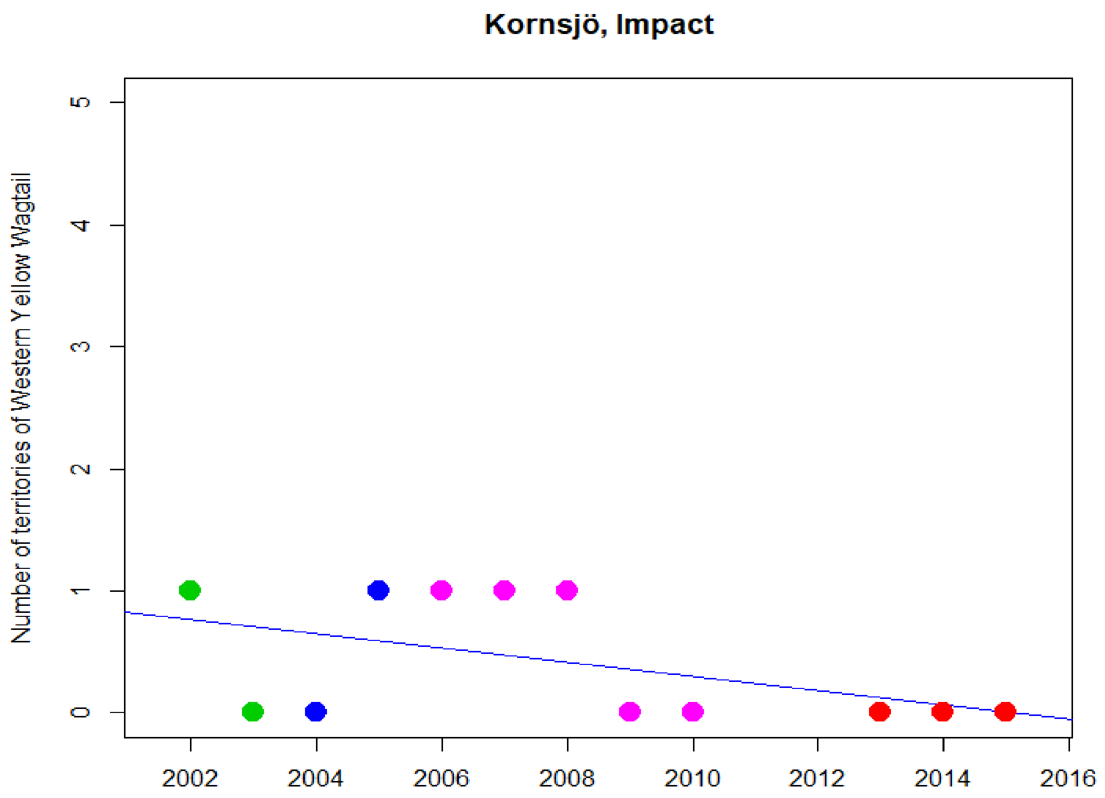

Kornsjö, Impact
Number of territories of Western Yellow Wagtail
0
1
2
3
4
5
2002
2004
2006
2008
2010
2012
2014
2016


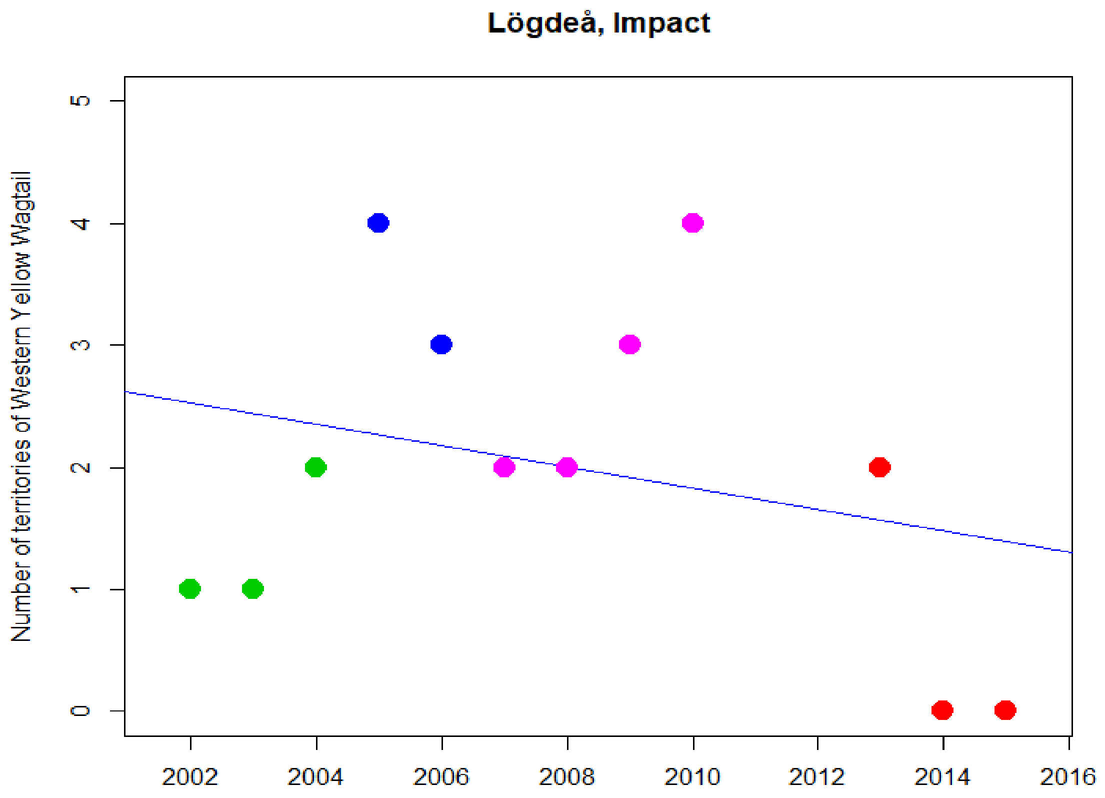

Lögdeå, Impact
Number of territories of Western Yellow Wagtail
0
1
2
3
4
5
2002
2004
2006
2008
2010
2012
2014
2016

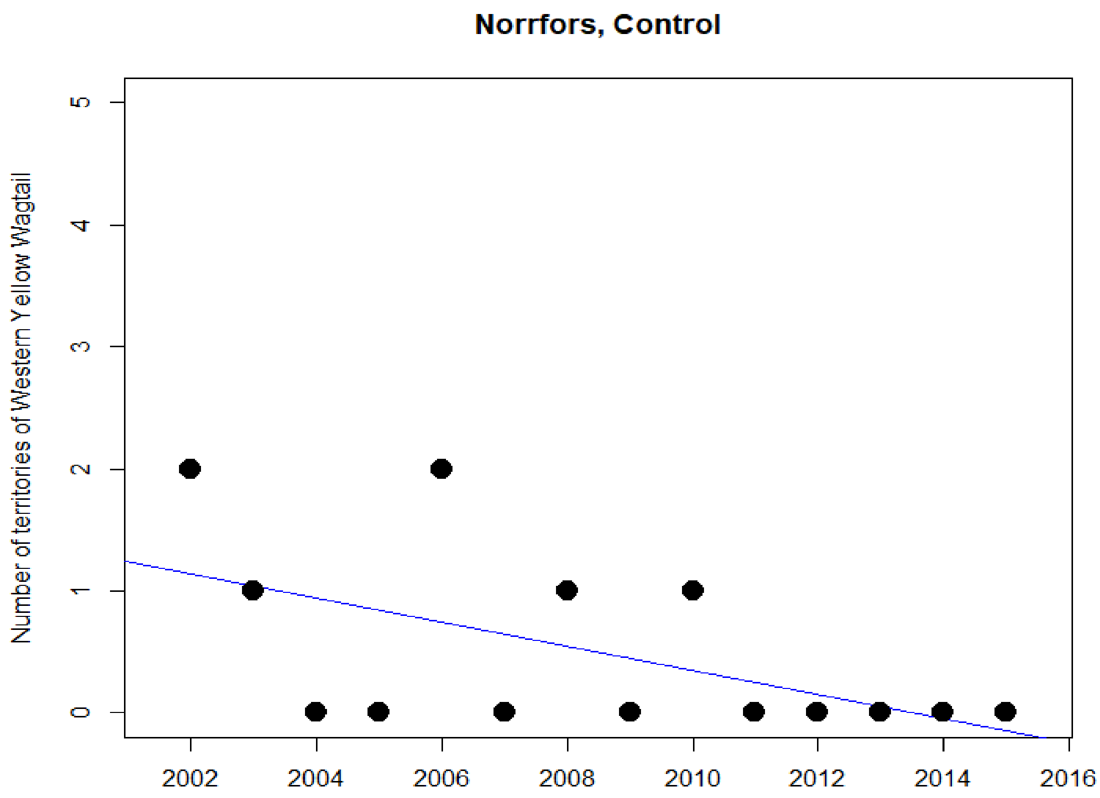
Norrfors, Control
Number of territories of Western Yellow Wagtail
0
1
2
3
4
5
2002
2004
2006
2008
2010
2012
2014
2016

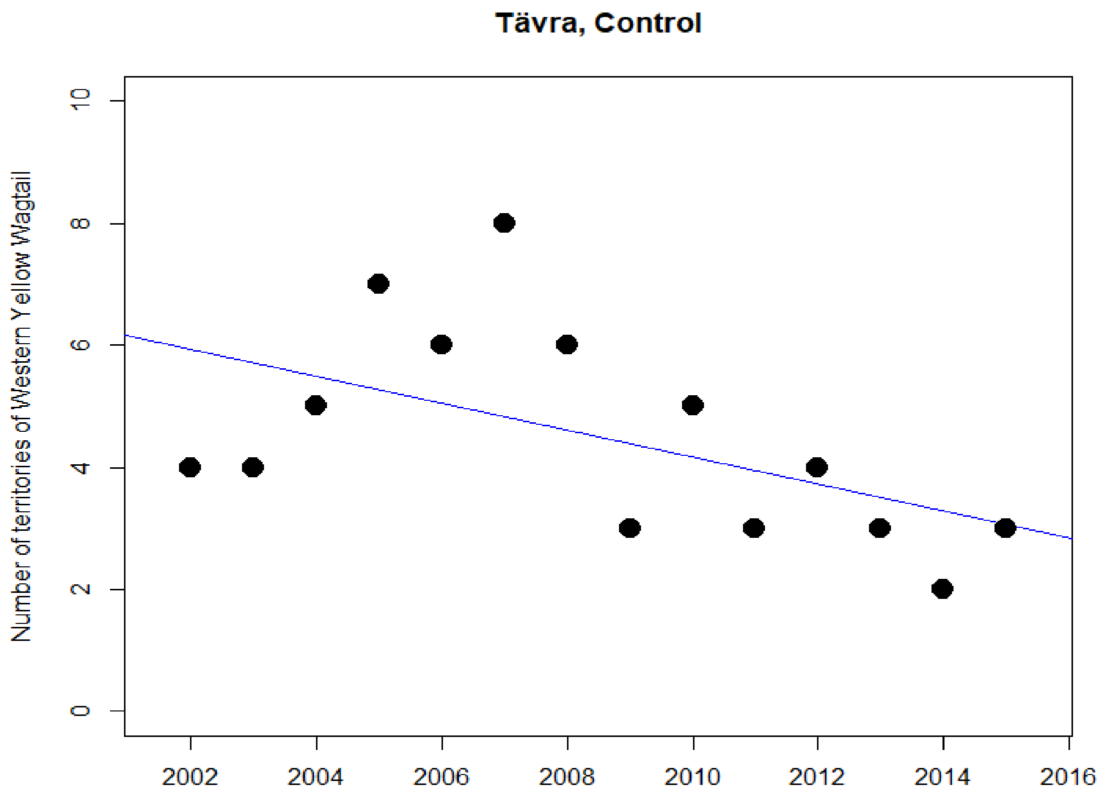
Tävra, Control
Number of territories of Western Yellow Wagtail
0
2
4
6
8
10
2002
2004
2006
2008
2010
2012
2014
2016

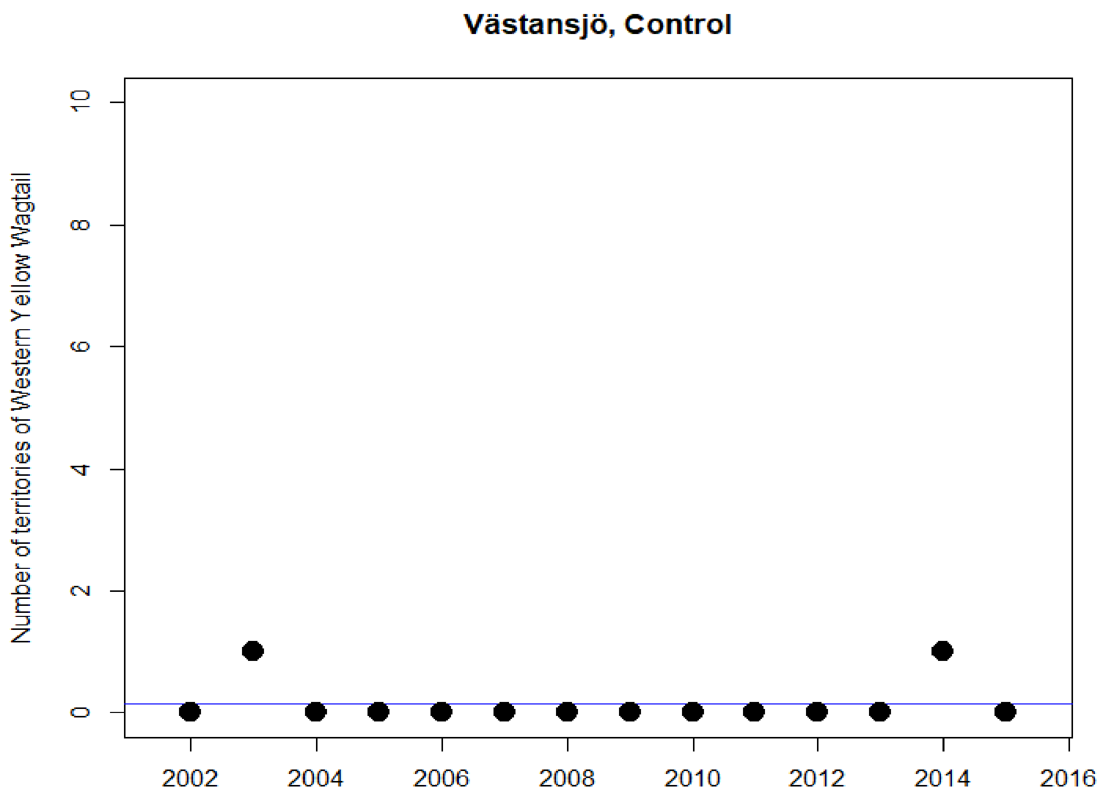
Västansjö, Control
Number of territories of Western Yellow Wagtail
0
2
4
6
8
10
2002
2004
2006
2008
2010
2012
2014
2016

### Whinchat

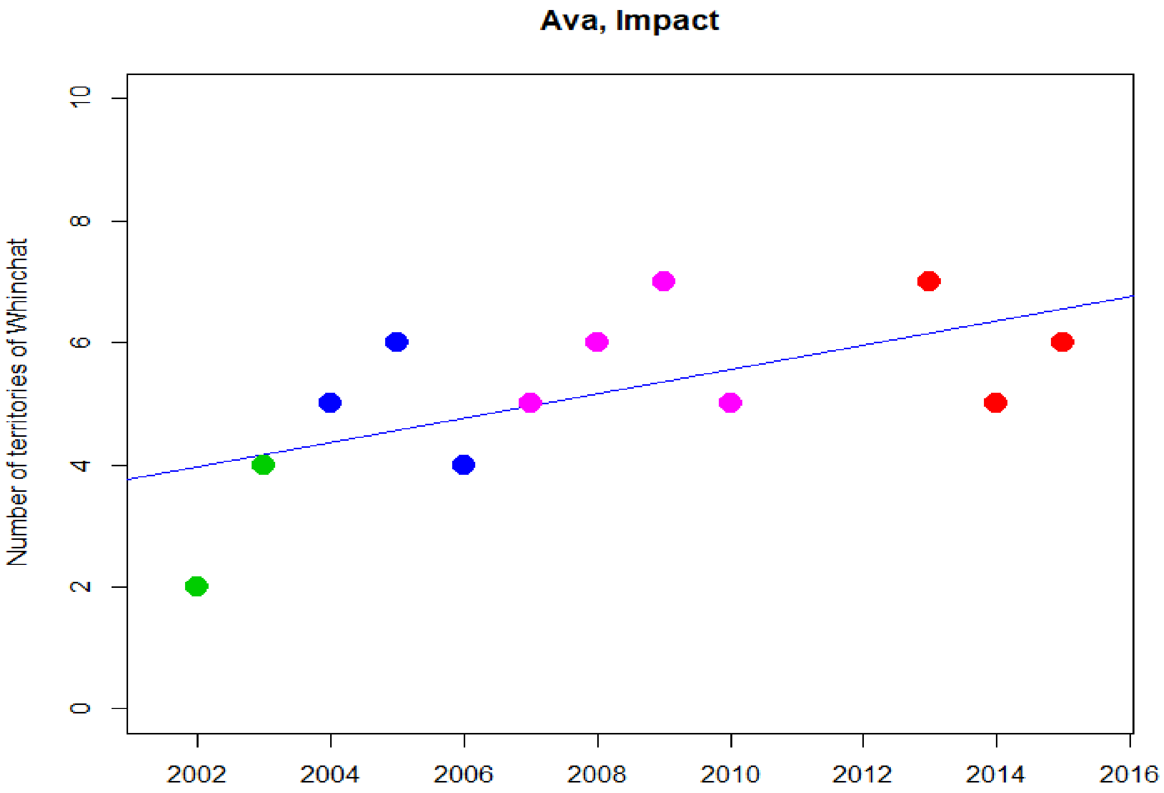
Ava, Impact
Number of territories of Whinchat
0
2
4
6
8
10
2002
2004
2006
2008
2010
2012
2014
2016

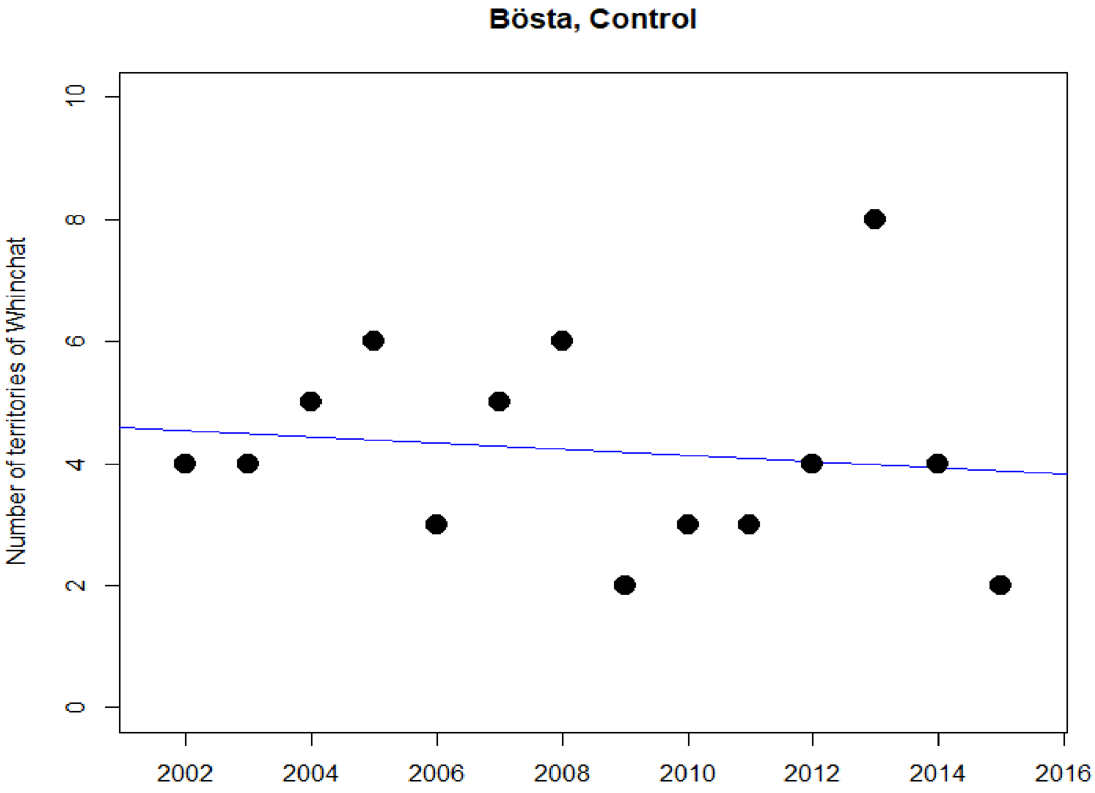
Bösta, Control
Number of territories of Whinchat
0
2
4
6
8
10
2002
2004
2006
2008
2010
2012
2014
2016

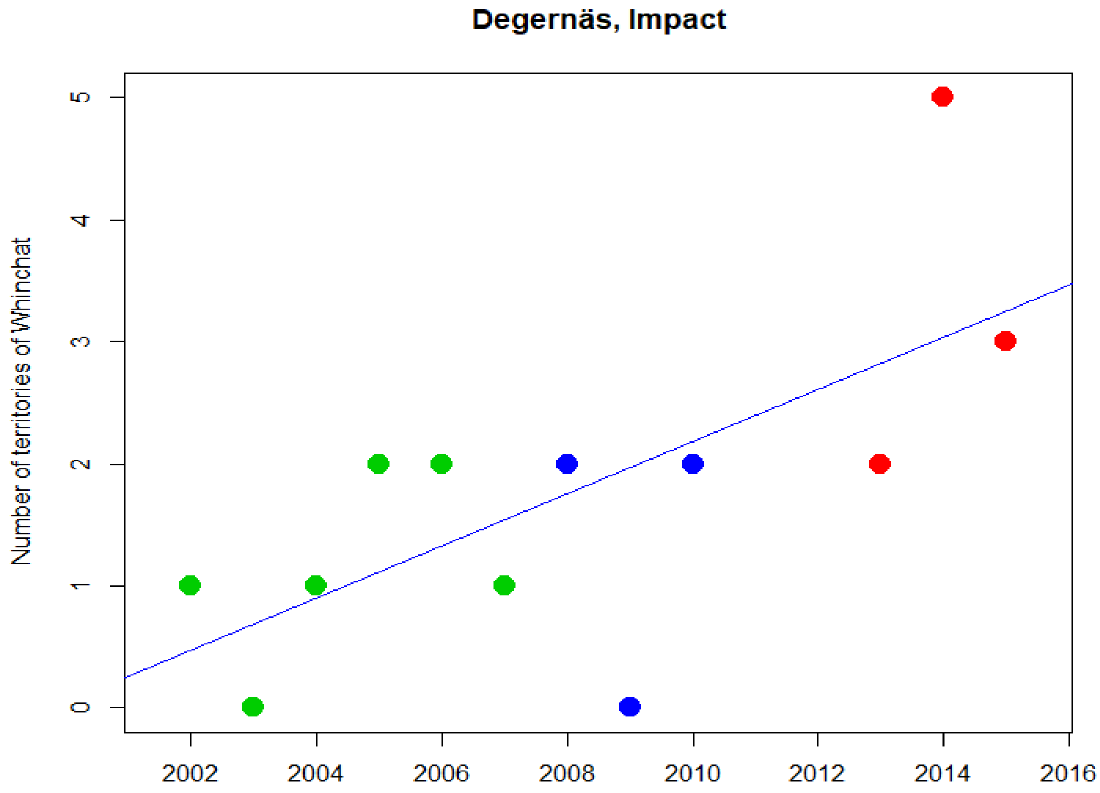
Degernäs, Impact
Number of territories of Whinchat
0
1
2
3
4
5
2002
2004
2006
2008
2010
2012
2014
2016

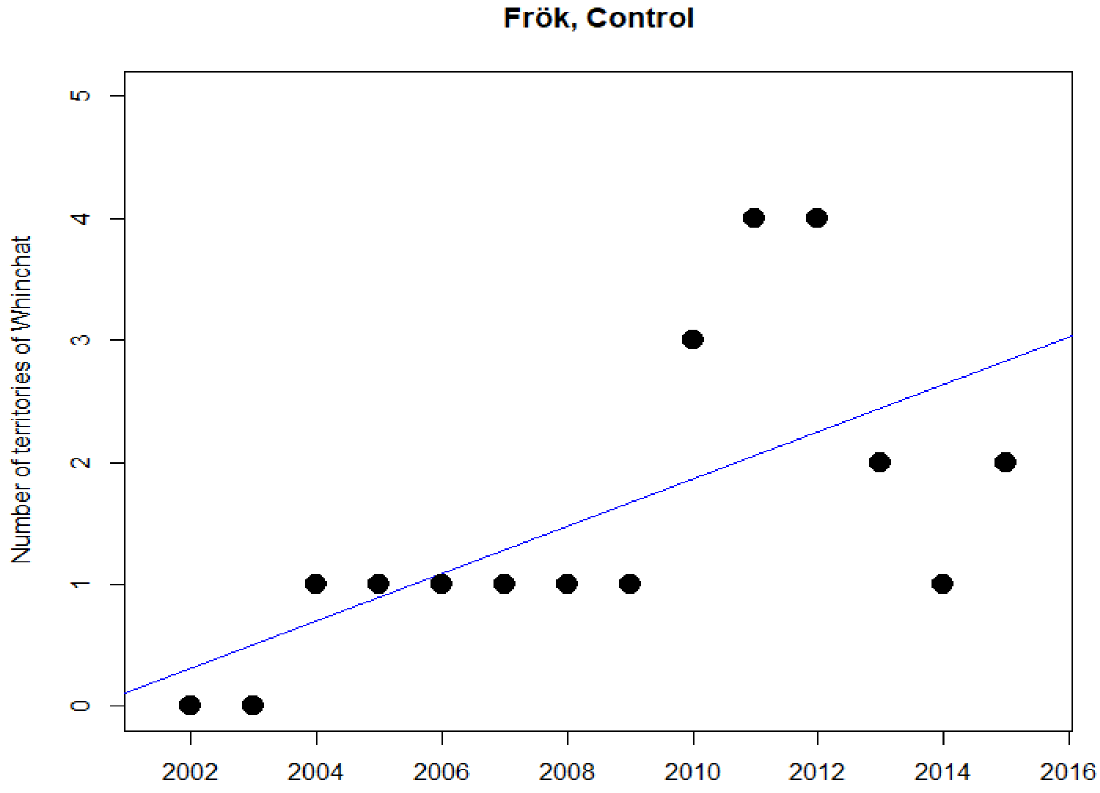
Frök, Control
Number of territories of Whinchat
0
1
2
3
4
5
2002
2004
2006
2008
2010
2012
2014
2016

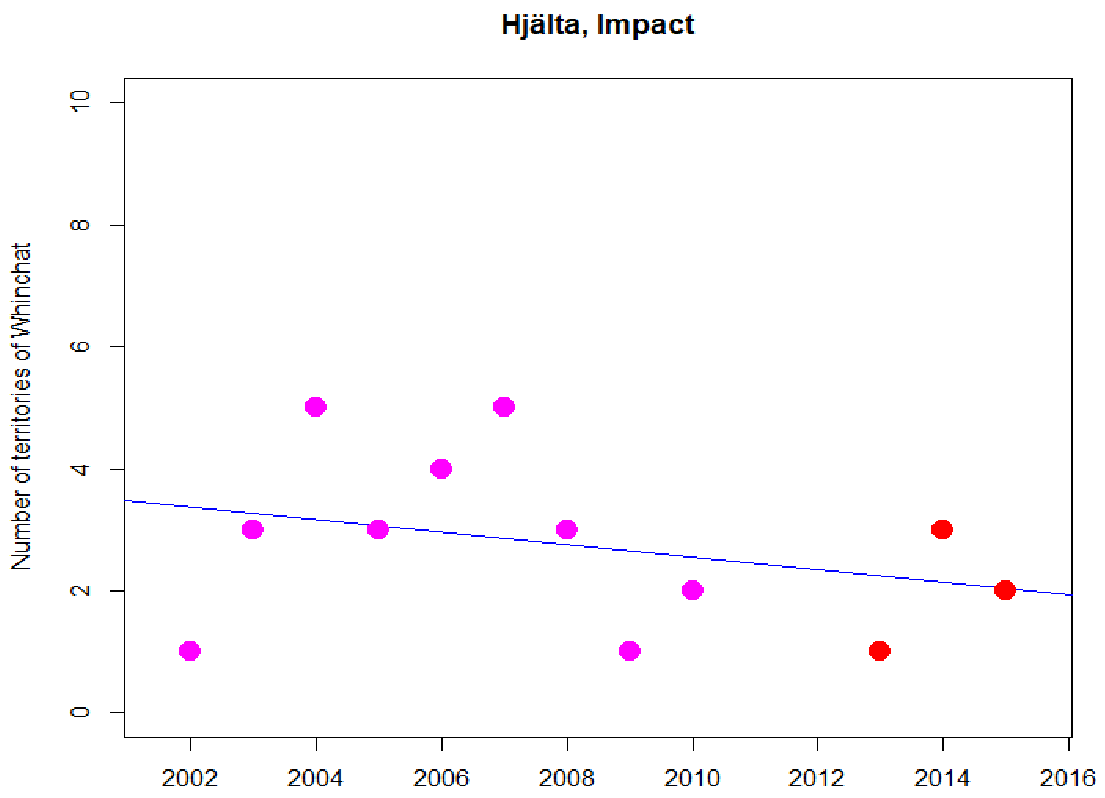
Hjälta, Impact
Number of territories of Whinchat
0
2
4
6
8
10
2002
2004
2006
2008
2010
2012
2014
2016

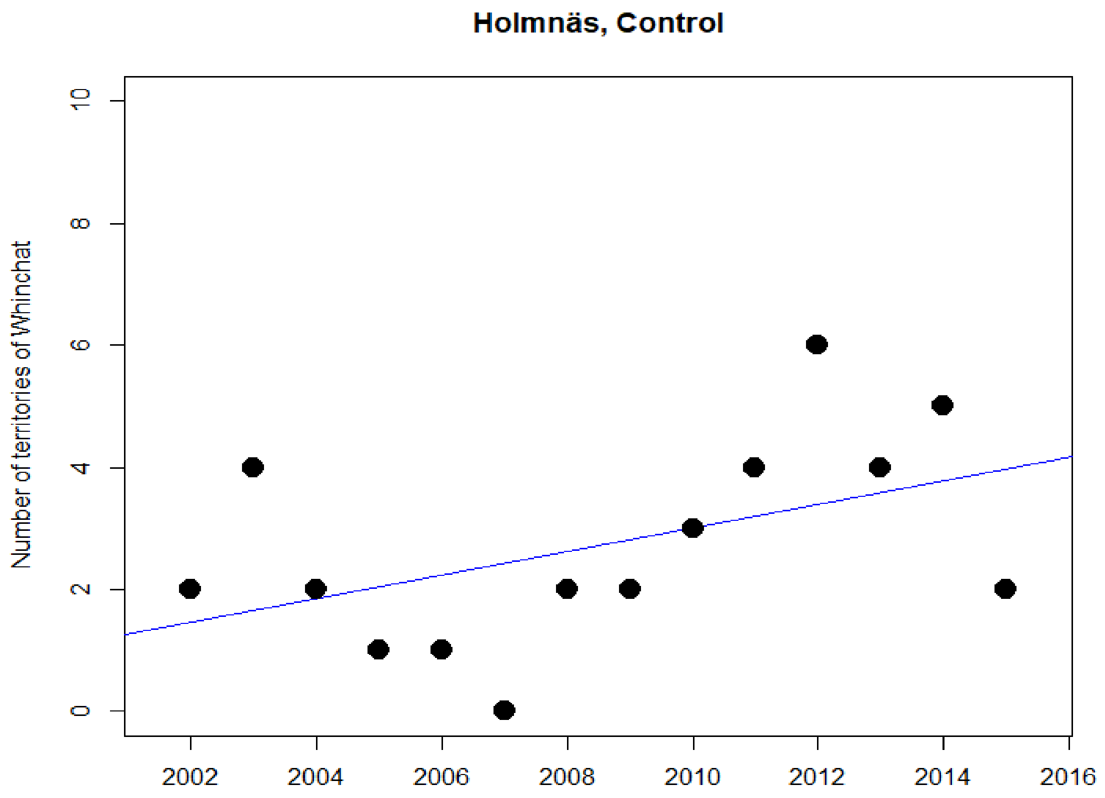
Holmnäs, Control
Number of territories of Whinchat
0
2
4
6
8
10
2002
2004
2006
2008
2010
2012
2014
2016

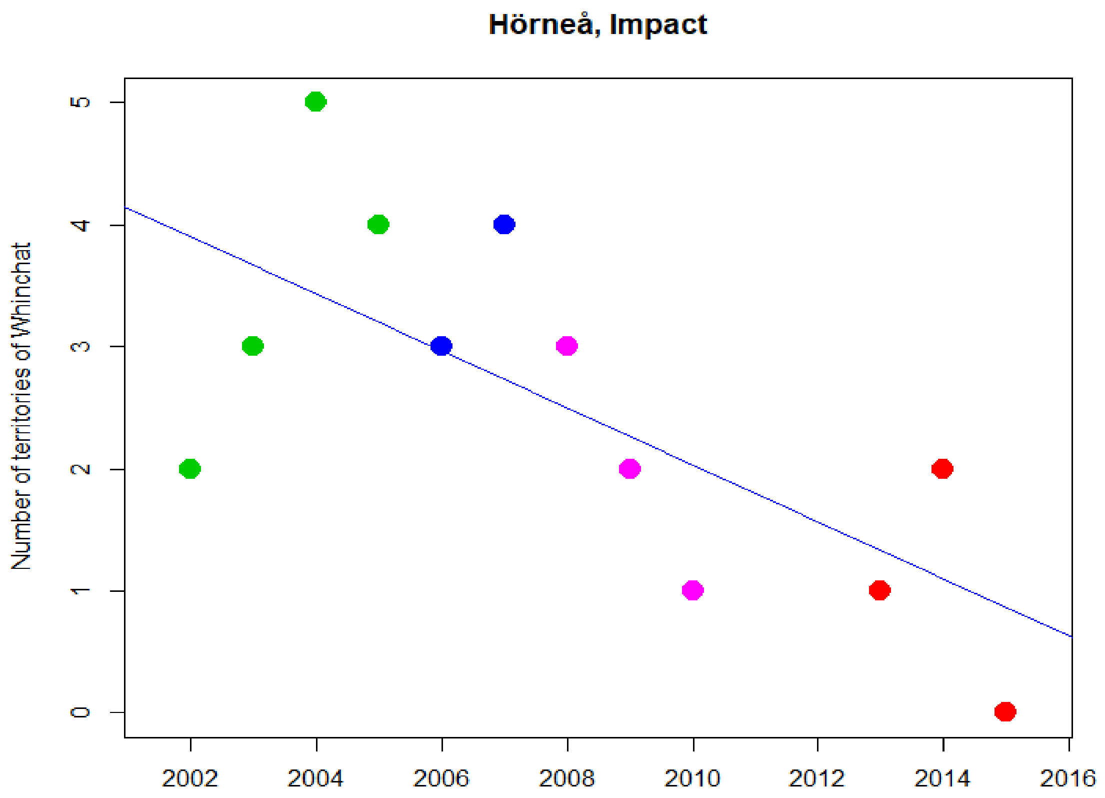
Hörneå, Impact
Number of territories of Whinchat
0
1
2
3
4
5
2002
2004
2006
2008
2010
2012
2014
2016

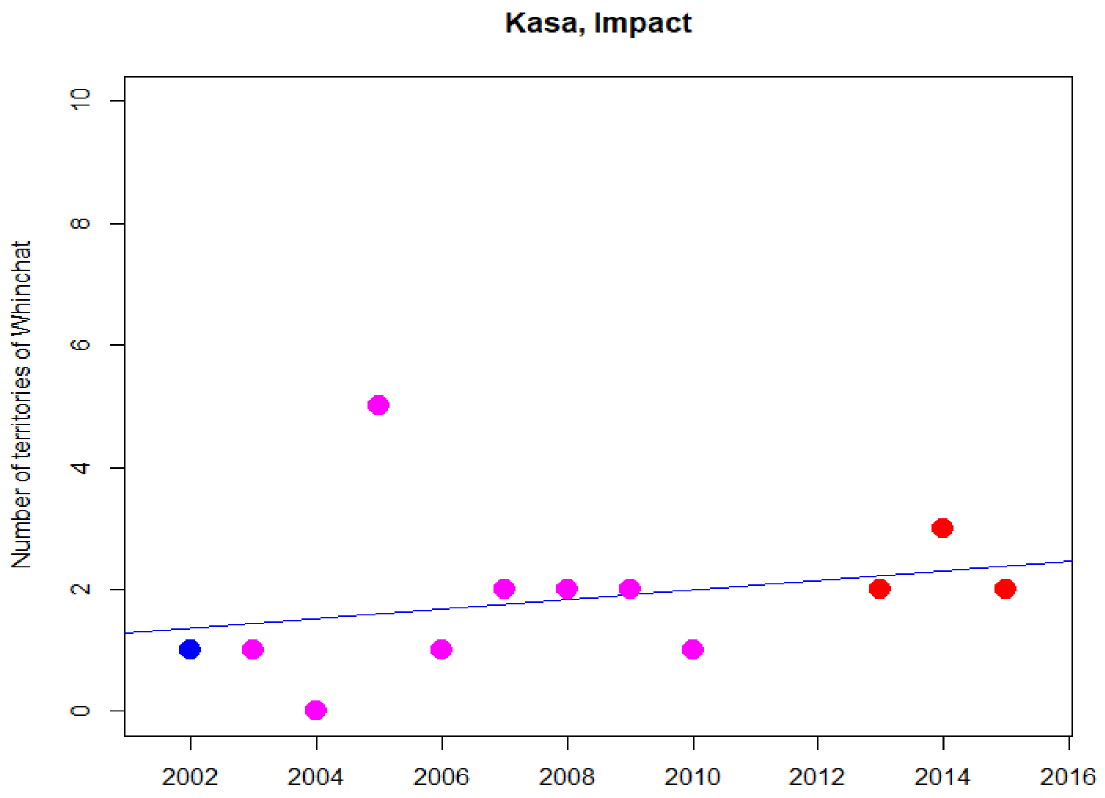
Kasa, Impact
Number of territories of Whinchat
0
2
4
6
8
10
2002
2004
2006
2008
2010
2012
2014
2016

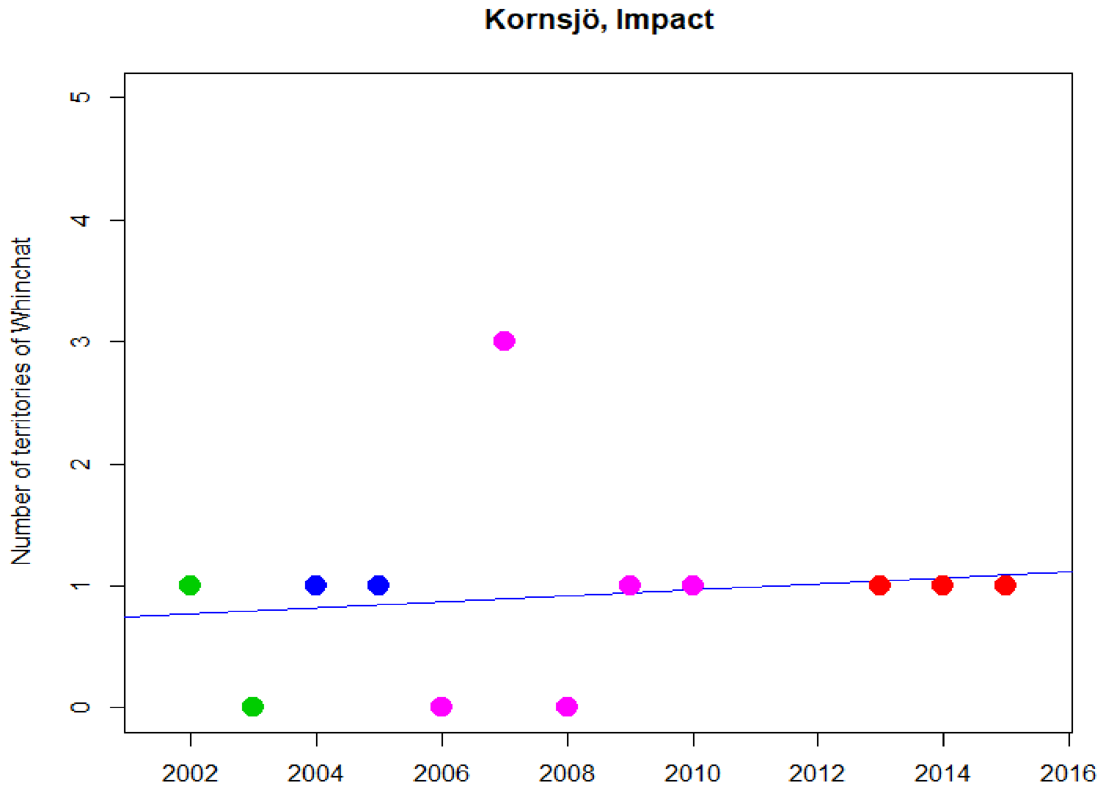
Kornsjö, Impact
Number of territories of Whinchat
0
1
2
3
4
5
2002
2004
2006
2008
2010
2012
2014
2016

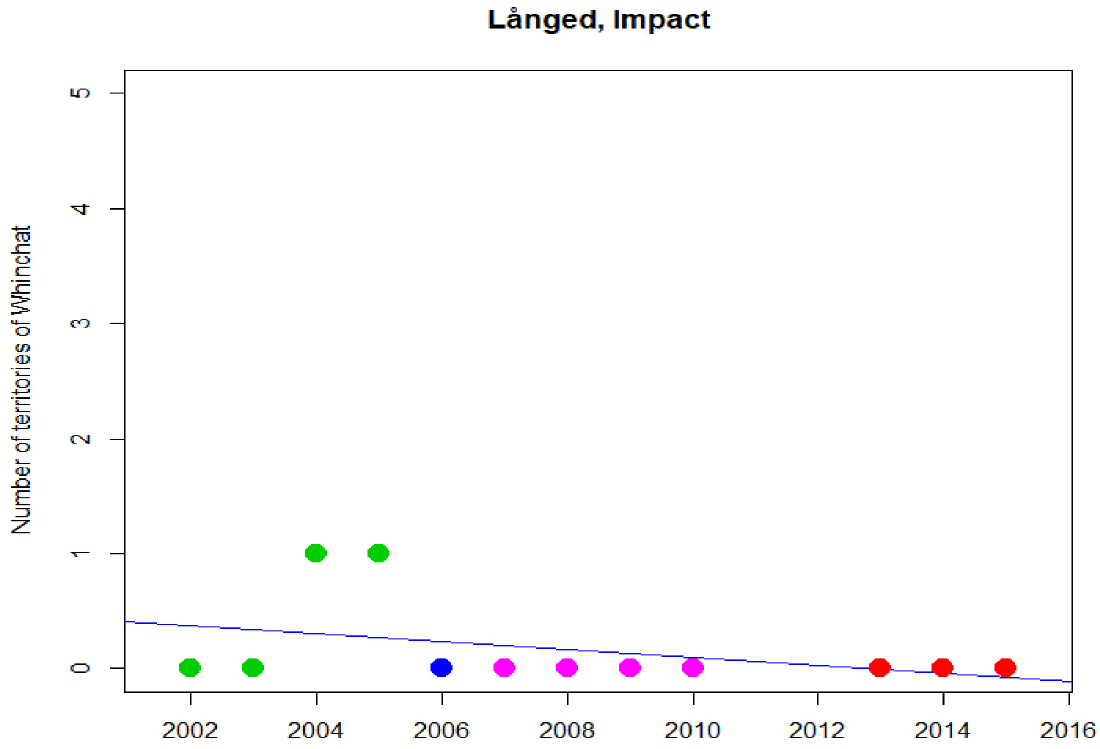
Långed, Impact
Number of territories of Whinchat
0
1
2
3
4
5
2002
2004
2006
2008
2010
2012
2014
2016

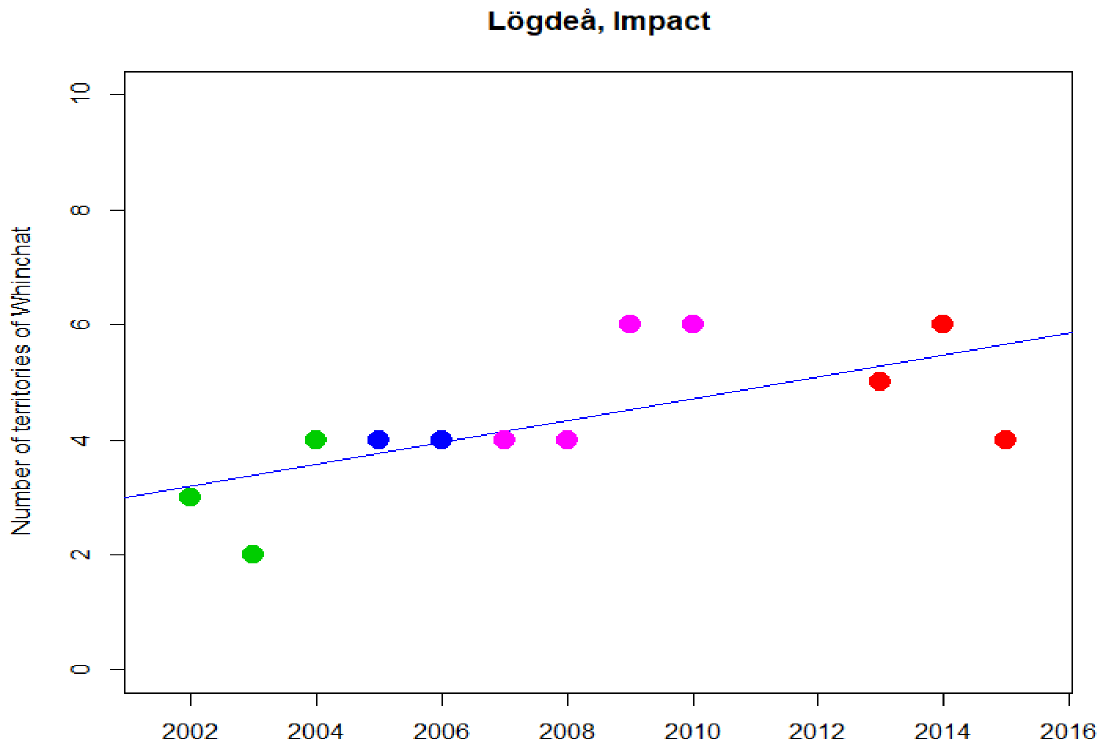
Lögdeå, Impact
Number of territories of Whinchat
0
2
4
6
8
10
2002
2004
2006
2008
2010
2012
2014
2016

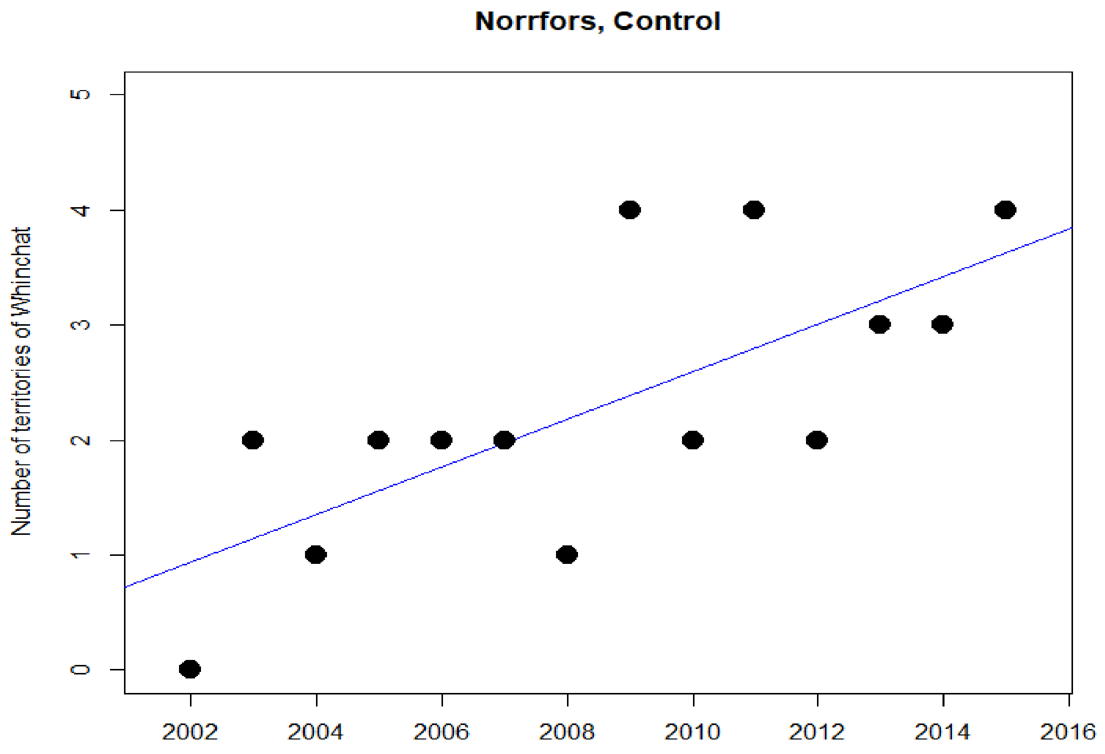
Norrfors, Control
Number of territories of Whinchat
0
1
2
3
4
5
2002
2004
2006
2008
2010
2012
2014
2016

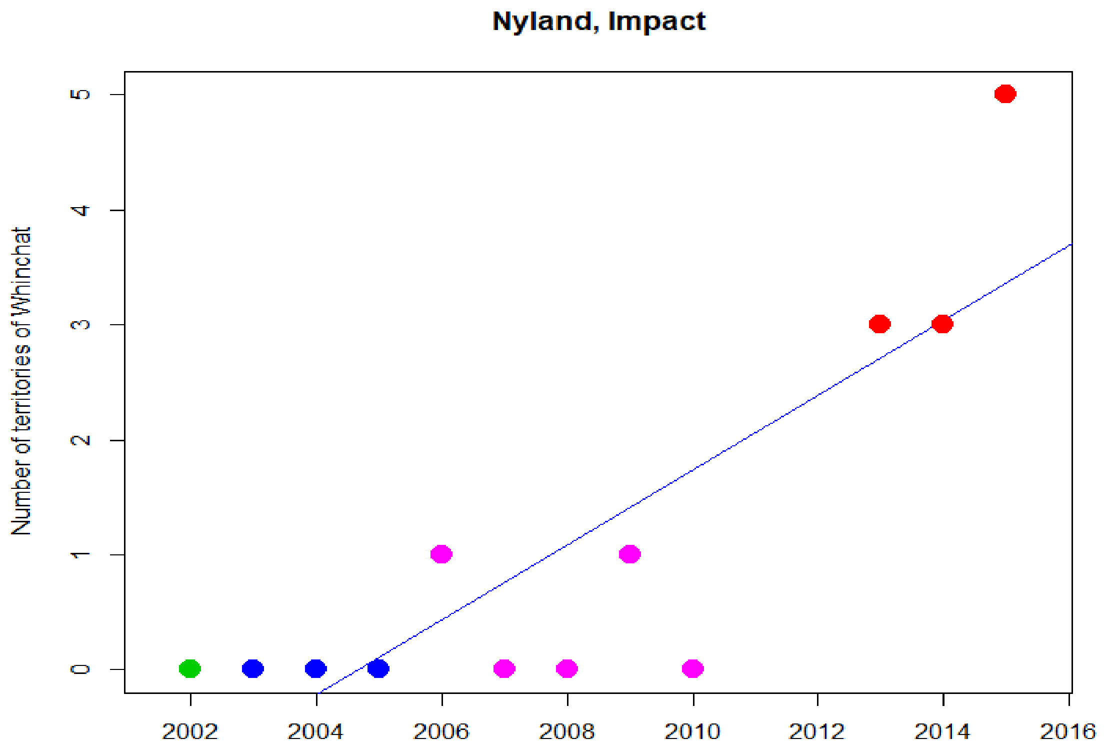

Nyland, Impact
Number of territories of Whinchat
0
1
2
3
4
5
2002
2004
2006
2008
2010
2012
2014
2016


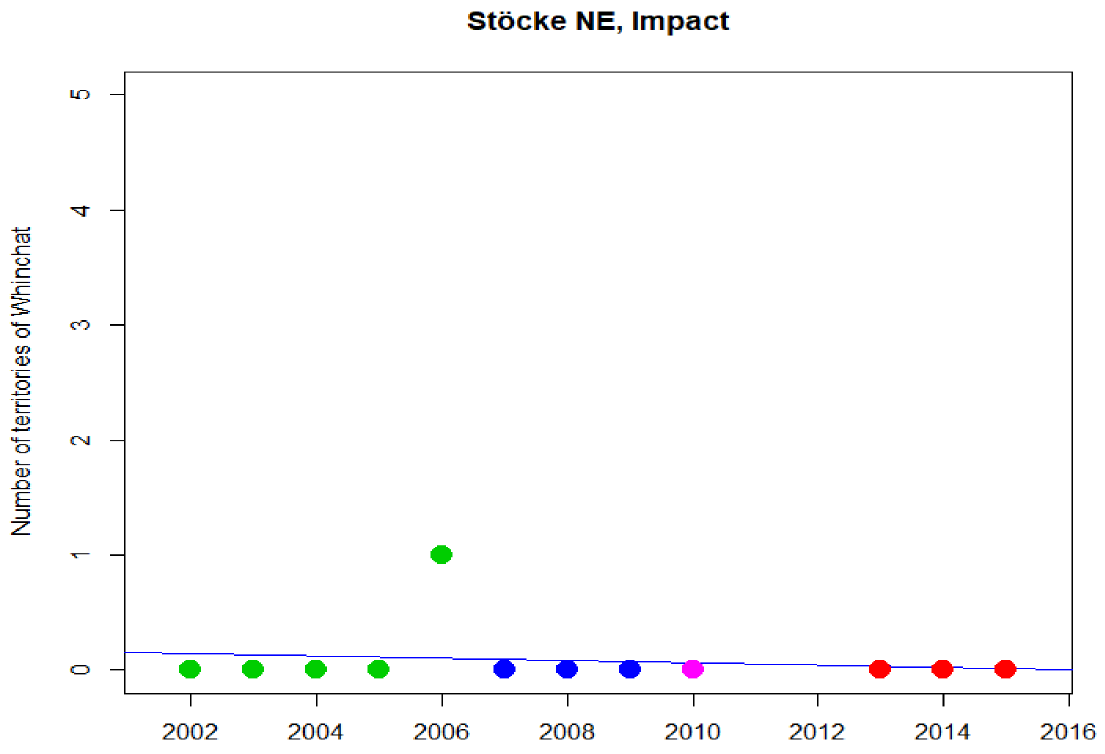

Stöcke NE, Impact
Number of territories of Whinchat
0
1
2
3
4
5
2002
2004
2006
2008
2010
2012
2014
2016

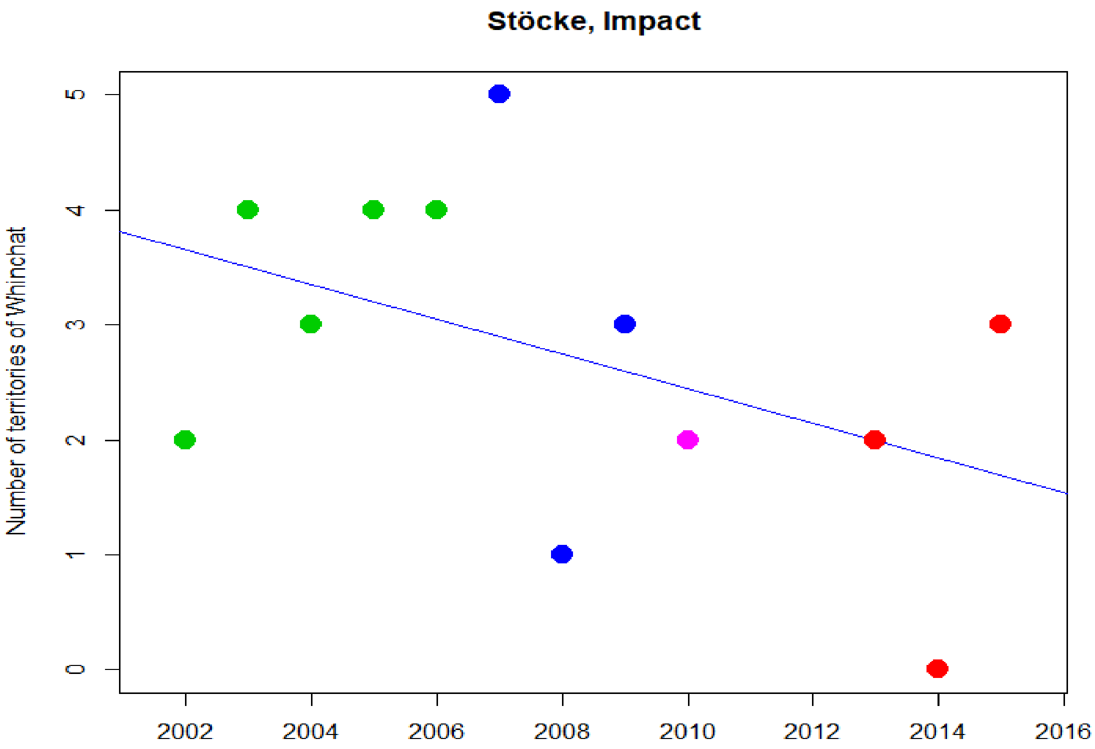
Stöcke, Impact
Number of territories of Whinchat
0
1
2
3
4
5
2002
2004
2006
2008
2010
2012
2014
2016

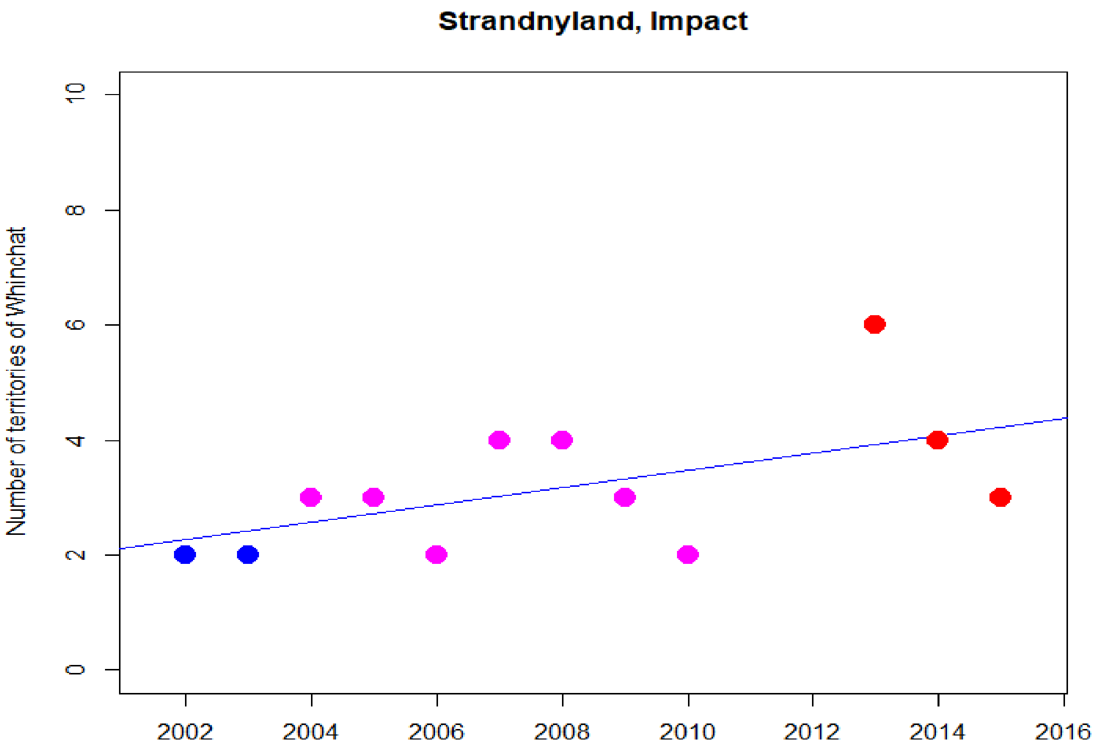
Strandnyland, Impact
Number of territories of Whinchat
0
2
4
6
8
10
2002
2004
2006
2008
2010
2012
2014
2016

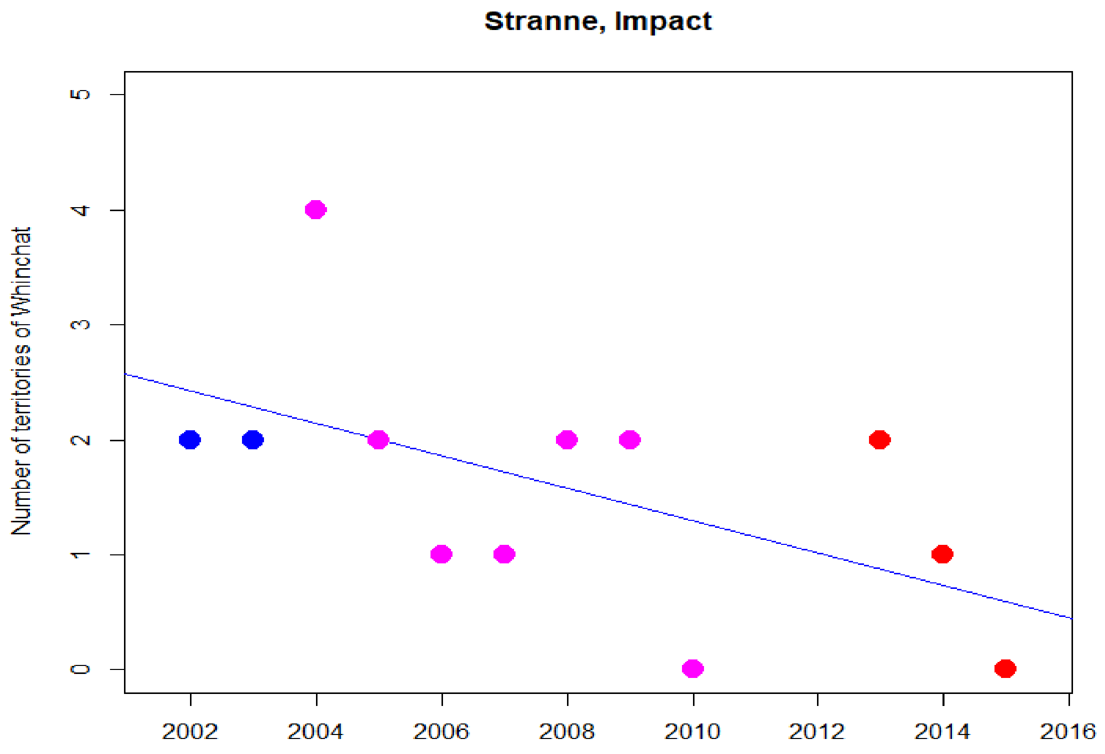
Stranne, Impact
Number of territories of Whinchat
0
1
2
3
4
5
2002
2004
2006
2008
2010
2012
2014
2016

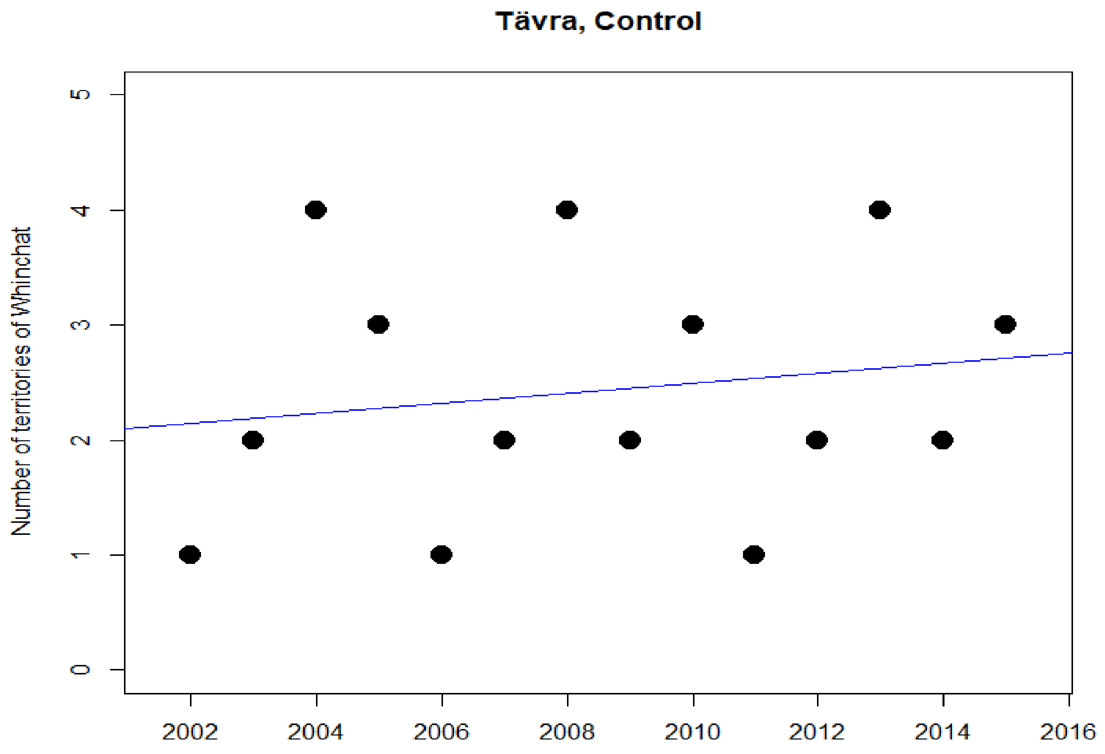
Tävra, Control
Number of territories of Whinchat
0
1
2
3
4
5
2002
2004
2006
2008
2010
2012
2014
2016

Västansjö, Control

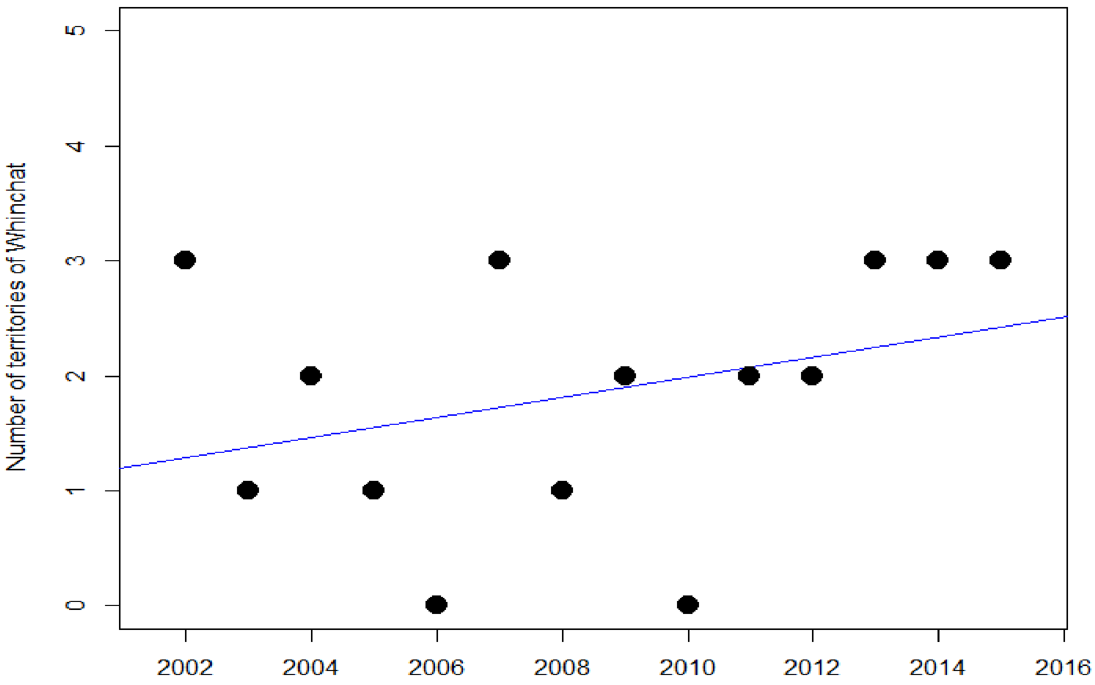
