## Supplementary material for "Are railways detrimental to bird populations? A BDACI study on the construction of the Bothnia Line Railway": Mixed effects model output for territory number data

### Supplementary material **Terr3**

### Mixed effects model output for numbers of territories of individual species across the phases of the railway construction process

Fixed effect components of GLMMadaptive models (family = poisson) with Site as random effect. All models were significantly better than the empty model (glm).

Intercept = Status *Before*

Positive estimates relate to larger numbers of territories. Significant ($P < 0.05$) estimates of Status levels vs Status *Before* are marked with yellow background.

Due to very low numbers of territories, binomial models were used for Red-backed Shrike and Common Rosefinch.

Northern Lapwing (Vv)

| | Estimate | Std.Err | z-value | p-value |
|---|---|---|---|---|
| (Intercept) | 1.23 | 0.294 | 4.17 | < 0.001 |
| StatusConstruction | -0.05 | 0.144 | -0.35 | 0.724 |
| StatusReady | -0.13 | 0.186 | -0.70 | 0.482 |
| StatusTraffic | -0.42 | 0.298 | -1.41 | 0.158 |
| YearS | 0.03 | 0.024 | 1.28 | 0.202 |
| log(NArea) | 1.43 | 0.571 | 2.50 | 0.0123 |

Eurasian Curlew (Na)

| | Estimate | Std.Err | z-value | p-value |
|---|---|---|---|---|
| (Intercept) | 1.24 | 0.172 | 7.19 | < 0.001 |
| StatusConstruction | 0.05 | 0.160 | 0.34 | 0.732 |
| StatusReady | 0.12 | 0.206 | 0.60 | 0.548 |
| StatusTraffic | 0.28 | 0.344 | 0.81 | 0.417 |
| YearS | -0.05 | 0.029 | -1.71 | 0.087 |
| log(NArea) | 1.54 | 0.259 | 5.95 | < 0.001 |

Eurasian Skylark (Aa)

| | Estimate | Std.Err | z-value | p-value |
|---|---|---|---|---|
| (Intercept) | 1.35 | 0.287 | 4.691 | < 0.001 |
| StatusConstruction | 0.33 | 0.165 | 2.020 | 0.043 |
| StatusReady | 0.40 | 0.228 | 1.750 | 0.080 |
| StatusTraffic | 0.47 | 0.358 | 1.302 | 0.193 |
| YearS | -0.08 | 0.030 | -2.610 | 0.009 |
| log(NArea) | 1.62 | 0.520 | 3.167 | 0.002 |

Barn Swallow (Hr)

| | Estimate | Std.Err | z-value | p-value |
|---|---|---|---|---|
| (Intercept) | 1.61 | 0.168 | 9.584 | < 0.001 |
| StatusConstruction | 0.02 | 0.143 | 0.138 | 0.890 |
| StatusReady | 0.19 | 0.178 | 1.073 | 0.283 |
| StatusTraffic | 0.17 | 0.306 | 0.567 | 0.571 |
| YearS | -0.02 | 0.025 | -0.792 | 0.428 |
| log(NArea) | 0.84 | 0.254 | 3.327 | < 0.001 |

Meadow Pipit (Ap)

| | Estimate | Std.Err | z-value | p-value |
|---|---|---|---|---|
| (Intercept) | 0.05 | 0.536 | 0.093 | 0.926 |
| StatusConstruction | 0.37 | 0.479 | 0.772 | 0.440 |
| StatusReady | 0.66 | 0.660 | 0.997 | 0.319 |
| StatusTraffic | 0.36 | 1.125 | 0.324 | 0.746 |
| YearS | -0.02 | 0.102 | -0.162 | 0.871 |
| log(NArea) | -0.24 | 0.772 | -0.317 | 0.752 |

Common Starling (Sv)

| | Estimate | Std.Err | z-value | p-value |
|---|---|---|---|---|
| (Intercept) | 0.50 | 0.373 | 1.339 | 0.181 |
| StatusConstruction | -0.80 | 0.458 | -1.747 | 0.081 |
| StatusReady | -0.89 | 0.511 | -1.744 | 0.081 |
| StatusTraffic | -0.53 | 0.851 | -0.624 | 0.532 |
| YearS | -0.01 | 0.065 | -0.107 | 0.915 |
| log(NArea) | 0.55 | 0.482 | 1.132 | 0.258 |

Common Snipe (Gg)

| | Estimate | Std.Err | z-value | p-value |
|---|---|---|---|---|
| (Intercept) | -0.40 | 0.375 | -1.060 | 0.289 |
| StatusConstruction | 0.59 | 0.489 | 1.211 | 0.226 |
| StatusReady | 0.43 | 0.513 | 0.842 | 0.400 |
| StatusTraffic | 0.74 | 0.897 | 0.830 | 0.407 |
| YearS | -0.005 | 0.077 | -0.062 | 0.951 |
| log(NArea) | -0.59 | 0.493 | -1.201 | 0.230 |

Green Sandpiper (To)

| | Estimate | Std.Err | z-value | p-value |
|---|---|---|---|---|
| (Intercept) | -0.30 | 0.819 | -0.365 | 0.715 |
| StatusConstruction | -0.05 | 1.158 | -0.041 | 0.967 |
| StatusReady | -0.12 | 1.835 | -0.067 | 0.947 |
| StatusTraffic | 0.20 | 3.500 | 0.058 | 0.954 |
| YearS | -0.08 | 0.311 | -0.258 | 0.797 |
| log(NArea) | -0.15 | 0.568 | -0.261 | 0.794 |

Western Yellow Wagtail (Mf)

| | Estimate | Std.Err | z-value | p-value |
|---|---|---|---|---|
| (Intercept) | -0.21 | 0.499 | -0.423 | 0.672 |
| StatusConstruction | 1.64 | 0.361 | 4.556 | < 0.001 |
| StatusReady | 2.51 | 0.411 | 6.123 | < 0.001 |
| StatusTraffic | 2.86 | 0.602 | 4.761 | < 0.001 |
| YearS | -0.20 | 0.042 | -4.846 | < 0.001 |
| log(NArea) | 0.12 | 1.991 | 0.062 | 0.950 |

Whinchat (Sr)

| | Estimate | Std.Err | z-value | p-value |
|---|---|---|---|---|
| (Intercept) | 0.69 | 0.205 | 3.372 | < 0.001 |
| StatusConstruction | 0.03 | 0.214 | 0.150 | 0.881 |
| StatusReady | 0.15 | 0.252 | 0.578 | 0.564 |
| StatusTraffic | 0.22 | 0.426 | 0.522 | 0.602 |
| YearS | 0.003 | 0.035 | 0.088 | 0.930 |
| log(NArea) | 0.44 | 0.288 | 1.519 | 0.129 |

Red-backed Shrike (Lc) binomial

| | Estimate | Std.Err | z-value | p-value |
|---|---|---|---|---|
| (Intercept) | -2.99 | 1.061 | -2.8154 | 0.005 |
| StatusConstruction | -0.25 | 1.197 | -0.2078 | 0.835 |
| StatusReady | -0.32 | 1.310 | -0.2475 | 0.805 |
| StatusTraffic | -4.93 | 2.563 | -1.9227 | 0.055 |
| YearS | 0.40 | 0.219 | 1.8479 | 0.065 |
| log(NArea) | 0.83 | 1.400 | 0.5912 | 0.554 |

Common Rosefinch (Ce), binomial

| | Estimate | Std.Err | z-value | p-value |
|---|---|---|---|---|
| (Intercept) | 1.76 | 1.345 | 1.310 | 0.190 |
| StatusConstruction | -0.20 | 1.489 | -0.132 | 0.895 |
| StatusReady | 1.73 | 1.823 | 0.948 | 0.343 |
| StatusTraffic | 3.56 | 3.193 | 1.114 | 0.265 |
| YearS | -0.57 | 0.251 | -2.260 | 0.024 |
| log(NArea) | -2.43 | 1.766 | -1.375 | 0.169 |
