## Supplementary figures and tables for "Are railways detrimental to bird populations? A BDACI study on the construction of the Bothnia Line Railway"

This document contains four figures and two tables that are referred to in the main text.

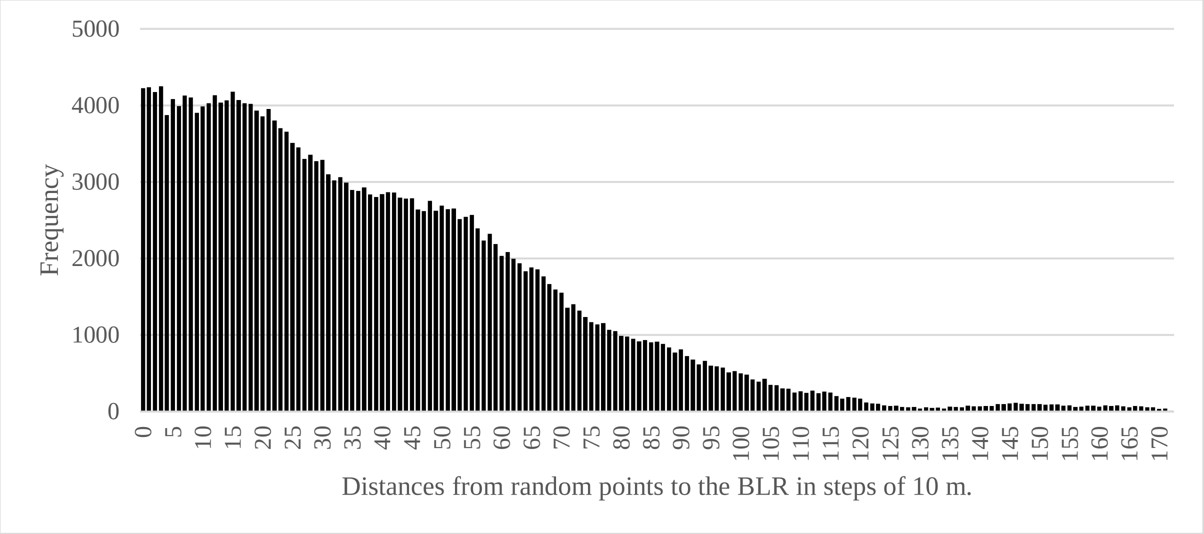


Figure S1. Frequency distribution of nearest distances between random points within the study sites and the BLR. This distribution reflects the uneven shape of the “agricultural land landmass” relative the BLR.

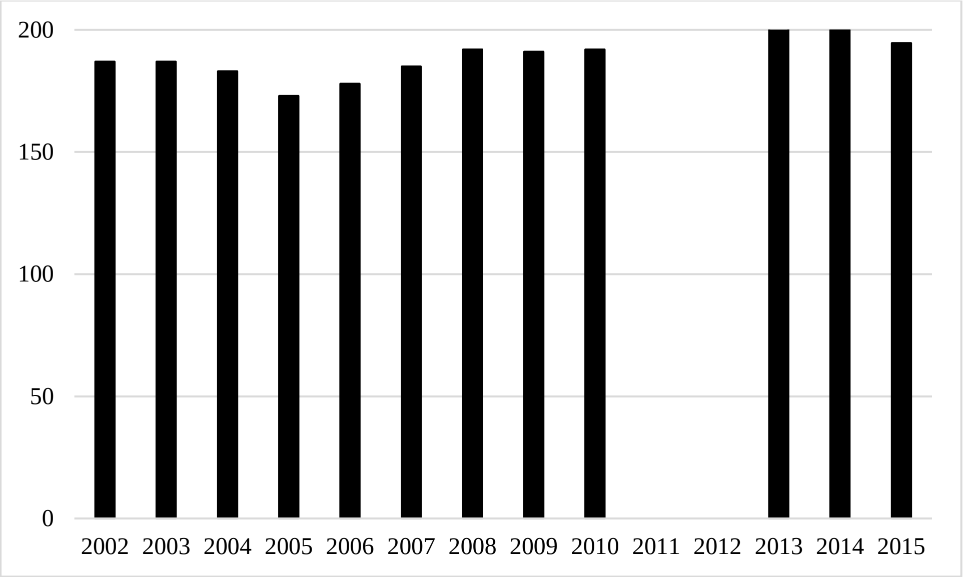


Figure S2. Yearly total effort (h) of the fieldwork for this study. In 2011 and 2012, only Control sites were surveyed.

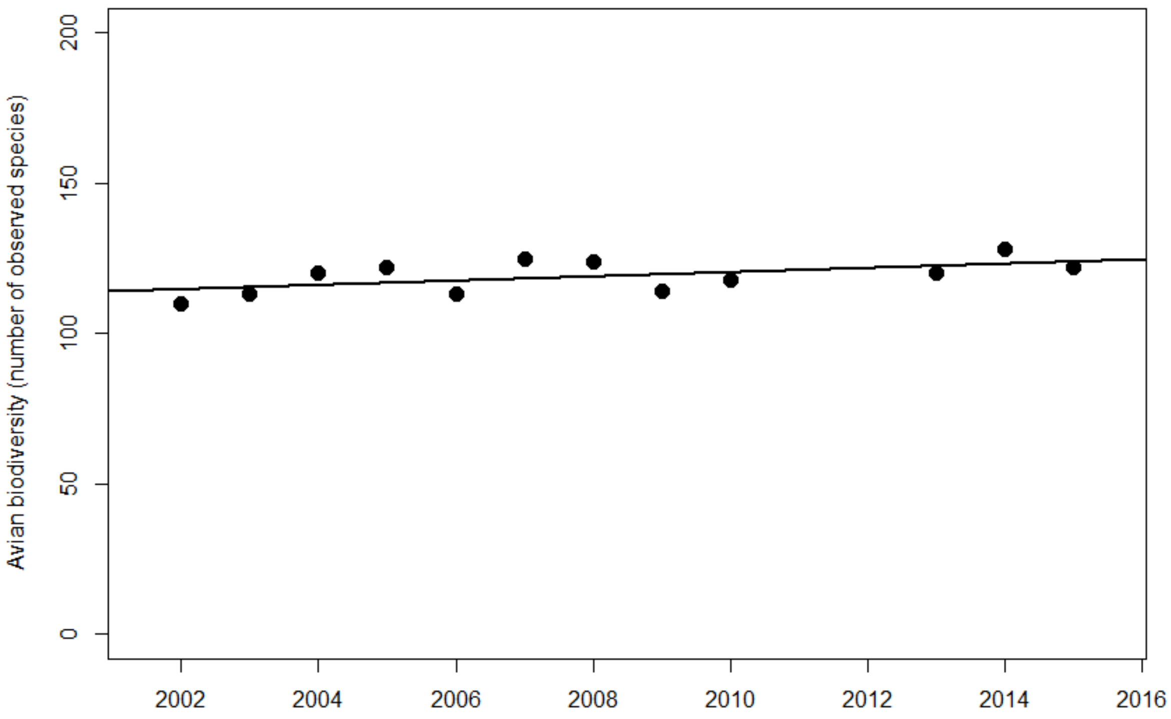


Figure S3. Yearly numbers of bird species observed in 17 study sites. Linear regression line added for visualization.

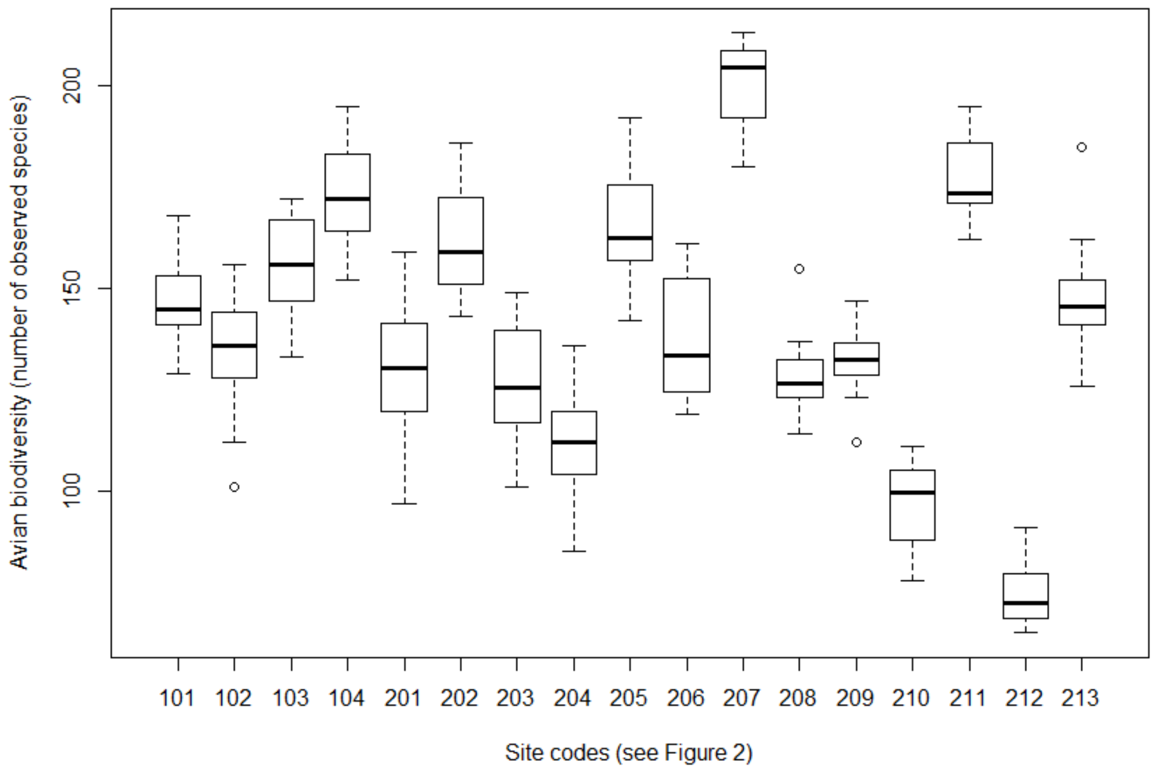


Figure S4. Boxplot of numbers of observed species during the study period in Control (101-104) and Impact (201-213) sites.

Table S1. Avian biodiversity index (*Traffic* = 100) for all 13 Impact sites during the various phases of the railway construction process. NA's indicate that the Status category was not represented during the study period.

| Impact site | Before | Construction | Ready | Traffic |
|---|---|---|---|---|
| 201 | 99 | 97 | 99 | 100 |
| 202 | 105 | 101 | 108 | 100 |
| 203 | NA | 105 | 101 | 100 |
| 204 | NA | 104 | 121 | 100 |
| 205 | NA | NA | 94 | 100 |
| 206 | NA | 115 | 100 | 100 |
| 207 | 93 | 95 | 102 | 100 |
| 208 | 103 | 98 | 107 | 100 |
| 209 | 99 | 89 | 102 | 100 |
| 210 | 96 | 96 | 98 | 100 |
| 211 | 93 | 96 | 95 | 100 |
| 212 | 97 | 116 | 100 | 100 |
| 213 | 93 | 97 | NA | 100 |

Table S2. Temporal trends of TMtR distances per species and site expressed in Kendall rank correlation sign and probability classes (Supplementary material Dist4 for details). Red background marks increasing distances (= territory midpoints move away from the railway) and yellow background decreasing distances (= midpoints move closer to the railway). Grey background = no significant trend. * = P<0.05, ** = P < 0.01 and *** = P < 0.001. Empty cells for cases where low numbers of territories prevented correlation tests. Table 2 for species name abbreviations.

| Site | Vv | Na | Aa | Ap | Hr | Sv | Eh | Cd | Gg | To | Mf | Sr | Lc | Ce |
|---|---|---|---|---|---|---|---|---|---|---|---|---|---|---|
| Nyland | | | | | | | | | | | | | | |
| Kornsjö | *** | | | | | | | | | | | | | |
| Stranne | | | | | | | | | | | | | | |
| Strandnyland | | | | | | | | | | | * | | | |
| Hjälta | | | | | | | | | | | | | | |
| Kasa | | | | | | | | | | | ** | | | |
| Ava | *** | | | | | | | | | | | *** | | |
| Lögdeå | | | | | | | | | | | | | | |
| Långed | | | | | *** | | | | | | | | | |
| Hörneå | | | | | | | | | | | | | | |
| Stöcke | | | | | | * | | | | | | | | |
| Stöcke NE | | | | | | | | | | | | | | |
| Degernäs | * | | | | | | | | | | | | | |
